# Winners and Losers Among European Arthropods over the Last Half Century of Global Change

**DOI:** 10.64898/2026.01.18.700202

**Authors:** Vaughn Shirey, Vasco Veiga Branco, Conrad P.D.T. Gillett, Marta Goula, Viktor Hartung, Axel Hochkirch, Lauri J. Kaila, Alain Maasri, Thomas Merrien, Marija Miličić, Juho Paukkunen, Sergei Tarasov, Artur Taszakowski, Veikko Yrjölä, Laura Melissa Guzman, Pedro Cardoso

## Abstract

Widespread concern over arthropod declines has raised questions about how global change is reshaping biodiversity, yet large-scale responses within the group remain poorly understood. Here, we analyse 50 years (1970-2019) of European arthropod occurrence data to quantify distributional changes across more than 700 species within 11 taxonomic groups. Using occupancy-detection models that account for imperfect detection, we compare time-only and environmentally mediated trends, including changes to temperature, precipitation, and agricultural and urban land cover. We find no universal signal of decline; responses are highly heterogeneous, revealing distinct winners and losers both within and among groups. Climate variables, particularly temperature and precipitation, are the strongest predictors of change, while agricultural intensification is broadly negative and urbanisation produces group-specific effects. Trait analyses indicate that warm- and dry-affiliated species and larger-bodied arthropods generally fare better. These results show that recent European arthropod change is dominated by redistribution rather than uniform collapse, with important implications for conservation and monitoring under ongoing global change.

## MAIN TEXT

The potential for severe declines in terrestrial arthropod biodiversity remains one of the most pressing challenges of our time^1,2^. This is because arthropods perform essential roles in nearly all ecosystems, including pollination, nutrient cycling, and pest control^3,4^. In addition to these critical roles, terrestrial arthropods are fundamental components of food webs and essential components of vertebrate diets, including those of birds and mammals^5–7^. Species traits can mediate these ecosystem services and the probability that arthropod species may experience population declines. Thus, declines in arthropod populations will undoubtedly impact the stability and function of ecosystems worldwide. In the face of global change, understanding the drivers of these losses and the traits that confer vulnerability or advantage to arthropod species is a defining goal of biodiversity science and conservation efforts.

Europe is where the most data on arthropod diversity has been collected across space and time, and, consequently, much of the current narrative around arthropod declines is based in the continent^8–11^. Over 600 global experts have identified the primary threats to arthropods, ranking intensive agriculture, climate change, and pollution as primary drivers of change^12,13^. These factors are important to European arthropods as the history of development and climatic changes across the continent has been highly variable. For example, land use change has shifted significantly, resulting in the expansion of forests, loss of overall agricultural area, intensification of agricultural land use in remaining farmland, increased urbanization, and suburban sprawl across the continent^14,15^. In contrast, many farmland areas have been abandoned, offering the opportunity for early- and late-successional habitats to establish on previously agricultural land^16^. These changes can have significant impacts on arthropods as modifications to the plant communities in a particular habitat can have considerable bottom-up impacts on insect communities^17^. Within urban contexts, organisms must contend with habitat fragmentation, potentially limited resources, and light, noise, and chemical pollution^18,19^. These stressors can have significant impacts on survival and the distribution of urban insect populations^2^.

The climate is also rapidly changing across Europe. Rising average temperatures have driven the poleward range expansion of several arthropod and non-arthropod groups^20–23^. Changing precipitation regimes, particularly droughts, are impacting hygrophilous species (Rohde et al. 2017) and are leading to the increasing threat of desertification under the Mediterranean climate. Here, increasingly hot, arid conditions and anomalous stressor events such as heatwaves are becoming more common^24^. One of the first notable signals of global change was detected in European butterflies and moths, which have long been observed to shift poleward^20^ and whose relative position within their climatic niche shifted substantially^22^.

While external threats are a significant contributing factor to patterns of distributional shifts among arthropods, some species may be at elevated risk of decline due to intrinsic characteristics, i.e., species traits. These traits can include narrow habitat breadth, slower life histories, body size, or limited dispersal ability^25^. A recent analysis of threatened invertebrates in North America revealed that such intrinsic factors are commonly cited as risks for designating species as threatened or endangered under the United States Endangered Species Act^26^. Comparatively, species traits may also benefit organisms as they grapple with changing environments. For example, elevated dispersal capability may provide species the ability to escape unsuitable habitats, such as those that are changing due to habitat destruction or climate change. These species may be able to disperse and colonise new, suitable habitats compared to others for which movement is more challenging^21^.

Despite their importance to life on Earth, arthropods remain among the most challenging groups to monitor and conserve. This is due to their relatively low detectability, exceptionally high diversity, and variable population sizes from year to year^27–30^. Many arthropod species are uncharismatic and unpopular to humans, except for a few notable groups such as butterflies and bees^31,32^. This may lead to their deprioritization for conservation from the public’s perspective and a reduction in buy-in to monitor them^33^. These challenges necessitate bringing the current status and trends of arthropod communities to the forefront of public attention, and reconstructing such trends demands the utilisation of all available historical biodiversity data.

Here, we used historical biodiversity occurrence data from the Global Biodiversity Information Facility (GBIF) to profile occupancy trends across over 700 arthropod species in Europe from 1970 to 2019. We conducted a two-stage analysis. First, using Bayesian occupancy-detection models, we reconstructed the occupancy probability for all species at a 100km resolution, considering (a) time as a sole predictor of change, and (b) minimum temperature, precipitation, urban, and agricultural land cover as predictors of occupancy shifts. We also compared the time-only and environmental predictor models to assess whether species’ traits might predict the sensitivity of arthropod species to changes in climate and land use over the last half-century. By examining the raw responses of arthropod species in relation to their traits, as well as the differences between time-only and environmental predictor models, we develop a sense of which species may have attributes that influence general sensitivity to change over time or modeled environmental predictors. In other words, we explore where our environmental model might break down in its ability to capture a time-dependent trend based on species traits.

Overall, we found no universal signal of species-level decline; instead, a mixture of winners and losers in occupancy trend across arthropod groups (Fig. 1). Across all species in all groups, the average change in occupancy probability was an increase of about 3.1% with high variability among species (SD = +/- 25%). On average, terrestrial arthropods shifted their distributions northward by about +0.47 degrees over the 50-year reference period, i.e., ca. 1km per year (Fig. 1b). These patterns are similar to those found in birds^23^. In tandem, elevational shifts are occurring^34–37^; however, the spatial scale of our present analysis is too coarse to assess these trends in detail. In general, the time-only model showed greater variation in occupancy responses, which may be attributable to unmodeled spatially heterogeneous predictors.

**Figure 1.**
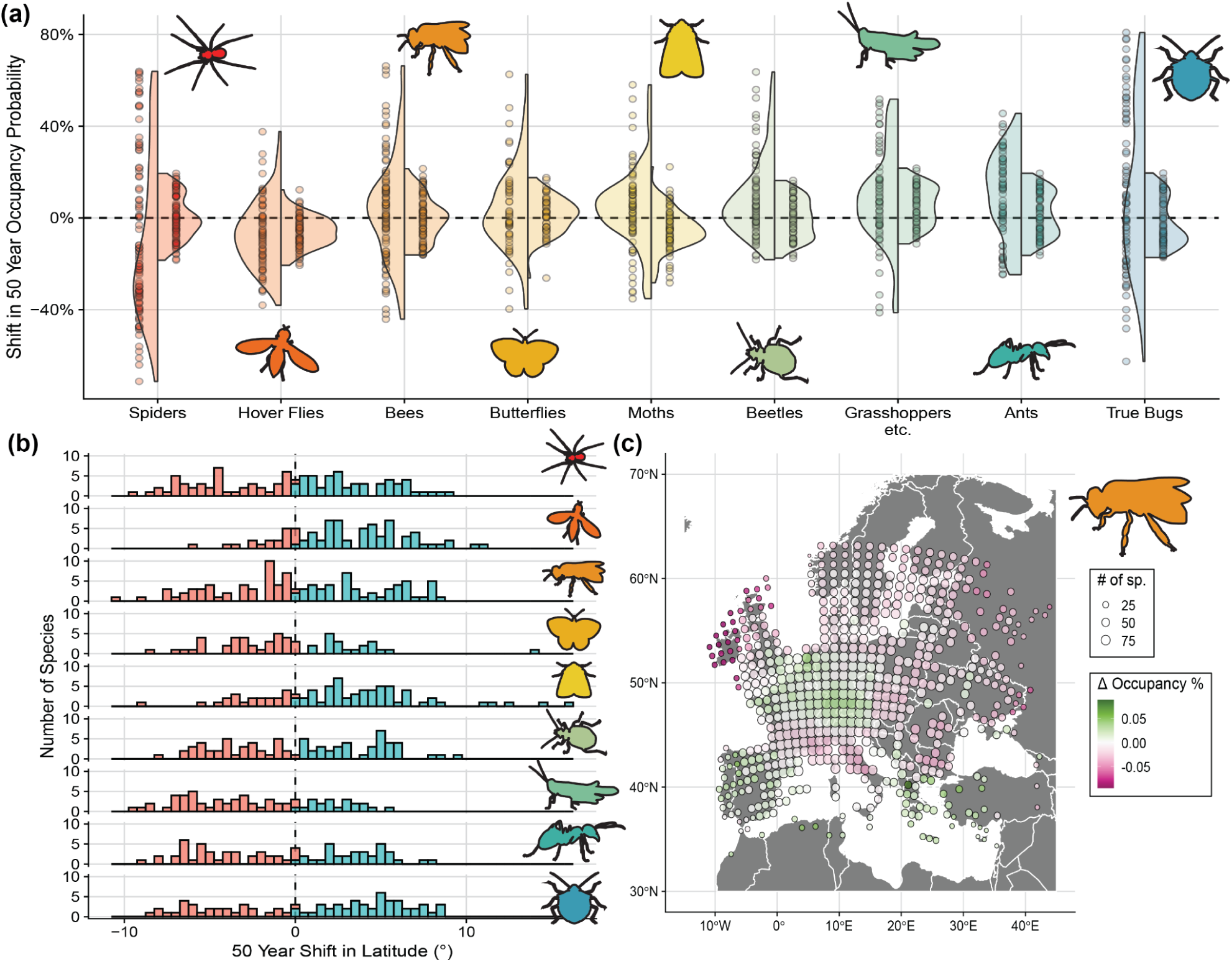
Panel (a) illustrates the estimates of 50-year shifts in occupancy probability across species (points representing species averages) and groups (shaded distributions). The left-hand distribution represents predictions from the time-only model. In contrast, the right-hand distribution represents predictions from the model based on environmental variables (climate and land-use change). Groups are ordered by mean trend from the time-only model, from those with the most decline to those with the most increase. Panel (b) categorises the number of species within each group by how far they have shifted in their latitudinal occupancy in decimal degrees from the environmental model. Finally, panel (c) illustrates the mean occupancy shift for all bees using the environmental model, where the size of the point indicates the number of species modelled at that location.

Spiders and hover flies had a large number of species in decline across the time-only and environmental models. Among spiders, *Hahnia montana*, *Oedotheorax fuscus*, and *Ceratinella scabrosa* are the most declining species. Notably, these spiders have not been assessed under the International Union for the Conservation of Nature (IUCN) Red List criteria, but they are all relatively widespread species, suggesting that common species may also be undergoing severe declines. Among hover flies, species like *Myathropa florea*, *Platycheirus manicatus*, and *Brachypalpus valgus* exhibit the most substantial decreases, yet these species have all been assessed as Least Concern^38^. Likewise, as for spiders, the inclusion of the first two species in this list is particularly alarming, as they have been considered quite common across their entire range, underscoring the potential for insect declines to cut across both rare and common taxa^39^. Other taxa also appear to be declining across our analysis. These include the red-backed furrow bee (*Lasioglossum laevigatum*), marbled skipper (*Carcharodus lavatherae*), and predatory bush cricket (*Saga pedo*)^40–42^.

In contrast to these declining species, several species are faring better over the 50-year reference period. Grasshoppers, ants, and true bugs appear to be among the groups with many species increasing in occupancy probability, underpinning the overall increase in their respective groups (Fig. 1a). Some notable species on the rise include the European firebug (*Pyrrhocoris apterus*), musk beetle (*Aromia moschata*), and heath fritillary (*Melitaea athalia*) (Fig. 2). Potential pest species, including the migratory locust, *Locusta migratoria*, are also increasing in occupancy probability (+41%), underscoring the potential for global change drivers to alter the distribution and prominence of potentially deleterious species.

**Figure 2.**
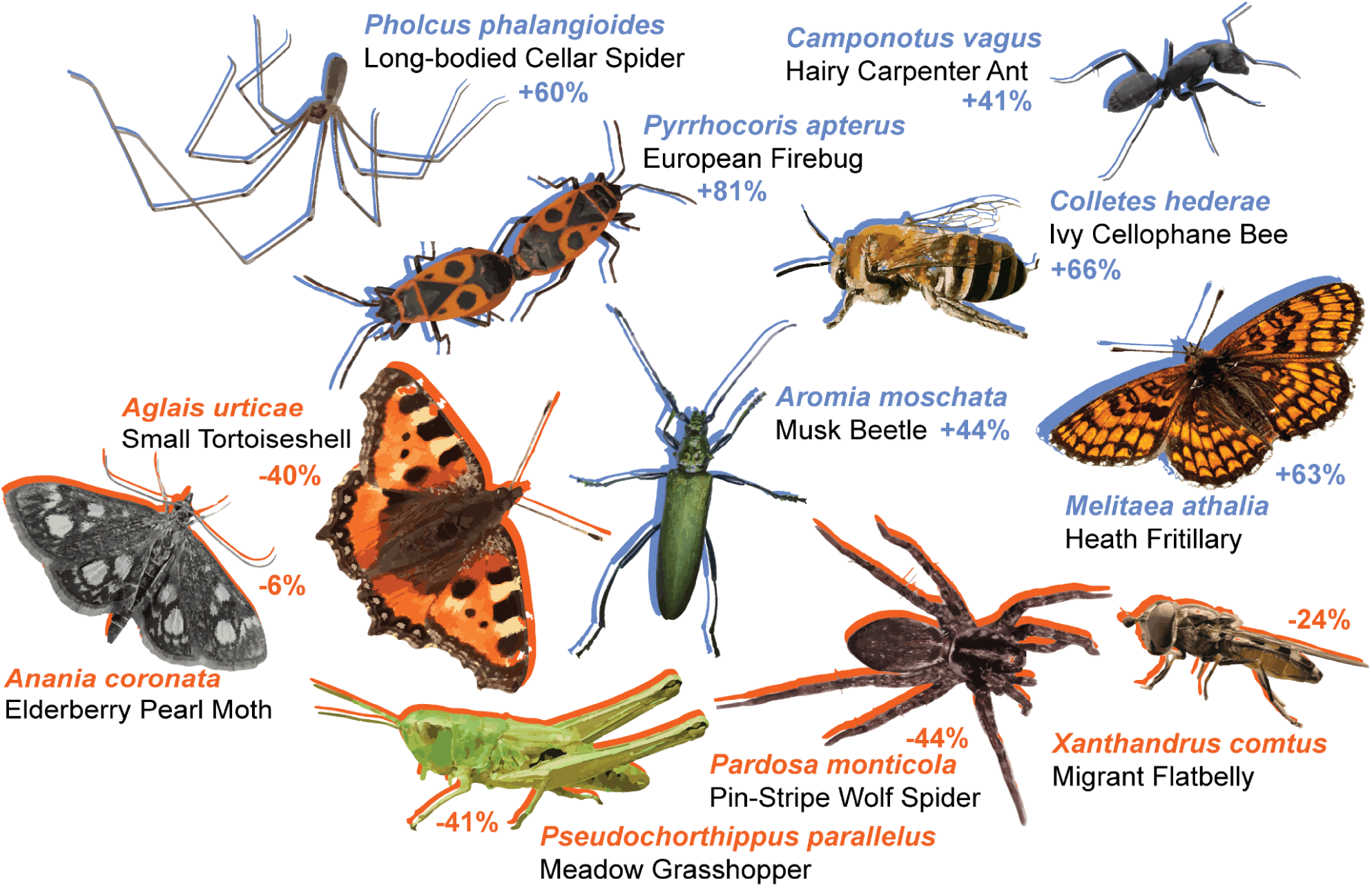
Selected arthropod winners and losers over the 50-year reference period. Images were obtained from iNaturalist observations of species under CC0 license.

Generally, temperature and precipitation were important predictors of changes in occupancy. Each focal group had a variable response to changes in minimum temperatures (Fig. 3a-b), with some groups, such as hoverflies and bees, responding negatively to temperature increases. On the other hand, orthopterans had a strong positive relationship between temperature and occupancy probability. Shifts to arid conditions at a given locality typically predict declines among arthropod species on average (Fig. 3b). The only groups for which this was not true were for moths and, to a lesser extent, beetles, which exhibited declines in occupancy probability with increasing precipitation. This strong pattern across most groups is notable since, across Europe, there is elevated concern of expanding desertification, a longstanding issue that has been identified over the last several decades^43–45^. Recent syntheses indicate that about 10% of Europe is experiencing some degree of desertification, suggesting that changes in precipitation will continue to play a pivotal role in structuring arthropod biodiversity across the continent^45^.

**Figure 3.**
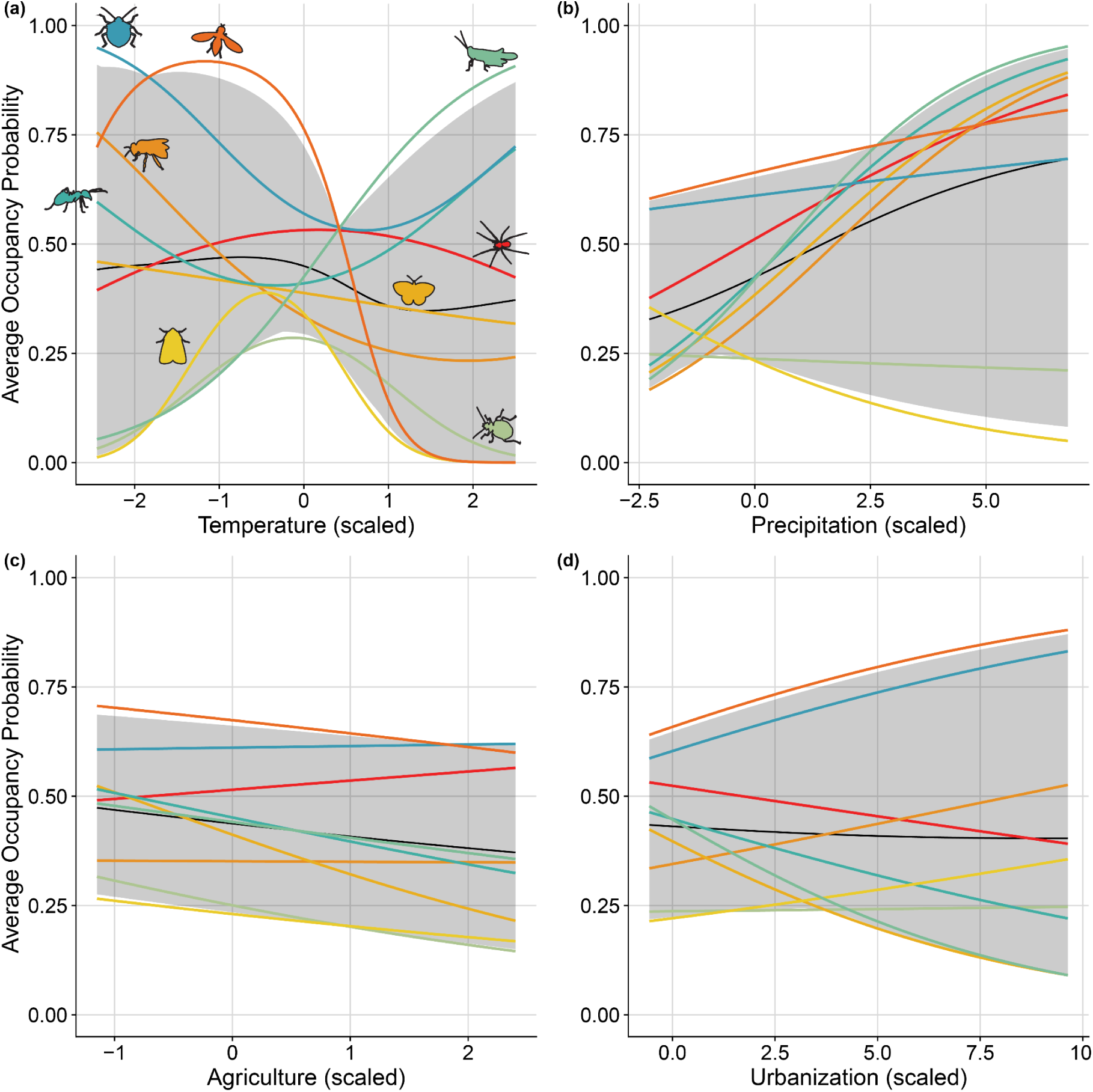
Average group-specific (colored) and overall average (black and shading) responses to the environmental factors considered in the study: (a) minimum temperature, (b) precipitation, (c) agricultural land cover, and (d) urbanised land cover. Shading represents the 95% Bayesian credible interval estimate across all groups.

Species-specific responses to land cover factors were also variable. Generally, there seemed to be a fairly universal decline related to increasing agricultural land cover on average occupancy probability, with occupancy probability declining across groups with increasing agricultural intensity (Fig. 3c). Urbanisation, on the other hand, had much stronger effects on average, with some groups showing increases in occupancy probability with greater urban cover (hoverflies and true bugs)(Fig. 3d). In contrast, others decreased in occupancy probability (bees, grasshoppers, and ants, for example). Some of the strongest responders to urbanisation included many butterflies known to inhabit high elevation/grassland habitats, such as *Spialia sertorius* and *Erebia cassioides* (both negatively impacted by urbanisation), and *Meliscaeva cinctella* (positively impacted by urbanisation), a hover fly commonly found in hedgerows and parks, feeding on aphids as larvae.

Clear winners and losers also emerged when considering species traits. Species with higher average temperature of occurrence (i.e., warm-tolerant species) and lower average precipitation of occurrence (i.e., arid-tolerant species) seemed to fare better across most groups (Fig. 4a). These results further reinforced our original occupancy analysis, which suggests that warmer and drier conditions are becoming more commonplace, resulting in occupancy decline for many species^45^.

**Figure 4.**
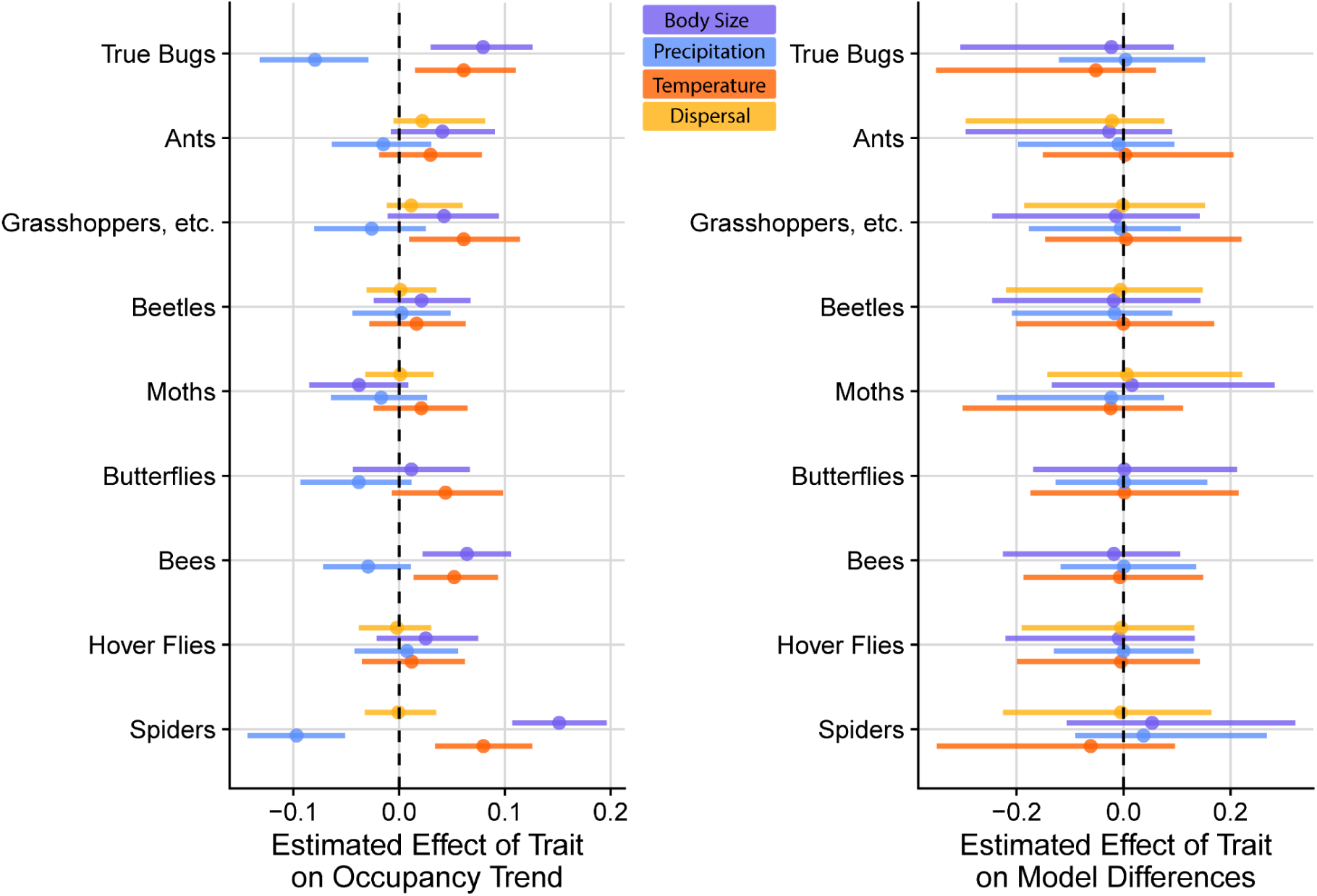
Modelled effect estimates for the species’ mean temperature of occurrences, mean precipitation of occurrences, body size, and dispersal capacity in predicting the average occupancy trend by the time-only model. Panel (b) uses the same traits in an attempt to explain the log-ratio difference in sensitivity between time-only and environmentally modelled trends. Error bars represent 95% Bayesian credible intervals.

Larger arthropod species are performing better across all groups, except for moths, where larger species exhibit a slightly more pronounced decline in occupancy probability (Fig. 4a). This finding contradicts our expectations and some prior simulation and empirical results on traits and extinction risk. For example, Chichorro *et al*. found that the response of extinction risk to body size was inconsistent across taxa and often not significant^25,46^. We hypothesise that these larger-bodied arthropods may be buffered against dessication. This interpretation meshes well with our general findings on the importance of precipitation in structuring arthropod occupancy in Europe; however, although this hypothesis has a physiological basis, it needs to be stringently tested in future work^47,48^. Contrary to our initial expectations, dispersal capacity was not strongly related to occupancy trends (Fig. 4a). While possibly indicative of a biological pattern, the relatively coarse spatial scale of our analysis (100km, with an average change of 1km/year) may limit our ability to detect a discernible effect of dispersal on overall occupancy trends. Future analyses that focus on scale should be conducted to determine whether this trait is important at finer resolutions. The relationship between species traits and the log-ratio between the change in occupancy of the time-only and environmental models revealed that for most groups, no single trait predicted the difference (Fig. 4b). Put simply, this means there is no obvious relation between particular combinations of traits, whether environmental predictors explain the modelled changes through time.

A caveat to the present study is that we limited our models to species with at least 250 reported occurrences over the 50-year reference period. This means that our predictions are based on the reporting of relatively common species across Europe. Lesser-known or rare species are certainly underrepresented in our dataset. Already rare species may be experiencing severe declines, although recent work also indicates that abundant species may make up a significant proportion of species composing global insect declines^39^. Our relatively coarse-grained analysis may also obscure local trends that are more relevant to range-restricted or rare species. Despite this, we believe our work represents a robust synthesis of trends across arthropod biodiversity within the continent.

## RECOMMENDATIONS

Although Europe, in general, and the European Union, in particular, are at the forefront of biodiversity conservation, arthropods have, in some respects, taken a backseat to more charismatic taxa^49,50^. This is especially true considering the magnitude of biological diversity represented by the group. We now have ample evidence that protected species and their habitats are only the tip of the iceberg in combating global declines in arthropods, underscoring the need for the urgent adoption of novel data, approaches, and results to inform further action to address longstanding challenges in arthropod conservation across Europe.

First, the vast amounts of data being produced must be reflected by conservation priorities and actions as soon as they are made available. Continued support of long-term arthropod monitoring programs is paramount. This can include existing schemes such as the European Butterfly Monitoring Scheme (EMBS)^51^ and the EU Pollinator Monitoring Scheme^52^. Both provide examples of robust frameworks for implementing wide-scale monitoring of terrestrial invertebrates. The emergence of automated monitoring methods will simplify processes for numerous taxa, and their increasingly rapid development and deployment will generate unprecedented amounts of data in relatively short periods^53–55^. The Habitats Directive, the Restoration Regulation, EU Nature Restoration Law, national legislation, as well as grassroots conservation initiatives, should be flexible enough to adapt to these emerging data streams; otherwise, they risk rapid obsolescence.

Following on this, conservation assessments of arthropods, which are challenging to undertake due to high species diversity, the difficulty of structured monitoring, incomplete knowledge, and the need for taxonomic expertise, must adapt and adopt novel approaches^27,56^. Occupancy-detection models provide a putative framework for addressing some of these challenges, and they are remarkably robust to the biases present in opportunistic data, provided a few key assumptions are met^57^. We recommend that conservation professionals seriously consider these models for assessing and prioritising species from diverse groups. They are already being proposed as a triage system for assessing hyperdiverse taxa and prioritising species for further investigation and conservation action. Finally, alternative data sources, such as species traits, should be used to infer species extinction risk in the absence of sufficient occurrence data across space and time^25,58^. Here, the cautionary principle should be adopted, and traits may provide a simple, yet effective, surrogate for population trends as demonstrated here and elsewhere.

The recent history of arthropod decline in Europe is characterised by winners and losers rather than by ubiquitous declines (see also Ogan *et al.* 2022). A critical distinction in our study is that we modelled occupancy trends rather than abundance. Therefore, species we find to have stable distributions may experience declines in abundance within those ranges. Any decline in arthropod species is cause for global alarm, as the functional roles of many remain poorly understood. Thus, the loss of arthropod biodiversity spells the loss of potentially unknown ecological interactions and ecosystem services. Our analysis highlights the inherent complexity and the need to decipher further declines in an era of increasing, intersecting global change drivers. In Europe, increased attention should be paid to the decline of spiders and hoverflies in particular, as these groups appear to be the most threatened overall; however, representative “losers” in each group are likely in need of both increased research and conservation attention. The need to establish a triage system for the vast diversity of arthropod species is ever-present as we grapple with the complexity of declines.

## METHODOLOGY

### Scope of Study

We sought to analyse trends in 50-100 species across 11 arthropod groups (mostly orders) over 50 years (1970-2019). Geographically, we constrained our analysis to a bounding box that encompassed the vast majority of European biodiversity data in GBIF (34° to 73° latitude and -17° to 43° longitude). We calculated the historical occupancy probability for all species within each taxonomic focal group every five years (referred to as “occupancy intervals,” during which each year constitutes a “visit interval”). Spatially, we calculated our model’s occupancy and detection components at a 100-by-100-kilometre grid resolution across the bounding box to ensure sufficient data were available in GBIF for each period.

### Environmental Predictor Data

We obtained four environmental predictors for our occupancy-detection models to inform our calculations of occupancy probability. First, we obtained monthly average surface temperature and precipitation data from Copernicus Climate^59^. We then averaged these climatic variables into the spatial grids for each occupancy interval. Second, we obtained land cover data from LUCAS LUC historical and future land cover dataset^60^. Since historical reconstructions of land cover data are available only on a decadal timescale, we used the same land cover information for our five-year historical periods. In other words, since data were only available for 1970-1979, we used the same land cover data for our 1970-1974 and 1975-1979 occupancy intervals. This treatment of the land cover data may reduce our statistical power in delineating land cover responses.

### Species Selection and Occurrence Data

Species within each broader taxonomic group were generally non-randomly selected to represent a diversity of ecological and life-history traits. Only species with at least 250 records in GBIF were selected, as models tend to perform better with a minimum number of occurrence records. In total, we retained 100 spider species, 79 hover flies, 100 bees, 56 butterflies, 80 moths, 80 beetles, 65 orthopterans, 73 ants, and 78 true bugs. Occurrences were filtered to include those that occurred on land (removing erroneous points over the ocean or other large bodies of water).

### Occupancy-Detection Modelling

We specified multispecies, multiseason, and static occupancy-detection models for each taxonomic focal group (two models per group)^57,61^. One model included all aforementioned environmental predictor variables to generate spatiotemporally explicit estimates of occupancy probability at the species level. The second model estimated only overall species-level trends using a linear and quadratic effect of occupancy interval as the sole predictor variable. Thus, it did not provide site-specific estimates of occupancy change based on environmental factors, but instead provided a mean estimate of occupancy probability for each interval over the duration of our study. We refer to this as our time-only model and use it to compare estimates of change from time alone with estimates that are captured by climate and land-use change covariates. The occupancy component of the time-only model was defined by Equation 1 below:

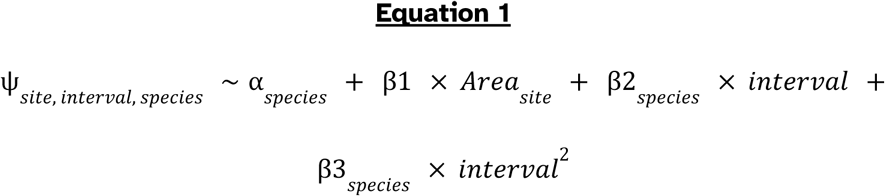

Where α _𝑠𝑝𝑒𝑐𝑖𝑒𝑠_ is the species-specific occupancy intercept, β1 is a fixed-effect of terrestrial surface area, β2_𝑠𝑝𝑒𝑐𝑖𝑒𝑠_ is the species-specific effect of occupancy interval, and β3_𝑠𝑝𝑒𝑐𝑖𝑒𝑠_ is the species-specific quadratic effect of occupancy interval. Conversely, the occupancy component of the environmental model was defined by Equation 2 below:

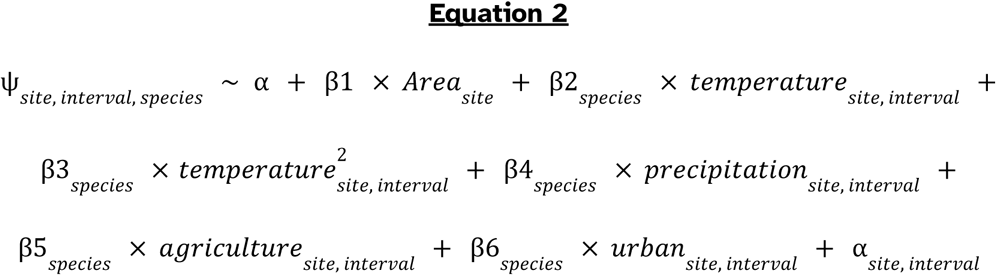

where α is a global intercept term, β1 is the fixed-effect of terrestrial surface area, β2_𝑠𝑝𝑒𝑐𝑖𝑒𝑠_is the species-specific effect of temperature,

β3_𝑠𝑝𝑒𝑐𝑖𝑒𝑠_ is the species-specific quadratic effect of temperature, β4_𝑠𝑝𝑒𝑐𝑖𝑒𝑠_ is the species-specific effect of precipitation, β5_𝑠𝑝𝑒𝑐𝑖𝑐𝑒𝑠_ is the species-specific effect of agricultural land cover, β6_𝑠𝑝𝑒𝑐𝑖𝑒𝑠_ is the species-specific effect of urban land cover, and α_𝑠𝑖𝑡𝑒, 𝑖𝑛𝑡𝑒𝑟𝑣𝑎𝑙_ is a site-by-interval intercept term. The global intercept, rather than the species-specific intercept, was used in this model due to convergence issues. Both models shared the same definition of the detection component, defined by Equation 3 below:

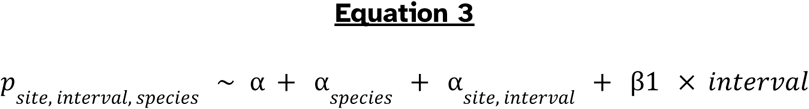

All models were run on either the Science Information Center (CSC) research computing infrastructure, personal desktop computers, or the University of Florida HiPerGator cluster for at least 25,000 sampling iterations, with 5,000 iterations used as burn-in. Thinning was performed by a factor of 20 across four chains, yielding 4,000 samples from the posterior. MCMC chains were run using Just Another Gibbs Sampler (JAGS)^62^. Additional iterations were added based on an assessment of convergence. We assessed the convergence of our models by examining the Gelman-Rubin diagnostic metric (“r-hat”) using an upper threshold of 1.2^63^. If the diagnostic metric exceeded this threshold, we also visually inspected the trace plots for each model using the R package “MCMCvis”^64^. All trace plots indicated that there was appropriate convergence of the model parameters, and only 0.4% model parameters required this more detailed inspection. MCMC trace plots are included in the supplemental material for all models.

### Trait Analysis

We collected trait data for species from each taxonomic focal group to determine how well certain species traits predict occupancy trends through time. We specified the following traits for inclusion in our post-hoc analysis: average temperature and precipitation at occurrence sites, body size, and dispersal capacity. These traits were aggregated from multiple published and unpublished sources^65^ and from averaged measurements of museum specimens. Further information about the types of measurements taken from museum specimens is provided in the Supplementary Material.

Using the trait data, we specified a taxonomic random-effects model in the R package “brms”^66^, which included simple linear effects of the traits above as single-trait models and an implicitly nested random-effects structure that accounted for taxonomy. Models were run across four chains for a total of 10,000 iterations. Our general model form is specified by Equation 4 below:

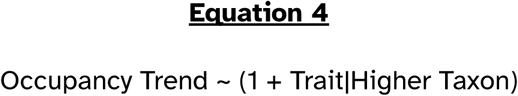

We used the same equation to examine differences between models, using the calculated difference as the response variable in place of occupancy trends. We calculated the log-ratio between the environmental and time-only models (absolute predicted trends) to serve as a response metric for the difference in sensitivity between models. Log-ratio values greater than zero indicate that the environmental model predicts a larger occupancy shift, while log-ratio values less than zero indicate that the time-only model predicts a larger shift.

## Authorship Contributions

VS and PC conceived and designed the original study. All authors contributed to selecting the appropriate species for analysis, aggregating relevant trait information, and critically assessing model results based on their respective taxonomic expertise. VS processed and analyzed the data. LMG and PC provided helpful feedback on the modeling process. VS wrote the initial manuscript, and all authors provided data, feedback, and edits throughout the drafting process. All authors approved the manuscript for submission and publication.

## Funding

The Finnish Museum of Natural History and the University of Helsinki (PC) provided funding for VS to travel to Finland to undertake this collaborative work. VS was also supported by the Department of Biology at Georgetown University, a Georgetown University Graduate Student Government STEM for the Public Good Award, a United States National Science Foundation Graduate Research Fellowship (#1937959), a David H. Smith Postdoctoral Conservation Research Fellowship, and start-up funding from the Florida Museum of Natural History, University of Florida, while conducting this research.

## Acknowledgements

Additionally, VS would like to thank the Fulbright Finland Foundation and Fulbright Program for a study/research grant in 2017-2018 that began a collaboration between VS and PC. The authors also wish to acknowledge CSC - IT Center for Science, Finland, and the University of Florida’s HiPerGator system for a generous allotment of computational resources used in this work.

## Use of Generative AI

The authors used Grammarly while writing this manuscript, which uses a generative AI model to suggest improvements to the text for grammar and clarity.

## Notes

### Competing Interest Statement

The authors have declared no competing interest.

