## Supplementary material for "Winners and Losers Among European Arthropods over the Last Half Century of Global Change": ants_timeTrace.pdf

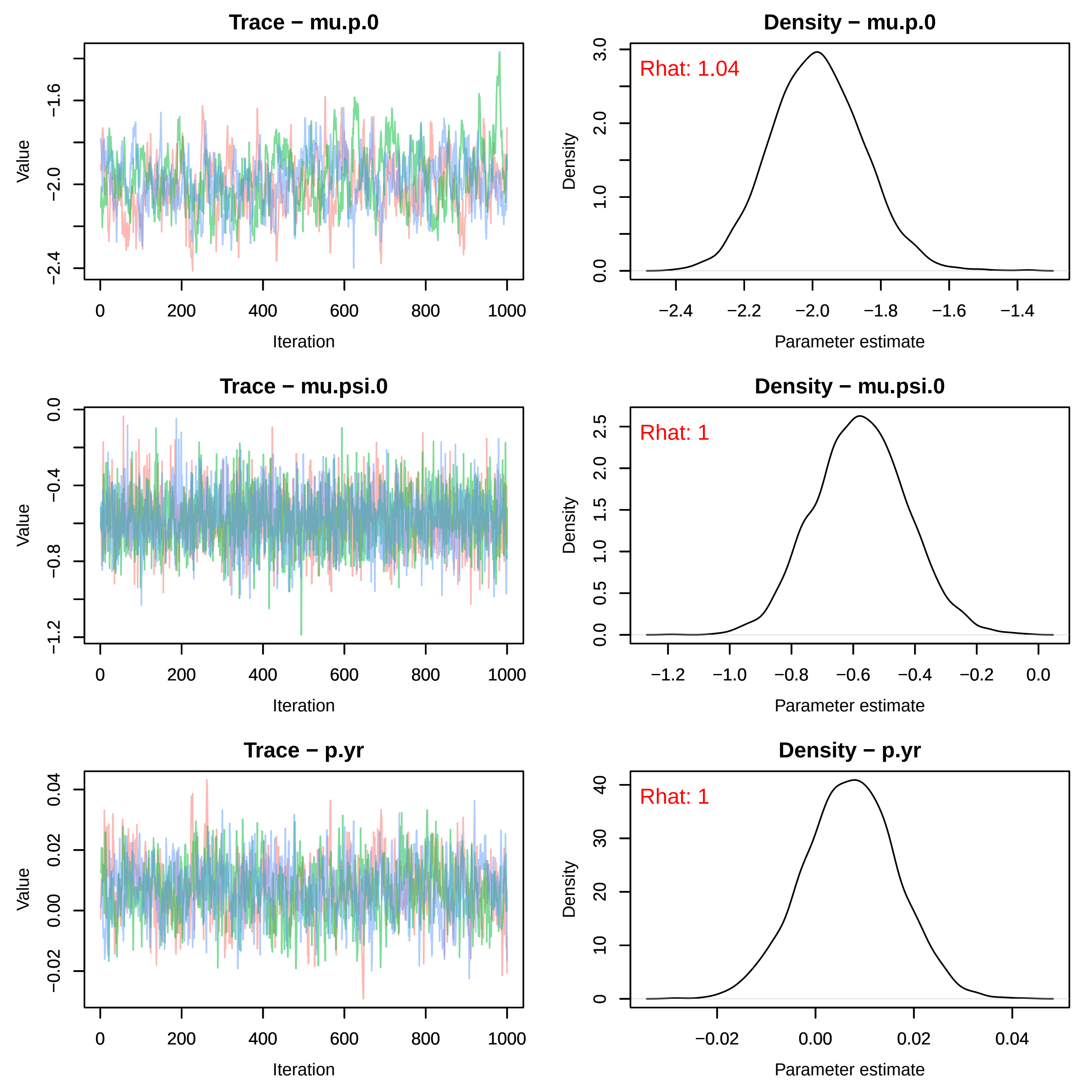
Trace − mu.p.0
Value
Iteration
−2.4
−2.0
−1.6
0
200
400
600
800
1000
Density − mu.p.0
Rhat: 1.04
Density
Parameter estimate
0.0
1.0
2.0
3.0
−2.4
−2.2
−2.0
−1.8
−1.6
−1.4
Trace − mu.psi.0
Value
Iteration
−1.2
−0.8
−0.4
0.0
0
200
400
600
800
1000
Density − mu.psi.0
Rhat: 1
Density
Parameter estimate
0.0
0.5
1.0
1.5
2.0
2.5
−1.2
−1.0
−0.8
−0.6
−0.4
−0.2
0.0
Trace − p.yr
Value
Iteration
−0.02
0.00
0.02
0.04
0
200
400
600
800
1000
Density − p.yr
Rhat: 1
Density
Parameter estimate
0
10
20
30
40
−0.02
0.00
0.02
0.04

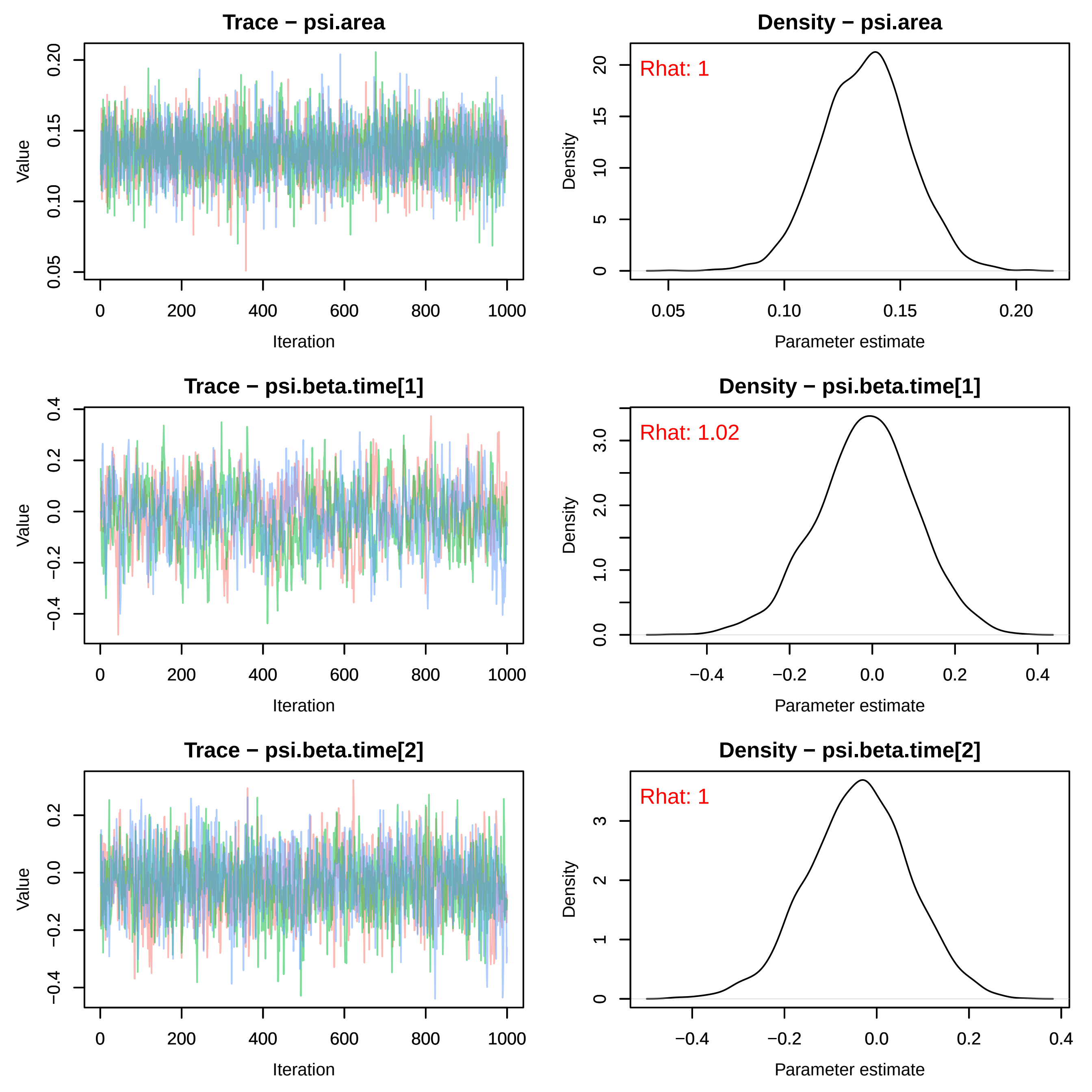

Trace − psi.area
Density − psi.area
Rhat: 1
Trace − psi.beta.time[1]
Density − psi.beta.time[1]
Rhat: 1.02
Trace − psi.beta.time[2]
Density − psi.beta.time[2]
Rhat: 1
Value
Iteration
Density
Parameter estimate

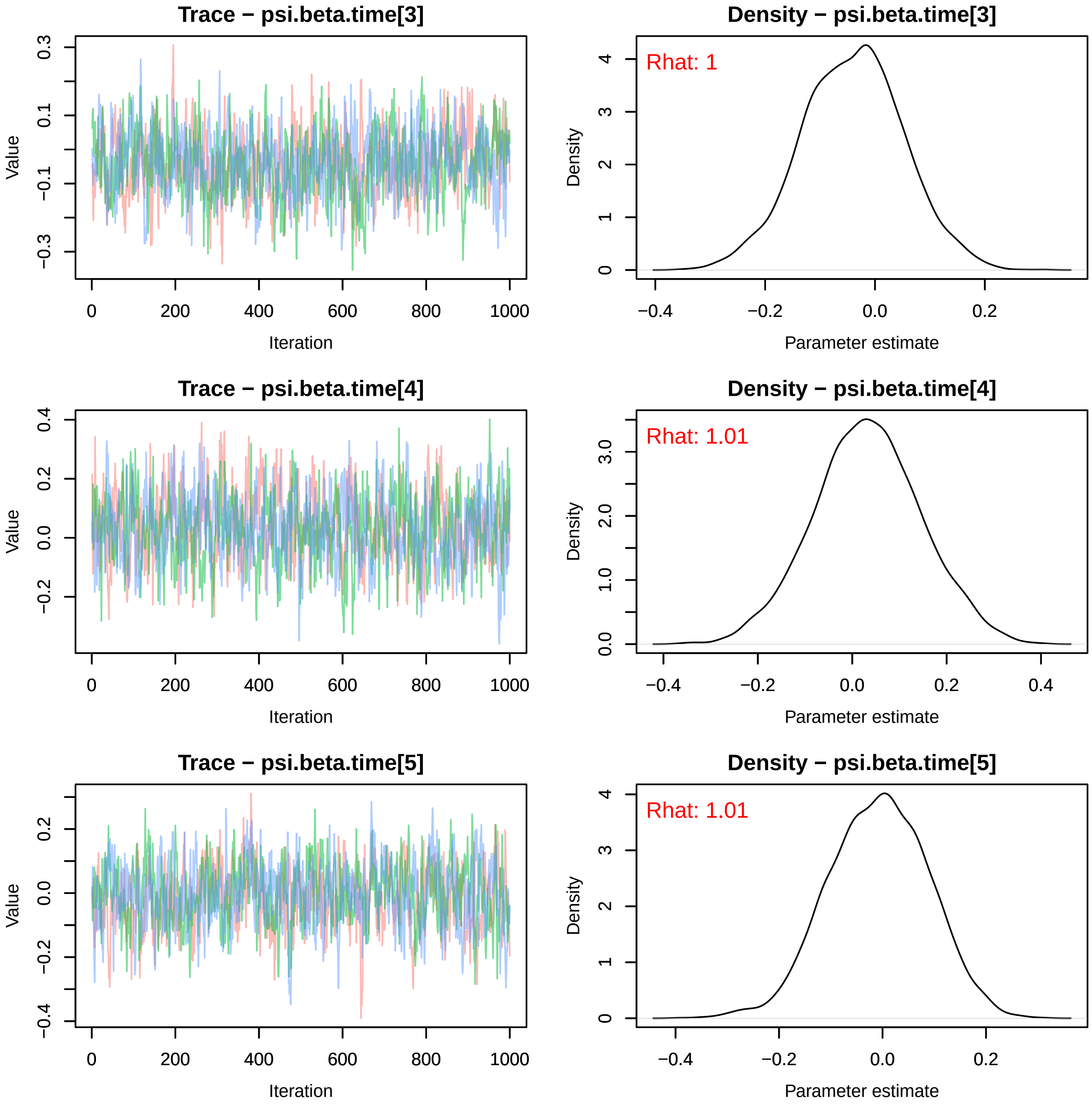
Trace − psi.beta.time[3]
Density − psi.beta.time[3]
Rhat: 1
Trace − psi.beta.time[4]
Density − psi.beta.time[4]
Rhat: 1.01
Trace − psi.beta.time[5]
Density − psi.beta.time[5]
Rhat: 1.01
Value
Iteration
Density
Parameter estimate

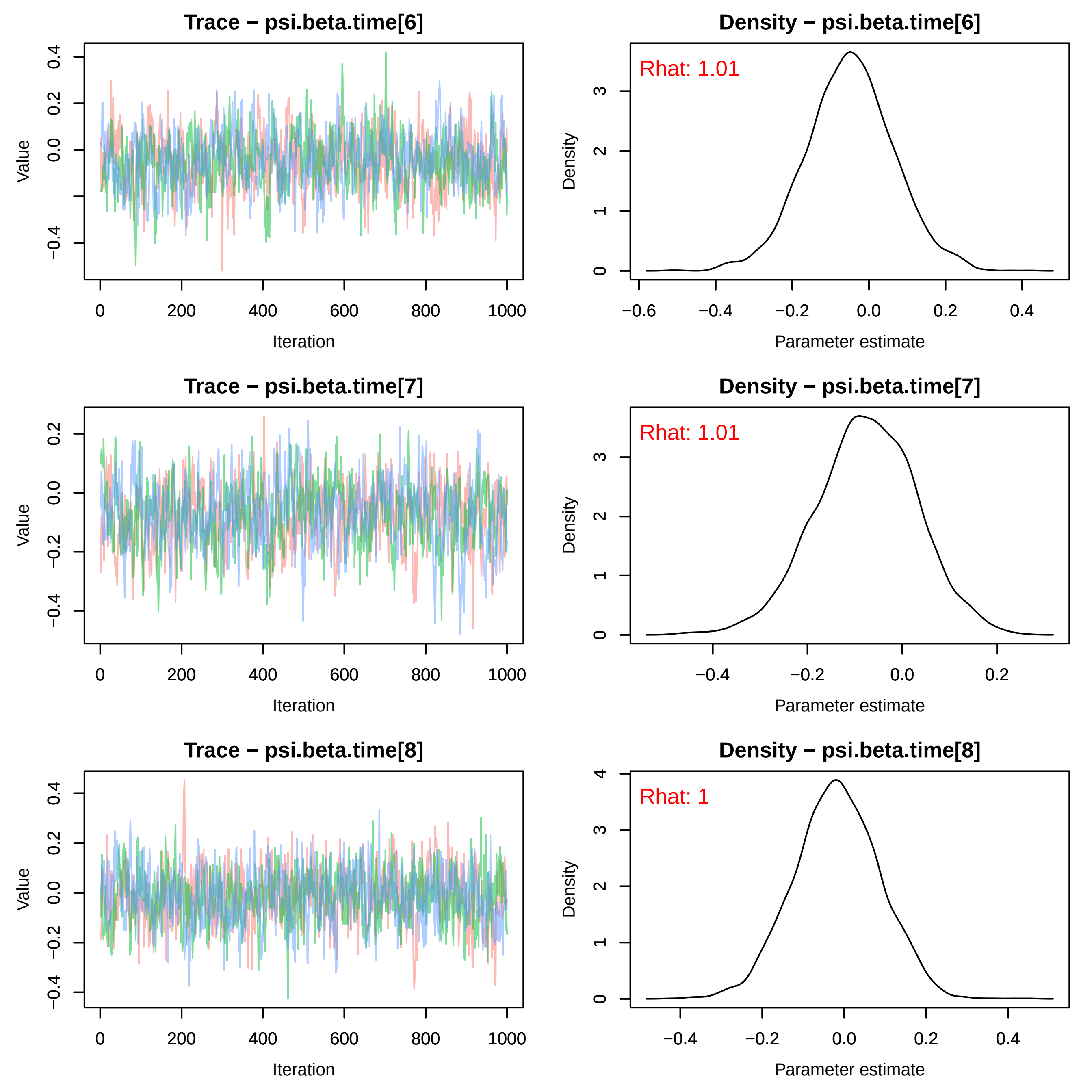
Trace − psi.beta.time[6]
Value
Iteration
Density − psi.beta.time[6]
Rhat: 1.01
Density
Parameter estimate
Trace − psi.beta.time[7]
Value
Iteration
Density − psi.beta.time[7]
Rhat: 1.01
Density
Parameter estimate
Trace − psi.beta.time[8]
Value
Iteration
Density − psi.beta.time[8]
Rhat: 1
Density
Parameter estimate

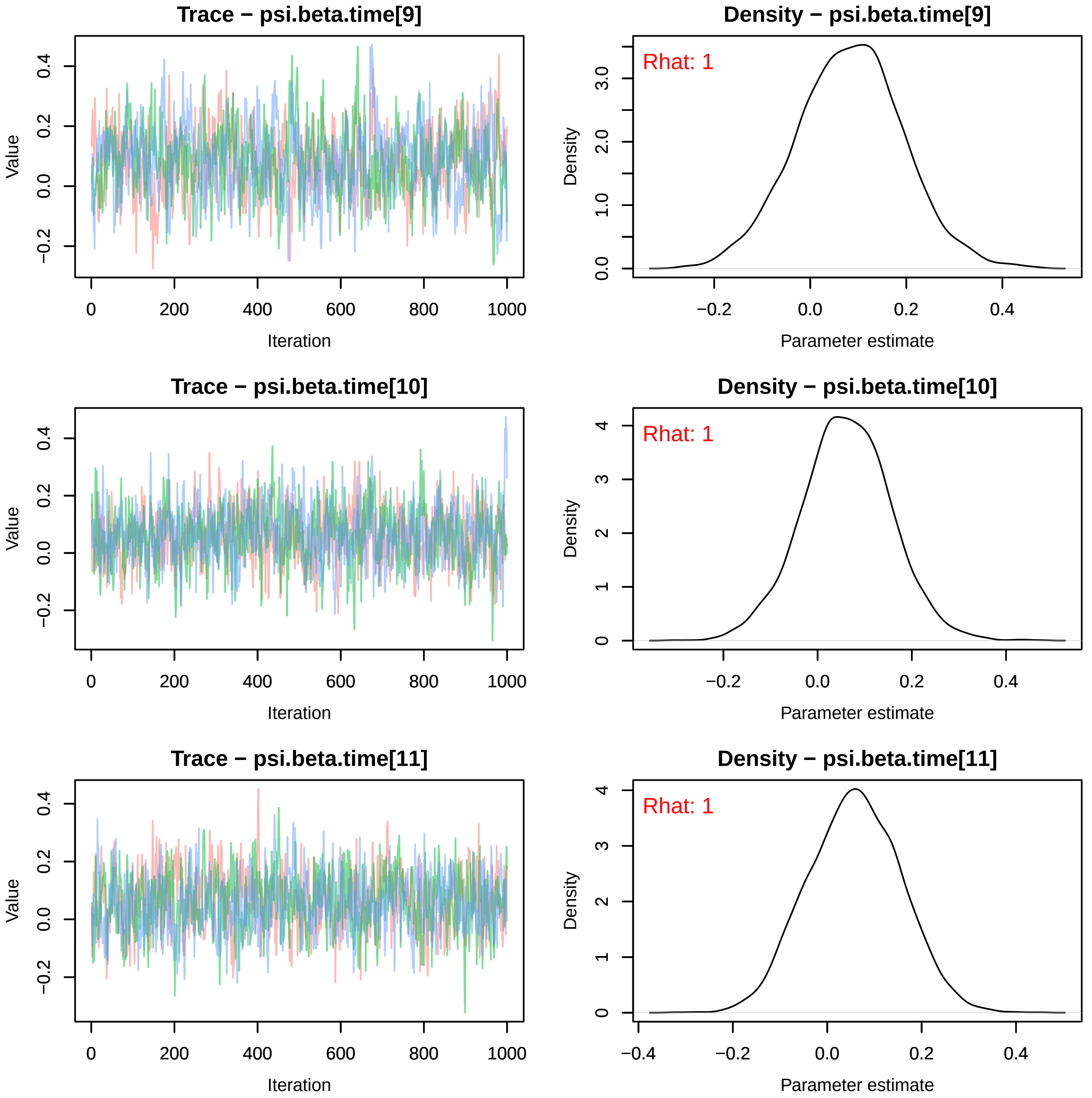

Trace − psi.beta.time[9]
Value
Iteration
Density − psi.beta.time[9]
Rhat: 1
Density
Parameter estimate
Trace − psi.beta.time[10]
Value
Iteration
Density − psi.beta.time[10]
Rhat: 1
Density
Parameter estimate
Trace − psi.beta.time[11]
Value
Iteration
Density − psi.beta.time[11]
Rhat: 1
Density
Parameter estimate

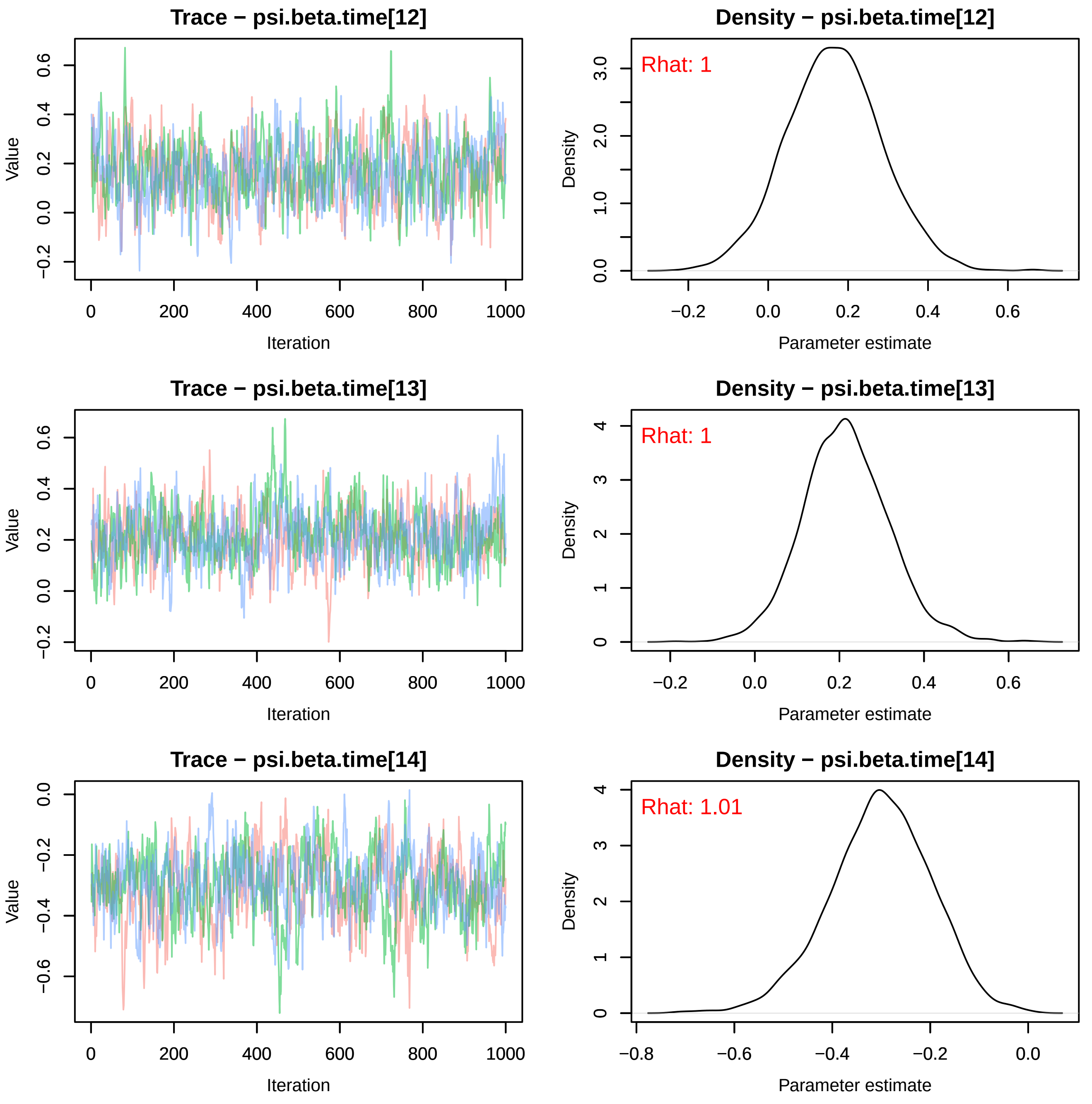
Trace − psi.beta.time[12]
Value
−0.2
0.0
0.2
0.4
0.6
0
200
400
600
800
1000
Iteration
Density − psi.beta.time[12]
Rhat: 1
Density
0.0
1.0
2.0
3.0
−0.2
0.0
0.2
0.4
0.6
Parameter estimate
Trace − psi.beta.time[13]
Value
−0.2
0.0
0.2
0.4
0.6
0
200
400
600
800
1000
Iteration
Density − psi.beta.time[13]
Rhat: 1
Density
0
1
2
3
4
−0.2
0.0
0.2
0.4
0.6
Parameter estimate
Trace − psi.beta.time[14]
Value
−0.6
−0.4
−0.2
0.0
0
200
400
600
800
1000
Iteration
Density − psi.beta.time[14]
Rhat: 1.01
Density
0
1
2
3
4
−0.8
−0.6
−0.4
−0.2
0.0
Parameter estimate

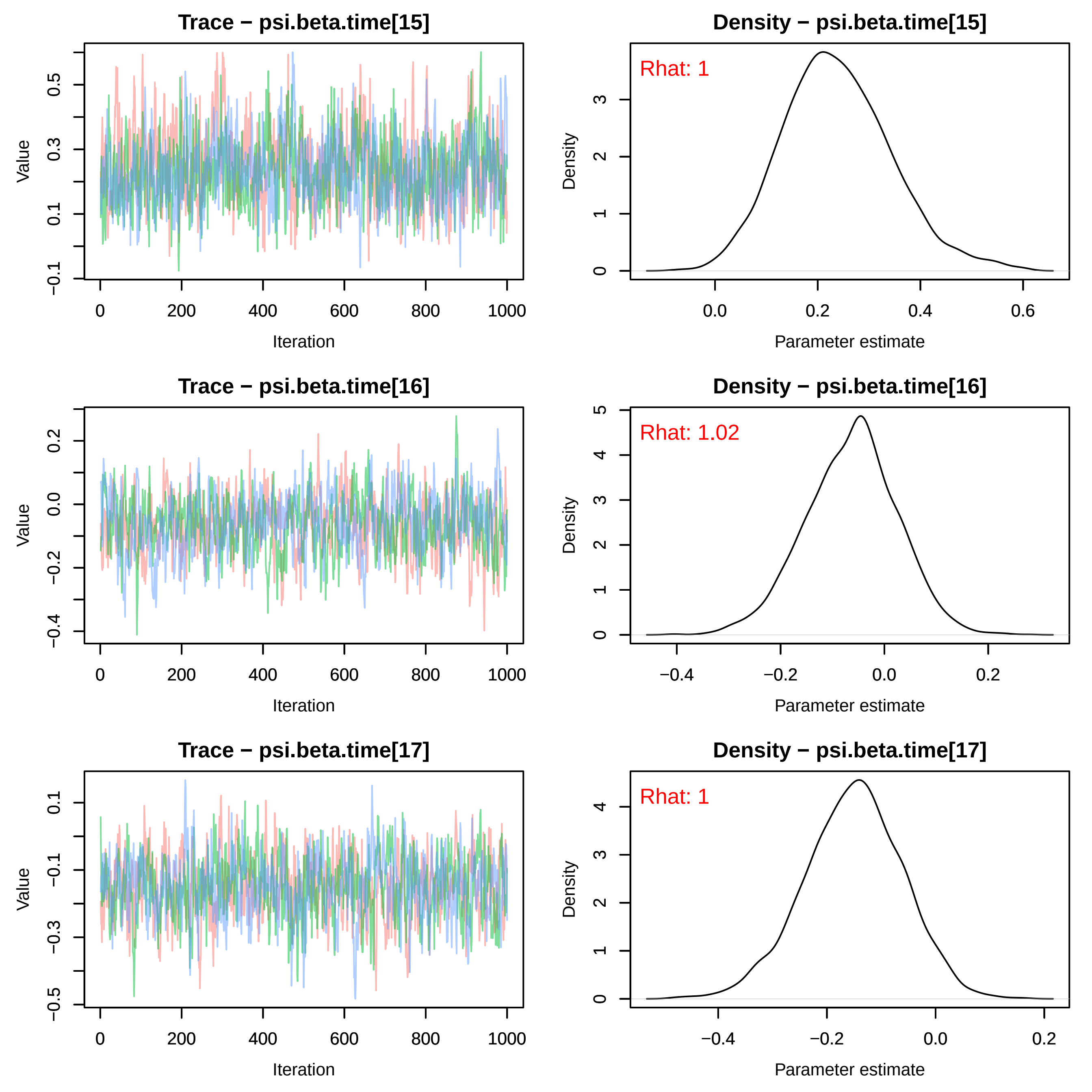
Trace − psi.beta.time[15]
Value
Iteration
Density − psi.beta.time[15]
Rhat: 1
Density
Parameter estimate
Trace − psi.beta.time[16]
Value
Iteration
Density − psi.beta.time[16]
Rhat: 1.02
Density
Parameter estimate
Trace − psi.beta.time[17]
Value
Iteration
Density − psi.beta.time[17]
Rhat: 1
Density
Parameter estimate

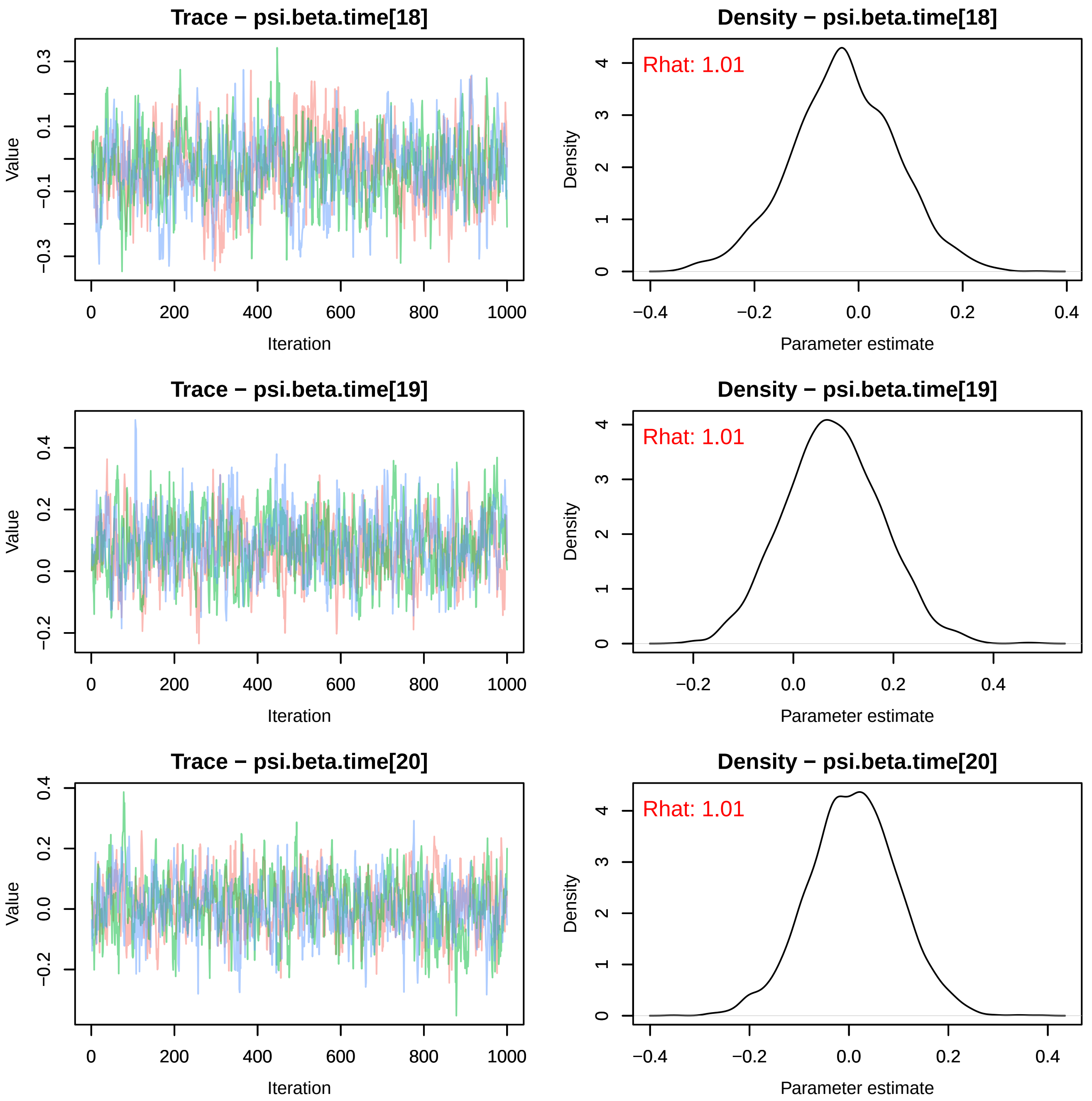
Trace − psi.beta.time[18]
Value
Iteration
0 200 400 600 800 1000
−0.3 −0.1 0.1 0.3
Density − psi.beta.time[18]
Rhat: 1.01
Density
Parameter estimate
−0.4 −0.2 0.0 0.2 0.4
0 1 2 3 4
Trace − psi.beta.time[19]
Value
Iteration
0 200 400 600 800 1000
−0.2 0.0 0.2 0.4
Density − psi.beta.time[19]
Rhat: 1.01
Density
Parameter estimate
−0.2 0.0 0.2 0.4
0 1 2 3 4
Trace − psi.beta.time[20]
Value
Iteration
0 200 400 600 800 1000
−0.2 0.0 0.2 0.4
Density − psi.beta.time[20]
Rhat: 1.01
Density
Parameter estimate
−0.4 −0.2 0.0 0.2 0.4
0 1 2 3 4

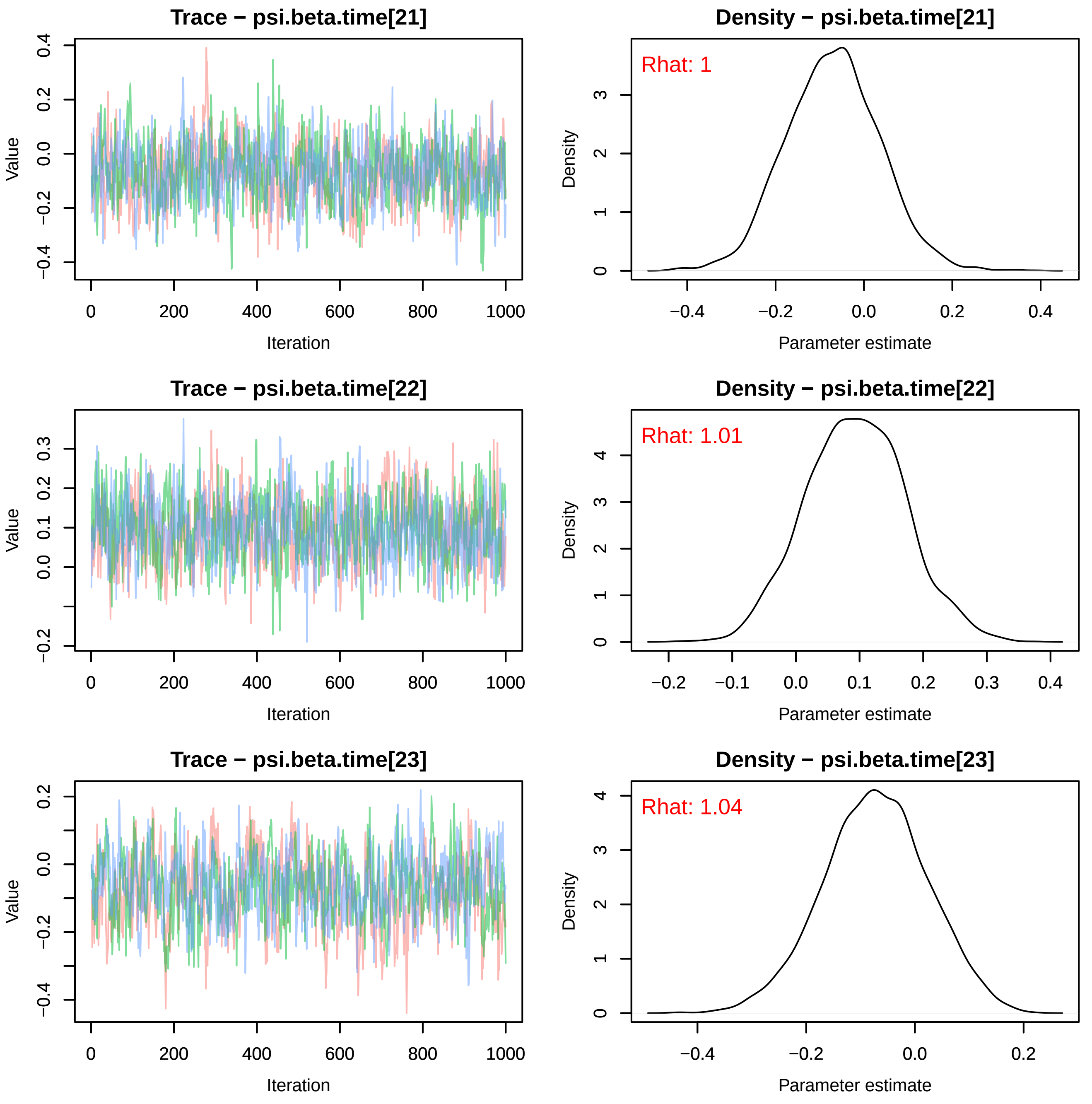

Trace − psi.beta.time[21]
Value
Iteration
0
200
400
600
800
1000
−0.4
−0.2
0.0
0.2
0.4
Density − psi.beta.time[21]
Rhat: 1
Density
Parameter estimate
−0.4
−0.2
0.0
0.2
0.4
0
1
2
3
Trace − psi.beta.time[22]
Value
Iteration
−0.2
0.0
0.1
0.2
0.3
Density − psi.beta.time[22]
Rhat: 1.01
Density
Parameter estimate
−0.2
−0.1
0.0
0.1
0.2
0.3
0.4
0
1
2
3
4
Trace − psi.beta.time[23]
Value
Iteration
−0.4
−0.2
0.0
0.2
Density − psi.beta.time[23]
Rhat: 1.04
Density
Parameter estimate
−0.4
−0.2
0.0
0.2
0
1
2
3
4

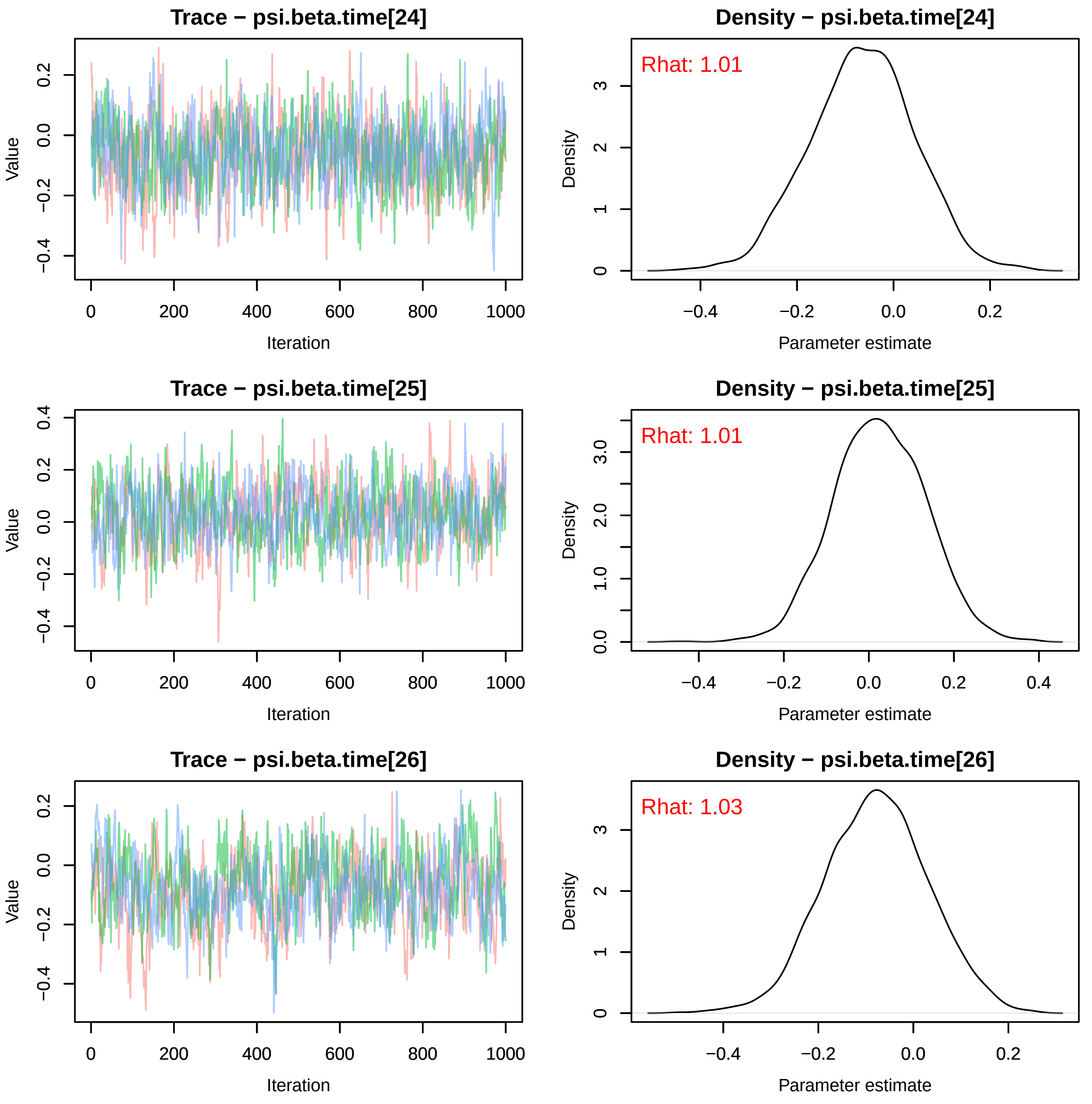
Trace − psi.beta.time[24]
Density − psi.beta.time[24]
Rhat: 1.01
Trace − psi.beta.time[25]
Density − psi.beta.time[25]
Rhat: 1.01
Trace − psi.beta.time[26]
Density − psi.beta.time[26]
Rhat: 1.03
Value
Iteration
Density
Parameter estimate
0
200
400
600
800
1000
−0.4
−0.2
0.0
0.2
0.4
1.0
2.0
3.0
1
2
3

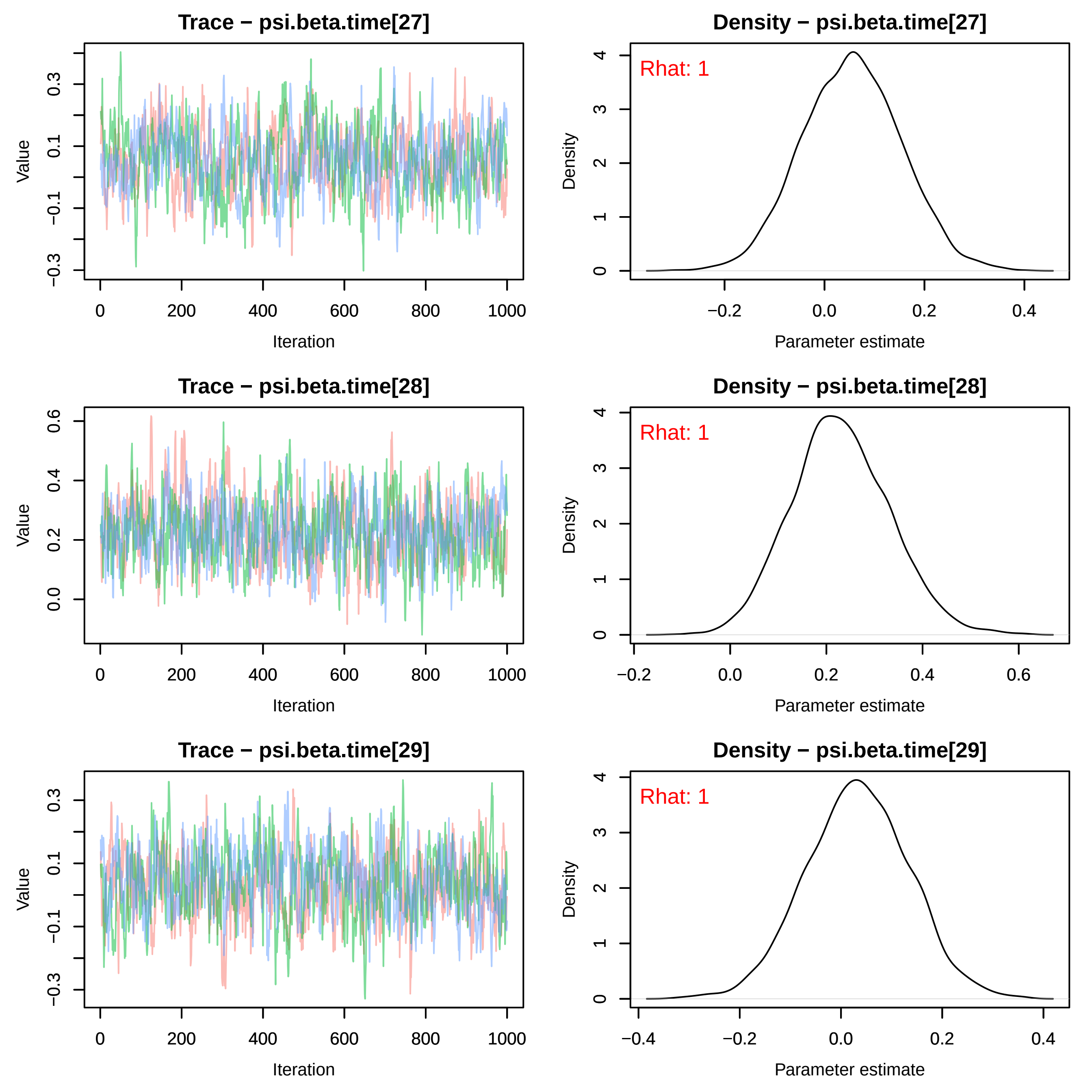
Trace − psi.beta.time[27]
Density − psi.beta.time[27]
Rhat: 1
Trace − psi.beta.time[28]
Density − psi.beta.time[28]
Rhat: 1
Trace − psi.beta.time[29]
Density − psi.beta.time[29]
Rhat: 1
Value
Iteration
Density
Parameter estimate

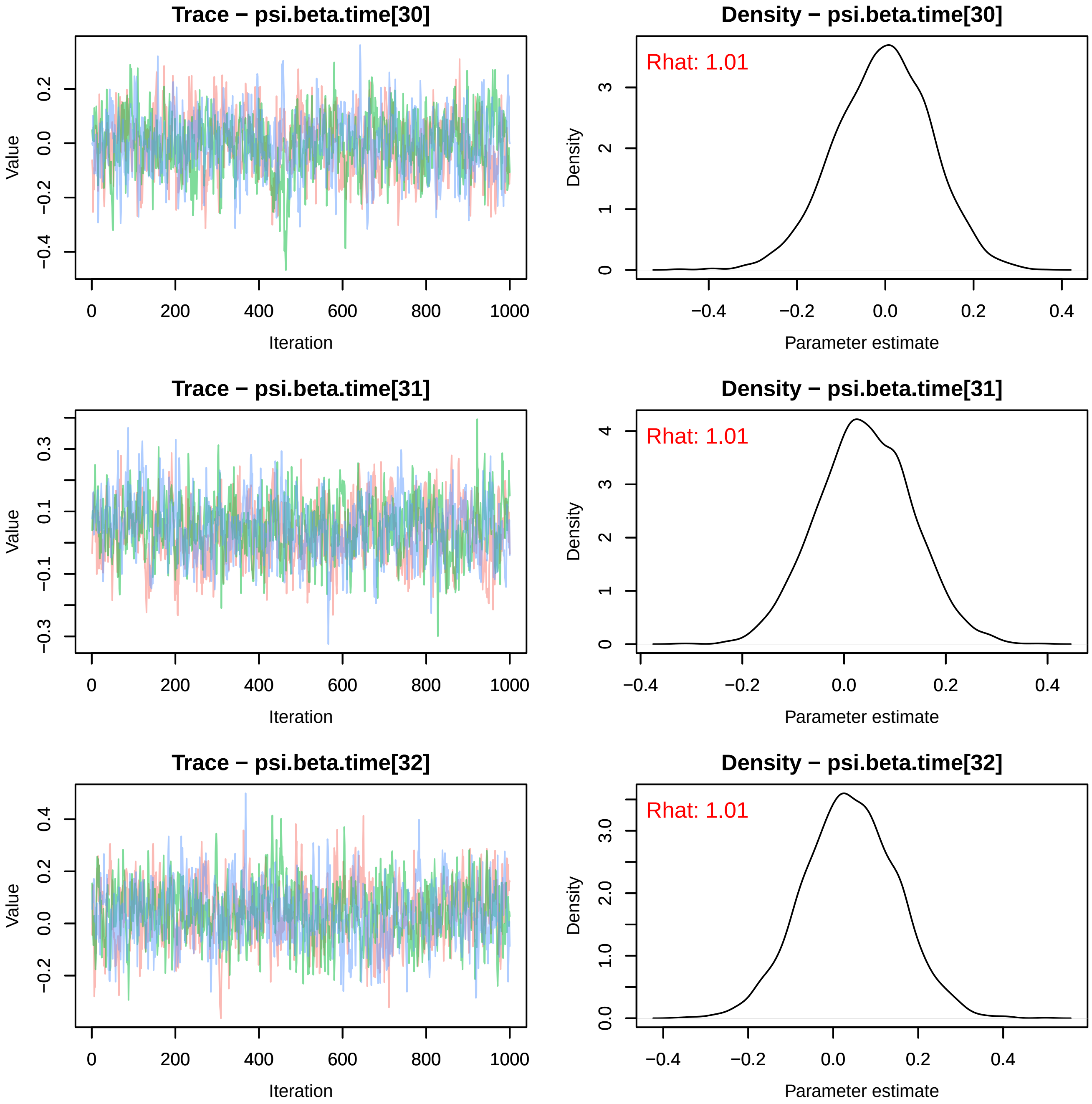
Trace − psi.beta.time[30]
Density − psi.beta.time[30]
Rhat: 1.01
Trace − psi.beta.time[31]
Density − psi.beta.time[31]
Rhat: 1.01
Trace − psi.beta.time[32]
Density − psi.beta.time[32]
Rhat: 1.01
Value
Iteration
Density
Parameter estimate
0
200
400
600
800
1000
−0.4
−0.2
0.0
0.2
0.4
−0.3
−0.1
0.1
0.3
1
2
3
4
1.0
2.0
3.0

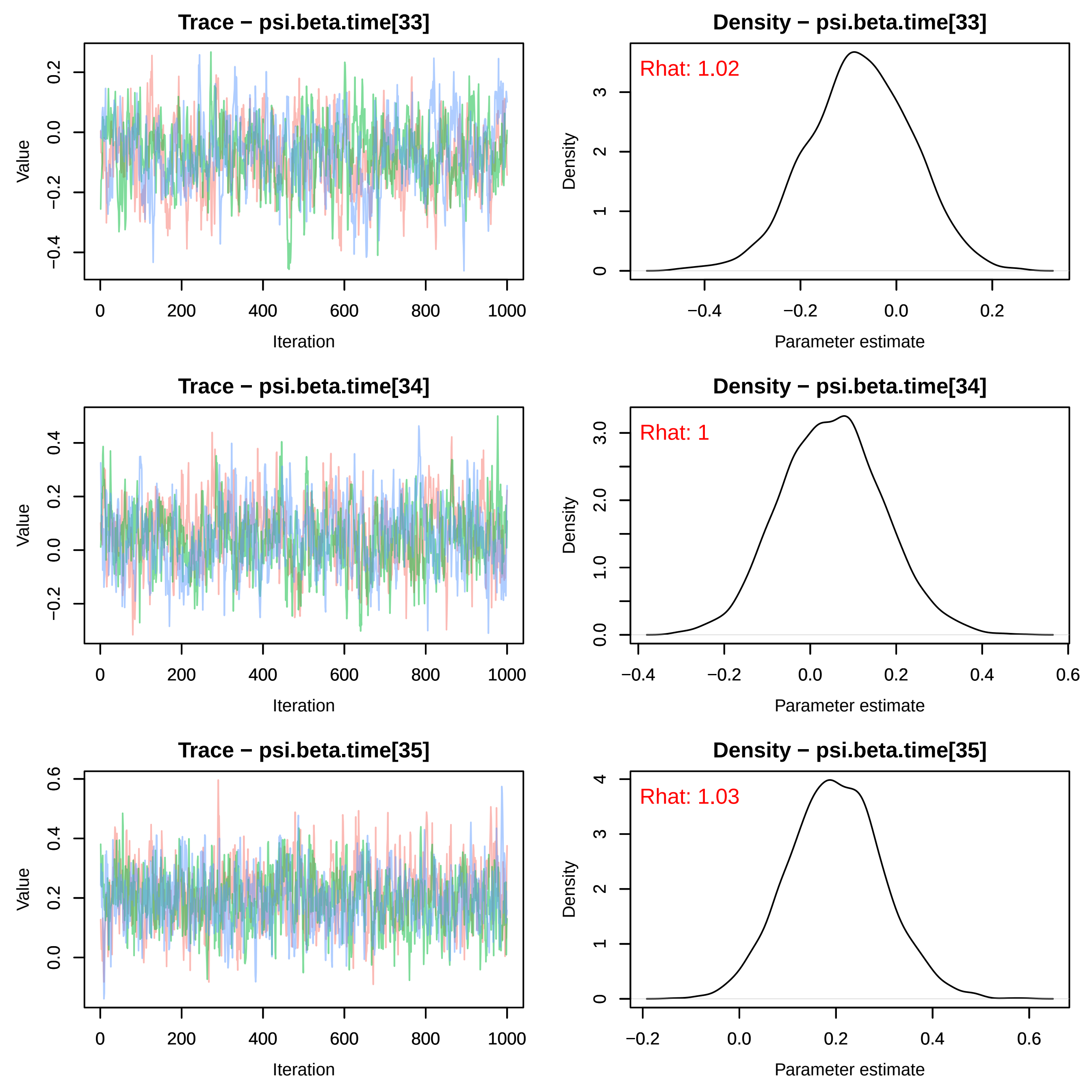

Trace − psi.beta.time[33]
Value
Iteration
Density − psi.beta.time[33]
Rhat: 1.02
Density
Parameter estimate
Trace − psi.beta.time[34]
Value
Iteration
Density − psi.beta.time[34]
Rhat: 1
Density
Parameter estimate
Trace − psi.beta.time[35]
Value
Iteration
Density − psi.beta.time[35]
Rhat: 1.03
Density
Parameter estimate

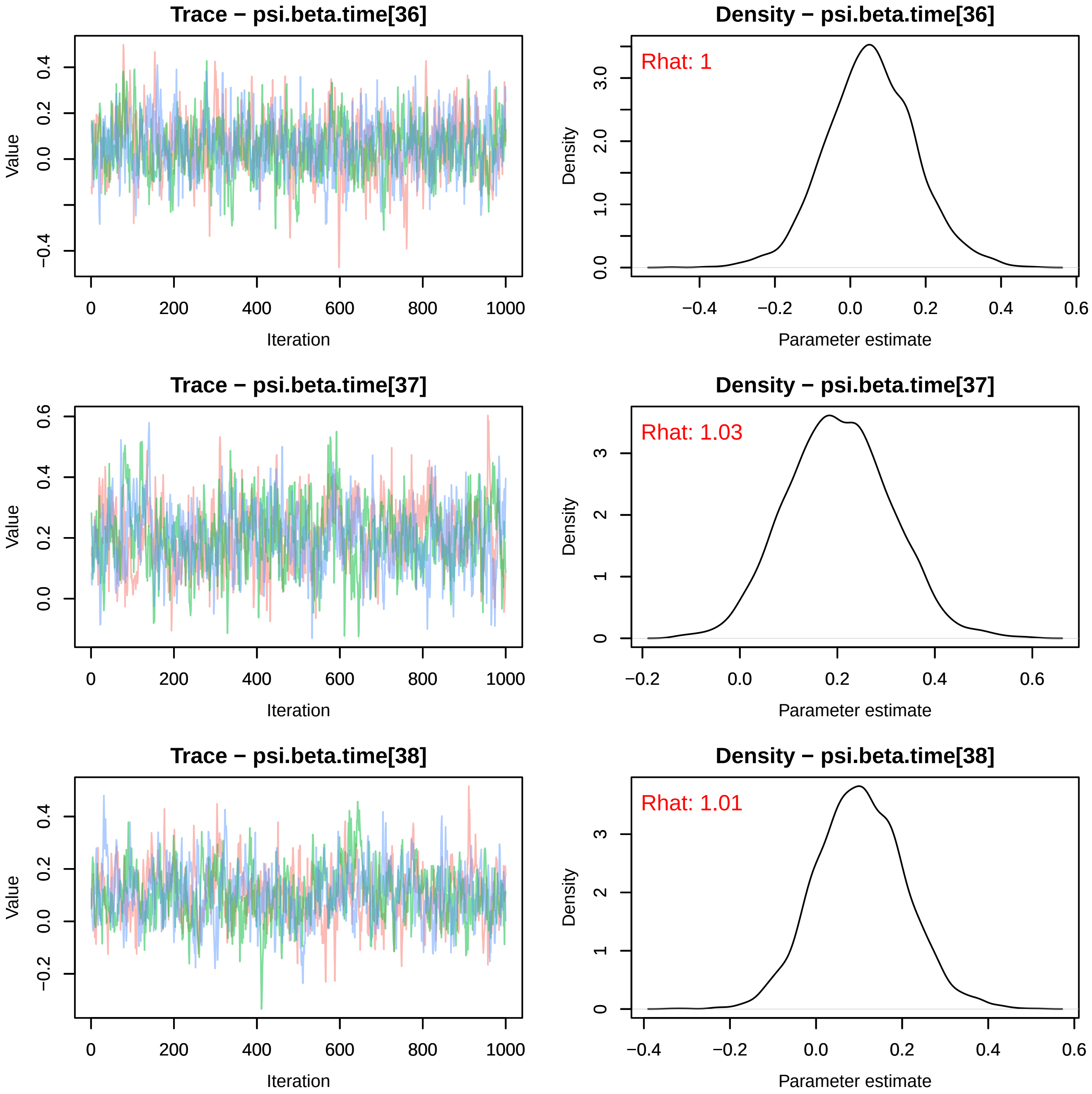
Trace − psi.beta.time[36]
Value
Iteration
Density − psi.beta.time[36]
Rhat: 1
Density
Parameter estimate
Trace − psi.beta.time[37]
Value
Iteration
Density − psi.beta.time[37]
Rhat: 1.03
Density
Parameter estimate
Trace − psi.beta.time[38]
Value
Iteration
Density − psi.beta.time[38]
Rhat: 1.01
Density
Parameter estimate

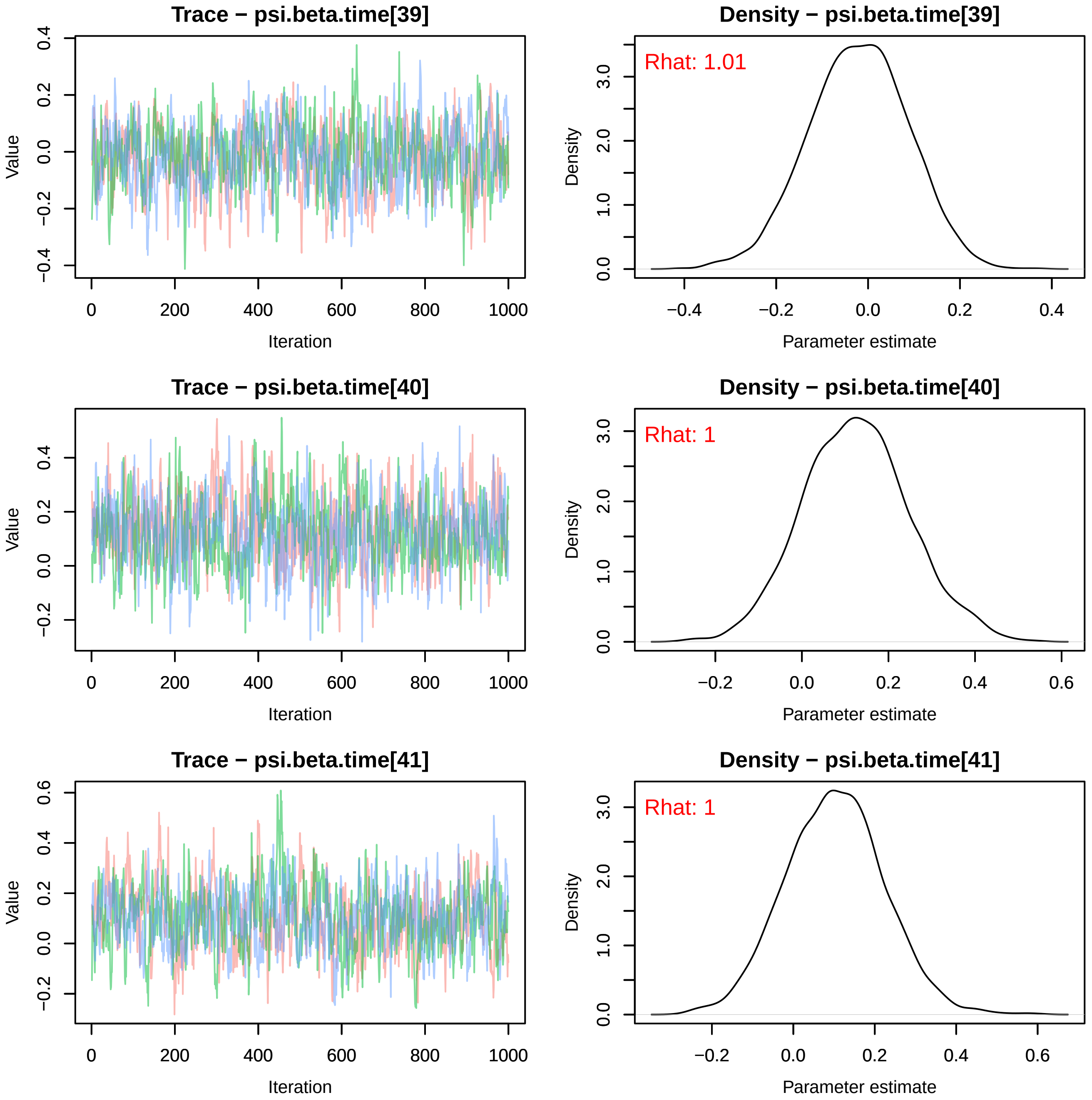
Trace − psi.beta.time[39]
Value
Iteration
Density − psi.beta.time[39]
Rhat: 1.01
Density
Parameter estimate
Trace − psi.beta.time[40]
Value
Iteration
Density − psi.beta.time[40]
Rhat: 1
Density
Parameter estimate
Trace − psi.beta.time[41]
Value
Iteration
Density − psi.beta.time[41]
Rhat: 1
Density
Parameter estimate

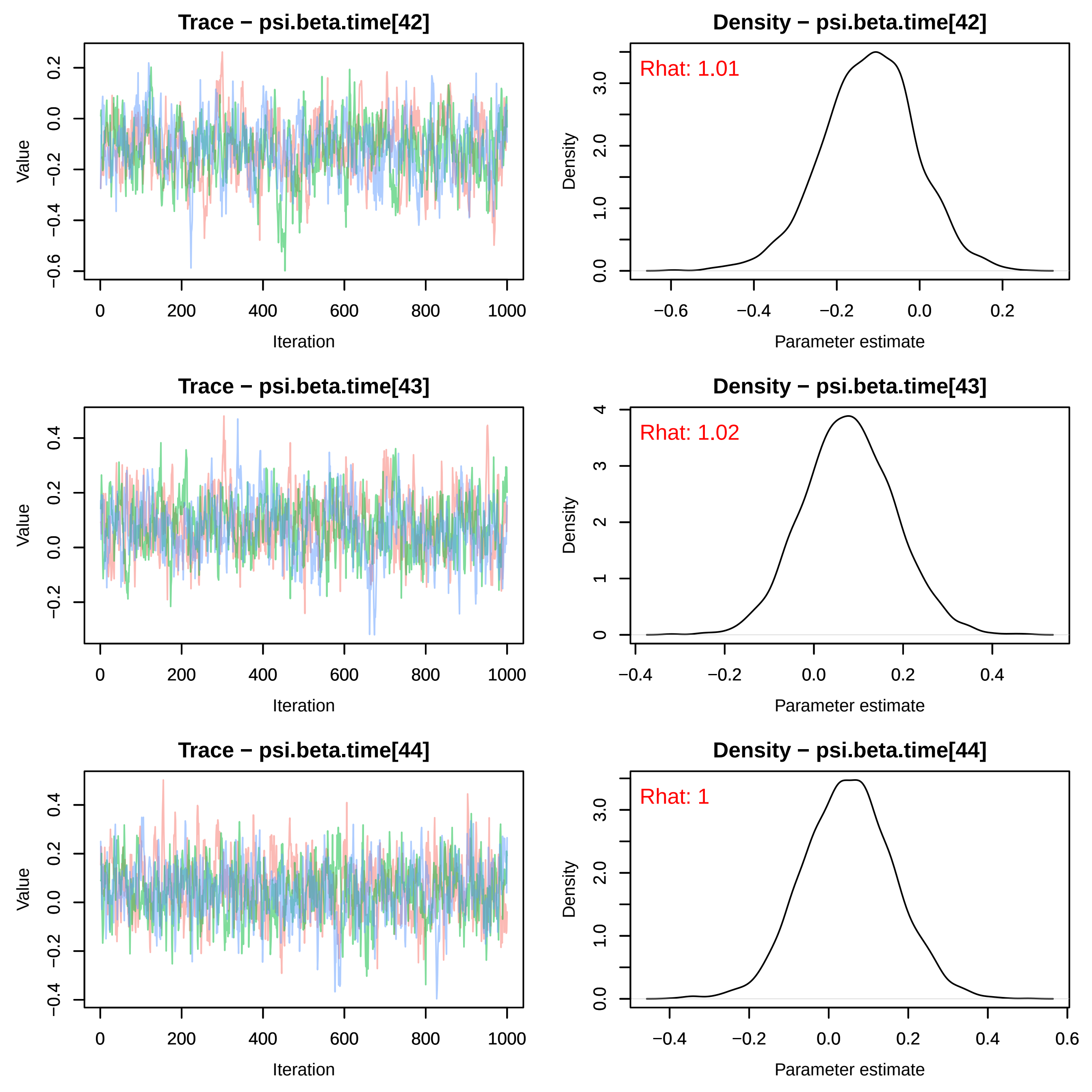
Trace − psi.beta.time[42]
Value
Iteration
Density − psi.beta.time[42]
Rhat: 1.01
Density
Parameter estimate
Trace − psi.beta.time[43]
Value
Iteration
Density − psi.beta.time[43]
Rhat: 1.02
Density
Parameter estimate
Trace − psi.beta.time[44]
Value
Iteration
Density − psi.beta.time[44]
Rhat: 1
Density
Parameter estimate

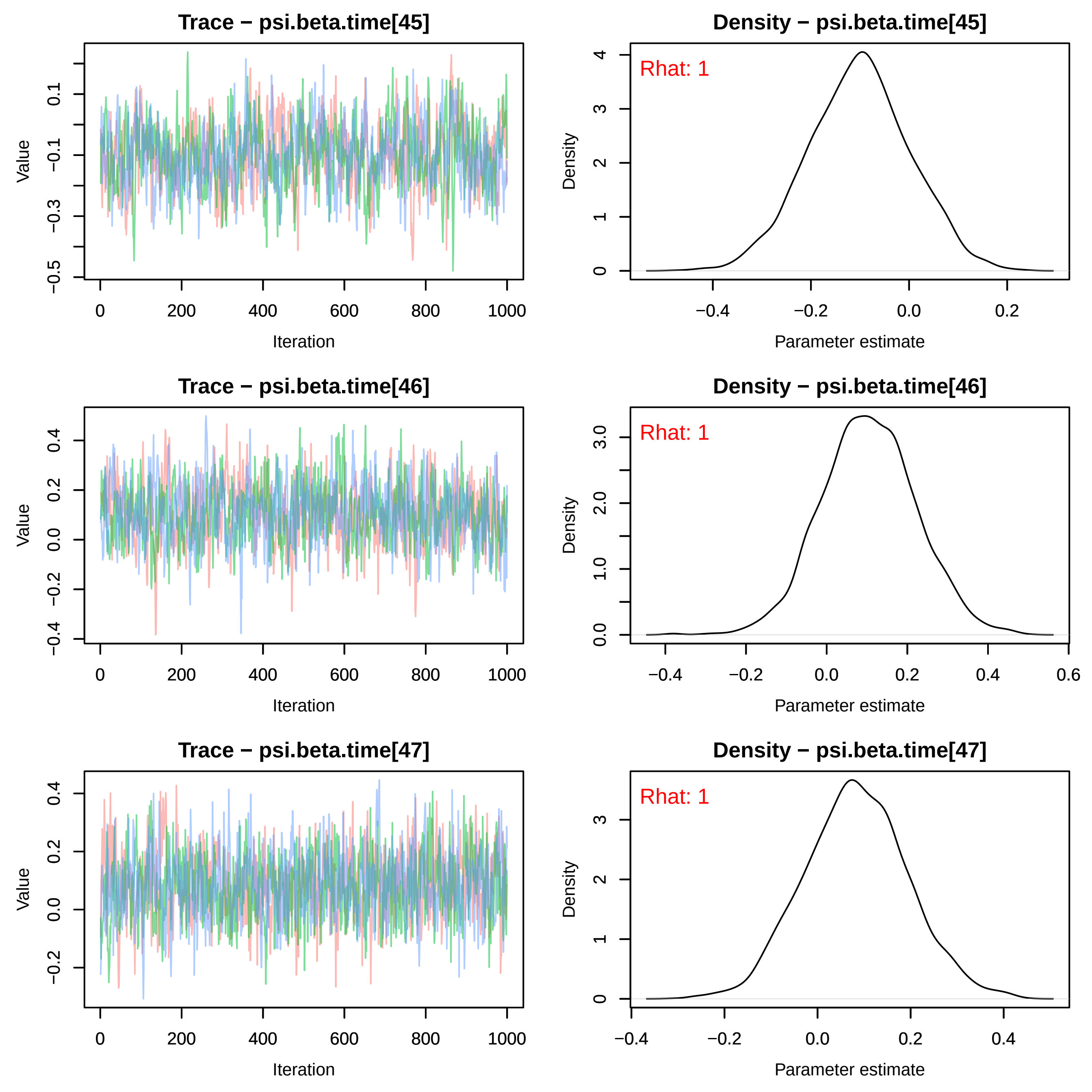

Trace − psi.beta.time[45]
Value
Iteration
Density − psi.beta.time[45]
Rhat: 1
Density
Parameter estimate
Trace − psi.beta.time[46]
Value
Iteration
Density − psi.beta.time[46]
Rhat: 1
Density
Parameter estimate
Trace − psi.beta.time[47]
Value
Iteration
Density − psi.beta.time[47]
Rhat: 1
Density
Parameter estimate

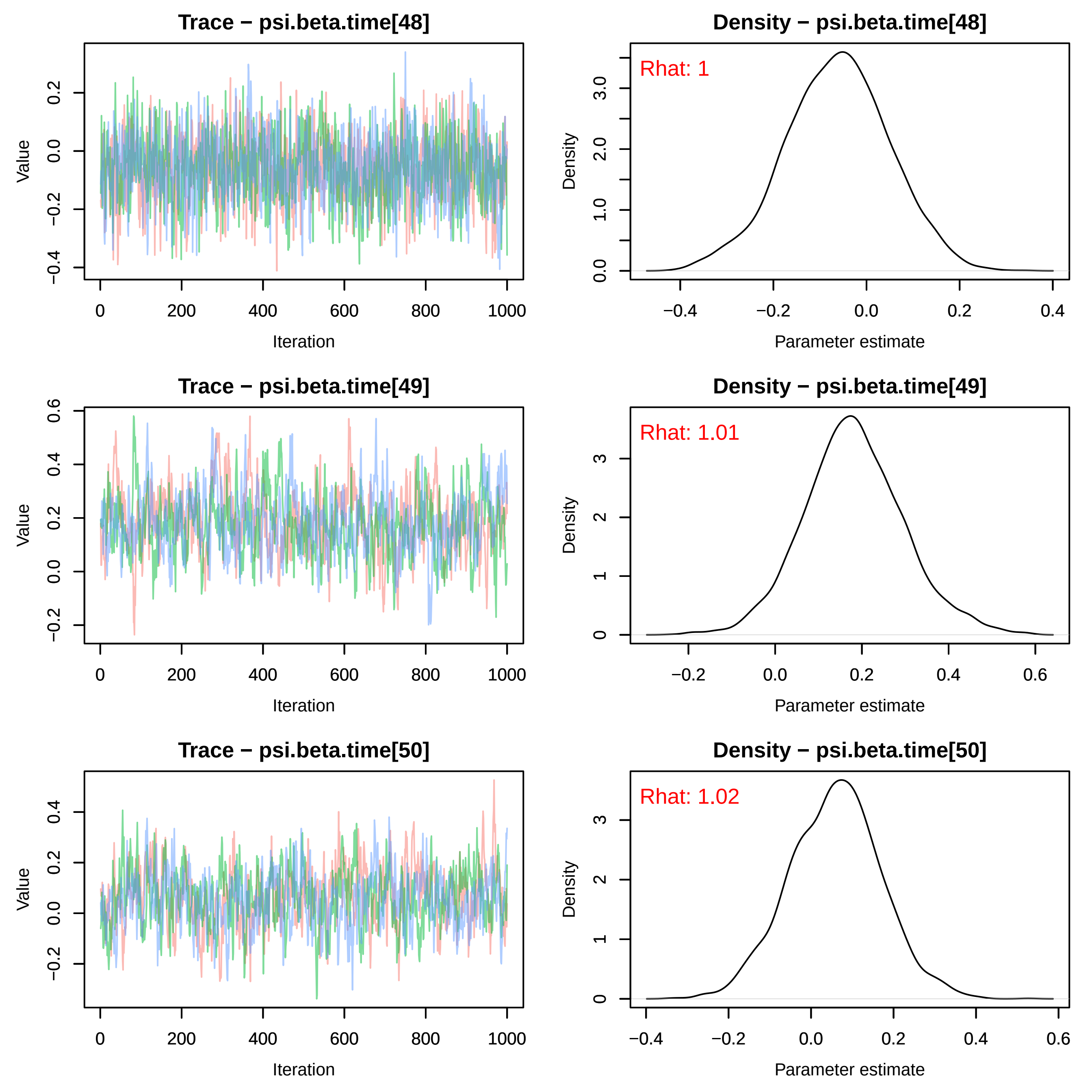
Trace − psi.beta.time[48]
Density − psi.beta.time[48]
Rhat: 1
Trace − psi.beta.time[49]
Density − psi.beta.time[49]
Rhat: 1.01
Trace − psi.beta.time[50]
Density − psi.beta.time[50]
Rhat: 1.02
Value
Iteration
Density
Parameter estimate

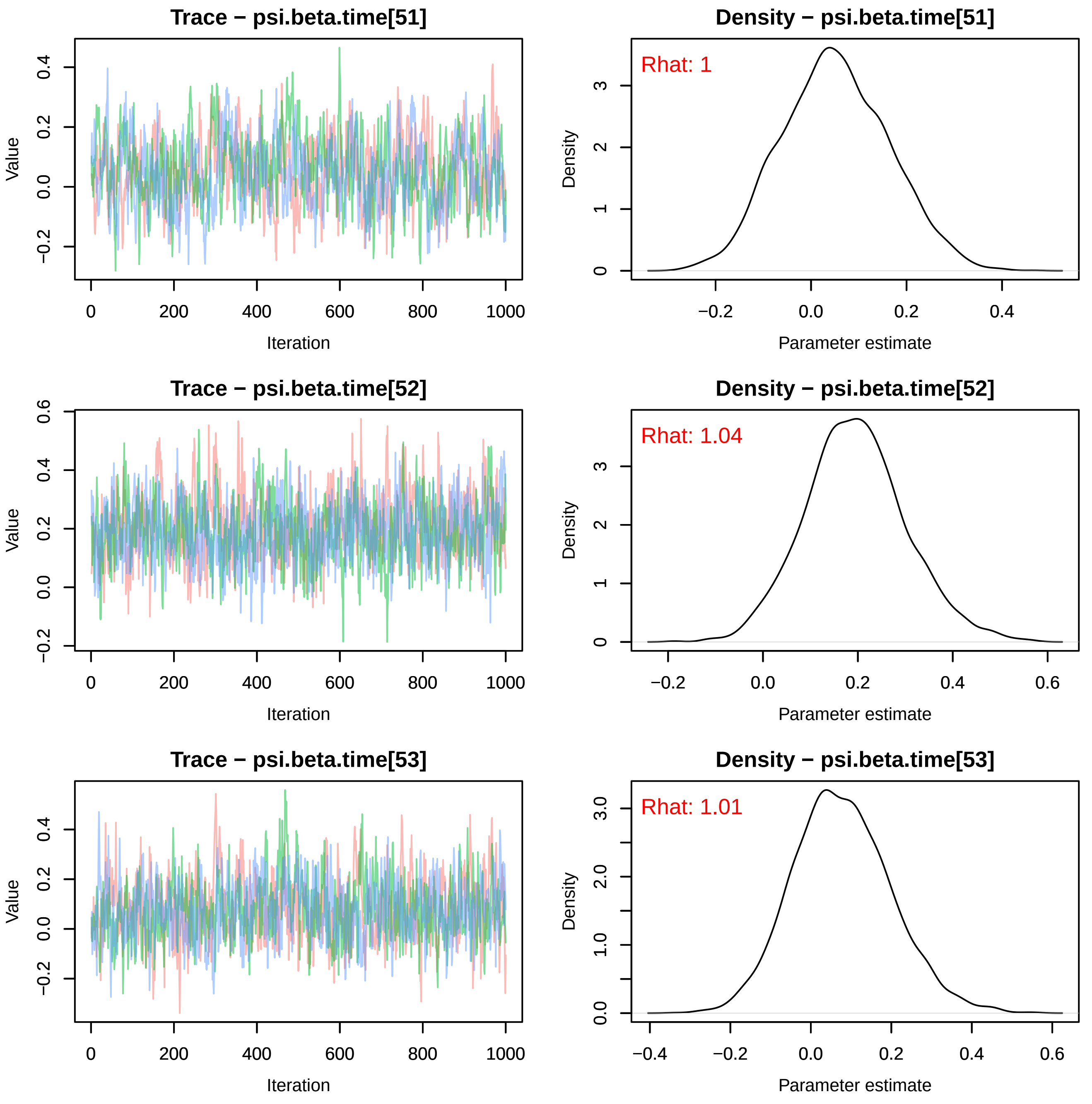
Trace − psi.beta.time[51]
Value
Iteration
Density − psi.beta.time[51]
Rhat: 1
Density
Parameter estimate
Trace − psi.beta.time[52]
Value
Iteration
Density − psi.beta.time[52]
Rhat: 1.04
Density
Parameter estimate
Trace − psi.beta.time[53]
Value
Iteration
Density − psi.beta.time[53]
Rhat: 1.01
Density
Parameter estimate

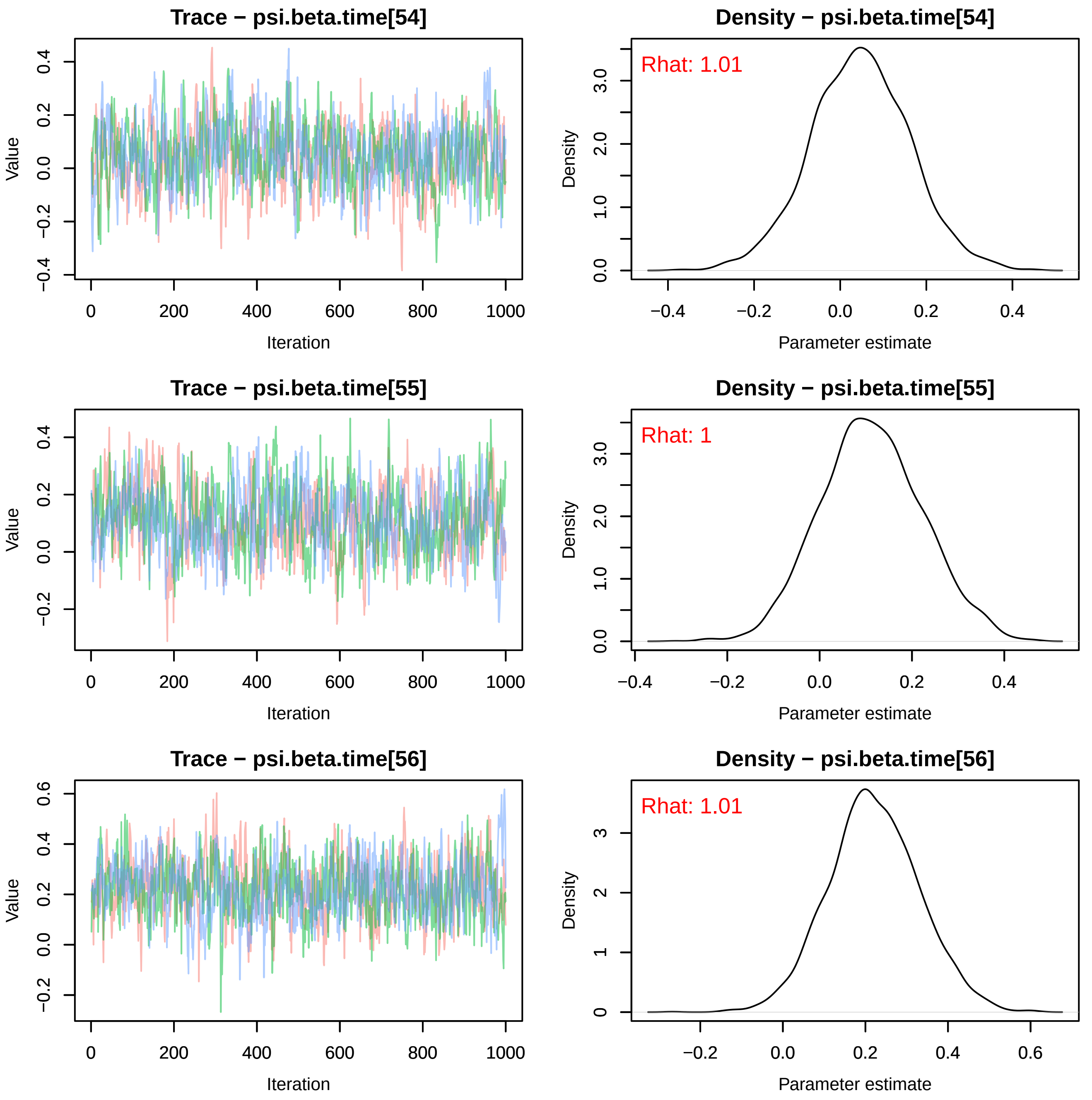
Trace − psi.beta.time[54]
Density − psi.beta.time[54]
Rhat: 1.01
Trace − psi.beta.time[55]
Density − psi.beta.time[55]
Rhat: 1
Trace − psi.beta.time[56]
Density − psi.beta.time[56]
Rhat: 1.01
Value
Iteration
Density
Parameter estimate

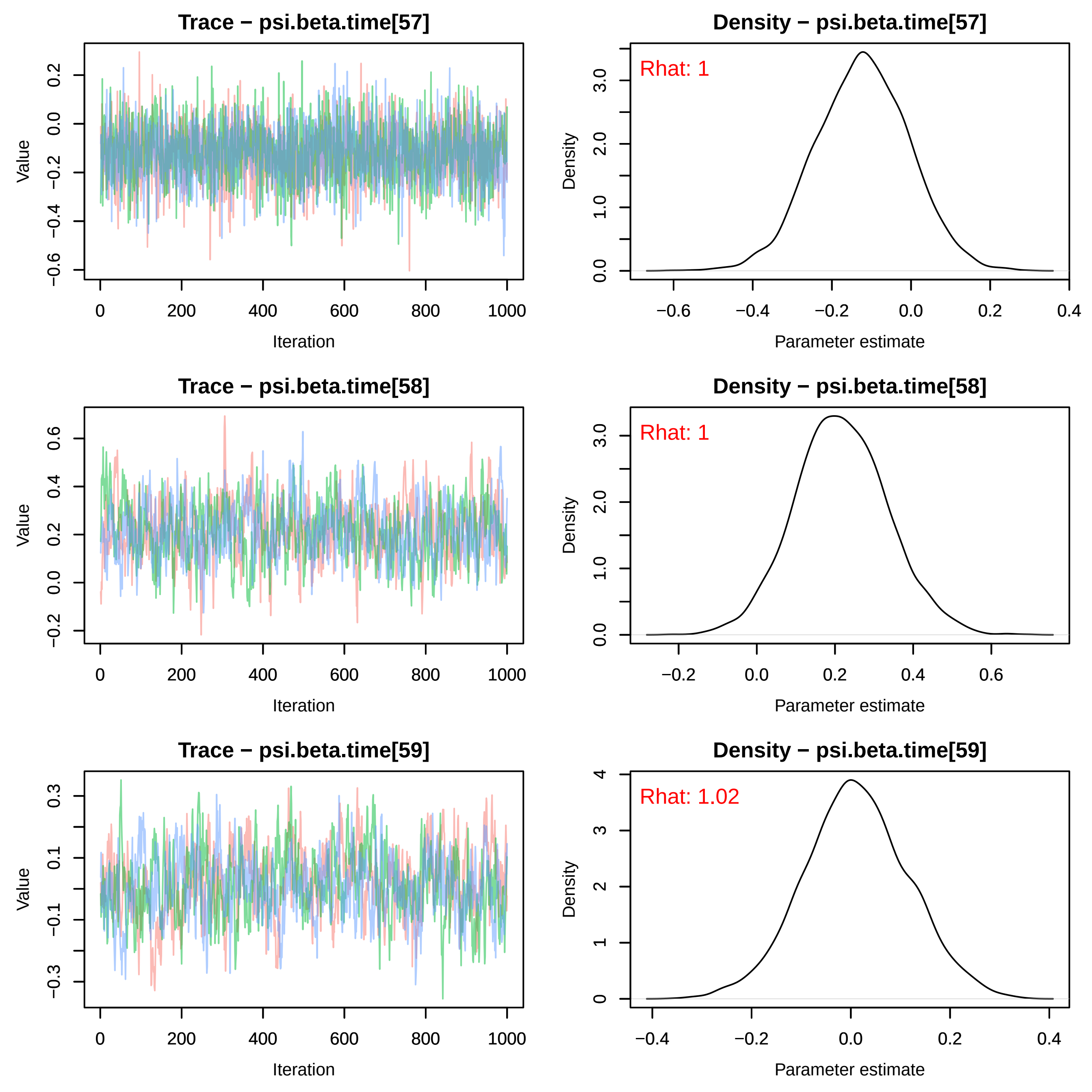
Trace − psi.beta.time[57]
Value
Iteration
Density − psi.beta.time[57]
Rhat: 1
Density
Parameter estimate
Trace − psi.beta.time[58]
Value
Iteration
Density − psi.beta.time[58]
Rhat: 1
Density
Parameter estimate
Trace − psi.beta.time[59]
Value
Iteration
Density − psi.beta.time[59]
Rhat: 1.02
Density
Parameter estimate

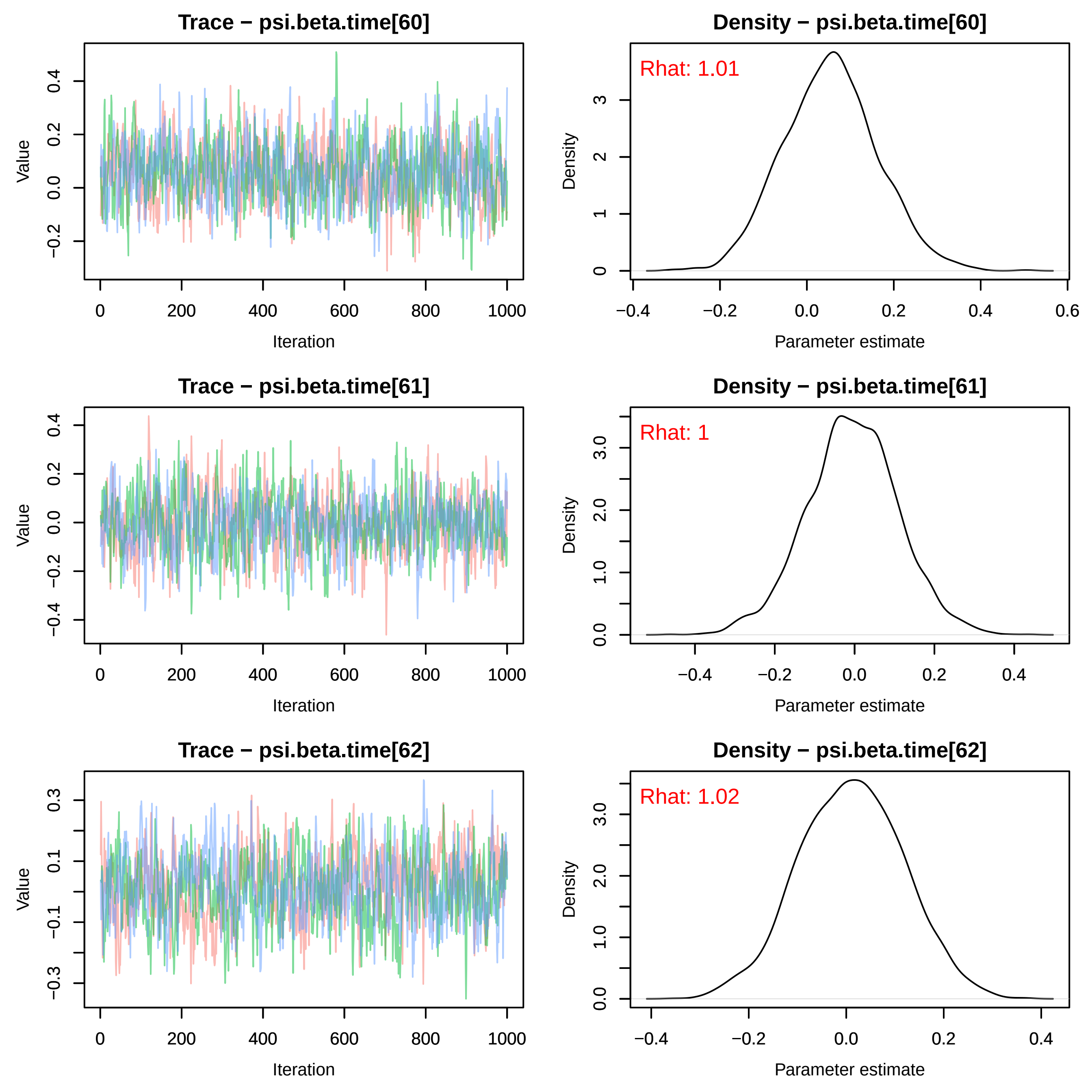

Trace − psi.beta.time[60]
Density − psi.beta.time[60]
Rhat: 1.01
Trace − psi.beta.time[61]
Density − psi.beta.time[61]
Rhat: 1
Trace − psi.beta.time[62]
Density − psi.beta.time[62]
Rhat: 1.02
Value
Iteration
Density
Parameter estimate

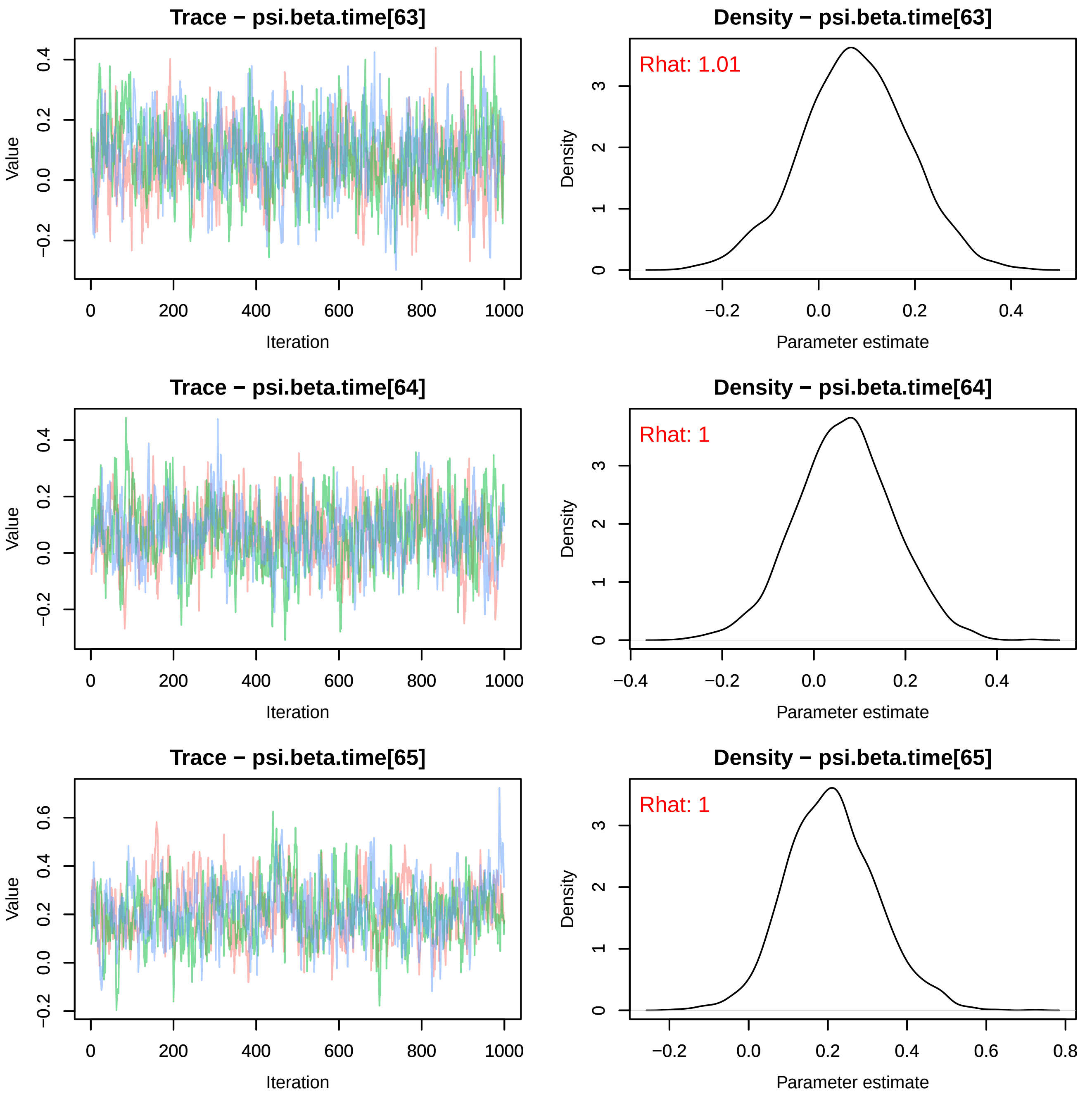
Trace − psi.beta.time[63]
Density − psi.beta.time[63]
Rhat: 1.01
Trace − psi.beta.time[64]
Density − psi.beta.time[64]
Rhat: 1
Trace − psi.beta.time[65]
Density − psi.beta.time[65]
Rhat: 1
Value
Iteration
Density
Parameter estimate
0
200
400
600
800
1000
−0.4
−0.2
0.0
0.2
0.4
0.6
0.8
0
1
2
3

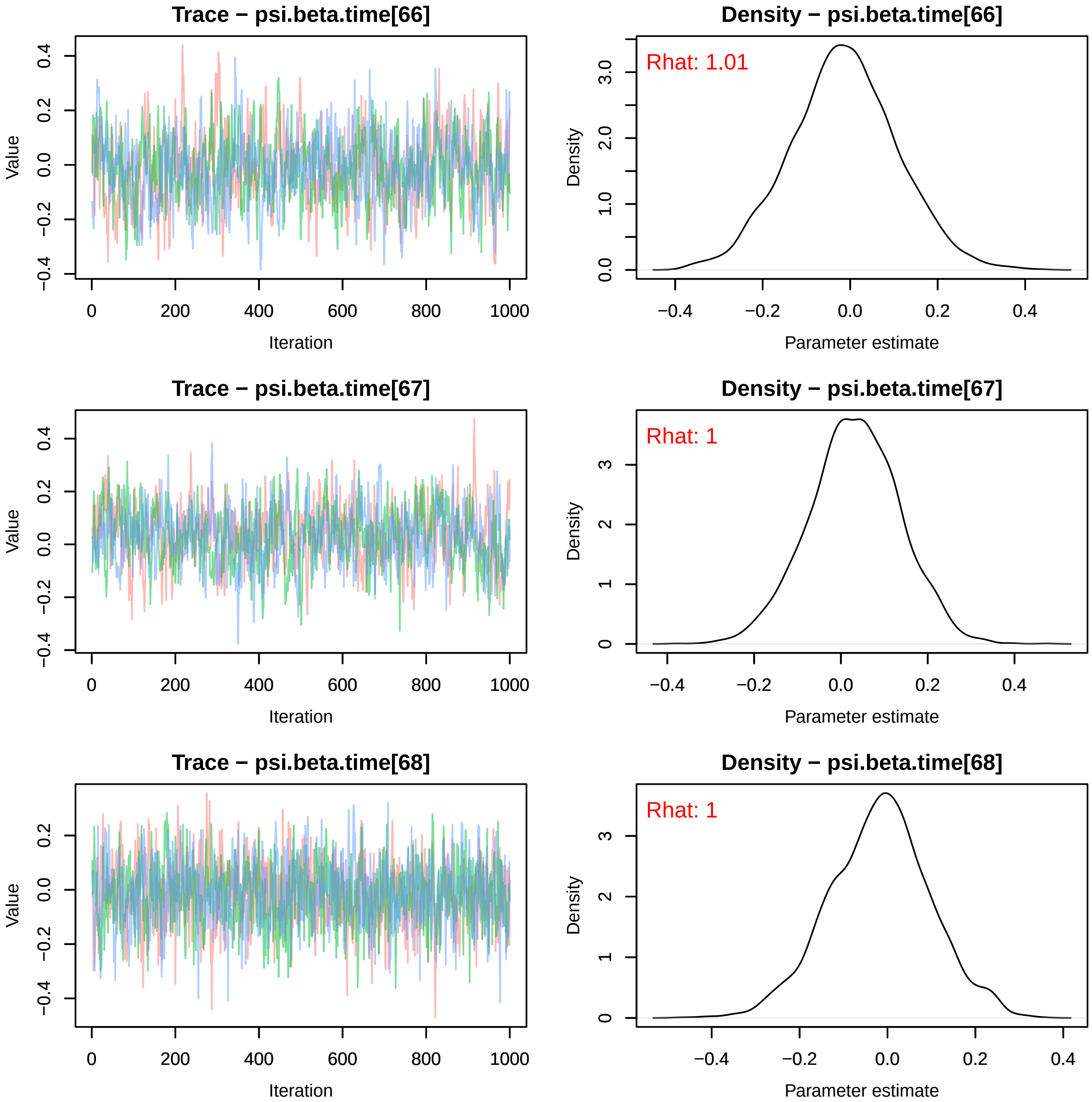

Trace − psi.beta.time[66]
Value
Iteration
0 200 400 600 800 1000
−0.4 −0.2 0.0 0.2 0.4
Density − psi.beta.time[66]
Rhat: 1.01
Density
0.0 1.0 2.0 3.0
−0.4 −0.2 0.0 0.2 0.4
Parameter estimate
Trace − psi.beta.time[67]
Value
Iteration
0 200 400 600 800 1000
−0.4 −0.2 0.0 0.2 0.4
Density − psi.beta.time[67]
Rhat: 1
Density
0 1 2 3
−0.4 −0.2 0.0 0.2 0.4
Parameter estimate
Trace − psi.beta.time[68]
Value
Iteration
0 200 400 600 800 1000
−0.4 −0.2 0.0 0.2
Density − psi.beta.time[68]
Rhat: 1
Density
0 1 2 3
−0.4 −0.2 0.0 0.2 0.4
Parameter estimate

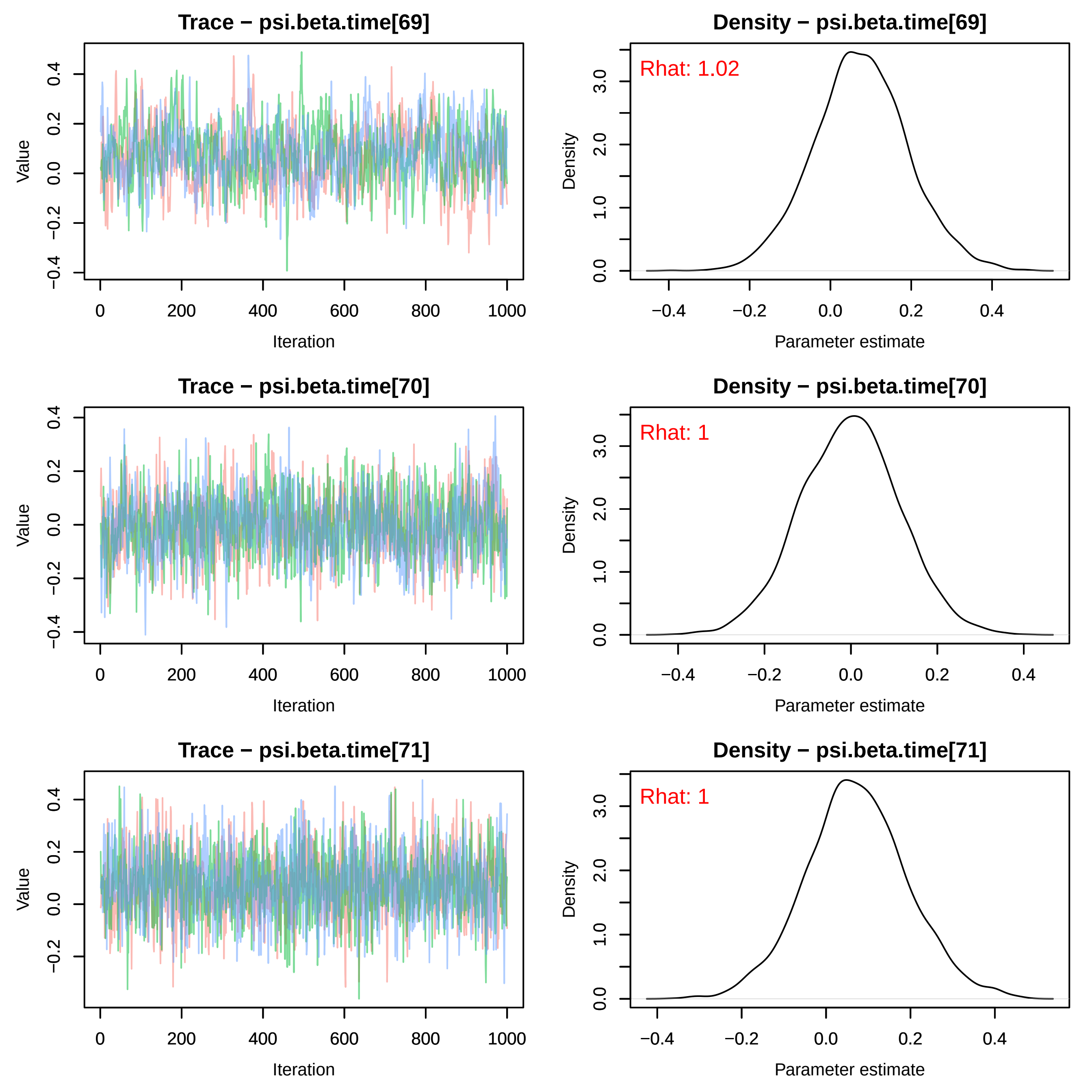
Trace − psi.beta.time[69]
Density − psi.beta.time[69]
Rhat: 1.02
Trace − psi.beta.time[70]
Density − psi.beta.time[70]
Rhat: 1
Trace − psi.beta.time[71]
Density − psi.beta.time[71]
Rhat: 1
Value
Iteration
Density
Parameter estimate

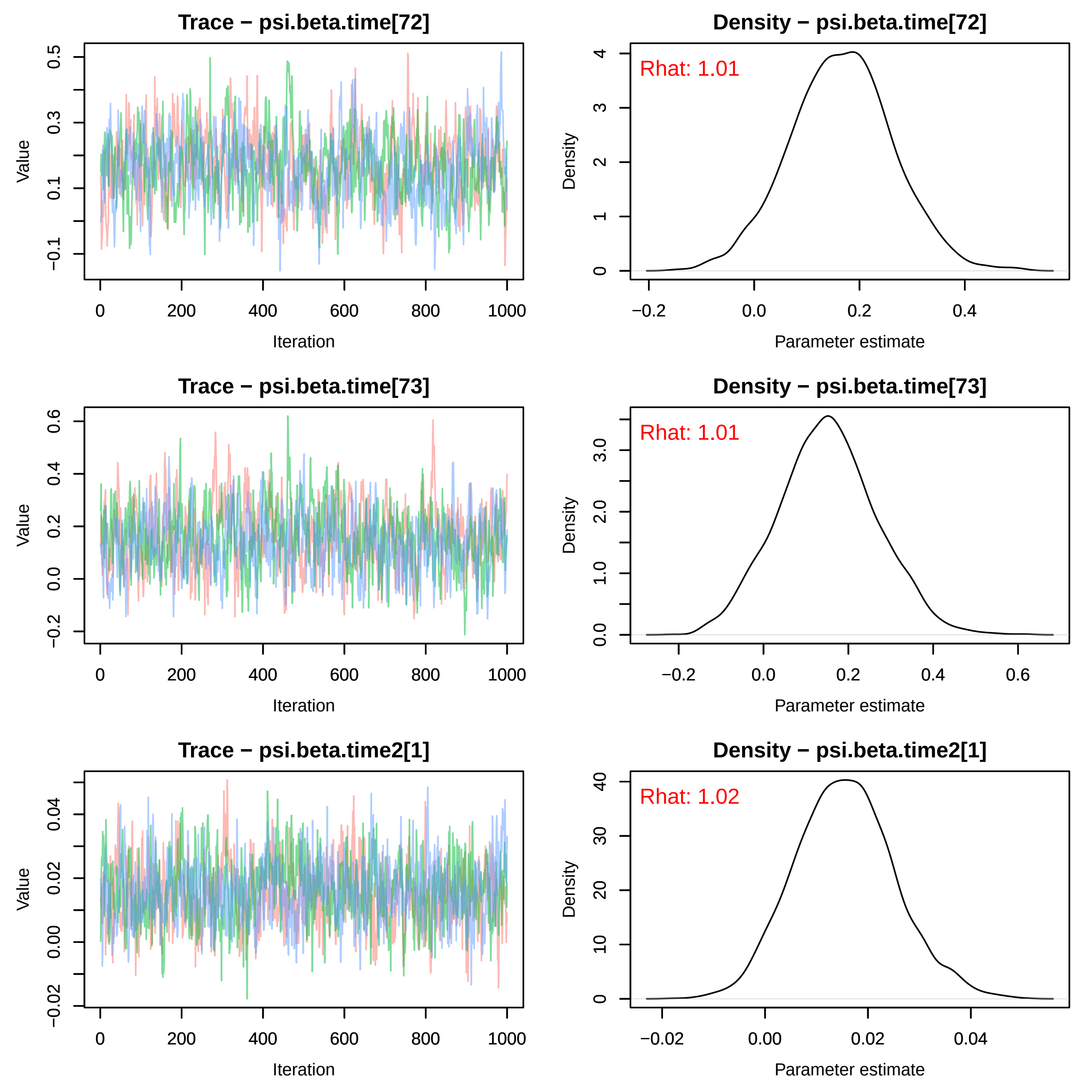
Trace − psi.beta.time[72]
Density − psi.beta.time[72]
Rhat: 1.01
Trace − psi.beta.time[73]
Density − psi.beta.time[73]
Rhat: 1.01
Trace − psi.beta.time2[1]
Density − psi.beta.time2[1]
Rhat: 1.02
Value
Iteration
Density
Parameter estimate

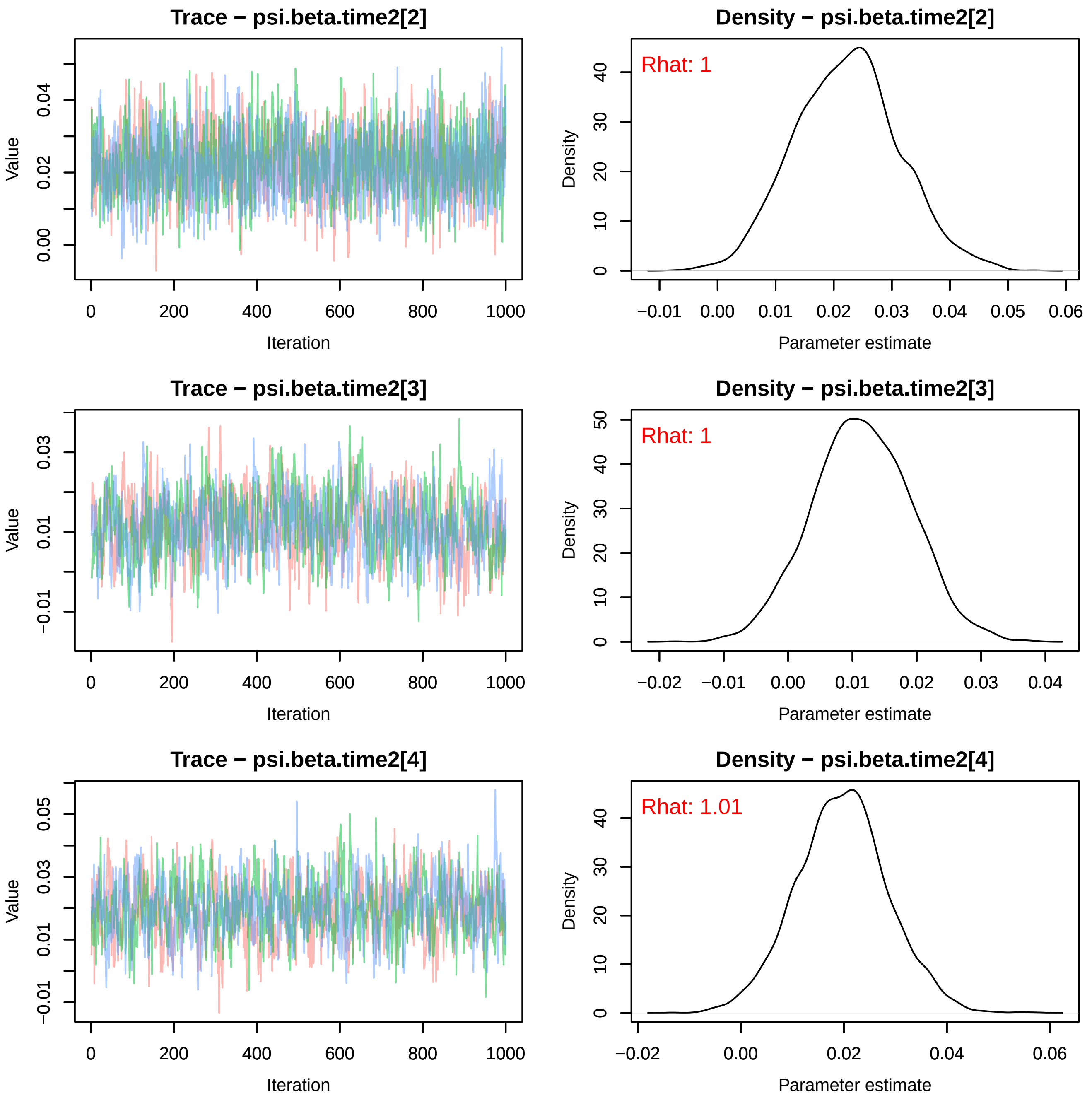
Trace − psi.beta.time2[2]
Value
0.00
0.02
0.04
0
200
400
600
800
1000
Iteration
Density − psi.beta.time2[2]
Rhat: 1
Density
0
10
20
30
40
−0.01
0.00
0.01
0.02
0.03
0.04
0.05
0.06
Parameter estimate
Trace − psi.beta.time2[3]
Value
−0.01
0.01
0.03
0
200
400
600
800
1000
Iteration
Density − psi.beta.time2[3]
Rhat: 1
Density
0
10
20
30
40
50
−0.02
−0.01
0.00
0.01
0.02
0.03
0.04
Parameter estimate
Trace − psi.beta.time2[4]
Value
−0.01
0.01
0.03
0.05
0
200
400
600
800
1000
Iteration
Density − psi.beta.time2[4]
Rhat: 1.01
Density
0
10
20
30
40
−0.02
0.00
0.02
0.04
0.06
Parameter estimate

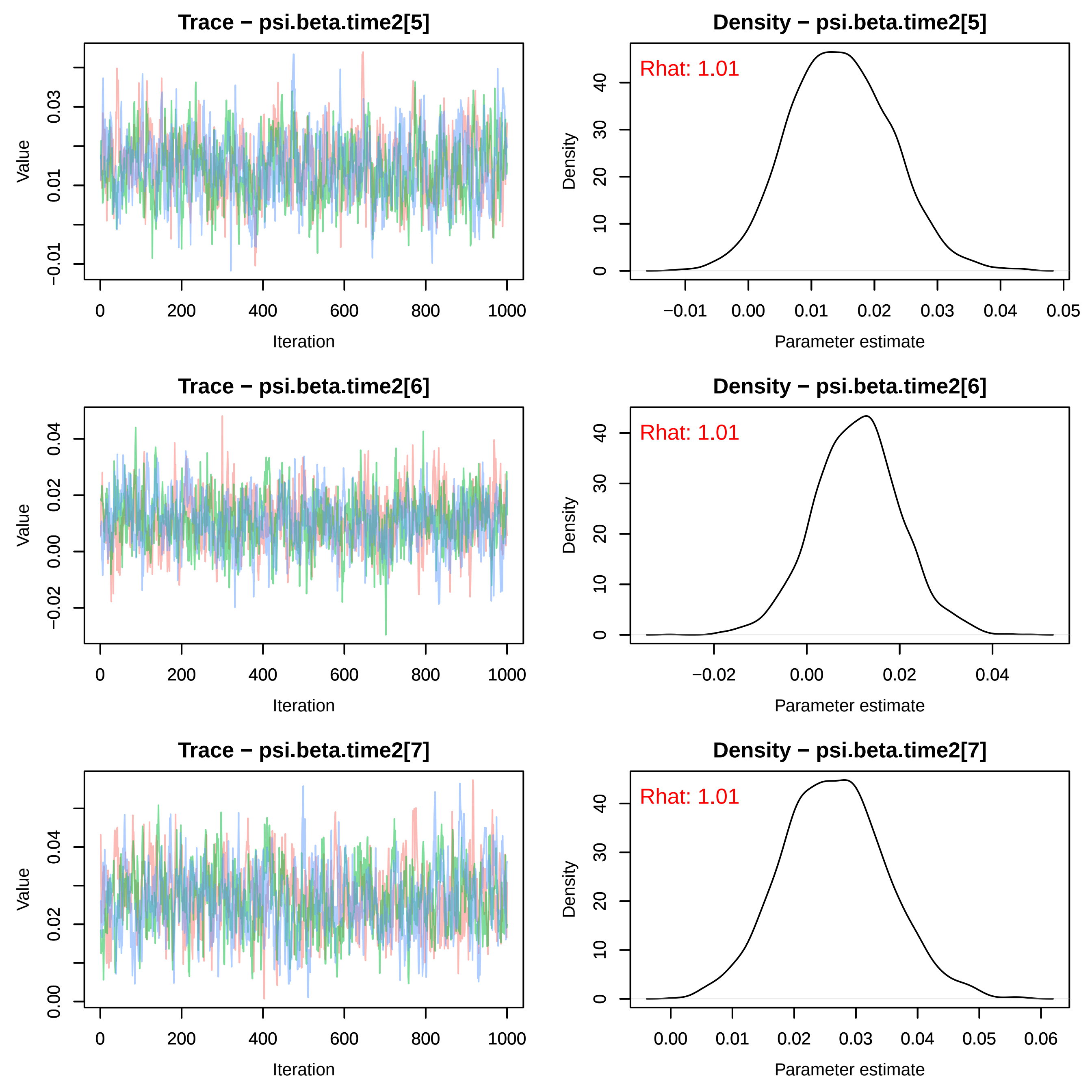
Trace − psi.beta.time2[5]
Value
−0.01
0.01
0.03
0
200
400
600
800
1000
Iteration
Density − psi.beta.time2[5]
Rhat: 1.01
Density
0
10
20
30
40
−0.01
0.00
0.01
0.02
0.03
0.04
0.05
Parameter estimate
Trace − psi.beta.time2[6]
Value
−0.02
0.00
0.02
0.04
0
200
400
600
800
1000
Iteration
Density − psi.beta.time2[6]
Rhat: 1.01
Density
0
10
20
30
40
−0.02
0.00
0.02
0.04
Parameter estimate
Trace − psi.beta.time2[7]
Value
0.00
0.02
0.04
0
200
400
600
800
1000
Iteration
Density − psi.beta.time2[7]
Rhat: 1.01
Density
0
10
20
30
40
0.00
0.01
0.02
0.03
0.04
0.05
0.06
Parameter estimate

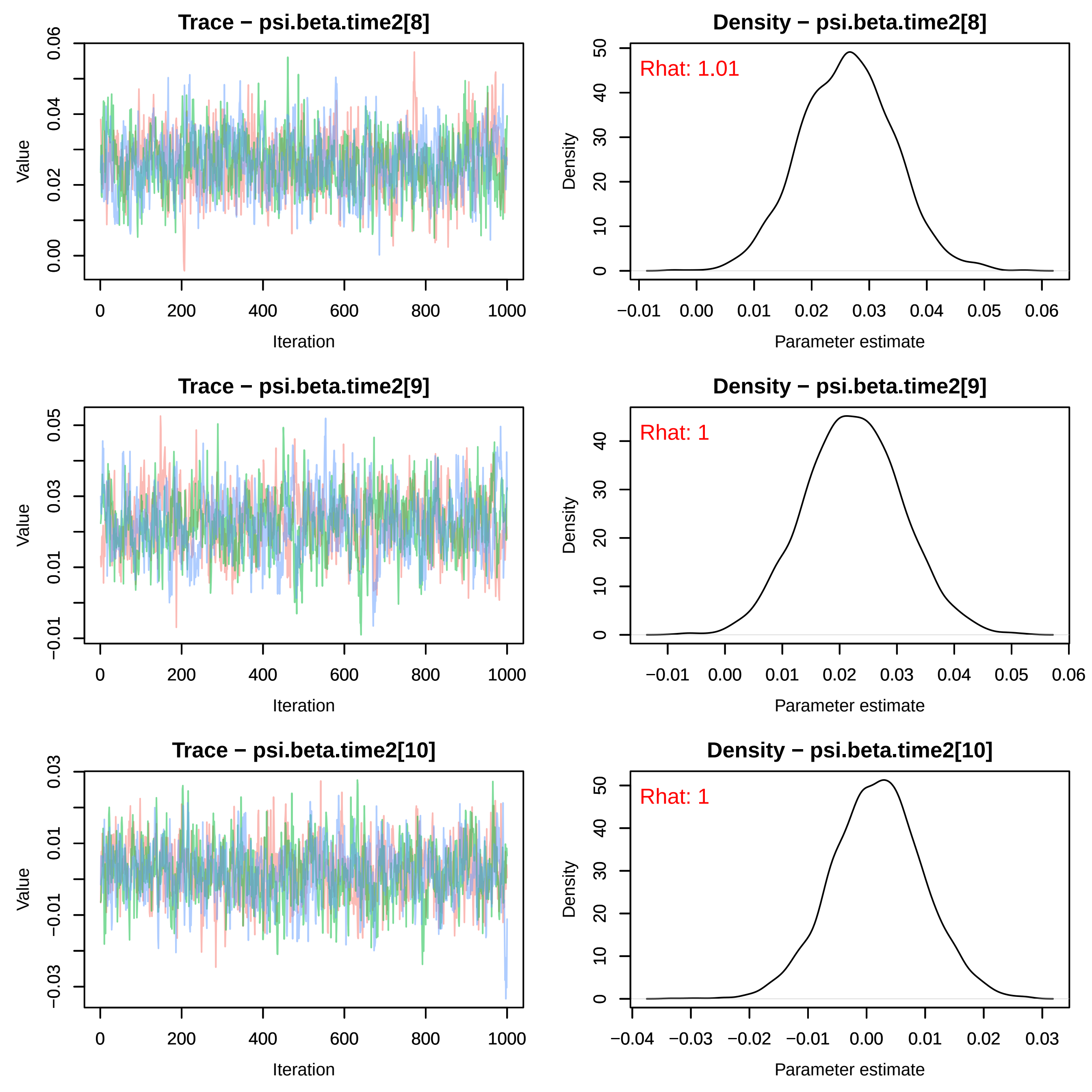
Trace − psi.beta.time2[8]
Density − psi.beta.time2[8]
Rhat: 1.01
Trace − psi.beta.time2[9]
Density − psi.beta.time2[9]
Rhat: 1
Trace − psi.beta.time2[10]
Density − psi.beta.time2[10]
Rhat: 1
Value
Iteration
Density
Parameter estimate

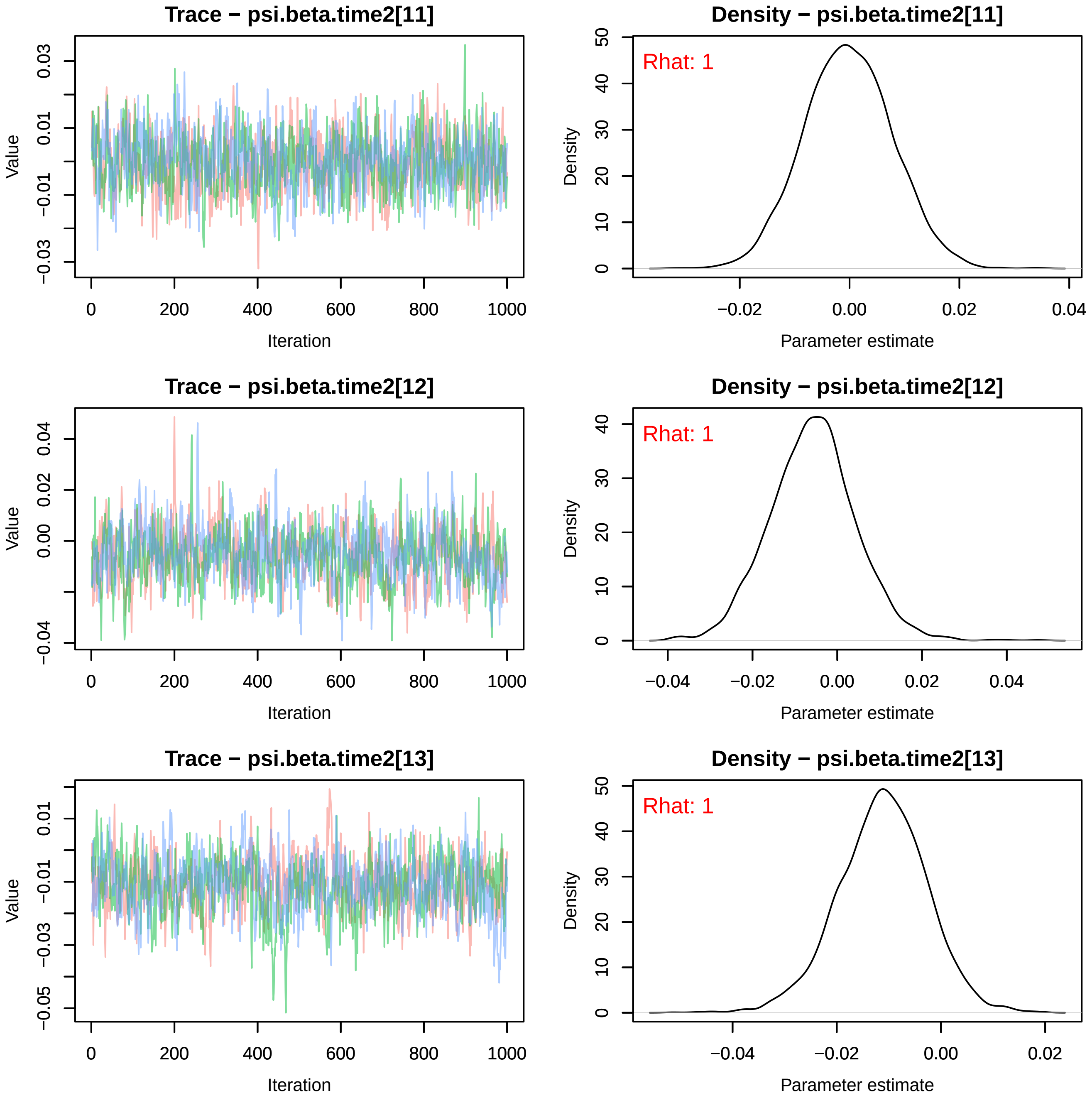

Trace − psi.beta.time2[11]
Value
Iteration
0
200
400
600
800
1000
0.03
0.01
−0.01
−0.03
Density − psi.beta.time2[11]
Rhat: 1
Density
Parameter estimate
50
40
30
20
10
0
−0.02
0.00
0.02
0.04
Trace − psi.beta.time2[12]
Value
Iteration
0.04
0.02
0.00
−0.04
Density − psi.beta.time2[12]
Rhat: 1
Density
Parameter estimate
40
30
20
10
0
−0.04
−0.02
0.00
0.02
0.04
Trace − psi.beta.time2[13]
Value
Iteration
0.01
−0.01
−0.03
−0.05
Density − psi.beta.time2[13]
Rhat: 1
Density
Parameter estimate
50
40
30
20
10
0
−0.04
−0.02
0.00
0.02

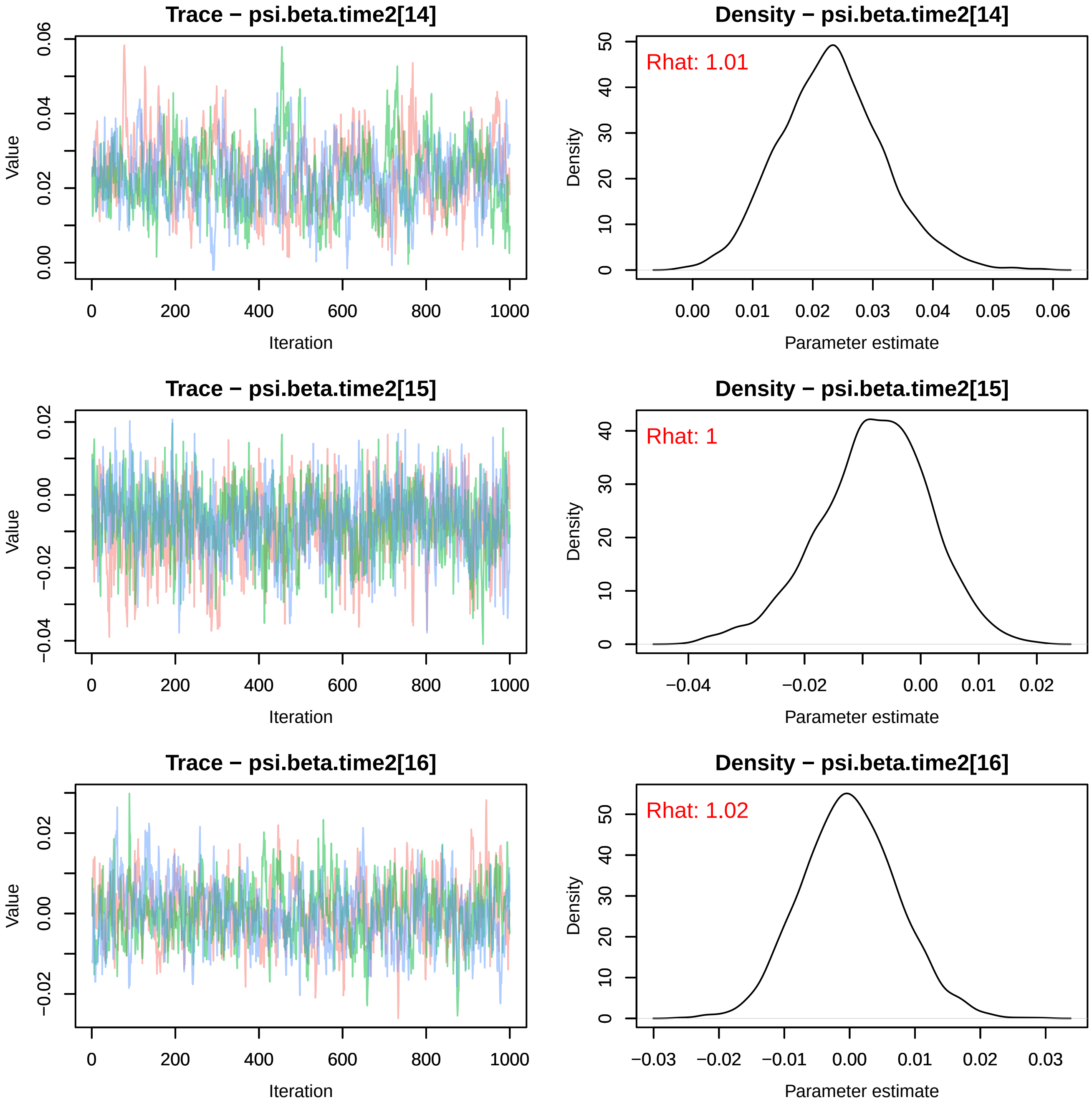

Trace − psi.beta.time2[14]
Value
Iteration
Density − psi.beta.time2[14]
Rhat: 1.01
Density
Parameter estimate
Trace − psi.beta.time2[15]
Value
Iteration
Density − psi.beta.time2[15]
Rhat: 1
Density
Parameter estimate
Trace − psi.beta.time2[16]
Value
Iteration
Density − psi.beta.time2[16]
Rhat: 1.02
Density
Parameter estimate

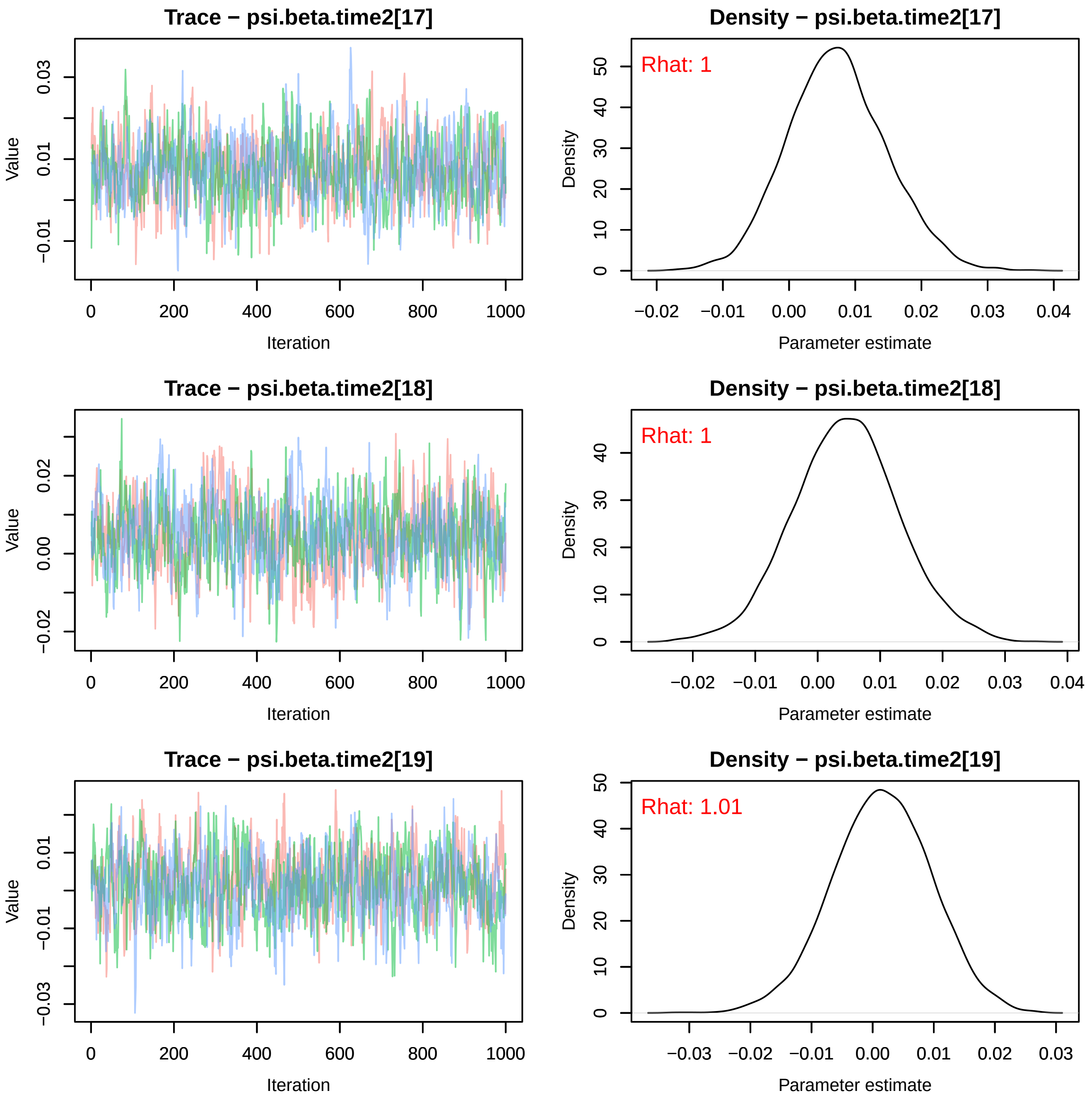
Trace − psi.beta.time2[17]
Value
Iteration
Density − psi.beta.time2[17]
Rhat: 1
Density
Parameter estimate
Trace − psi.beta.time2[18]
Value
Iteration
Density − psi.beta.time2[18]
Rhat: 1
Density
Parameter estimate
Trace − psi.beta.time2[19]
Value
Iteration
Density − psi.beta.time2[19]
Rhat: 1.01
Density
Parameter estimate

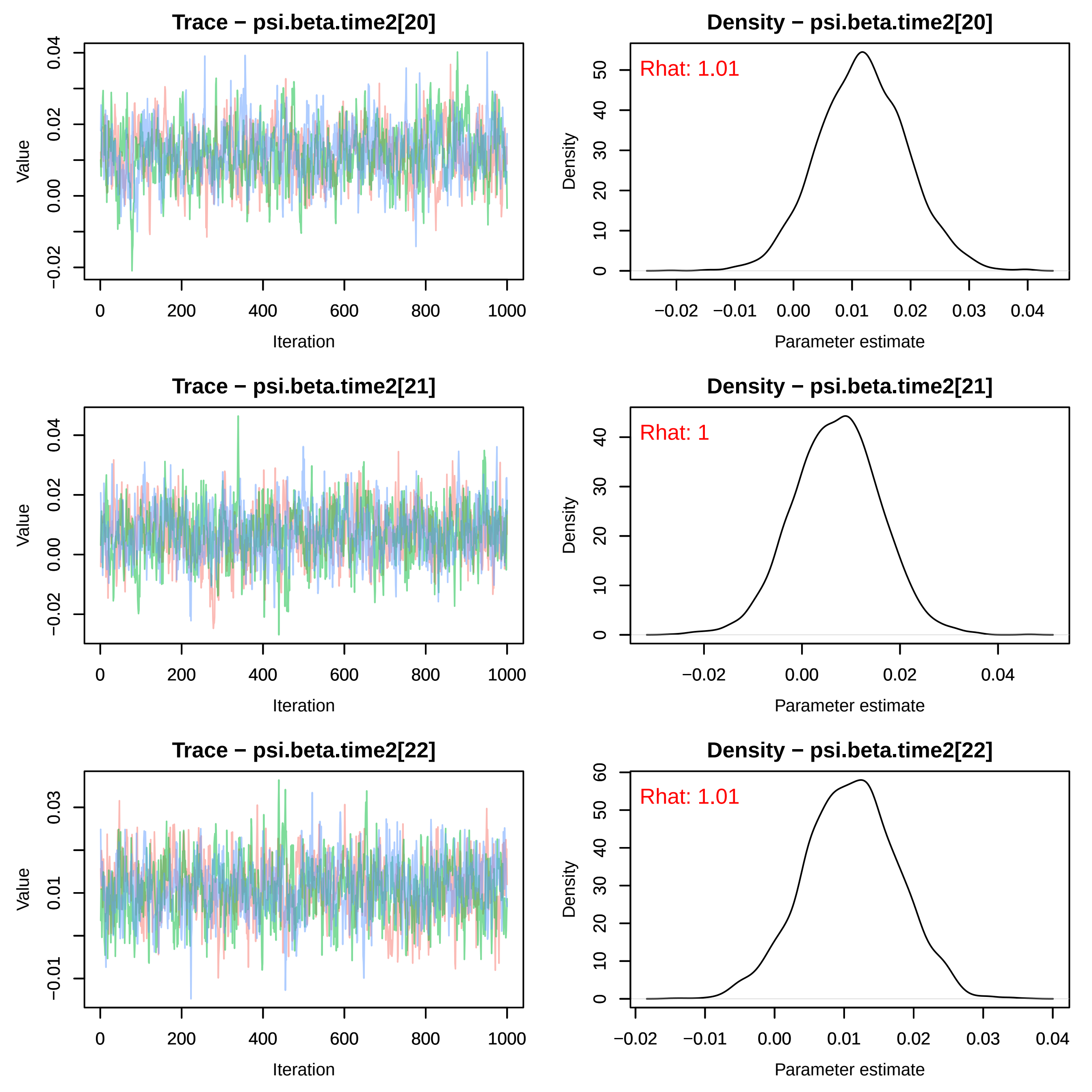
Trace − psi.beta.time2[20]
Value
Iteration
Density − psi.beta.time2[20]
Rhat: 1.01
Density
Parameter estimate
Trace − psi.beta.time2[21]
Value
Iteration
Density − psi.beta.time2[21]
Rhat: 1
Density
Parameter estimate
Trace − psi.beta.time2[22]
Value
Iteration
Density − psi.beta.time2[22]
Rhat: 1.01
Density
Parameter estimate

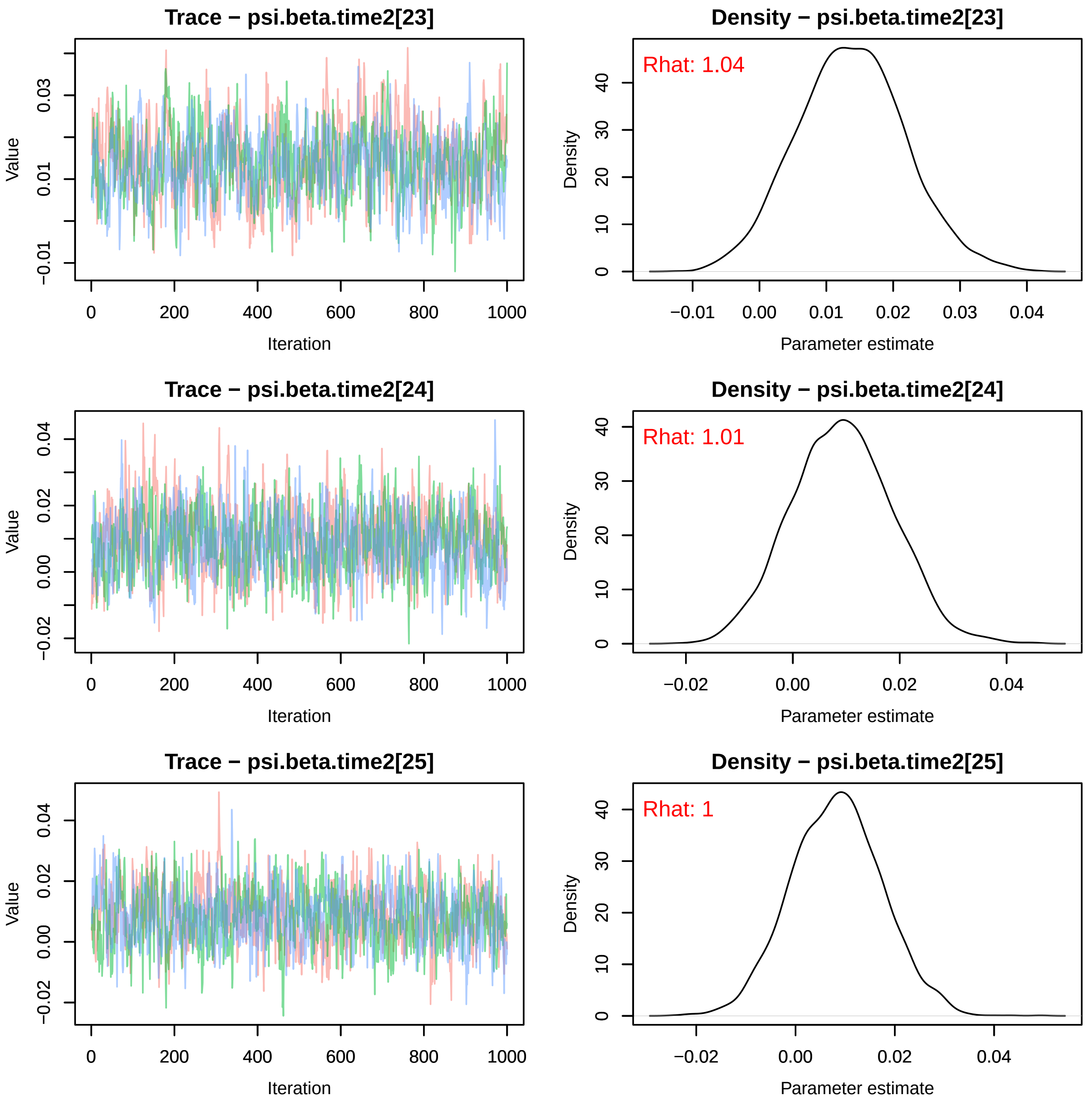
Trace − psi.beta.time2[23]
Value
Iteration
0 200 400 600 800 1000
−0.01 0.01 0.03
Density − psi.beta.time2[23]
Rhat: 1.04
Density
0 10 20 30 40
−0.01 0.00 0.01 0.02 0.03 0.04
Parameter estimate
Trace − psi.beta.time2[24]
Value
Iteration
0 200 400 600 800 1000
−0.02 0.00 0.02 0.04
Density − psi.beta.time2[24]
Rhat: 1.01
Density
0 10 20 30 40
−0.02 0.00 0.02 0.04
Parameter estimate
Trace − psi.beta.time2[25]
Value
Iteration
0 200 400 600 800 1000
−0.02 0.00 0.02 0.04
Density − psi.beta.time2[25]
Rhat: 1
Density
0 10 20 30 40
−0.02 0.00 0.02 0.04
Parameter estimate

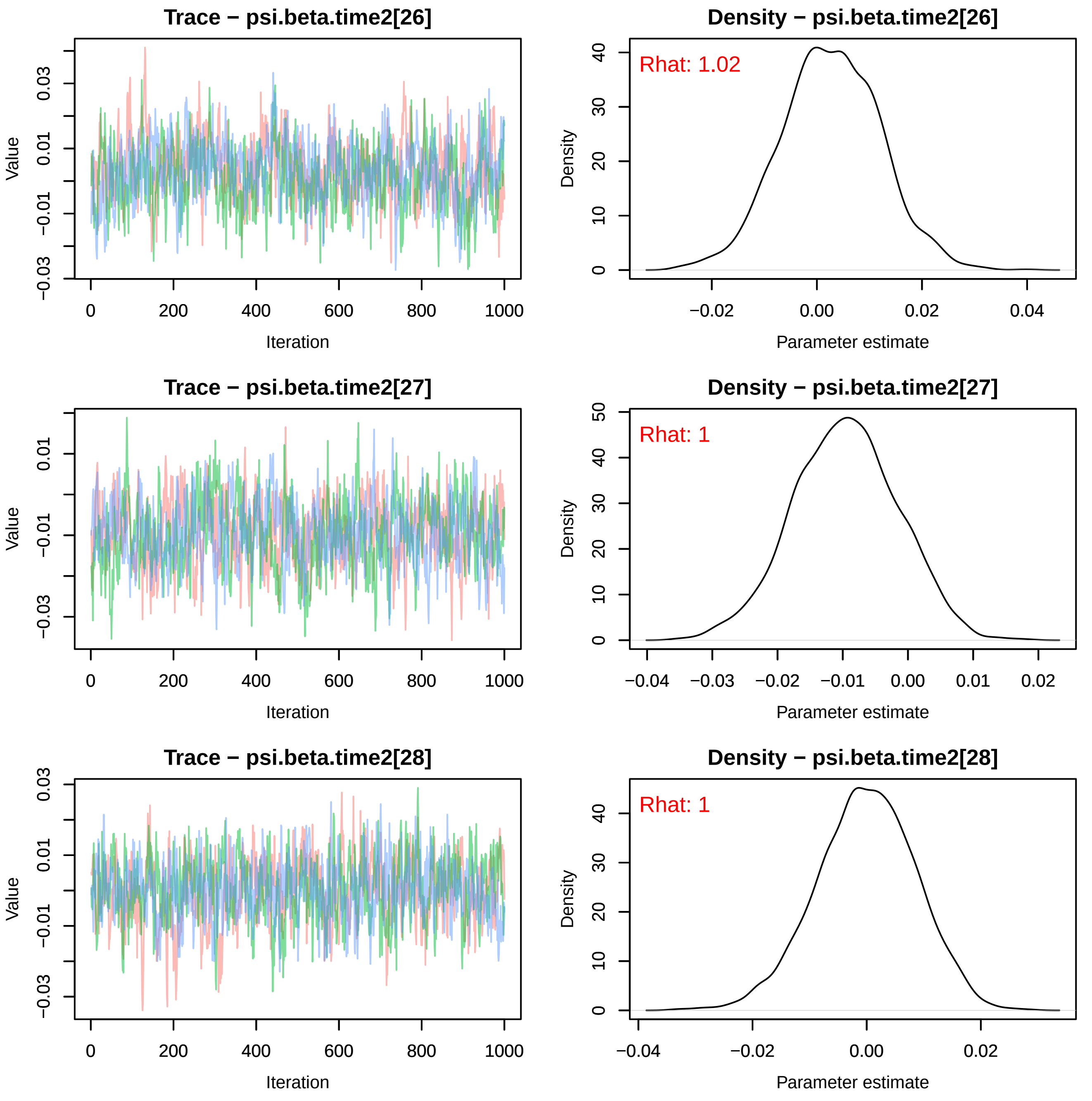
Trace − psi.beta.time2[26]
Value
Iteration
Density − psi.beta.time2[26]
Rhat: 1.02
Density
Parameter estimate
Trace − psi.beta.time2[27]
Value
Iteration
Density − psi.beta.time2[27]
Rhat: 1
Density
Parameter estimate
Trace − psi.beta.time2[28]
Value
Iteration
Density − psi.beta.time2[28]
Rhat: 1
Density
Parameter estimate

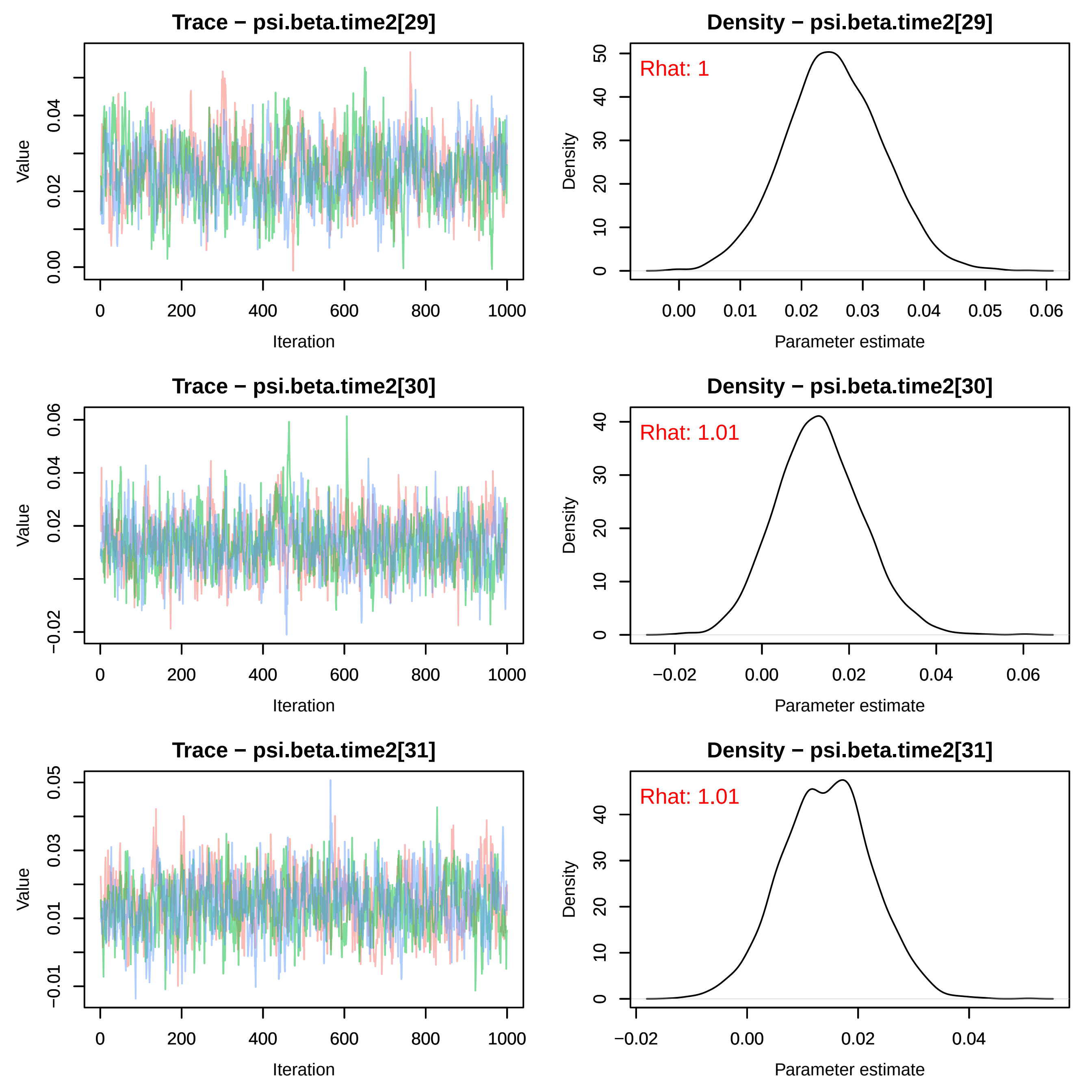
Trace − psi.beta.time2[29]
Value
Iteration
Density − psi.beta.time2[29]
Rhat: 1
Density
Parameter estimate
Trace − psi.beta.time2[30]
Value
Iteration
Density − psi.beta.time2[30]
Rhat: 1.01
Density
Parameter estimate
Trace − psi.beta.time2[31]
Value
Iteration
Density − psi.beta.time2[31]
Rhat: 1.01
Density
Parameter estimate

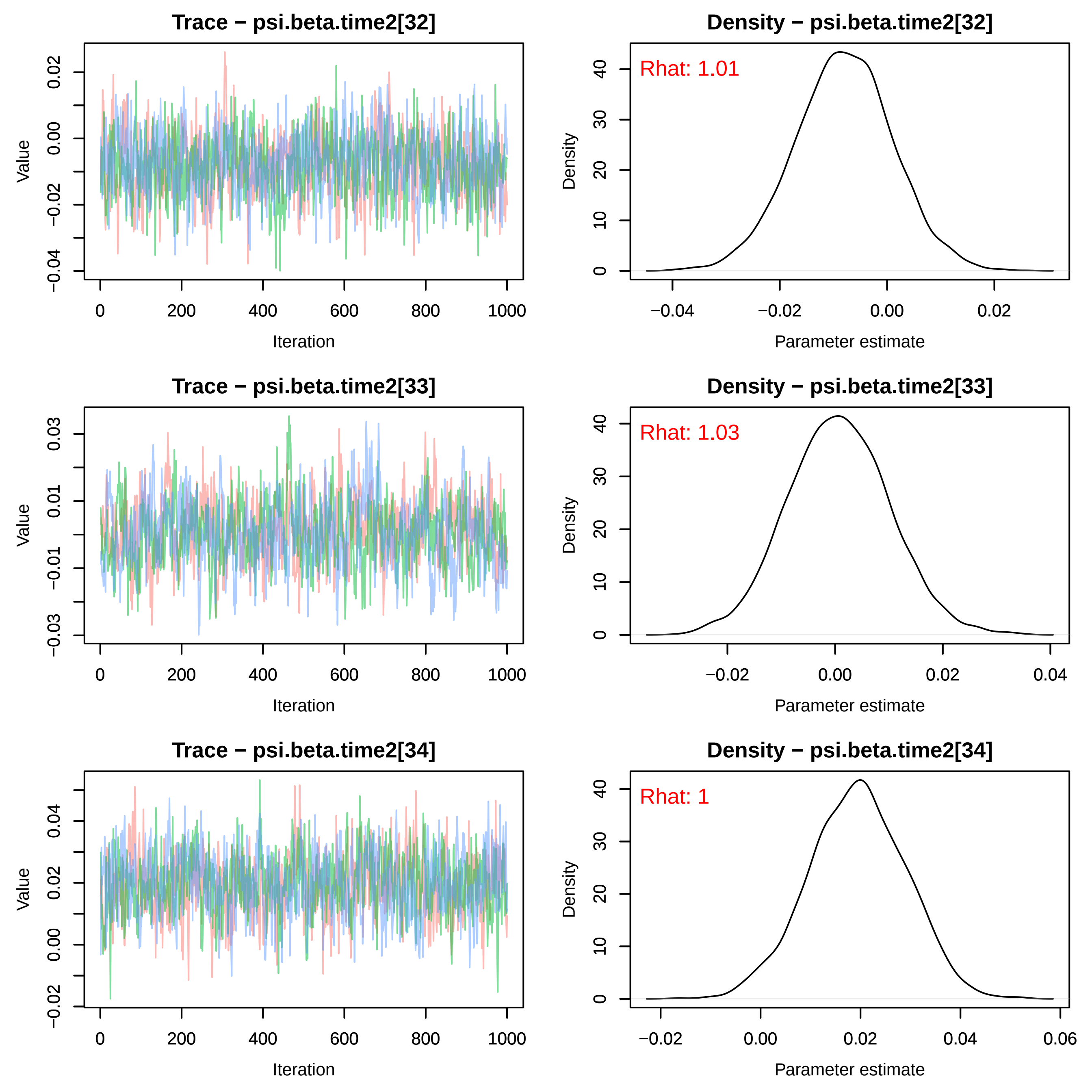

Trace − psi.beta.time2[32]
Density − psi.beta.time2[32]
Rhat: 1.01
Trace − psi.beta.time2[33]
Density − psi.beta.time2[33]
Rhat: 1.03
Trace − psi.beta.time2[34]
Density − psi.beta.time2[34]
Rhat: 1
Value
Iteration
Density
Parameter estimate

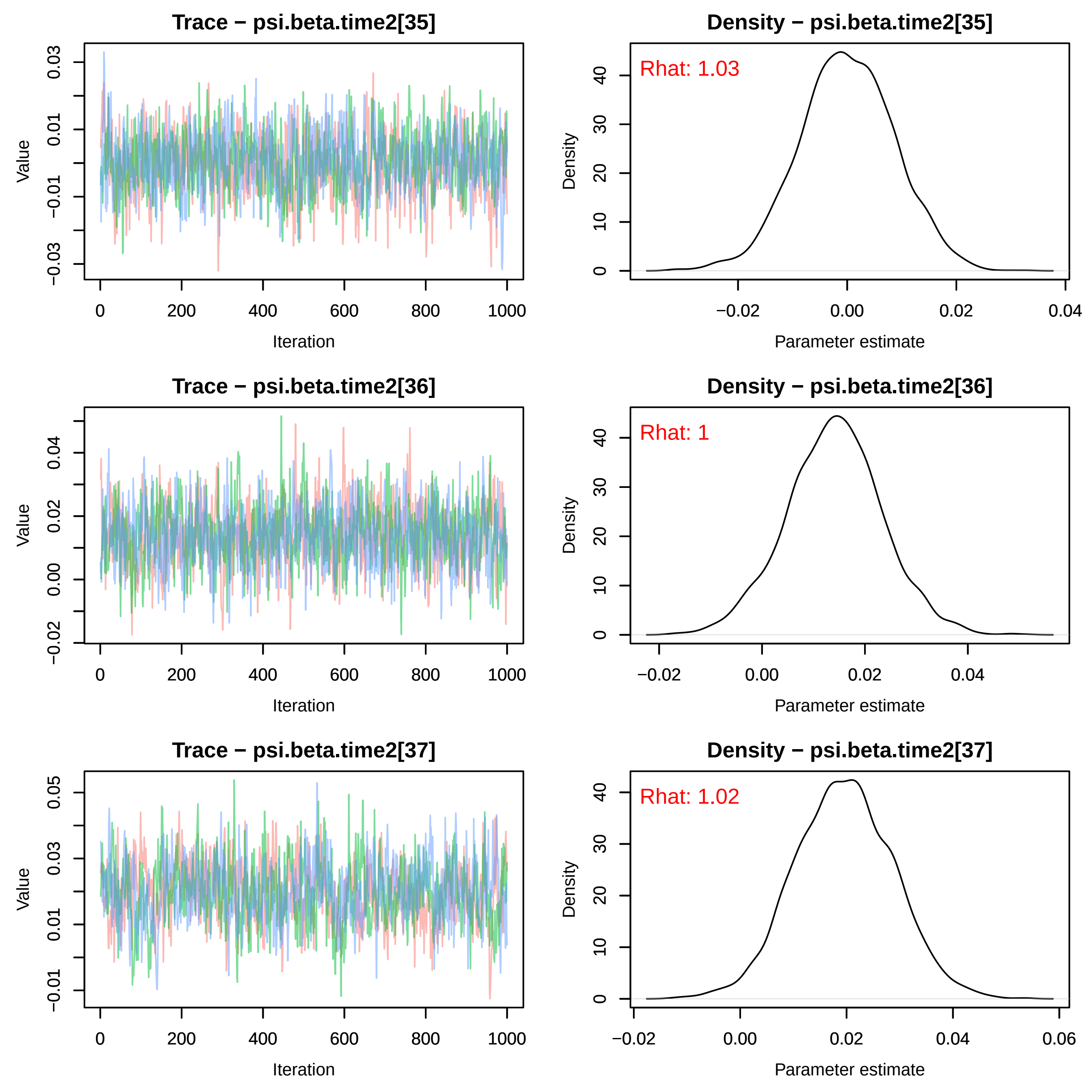
Trace − psi.beta.time2[35]
Value
0.03
0.01
−0.01
−0.03
0
200
400
600
800
1000
Iteration
Density − psi.beta.time2[35]
Rhat: 1.03
Density
0
10
20
30
40
−0.02
0.00
0.02
0.04
Parameter estimate
Trace − psi.beta.time2[36]
Value
0.04
0.02
0.00
−0.02
0
200
400
600
800
1000
Iteration
Density − psi.beta.time2[36]
Rhat: 1
Density
0
10
20
30
40
−0.02
0.00
0.02
0.04
Parameter estimate
Trace − psi.beta.time2[37]
Value
0.05
0.03
0.01
−0.01
0
200
400
600
800
1000
Iteration
Density − psi.beta.time2[37]
Rhat: 1.02
Density
0
10
20
30
40
−0.02
0.00
0.02
0.04
0.06
Parameter estimate

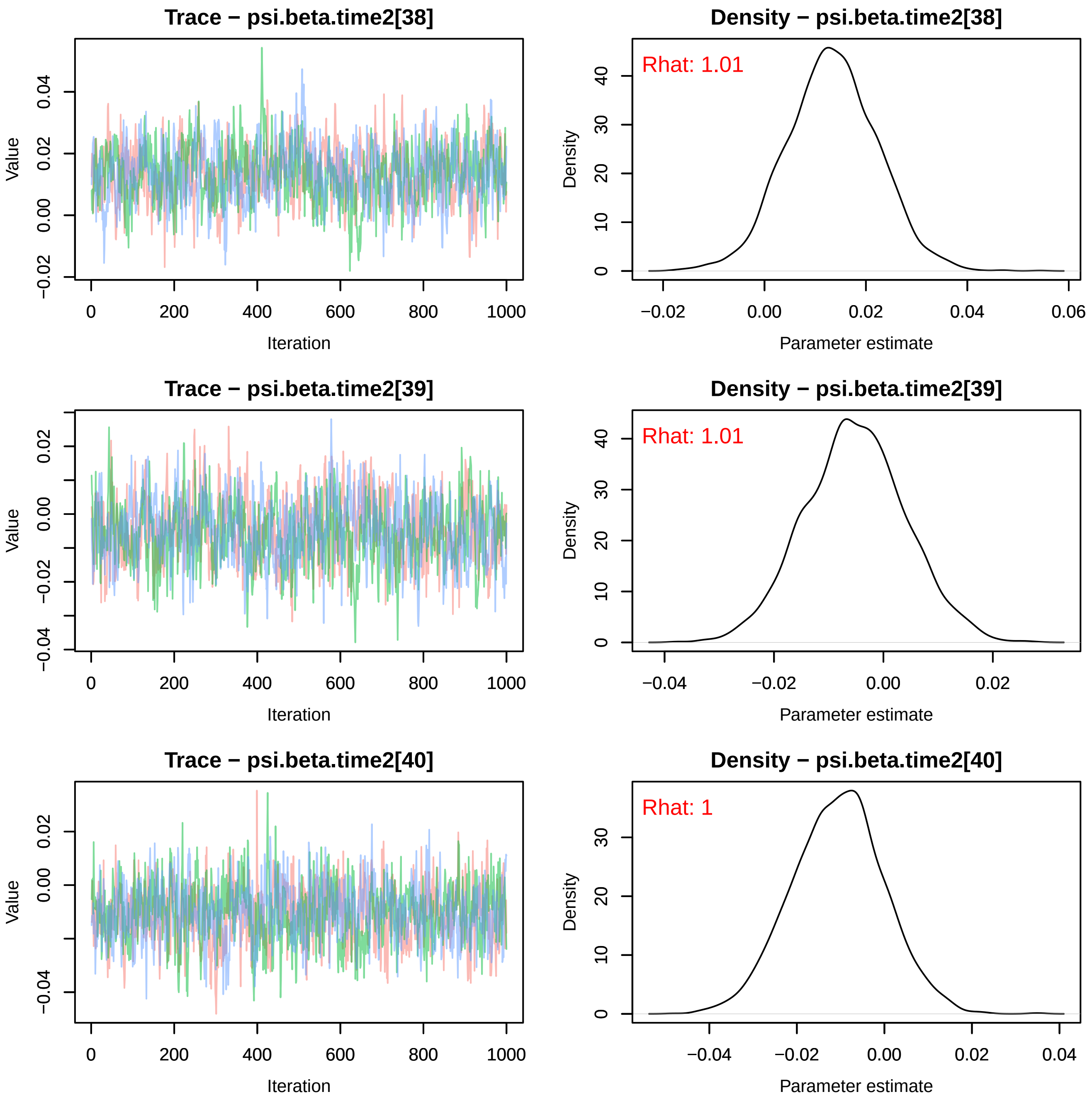
Trace − psi.beta.time2[38]
Value
Iteration
Density − psi.beta.time2[38]
Rhat: 1.01
Density
Parameter estimate
Trace − psi.beta.time2[39]
Value
Iteration
Density − psi.beta.time2[39]
Rhat: 1.01
Density
Parameter estimate
Trace − psi.beta.time2[40]
Value
Iteration
Density − psi.beta.time2[40]
Rhat: 1
Density
Parameter estimate

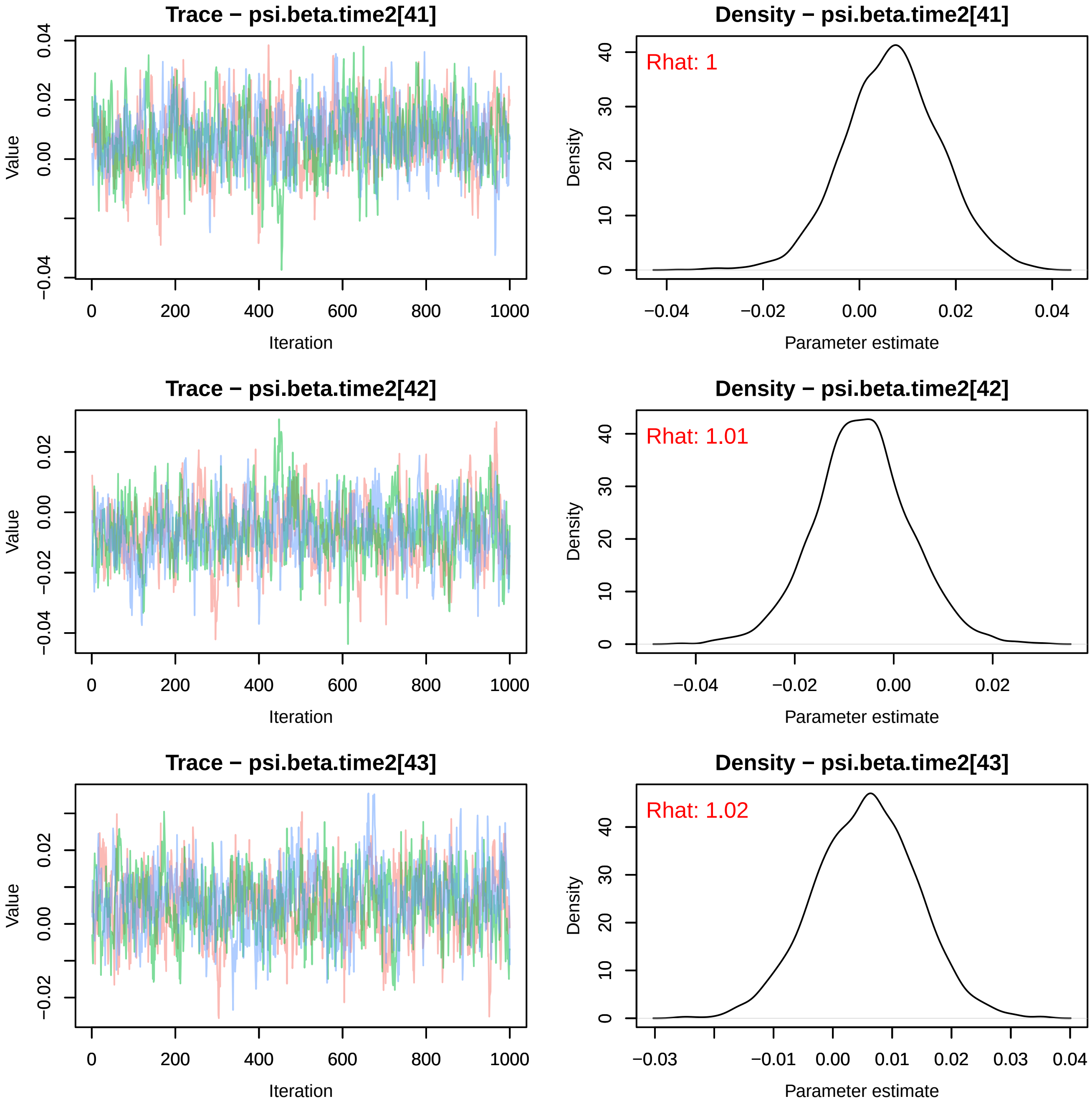
Trace − psi.beta.time2[41]
Value
Iteration
0.04
0.02
0.00
−0.04
0
200
400
600
800
1000
Density − psi.beta.time2[41]
Rhat: 1
Density
Parameter estimate
0
10
20
30
40
−0.04
−0.02
0.00
0.02
0.04
Trace − psi.beta.time2[42]
Value
Iteration
0.02
0.00
−0.02
−0.04
0
200
400
600
800
1000
Density − psi.beta.time2[42]
Rhat: 1.01
Density
Parameter estimate
0
10
20
30
40
−0.04
−0.02
0.00
0.02
Trace − psi.beta.time2[43]
Value
Iteration
0.02
0.00
−0.02
0
200
400
600
800
1000
Density − psi.beta.time2[43]
Rhat: 1.02
Density
Parameter estimate
0
10
20
30
40
−0.03
−0.01
0.00
0.01
0.02
0.03
0.04

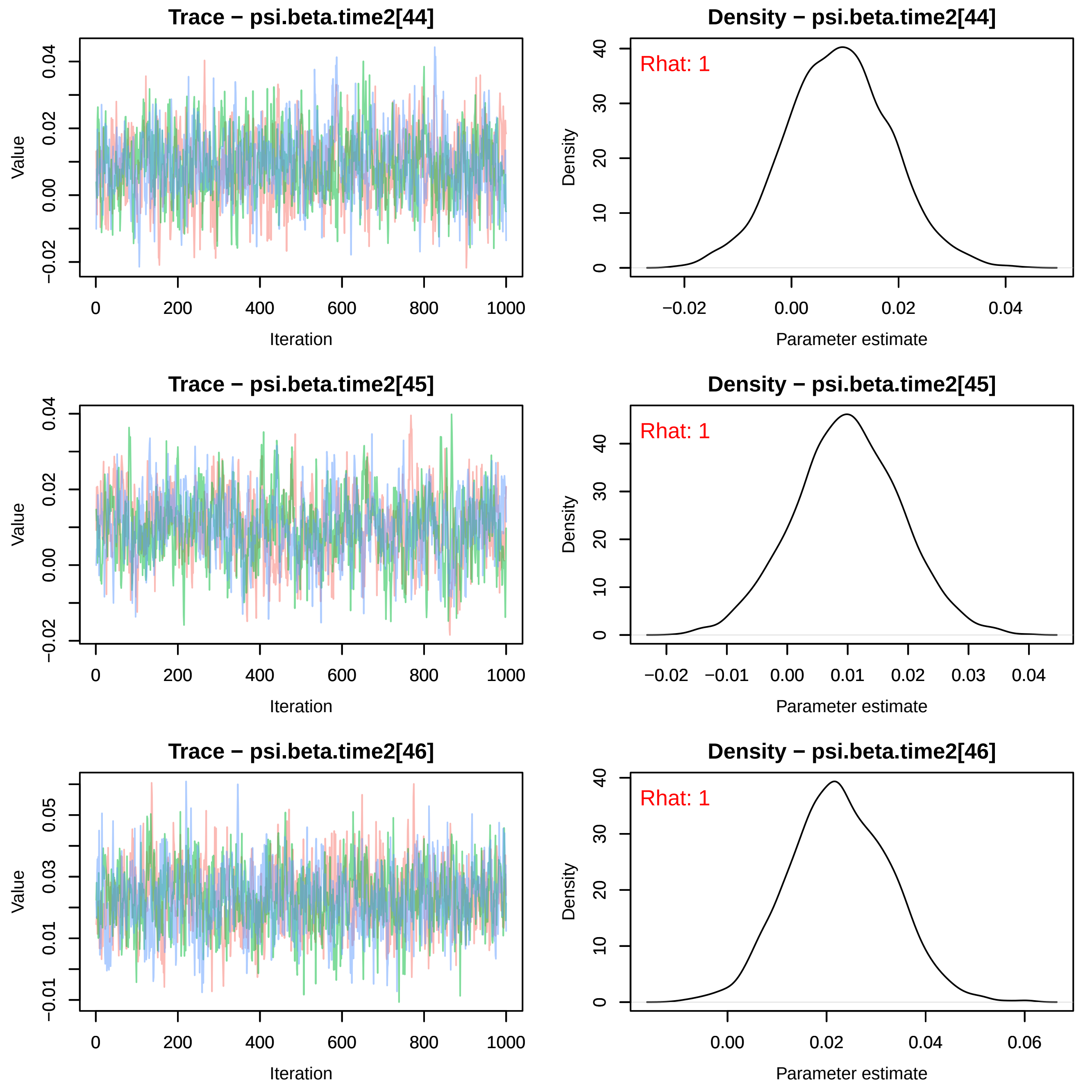
Trace − psi.beta.time2[44]
Value
Iteration
Density − psi.beta.time2[44]
Rhat: 1
Density
Parameter estimate
Trace − psi.beta.time2[45]
Value
Iteration
Density − psi.beta.time2[45]
Rhat: 1
Density
Parameter estimate
Trace − psi.beta.time2[46]
Value
Iteration
Density − psi.beta.time2[46]
Rhat: 1
Density
Parameter estimate

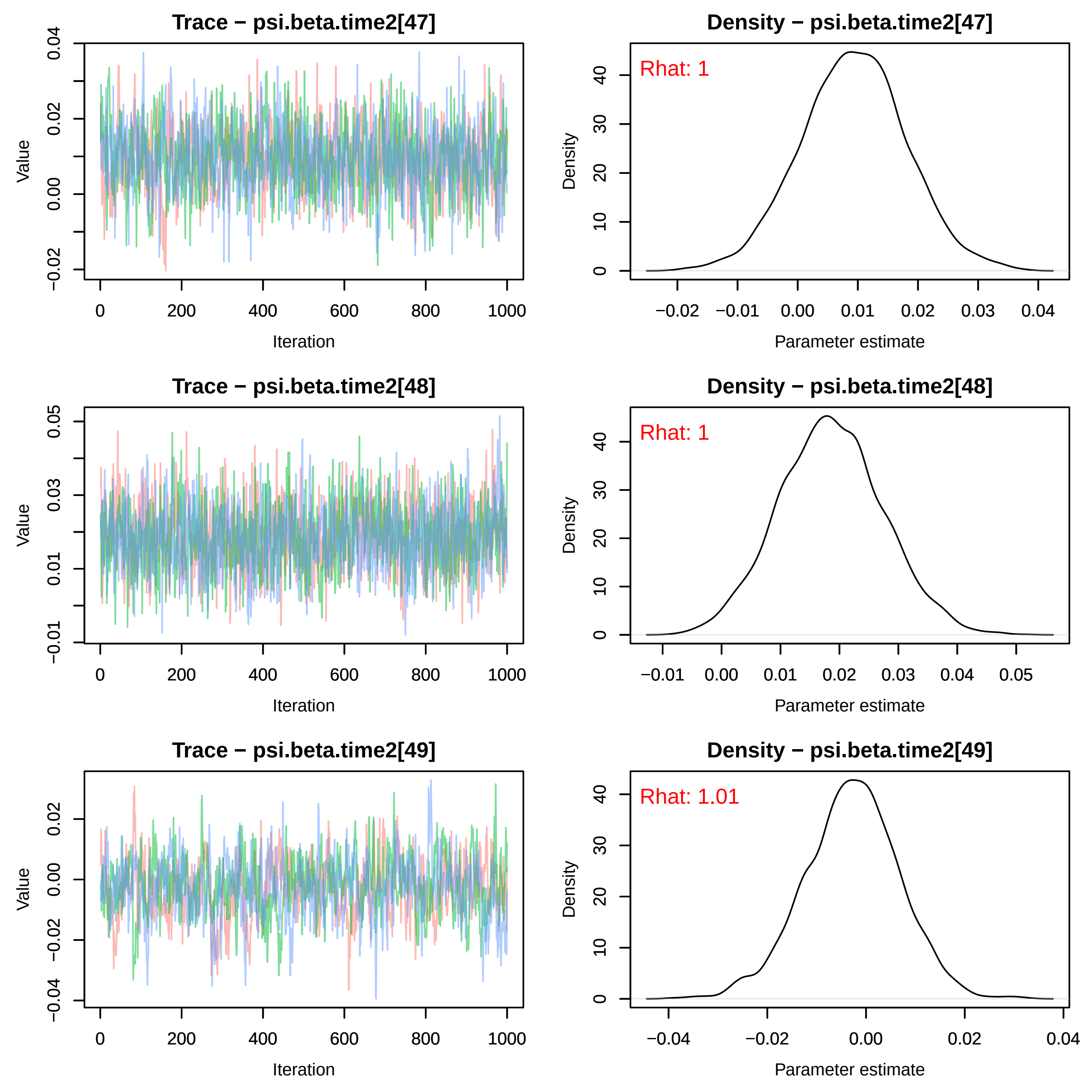
Trace − psi.beta.time2[47]
Value
Iteration
Density − psi.beta.time2[47]
Rhat: 1
Density
Parameter estimate
Trace − psi.beta.time2[48]
Value
Iteration
Density − psi.beta.time2[48]
Rhat: 1
Density
Parameter estimate
Trace − psi.beta.time2[49]
Value
Iteration
Density − psi.beta.time2[49]
Rhat: 1.01
Density
Parameter estimate

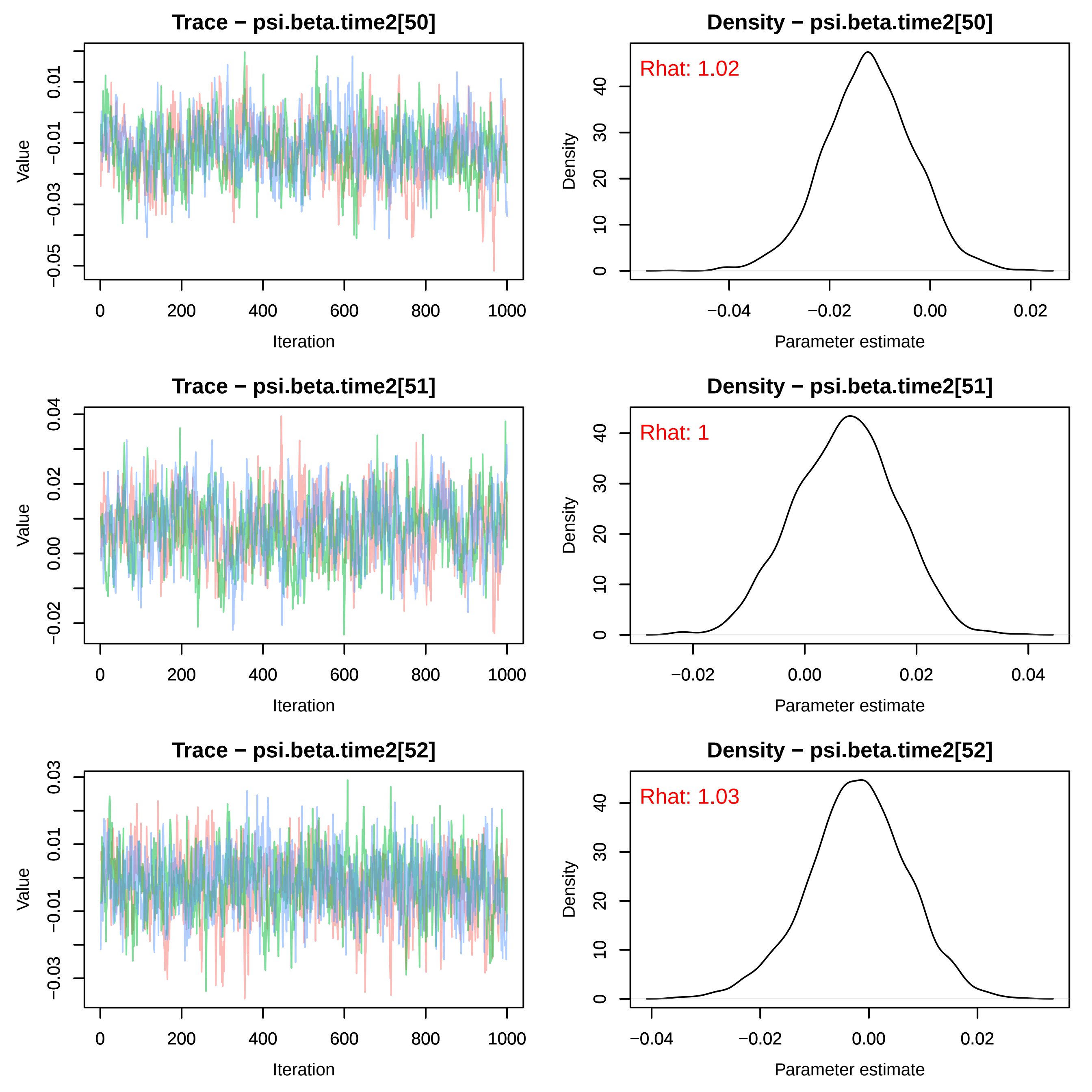
Trace − psi.beta.time2[50]
Value
Iteration
Density − psi.beta.time2[50]
Rhat: 1.02
Density
Parameter estimate
Trace − psi.beta.time2[51]
Value
Iteration
Density − psi.beta.time2[51]
Rhat: 1
Density
Parameter estimate
Trace − psi.beta.time2[52]
Value
Iteration
Density − psi.beta.time2[52]
Rhat: 1.03
Density
Parameter estimate

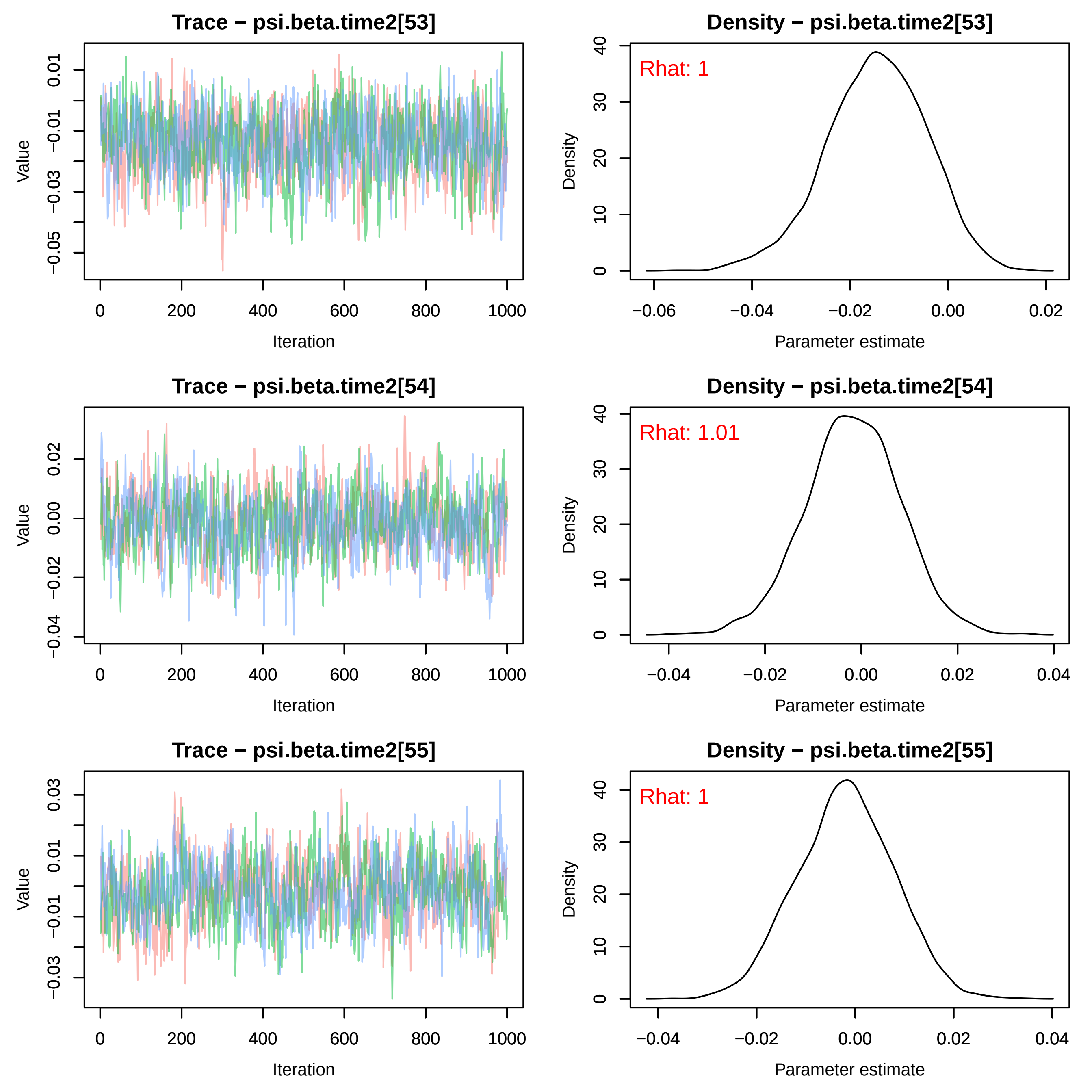
Trace − psi.beta.time2[53]
Density − psi.beta.time2[53]
Rhat: 1
Trace − psi.beta.time2[54]
Density − psi.beta.time2[54]
Rhat: 1.01
Trace − psi.beta.time2[55]
Density − psi.beta.time2[55]
Rhat: 1
Value
Iteration
Density
Parameter estimate

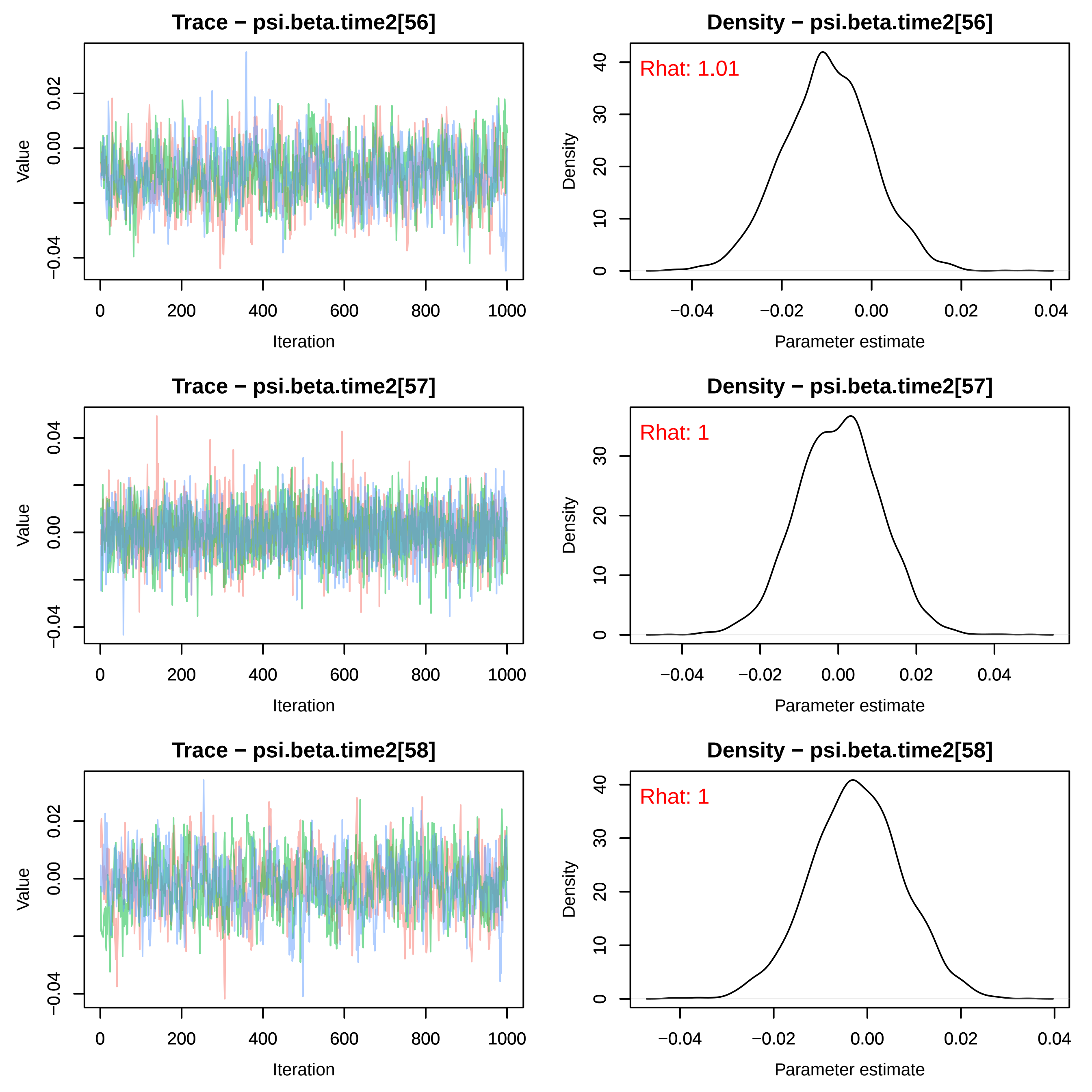

Trace − psi.beta.time2[56]
Value
0.02
0.00
−0.04
0
200
400
600
800
1000
Iteration
Density − psi.beta.time2[56]
Rhat: 1.01
Density
40
30
20
10
0
−0.04
−0.02
0.00
0.02
0.04
Parameter estimate
Trace − psi.beta.time2[57]
Value
0.04
0.00
−0.04
0
200
400
600
800
1000
Iteration
Density − psi.beta.time2[57]
Rhat: 1
Density
30
20
10
0
−0.04
−0.02
0.00
0.02
0.04
Parameter estimate
Trace − psi.beta.time2[58]
Value
0.02
0.00
−0.04
0
200
400
600
800
1000
Iteration
Density − psi.beta.time2[58]
Rhat: 1
Density
40
30
20
10
0
−0.04
−0.02
0.00
0.02
0.04
Parameter estimate

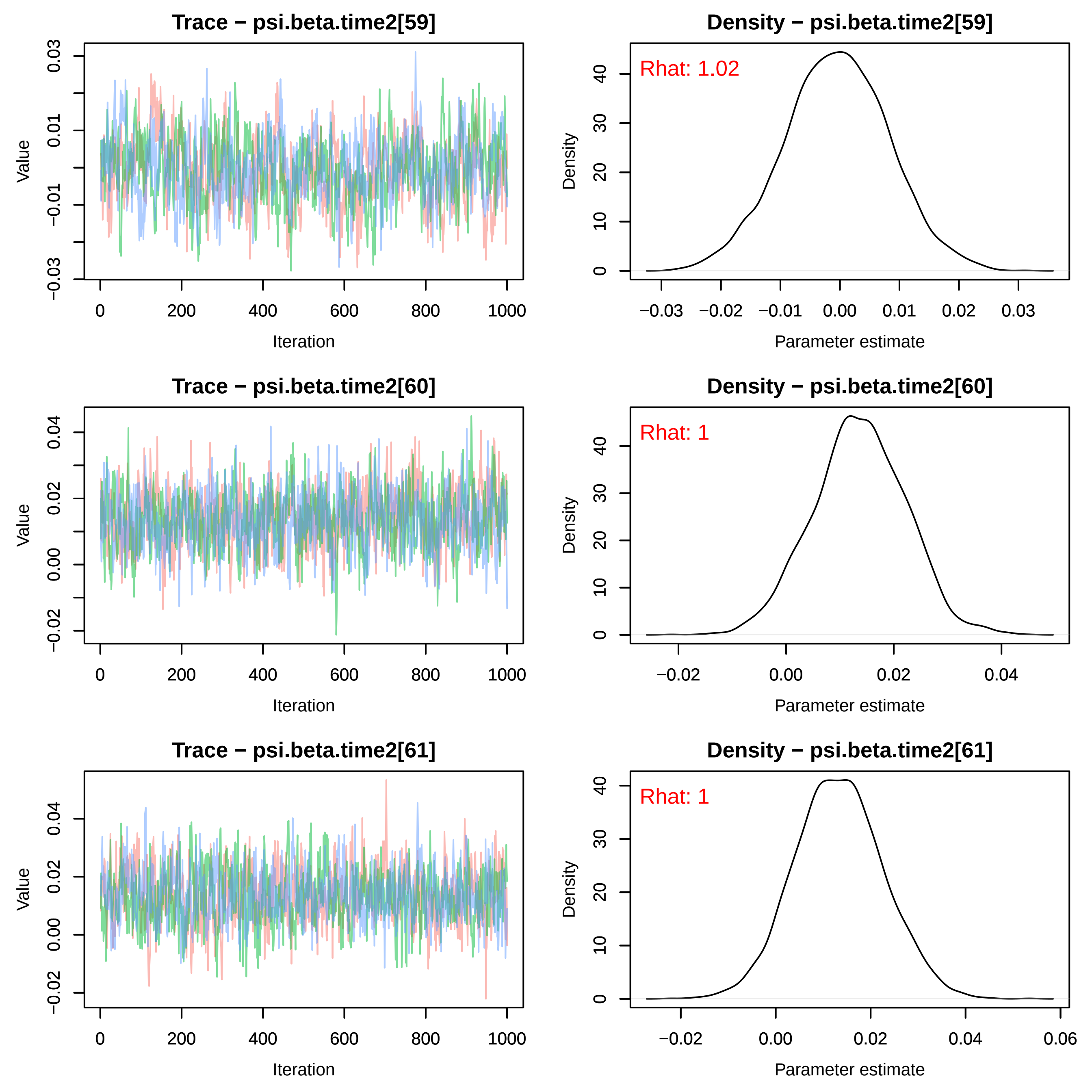
Trace − psi.beta.time2[59]
Density − psi.beta.time2[59]
Rhat: 1.02
Trace − psi.beta.time2[60]
Density − psi.beta.time2[60]
Rhat: 1
Trace − psi.beta.time2[61]
Density − psi.beta.time2[61]
Rhat: 1
Value
Iteration
Density
Parameter estimate

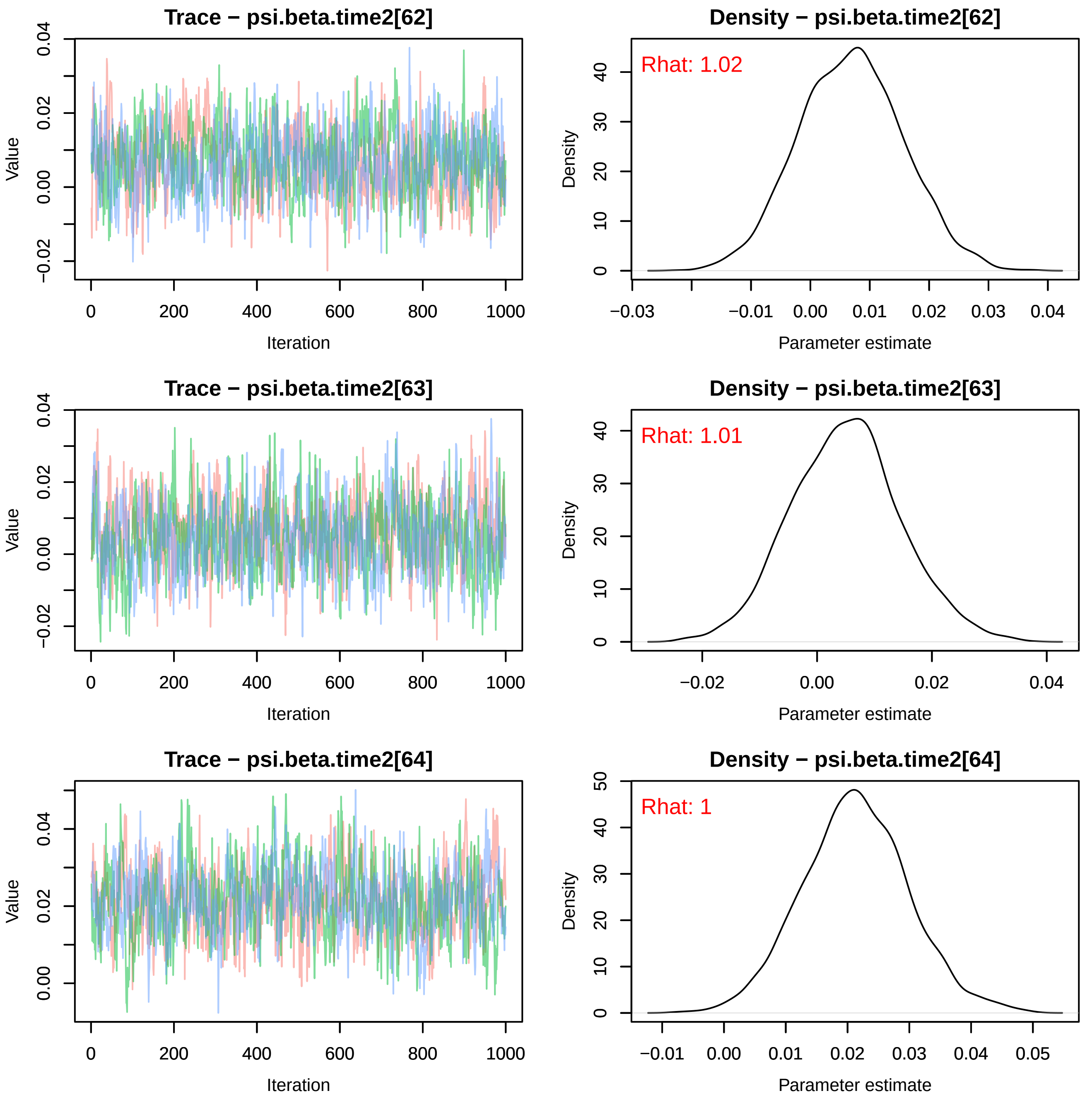
Trace − psi.beta.time2[62]
Value
−0.02
0.00
0.02
0.04
0
200
400
600
800
1000
Iteration
Density − psi.beta.time2[62]
Rhat: 1.02
Density
0
10
20
30
40
−0.03
−0.01
0.00
0.01
0.02
0.03
0.04
Parameter estimate
Trace − psi.beta.time2[63]
Value
−0.02
0.00
0.02
0.04
0
200
400
600
800
1000
Iteration
Density − psi.beta.time2[63]
Rhat: 1.01
Density
0
10
20
30
40
−0.02
0.00
0.02
0.04
Parameter estimate
Trace − psi.beta.time2[64]
Value
0.00
0.02
0.04
0
200
400
600
800
1000
Iteration
Density − psi.beta.time2[64]
Rhat: 1
Density
0
10
20
30
40
50
−0.01
0.00
0.01
0.02
0.03
0.04
0.05
Parameter estimate

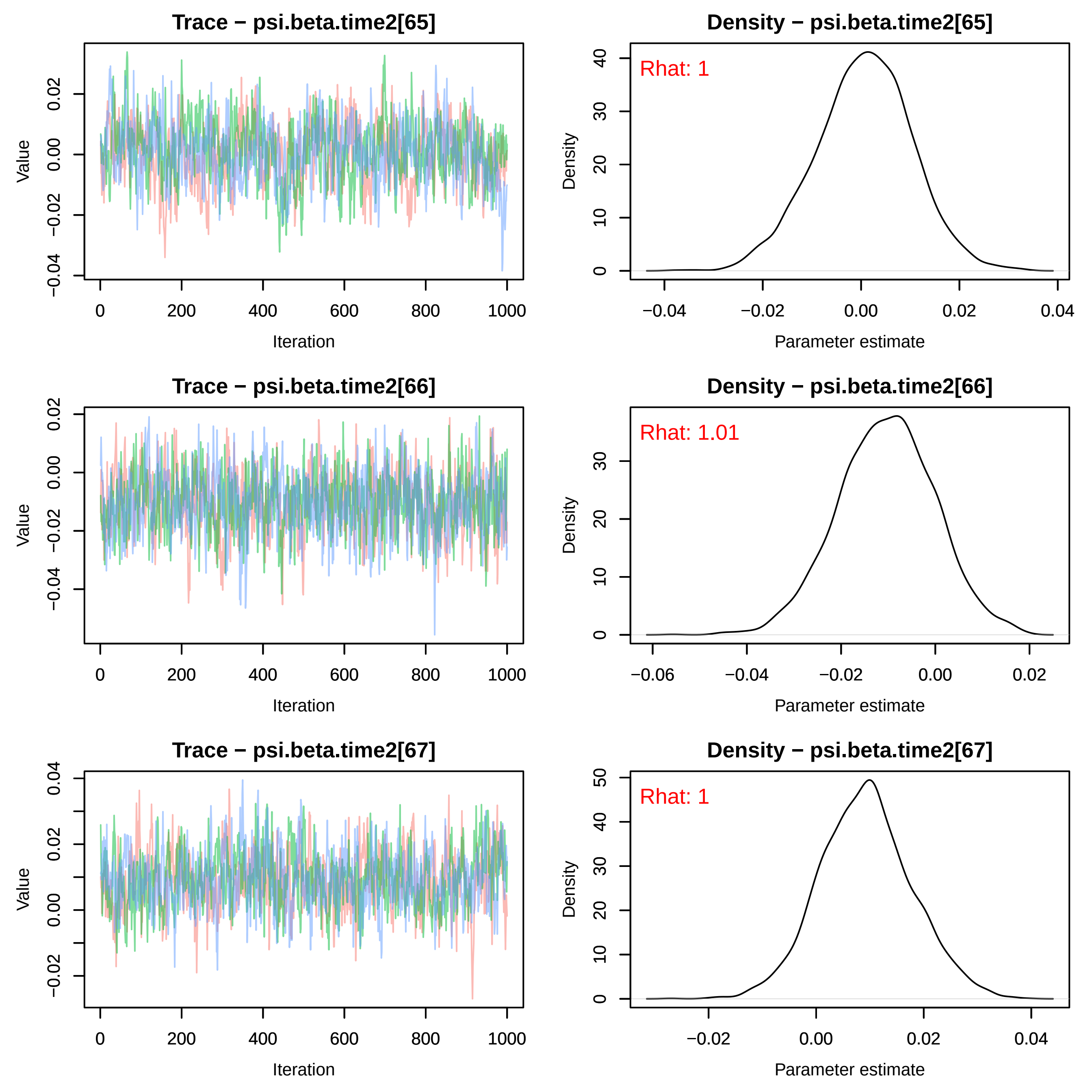

Trace − psi.beta.time2[65]
Value
Iteration
Density − psi.beta.time2[65]
Rhat: 1
Density
Parameter estimate
Trace − psi.beta.time2[66]
Value
Iteration
Density − psi.beta.time2[66]
Rhat: 1.01
Density
Parameter estimate
Trace − psi.beta.time2[67]
Value
Iteration
Density − psi.beta.time2[67]
Rhat: 1
Density
Parameter estimate

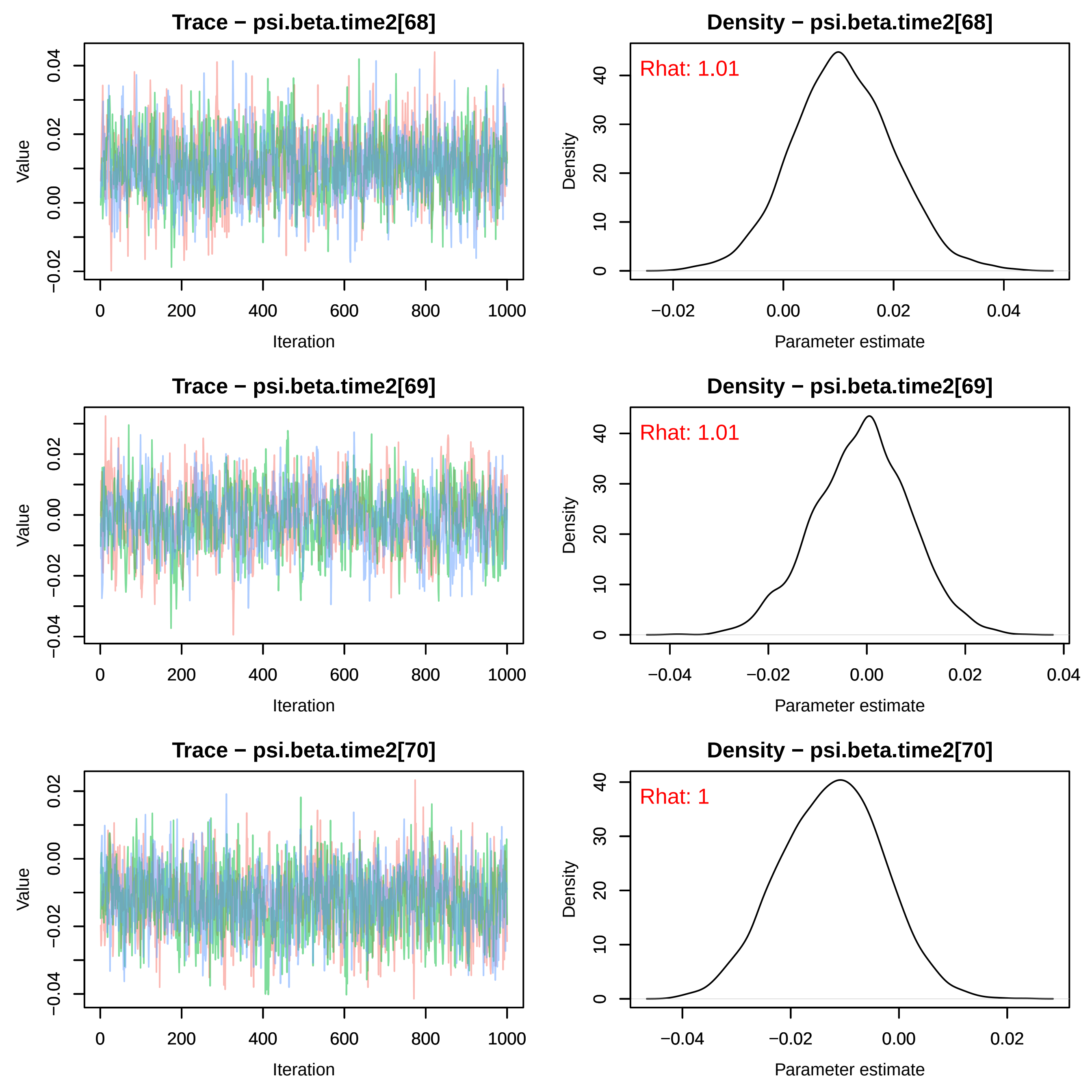
Trace − psi.beta.time2[68]
Density − psi.beta.time2[68]
Rhat: 1.01
Trace − psi.beta.time2[69]
Density − psi.beta.time2[69]
Rhat: 1.01
Trace − psi.beta.time2[70]
Density − psi.beta.time2[70]
Rhat: 1
Value
Iteration
Density
Parameter estimate

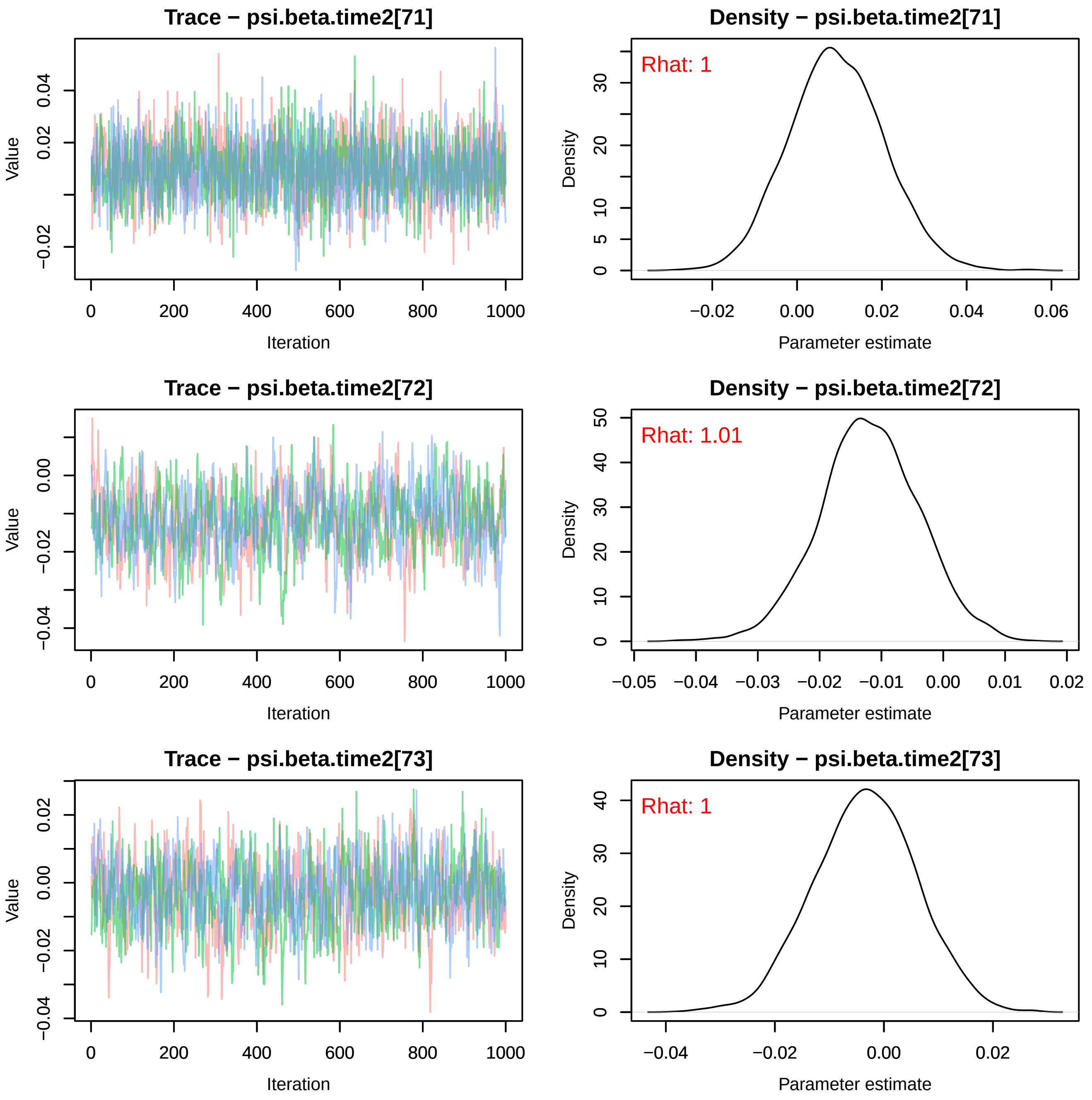
Trace − psi.beta.time2[71]
Value
Iteration
Density − psi.beta.time2[71]
Rhat: 1
Density
Parameter estimate
Trace − psi.beta.time2[72]
Value
Iteration
Density − psi.beta.time2[72]
Rhat: 1.01
Density
Parameter estimate
Trace − psi.beta.time2[73]
Value
Iteration
Density − psi.beta.time2[73]
Rhat: 1
Density
Parameter estimate

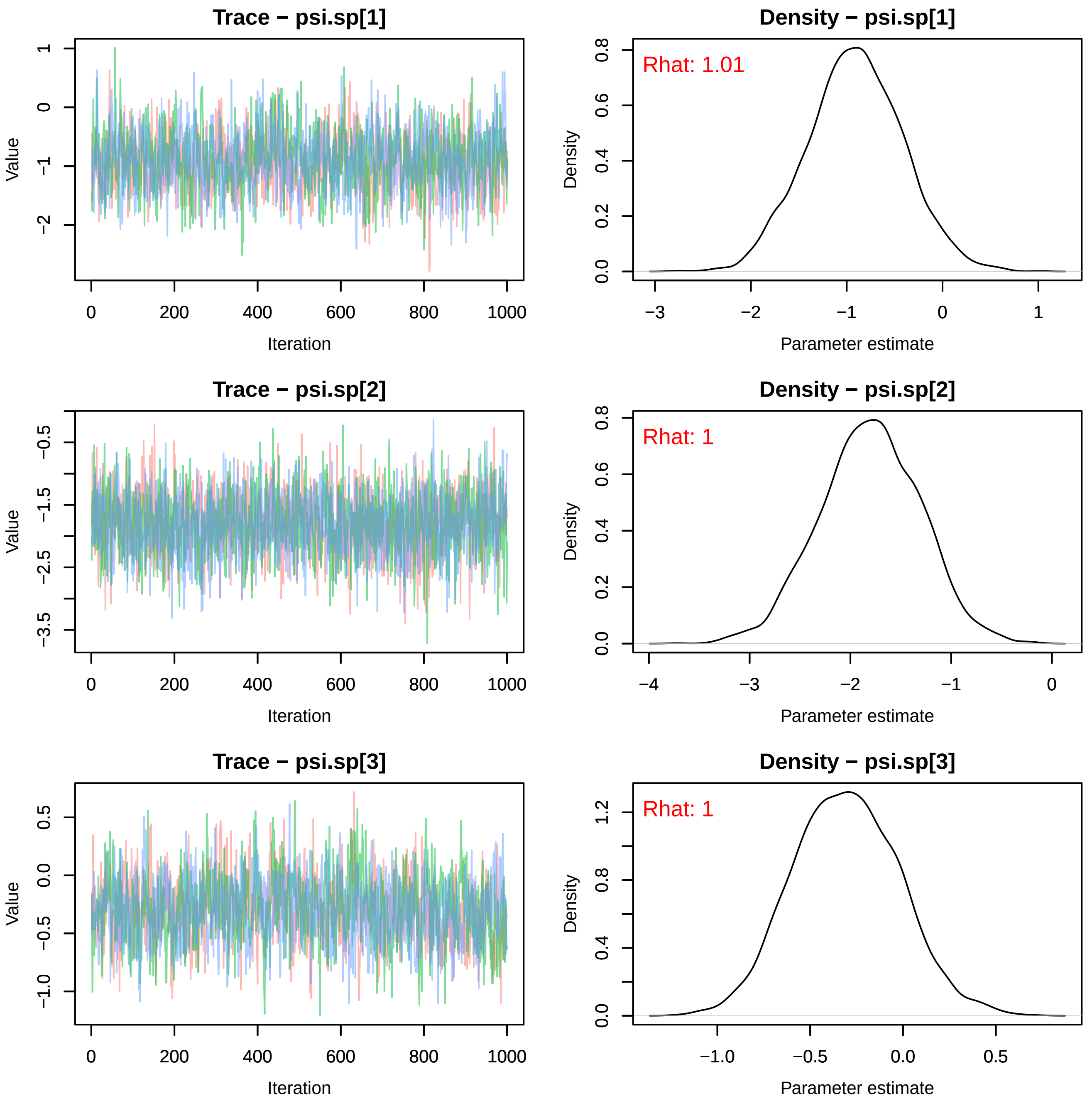
Trace − psi.sp[1]
Value
Iteration
0 200 400 600 800 1000
1 0 −1 −2
Density − psi.sp[1]
Rhat: 1.01
Density
Parameter estimate
−3 −2 −1 0 1
0.0 0.2 0.4 0.6 0.8
Trace − psi.sp[2]
Value
Iteration
−0.5 −1.5 −2.5 −3.5
Density − psi.sp[2]
Rhat: 1
Density
Parameter estimate
−4 −3 −2 −1 0
Trace − psi.sp[3]
Value
Iteration
0.5 0.0 −0.5 −1.0
Density − psi.sp[3]
Rhat: 1
Density
Parameter estimate
−1.0 −0.5 0.0 0.5
0.0 0.4 0.8 1.2

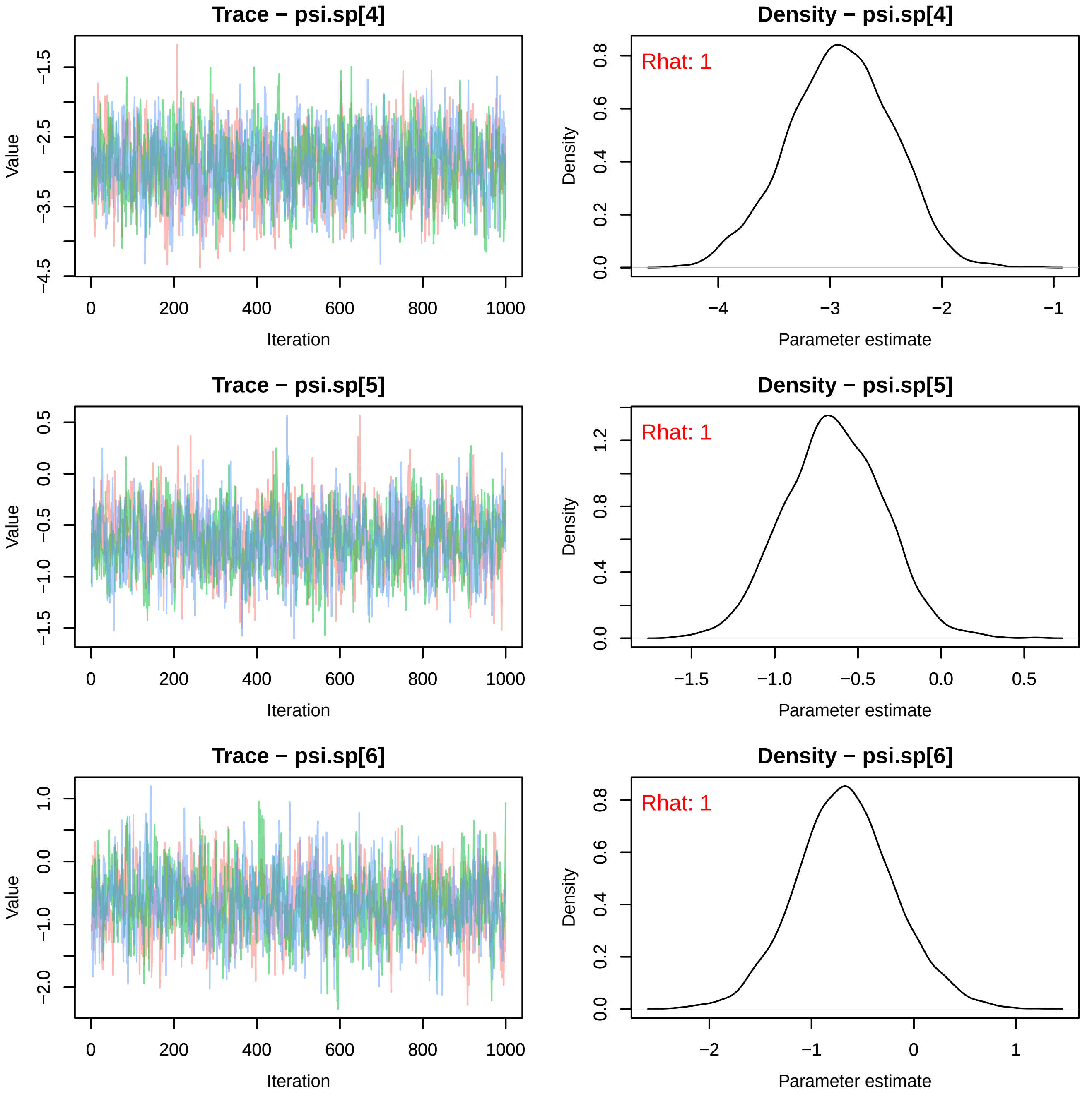

Trace − psi.sp[4]
Value
Iteration
−1.5
−2.5
−3.5
−4.5
0
200
400
600
800
1000
Density − psi.sp[4]
Rhat: 1
Density
0.0
0.2
0.4
0.6
0.8
−4
−3
−2
−1
Parameter estimate
Trace − psi.sp[5]
Value
0.5
0.0
−0.5
−1.0
−1.5
Iteration
Density − psi.sp[5]
Rhat: 1
Density
0.0
0.4
0.8
1.2
−1.5
−1.0
−0.5
0.0
0.5
Parameter estimate
Trace − psi.sp[6]
Value
1.0
0.0
−1.0
−2.0
Iteration
Density − psi.sp[6]
Rhat: 1
Density
0.0
0.2
0.4
0.6
0.8
−2
−1
0
1
Parameter estimate

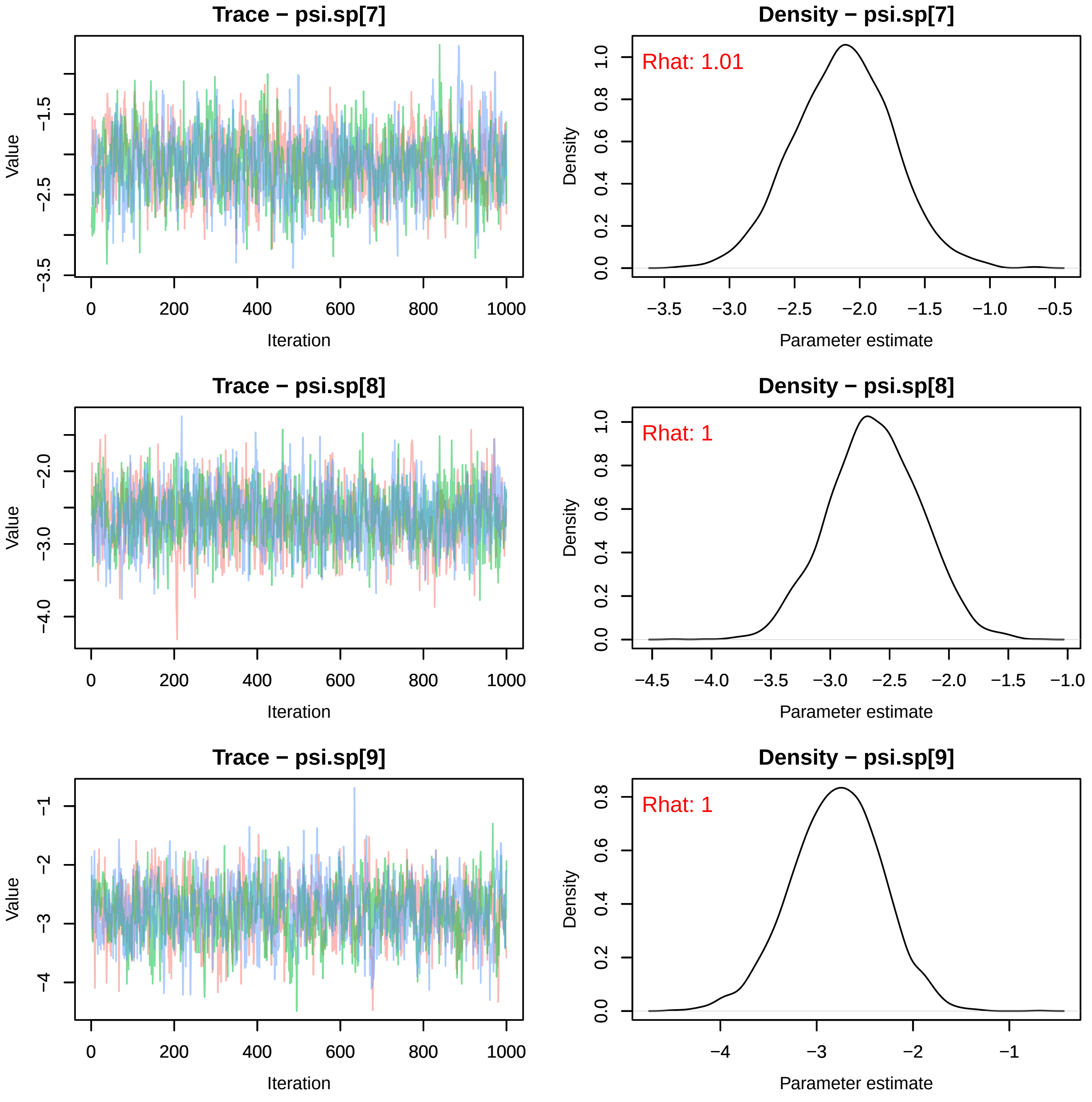
Trace − psi.sp[7]
Density − psi.sp[7]
Rhat: 1.01
Trace − psi.sp[8]
Density − psi.sp[8]
Rhat: 1
Trace − psi.sp[9]
Density − psi.sp[9]
Rhat: 1
Value
Density
Iteration
Parameter estimate
0
200
400
600
800
1000
−1.5
−2.5
−3.5
−2.0
−3.0
−4.0
−1
−2
−3
−4
0.0
0.2
0.4
0.6
0.8
1.0
−3.5
−3.0
−2.5
−2.0
−1.5
−1.0
−0.5
−4.5
−4.0

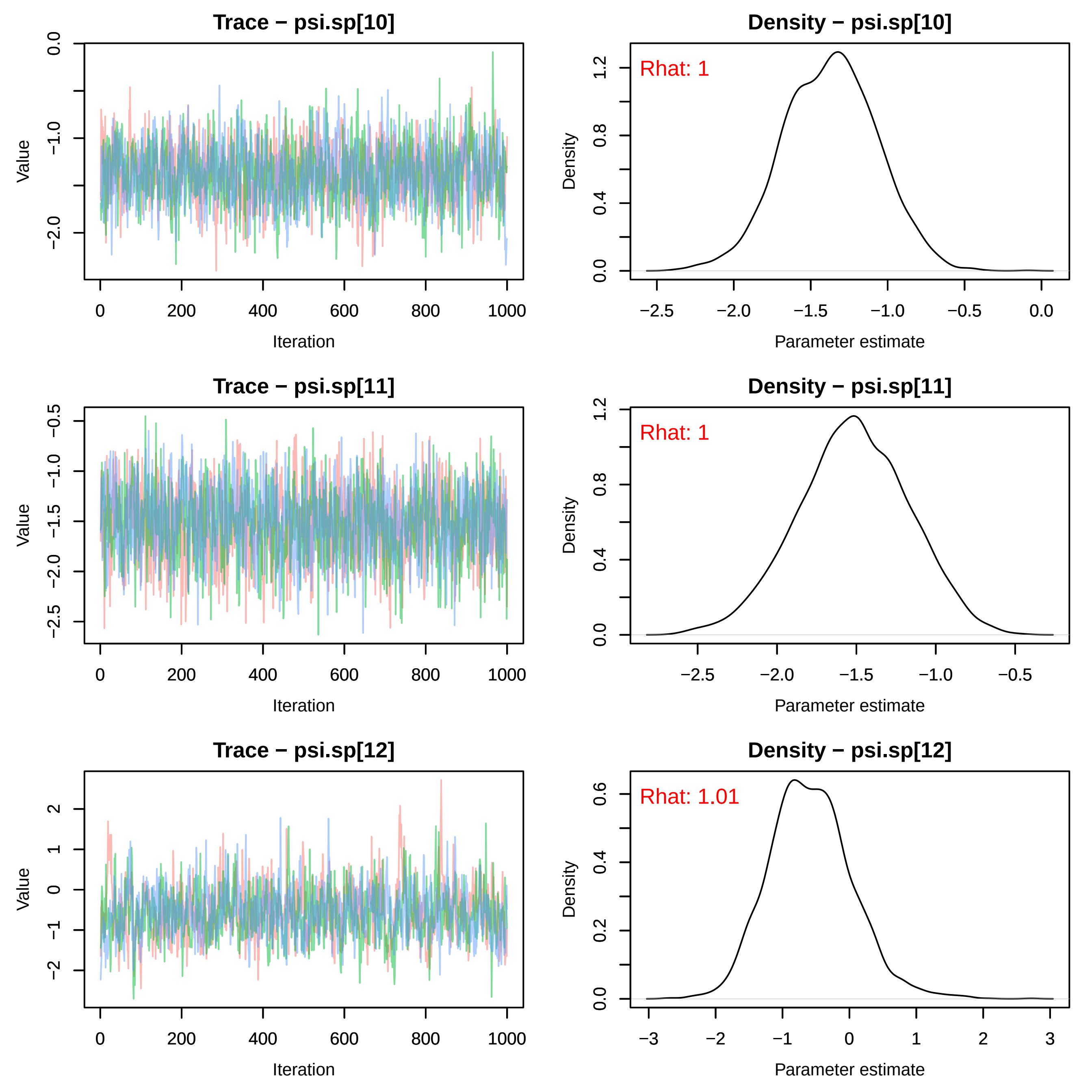
Trace − psi.sp[10]
Value
Iteration
Density − psi.sp[10]
Rhat: 1
Density
Parameter estimate
Trace − psi.sp[11]
Value
Iteration
Density − psi.sp[11]
Rhat: 1
Density
Parameter estimate
Trace − psi.sp[12]
Value
Iteration
Density − psi.sp[12]
Rhat: 1.01
Density
Parameter estimate

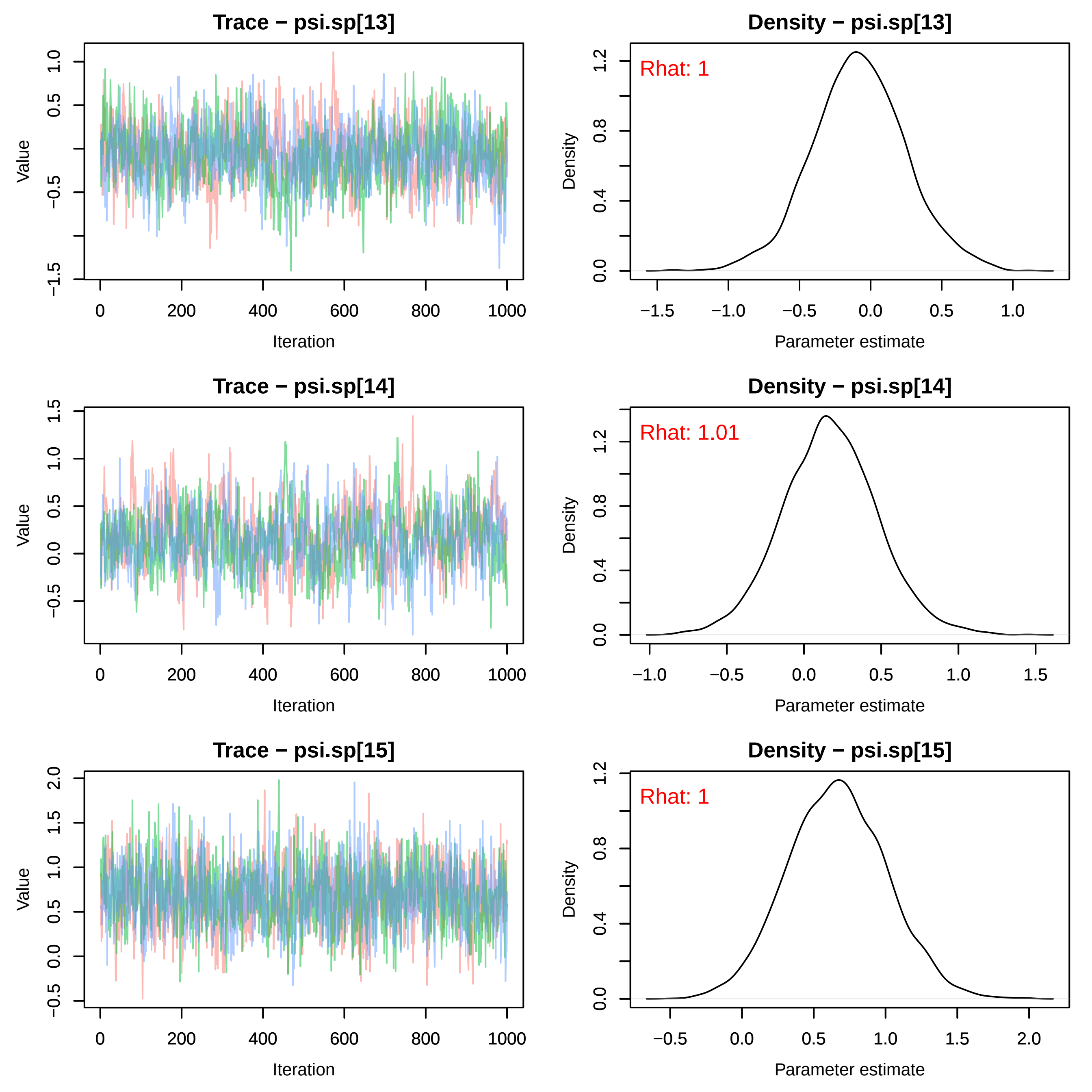
Trace − psi.sp[13]
Density − psi.sp[13]
Rhat: 1
Trace − psi.sp[14]
Density − psi.sp[14]
Rhat: 1.01
Trace − psi.sp[15]
Density − psi.sp[15]
Rhat: 1
Value
Iteration
Density
Parameter estimate

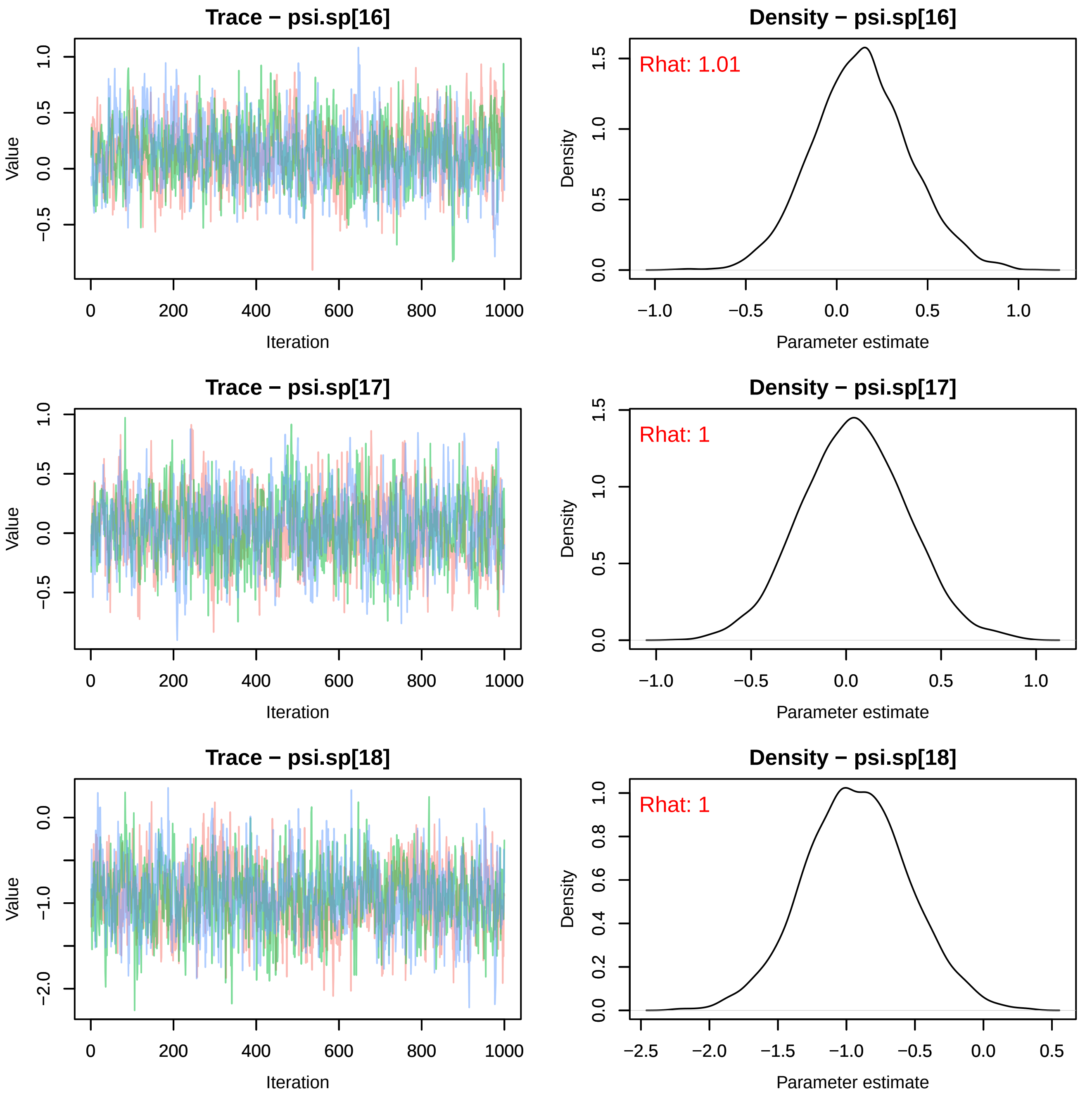

Trace − psi.sp[16]
Value
1.0
0.5
0.0
−0.5
0
200
400
600
800
1000
Iteration
Density − psi.sp[16]
Rhat: 1.01
Density
1.5
1.0
0.5
0.0
−1.0
−0.5
0.0
0.5
1.0
Parameter estimate
Trace − psi.sp[17]
Value
1.0
0.5
0.0
−0.5
0
200
400
600
800
1000
Iteration
Density − psi.sp[17]
Rhat: 1
Density
1.5
1.0
0.5
0.0
−1.0
−0.5
0.0
0.5
1.0
Parameter estimate
Trace − psi.sp[18]
Value
0.0
−1.0
−2.0
0
200
400
600
800
1000
Iteration
Density − psi.sp[18]
Rhat: 1
Density
1.0
0.8
0.6
0.4
0.2
0.0
−2.5
−2.0
−1.5
−1.0
−0.5
0.0
0.5
Parameter estimate

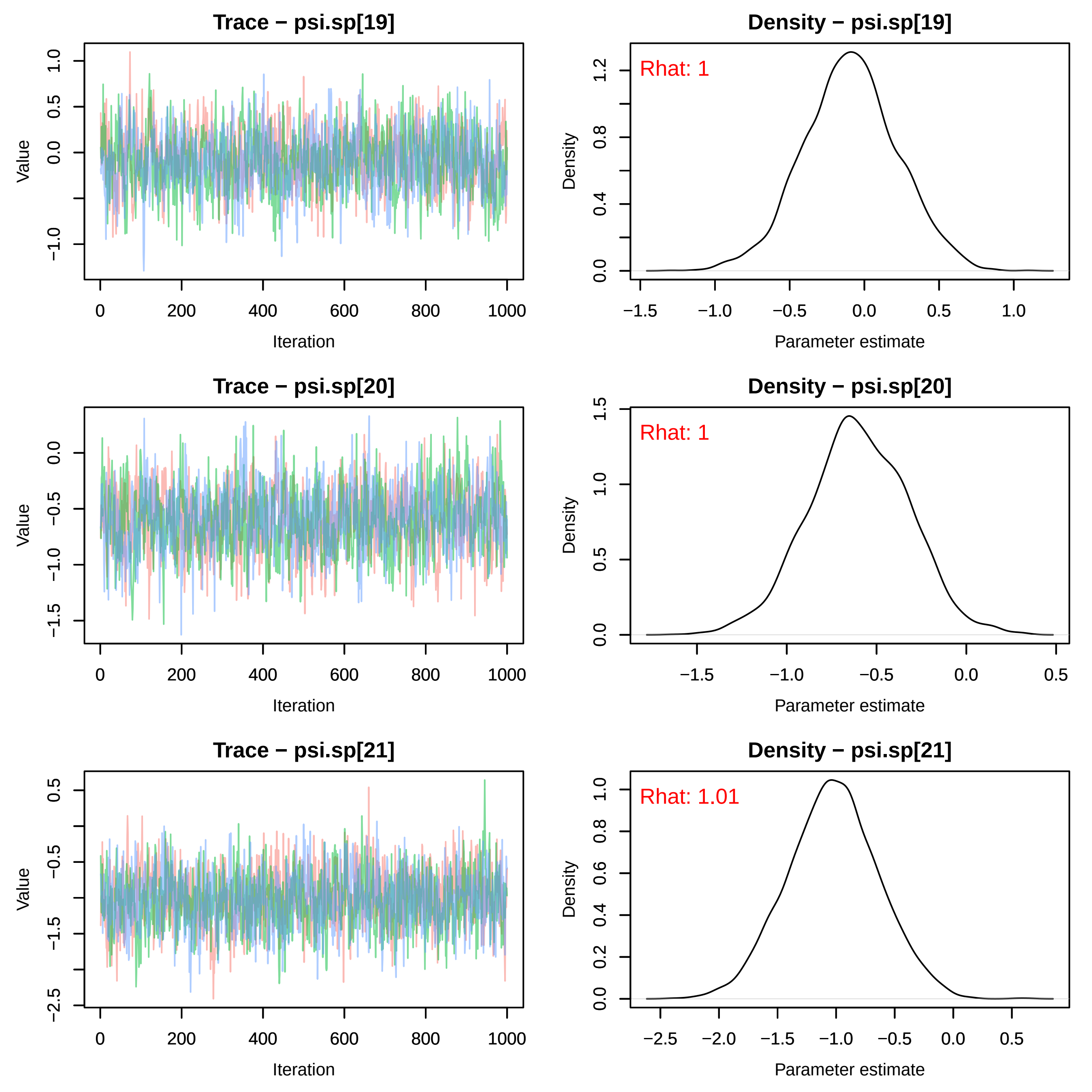

Trace − psi.sp[19]
Value
Iteration
Density − psi.sp[19]
Rhat: 1
Density
Parameter estimate
Trace − psi.sp[20]
Value
Iteration
Density − psi.sp[20]
Rhat: 1
Density
Parameter estimate
Trace − psi.sp[21]
Value
Iteration
Density − psi.sp[21]
Rhat: 1.01
Density
Parameter estimate

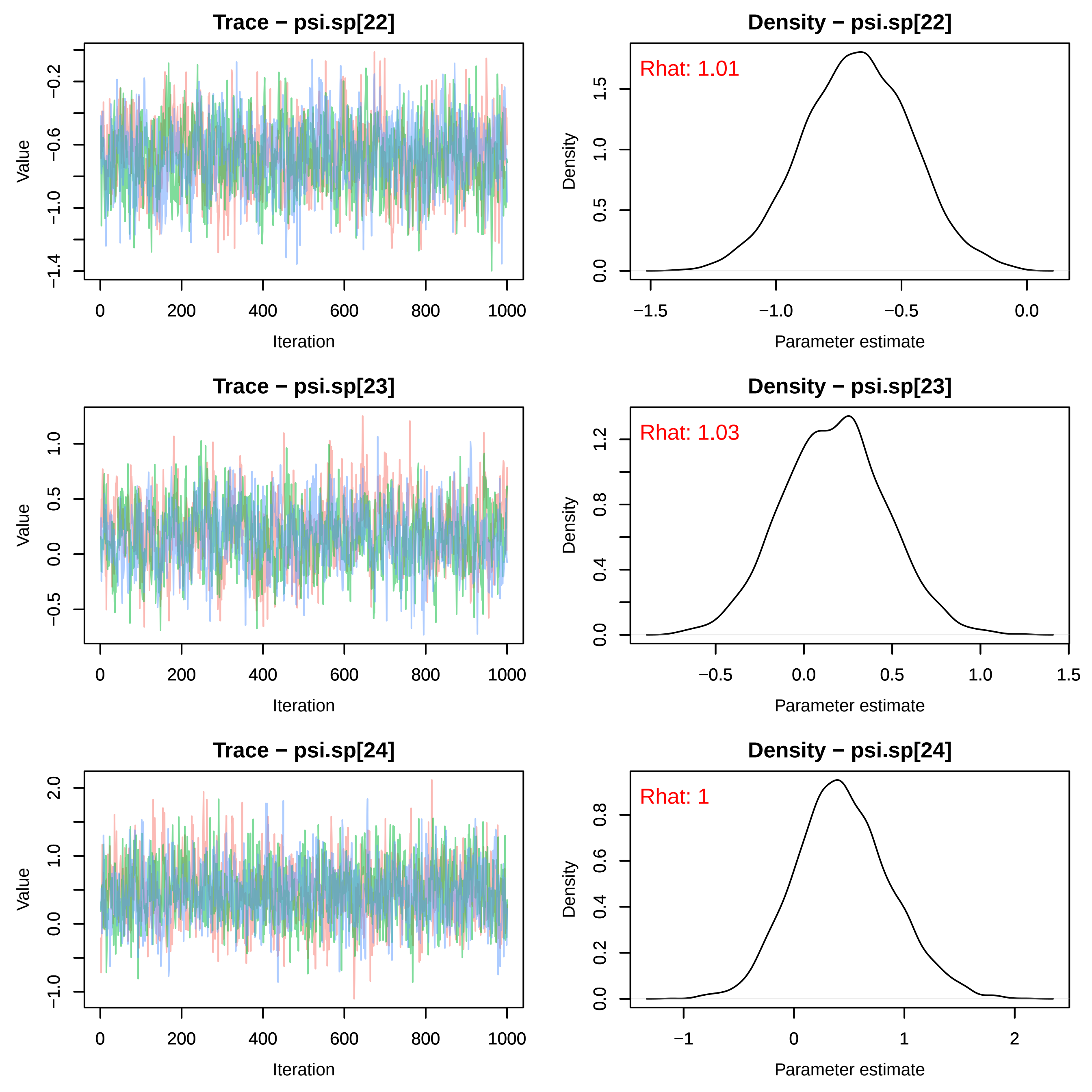
Trace − psi.sp[22]
Density − psi.sp[22]
Rhat: 1.01
Trace − psi.sp[23]
Density − psi.sp[23]
Rhat: 1.03
Trace − psi.sp[24]
Density − psi.sp[24]
Rhat: 1
Value
Iteration
Density
Parameter estimate

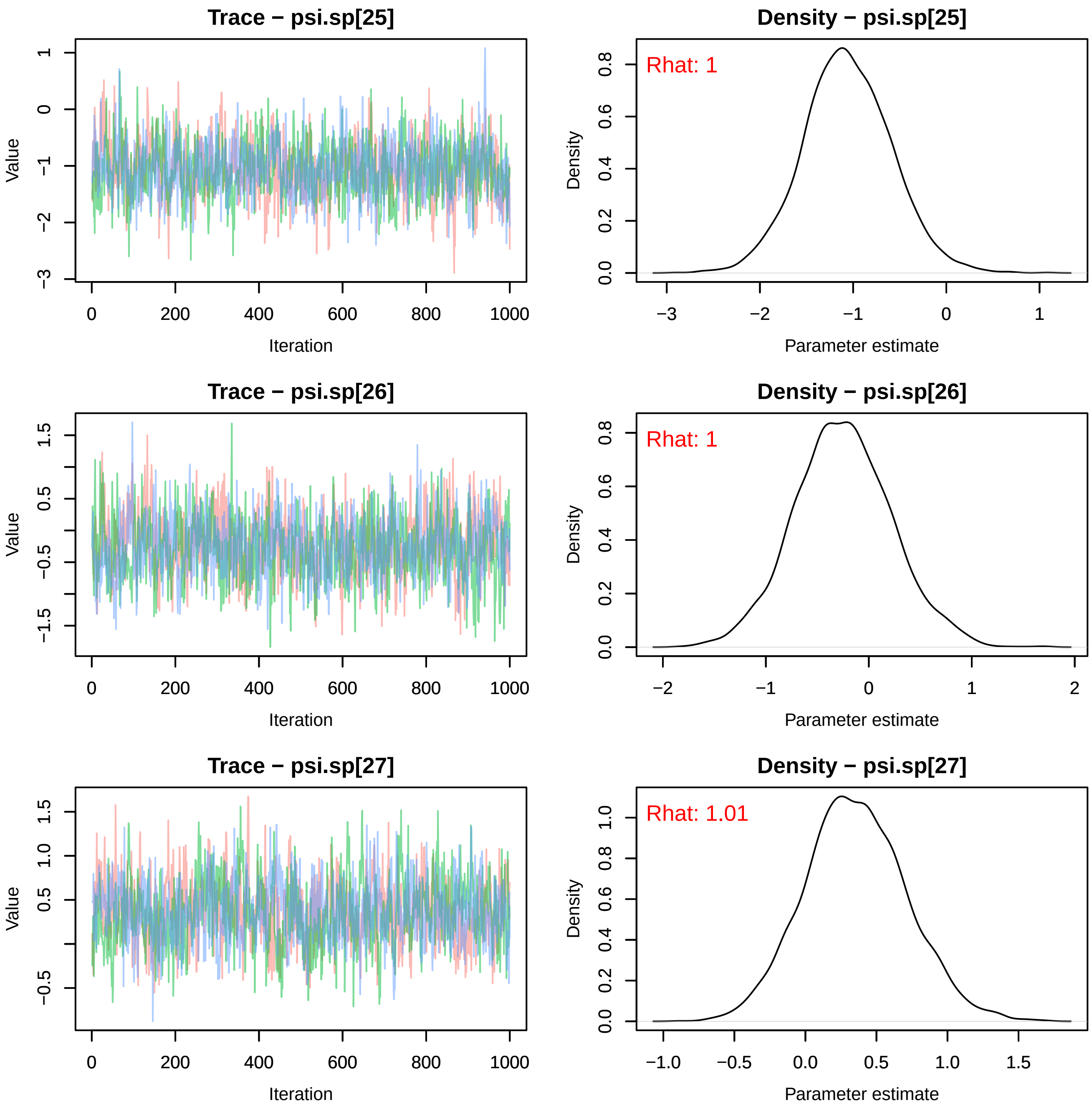
Trace − psi.sp[25]
Density − psi.sp[25]
Rhat: 1
Trace − psi.sp[26]
Density − psi.sp[26]
Rhat: 1
Trace − psi.sp[27]
Density − psi.sp[27]
Rhat: 1.01
Value
Iteration
Density
Parameter estimate

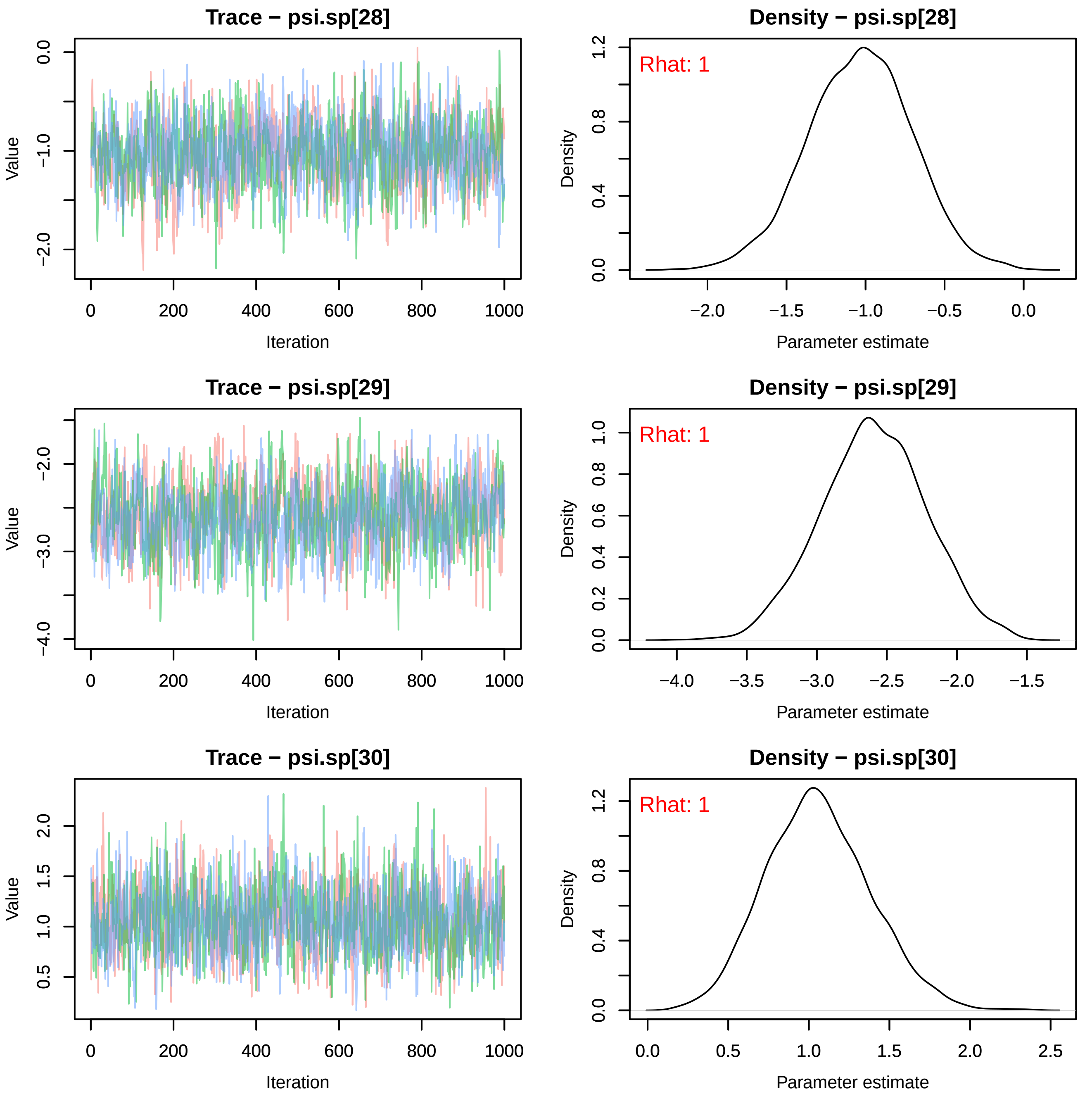
Trace − psi.sp[28]
Value
Iteration
Density − psi.sp[28]
Rhat: 1
Density
Parameter estimate
Trace − psi.sp[29]
Value
Iteration
Density − psi.sp[29]
Rhat: 1
Density
Parameter estimate
Trace − psi.sp[30]
Value
Iteration
Density − psi.sp[30]
Rhat: 1
Density
Parameter estimate

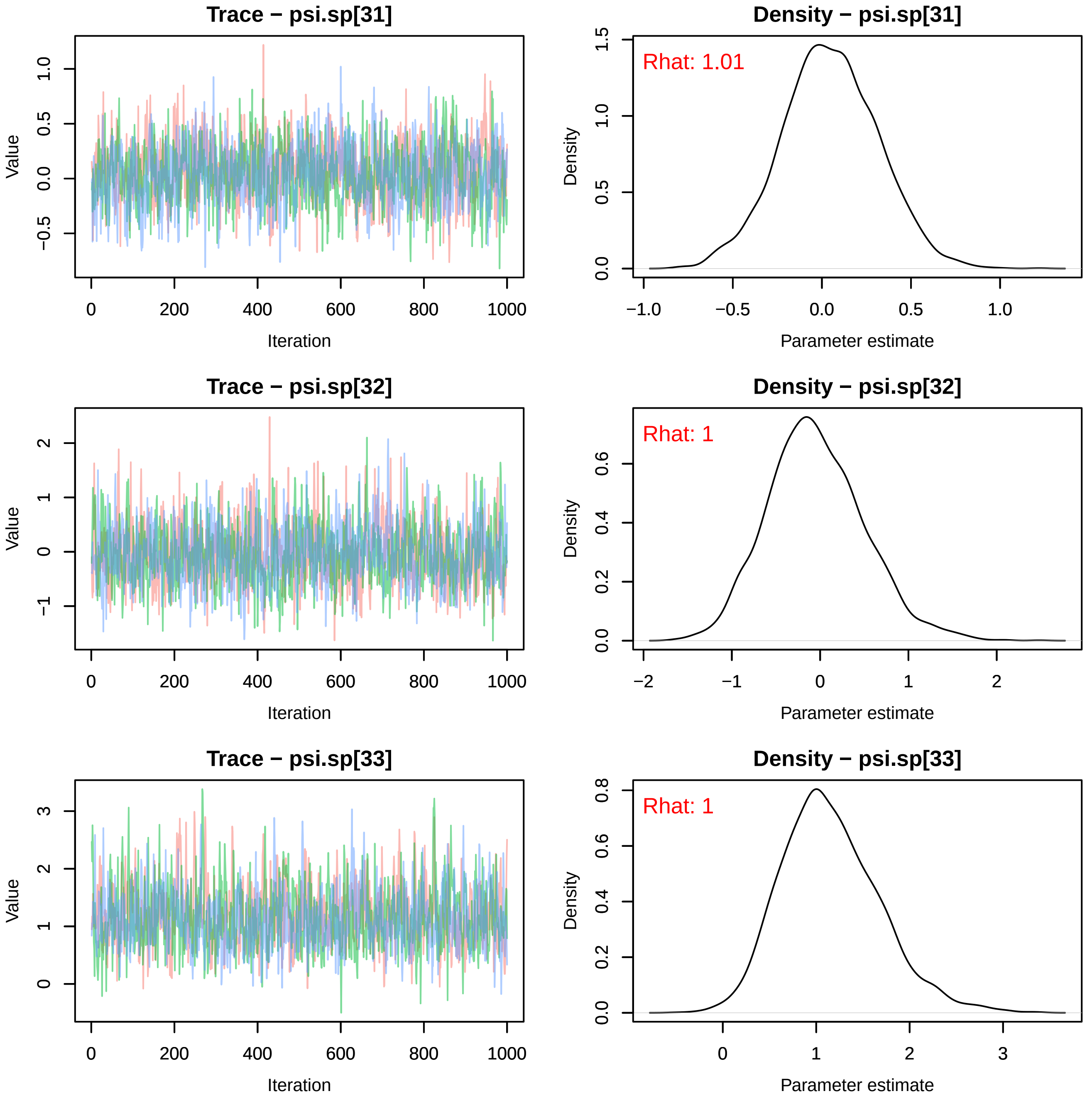

Trace − psi.sp[31]
Value
Iteration
Density − psi.sp[31]
Rhat: 1.01
Density
Parameter estimate
Trace − psi.sp[32]
Value
Iteration
Density − psi.sp[32]
Rhat: 1
Density
Parameter estimate
Trace − psi.sp[33]
Value
Iteration
Density − psi.sp[33]
Rhat: 1
Density
Parameter estimate

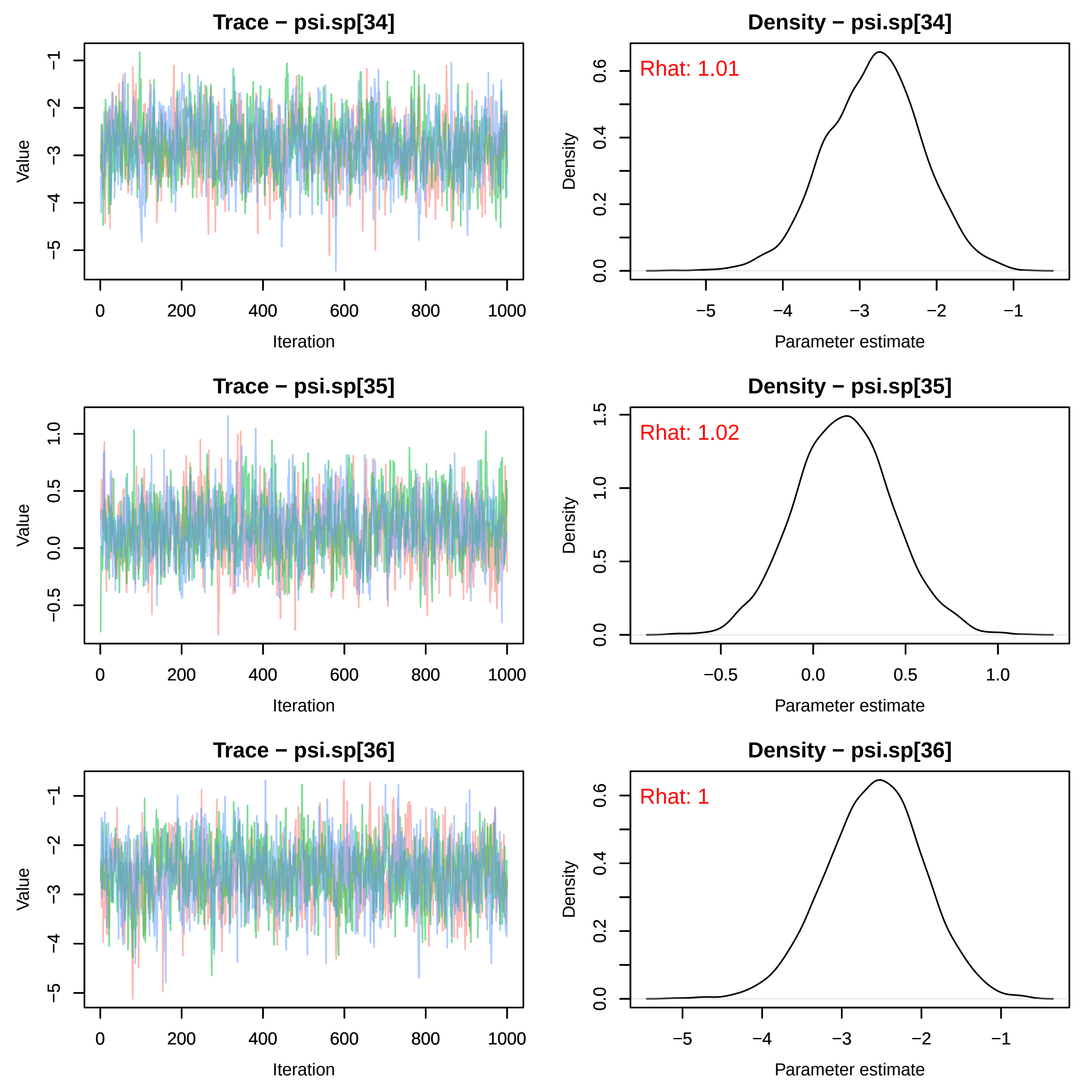

Trace − psi.sp[34]
Value
Iteration
Density − psi.sp[34]
Rhat: 1.01
Density
Parameter estimate
Trace − psi.sp[35]
Value
Iteration
Density − psi.sp[35]
Rhat: 1.02
Density
Parameter estimate
Trace − psi.sp[36]
Value
Iteration
Density − psi.sp[36]
Rhat: 1
Density
Parameter estimate

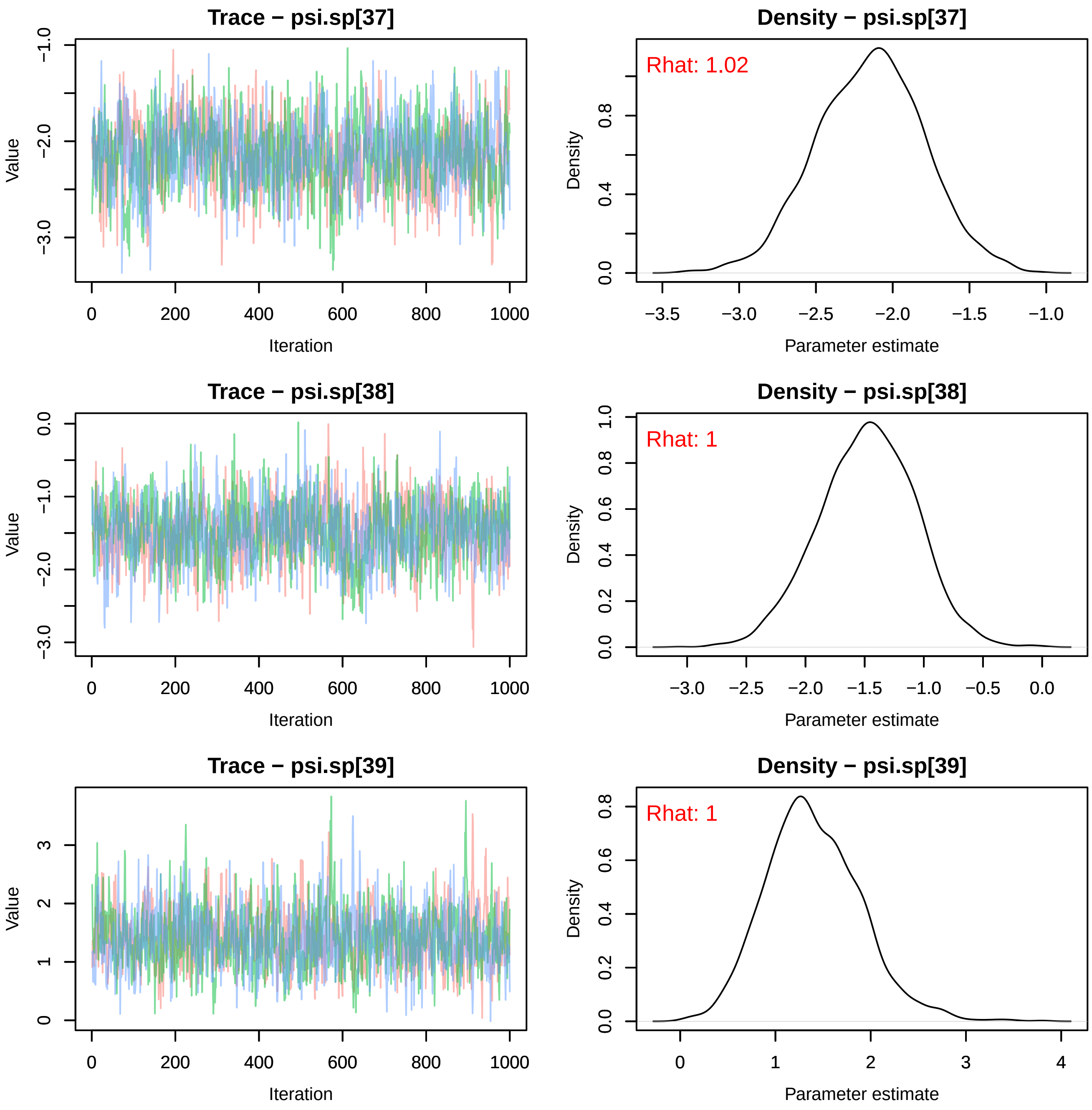
Trace − psi.sp[37]
Density − psi.sp[37]
Rhat: 1.02
Trace − psi.sp[38]
Density − psi.sp[38]
Rhat: 1
Trace − psi.sp[39]
Density − psi.sp[39]
Rhat: 1
Value
Iteration
Density
Parameter estimate
0
200
400
600
800
1000
−1.0
−2.0
−3.0
0.0
0.4
0.8
−3.5
−3.0
−2.5
−2.0
−1.5
−1.0
0.0
−1.0
−2.0
−3.0
0.0
0.2
0.4
0.6
0.8
1.0
−3.0
−2.5
−2.0
−1.5
−1.0
−0.5
0.0
0
1
2
3
0.0
0.2
0.4
0.6
0.8
0
1
2
3
4

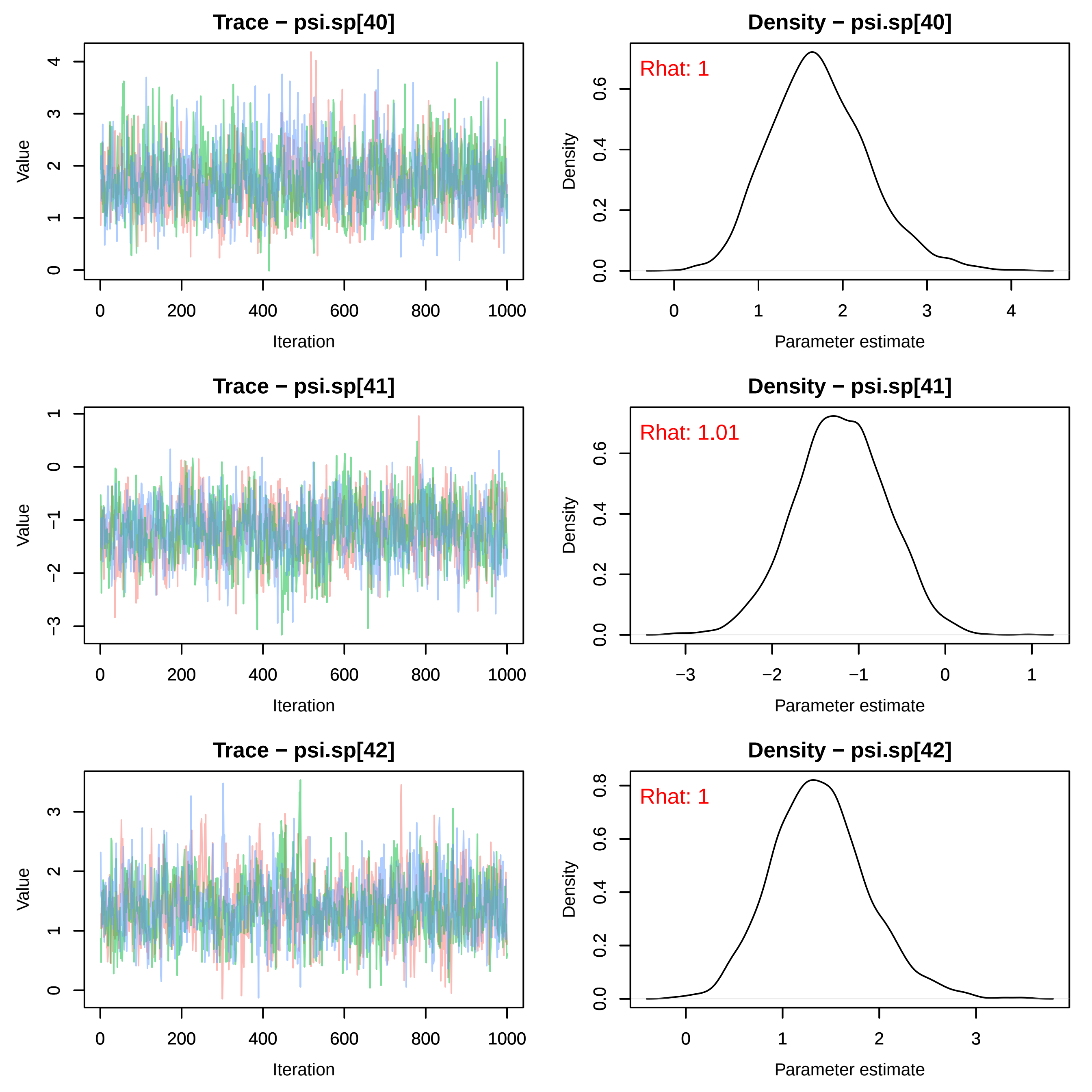
Trace − psi.sp[40]
Value
Iteration
Density − psi.sp[40]
Rhat: 1
Density
Parameter estimate
Trace − psi.sp[41]
Value
Iteration
Density − psi.sp[41]
Rhat: 1.01
Density
Parameter estimate
Trace − psi.sp[42]
Value
Iteration
Density − psi.sp[42]
Rhat: 1
Density
Parameter estimate

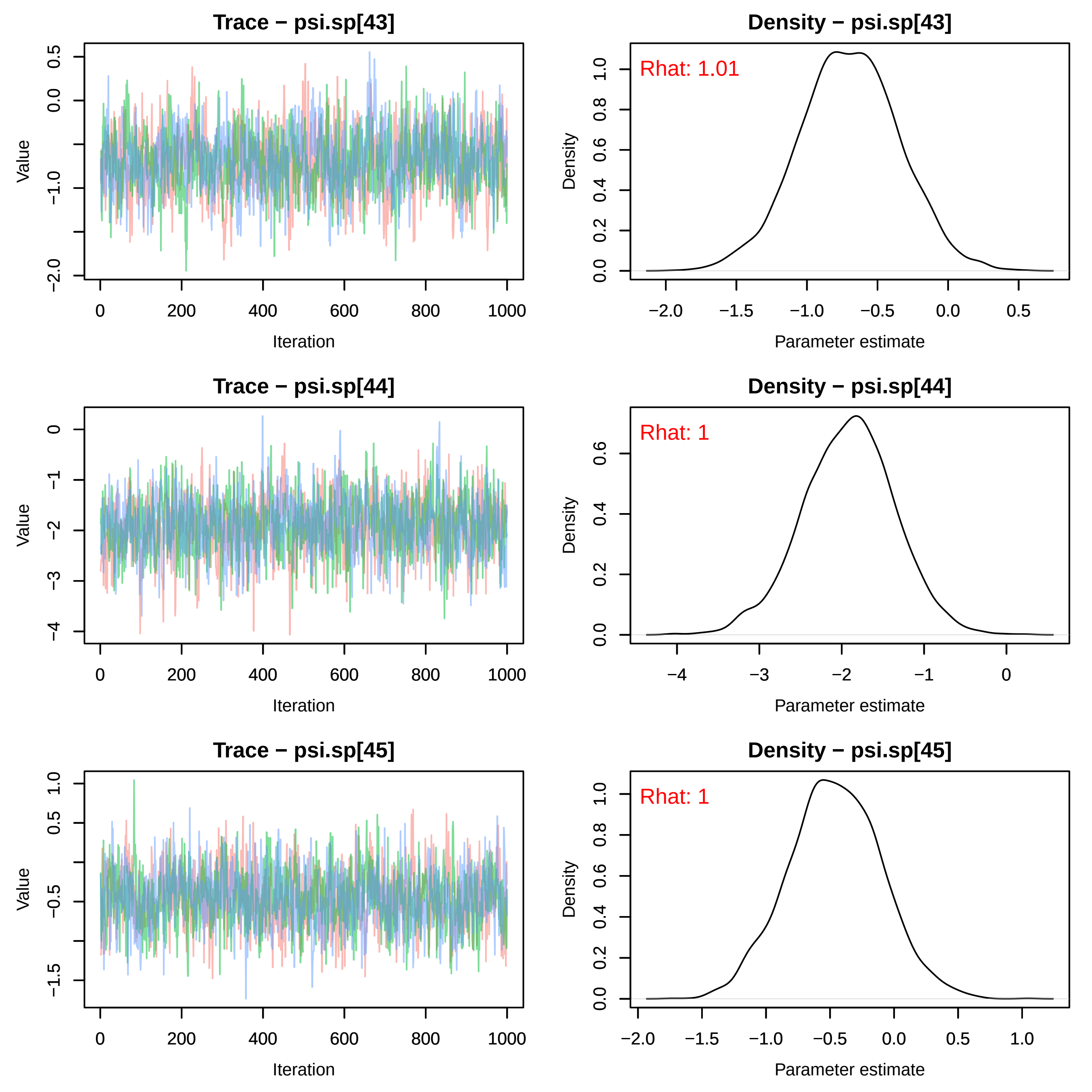
Trace − psi.sp[43]
Density − psi.sp[43]
Rhat: 1.01
Trace − psi.sp[44]
Density − psi.sp[44]
Rhat: 1
Trace − psi.sp[45]
Density − psi.sp[45]
Rhat: 1
Value
Iteration
Density
Parameter estimate

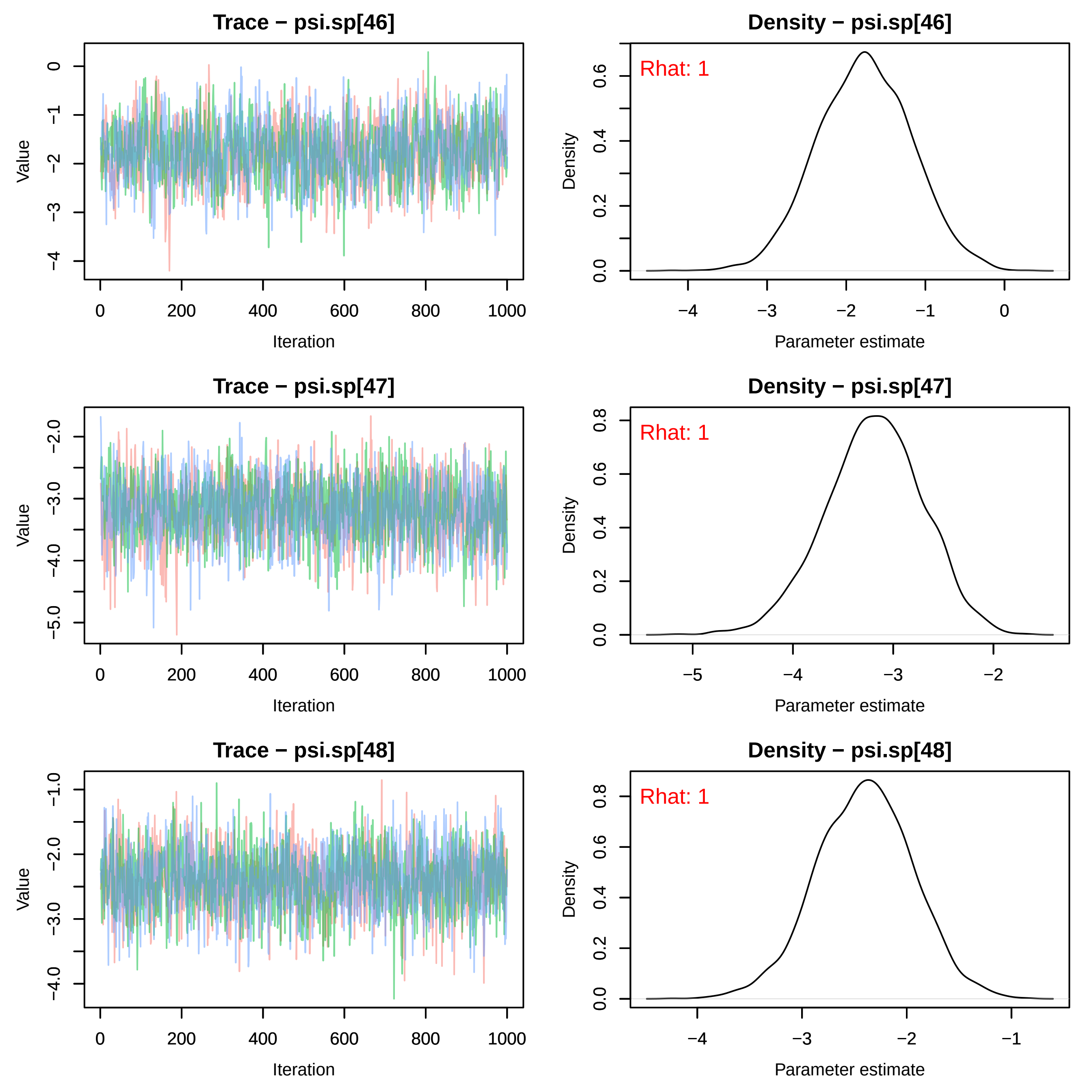
Trace − psi.sp[46]
Value
Iteration
0
200
400
600
800
1000
0
−1
−2
−3
−4
Density − psi.sp[46]
Rhat: 1
Density
Parameter estimate
0.0
0.2
0.4
0.6
−4
−3
−2
−1
0
Trace − psi.sp[47]
Value
Iteration
−2.0
−3.0
−4.0
−5.0
0
200
400
600
800
1000
Density − psi.sp[47]
Rhat: 1
Density
Parameter estimate
0.0
0.2
0.4
0.6
0.8
−5
−4
−3
−2
Trace − psi.sp[48]
Value
Iteration
−1.0
−2.0
−3.0
−4.0
0
200
400
600
800
1000
Density − psi.sp[48]
Rhat: 1
Density
Parameter estimate
0.0
0.2
0.4
0.6
0.8
−4
−3
−2
−1

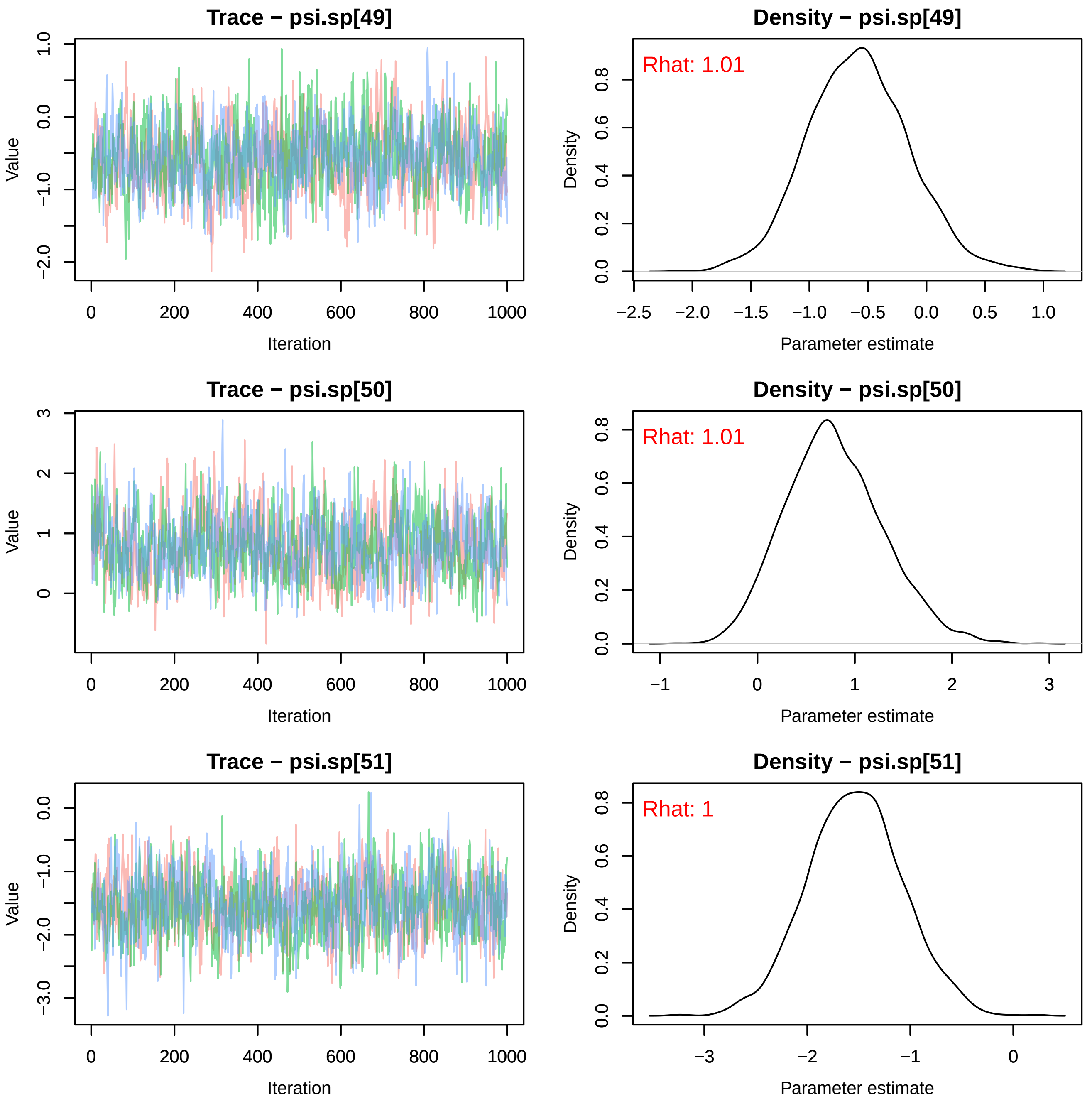

Trace − psi.sp[49]
Value
Iteration
Density − psi.sp[49]
Rhat: 1.01
Density
Parameter estimate
Trace − psi.sp[50]
Value
Iteration
Density − psi.sp[50]
Rhat: 1.01
Density
Parameter estimate
Trace − psi.sp[51]
Value
Iteration
Density − psi.sp[51]
Rhat: 1
Density
Parameter estimate

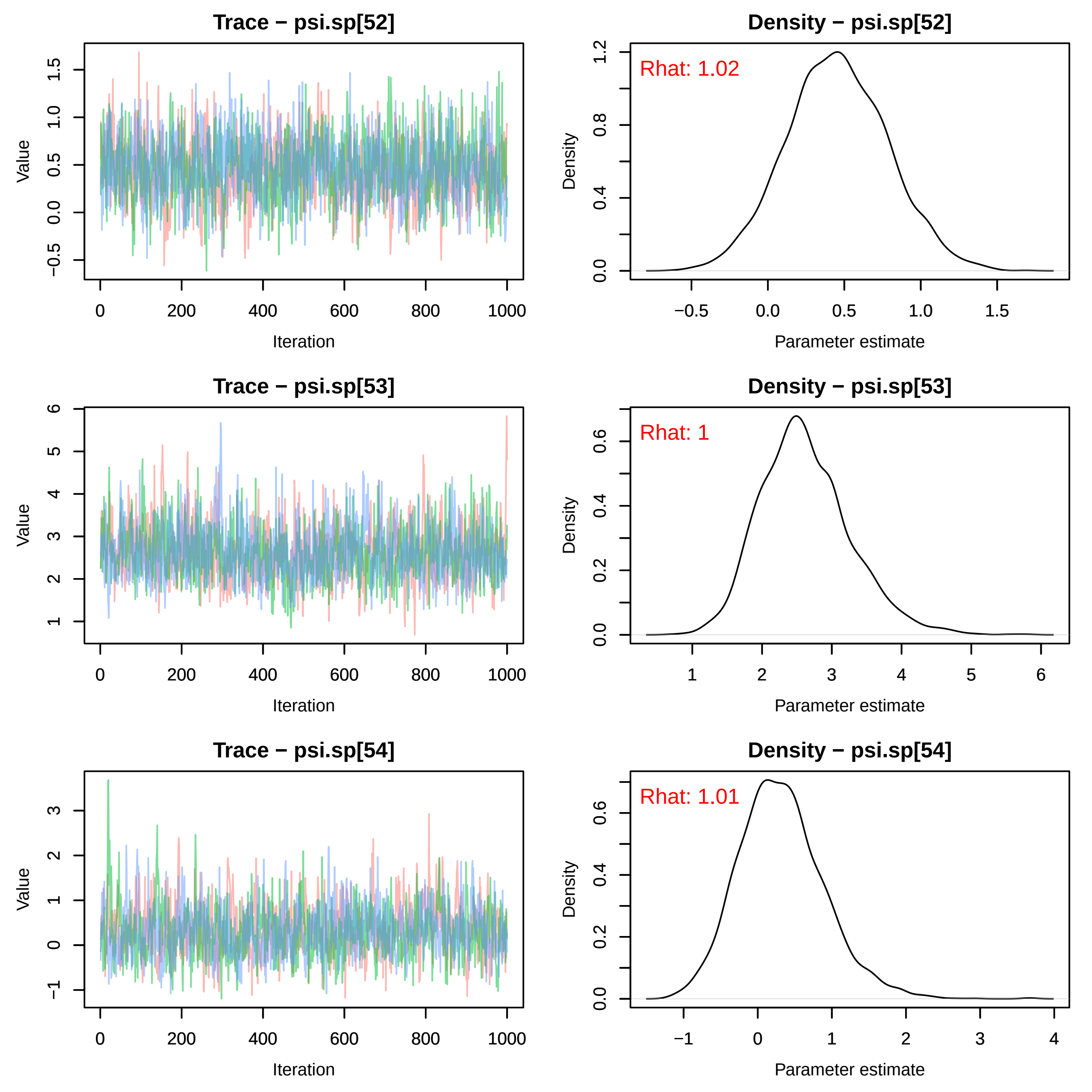

Trace − psi.sp[52]
Value
Iteration
Density − psi.sp[52]
Rhat: 1.02
Density
Parameter estimate
Trace − psi.sp[53]
Value
Iteration
Density − psi.sp[53]
Rhat: 1
Density
Parameter estimate
Trace − psi.sp[54]
Value
Iteration
Density − psi.sp[54]
Rhat: 1.01
Density
Parameter estimate

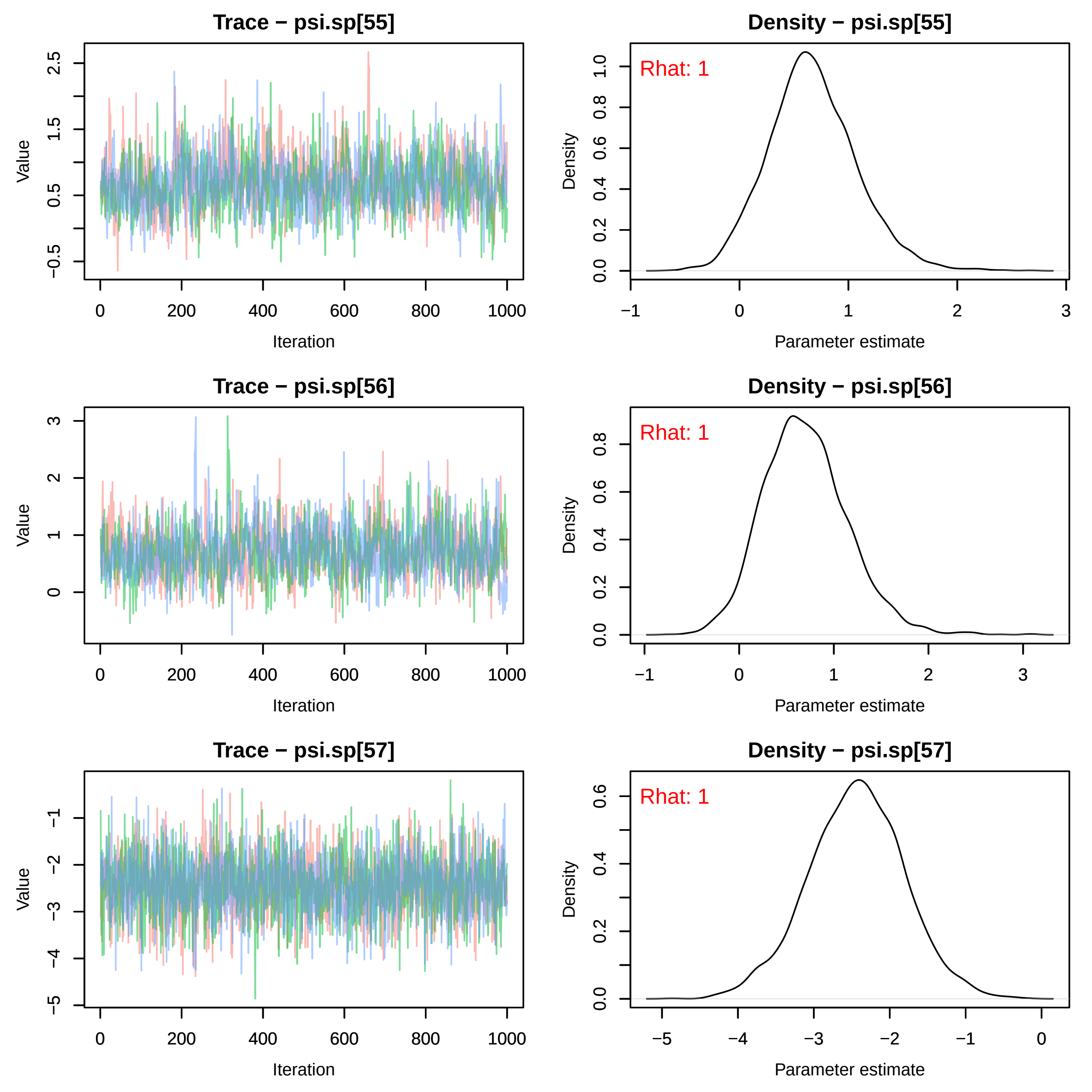

Trace − psi.sp[55]
Value
Iteration
Density − psi.sp[55]
Rhat: 1
Density
Parameter estimate
Trace − psi.sp[56]
Value
Iteration
Density − psi.sp[56]
Rhat: 1
Density
Parameter estimate
Trace − psi.sp[57]
Value
Iteration
Density − psi.sp[57]
Rhat: 1
Density
Parameter estimate

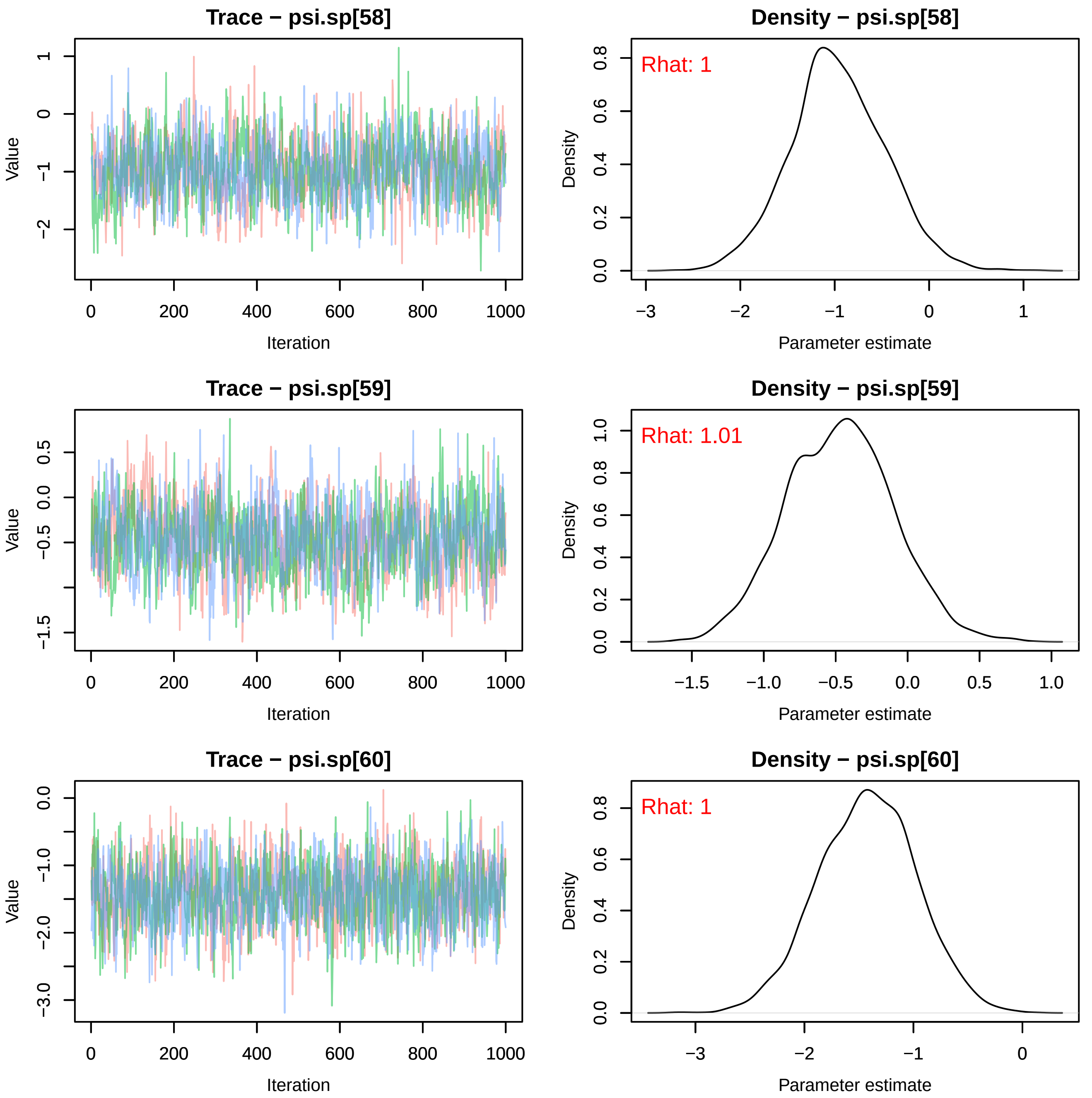
Trace − psi.sp[58]
Value
Iteration
Density − psi.sp[58]
Rhat: 1
Density
Parameter estimate
Trace − psi.sp[59]
Value
Iteration
Density − psi.sp[59]
Rhat: 1.01
Density
Parameter estimate
Trace − psi.sp[60]
Value
Iteration
Density − psi.sp[60]
Rhat: 1
Density
Parameter estimate

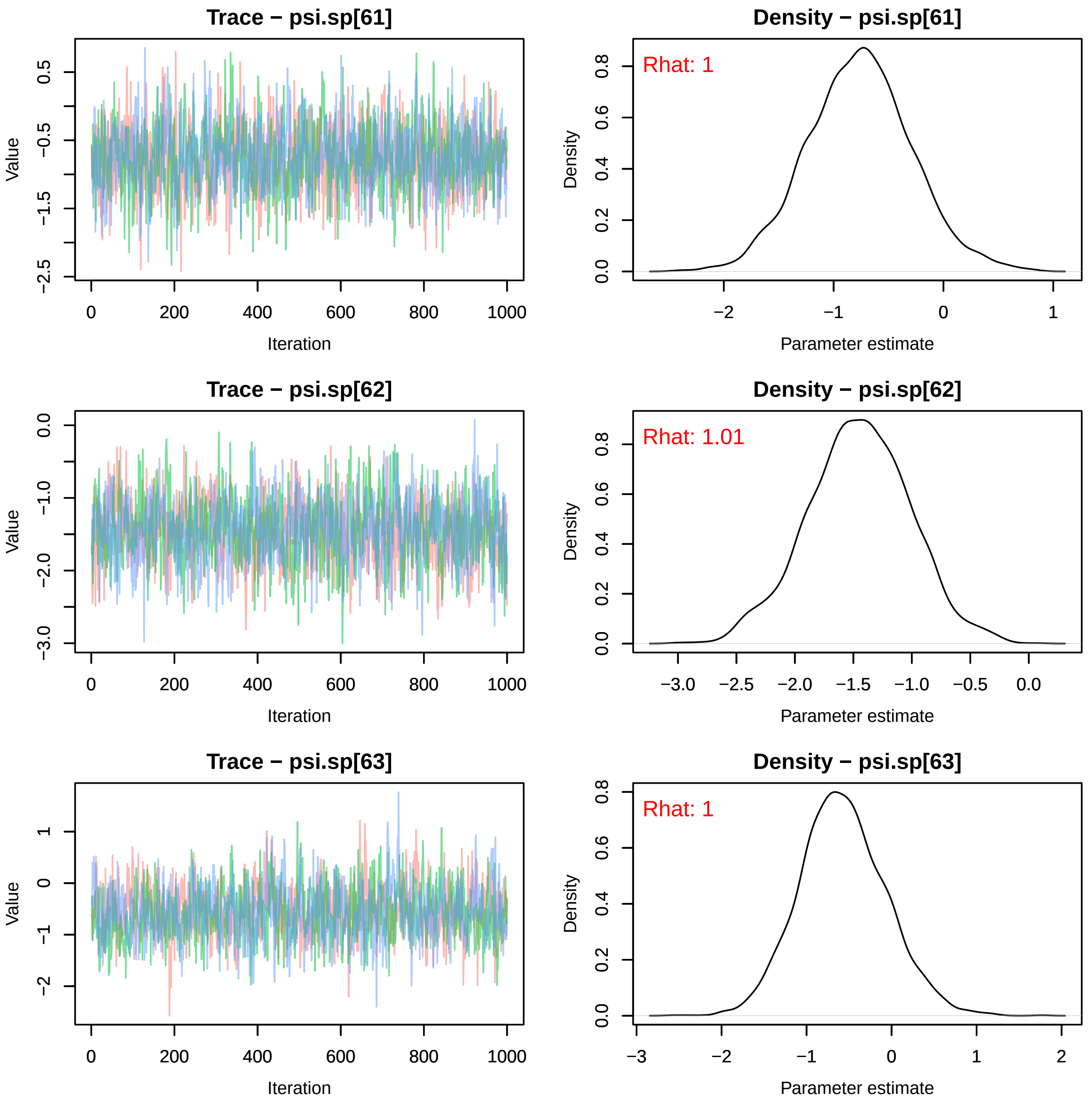
Trace − psi.sp[61]
Value
Iteration
Density − psi.sp[61]
Rhat: 1
Density
Parameter estimate
Trace − psi.sp[62]
Value
Iteration
Density − psi.sp[62]
Rhat: 1.01
Density
Parameter estimate
Trace − psi.sp[63]
Value
Iteration
Density − psi.sp[63]
Rhat: 1
Density
Parameter estimate

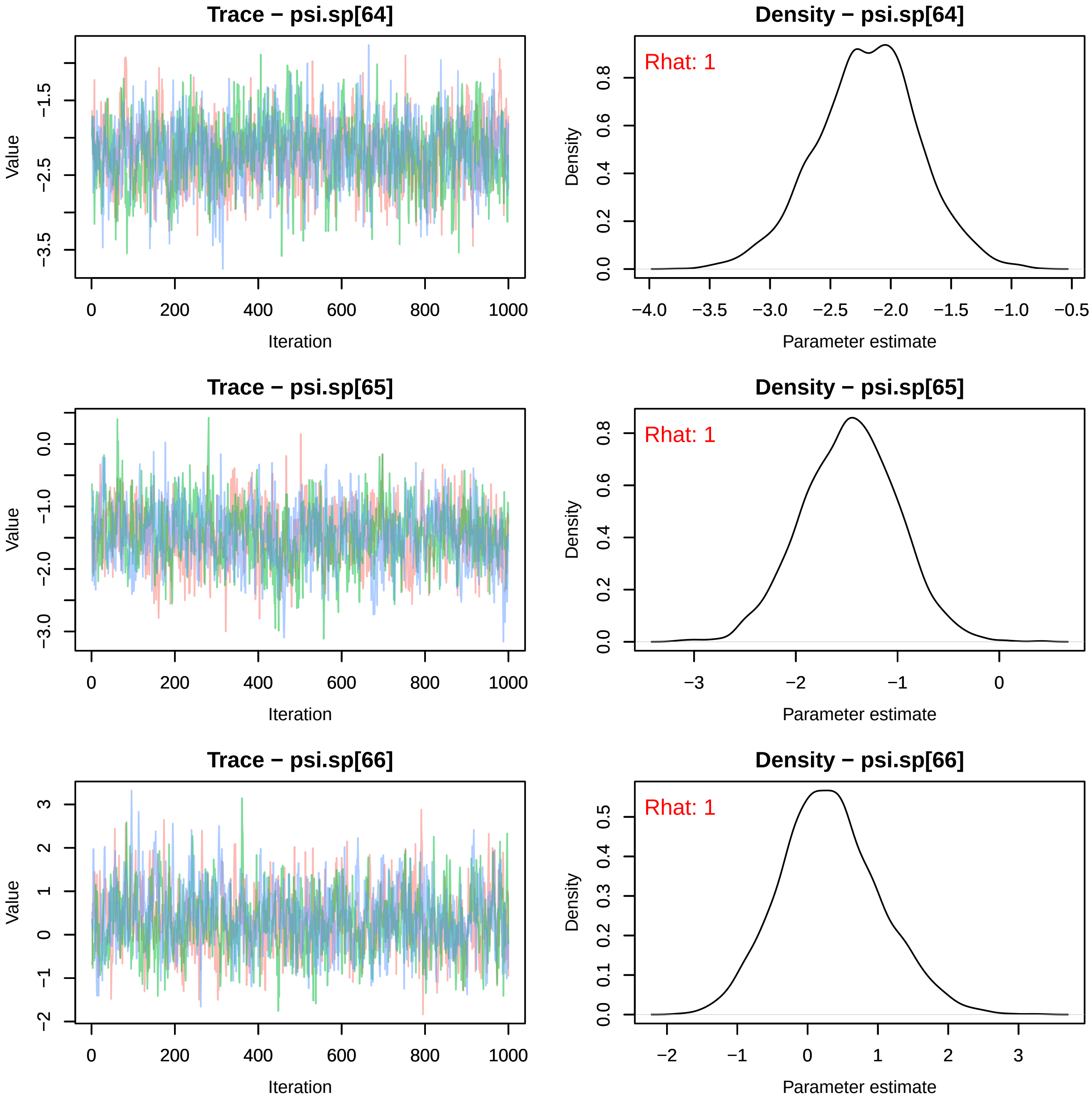
Trace − psi.sp[64]
Density − psi.sp[64]
Rhat: 1
Trace − psi.sp[65]
Density − psi.sp[65]
Rhat: 1
Trace − psi.sp[66]
Density − psi.sp[66]
Rhat: 1
Value
Iteration
Density
Parameter estimate

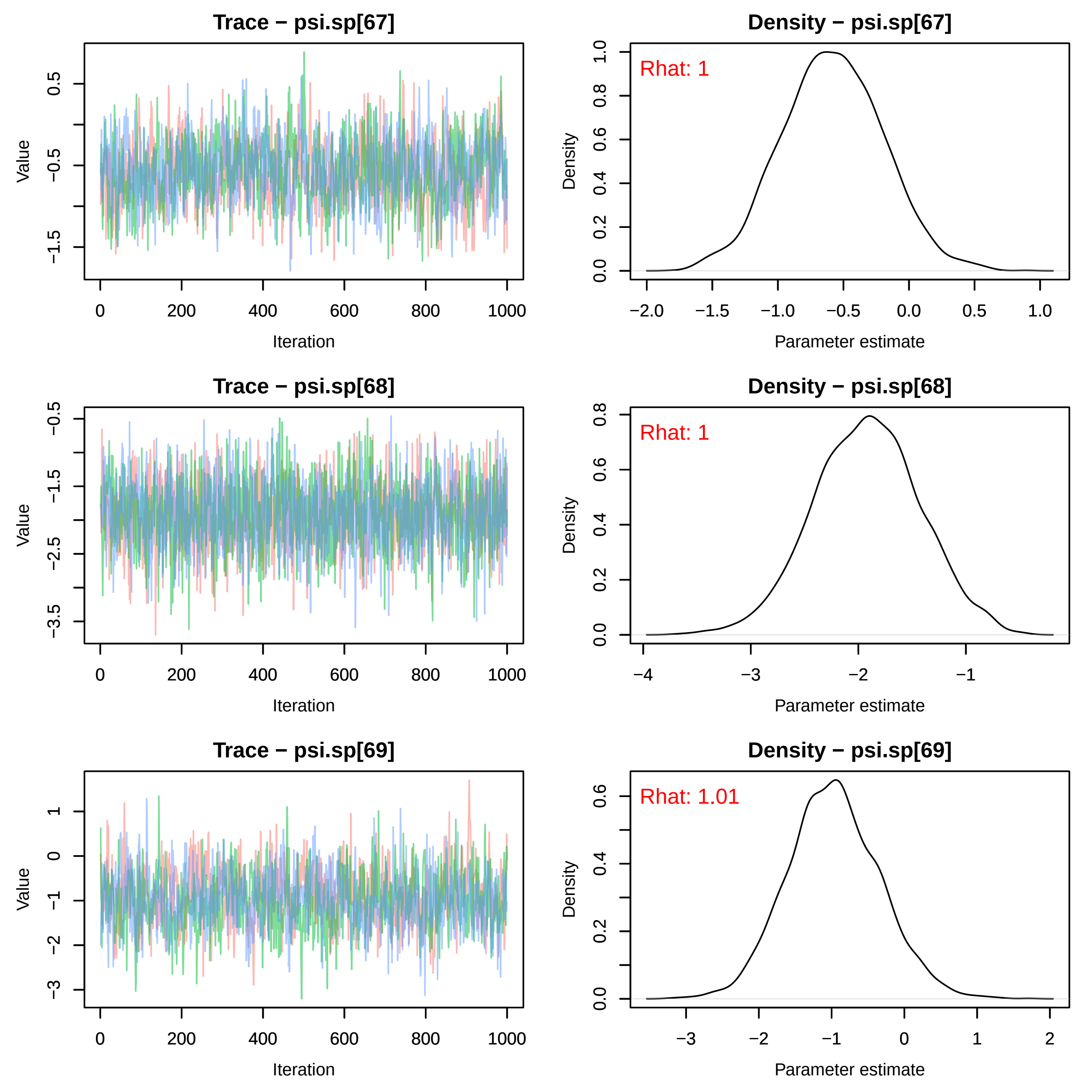
Trace − psi.sp[67]
Value
Iteration
Density − psi.sp[67]
Rhat: 1
Density
Parameter estimate
Trace − psi.sp[68]
Value
Iteration
Density − psi.sp[68]
Rhat: 1
Density
Parameter estimate
Trace − psi.sp[69]
Value
Iteration
Density − psi.sp[69]
Rhat: 1.01
Density
Parameter estimate

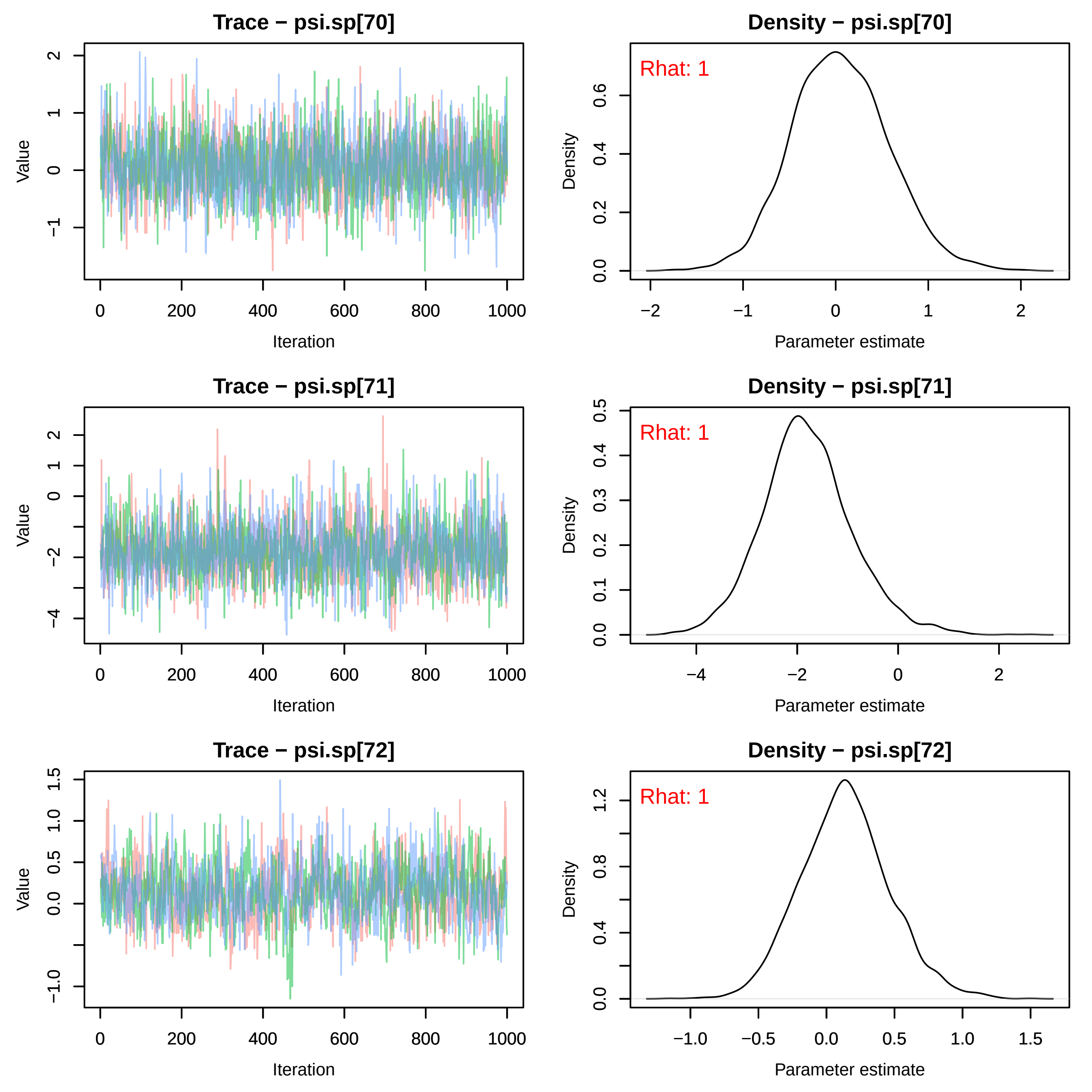

Trace − psi.sp[70]
Value
Iteration
0
200
400
600
800
1000
−1
0
1
2
Density − psi.sp[70]
Rhat: 1
Density
Parameter estimate
0.0
0.2
0.4
0.6
−2
−1
0
1
2
Trace − psi.sp[71]
Value
Iteration
−4
−2
0
1
2
Density − psi.sp[71]
Rhat: 1
Density
Parameter estimate
0.0
0.1
0.2
0.3
0.4
0.5
−4
−2
0
2
Trace − psi.sp[72]
Value
Iteration
−1.0
0.0
0.5
1.0
1.5
Density − psi.sp[72]
Rhat: 1
Density
Parameter estimate
0.0
0.4
0.8
1.2
−1.0
−0.5
0.0
0.5
1.0
1.5

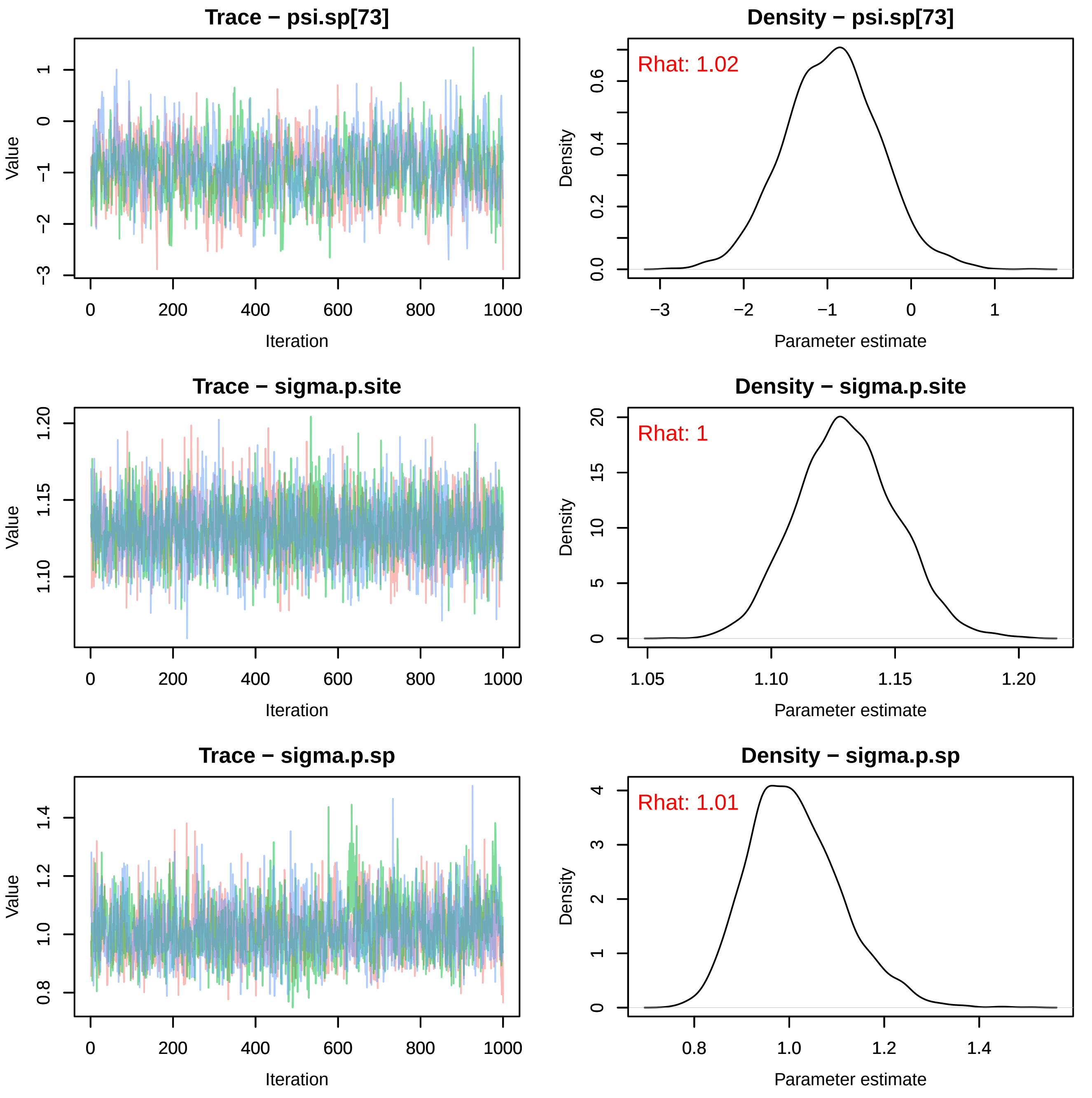
Trace − psi.sp[73]
Density − psi.sp[73]
Rhat: 1.02
Trace − sigma.p.site
Density − sigma.p.site
Rhat: 1
Trace − sigma.p.sp
Density − sigma.p.sp
Rhat: 1.01
Value
Iteration
Density
Parameter estimate

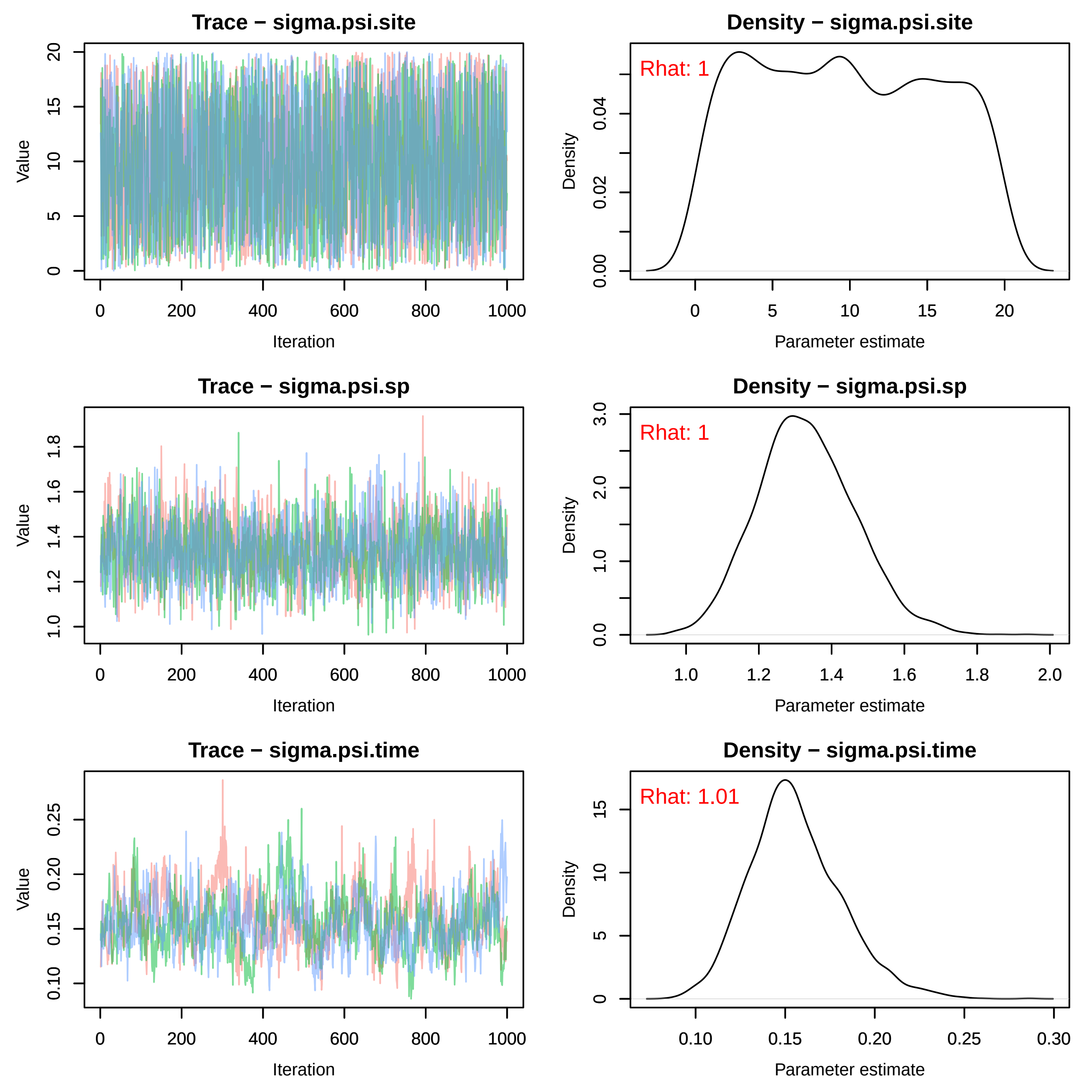
Trace − sigma.psi.site
Value
Iteration
Density − sigma.psi.site
Rhat: 1
Density
Parameter estimate
Trace − sigma.psi.sp
Value
Iteration
Density − sigma.psi.sp
Rhat: 1
Density
Parameter estimate
Trace − sigma.psi.time
Value
Iteration
Density − sigma.psi.time
Rhat: 1.01
Density
Parameter estimate

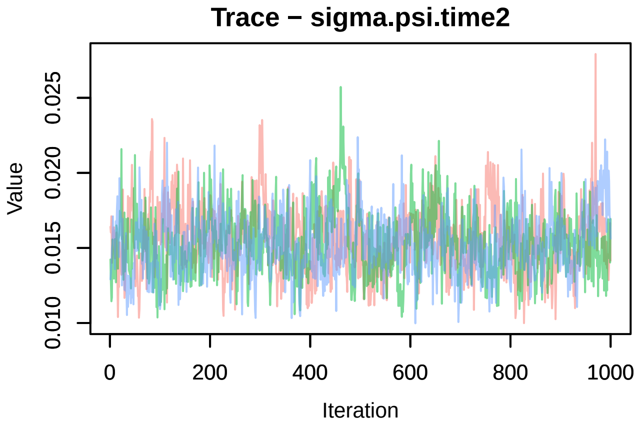
Trace − sigma.psi.time2
Value
0.010
0.015
0.020
0.025
0
200
400
600
800
1000
Iteration

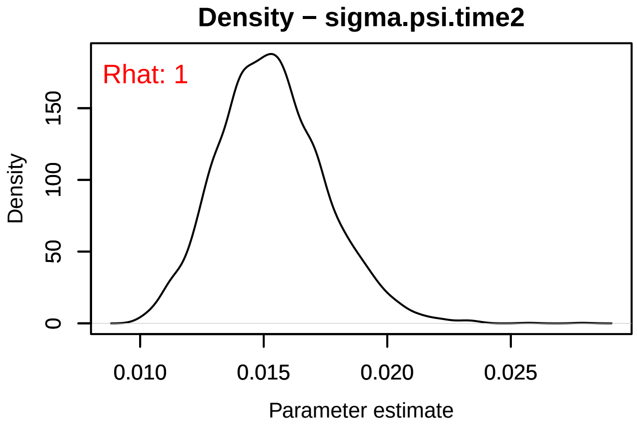
Density − sigma.psi.time2
Rhat: 1
Density
0
50
100
150
0.010
0.015
0.020
0.025
Parameter estimate
