## Supplementary material for "Winners and Losers Among European Arthropods over the Last Half Century of Global Change": araneae_timeTrace.pdf

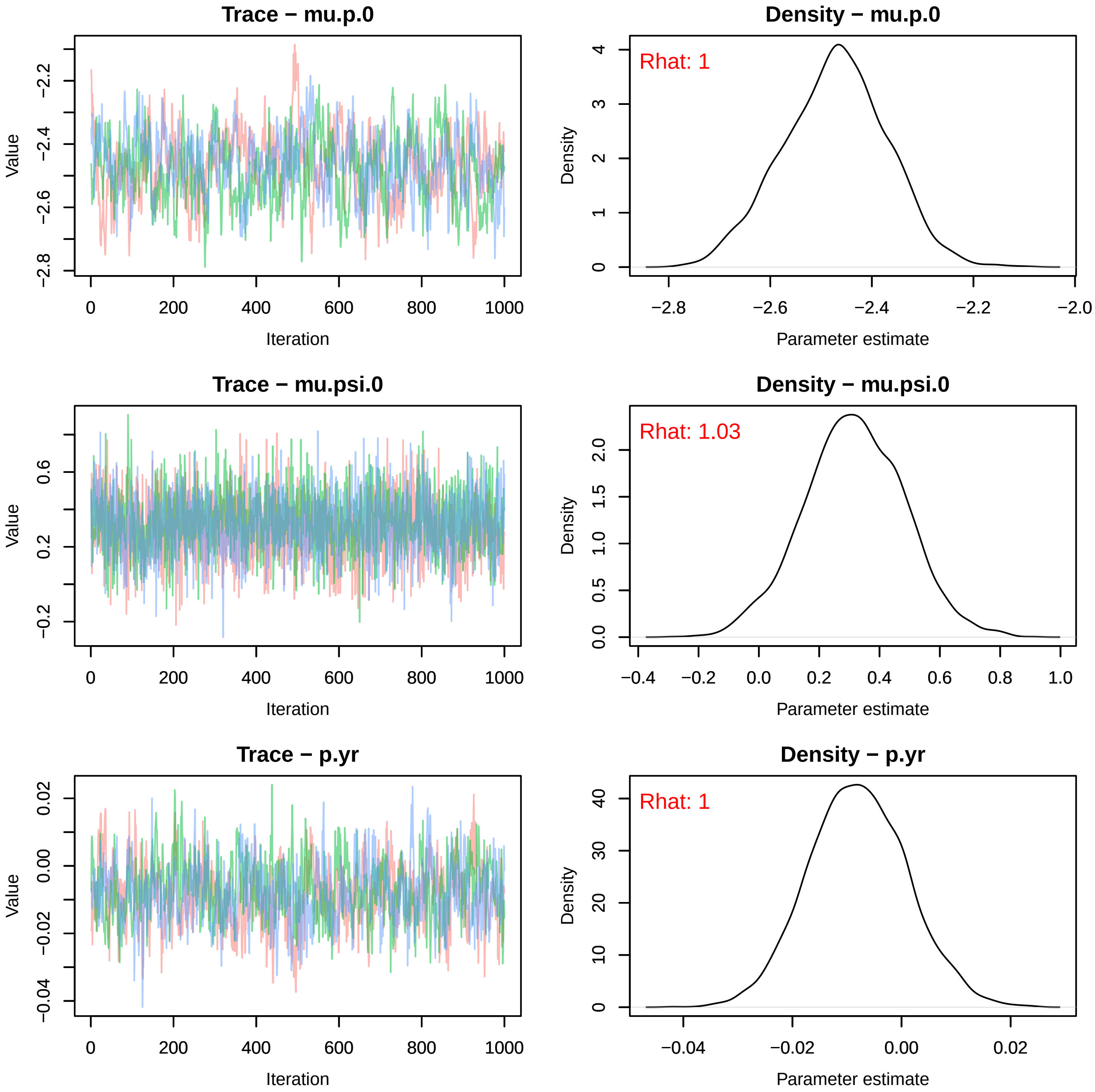

Trace − mu.p.0
Value
−2.8
−2.6
−2.4
−2.2
Iteration
0
200
400
600
800
1000
Density − mu.p.0
Rhat: 1
Density
0
1
2
3
4
Parameter estimate
−2.8
−2.6
−2.4
−2.2
−2.0
Trace − mu.psi.0
Value
−0.2
0.2
0.6
Iteration
0
200
400
600
800
1000
Density − mu.psi.0
Rhat: 1.03
Density
0.0
0.5
1.0
1.5
2.0
Parameter estimate
−0.4
−0.2
0.0
0.2
0.4
0.6
0.8
1.0
Trace − p.yr
Value
−0.04
−0.02
0.00
0.02
Iteration
0
200
400
600
800
1000
Density − p.yr
Rhat: 1
Density
0
10
20
30
40
Parameter estimate
−0.04
−0.02
0.00
0.02

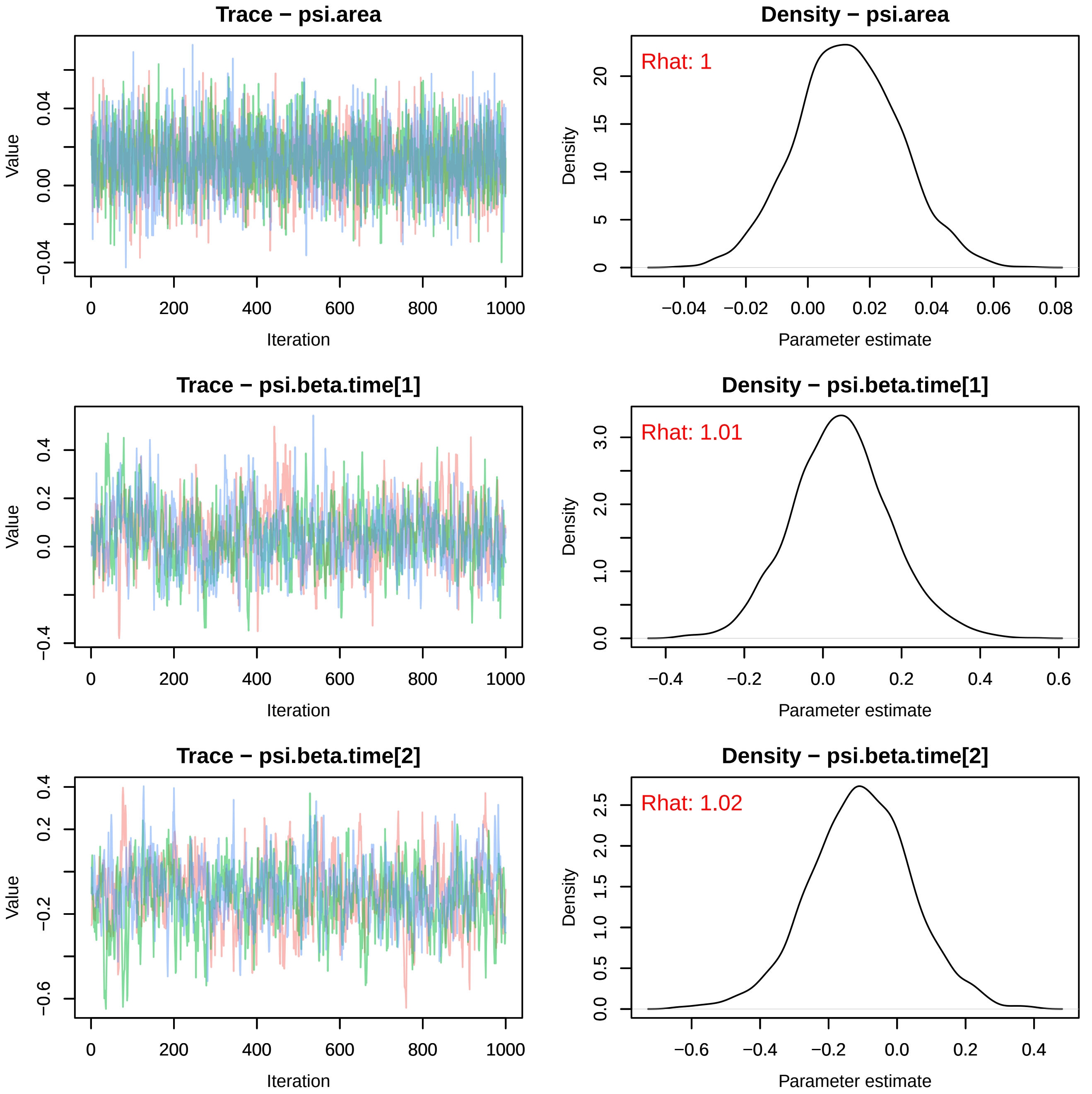

Trace − psi.area
Density − psi.area
Rhat: 1
Trace − psi.beta.time[1]
Density − psi.beta.time[1]
Rhat: 1.01
Trace − psi.beta.time[2]
Density − psi.beta.time[2]
Rhat: 1.02
Value
Iteration
Density
Parameter estimate

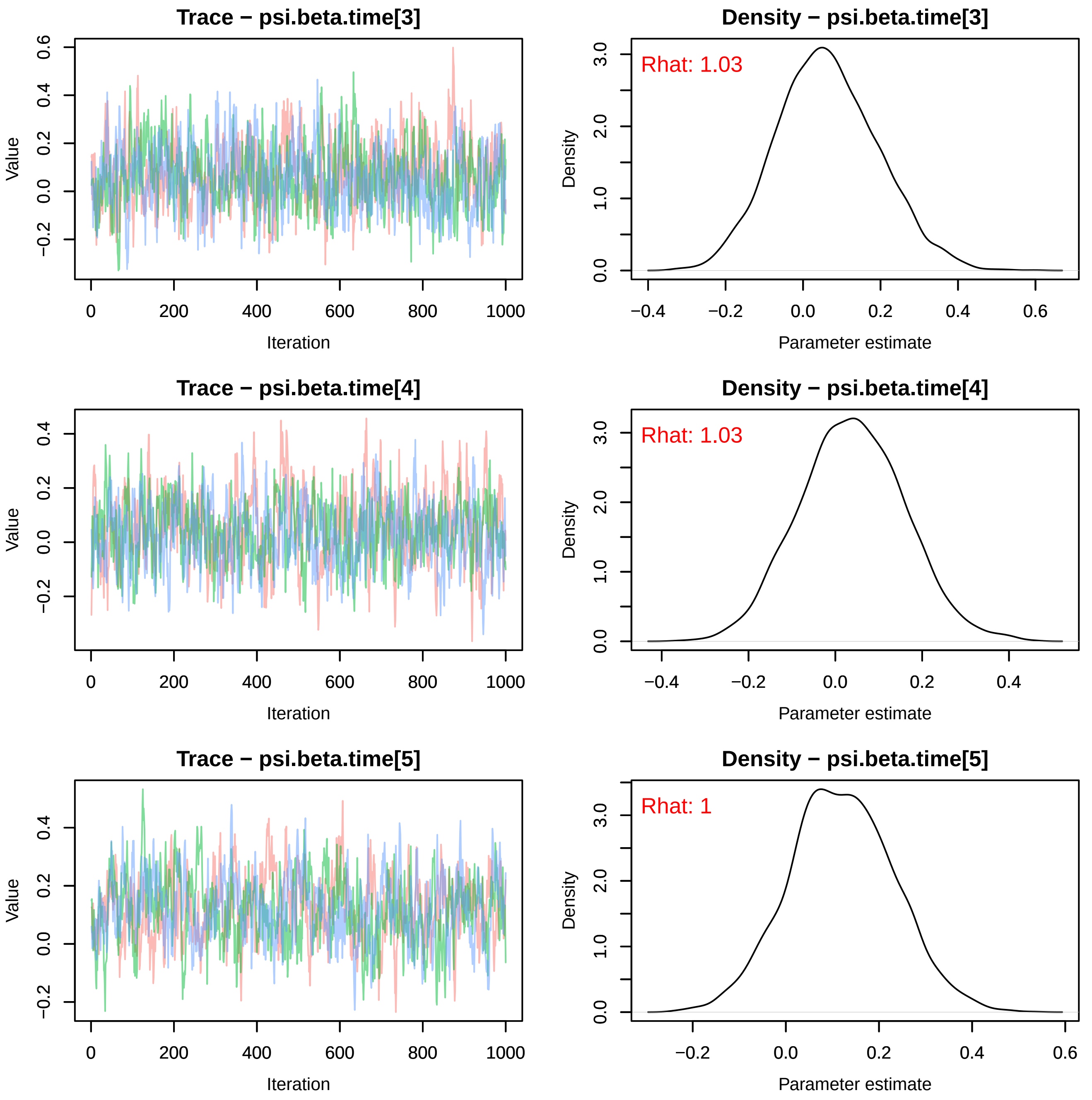
Trace − psi.beta.time[3]
Value
Iteration
Density − psi.beta.time[3]
Rhat: 1.03
Density
Parameter estimate
Trace − psi.beta.time[4]
Value
Iteration
Density − psi.beta.time[4]
Rhat: 1.03
Density
Parameter estimate
Trace − psi.beta.time[5]
Value
Iteration
Density − psi.beta.time[5]
Rhat: 1
Density
Parameter estimate

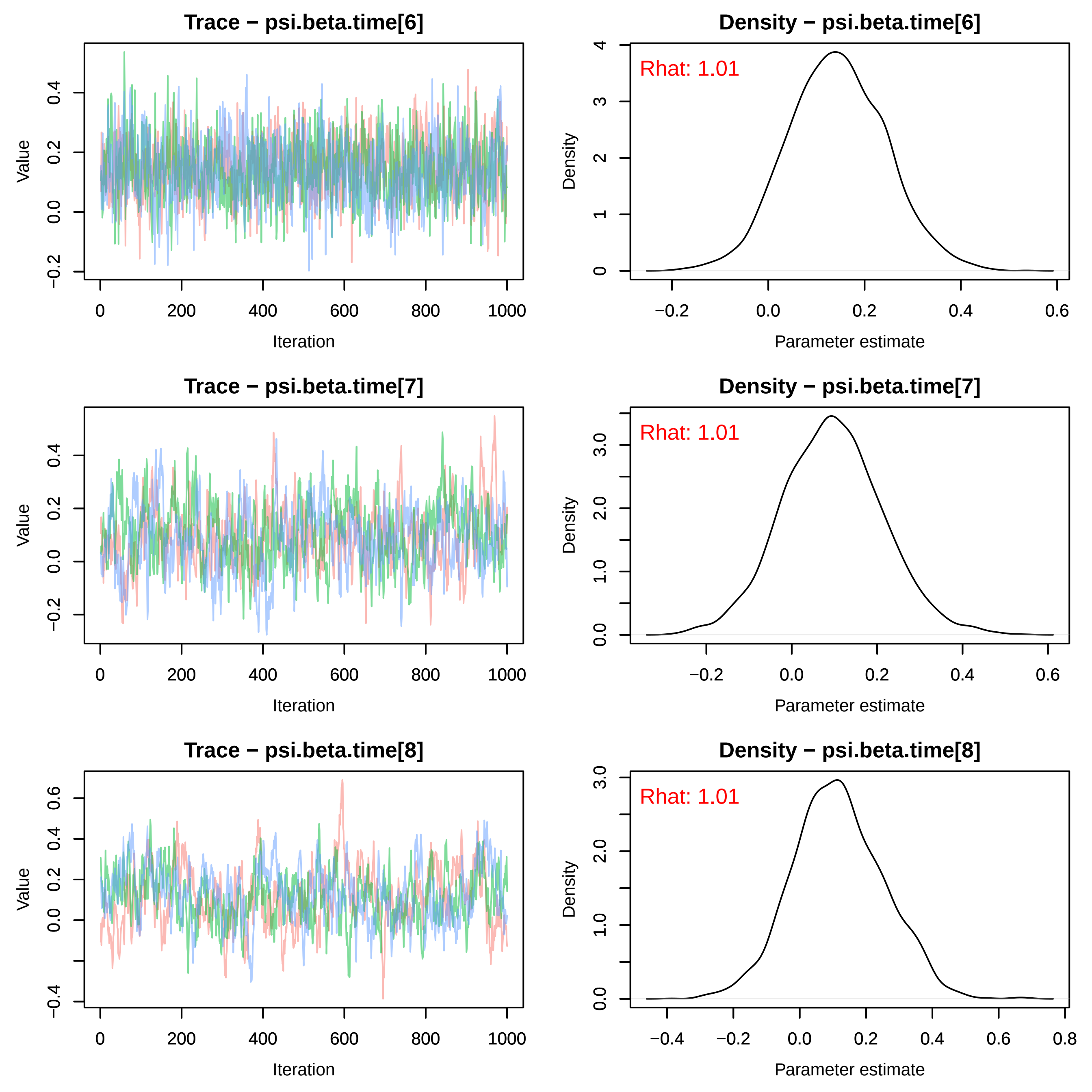
Trace − psi.beta.time[6]
Value
Iteration
Density − psi.beta.time[6]
Rhat: 1.01
Density
Parameter estimate
Trace − psi.beta.time[7]
Value
Iteration
Density − psi.beta.time[7]
Rhat: 1.01
Density
Parameter estimate
Trace − psi.beta.time[8]
Value
Iteration
Density − psi.beta.time[8]
Rhat: 1.01
Density
Parameter estimate

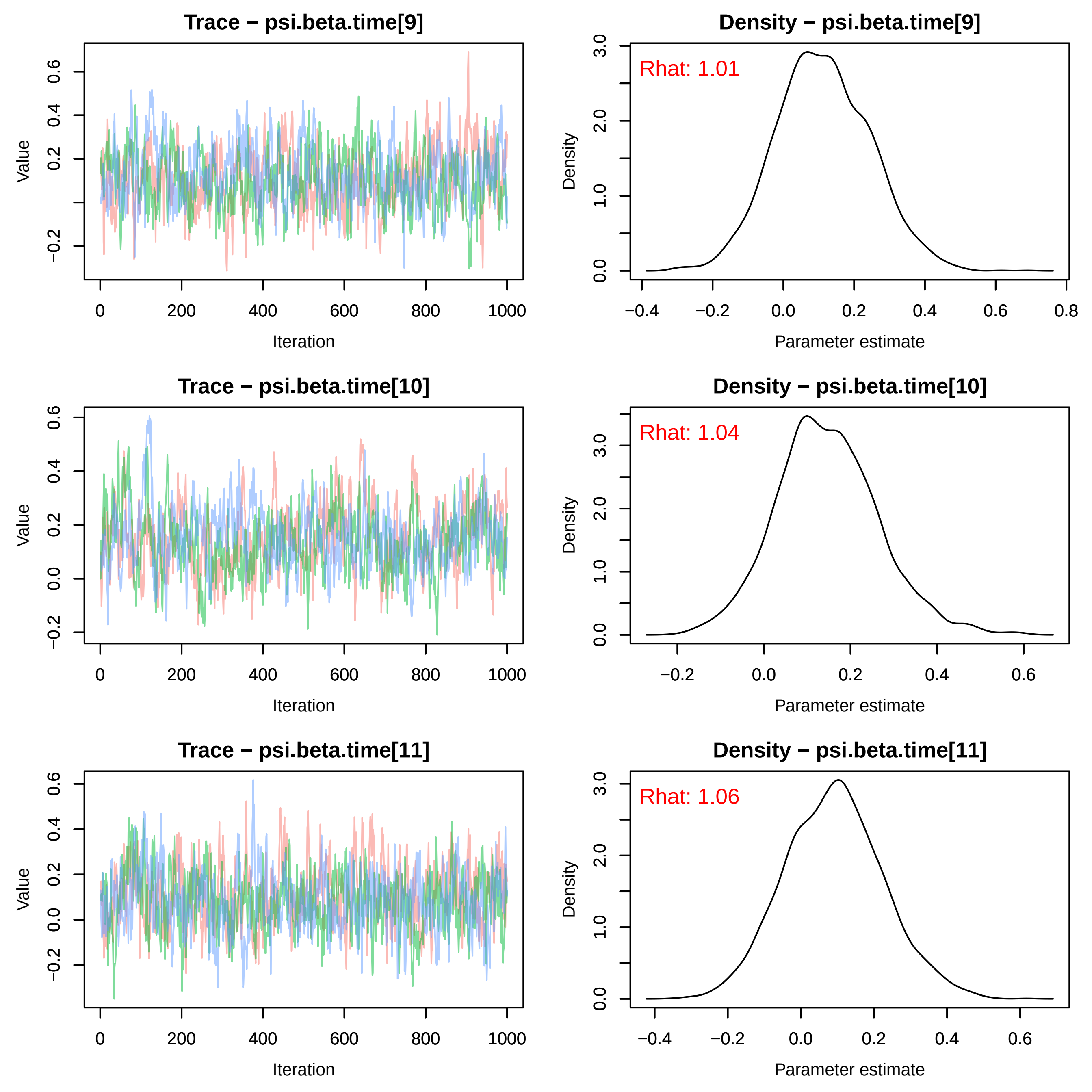
Trace − psi.beta.time[9]
Value
Iteration
Density − psi.beta.time[9]
Rhat: 1.01
Density
Parameter estimate
Trace − psi.beta.time[10]
Value
Iteration
Density − psi.beta.time[10]
Rhat: 1.04
Density
Parameter estimate
Trace − psi.beta.time[11]
Value
Iteration
Density − psi.beta.time[11]
Rhat: 1.06
Density
Parameter estimate

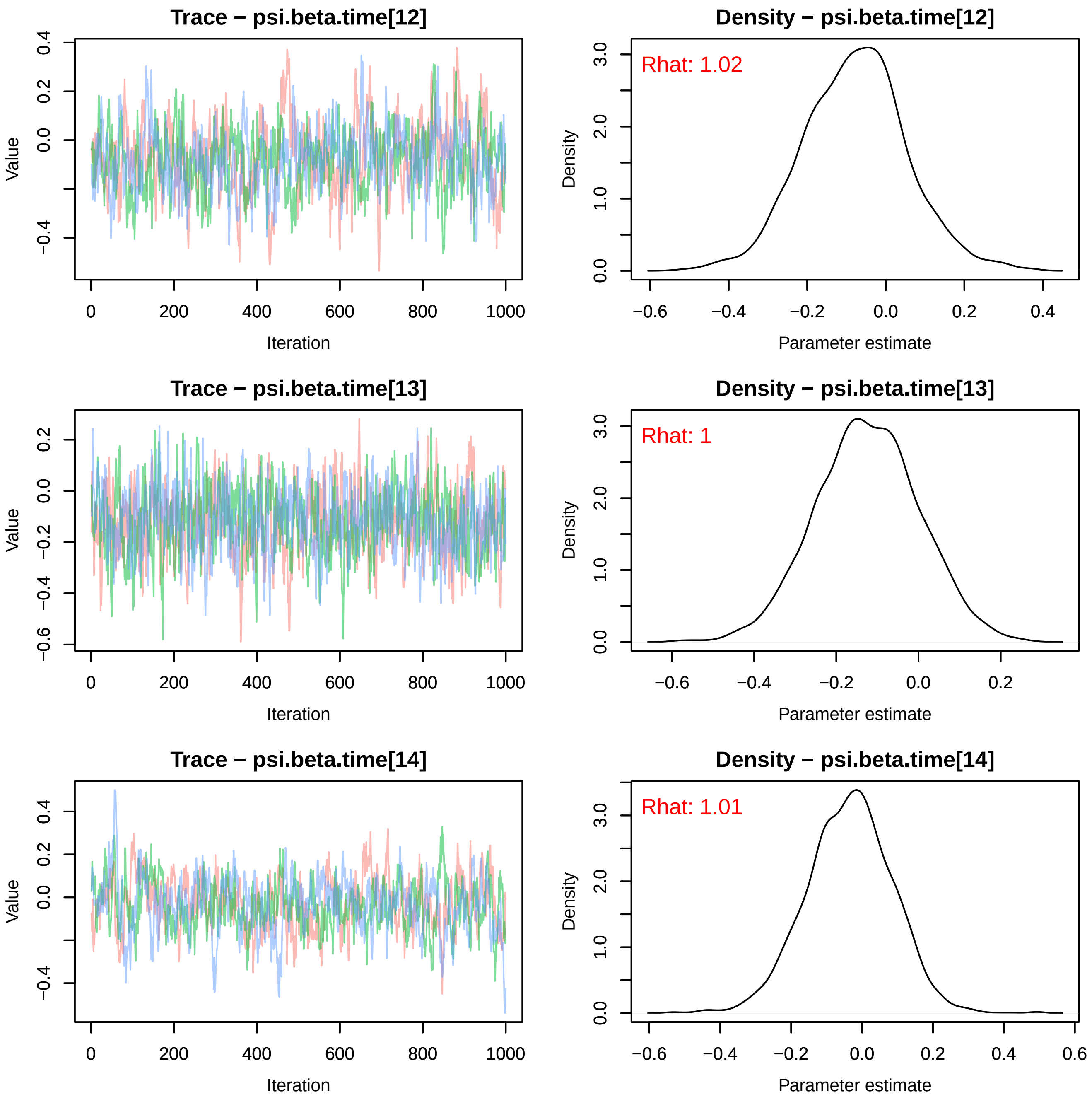

Trace − psi.beta.time[12]
Value
Iteration
0
200
400
600
800
1000
−0.4
0.0
0.2
0.4
Density − psi.beta.time[12]
Rhat: 1.02
Density
Parameter estimate
−0.6
−0.4
−0.2
0.0
0.2
0.4
0.0
1.0
2.0
3.0
Trace − psi.beta.time[13]
Value
Iteration
−0.6
−0.4
−0.2
0.0
0.2
Density − psi.beta.time[13]
Rhat: 1
Density
Parameter estimate
Trace − psi.beta.time[14]
Value
Iteration
Density − psi.beta.time[14]
Rhat: 1.01
Density
Parameter estimate
0.6

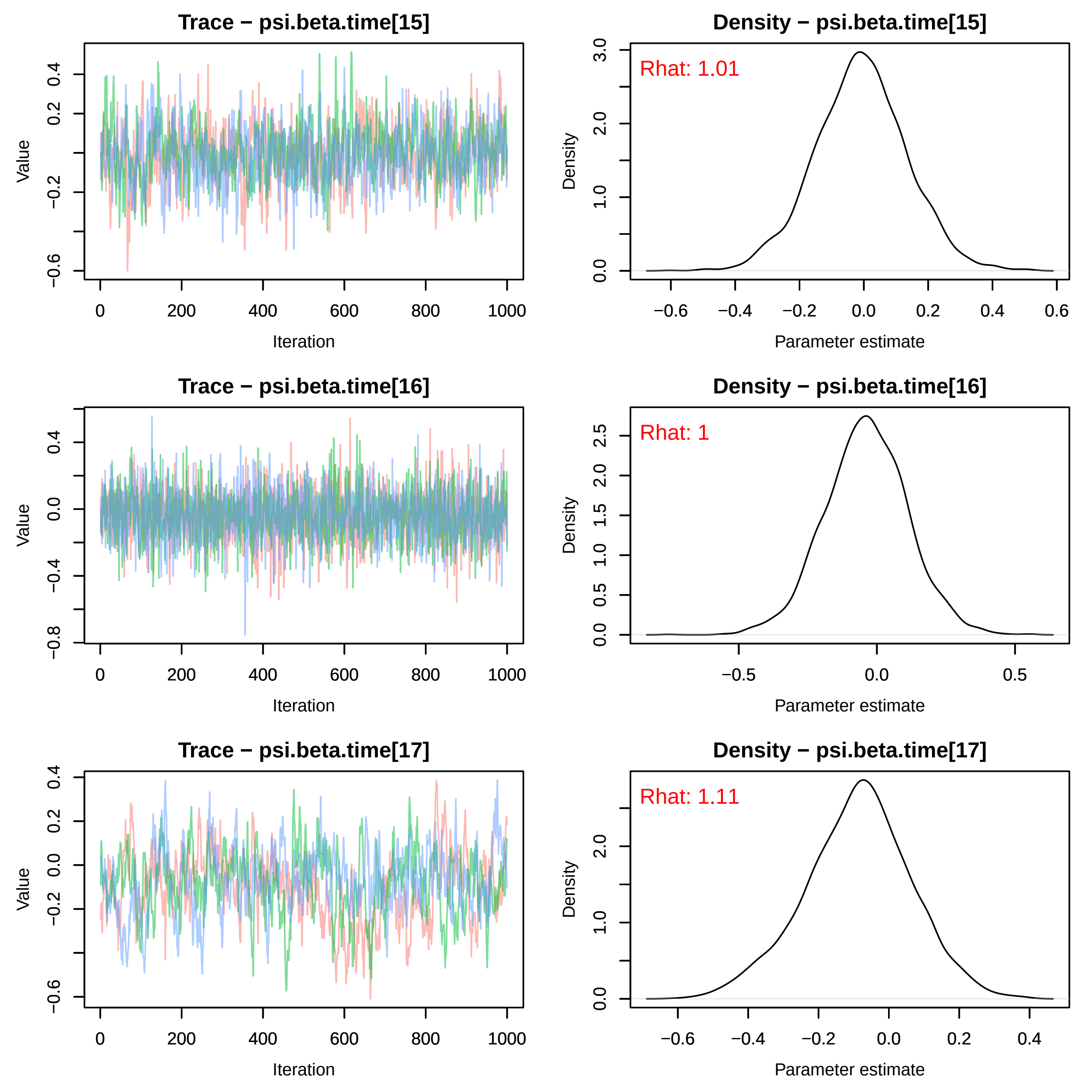

Trace − psi.beta.time[15]
Value
Iteration
Density − psi.beta.time[15]
Rhat: 1.01
Density
Parameter estimate
Trace − psi.beta.time[16]
Value
Iteration
Density − psi.beta.time[16]
Rhat: 1
Density
Parameter estimate
Trace − psi.beta.time[17]
Value
Iteration
Density − psi.beta.time[17]
Rhat: 1.11
Density
Parameter estimate

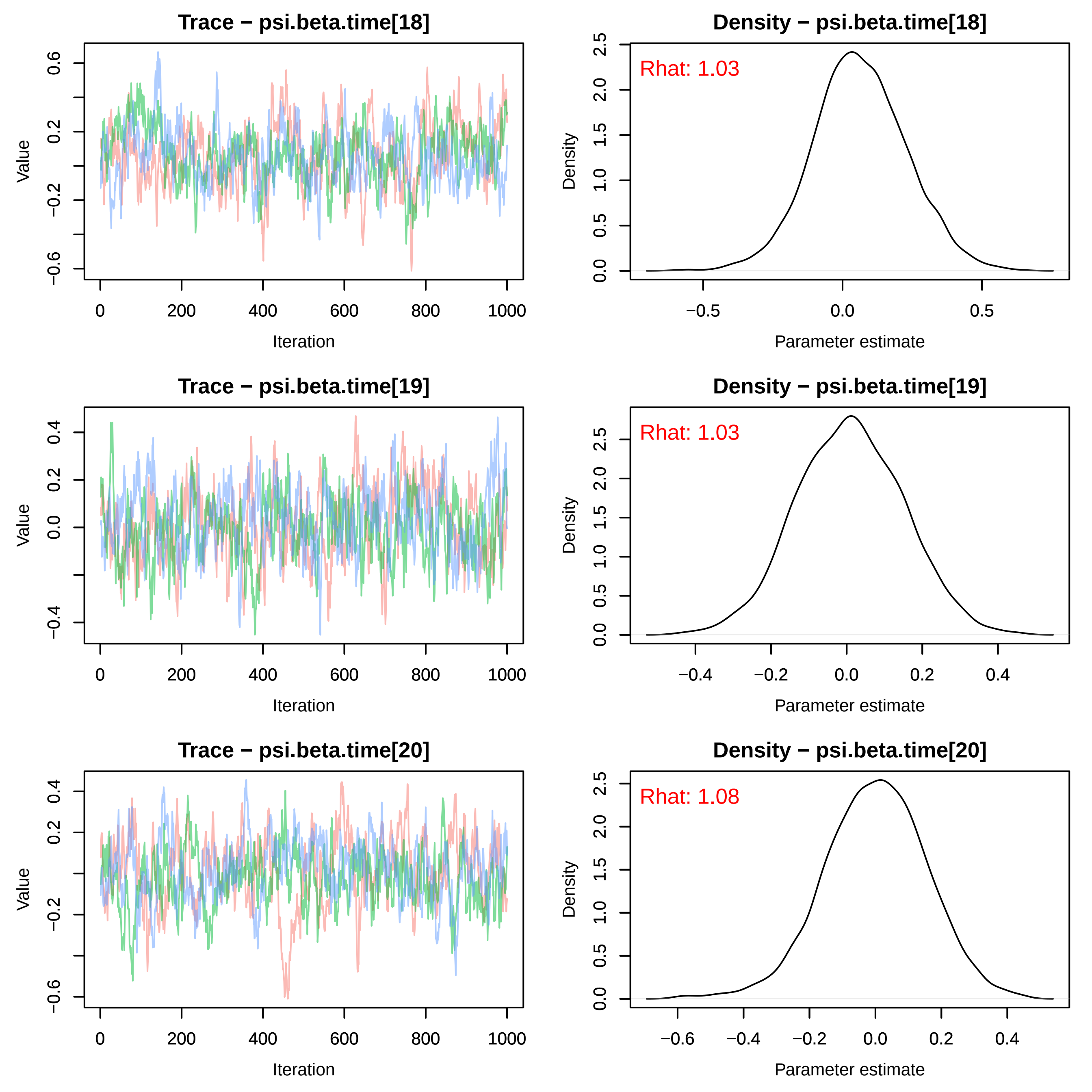

Trace − psi.beta.time[18]
Value
Iteration
Density − psi.beta.time[18]
Rhat: 1.03
Density
Parameter estimate
Trace − psi.beta.time[19]
Value
Iteration
Density − psi.beta.time[19]
Rhat: 1.03
Density
Parameter estimate
Trace − psi.beta.time[20]
Value
Iteration
Density − psi.beta.time[20]
Rhat: 1.08
Density
Parameter estimate

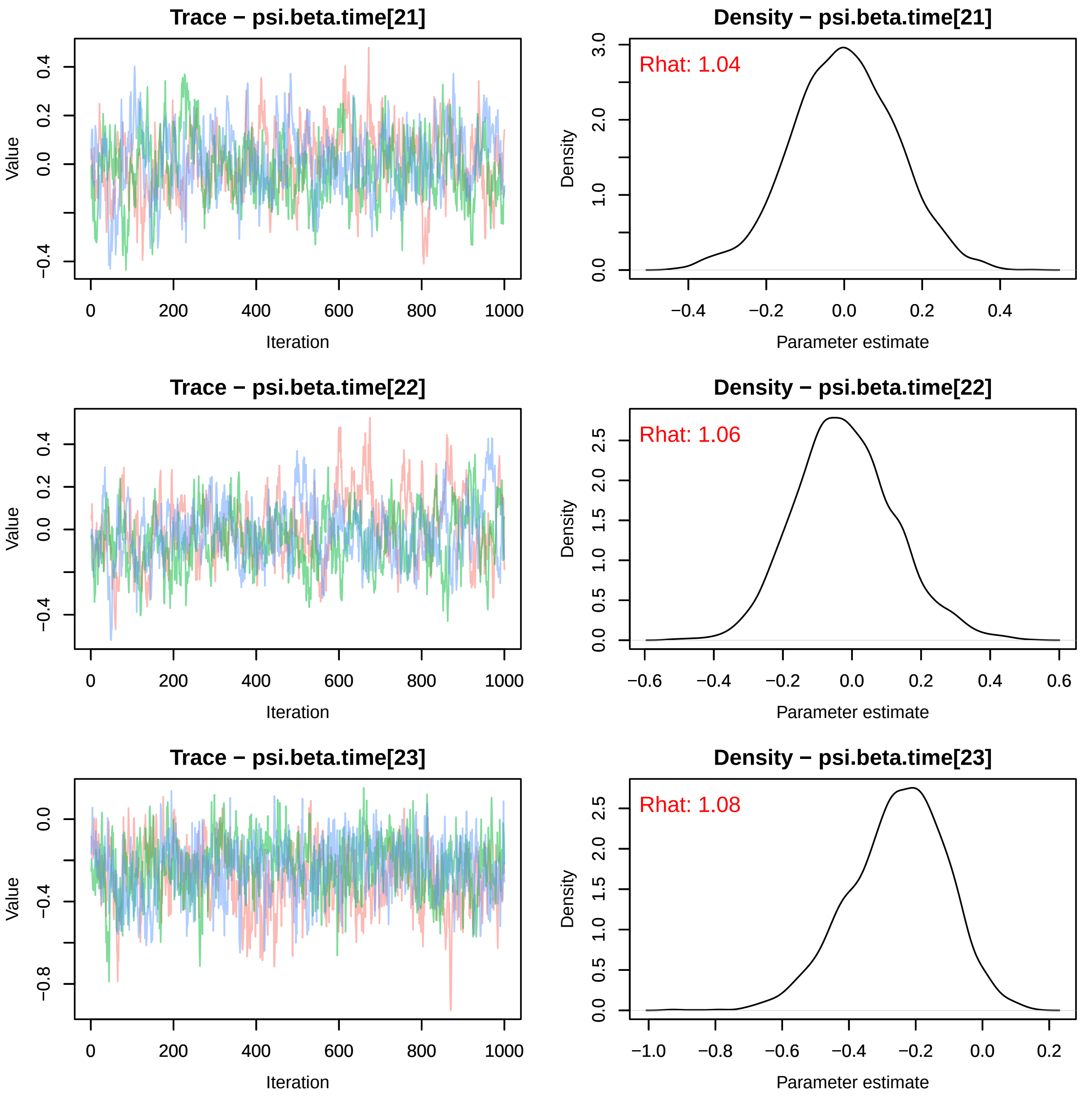
Trace − psi.beta.time[21]
Value
Iteration
Density − psi.beta.time[21]
Rhat: 1.04
Density
Parameter estimate
Trace − psi.beta.time[22]
Value
Iteration
Density − psi.beta.time[22]
Rhat: 1.06
Density
Parameter estimate
Trace − psi.beta.time[23]
Value
Iteration
Density − psi.beta.time[23]
Rhat: 1.08
Density
Parameter estimate

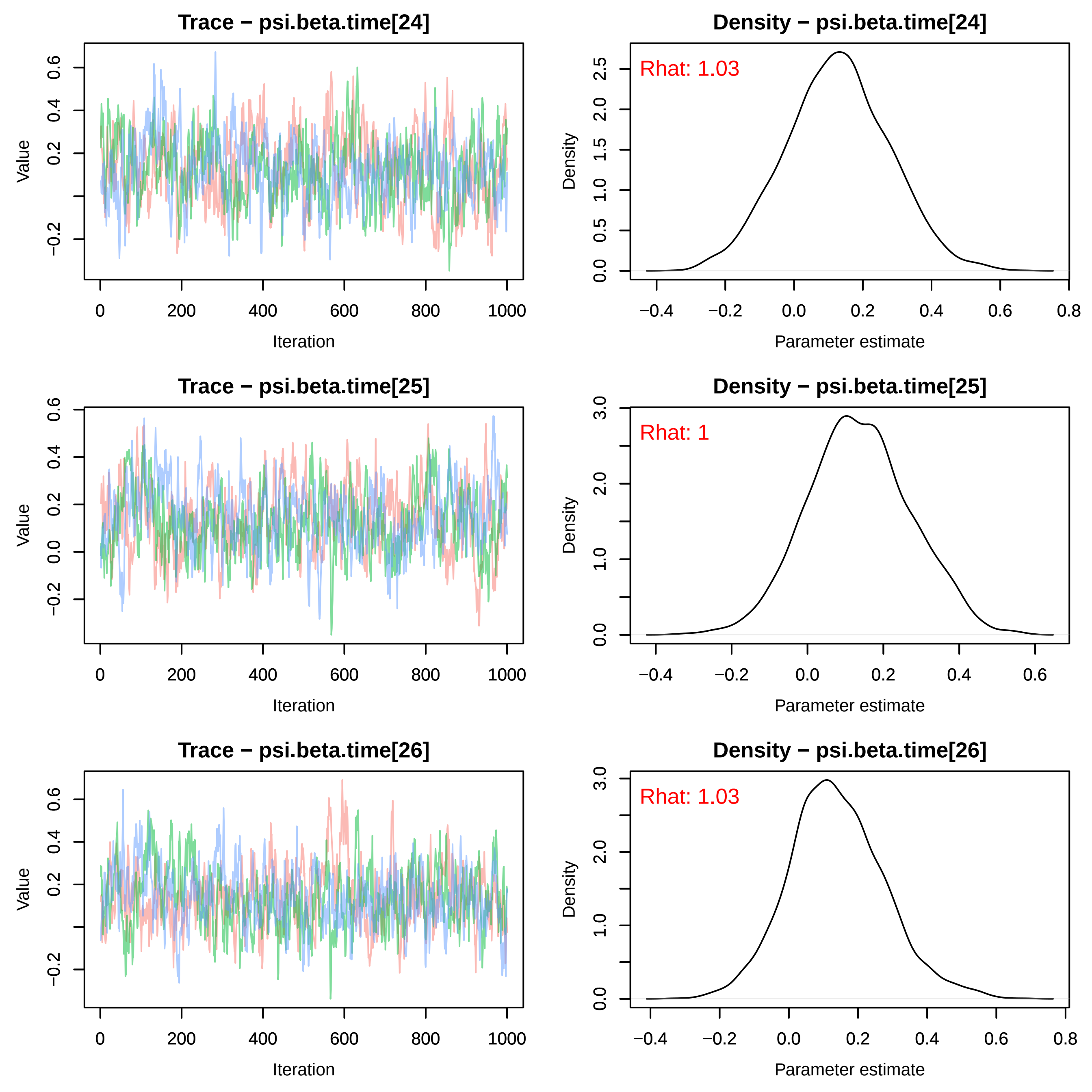
Trace − psi.beta.time[24]
Density − psi.beta.time[24]
Rhat: 1.03
Trace − psi.beta.time[25]
Density − psi.beta.time[25]
Rhat: 1
Trace − psi.beta.time[26]
Density − psi.beta.time[26]
Rhat: 1.03
Value
Iteration
Density
Parameter estimate

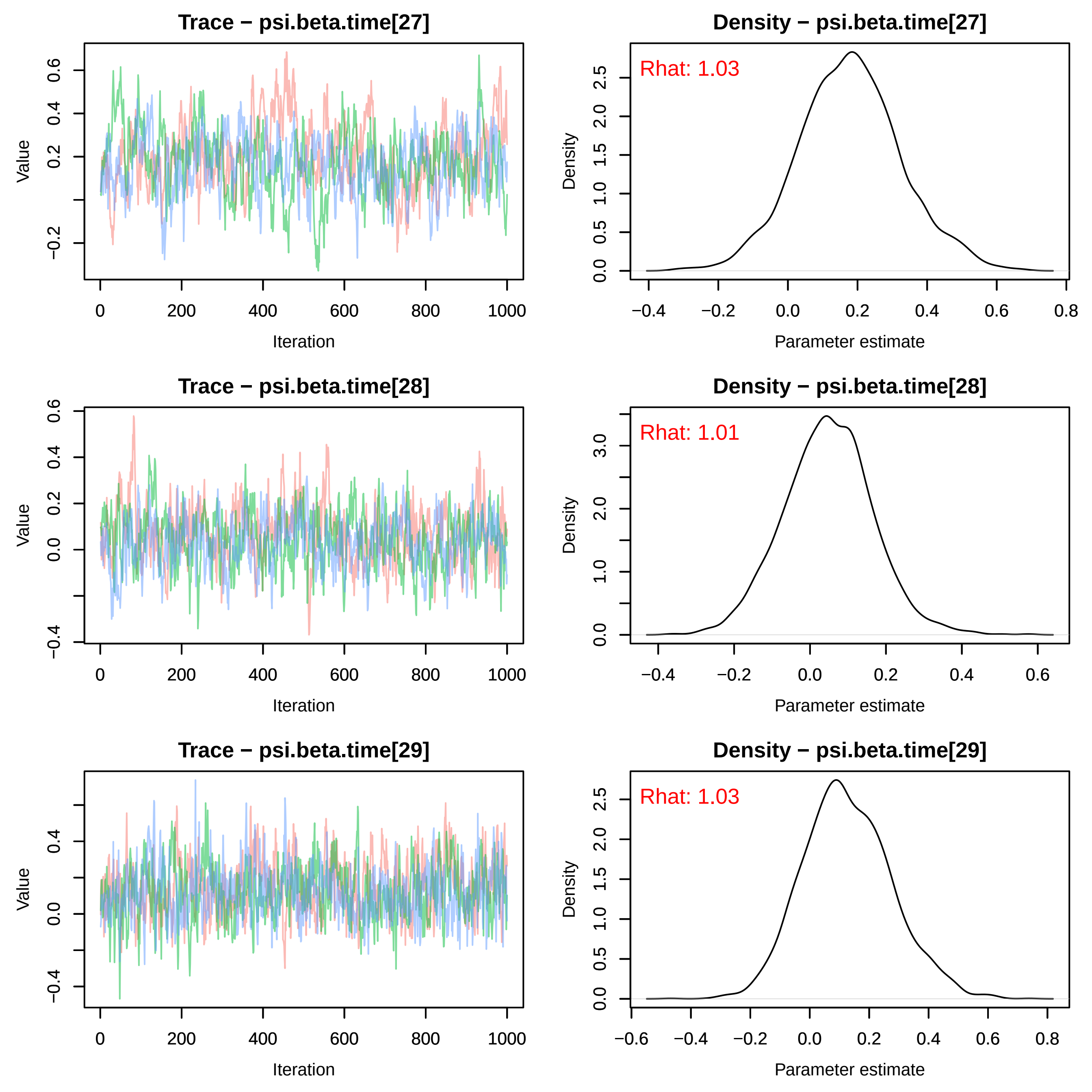
Trace − psi.beta.time[27]
Density − psi.beta.time[27]
Rhat: 1.03
Trace − psi.beta.time[28]
Density − psi.beta.time[28]
Rhat: 1.01
Trace − psi.beta.time[29]
Density − psi.beta.time[29]
Rhat: 1.03
Value
Iteration
Density
Parameter estimate

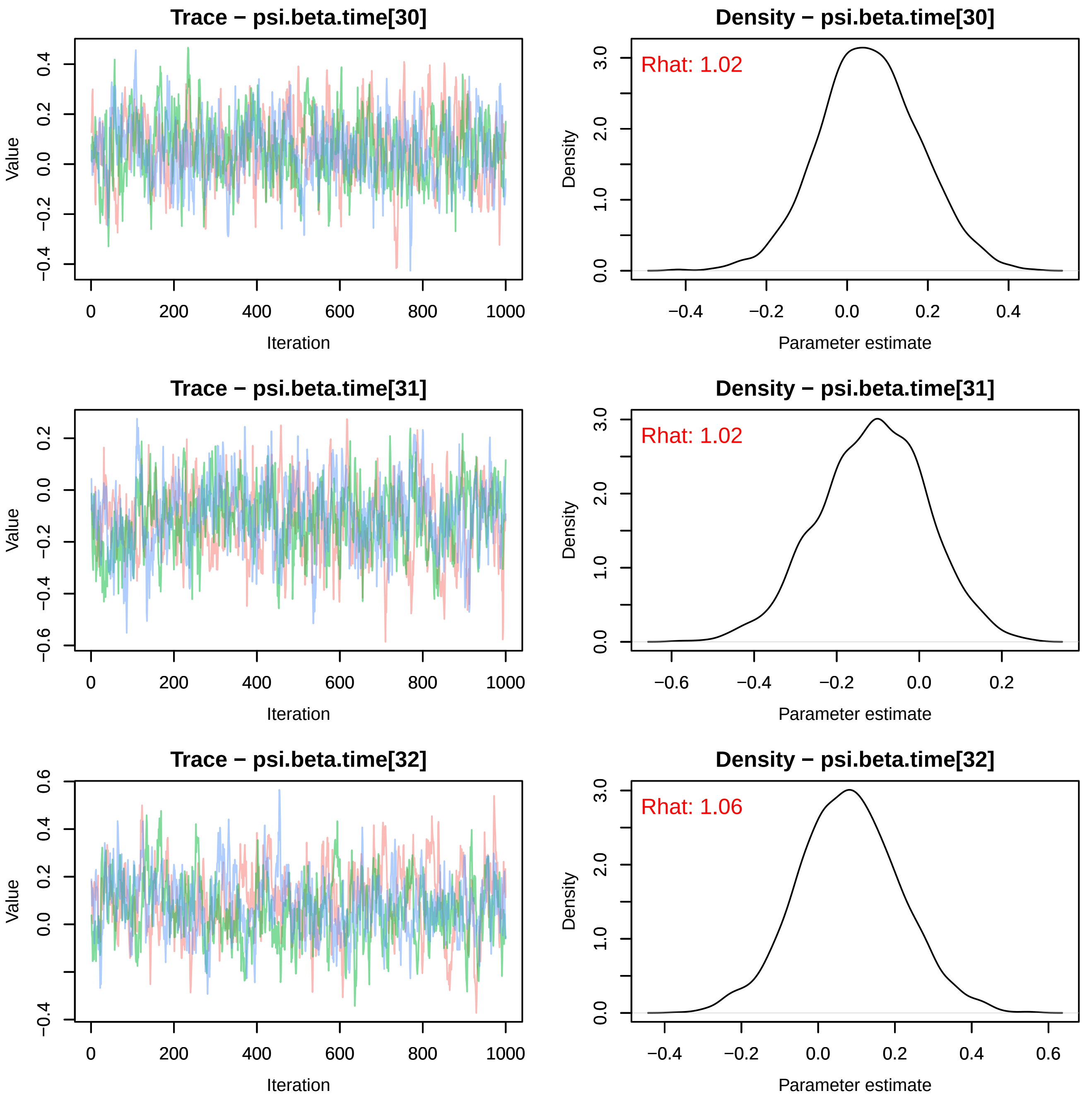
Trace − psi.beta.time[30]
Value
Iteration
0
200
400
600
800
1000
−0.4
−0.2
0.0
0.2
0.4
Density − psi.beta.time[30]
Rhat: 1.02
Density
Parameter estimate
0.0
1.0
2.0
3.0
−0.4
−0.2
0.0
0.2
0.4
Trace − psi.beta.time[31]
Value
Iteration
0
200
400
600
800
1000
−0.6
−0.4
−0.2
0.0
0.2
Density − psi.beta.time[31]
Rhat: 1.02
Density
Parameter estimate
0.0
1.0
2.0
3.0
−0.6
−0.4
−0.2
0.0
0.2
Trace − psi.beta.time[32]
Value
Iteration
0
200
400
600
800
1000
−0.4
0.0
0.2
0.4
0.6
Density − psi.beta.time[32]
Rhat: 1.06
Density
Parameter estimate
0.0
1.0
2.0
3.0
−0.4
−0.2
0.0
0.2
0.4
0.6

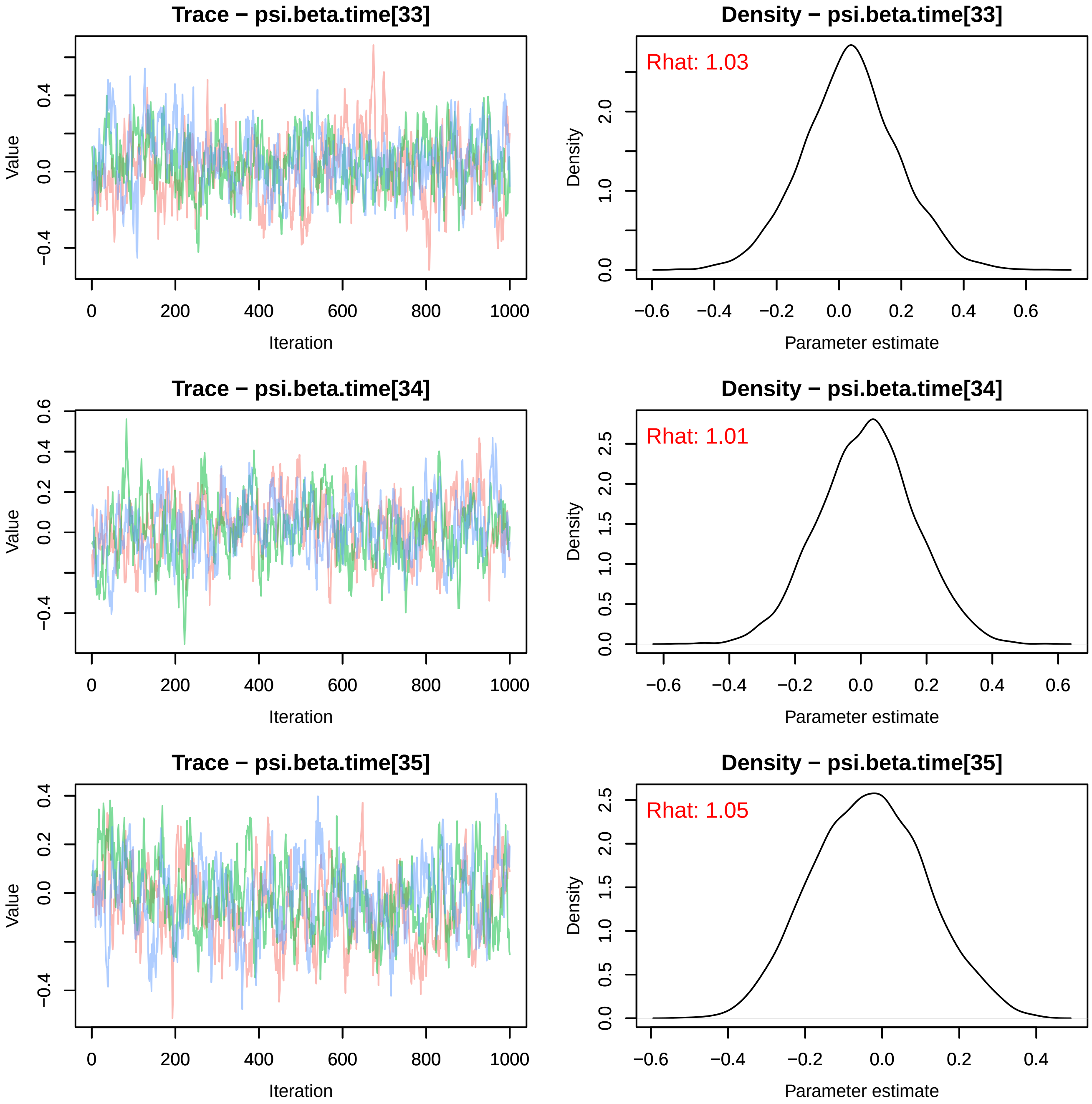

Trace − psi.beta.time[33]
Value
Iteration
Density − psi.beta.time[33]
Rhat: 1.03
Density
Parameter estimate
Trace − psi.beta.time[34]
Value
Iteration
Density − psi.beta.time[34]
Rhat: 1.01
Density
Parameter estimate
Trace − psi.beta.time[35]
Value
Iteration
Density − psi.beta.time[35]
Rhat: 1.05
Density
Parameter estimate

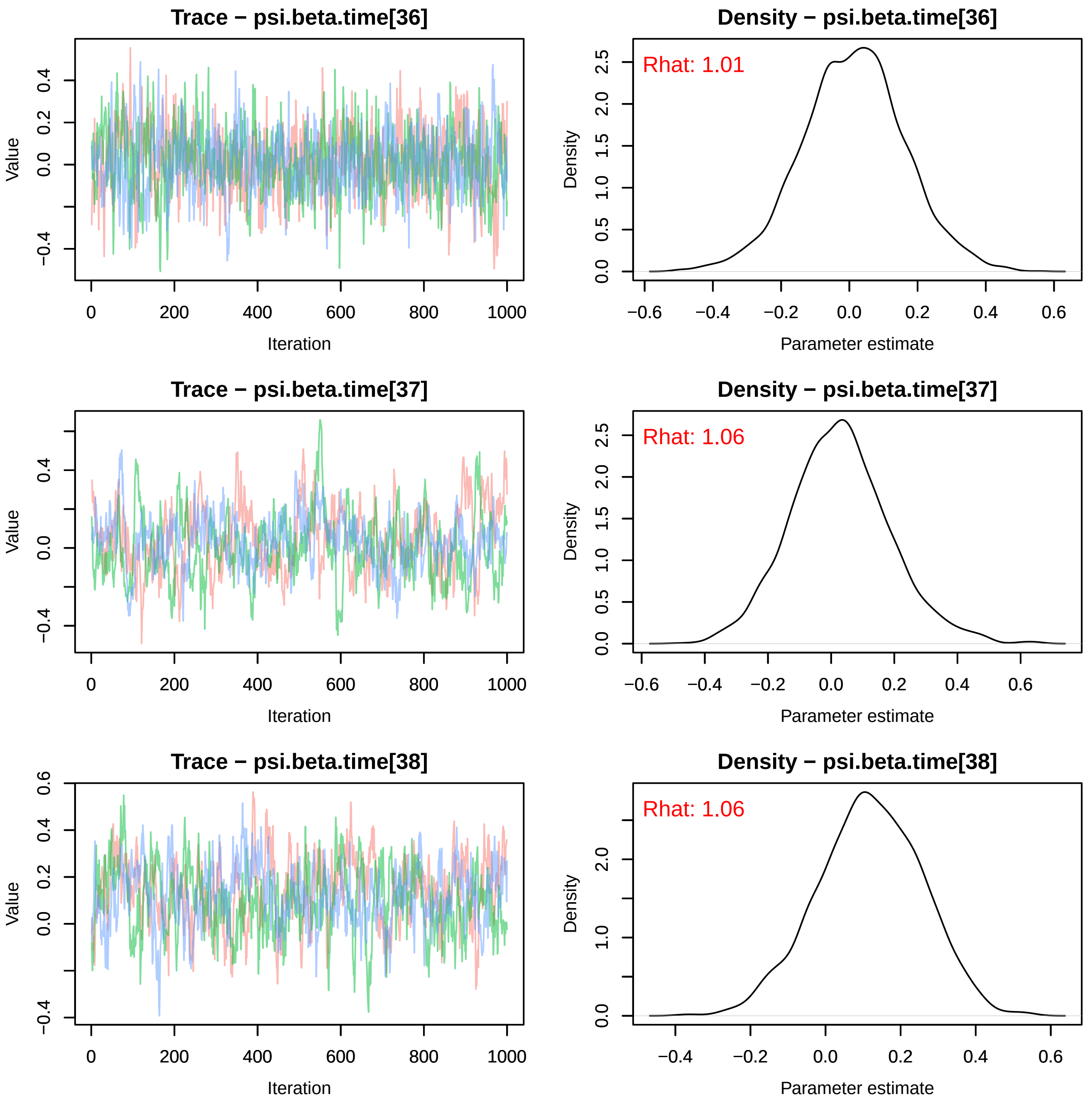
Trace − psi.beta.time[36]
Value
Iteration
Density − psi.beta.time[36]
Rhat: 1.01
Density
Parameter estimate
Trace − psi.beta.time[37]
Value
Iteration
Density − psi.beta.time[37]
Rhat: 1.06
Density
Parameter estimate
Trace − psi.beta.time[38]
Value
Iteration
Density − psi.beta.time[38]
Rhat: 1.06
Density
Parameter estimate

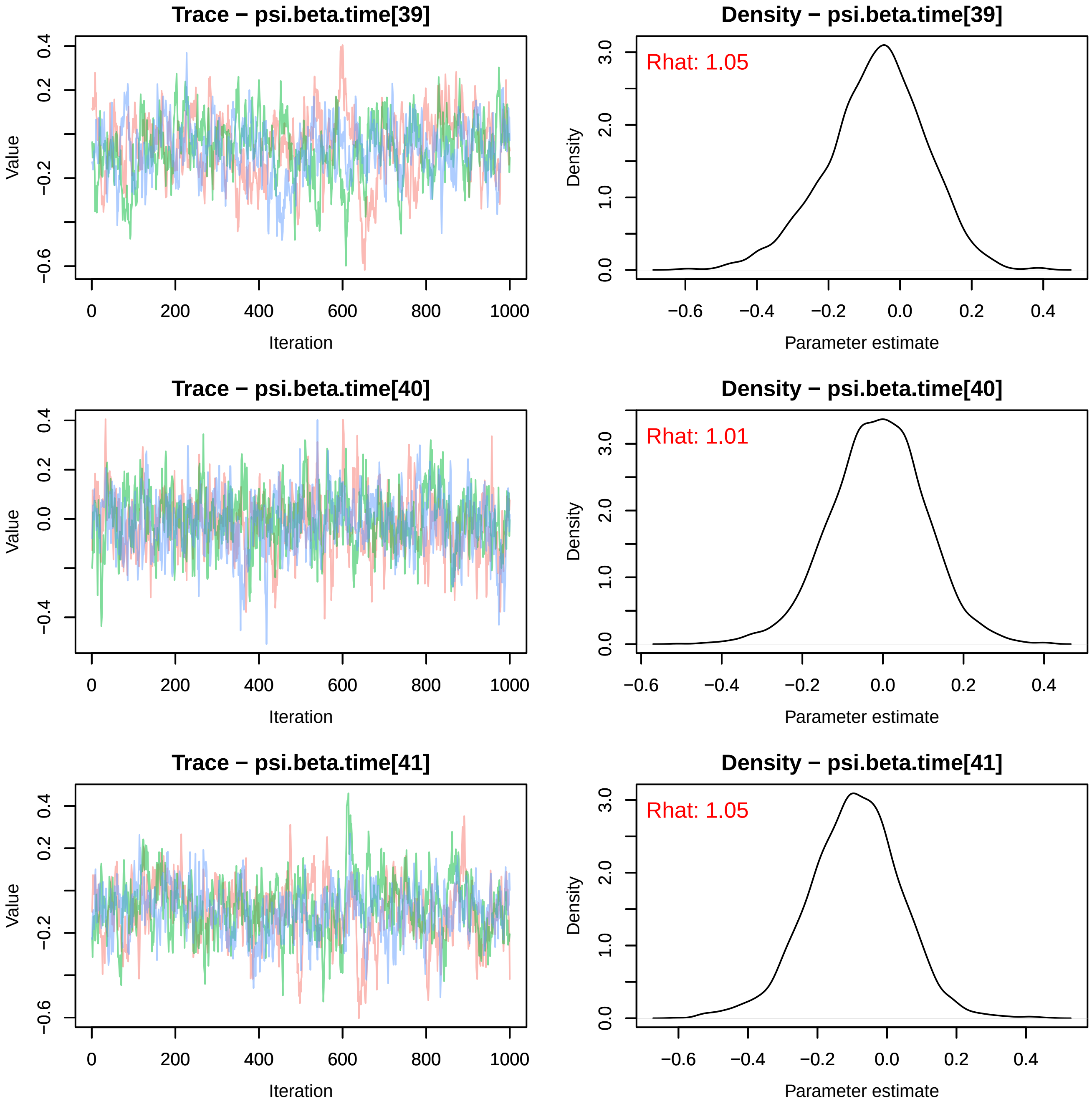
Trace − psi.beta.time[39]
Value
Iteration
Density − psi.beta.time[39]
Rhat: 1.05
Density
Parameter estimate
Trace − psi.beta.time[40]
Value
Iteration
Density − psi.beta.time[40]
Rhat: 1.01
Density
Parameter estimate
Trace − psi.beta.time[41]
Value
Iteration
Density − psi.beta.time[41]
Rhat: 1.05
Density
Parameter estimate

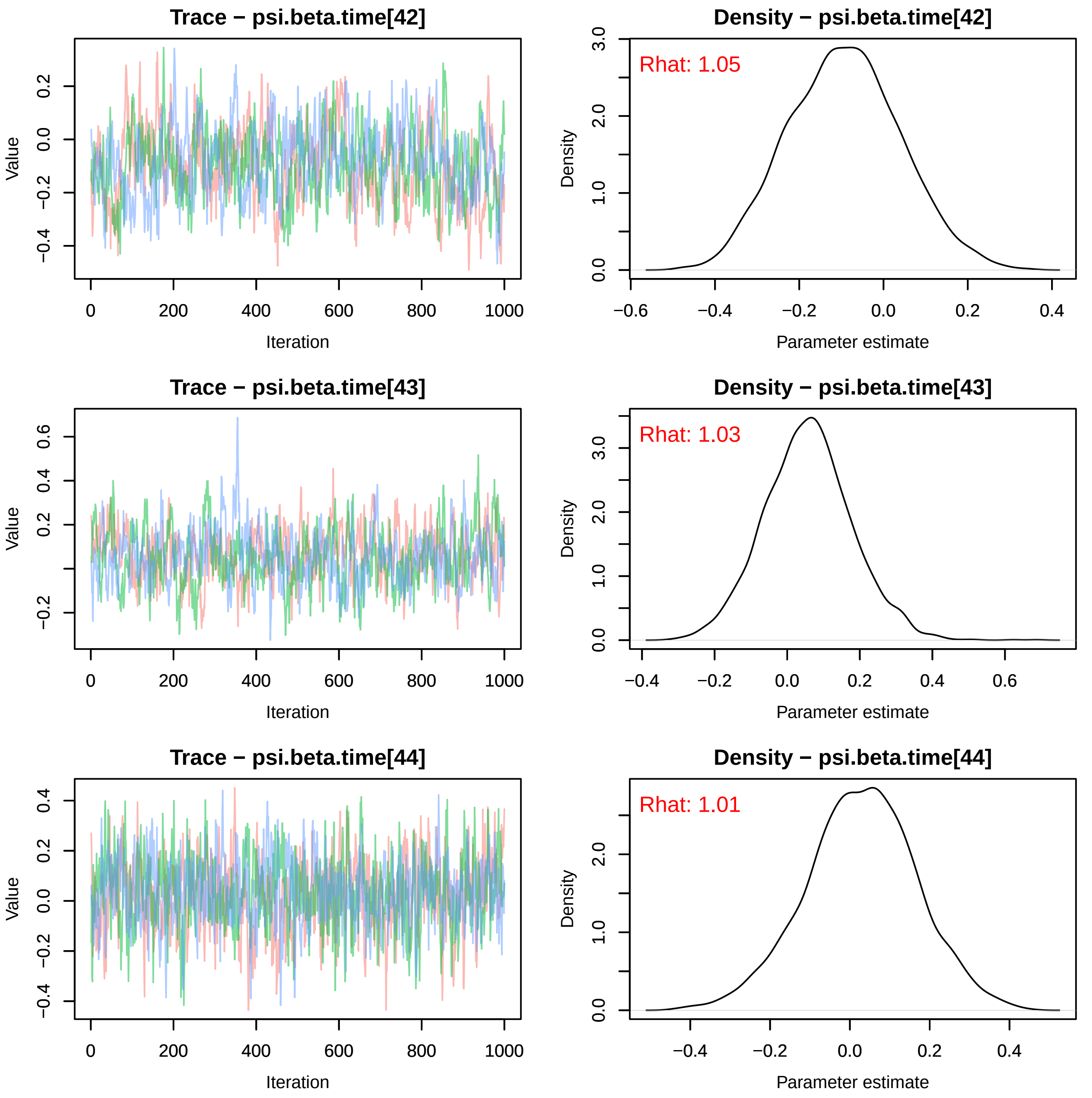
Trace − psi.beta.time[42]
Density − psi.beta.time[42]
Rhat: 1.05
Trace − psi.beta.time[43]
Density − psi.beta.time[43]
Rhat: 1.03
Trace − psi.beta.time[44]
Density − psi.beta.time[44]
Rhat: 1.01
Value
Iteration
Density
Parameter estimate
0
200
400
600
800
1000
−0.6
−0.4
−0.2
0.0
0.2
0.4
0.6
1.0
2.0
3.0

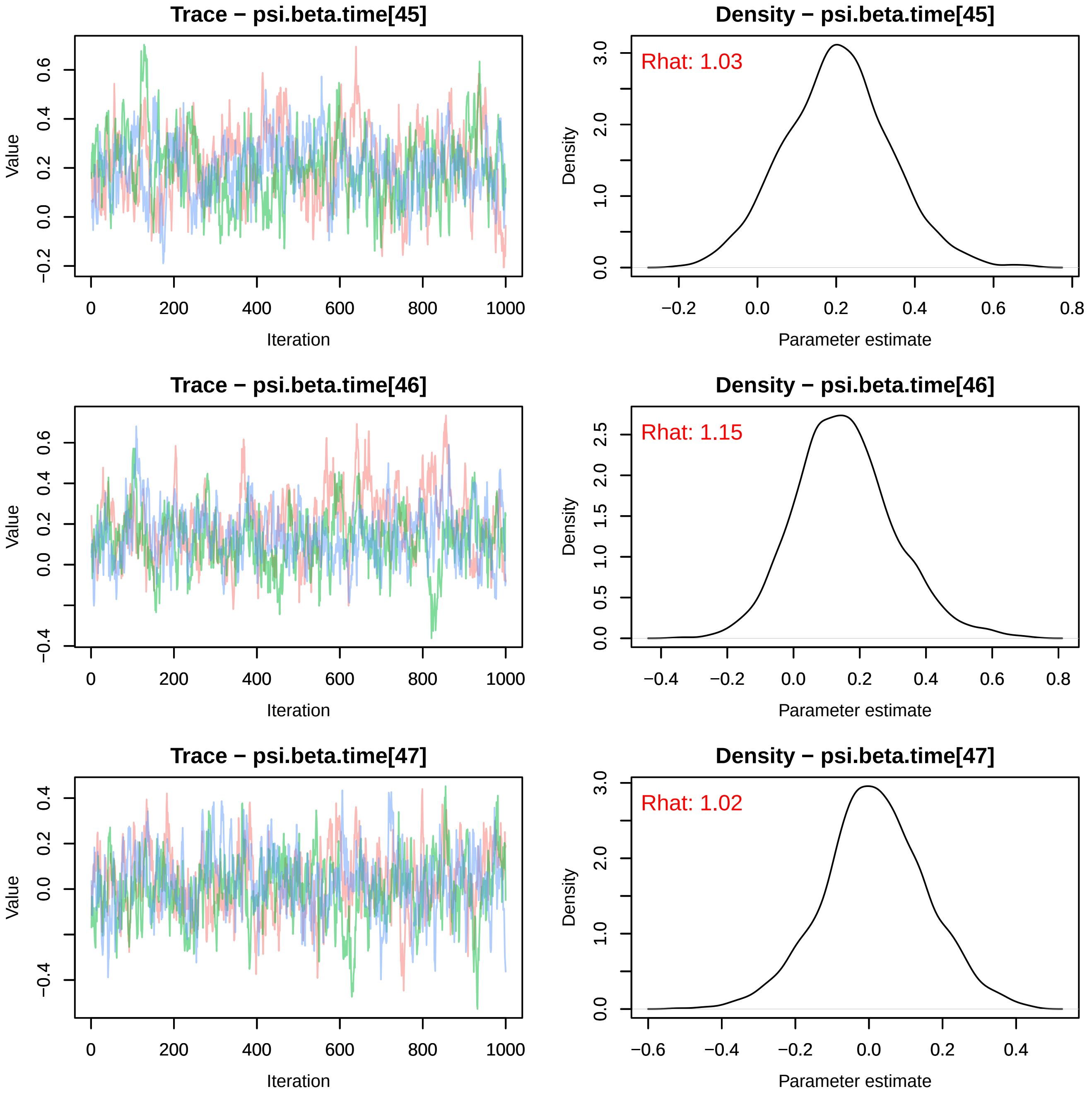
Trace − psi.beta.time[45]
Density − psi.beta.time[45]
Rhat: 1.03
Trace − psi.beta.time[46]
Density − psi.beta.time[46]
Rhat: 1.15
Trace − psi.beta.time[47]
Density − psi.beta.time[47]
Rhat: 1.02
Value
Iteration
Density
Parameter estimate

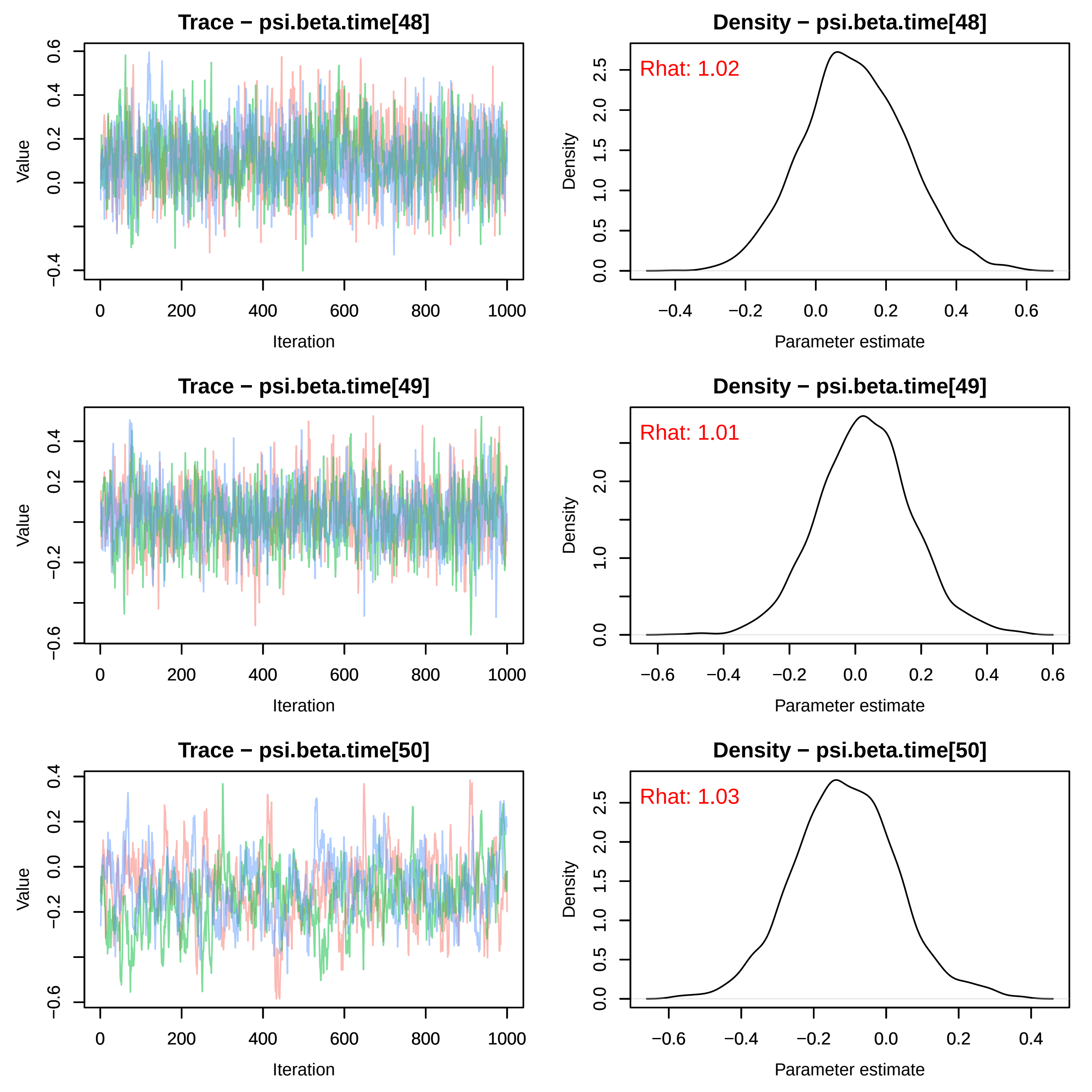

Trace − psi.beta.time[48]
Value
Iteration
Density − psi.beta.time[48]
Rhat: 1.02
Density
Parameter estimate
Trace − psi.beta.time[49]
Value
Iteration
Density − psi.beta.time[49]
Rhat: 1.01
Density
Parameter estimate
Trace − psi.beta.time[50]
Value
Iteration
Density − psi.beta.time[50]
Rhat: 1.03
Density
Parameter estimate

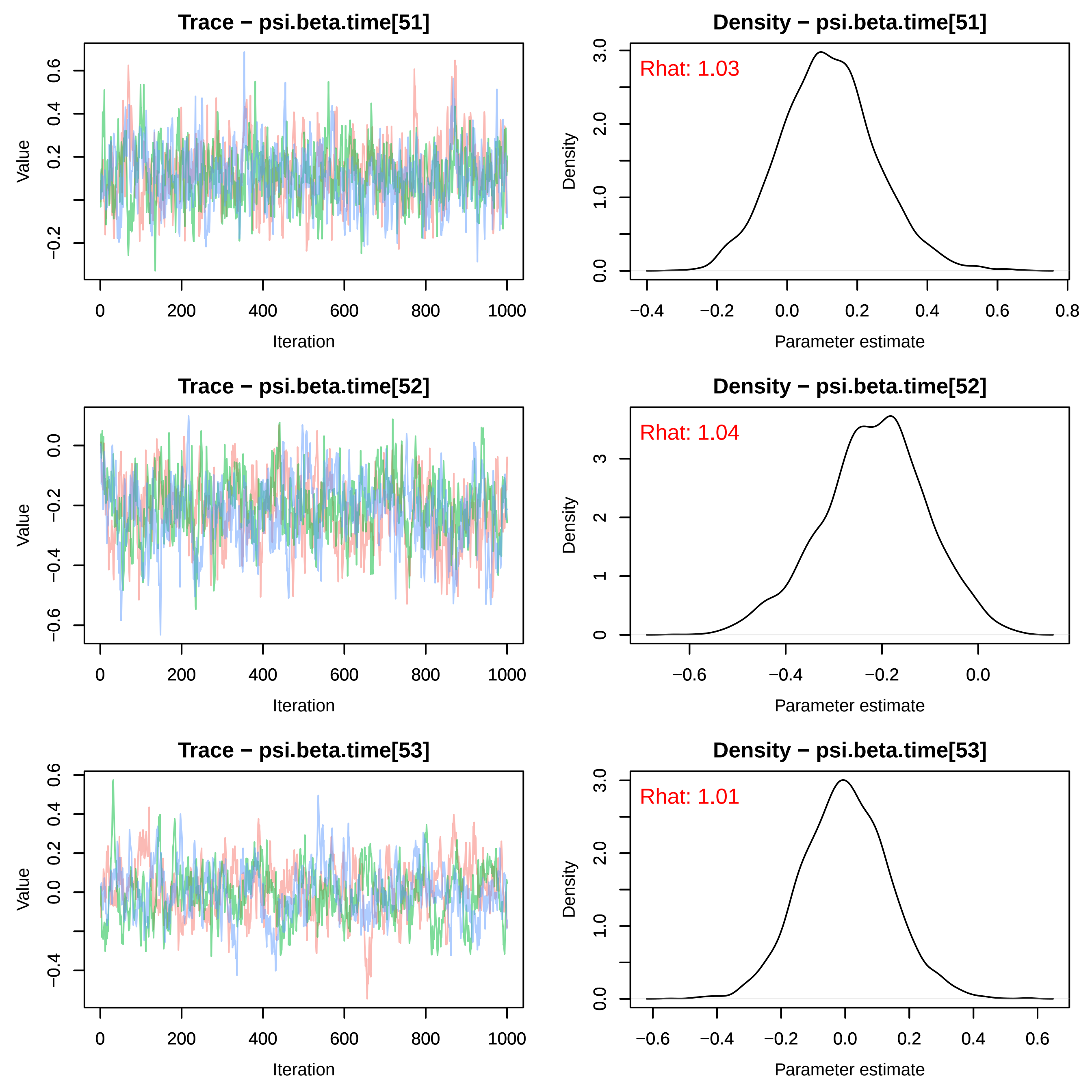
Trace − psi.beta.time[51]
Value
Iteration
Density − psi.beta.time[51]
Rhat: 1.03
Density
Parameter estimate
Trace − psi.beta.time[52]
Value
Iteration
Density − psi.beta.time[52]
Rhat: 1.04
Density
Parameter estimate
Trace − psi.beta.time[53]
Value
Iteration
Density − psi.beta.time[53]
Rhat: 1.01
Density
Parameter estimate

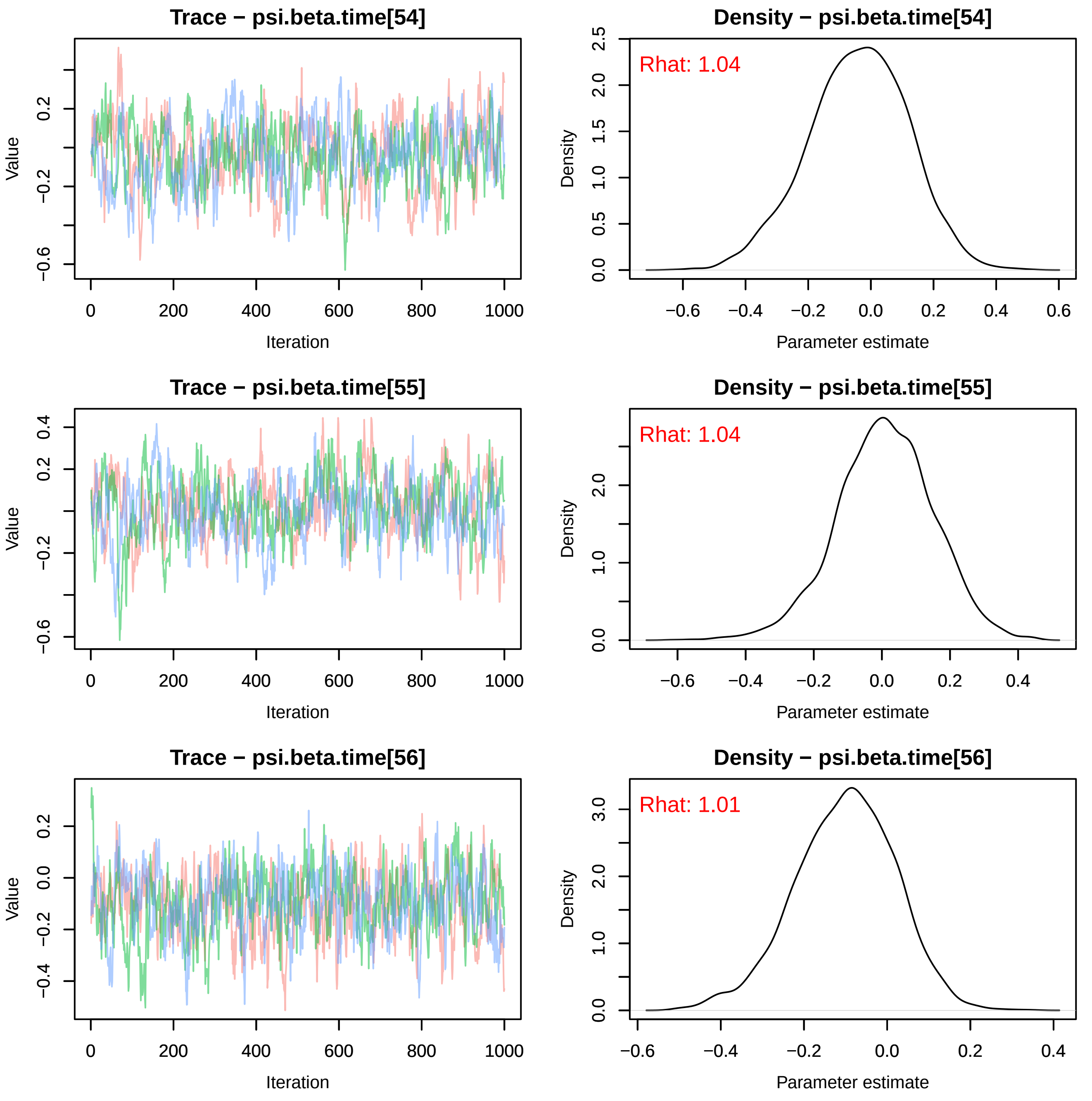
Trace − psi.beta.time[54]
Value
Iteration
Density − psi.beta.time[54]
Rhat: 1.04
Density
Parameter estimate
Trace − psi.beta.time[55]
Value
Iteration
Density − psi.beta.time[55]
Rhat: 1.04
Density
Parameter estimate
Trace − psi.beta.time[56]
Value
Iteration
Density − psi.beta.time[56]
Rhat: 1.01
Density
Parameter estimate

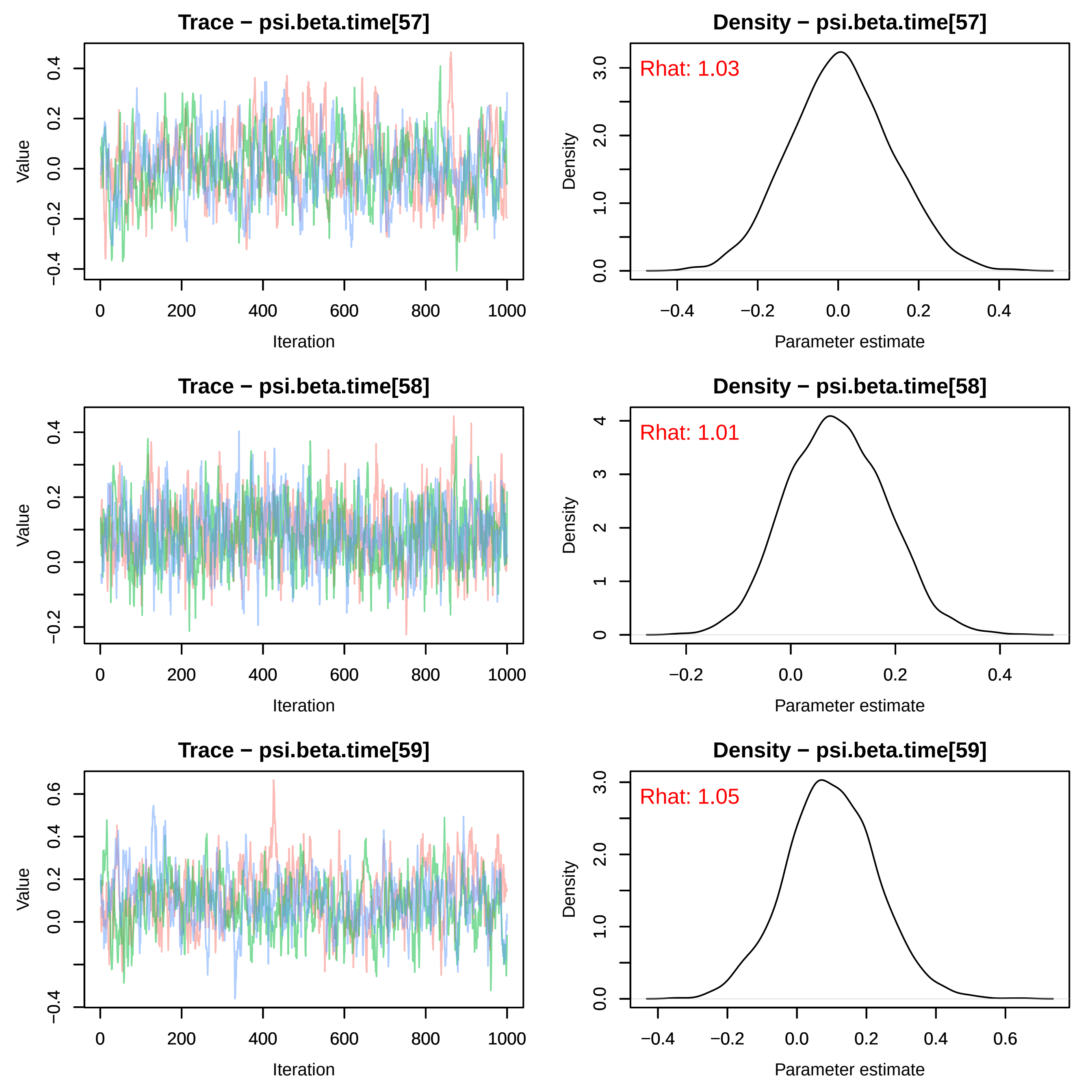
Trace − psi.beta.time[57]
Density − psi.beta.time[57]
Rhat: 1.03
Trace − psi.beta.time[58]
Density − psi.beta.time[58]
Rhat: 1.01
Trace − psi.beta.time[59]
Density − psi.beta.time[59]
Rhat: 1.05
Value
Iteration
Density
Parameter estimate

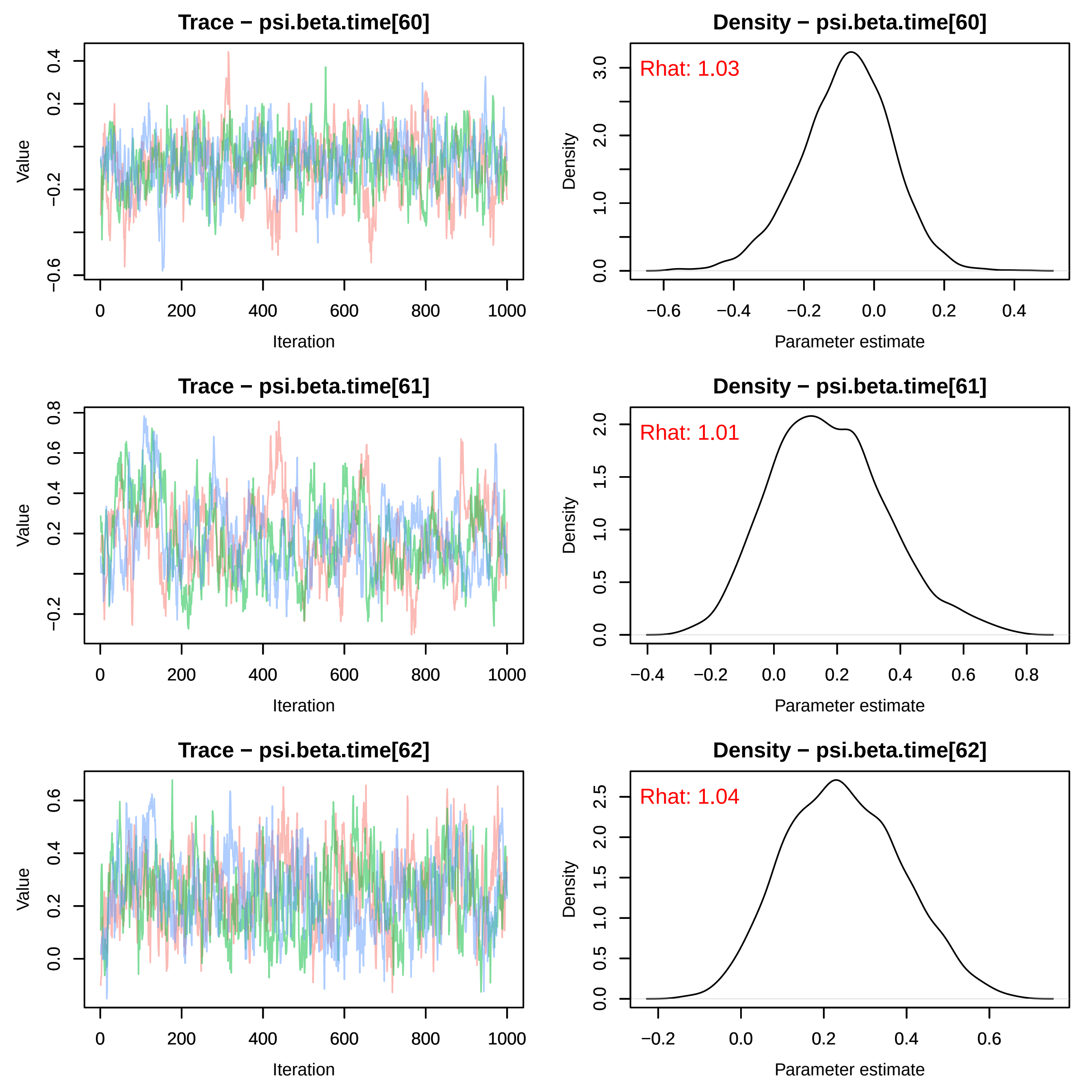
Trace − psi.beta.time[60]
Value
Iteration
0
200
400
600
800
1000
−0.6
−0.2
0.2
0.4
Density − psi.beta.time[60]
Rhat: 1.03
Density
Parameter estimate
−0.6
−0.4
−0.2
0.0
0.2
0.4
0.0
1.0
2.0
3.0
Trace − psi.beta.time[61]
Value
Iteration
−0.2
0.2
0.4
0.6
0.8
Density − psi.beta.time[61]
Rhat: 1.01
Density
Parameter estimate
−0.4
−0.2
0.0
0.2
0.4
0.6
0.8
0.0
0.5
1.0
1.5
2.0
Trace − psi.beta.time[62]
Value
Iteration
0.0
0.2
0.4
0.6
Density − psi.beta.time[62]
Rhat: 1.04
Density
Parameter estimate
−0.2
0.0
0.2
0.4
0.6
0.0
0.5
1.0
1.5
2.0
2.5

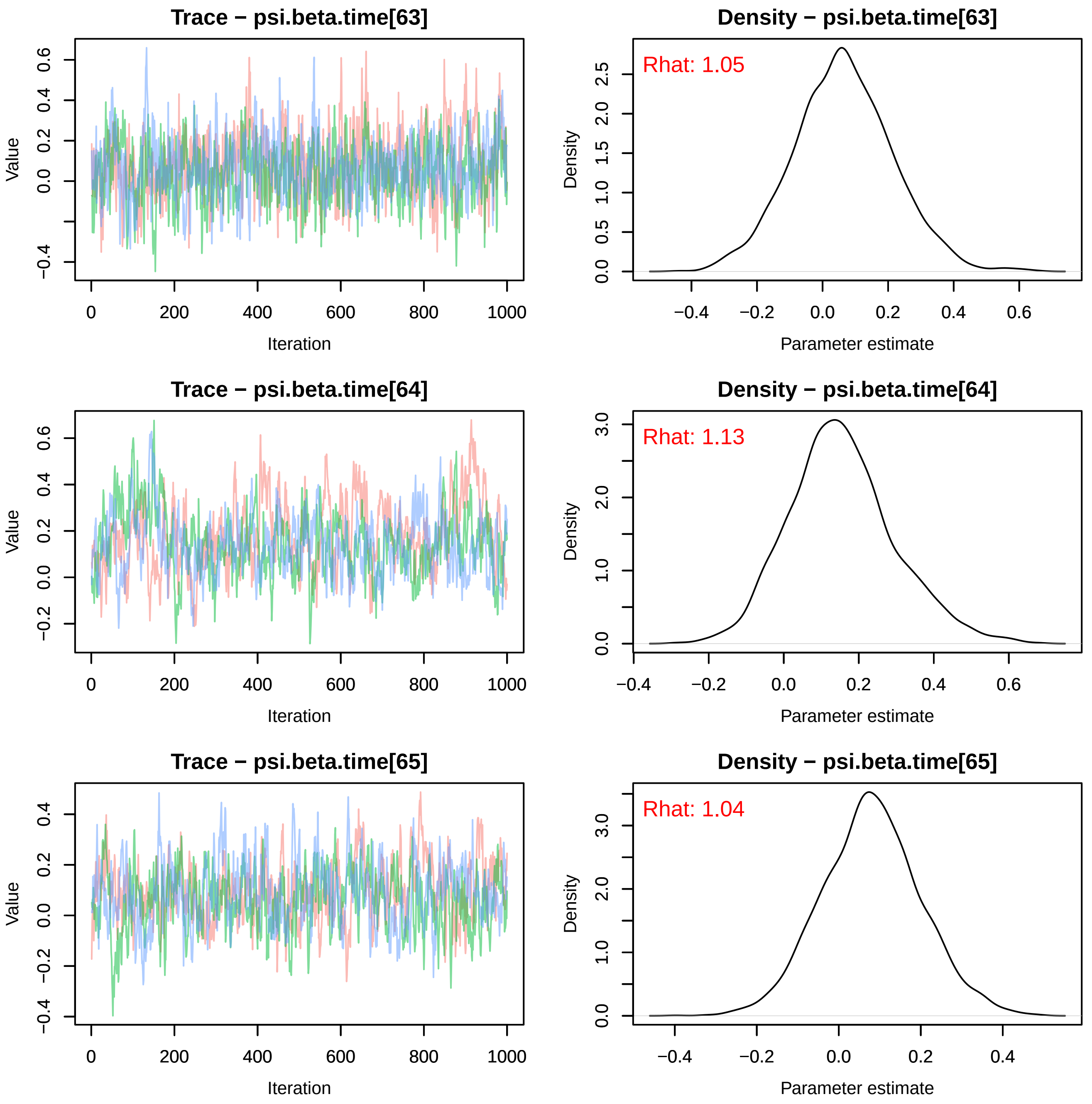
Trace − psi.beta.time[63]
Density − psi.beta.time[63]
Rhat: 1.05
Trace − psi.beta.time[64]
Density − psi.beta.time[64]
Rhat: 1.13
Trace − psi.beta.time[65]
Density − psi.beta.time[65]
Rhat: 1.04
Value
Iteration
Density
Parameter estimate

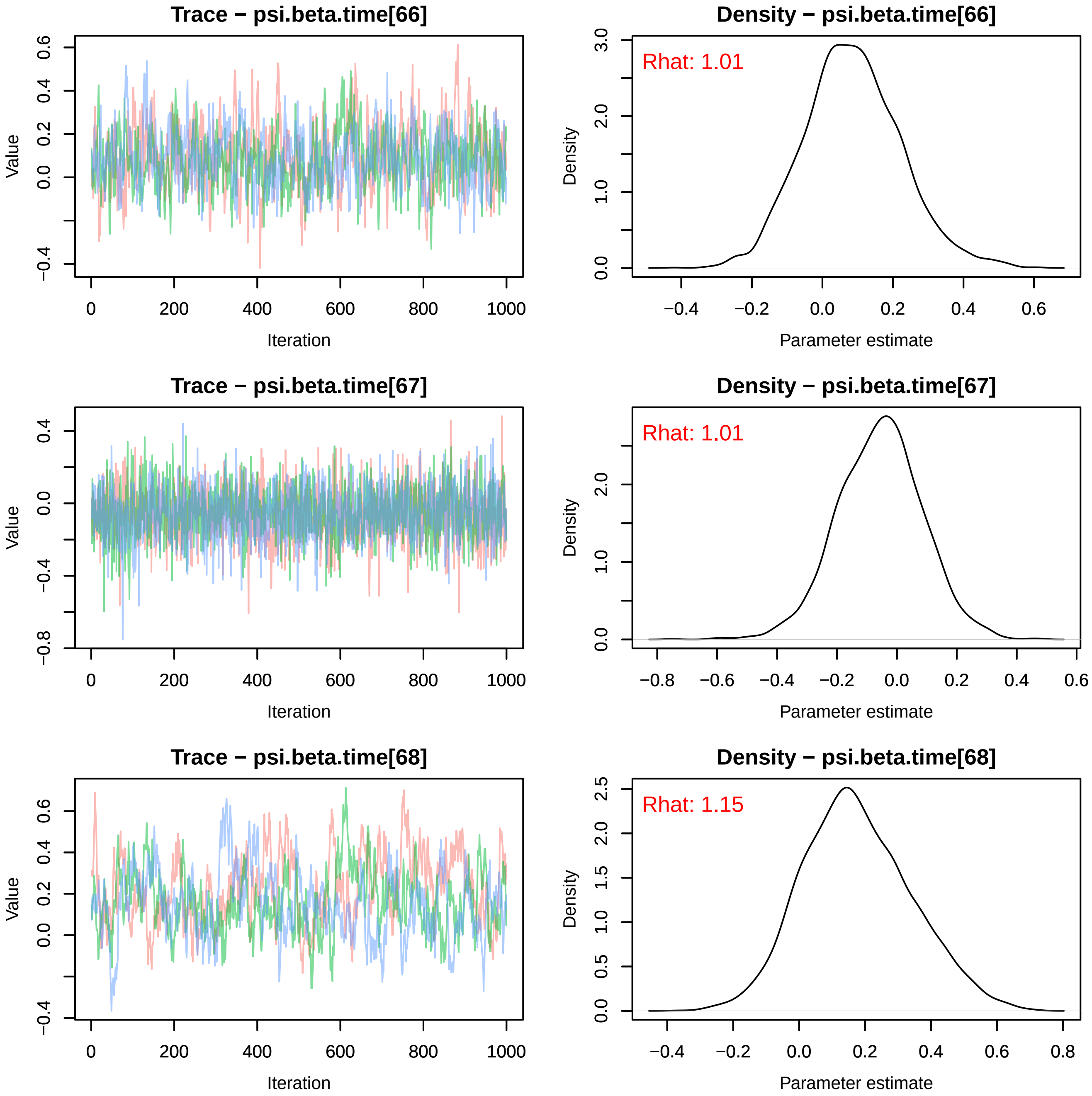

Trace − psi.beta.time[66]
Value
Iteration
Density − psi.beta.time[66]
Rhat: 1.01
Density
Parameter estimate
Trace − psi.beta.time[67]
Value
Iteration
Density − psi.beta.time[67]
Rhat: 1.01
Density
Parameter estimate
Trace − psi.beta.time[68]
Value
Iteration
Density − psi.beta.time[68]
Rhat: 1.15
Density
Parameter estimate

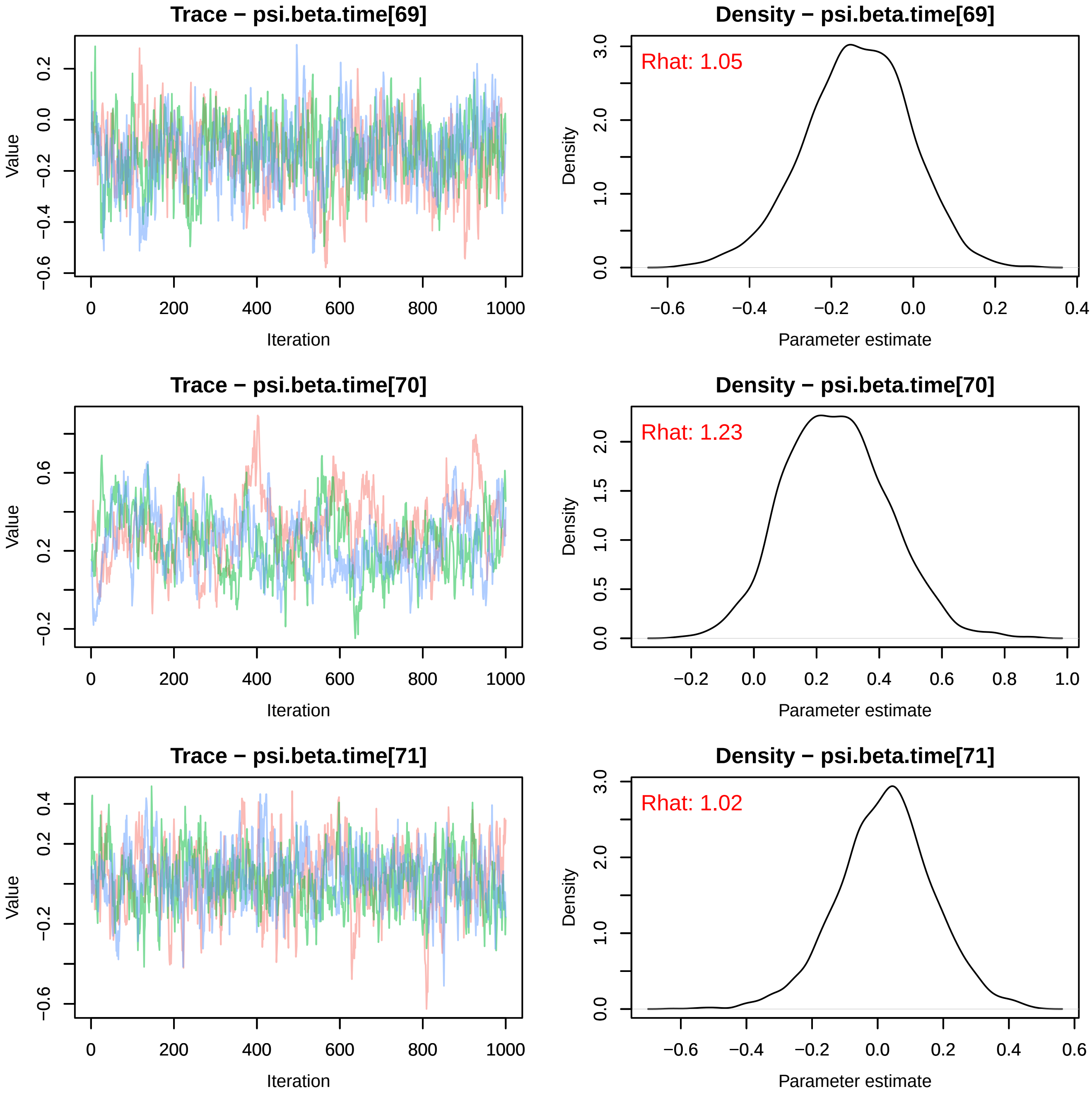
Trace − psi.beta.time[69]
Value
−0.6
−0.4
−0.2
0.0
0.2
0
200
400
600
800
1000
Iteration
Density − psi.beta.time[69]
Rhat: 1.05
Density
0.0
1.0
2.0
3.0
−0.6
−0.4
−0.2
0.0
0.2
0.4
Parameter estimate
Trace − psi.beta.time[70]
Value
−0.2
0.2
0.6
0
200
400
600
800
1000
Iteration
Density − psi.beta.time[70]
Rhat: 1.23
Density
0.0
0.5
1.0
1.5
2.0
−0.2
0.0
0.2
0.4
0.6
0.8
1.0
Parameter estimate
Trace − psi.beta.time[71]
Value
−0.6
−0.2
0.2
0.4
0
200
400
600
800
1000
Iteration
Density − psi.beta.time[71]
Rhat: 1.02
Density
0.0
1.0
2.0
3.0
−0.6
−0.4
−0.2
0.0
0.2
0.4
0.6
Parameter estimate

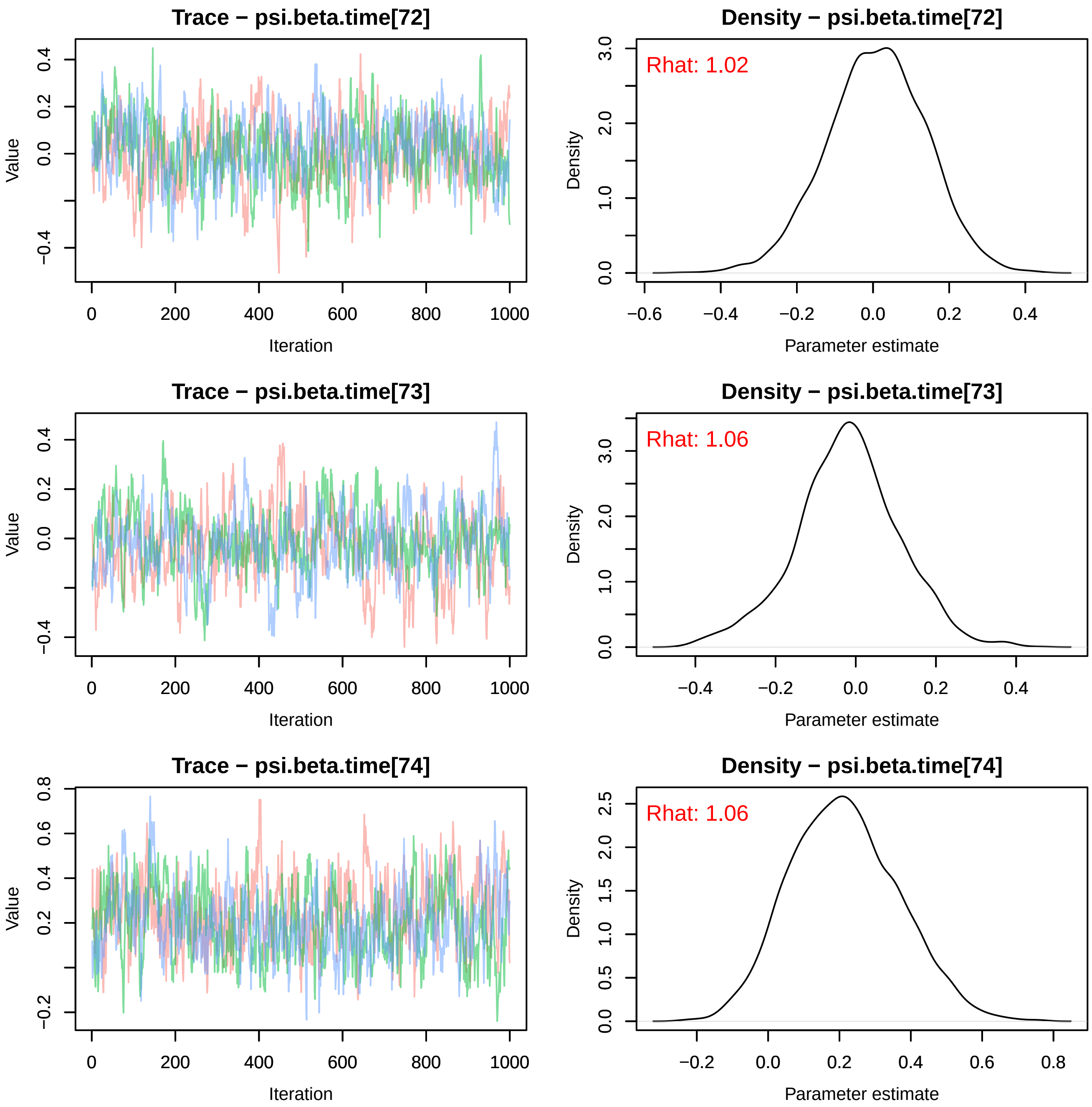

Trace − psi.beta.time[72]
Density − psi.beta.time[72]
Rhat: 1.02
Trace − psi.beta.time[73]
Density − psi.beta.time[73]
Rhat: 1.06
Trace − psi.beta.time[74]
Density − psi.beta.time[74]
Rhat: 1.06
Value
Iteration
Density
Parameter estimate

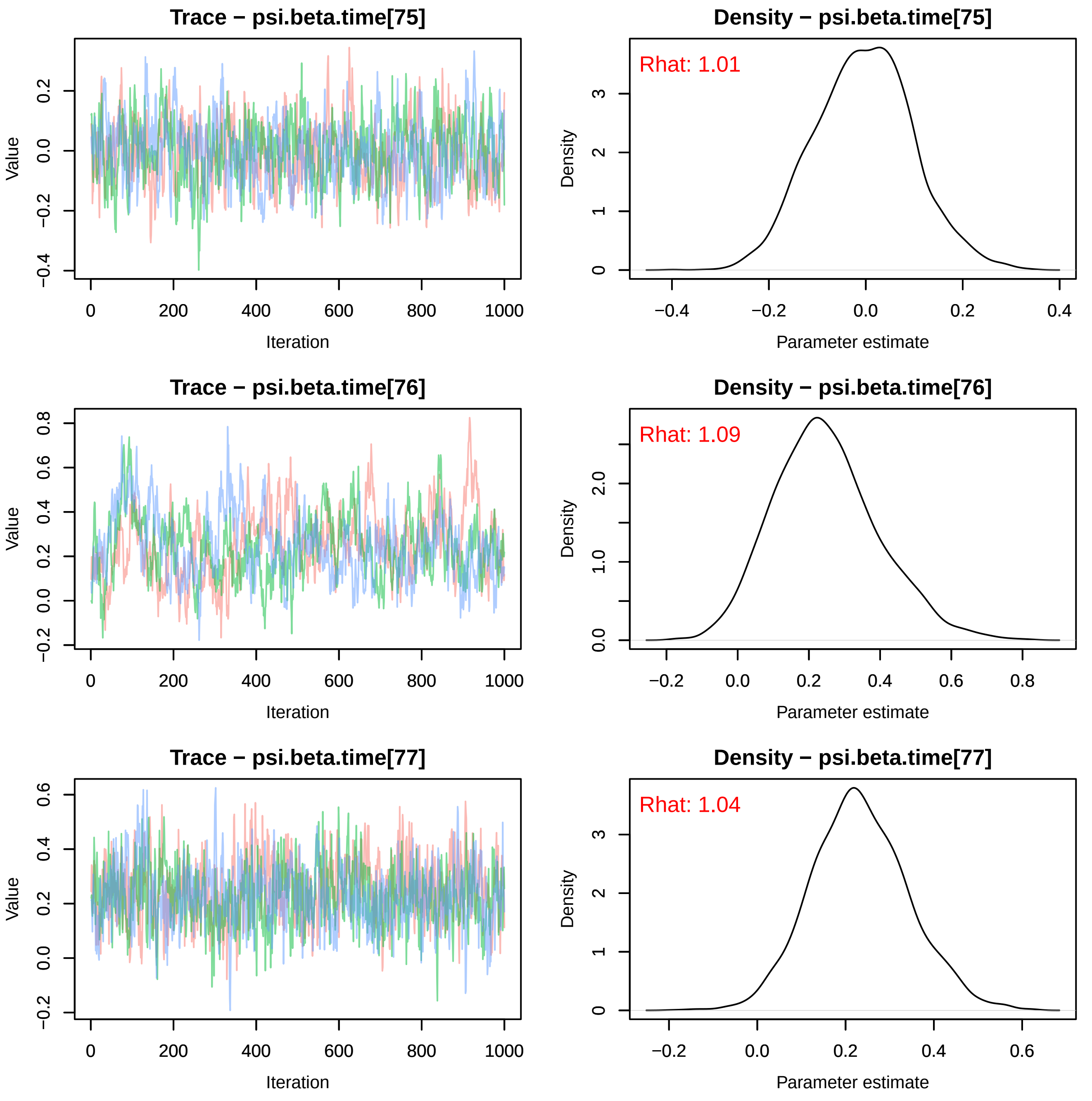

Trace − psi.beta.time[75]
Density − psi.beta.time[75]
Rhat: 1.01
Trace − psi.beta.time[76]
Density − psi.beta.time[76]
Rhat: 1.09
Trace − psi.beta.time[77]
Density − psi.beta.time[77]
Rhat: 1.04
Value
Iteration
Density
Parameter estimate
0
200
400
600
800
1000
−0.4
−0.2
0.0
0.2
0.4
0.6
0.8
1.0
2.0
1
2
3

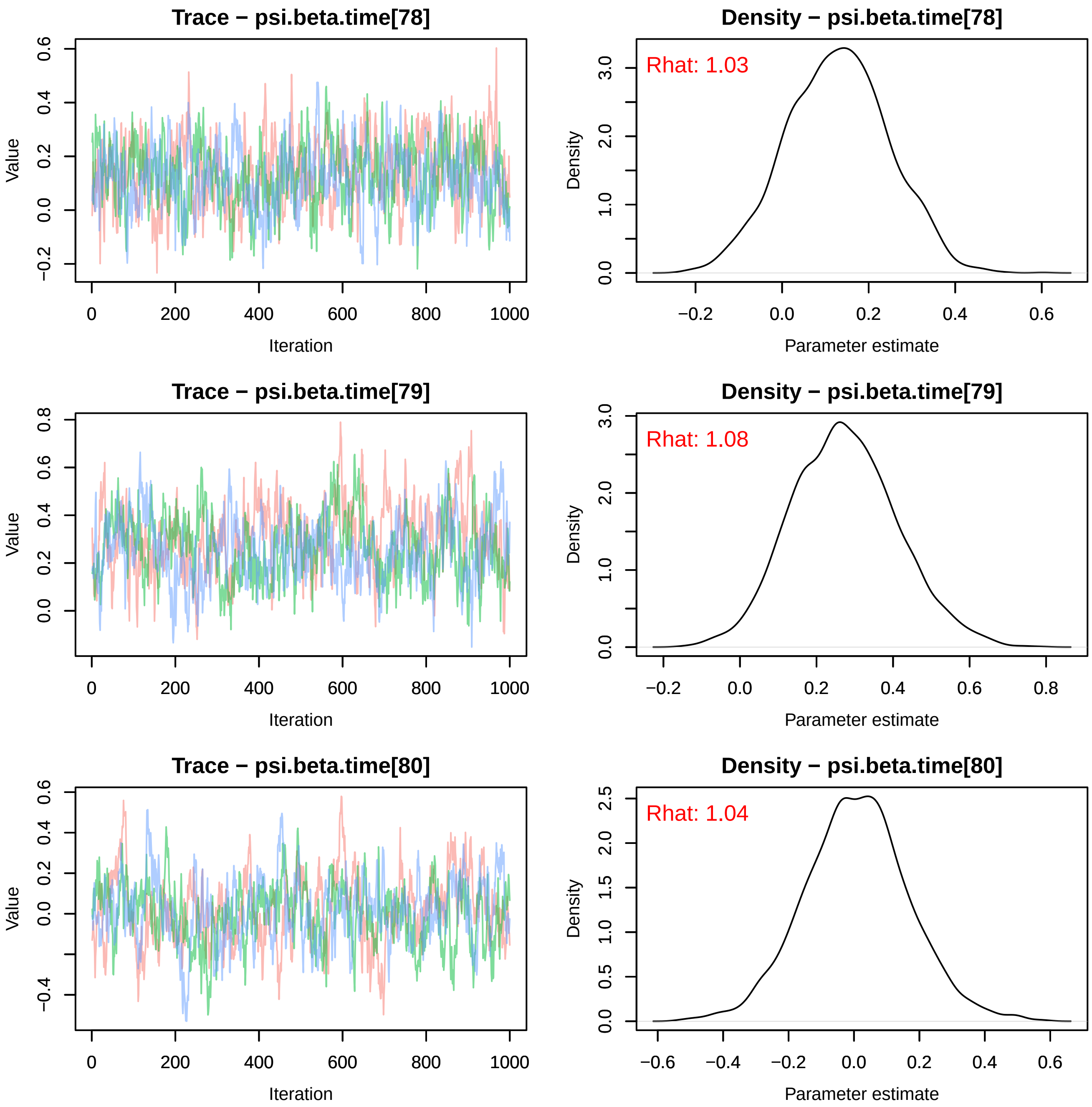
Trace − psi.beta.time[78]
Value
Iteration
0
200
400
600
800
1000
−0.2
0.0
0.2
0.4
0.6
Density − psi.beta.time[78]
Rhat: 1.03
Density
Parameter estimate
0.0
1.0
2.0
3.0
−0.2
0.0
0.2
0.4
0.6
Trace − psi.beta.time[79]
Value
Iteration
0
200
400
600
800
1000
0.0
0.2
0.4
0.6
0.8
Density − psi.beta.time[79]
Rhat: 1.08
Density
Parameter estimate
0.0
1.0
2.0
3.0
−0.2
0.0
0.2
0.4
0.6
0.8
Trace − psi.beta.time[80]
Value
Iteration
0
200
400
600
800
1000
−0.4
0.0
0.2
0.4
0.6
Density − psi.beta.time[80]
Rhat: 1.04
Density
Parameter estimate
0.0
0.5
1.0
1.5
2.0
2.5
−0.6
−0.4
−0.2
0.0
0.2
0.4
0.6

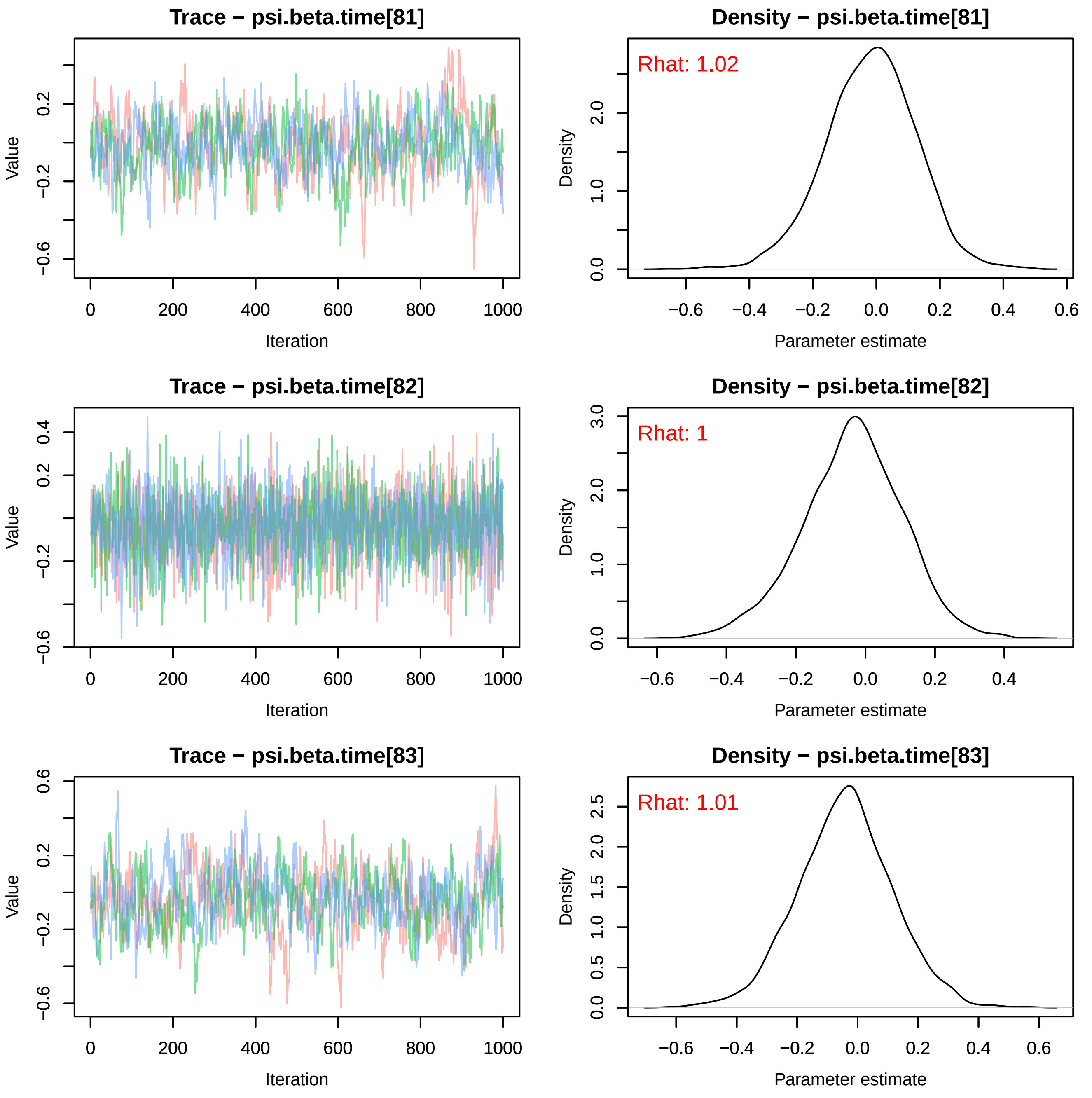
Trace − psi.beta.time[81]
Density − psi.beta.time[81]
Rhat: 1.02
Trace − psi.beta.time[82]
Density − psi.beta.time[82]
Rhat: 1
Trace − psi.beta.time[83]
Density − psi.beta.time[83]
Rhat: 1.01
Value
Iteration
Density
Parameter estimate

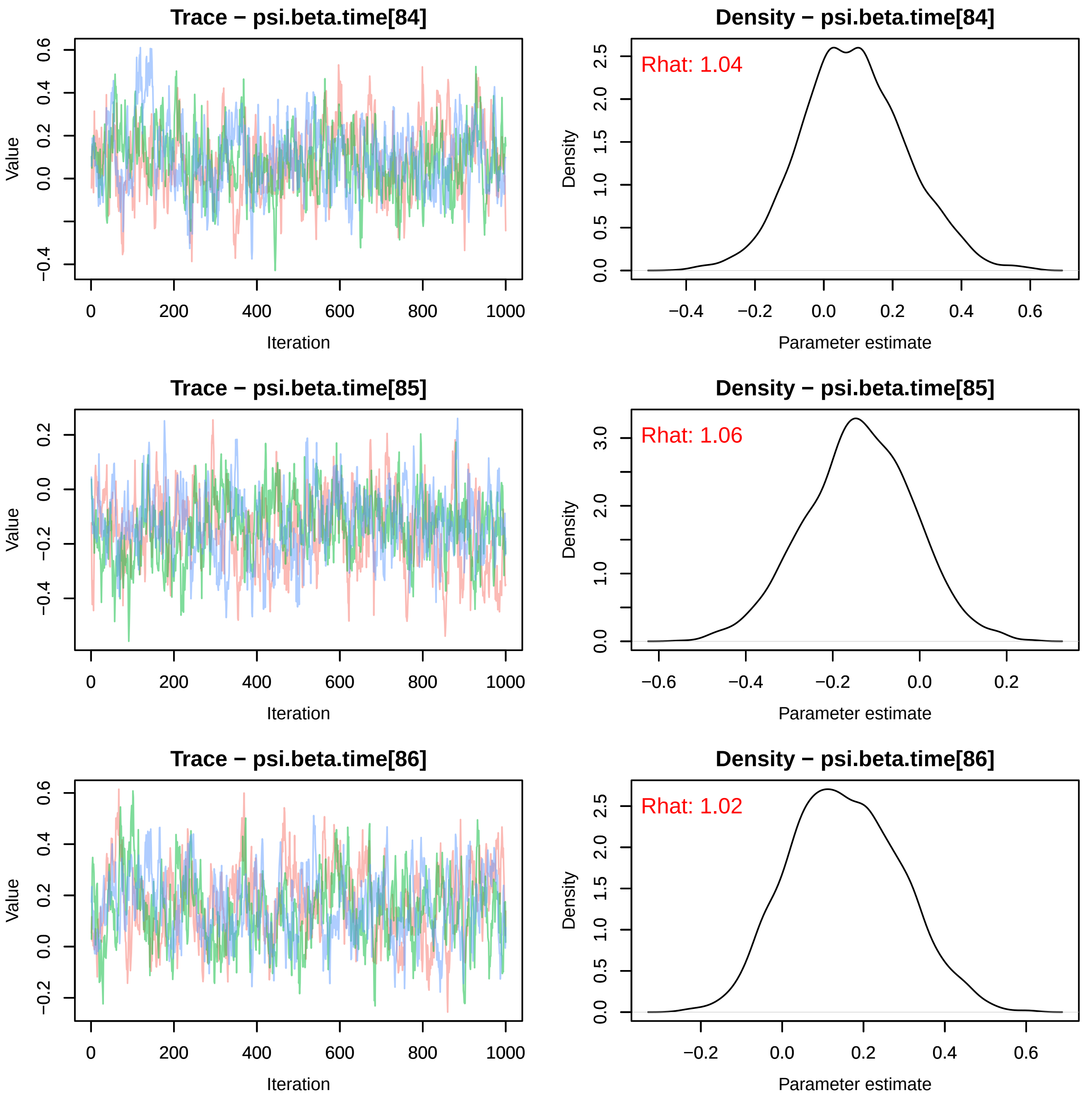
Trace − psi.beta.time[84]
Value
Iteration
Density − psi.beta.time[84]
Rhat: 1.04
Density
Parameter estimate
Trace − psi.beta.time[85]
Value
Iteration
Density − psi.beta.time[85]
Rhat: 1.06
Density
Parameter estimate
Trace − psi.beta.time[86]
Value
Iteration
Density − psi.beta.time[86]
Rhat: 1.02
Density
Parameter estimate

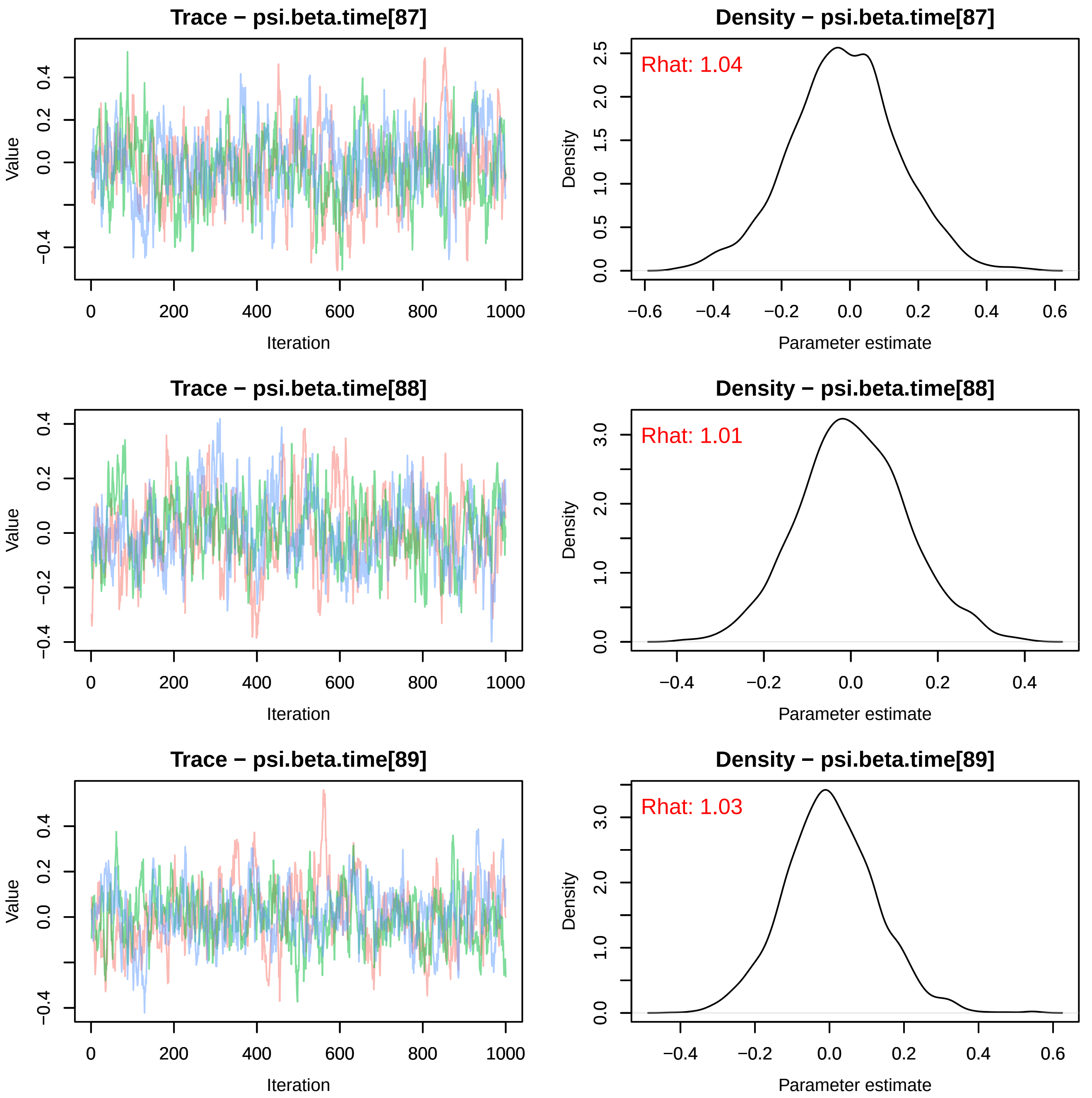

Trace − psi.beta.time[87]
Value
Iteration
Density − psi.beta.time[87]
Rhat: 1.04
Density
Parameter estimate
Trace − psi.beta.time[88]
Value
Iteration
Density − psi.beta.time[88]
Rhat: 1.01
Density
Parameter estimate
Trace − psi.beta.time[89]
Value
Iteration
Density − psi.beta.time[89]
Rhat: 1.03
Density
Parameter estimate

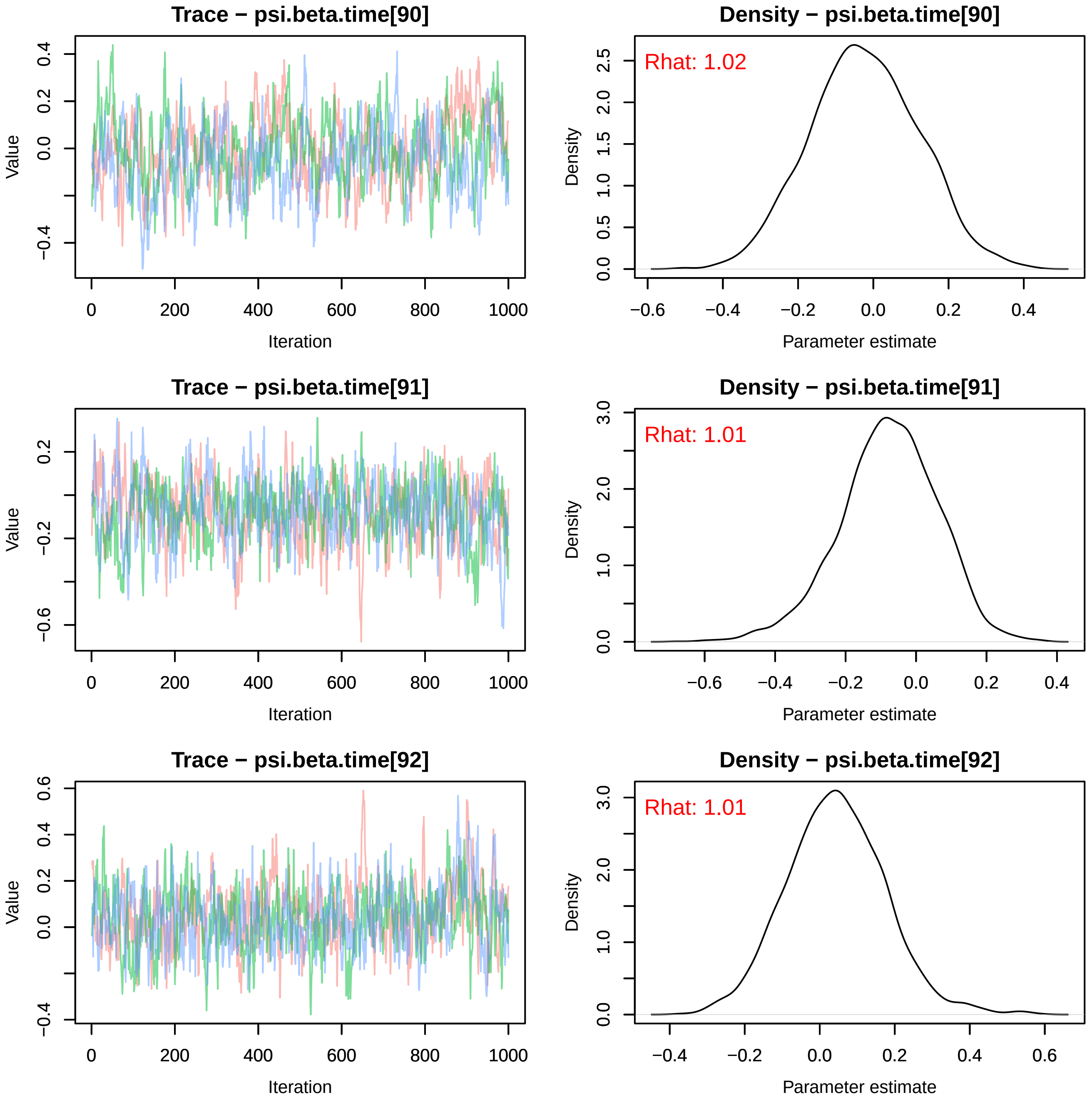
Trace − psi.beta.time[90]
Value
Iteration
Density − psi.beta.time[90]
Rhat: 1.02
Density
Parameter estimate
Trace − psi.beta.time[91]
Value
Iteration
Density − psi.beta.time[91]
Rhat: 1.01
Density
Parameter estimate
Trace − psi.beta.time[92]
Value
Iteration
Density − psi.beta.time[92]
Rhat: 1.01
Density
Parameter estimate

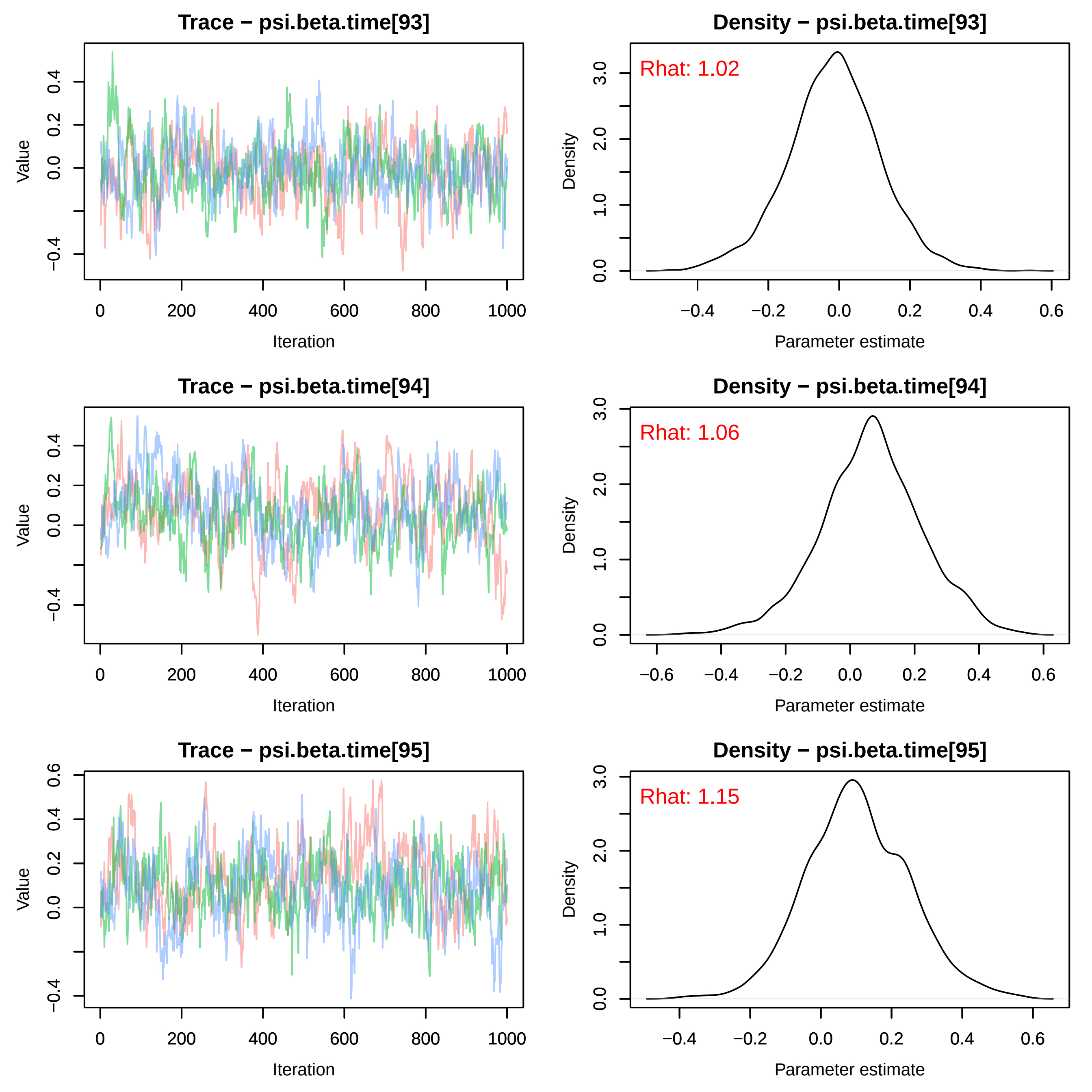
Trace − psi.beta.time[93]
Value
Iteration
Density − psi.beta.time[93]
Rhat: 1.02
Density
Parameter estimate
Trace − psi.beta.time[94]
Value
Iteration
Density − psi.beta.time[94]
Rhat: 1.06
Density
Parameter estimate
Trace − psi.beta.time[95]
Value
Iteration
Density − psi.beta.time[95]
Rhat: 1.15
Density
Parameter estimate

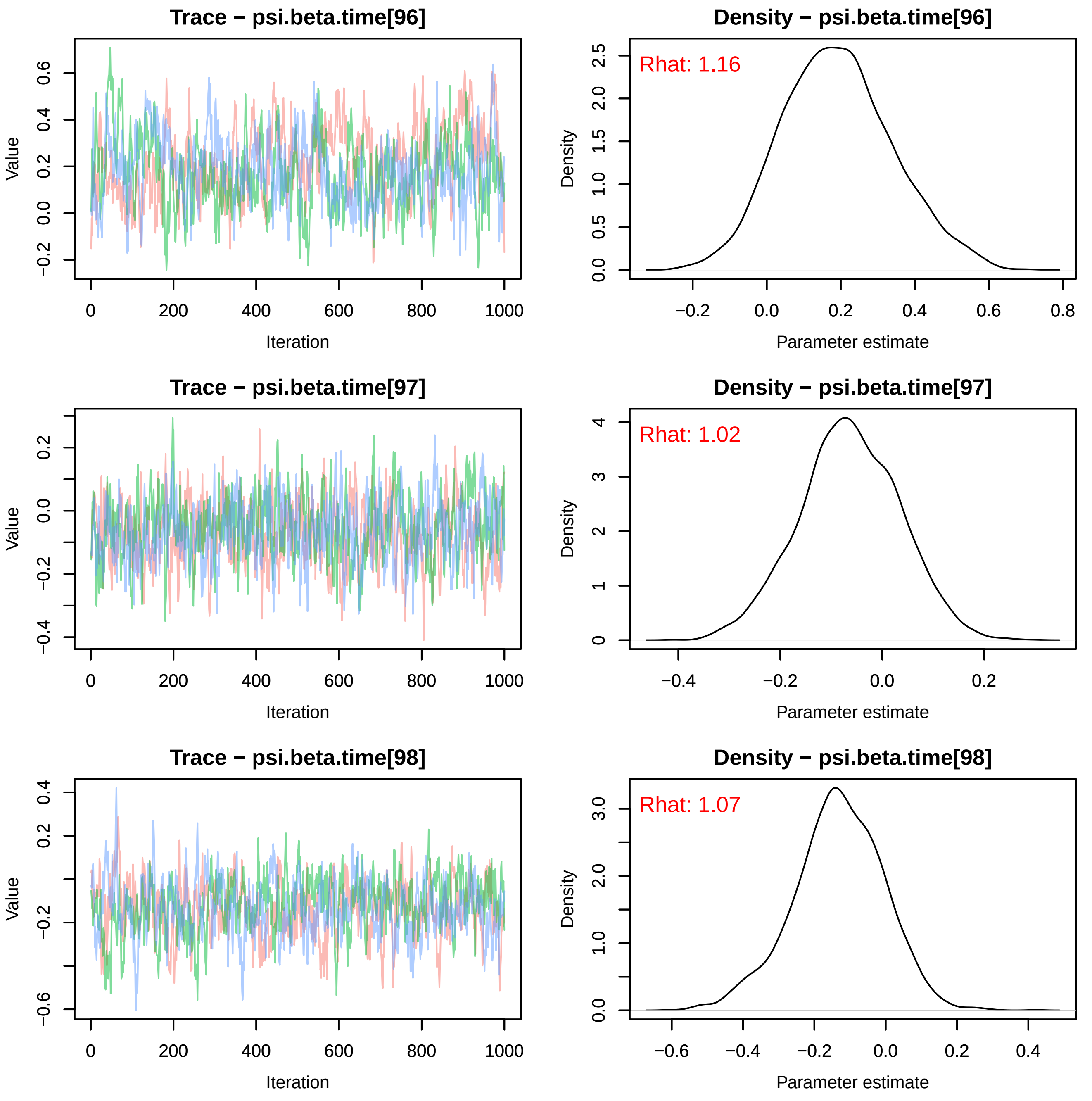
Trace − psi.beta.time[96]
Density − psi.beta.time[96]
Rhat: 1.16
Trace − psi.beta.time[97]
Density − psi.beta.time[97]
Rhat: 1.02
Trace − psi.beta.time[98]
Density − psi.beta.time[98]
Rhat: 1.07
Value
Iteration
Density
Parameter estimate

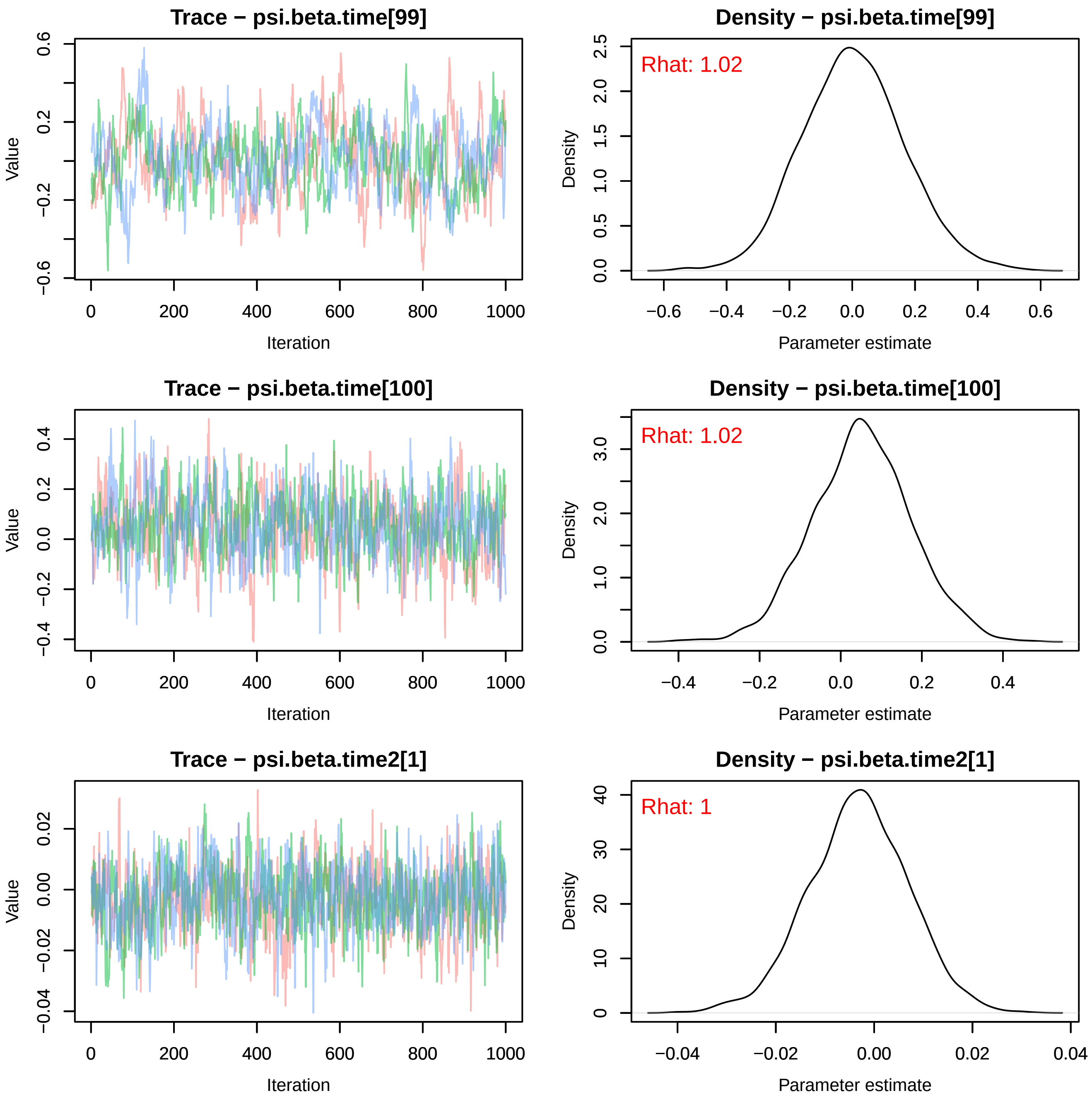
Trace − psi.beta.time[99]
Value
Iteration
Density − psi.beta.time[99]
Rhat: 1.02
Density
Parameter estimate
Trace − psi.beta.time[100]
Value
Iteration
Density − psi.beta.time[100]
Rhat: 1.02
Density
Parameter estimate
Trace − psi.beta.time2[1]
Value
Iteration
Density − psi.beta.time2[1]
Rhat: 1
Density
Parameter estimate

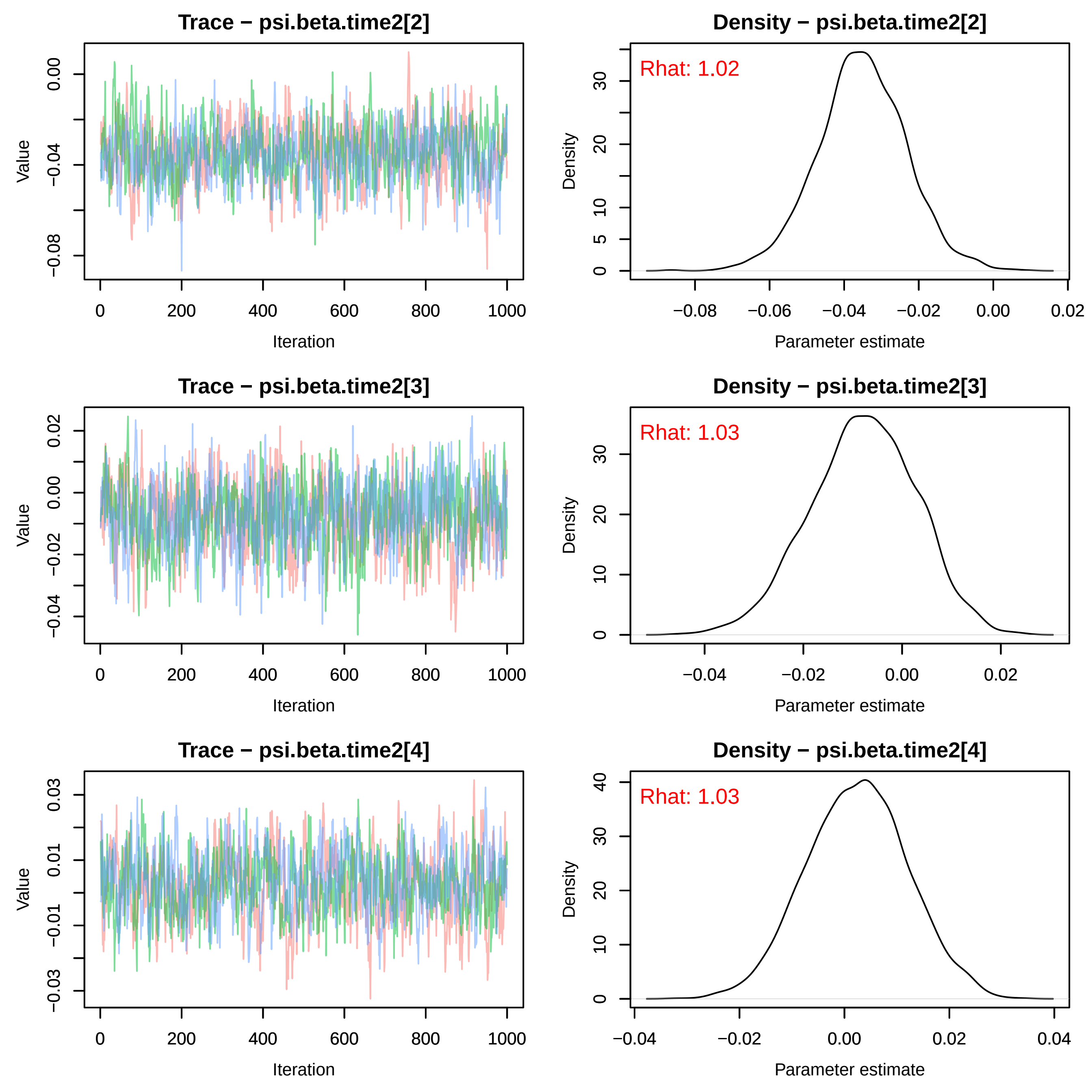
Trace − psi.beta.time2[2]
Value
Iteration
Density − psi.beta.time2[2]
Rhat: 1.02
Density
Parameter estimate
Trace − psi.beta.time2[3]
Value
Iteration
Density − psi.beta.time2[3]
Rhat: 1.03
Density
Parameter estimate
Trace − psi.beta.time2[4]
Value
Iteration
Density − psi.beta.time2[4]
Rhat: 1.03
Density
Parameter estimate

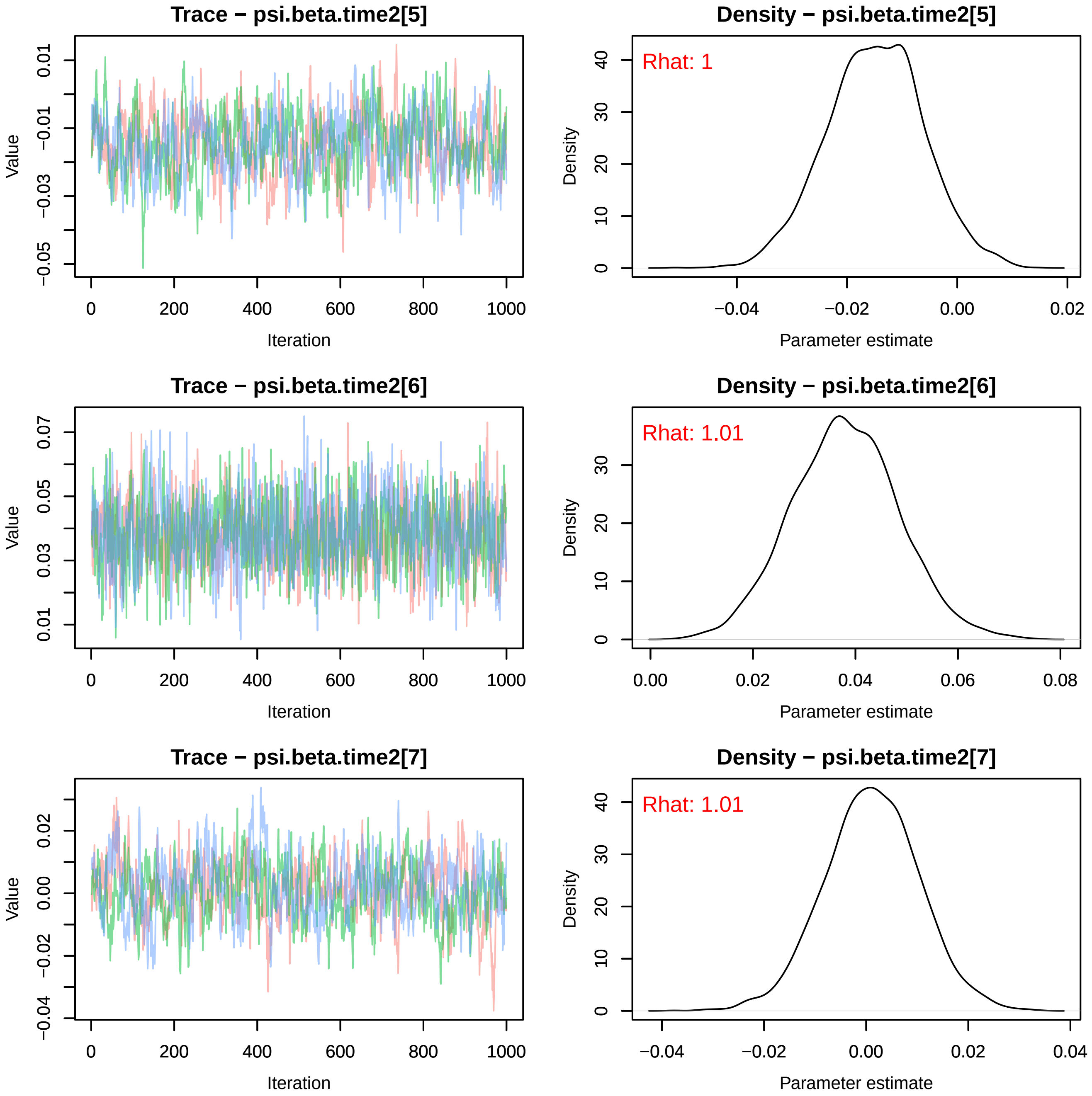

Trace − psi.beta.time2[5]
Value
0.01
−0.01
−0.03
−0.05
0
200
400
600
800
1000
Iteration
Density − psi.beta.time2[5]
Rhat: 1
Density
0
10
20
30
40
−0.04
−0.02
0.00
0.02
Parameter estimate
Trace − psi.beta.time2[6]
Value
0.01
0.03
0.05
0.07
0
200
400
600
800
1000
Iteration
Density − psi.beta.time2[6]
Rhat: 1.01
Density
0
10
20
30
0.00
0.02
0.04
0.06
0.08
Parameter estimate
Trace − psi.beta.time2[7]
Value
−0.04
−0.02
0.00
0.02
0
200
400
600
800
1000
Iteration
Density − psi.beta.time2[7]
Rhat: 1.01
Density
0
10
20
30
40
−0.04
−0.02
0.00
0.02
0.04
Parameter estimate

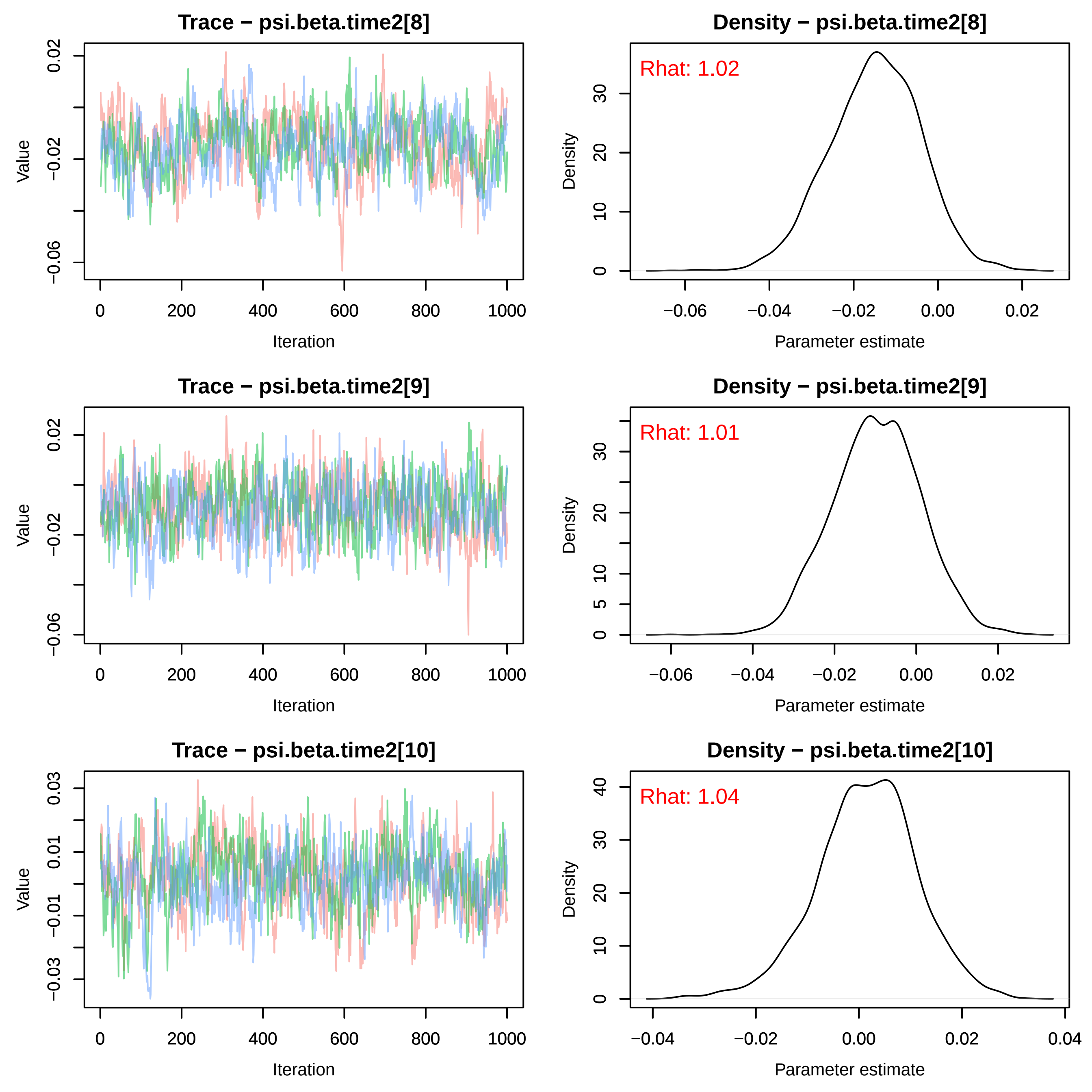

Trace − psi.beta.time2[8]
Density − psi.beta.time2[8]
Rhat: 1.02
Trace − psi.beta.time2[9]
Density − psi.beta.time2[9]
Rhat: 1.01
Trace − psi.beta.time2[10]
Density − psi.beta.time2[10]
Rhat: 1.04
Value
Iteration
Density
Parameter estimate

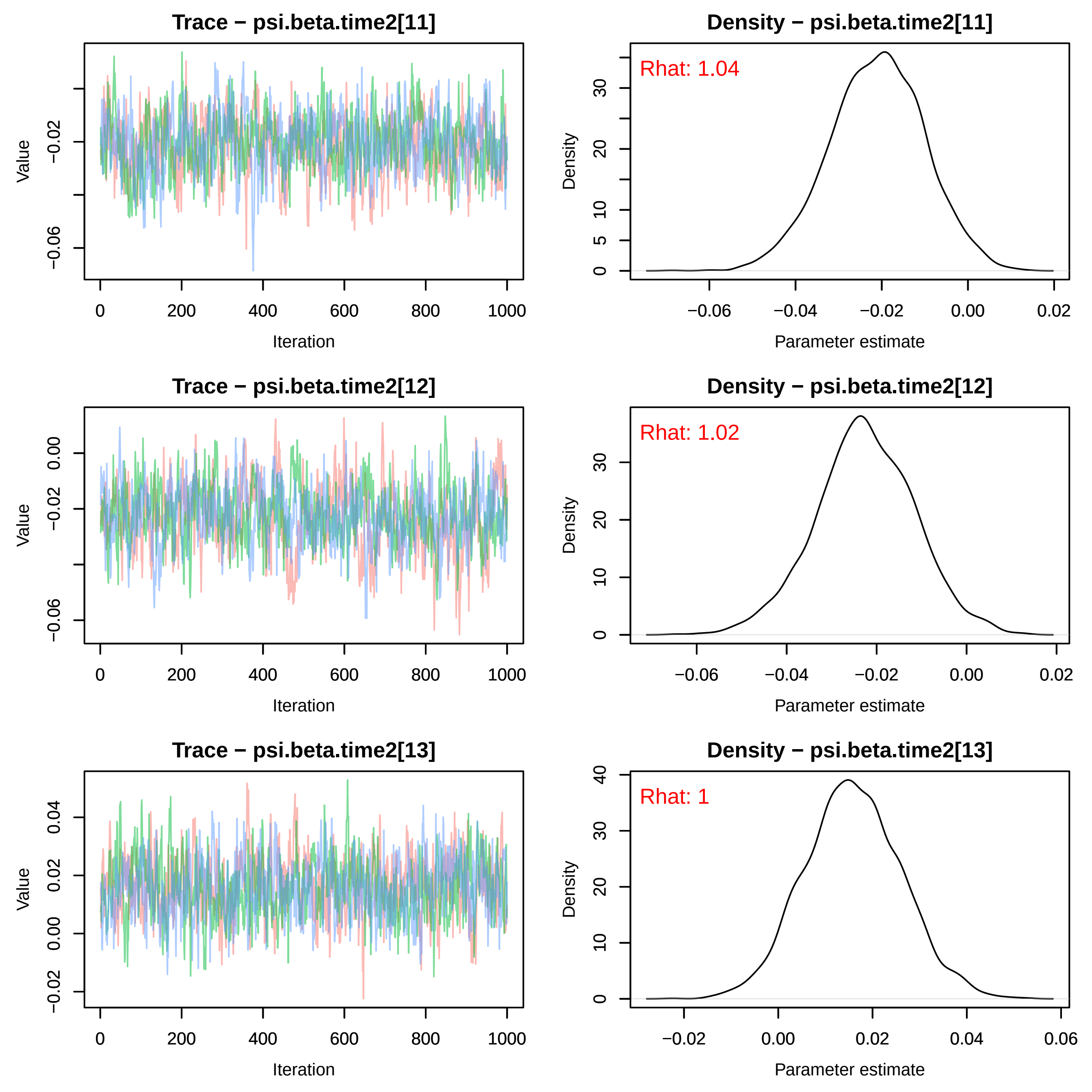

Trace − psi.beta.time2[11]
Value
−0.06
−0.02
0
200
400
600
800
1000
Iteration
Density − psi.beta.time2[11]
Rhat: 1.04
Density
0
5
10
20
30
−0.06
−0.04
−0.02
0.00
0.02
Parameter estimate
Trace − psi.beta.time2[12]
Value
−0.06
−0.02
0.00
0
200
400
600
800
1000
Iteration
Density − psi.beta.time2[12]
Rhat: 1.02
Density
0
10
20
30
−0.06
−0.04
−0.02
0.00
0.02
Parameter estimate
Trace − psi.beta.time2[13]
Value
−0.02
0.00
0.02
0.04
0
200
400
600
800
1000
Iteration
Density − psi.beta.time2[13]
Rhat: 1
Density
0
10
20
30
40
−0.02
0.00
0.02
0.04
0.06
Parameter estimate

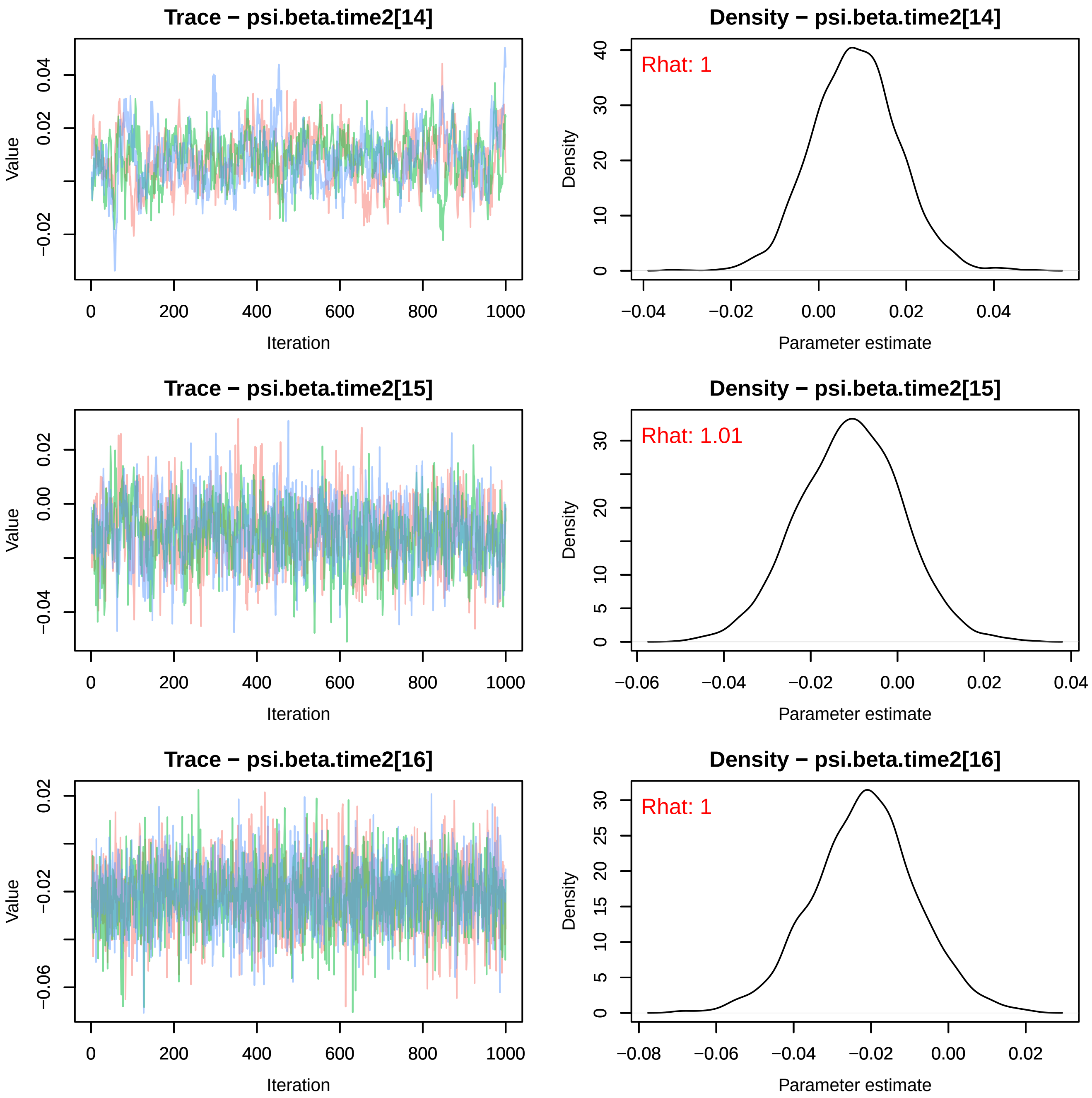
Trace − psi.beta.time2[14]
Value
Iteration
Density − psi.beta.time2[14]
Rhat: 1
Density
Parameter estimate
Trace − psi.beta.time2[15]
Value
Iteration
Density − psi.beta.time2[15]
Rhat: 1.01
Density
Parameter estimate
Trace − psi.beta.time2[16]
Value
Iteration
Density − psi.beta.time2[16]
Rhat: 1
Density
Parameter estimate

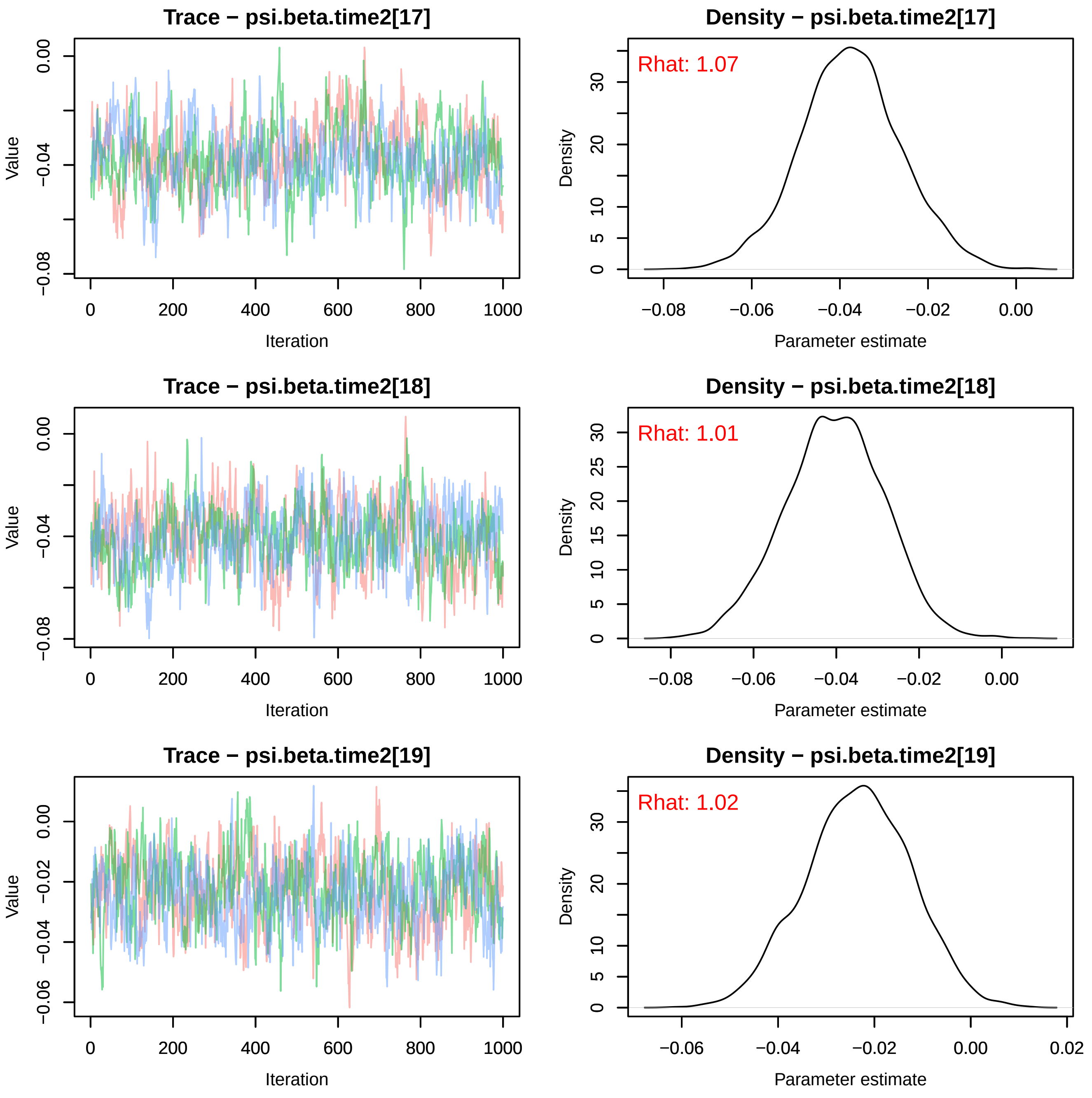

Trace − psi.beta.time2[17]
Value
0.00
−0.04
−0.08
0
200
400
600
800
1000
Iteration
Density − psi.beta.time2[17]
Rhat: 1.07
Density
0
5
10
20
30
−0.08
−0.06
−0.04
−0.02
0.00
Parameter estimate
Trace − psi.beta.time2[18]
Value
0.00
−0.04
−0.08
0
200
400
600
800
1000
Iteration
Density − psi.beta.time2[18]
Rhat: 1.01
Density
0
5
10
15
20
25
30
−0.08
−0.06
−0.04
−0.02
0.00
Parameter estimate
Trace − psi.beta.time2[19]
Value
0.00
−0.02
−0.04
−0.06
0
200
400
600
800
1000
Iteration
Density − psi.beta.time2[19]
Rhat: 1.02
Density
0
5
10
20
30
−0.06
−0.04
−0.02
0.00
0.02
Parameter estimate

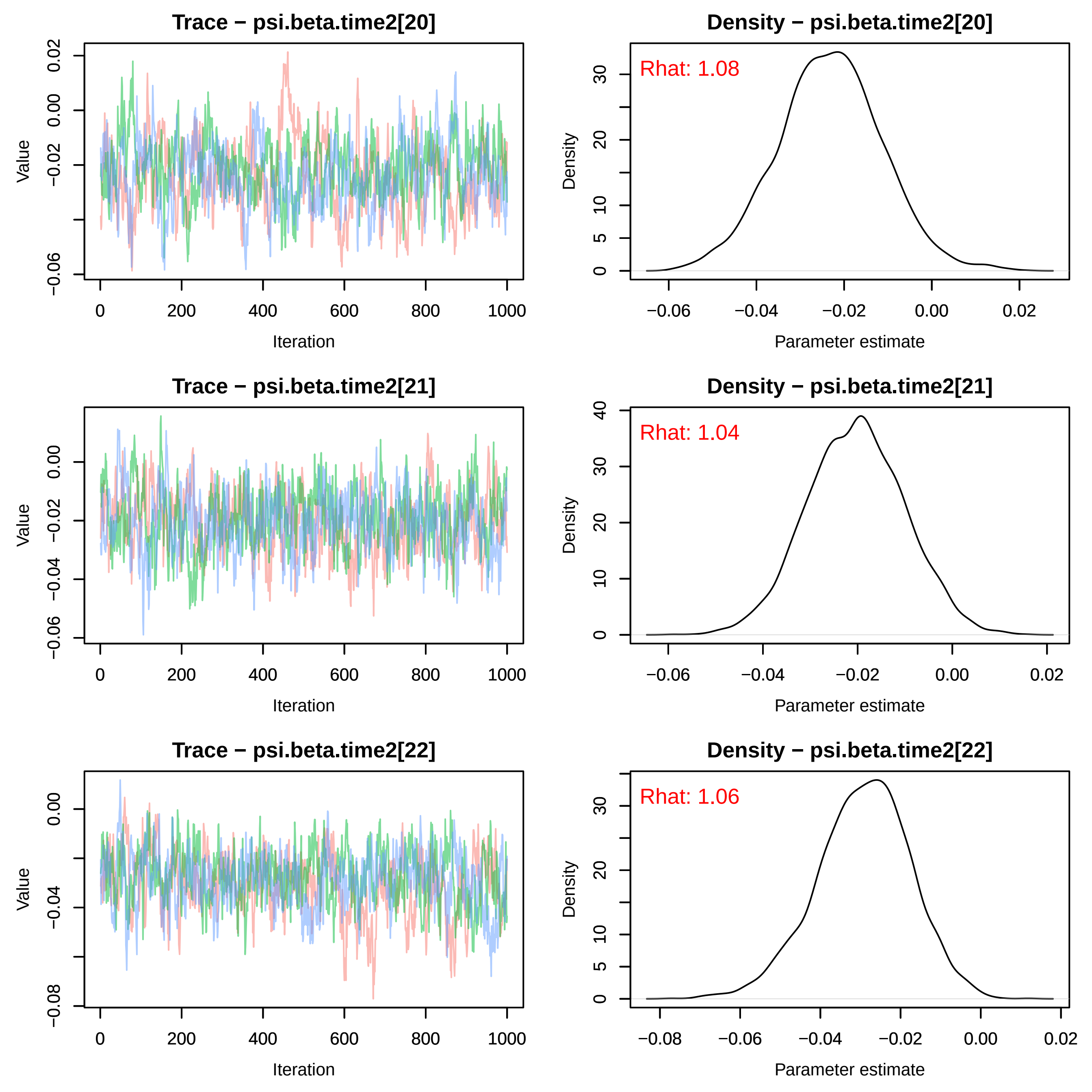
Trace − psi.beta.time2[20]
Value
Iteration
0
200
400
600
800
1000
−0.06
−0.02
0.00
0.02
Density − psi.beta.time2[20]
Rhat: 1.08
Density
Parameter estimate
−0.06
−0.04
−0.02
0.00
0.02
0
5
10
20
30
Trace − psi.beta.time2[21]
Value
Iteration
−0.06
−0.04
−0.02
0.00
Density − psi.beta.time2[21]
Rhat: 1.04
Density
Parameter estimate
0
10
20
30
40
Trace − psi.beta.time2[22]
Value
Iteration
−0.08
−0.04
0.00
Density − psi.beta.time2[22]
Rhat: 1.06
Density
Parameter estimate
−0.08
−0.06
−0.04
−0.02
0.00
0.02

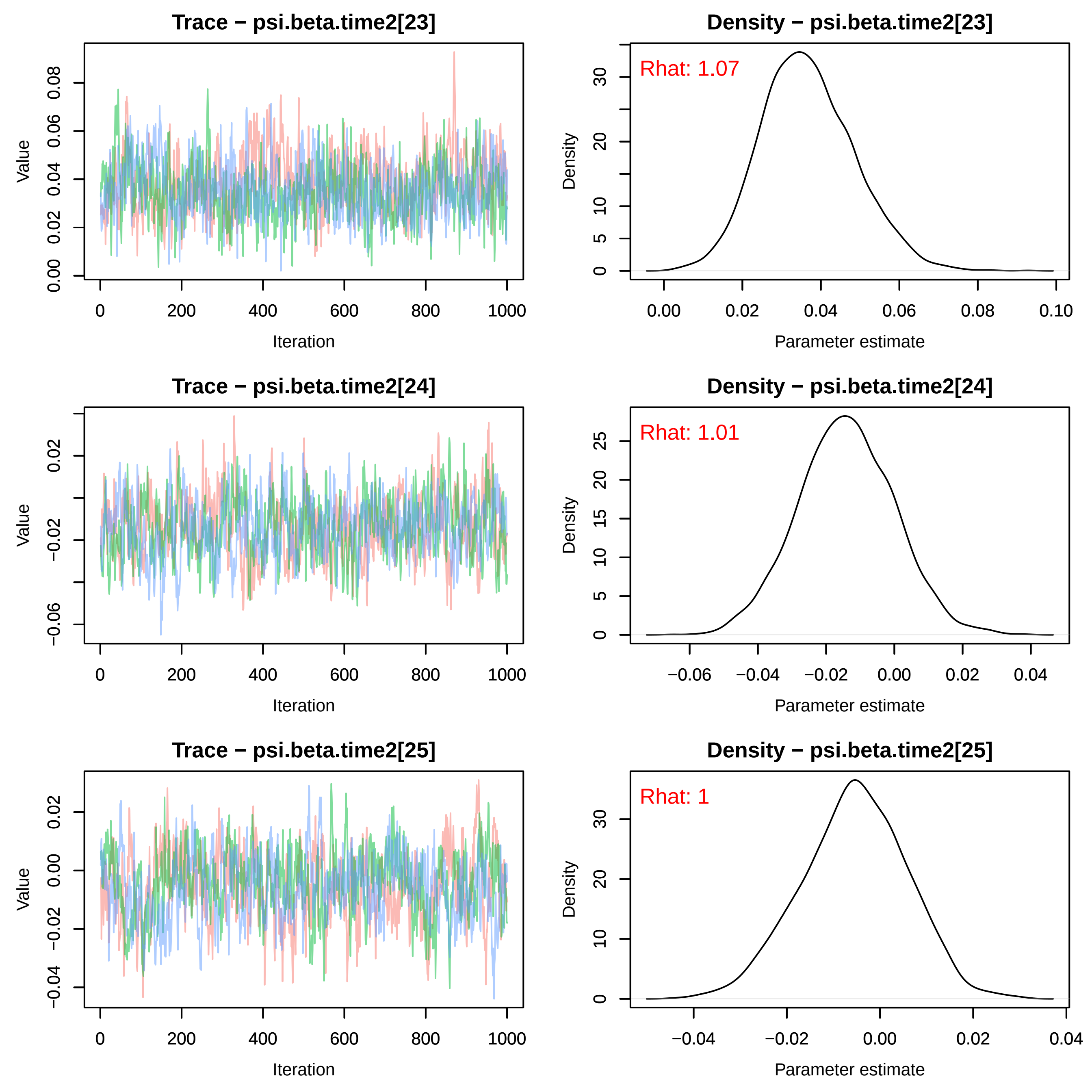

Trace − psi.beta.time2[23]
Value
Iteration
Density − psi.beta.time2[23]
Rhat: 1.07
Density
Parameter estimate
Trace − psi.beta.time2[24]
Value
Iteration
Density − psi.beta.time2[24]
Rhat: 1.01
Density
Parameter estimate
Trace − psi.beta.time2[25]
Value
Iteration
Density − psi.beta.time2[25]
Rhat: 1
Density
Parameter estimate

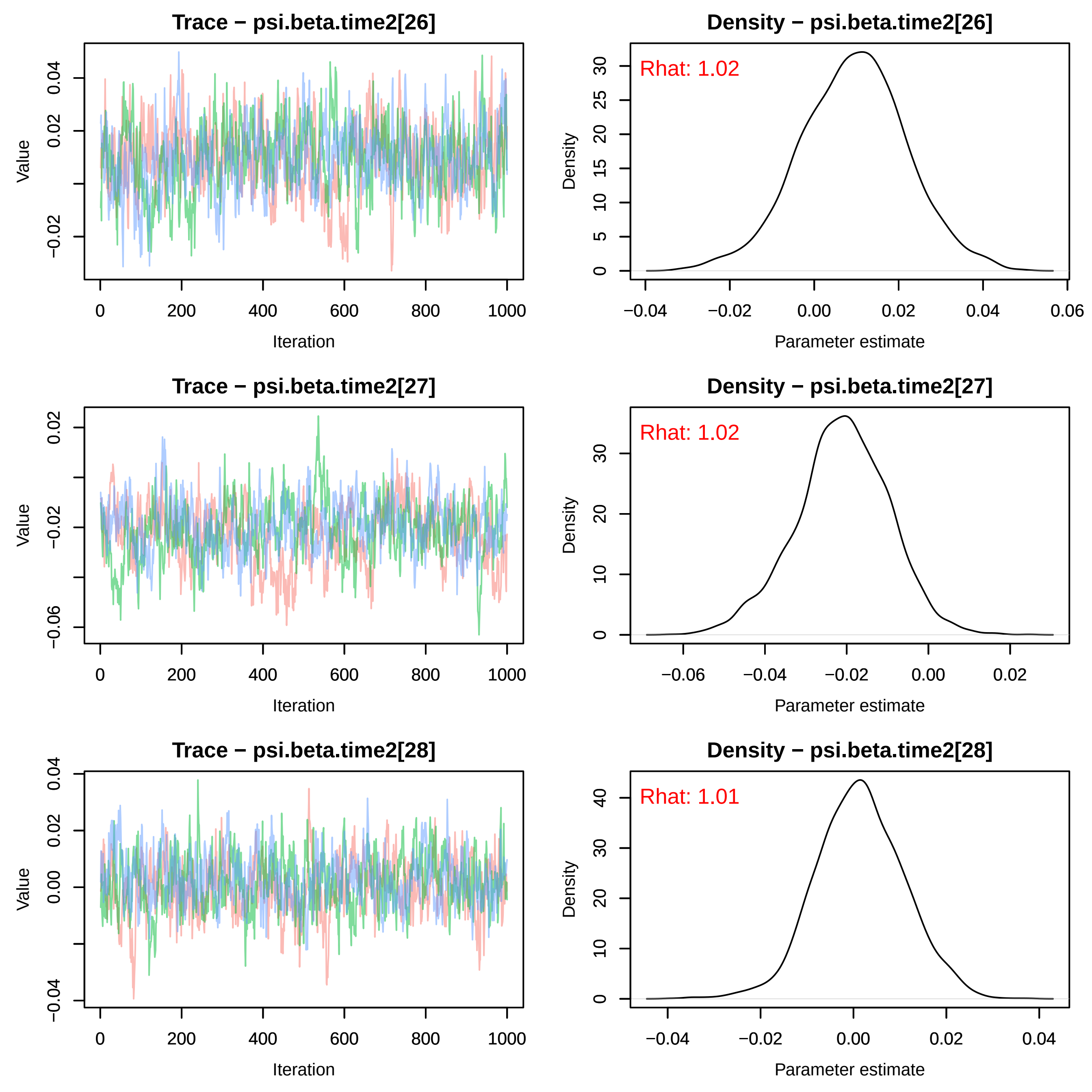

Trace − psi.beta.time2[26]
Value
Iteration
Density − psi.beta.time2[26]
Rhat: 1.02
Density
Parameter estimate
Trace − psi.beta.time2[27]
Value
Iteration
Density − psi.beta.time2[27]
Rhat: 1.02
Density
Parameter estimate
Trace − psi.beta.time2[28]
Value
Iteration
Density − psi.beta.time2[28]
Rhat: 1.01
Density
Parameter estimate

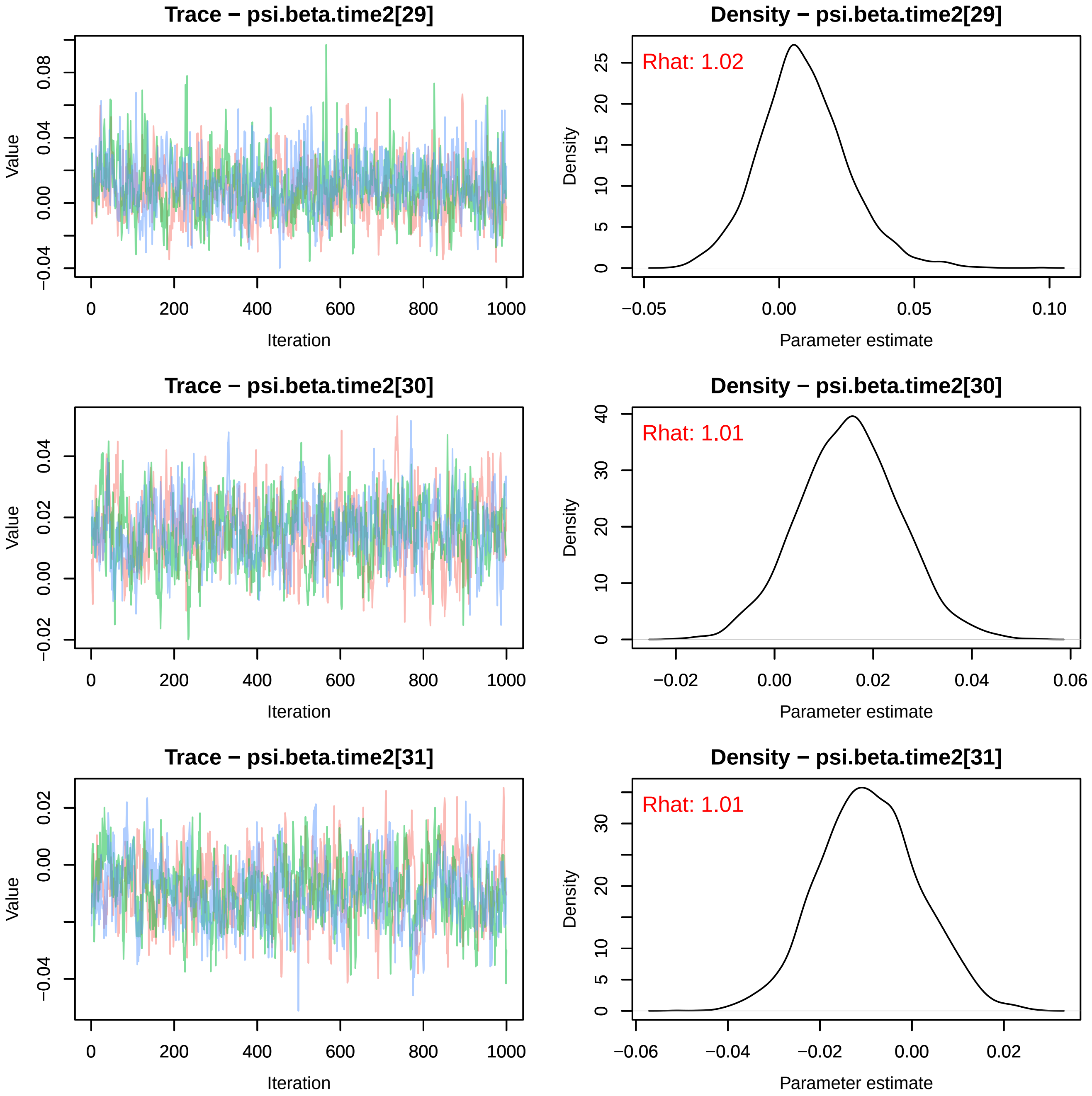

Trace − psi.beta.time2[29]
Value
Iteration
Density − psi.beta.time2[29]
Rhat: 1.02
Density
Parameter estimate
Trace − psi.beta.time2[30]
Value
Iteration
Density − psi.beta.time2[30]
Rhat: 1.01
Density
Parameter estimate
Trace − psi.beta.time2[31]
Value
Iteration
Density − psi.beta.time2[31]
Rhat: 1.01
Density
Parameter estimate

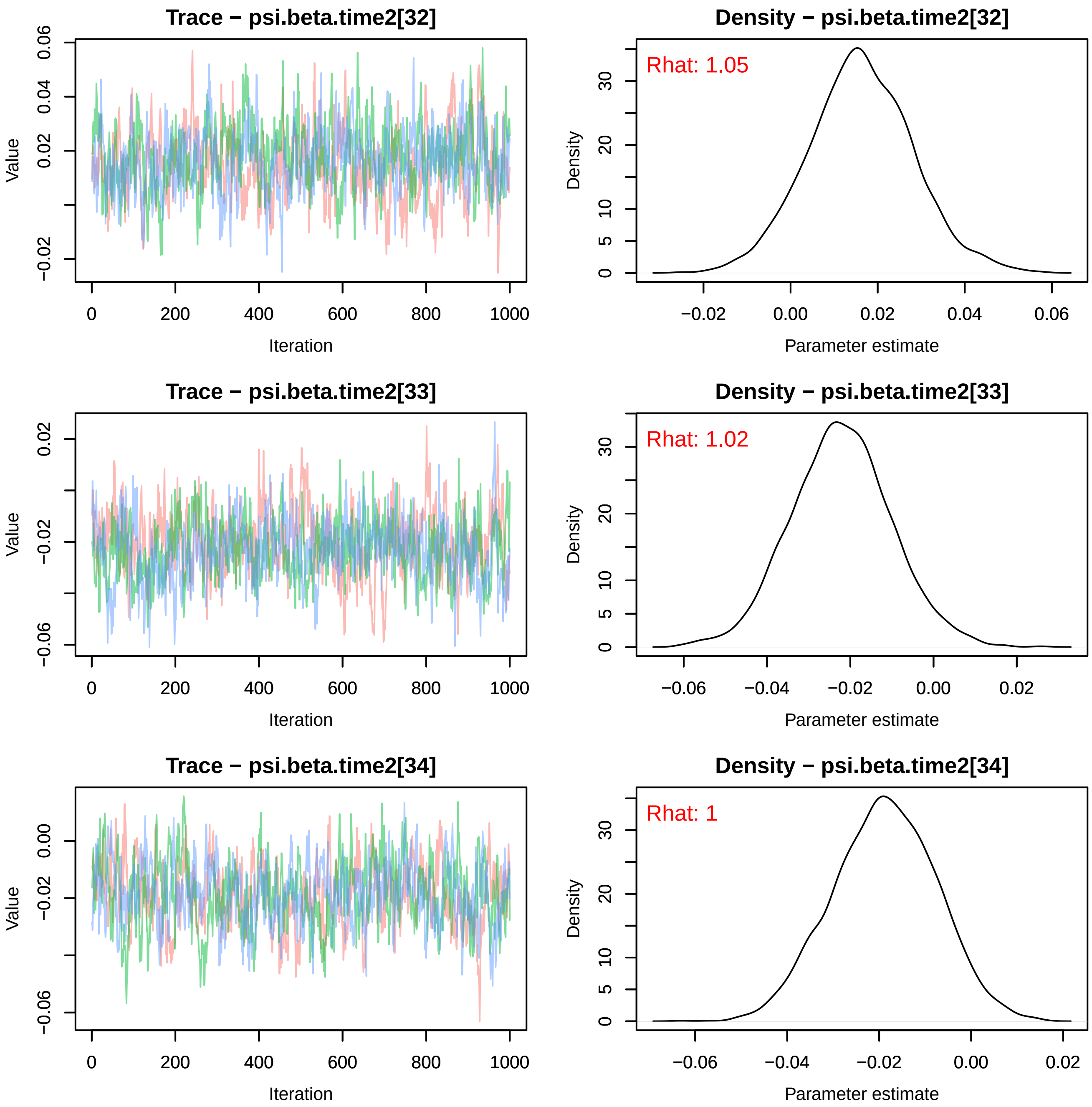
Trace − psi.beta.time2[32]
Value
−0.02
0.02
0.04
0.06
0
200
400
600
800
1000
Iteration
Density − psi.beta.time2[32]
Rhat: 1.05
Density
0
5
10
20
30
−0.02
0.00
0.02
0.04
0.06
Parameter estimate
Trace − psi.beta.time2[33]
Value
−0.06
−0.02
0.02
0
200
400
600
800
1000
Iteration
Density − psi.beta.time2[33]
Rhat: 1.02
Density
0
5
10
20
30
−0.06
−0.04
−0.02
0.00
0.02
Parameter estimate
Trace − psi.beta.time2[34]
Value
−0.06
−0.02
0.00
0
200
400
600
800
1000
Iteration
Density − psi.beta.time2[34]
Rhat: 1
Density
0
5
10
20
30
−0.06
−0.04
−0.02
0.00
0.02
Parameter estimate

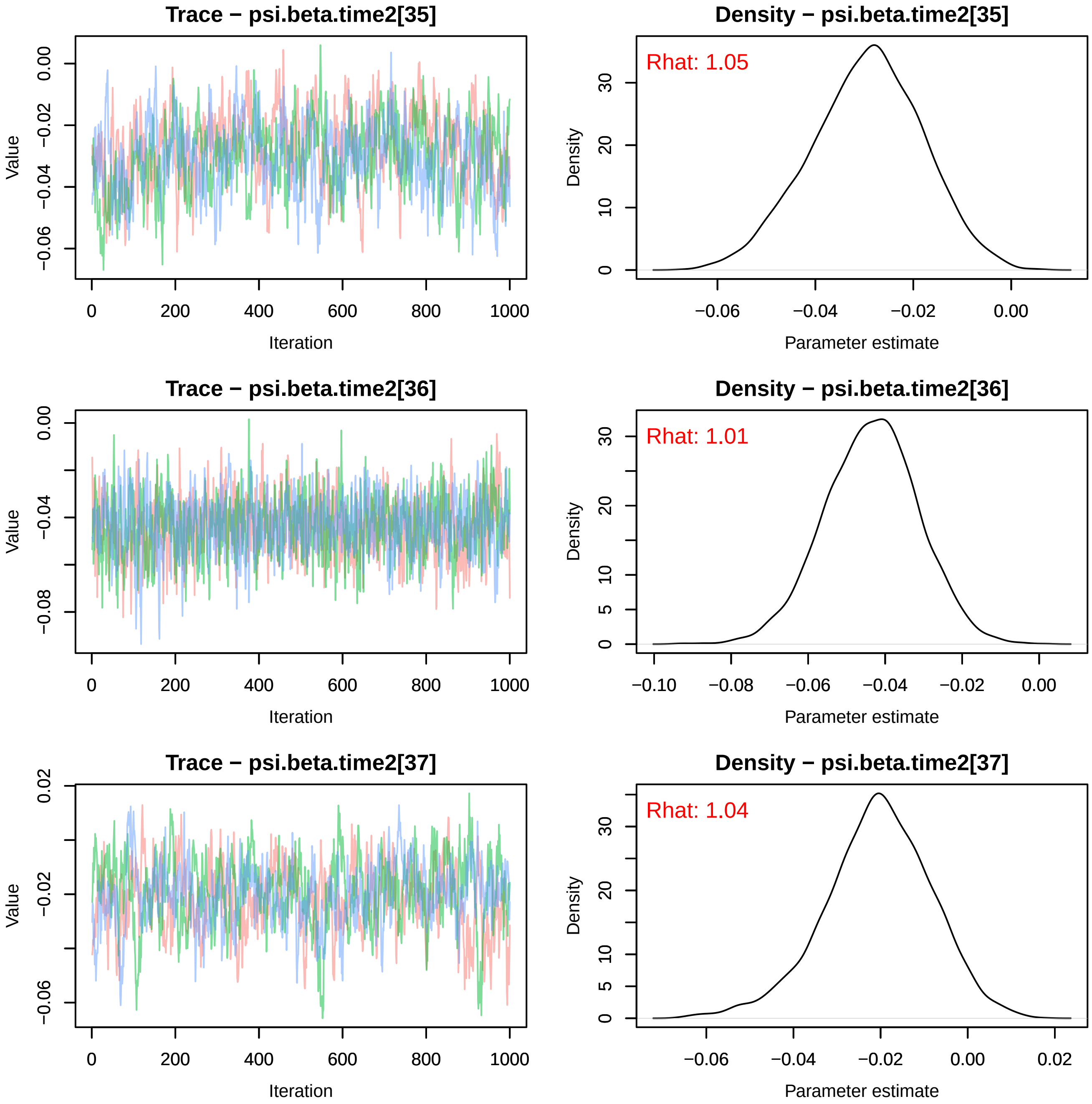
Trace − psi.beta.time2[35]
Value
Iteration
0.00
−0.02
−0.04
−0.06
0
200
400
600
800
1000
Density − psi.beta.time2[35]
Rhat: 1.05
Density
Parameter estimate
0
10
20
30
−0.06
−0.04
−0.02
0.00
Trace − psi.beta.time2[36]
Value
Iteration
0.00
−0.04
−0.08
0
200
400
600
800
1000
Density − psi.beta.time2[36]
Rhat: 1.01
Density
Parameter estimate
0
5
10
20
30
−0.10
−0.08
−0.06
−0.04
−0.02
0.00
Trace − psi.beta.time2[37]
Value
Iteration
0.02
−0.02
−0.06
0
200
400
600
800
1000
Density − psi.beta.time2[37]
Rhat: 1.04
Density
Parameter estimate
0
5
10
20
30
−0.06
−0.04
−0.02
0.00
0.02

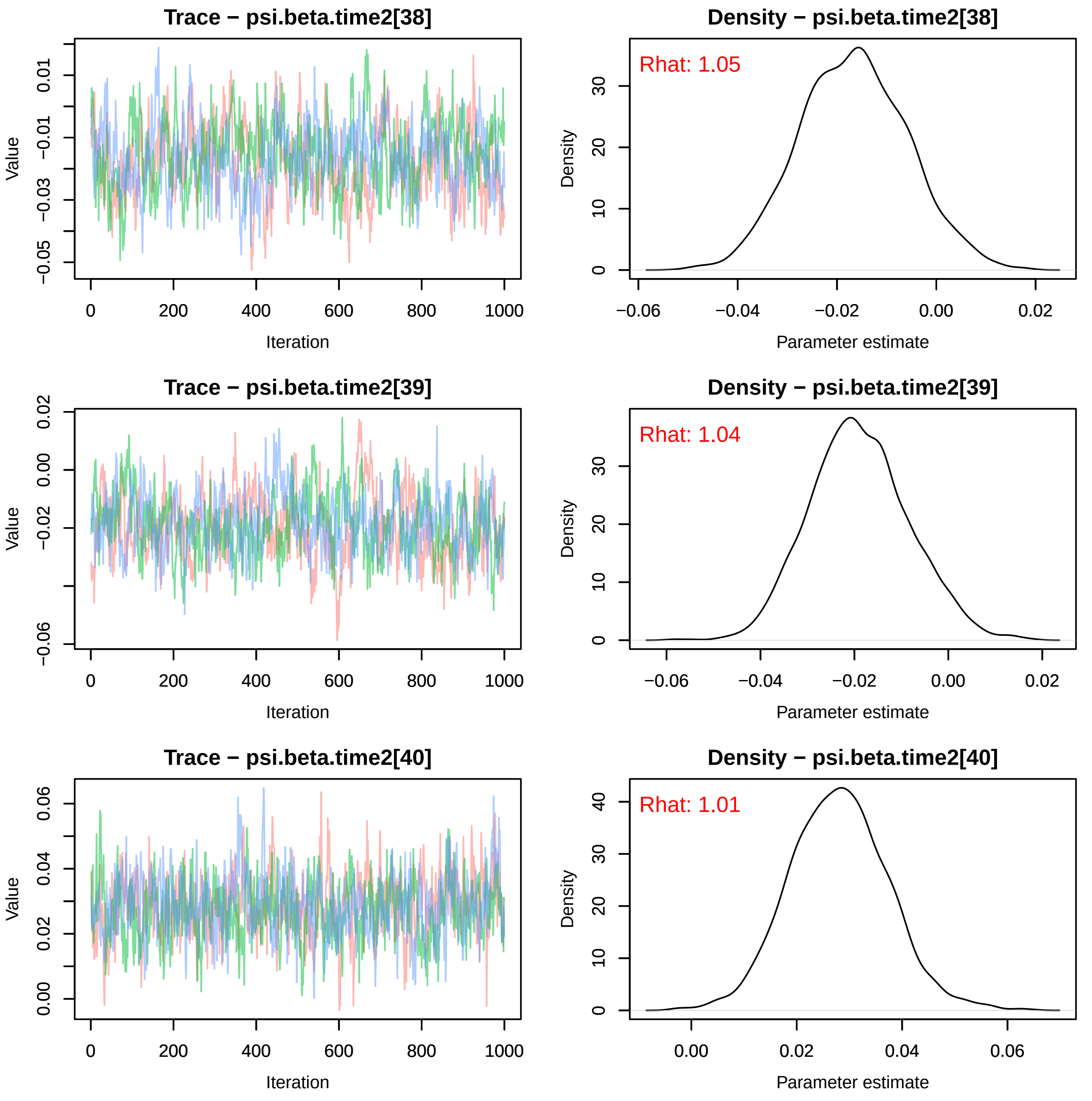
Trace − psi.beta.time2[38]
Value
Iteration
Density − psi.beta.time2[38]
Rhat: 1.05
Density
Parameter estimate
Trace − psi.beta.time2[39]
Value
Iteration
Density − psi.beta.time2[39]
Rhat: 1.04
Density
Parameter estimate
Trace − psi.beta.time2[40]
Value
Iteration
Density − psi.beta.time2[40]
Rhat: 1.01
Density
Parameter estimate

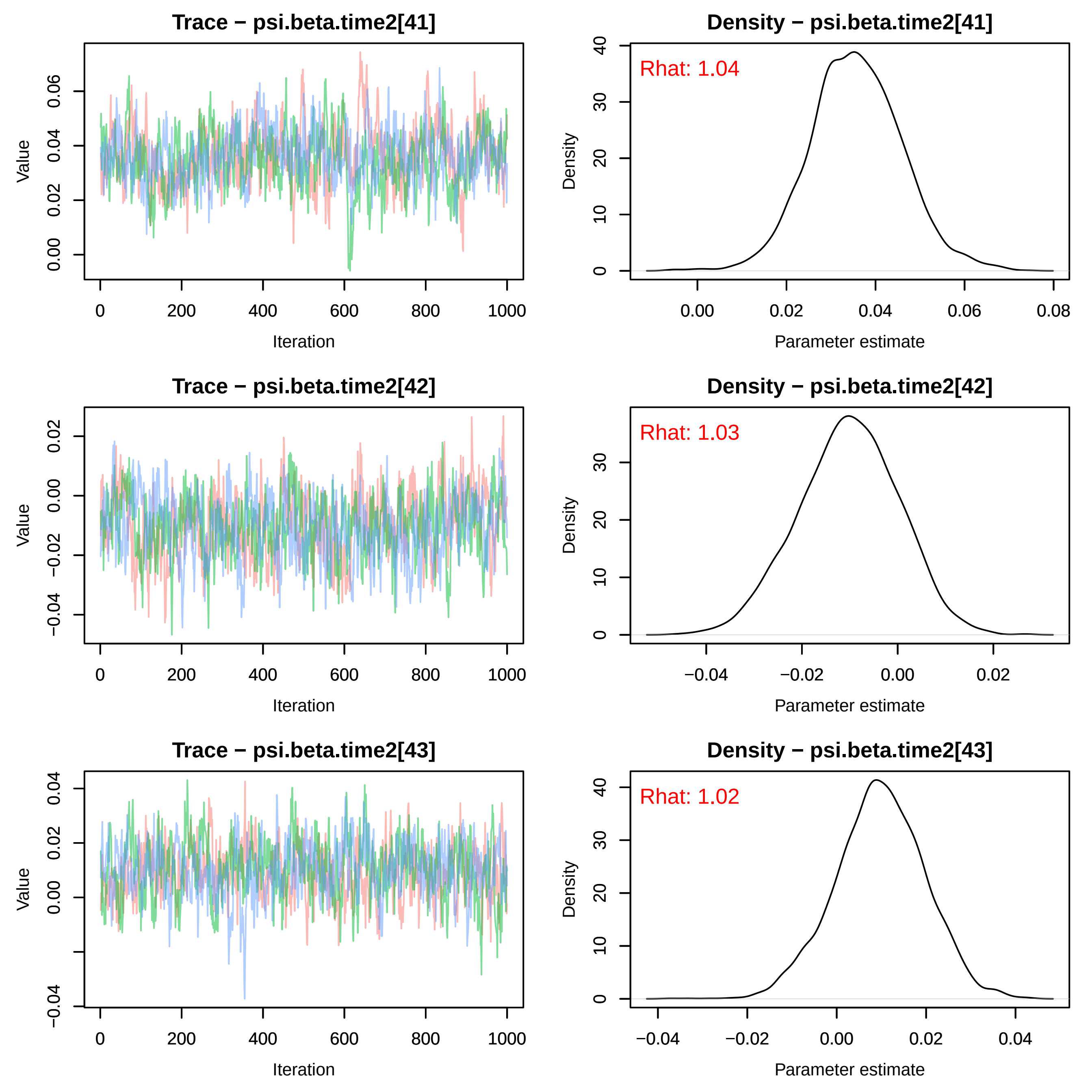
Trace − psi.beta.time2[41]
Value
Iteration
Density − psi.beta.time2[41]
Rhat: 1.04
Density
Parameter estimate
Trace − psi.beta.time2[42]
Value
Iteration
Density − psi.beta.time2[42]
Rhat: 1.03
Density
Parameter estimate
Trace − psi.beta.time2[43]
Value
Iteration
Density − psi.beta.time2[43]
Rhat: 1.02
Density
Parameter estimate

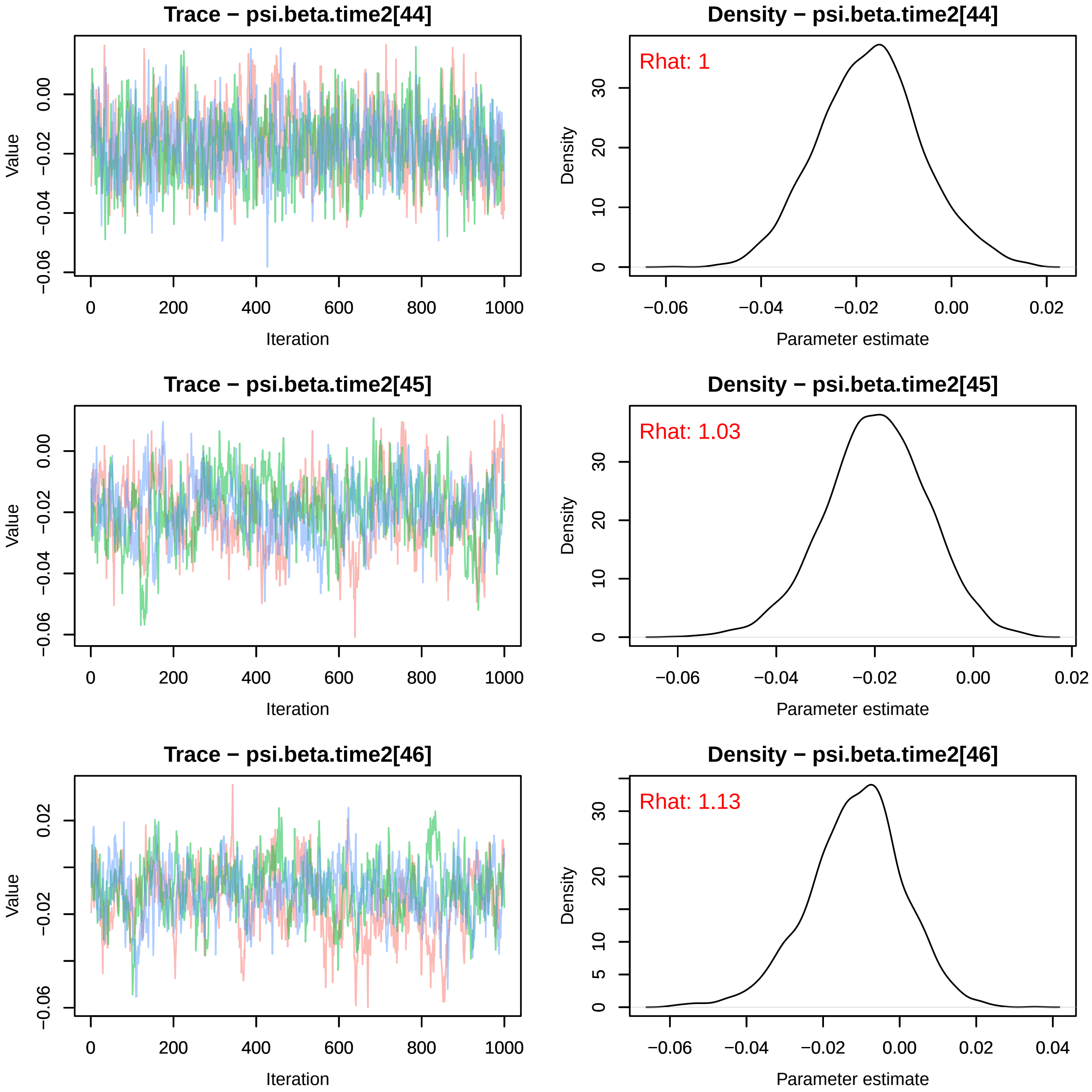
Trace − psi.beta.time2[44]
Density − psi.beta.time2[44]
Rhat: 1
Trace − psi.beta.time2[45]
Density − psi.beta.time2[45]
Rhat: 1.03
Trace − psi.beta.time2[46]
Density − psi.beta.time2[46]
Rhat: 1.13
Value
Iteration
Density
Parameter estimate
0
200
400
600
800
1000
−0.06
−0.04
−0.02
0.00
0.02
0.04
0
5
10
20
30

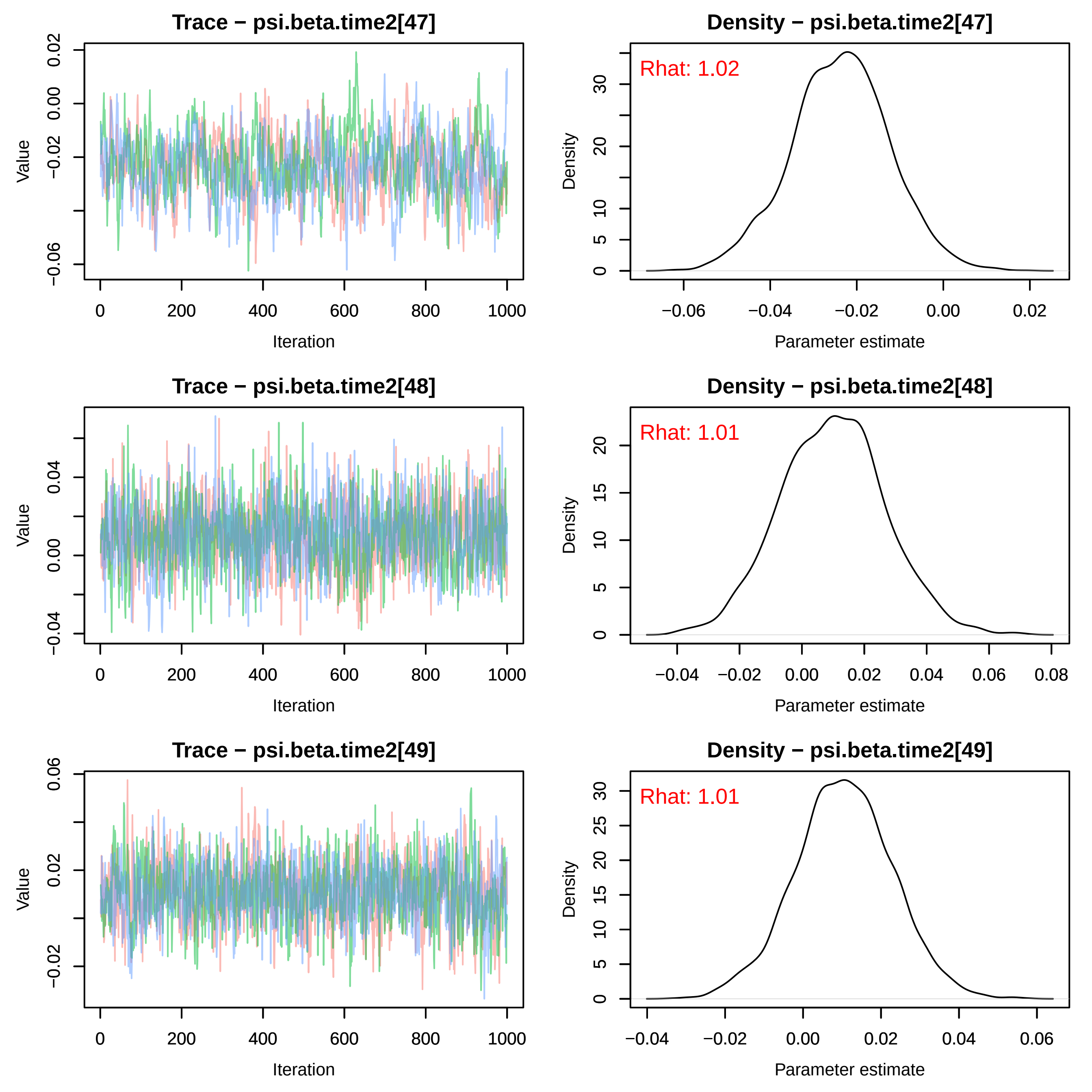
Trace − psi.beta.time2[47]
Value
Iteration
Density − psi.beta.time2[47]
Rhat: 1.02
Density
Parameter estimate
Trace − psi.beta.time2[48]
Value
Iteration
Density − psi.beta.time2[48]
Rhat: 1.01
Density
Parameter estimate
Trace − psi.beta.time2[49]
Value
Iteration
Density − psi.beta.time2[49]
Rhat: 1.01
Density
Parameter estimate

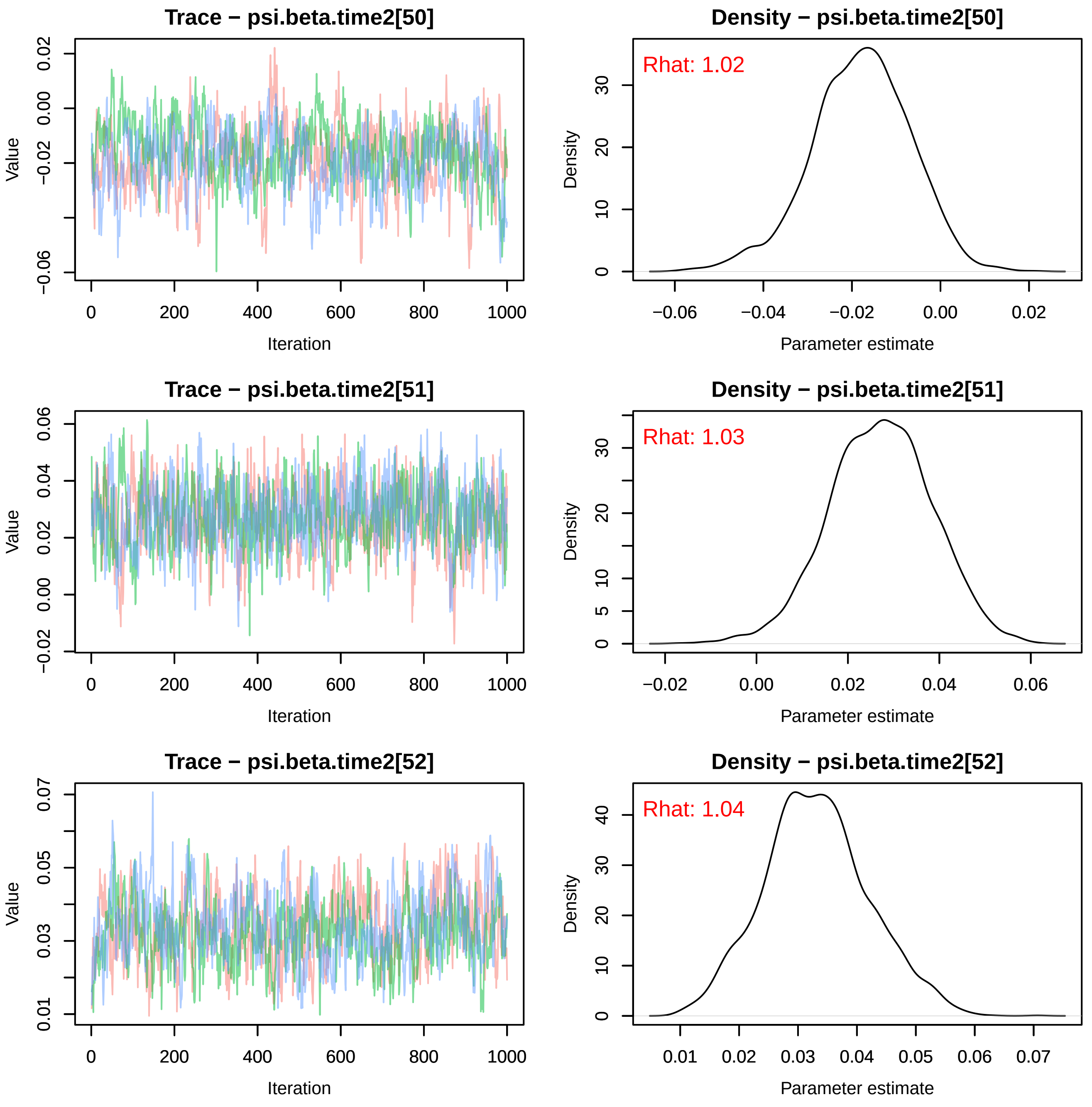
Trace − psi.beta.time2[50]
Value
Iteration
Density − psi.beta.time2[50]
Rhat: 1.02
Density
Parameter estimate
Trace − psi.beta.time2[51]
Value
Iteration
Density − psi.beta.time2[51]
Rhat: 1.03
Density
Parameter estimate
Trace − psi.beta.time2[52]
Value
Iteration
Density − psi.beta.time2[52]
Rhat: 1.04
Density
Parameter estimate

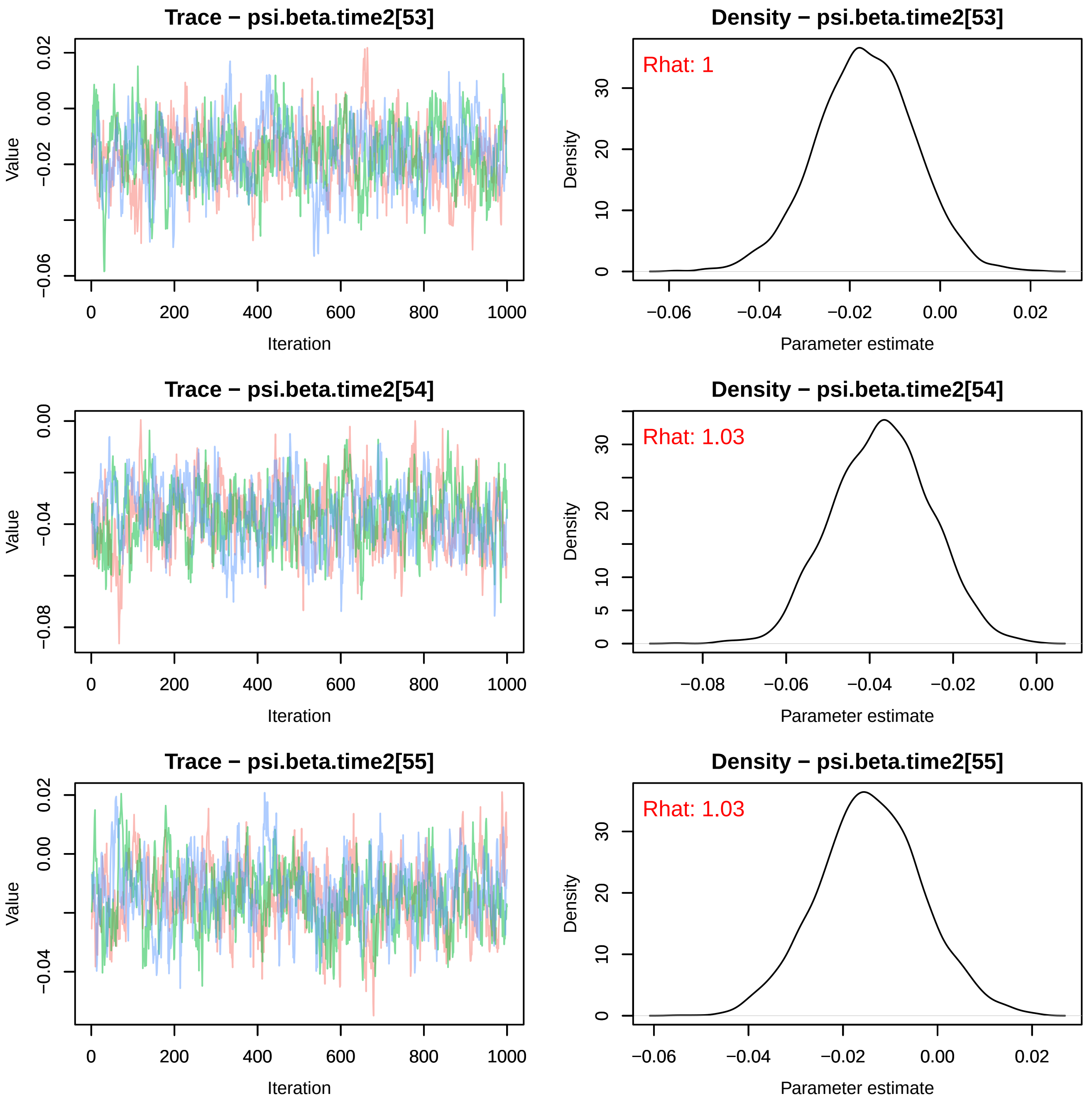

Trace − psi.beta.time2[53]
Density − psi.beta.time2[53]
Rhat: 1
Trace − psi.beta.time2[54]
Density − psi.beta.time2[54]
Rhat: 1.03
Trace − psi.beta.time2[55]
Density − psi.beta.time2[55]
Rhat: 1.03
Value
Iteration
Density
Parameter estimate
0
200
400
600
800
1000
−0.06
−0.04
−0.02
0.00
0.02
−0.08
0
5
10
20
30

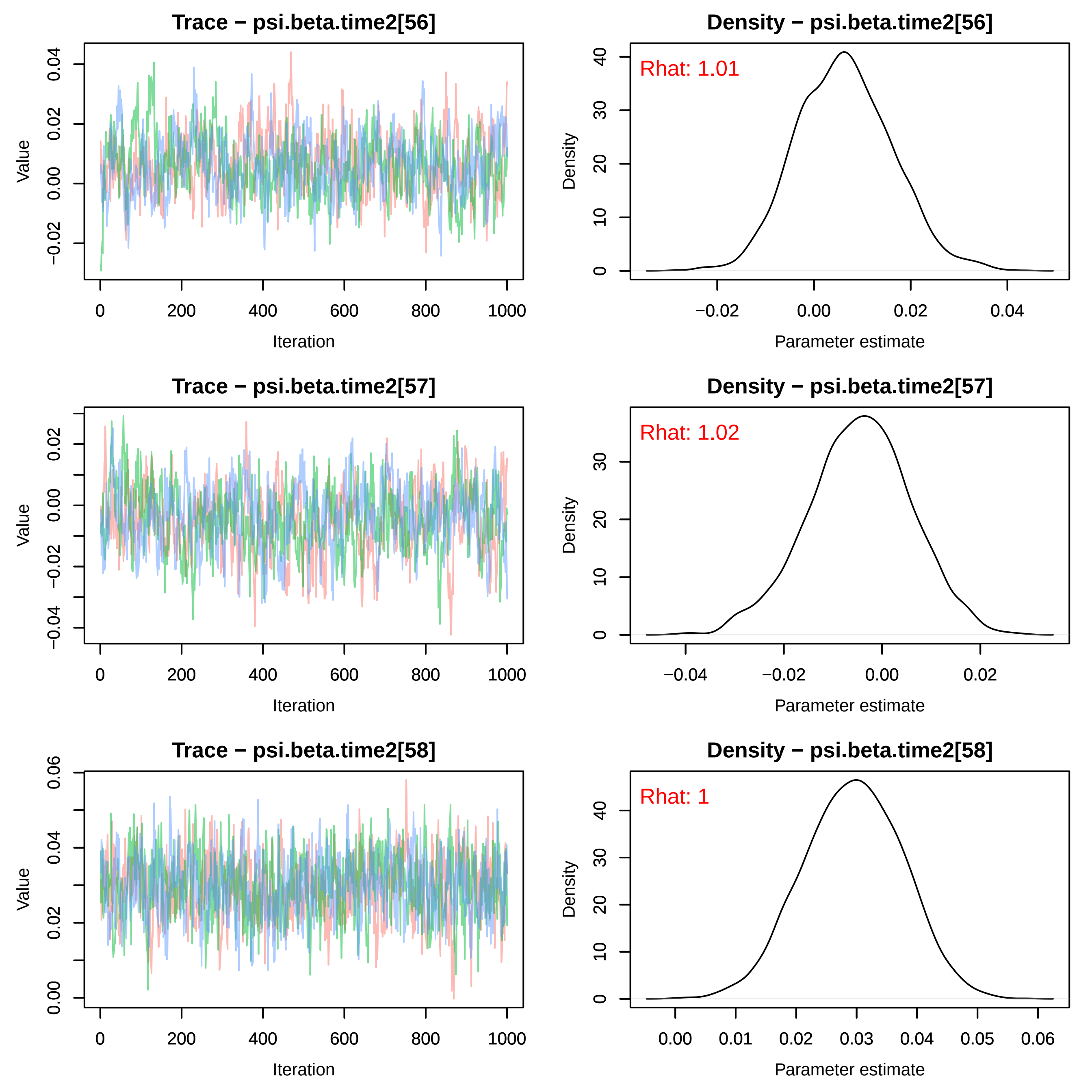

Trace − psi.beta.time2[56]
Value
Iteration
Density − psi.beta.time2[56]
Rhat: 1.01
Density
Parameter estimate
Trace − psi.beta.time2[57]
Value
Iteration
Density − psi.beta.time2[57]
Rhat: 1.02
Density
Parameter estimate
Trace − psi.beta.time2[58]
Value
Iteration
Density − psi.beta.time2[58]
Rhat: 1
Density
Parameter estimate

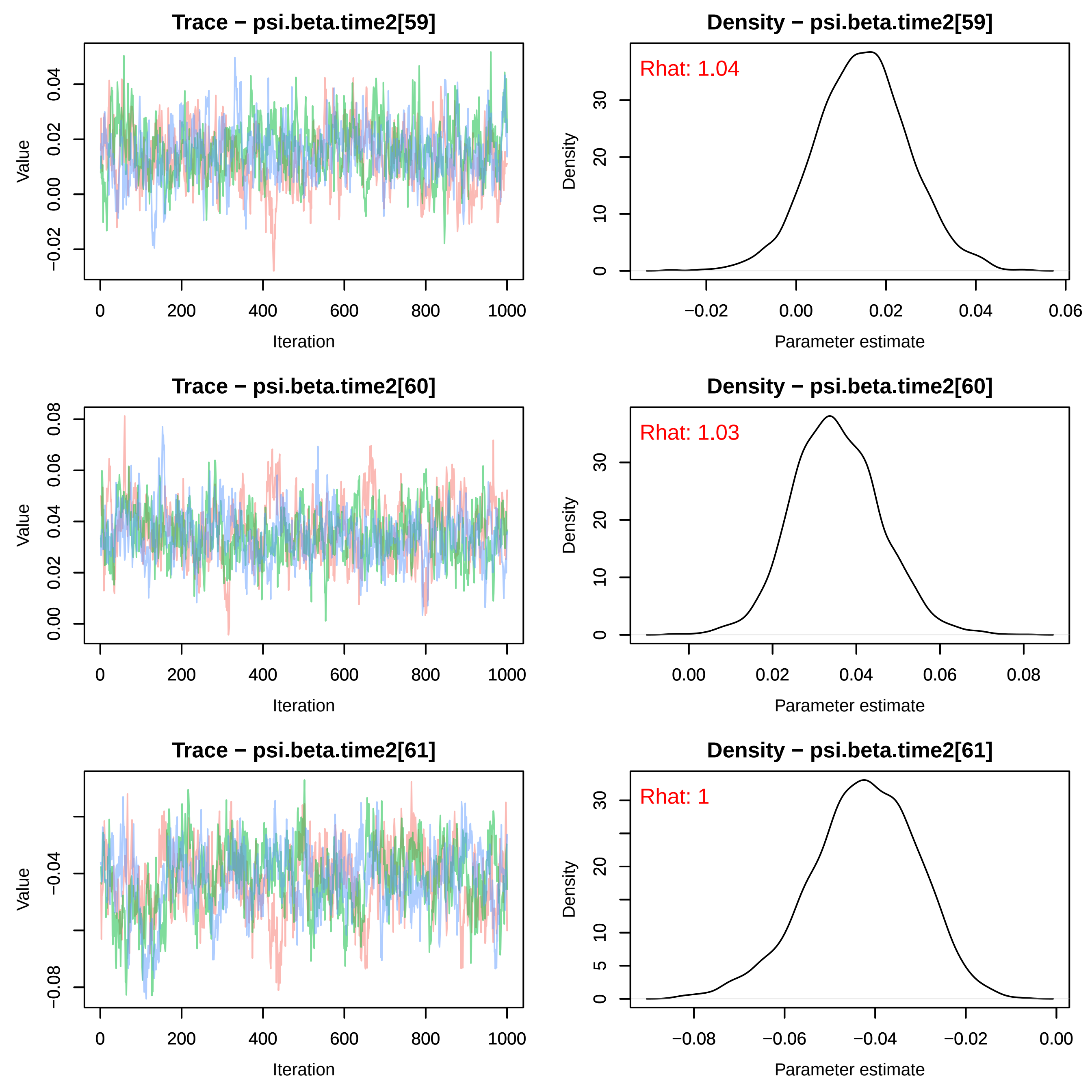

Trace − psi.beta.time2[59]
Density − psi.beta.time2[59]
Rhat: 1.04
Trace − psi.beta.time2[60]
Density − psi.beta.time2[60]
Rhat: 1.03
Trace − psi.beta.time2[61]
Density − psi.beta.time2[61]
Rhat: 1
Value
Iteration
Density
Parameter estimate

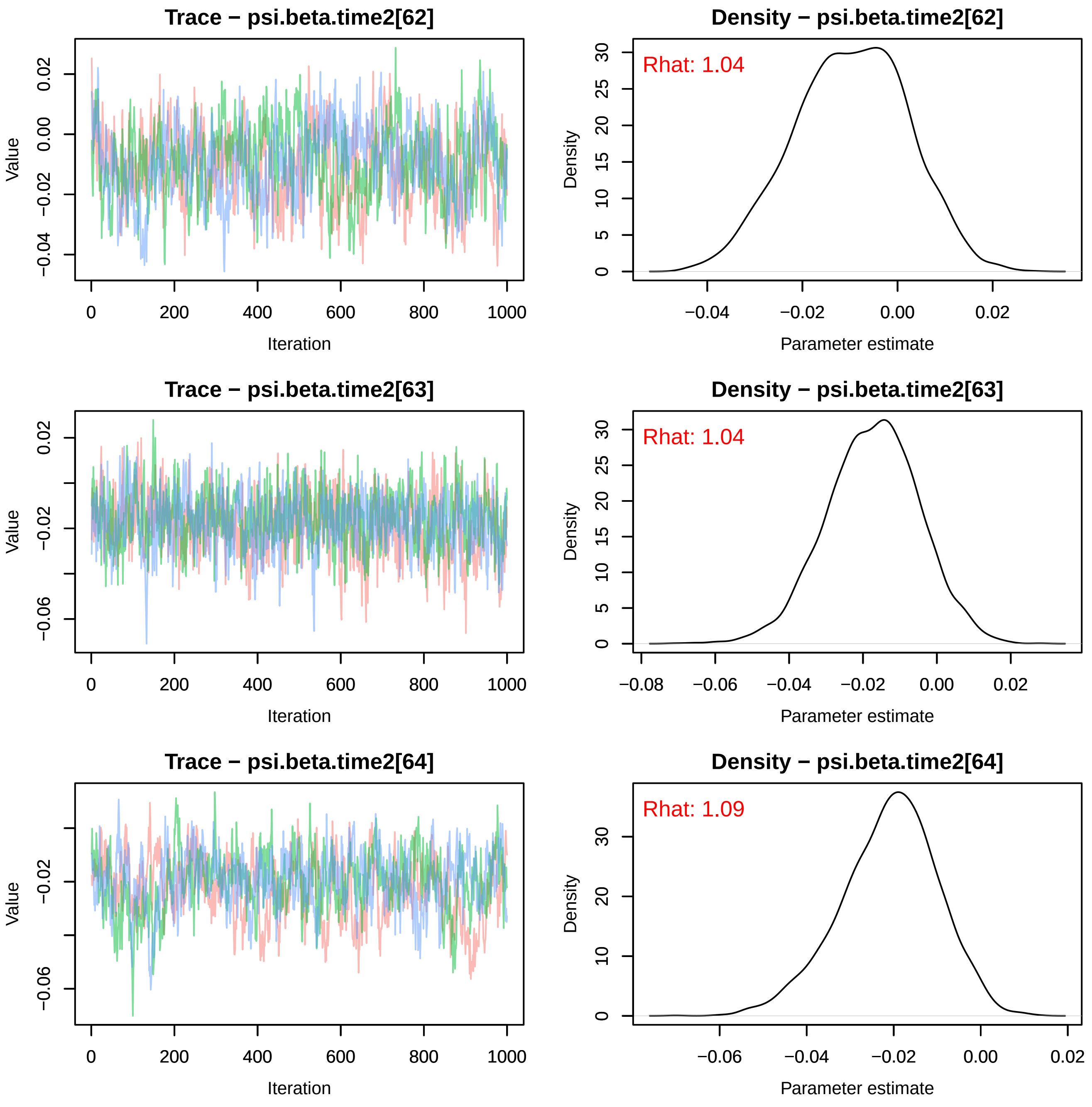

Trace − psi.beta.time2[62]
Value
Iteration
Density − psi.beta.time2[62]
Rhat: 1.04
Density
Parameter estimate
Trace − psi.beta.time2[63]
Value
Iteration
Density − psi.beta.time2[63]
Rhat: 1.04
Density
Parameter estimate
Trace − psi.beta.time2[64]
Value
Iteration
Density − psi.beta.time2[64]
Rhat: 1.09
Density
Parameter estimate

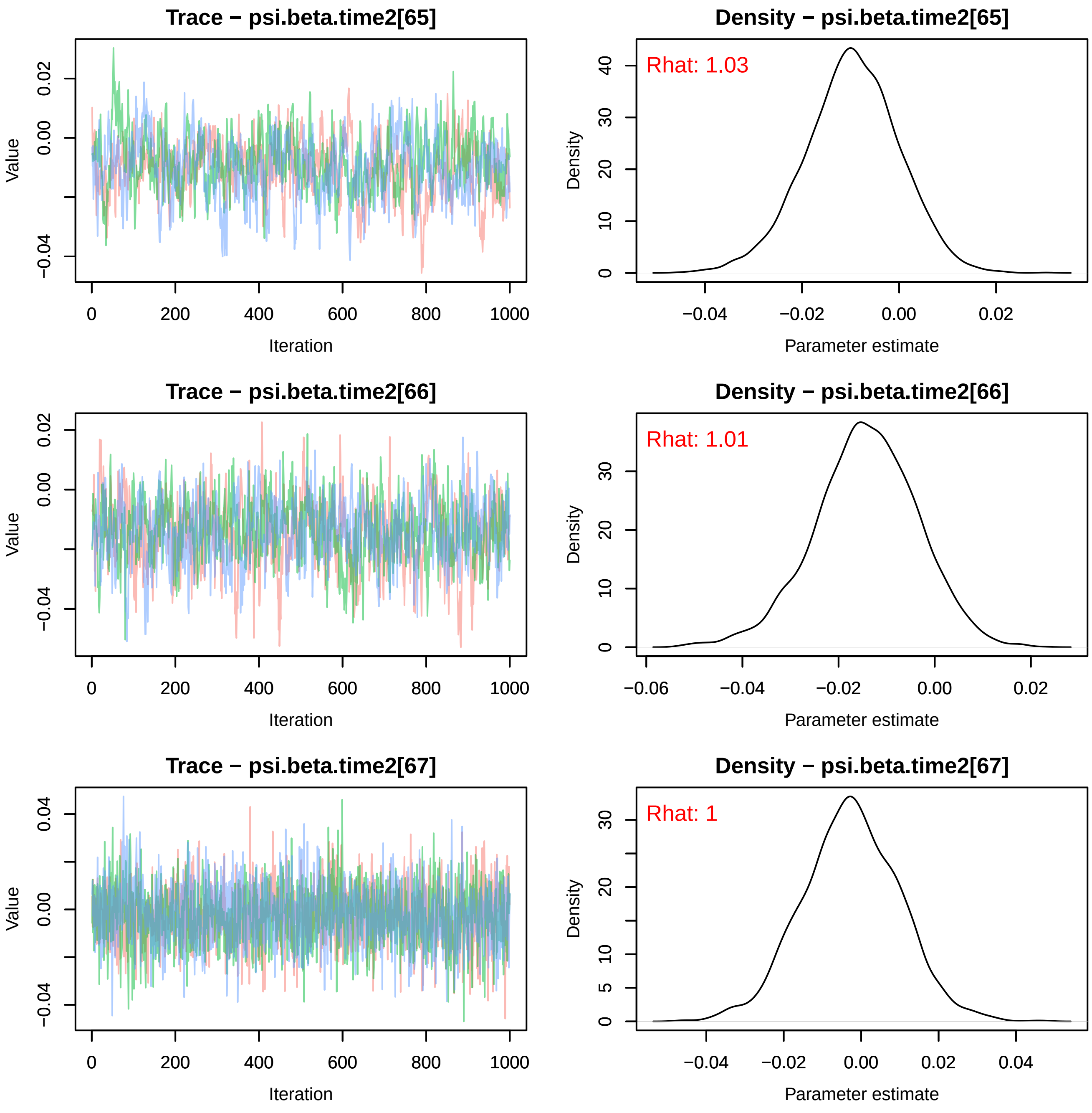
Trace − psi.beta.time2[65]
Value
0.02
0.00
−0.04
0
200
400
600
800
1000
Iteration
Density − psi.beta.time2[65]
Rhat: 1.03
Density
0
10
20
30
40
−0.04
−0.02
0.00
0.02
Parameter estimate
Trace − psi.beta.time2[66]
Value
0.02
0.00
−0.04
0
200
400
600
800
1000
Iteration
Density − psi.beta.time2[66]
Rhat: 1.01
Density
0
10
20
30
−0.06
−0.04
−0.02
0.00
0.02
Parameter estimate
Trace − psi.beta.time2[67]
Value
0.04
0.00
−0.04
0
200
400
600
800
1000
Iteration
Density − psi.beta.time2[67]
Rhat: 1
Density
0
5
10
20
30
−0.04
−0.02
0.00
0.02
0.04
Parameter estimate

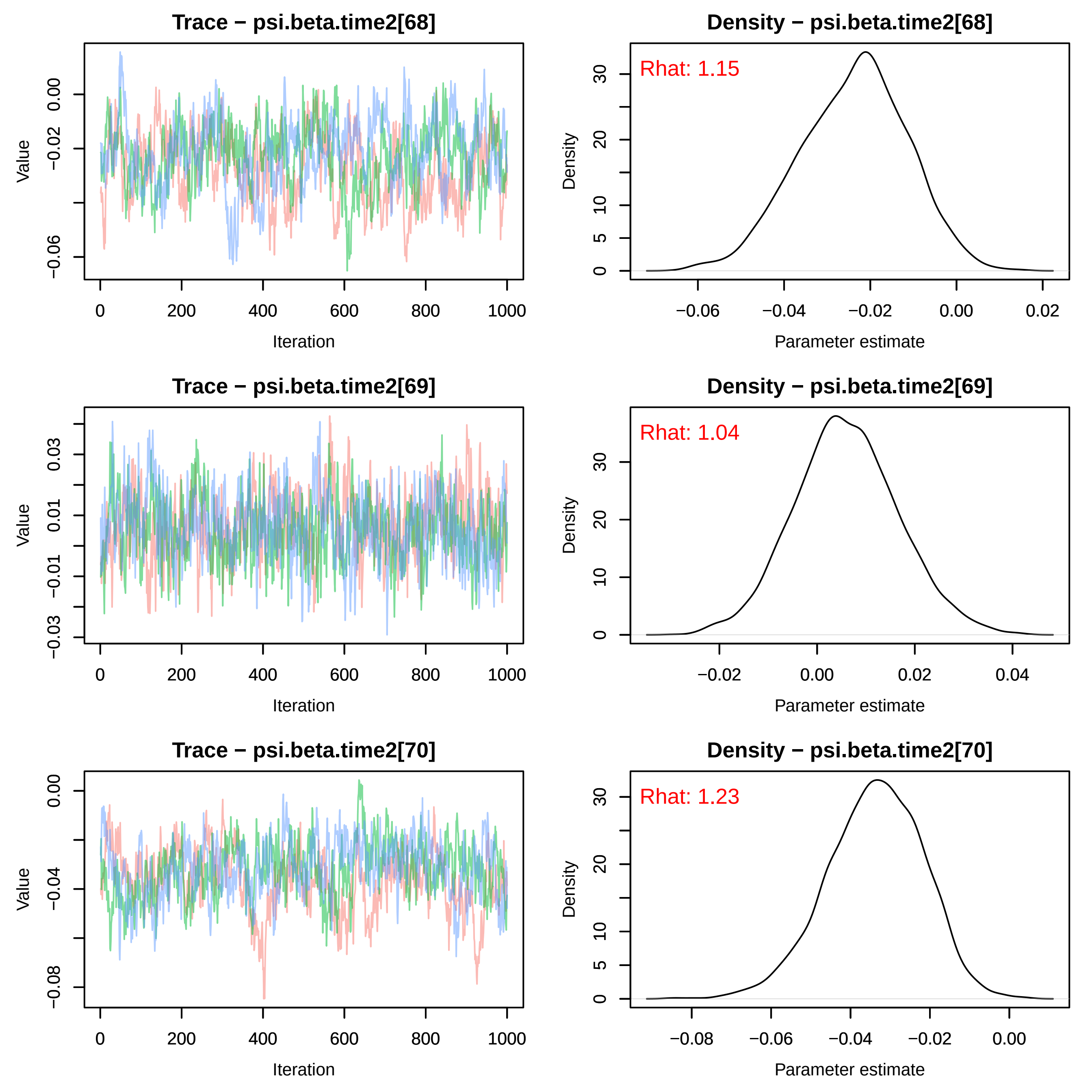
Trace − psi.beta.time2[68]
Density − psi.beta.time2[68]
Rhat: 1.15
Trace − psi.beta.time2[69]
Density − psi.beta.time2[69]
Rhat: 1.04
Trace − psi.beta.time2[70]
Density − psi.beta.time2[70]
Rhat: 1.23
Value
Iteration
Density
Parameter estimate
0
200
400
600
800
1000
0.00
−0.02
−0.06
0.03
0.01
−0.01
−0.03
−0.04
−0.08
0.02
0.04
−0.06
−0.04
−0.02
0
5
10
20
30

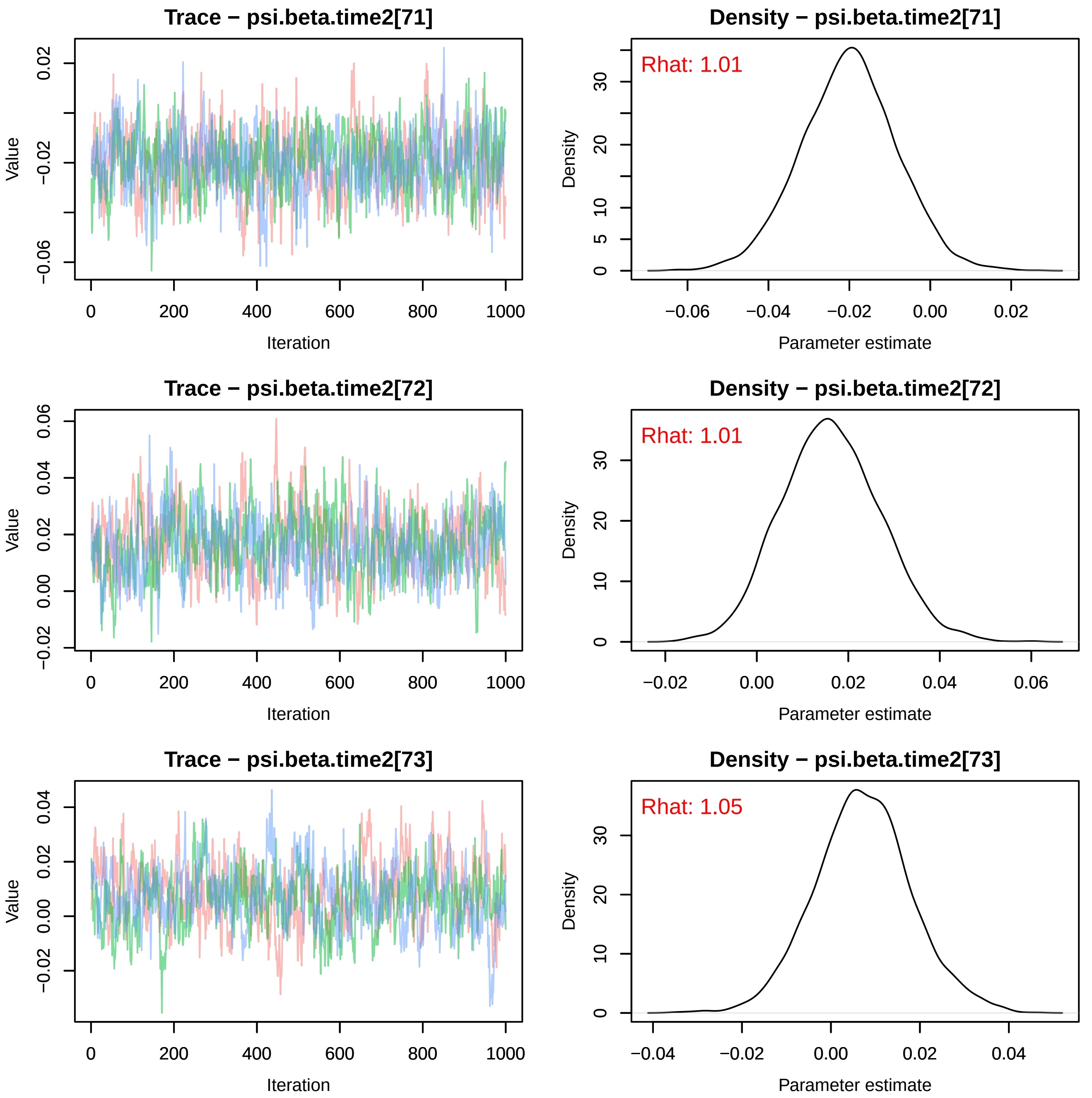
Trace − psi.beta.time2[71]
Value
Iteration
Density − psi.beta.time2[71]
Rhat: 1.01
Density
Parameter estimate
Trace − psi.beta.time2[72]
Value
Iteration
Density − psi.beta.time2[72]
Rhat: 1.01
Density
Parameter estimate
Trace − psi.beta.time2[73]
Value
Iteration
Density − psi.beta.time2[73]
Rhat: 1.05
Density
Parameter estimate

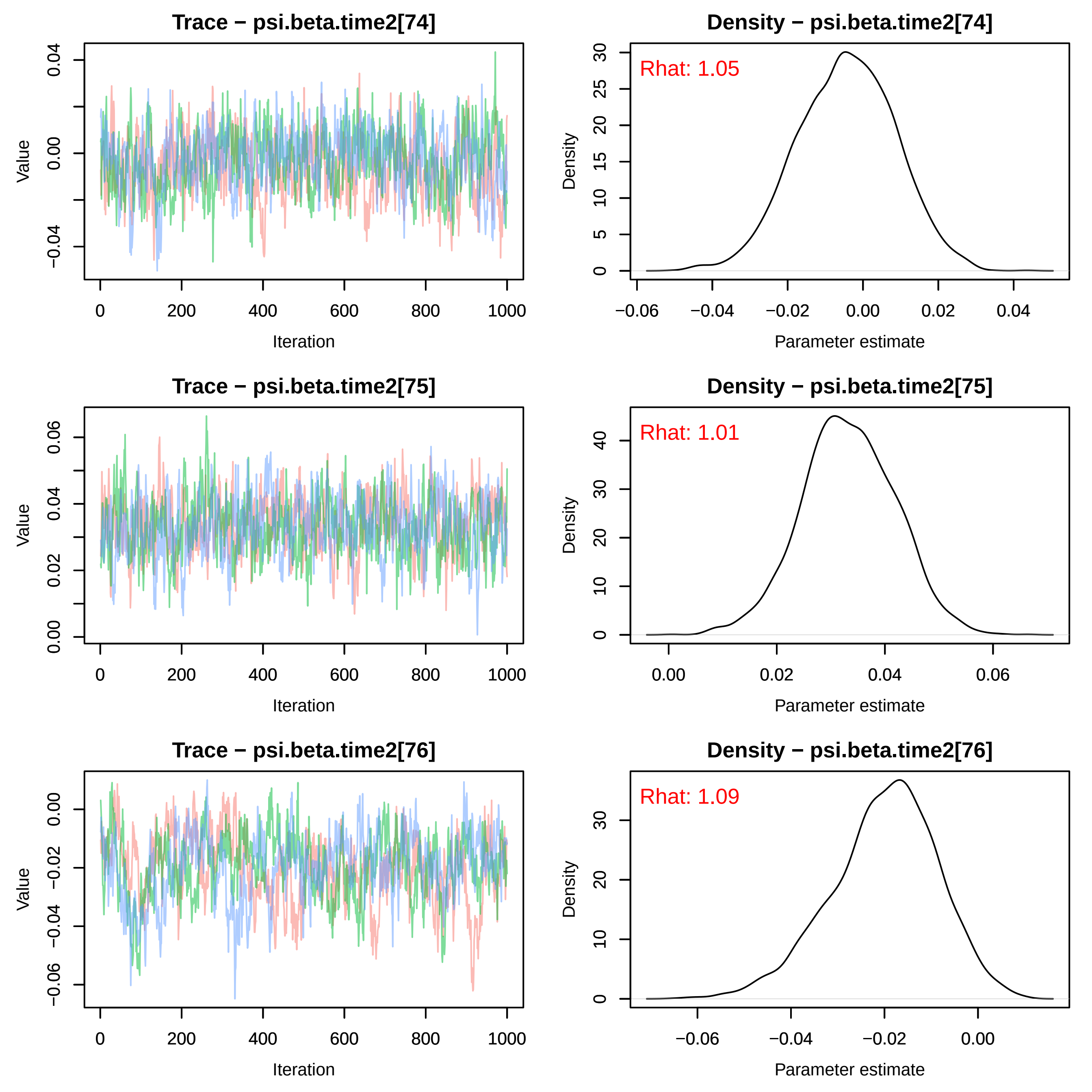
Trace − psi.beta.time2[74]
Value
Iteration
Density − psi.beta.time2[74]
Rhat: 1.05
Density
Parameter estimate
Trace − psi.beta.time2[75]
Value
Iteration
Density − psi.beta.time2[75]
Rhat: 1.01
Density
Parameter estimate
Trace − psi.beta.time2[76]
Value
Iteration
Density − psi.beta.time2[76]
Rhat: 1.09
Density
Parameter estimate

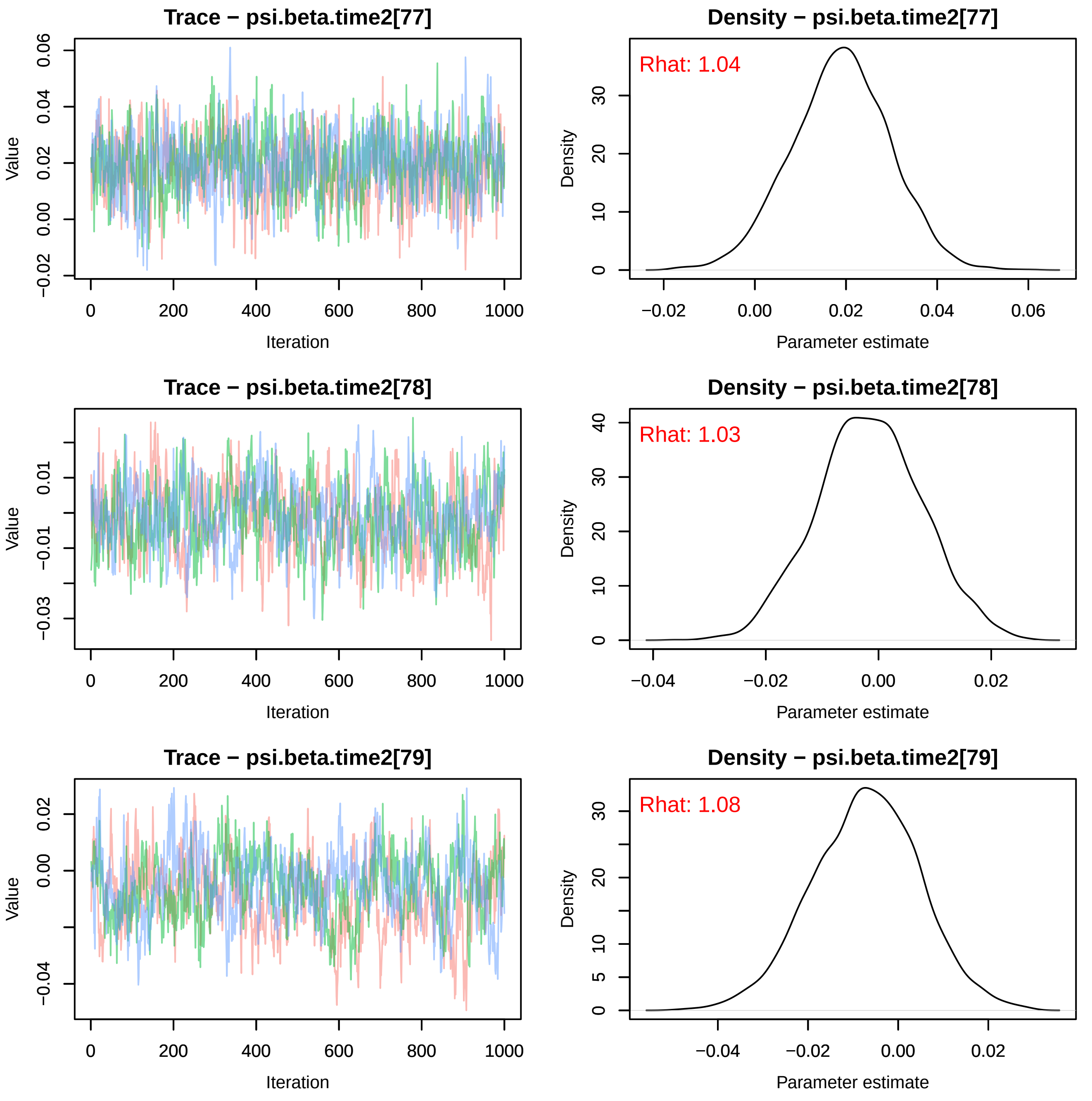

Trace − psi.beta.time2[77]
Value
Iteration
Density − psi.beta.time2[77]
Rhat: 1.04
Density
Parameter estimate
Trace − psi.beta.time2[78]
Value
Iteration
Density − psi.beta.time2[78]
Rhat: 1.03
Density
Parameter estimate
Trace − psi.beta.time2[79]
Value
Iteration
Density − psi.beta.time2[79]
Rhat: 1.08
Density
Parameter estimate

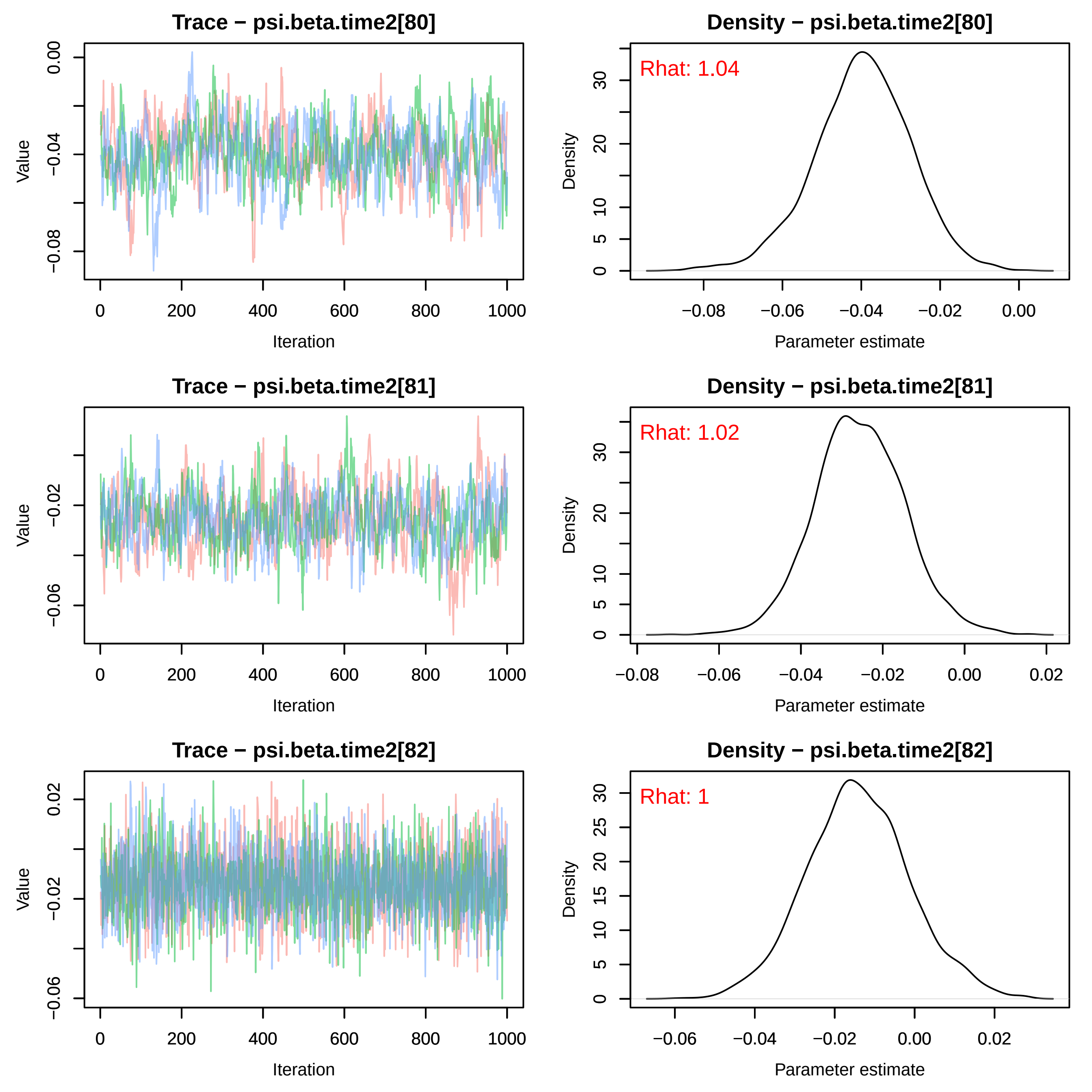

Trace − psi.beta.time2[80]
Value
Iteration
Density − psi.beta.time2[80]
Rhat: 1.04
Density
Parameter estimate
Trace − psi.beta.time2[81]
Value
Iteration
Density − psi.beta.time2[81]
Rhat: 1.02
Density
Parameter estimate
Trace − psi.beta.time2[82]
Value
Iteration
Density − psi.beta.time2[82]
Rhat: 1
Density
Parameter estimate

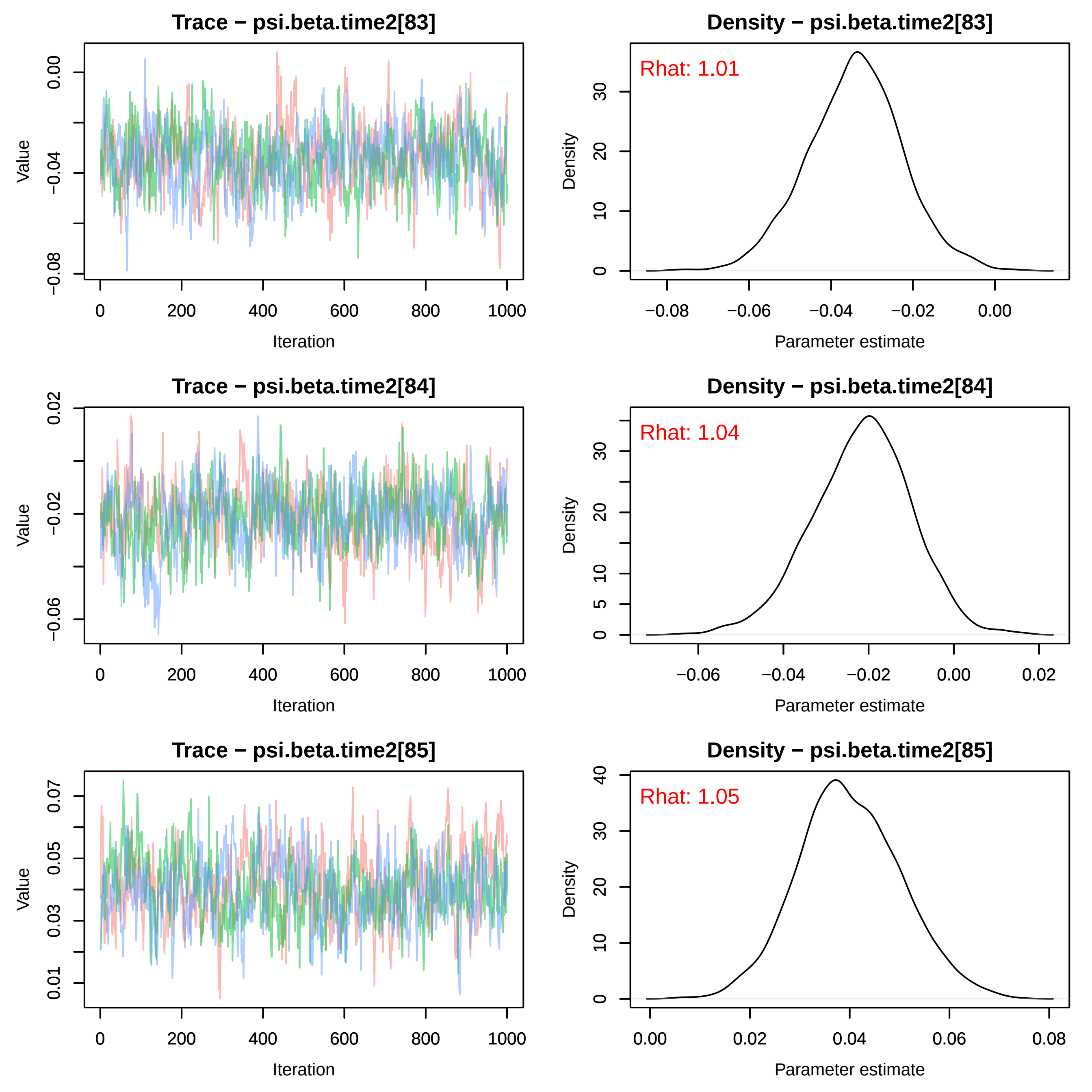

Trace − psi.beta.time2[83]
Value
−0.08
−0.04
0.00
0
200
400
600
800
1000
Iteration
Density − psi.beta.time2[83]
Rhat: 1.01
Density
0
10
20
30
−0.08
−0.06
−0.04
−0.02
0.00
Parameter estimate
Trace − psi.beta.time2[84]
Value
−0.06
−0.02
0.02
0
200
400
600
800
1000
Iteration
Density − psi.beta.time2[84]
Rhat: 1.04
Density
0
5
10
20
30
−0.06
−0.04
−0.02
0.00
0.02
Parameter estimate
Trace − psi.beta.time2[85]
Value
0.01
0.03
0.05
0.07
0
200
400
600
800
1000
Iteration
Density − psi.beta.time2[85]
Rhat: 1.05
Density
0
10
20
30
40
0.00
0.02
0.04
0.06
0.08
Parameter estimate

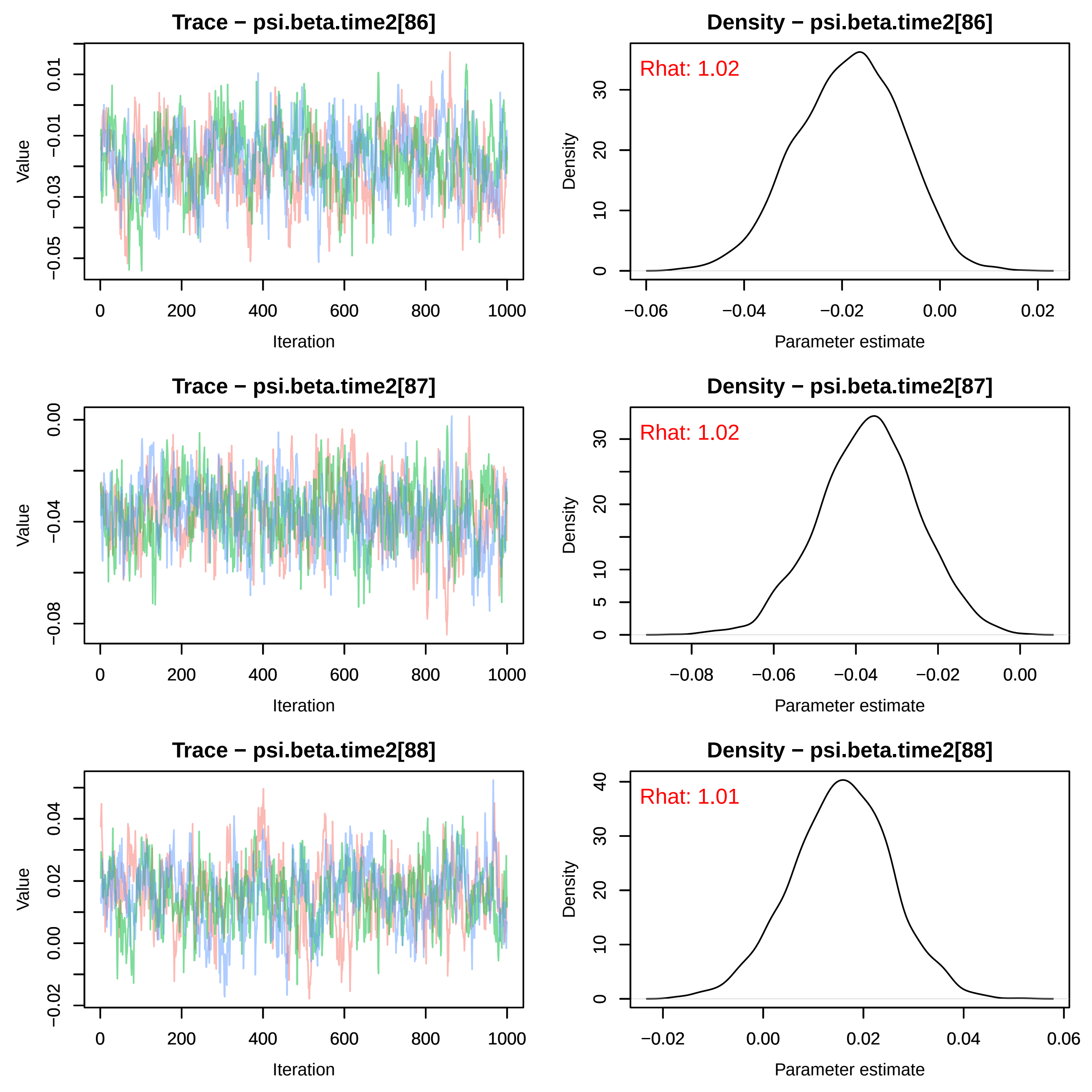

Trace − psi.beta.time2[86]
Value
Iteration
Density − psi.beta.time2[86]
Rhat: 1.02
Density
Parameter estimate
Trace − psi.beta.time2[87]
Value
Iteration
Density − psi.beta.time2[87]
Rhat: 1.02
Density
Parameter estimate
Trace − psi.beta.time2[88]
Value
Iteration
Density − psi.beta.time2[88]
Rhat: 1.01
Density
Parameter estimate

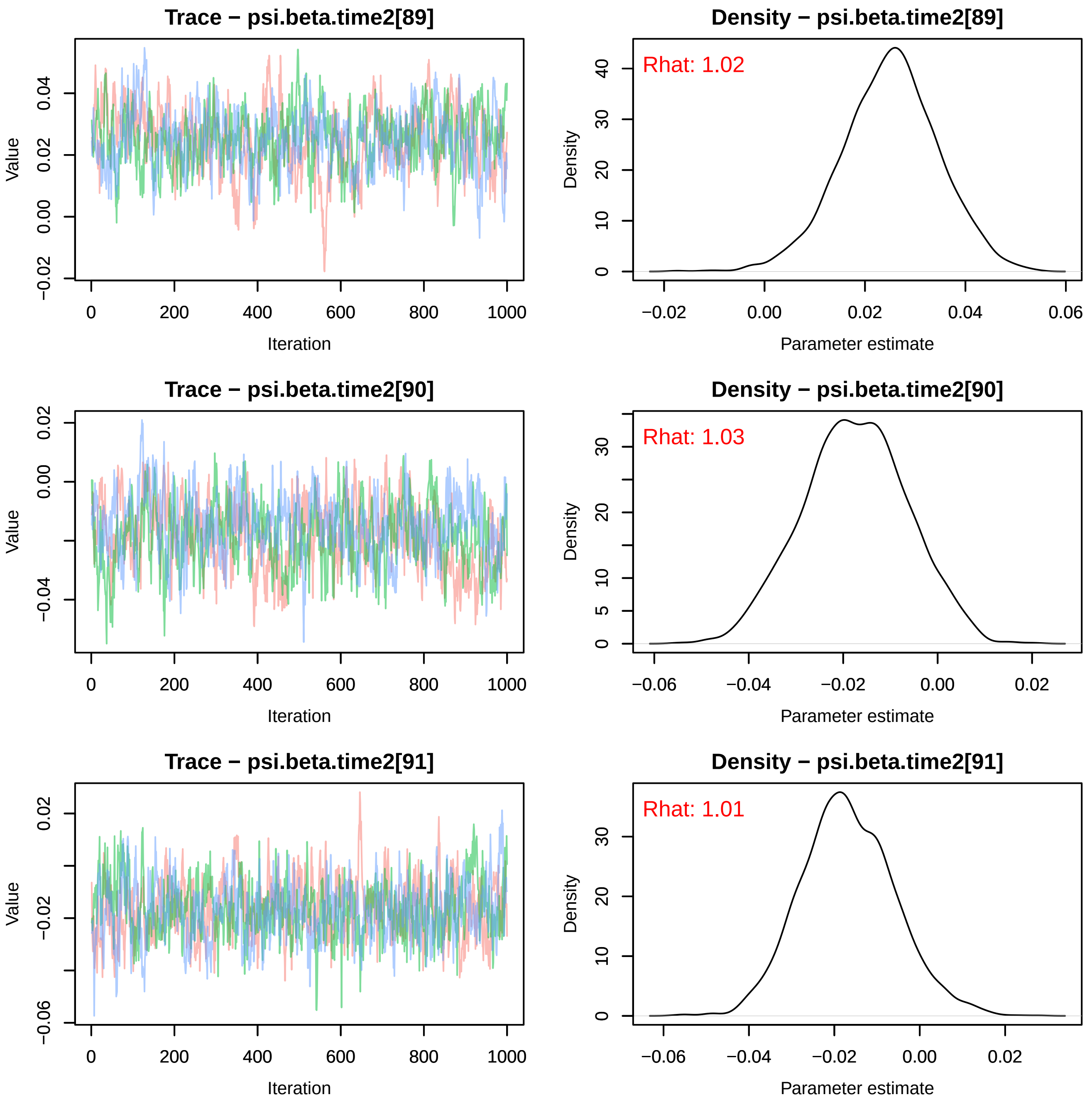
Trace − psi.beta.time2[89]
Density − psi.beta.time2[89]
Rhat: 1.02
Trace − psi.beta.time2[90]
Density − psi.beta.time2[90]
Rhat: 1.03
Trace − psi.beta.time2[91]
Density − psi.beta.time2[91]
Rhat: 1.01
Value
Iteration
Density
Parameter estimate

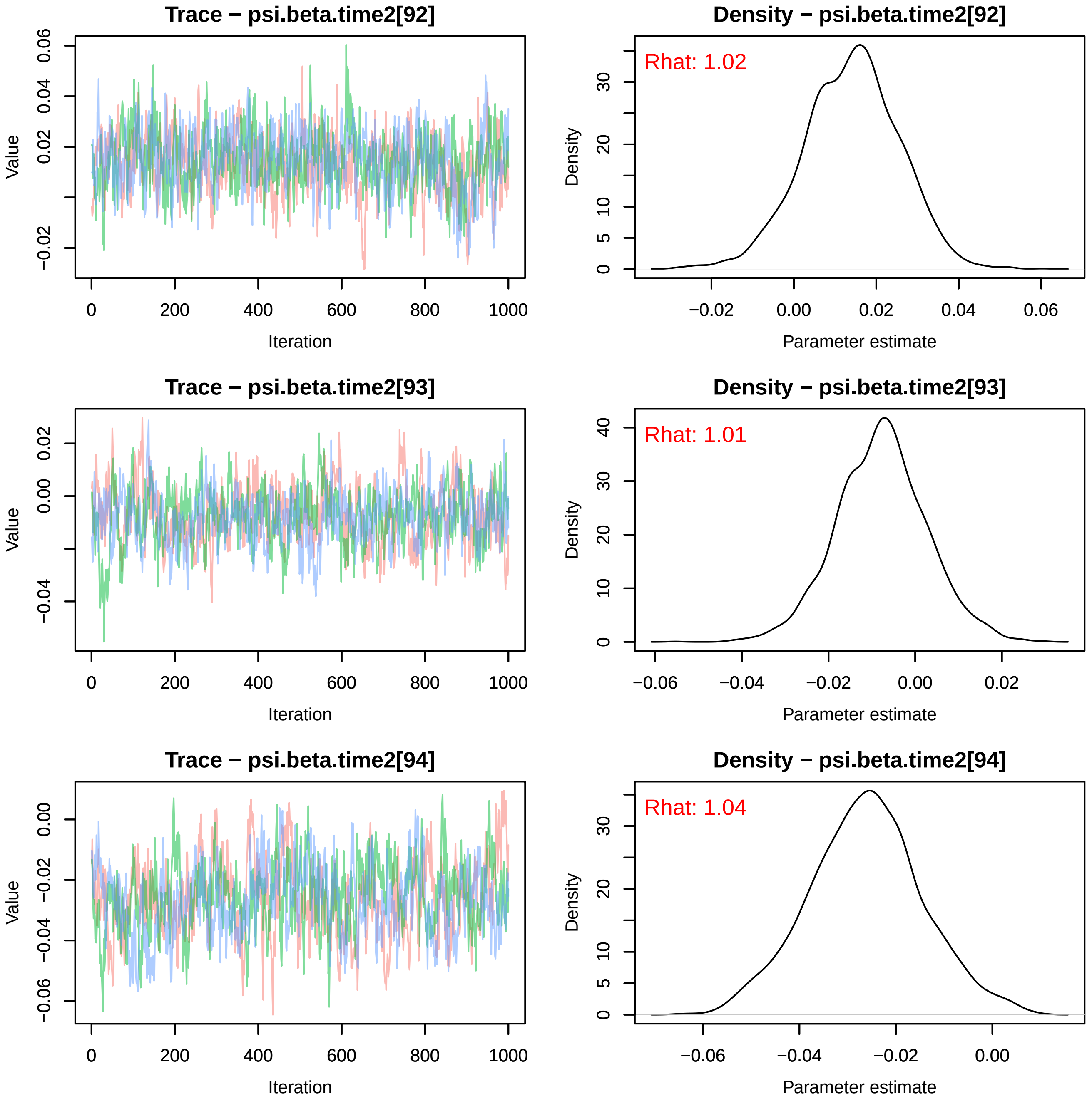

Trace − psi.beta.time2[92]
Density − psi.beta.time2[92]
Rhat: 1.02
Trace − psi.beta.time2[93]
Density − psi.beta.time2[93]
Rhat: 1.01
Trace − psi.beta.time2[94]
Density − psi.beta.time2[94]
Rhat: 1.04
Value
Iteration
Density
Parameter estimate
0
200
400
600
800
1000
−0.06
−0.04
−0.02
0.00
0.02
0.04
0.06
5
10
20
30
40

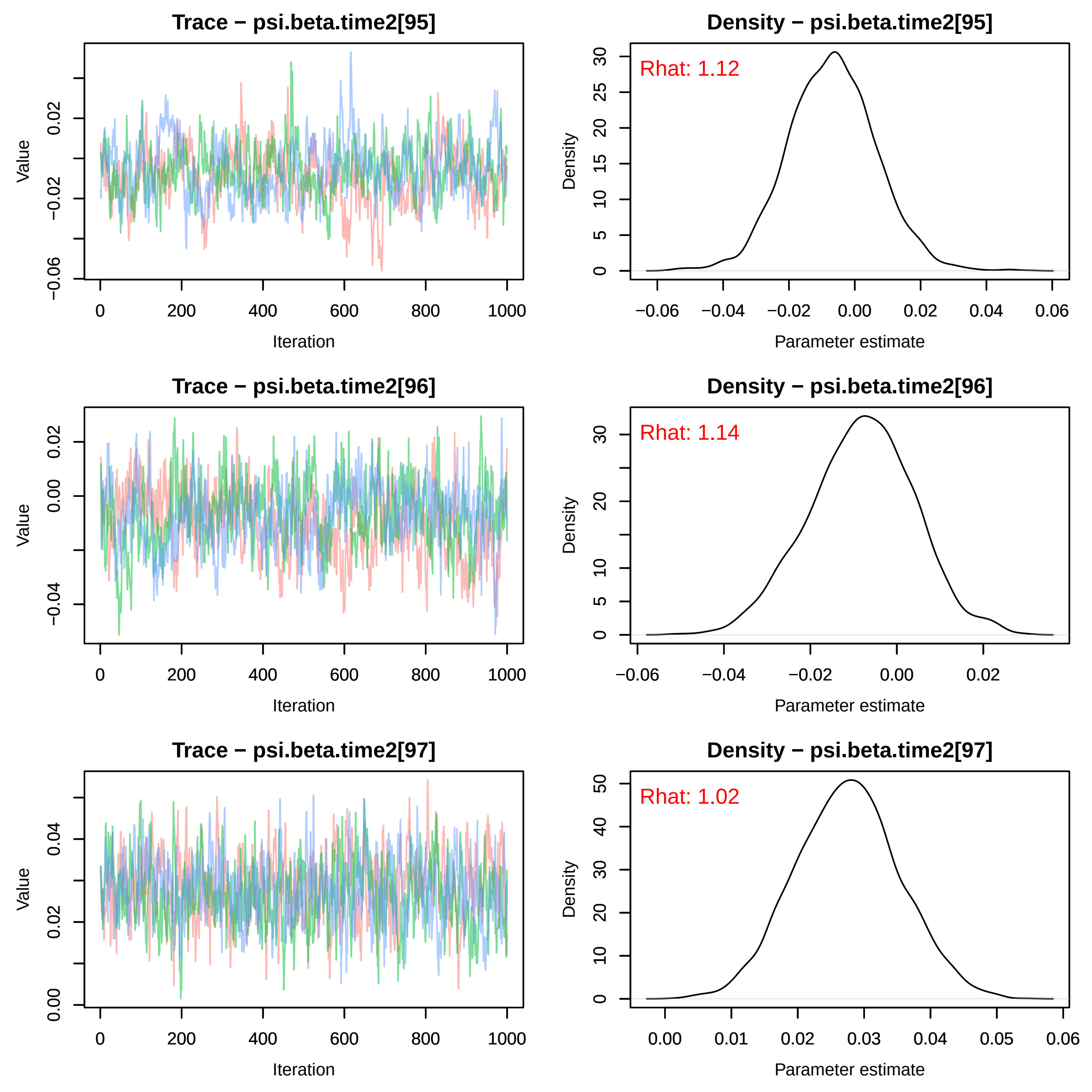
Trace − psi.beta.time2[95]
Value
0.02
−0.02
−0.06
0
200
400
600
800
1000
Iteration
Density − psi.beta.time2[95]
Rhat: 1.12
Density
0
5
10
15
20
25
30
−0.06
−0.04
−0.02
0.00
0.02
0.04
0.06
Parameter estimate
Trace − psi.beta.time2[96]
Value
0.02
0.00
−0.04
0
200
400
600
800
1000
Iteration
Density − psi.beta.time2[96]
Rhat: 1.14
Density
0
5
10
20
30
−0.06
−0.04
−0.02
0.00
0.02
Parameter estimate
Trace − psi.beta.time2[97]
Value
0.00
0.02
0.04
0
200
400
600
800
1000
Iteration
Density − psi.beta.time2[97]
Rhat: 1.02
Density
0
10
20
30
40
50
0.00
0.01
0.02
0.03
0.04
0.05
0.06
Parameter estimate

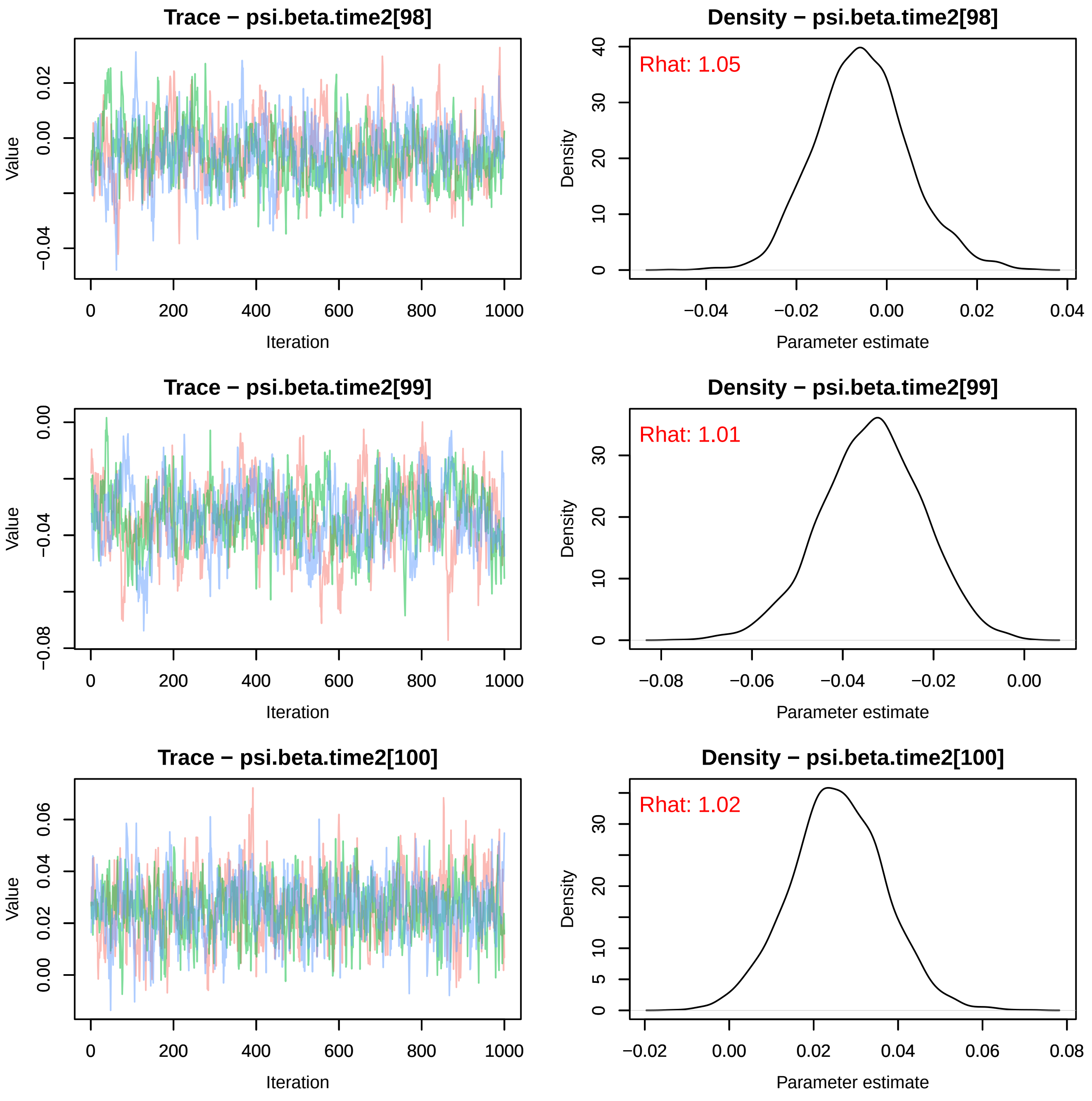

Trace − psi.beta.time2[98]
Value
Iteration
Density − psi.beta.time2[98]
Rhat: 1.05
Density
Parameter estimate
Trace − psi.beta.time2[99]
Value
Iteration
Density − psi.beta.time2[99]
Rhat: 1.01
Density
Parameter estimate
Trace − psi.beta.time2[100]
Value
Iteration
Density − psi.beta.time2[100]
Rhat: 1.02
Density
Parameter estimate

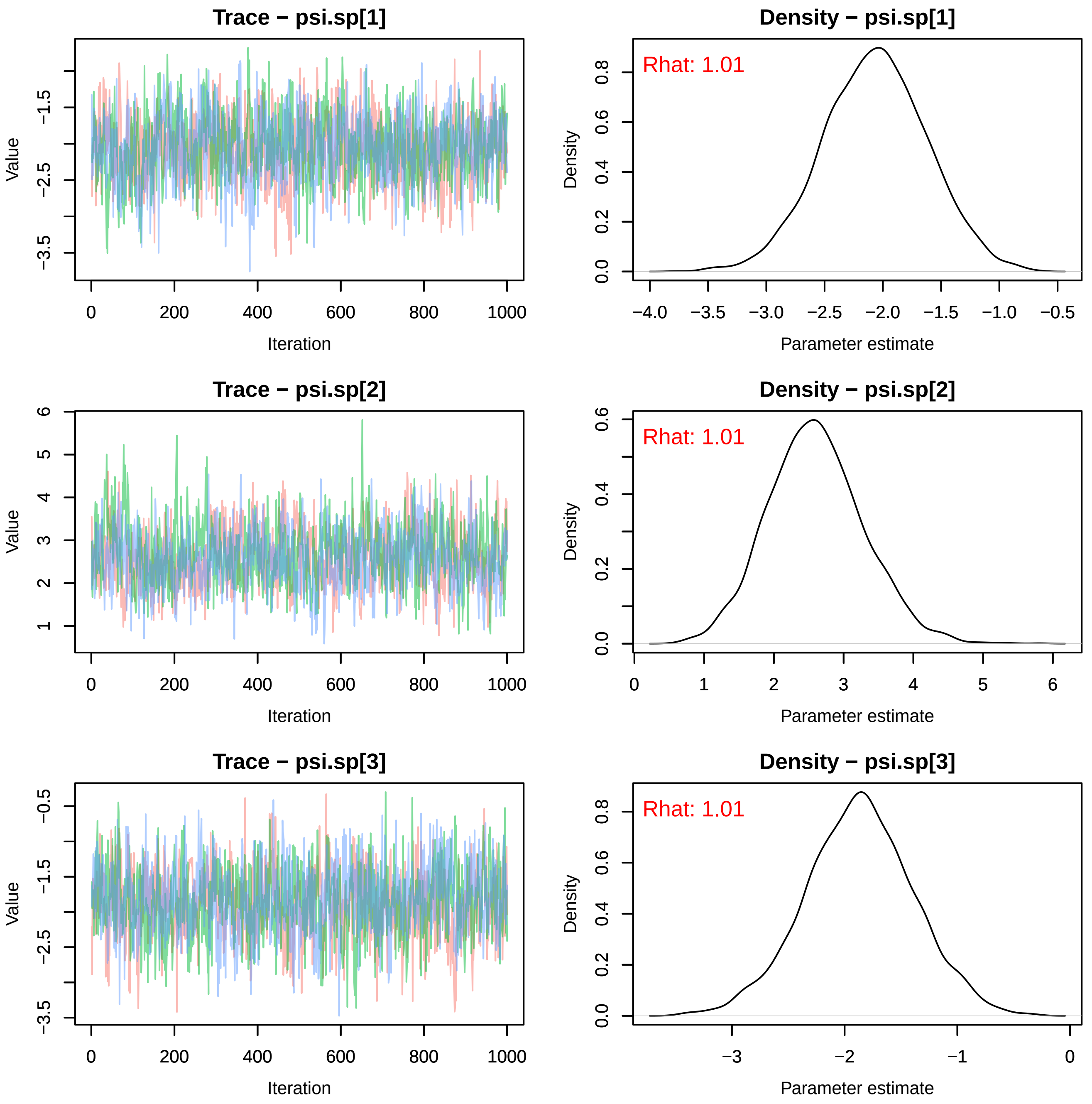
Trace − psi.sp[1]
Density − psi.sp[1]
Rhat: 1.01
Value
Density
Iteration
Parameter estimate
Trace − psi.sp[2]
Density − psi.sp[2]
Rhat: 1.01
Value
Density
Iteration
Parameter estimate
Trace − psi.sp[3]
Density − psi.sp[3]
Rhat: 1.01
Value
Density
Iteration
Parameter estimate

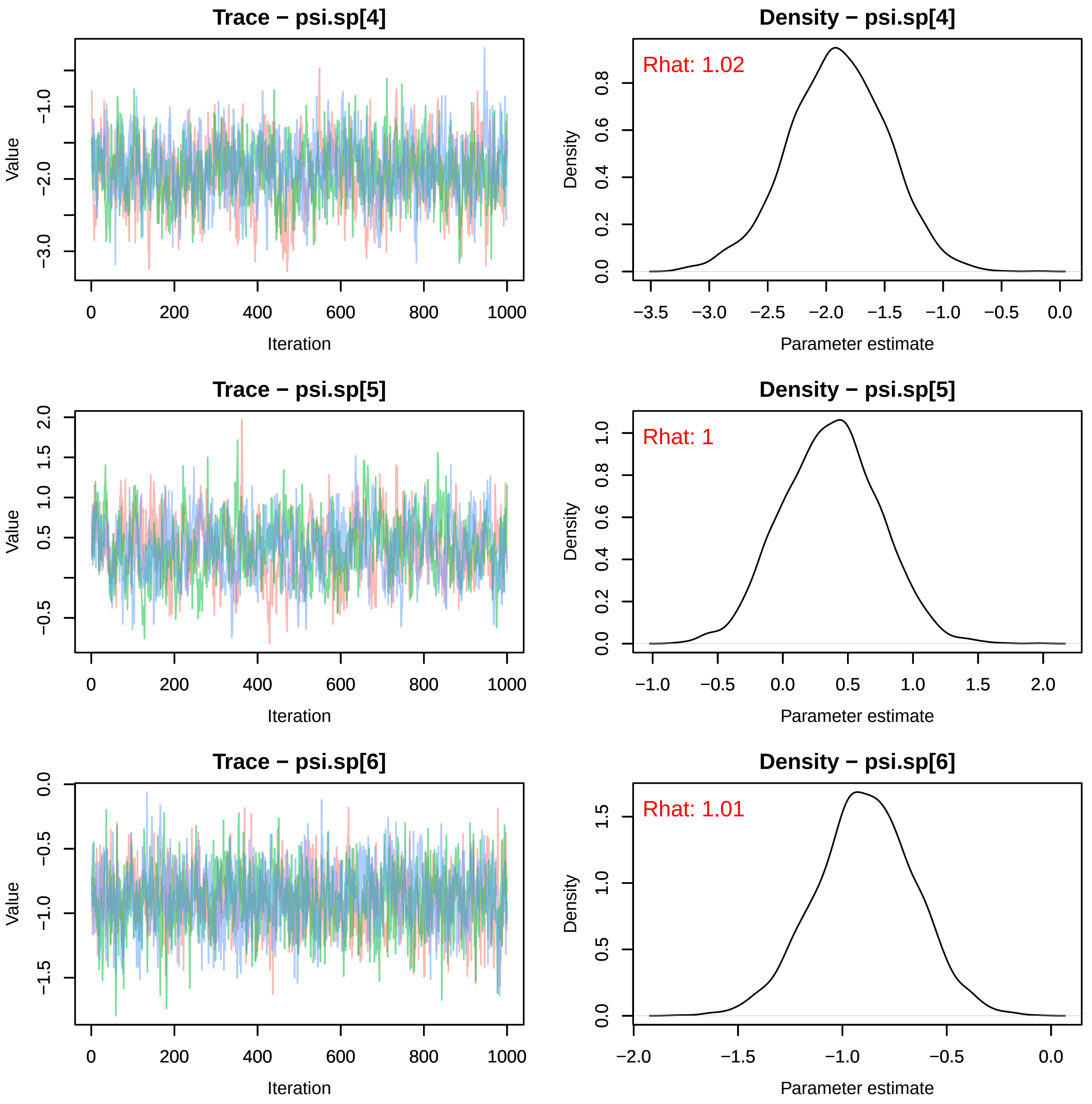
Trace − psi.sp[4]
Value
Iteration
Density − psi.sp[4]
Rhat: 1.02
Density
Parameter estimate
Trace − psi.sp[5]
Value
Iteration
Density − psi.sp[5]
Rhat: 1
Density
Parameter estimate
Trace − psi.sp[6]
Value
Iteration
Density − psi.sp[6]
Rhat: 1.01
Density
Parameter estimate

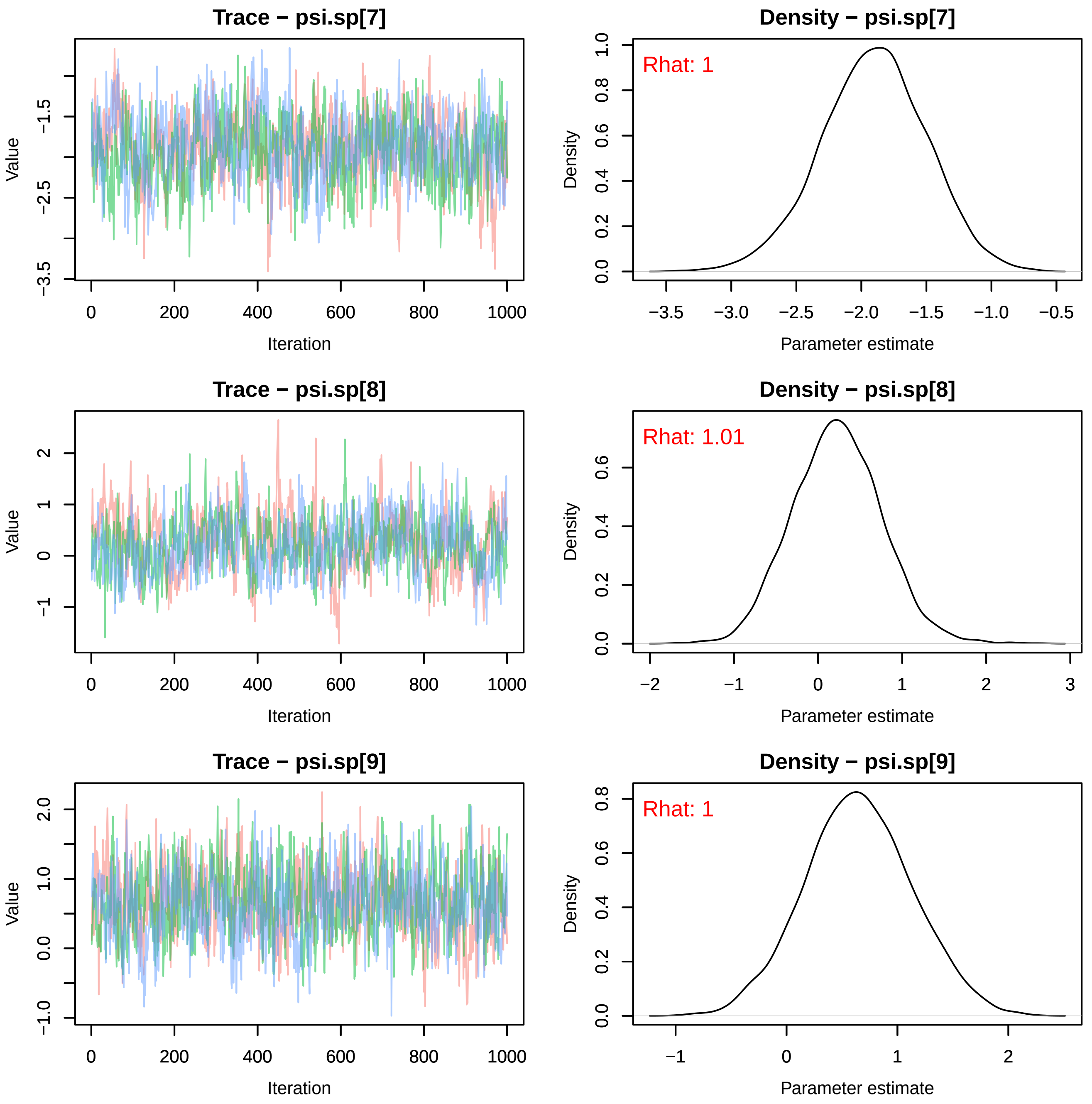

Trace − psi.sp[7]
Density − psi.sp[7]
Rhat: 1
Trace − psi.sp[8]
Density − psi.sp[8]
Rhat: 1.01
Trace − psi.sp[9]
Density − psi.sp[9]
Rhat: 1
Value
Iteration
Density
Parameter estimate

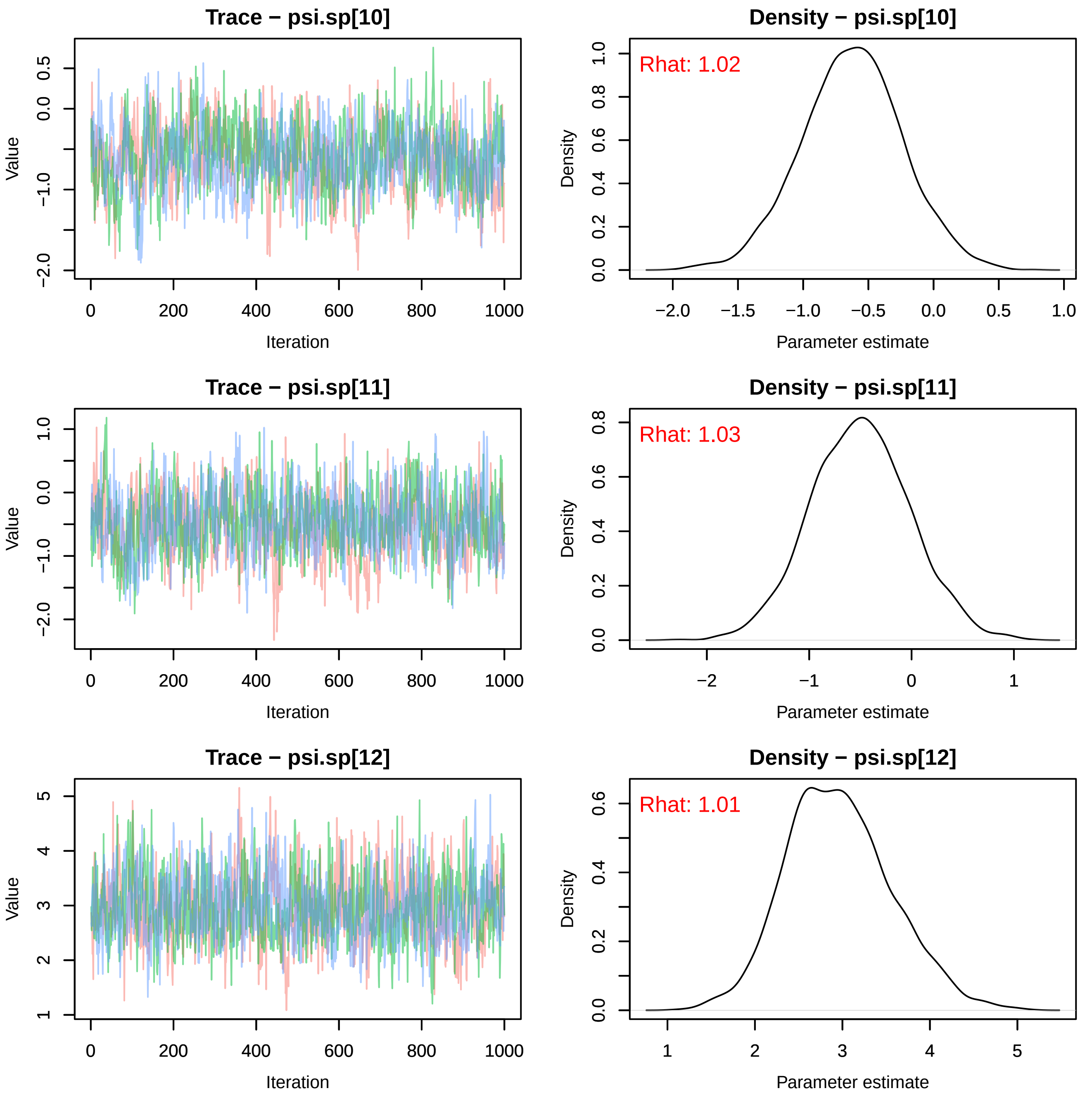
Trace − psi.sp[10]
Value
Iteration
0 200 400 600 800 1000
−2.0 −1.0 0.0 0.5
Density − psi.sp[10]
Rhat: 1.02
Density
Parameter estimate
−2.0 −1.5 −1.0 −0.5 0.0 0.5 1.0
0.0 0.2 0.4 0.6 0.8 1.0
Trace − psi.sp[11]
Value
Iteration
0 200 400 600 800 1000
−2.0 −1.0 0.0 1.0
Density − psi.sp[11]
Rhat: 1.03
Density
Parameter estimate
−2 −1 0 1
0.0 0.2 0.4 0.6 0.8
Trace − psi.sp[12]
Value
Iteration
0 200 400 600 800 1000
1 2 3 4 5
Density − psi.sp[12]
Rhat: 1.01
Density
Parameter estimate
1 2 3 4 5
0.0 0.2 0.4 0.6

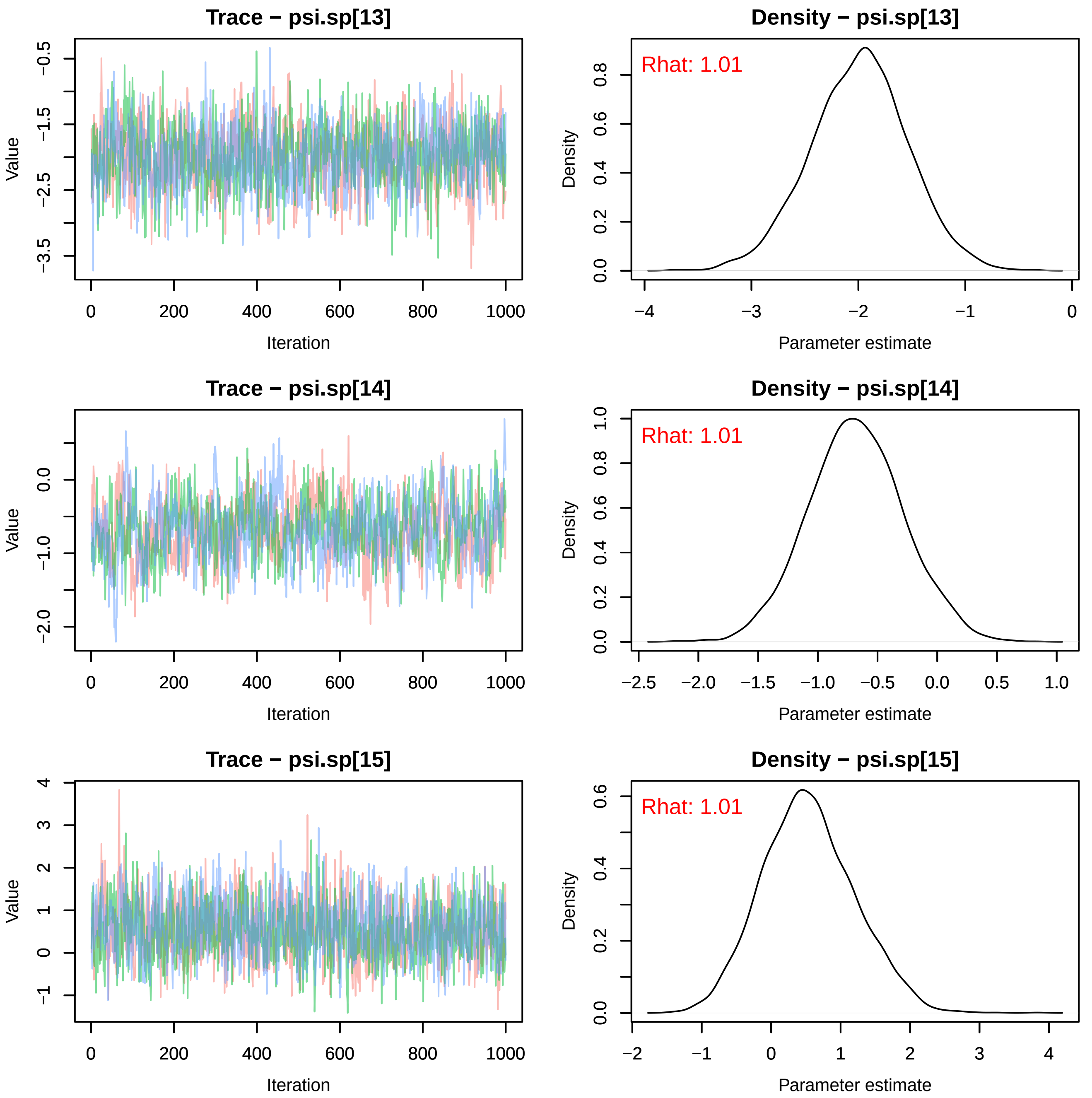
Trace − psi.sp[13]
Value
Iteration
Density − psi.sp[13]
Rhat: 1.01
Density
Parameter estimate
Trace − psi.sp[14]
Value
Iteration
Density − psi.sp[14]
Rhat: 1.01
Density
Parameter estimate
Trace − psi.sp[15]
Value
Iteration
Density − psi.sp[15]
Rhat: 1.01
Density
Parameter estimate

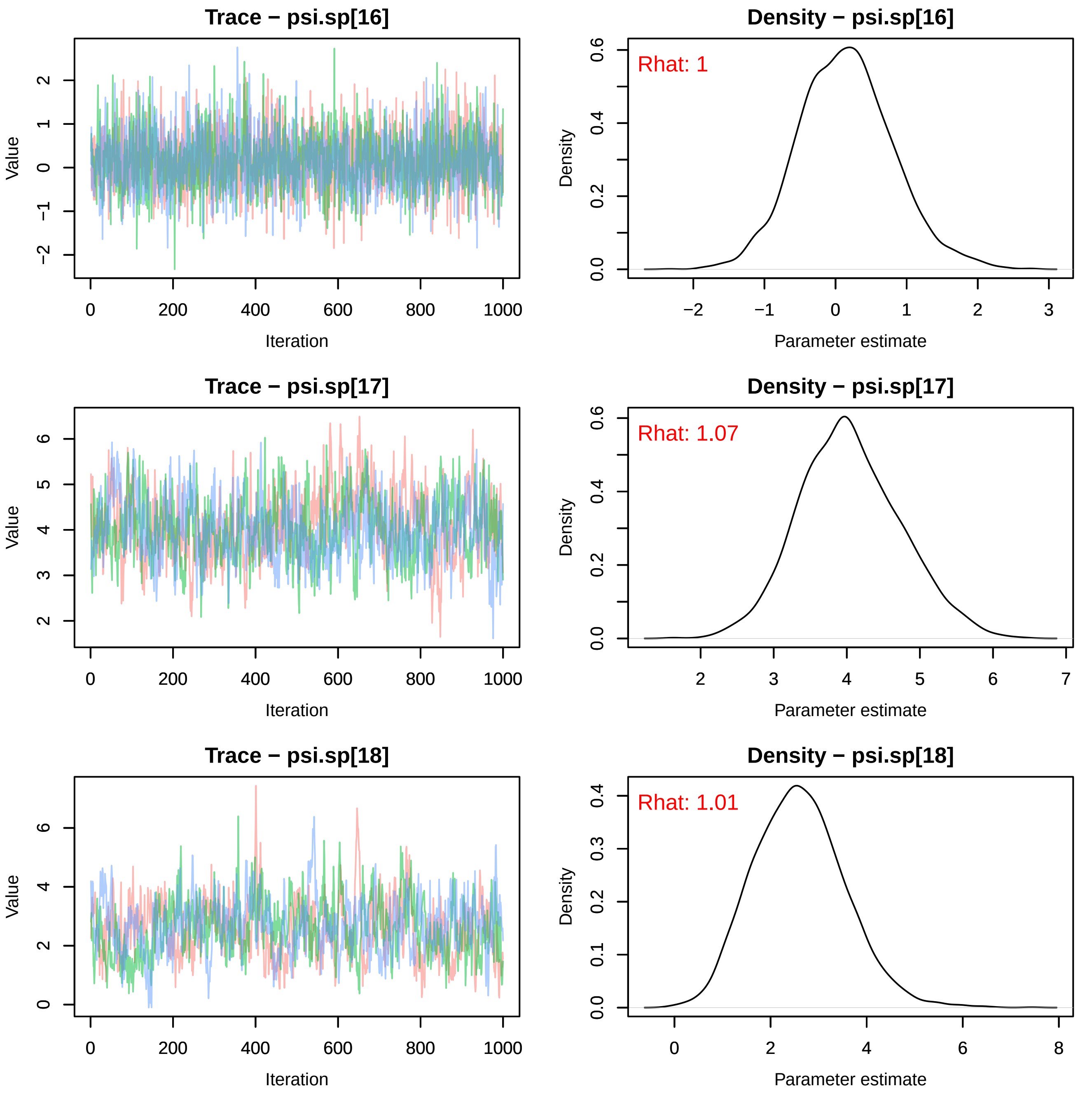
Trace − psi.sp[16]
Value
Iteration
Density − psi.sp[16]
Rhat: 1
Density
Parameter estimate
Trace − psi.sp[17]
Value
Iteration
Density − psi.sp[17]
Rhat: 1.07
Density
Parameter estimate
Trace − psi.sp[18]
Value
Iteration
Density − psi.sp[18]
Rhat: 1.01
Density
Parameter estimate

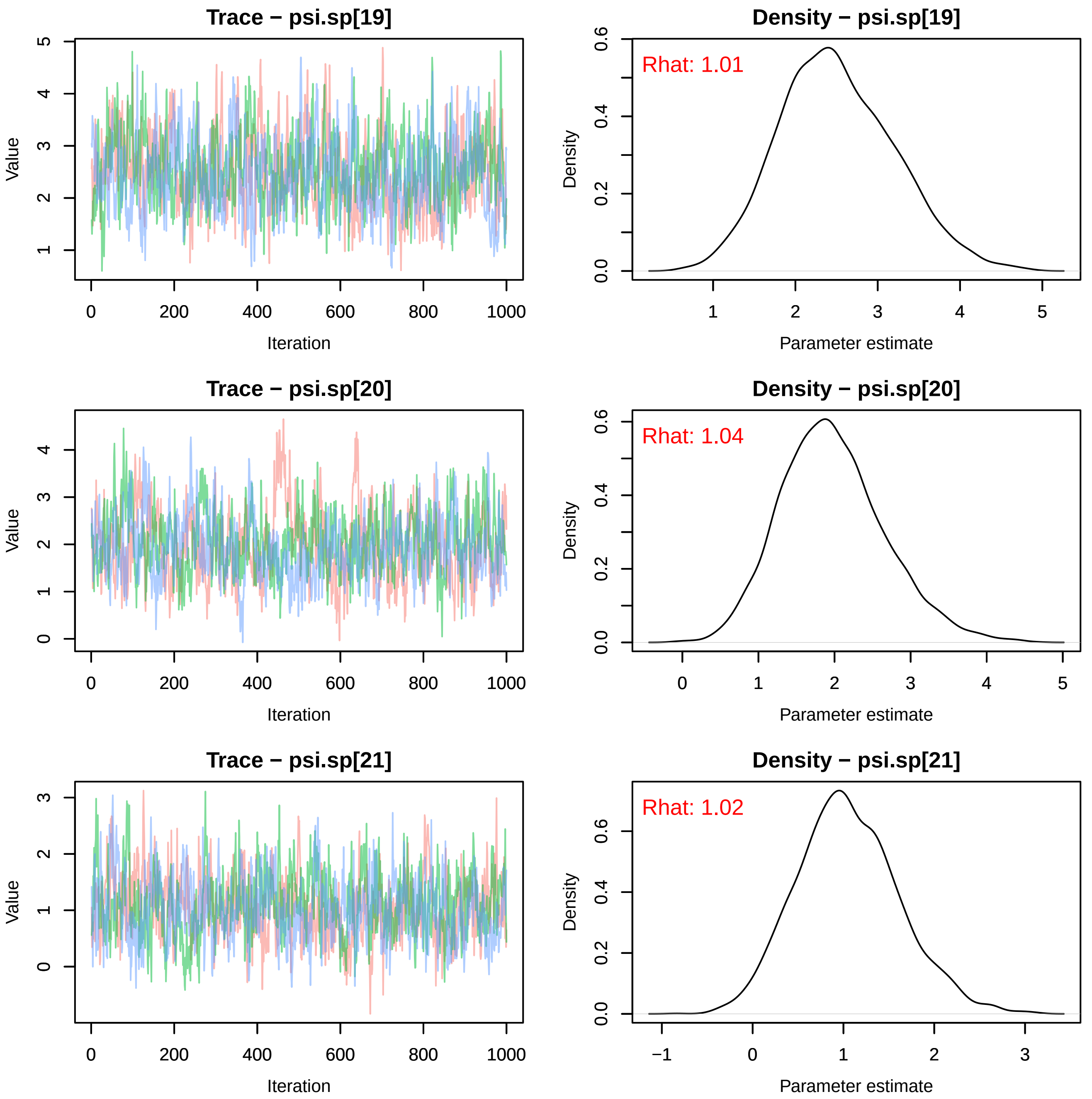

Trace − psi.sp[19]
Value
Iteration
Density − psi.sp[19]
Rhat: 1.01
Density
Parameter estimate
Trace − psi.sp[20]
Value
Iteration
Density − psi.sp[20]
Rhat: 1.04
Density
Parameter estimate
Trace − psi.sp[21]
Value
Iteration
Density − psi.sp[21]
Rhat: 1.02
Density
Parameter estimate

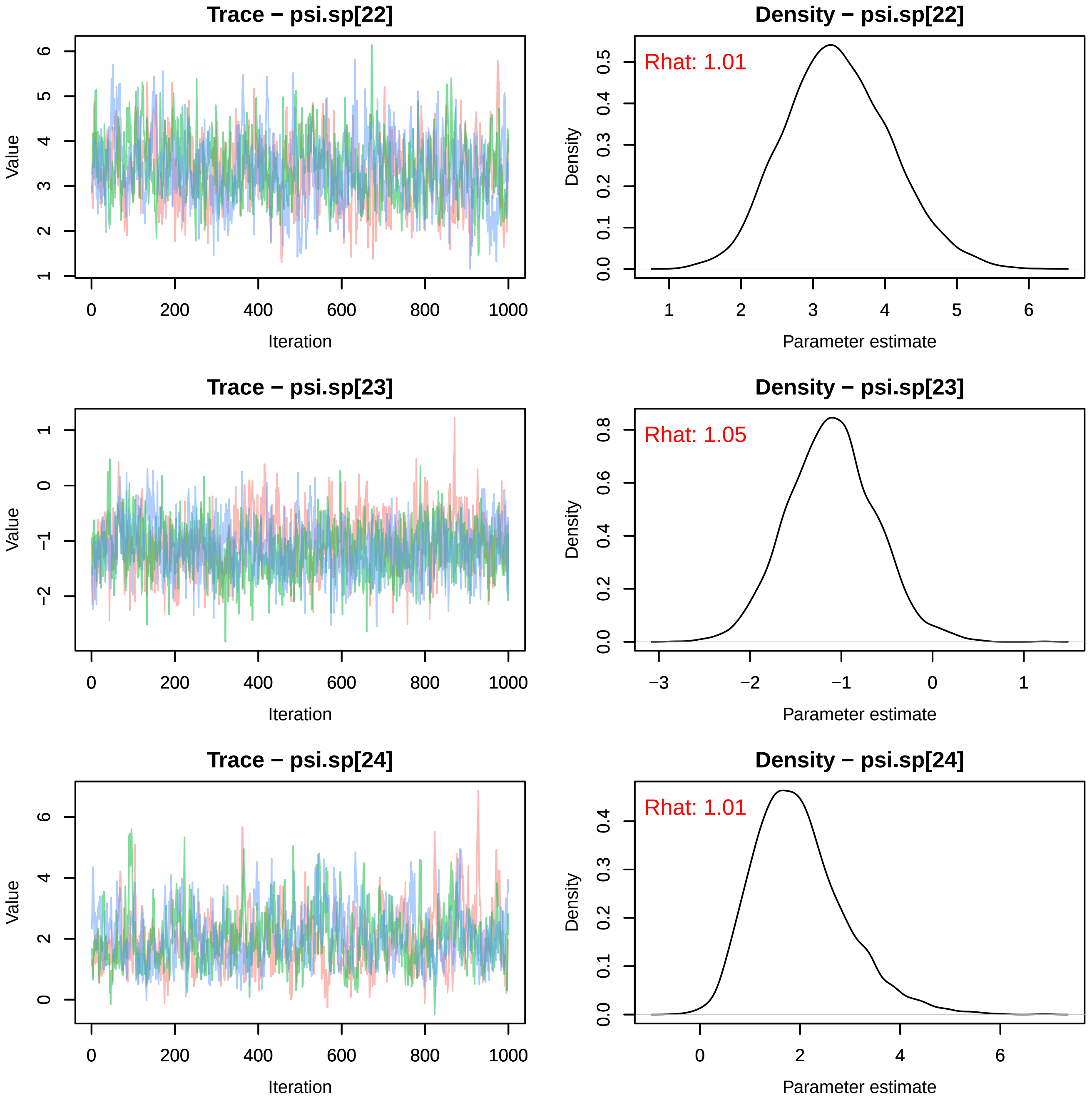
Trace − psi.sp[22]
Density − psi.sp[22]
Rhat: 1.01
Trace − psi.sp[23]
Density − psi.sp[23]
Rhat: 1.05
Trace − psi.sp[24]
Density − psi.sp[24]
Rhat: 1.01
Value
Iteration
Density
Parameter estimate

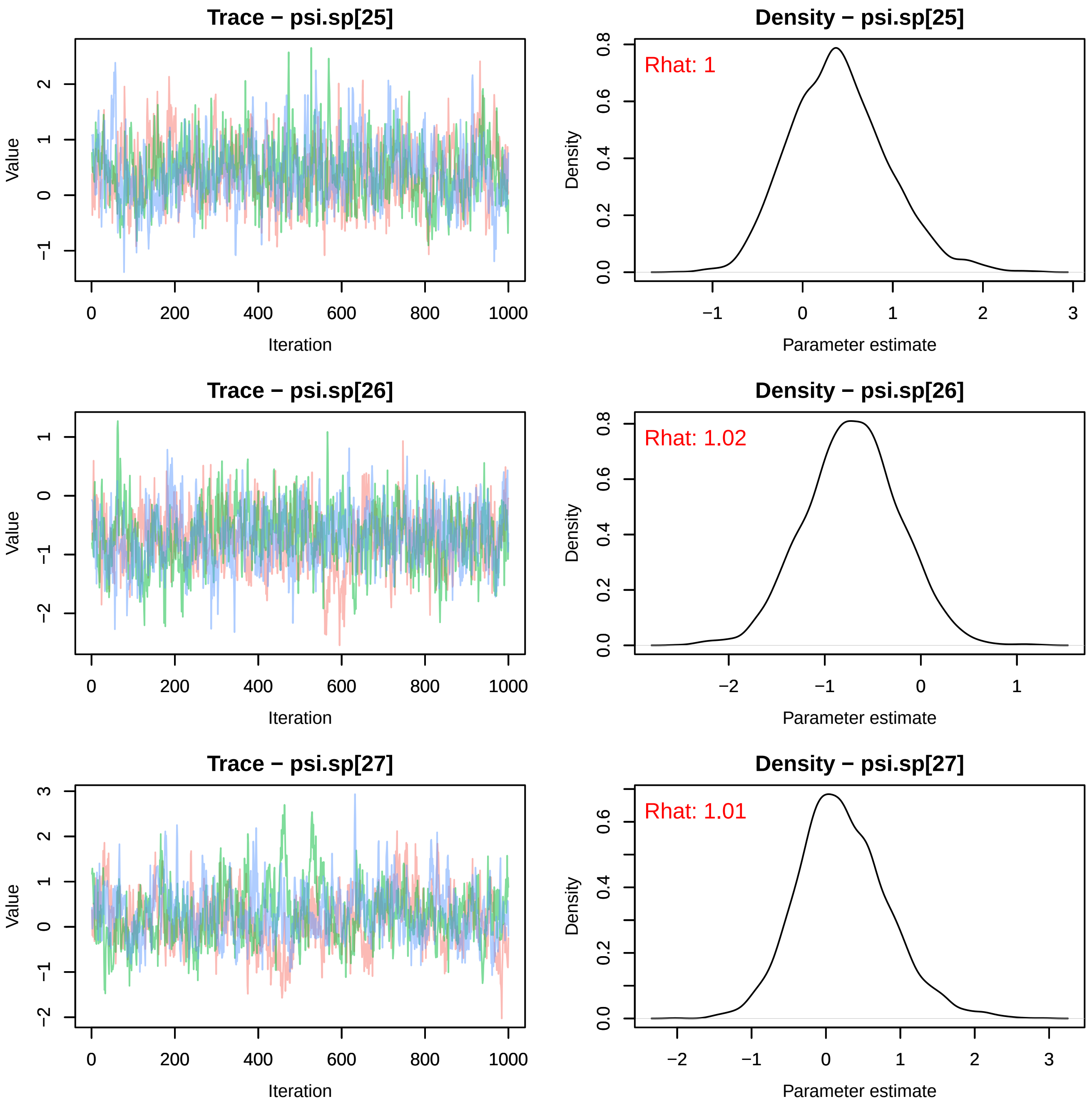
Trace − psi.sp[25]
Value
Iteration
Density − psi.sp[25]
Rhat: 1
Density
Parameter estimate
Trace − psi.sp[26]
Value
Iteration
Density − psi.sp[26]
Rhat: 1.02
Density
Parameter estimate
Trace − psi.sp[27]
Value
Iteration
Density − psi.sp[27]
Rhat: 1.01
Density
Parameter estimate

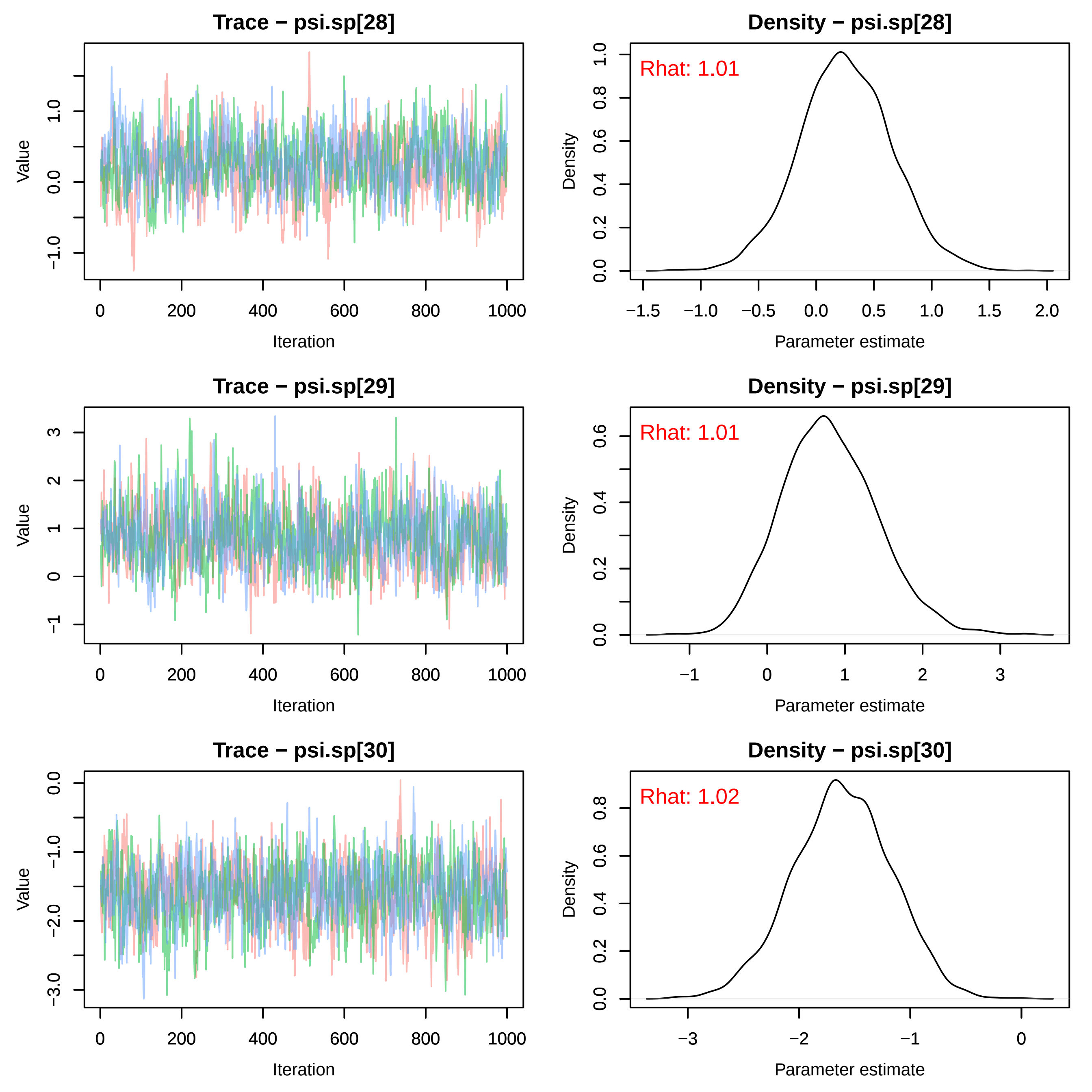

Trace − psi.sp[28]
Value
Iteration
Density − psi.sp[28]
Rhat: 1.01
Density
Parameter estimate
Trace − psi.sp[29]
Value
Iteration
Density − psi.sp[29]
Rhat: 1.01
Density
Parameter estimate
Trace − psi.sp[30]
Value
Iteration
Density − psi.sp[30]
Rhat: 1.02
Density
Parameter estimate

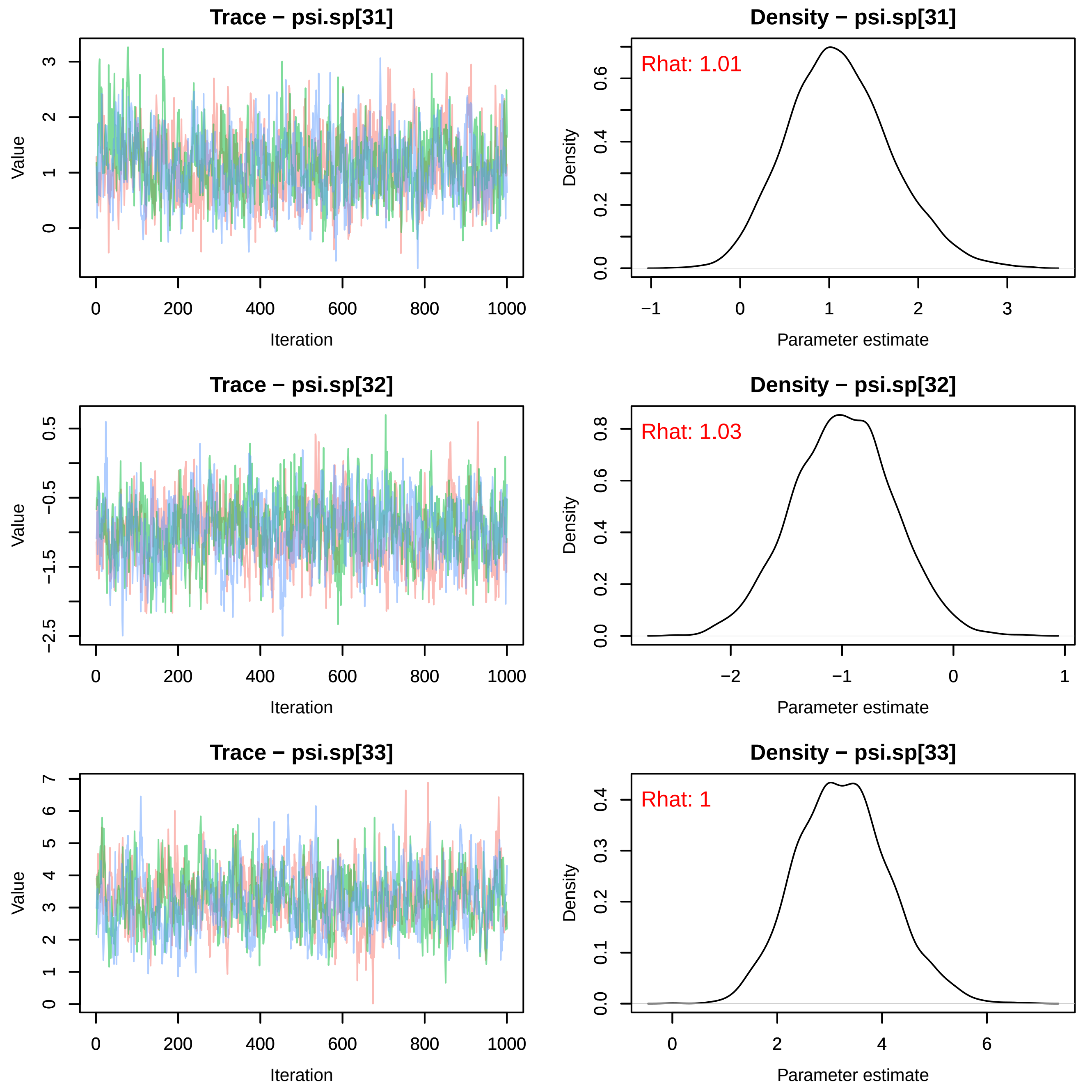

Trace − psi.sp[31]
Density − psi.sp[31]
Rhat: 1.01
Trace − psi.sp[32]
Density − psi.sp[32]
Rhat: 1.03
Trace − psi.sp[33]
Density − psi.sp[33]
Rhat: 1
Value
Iteration
Density
Parameter estimate

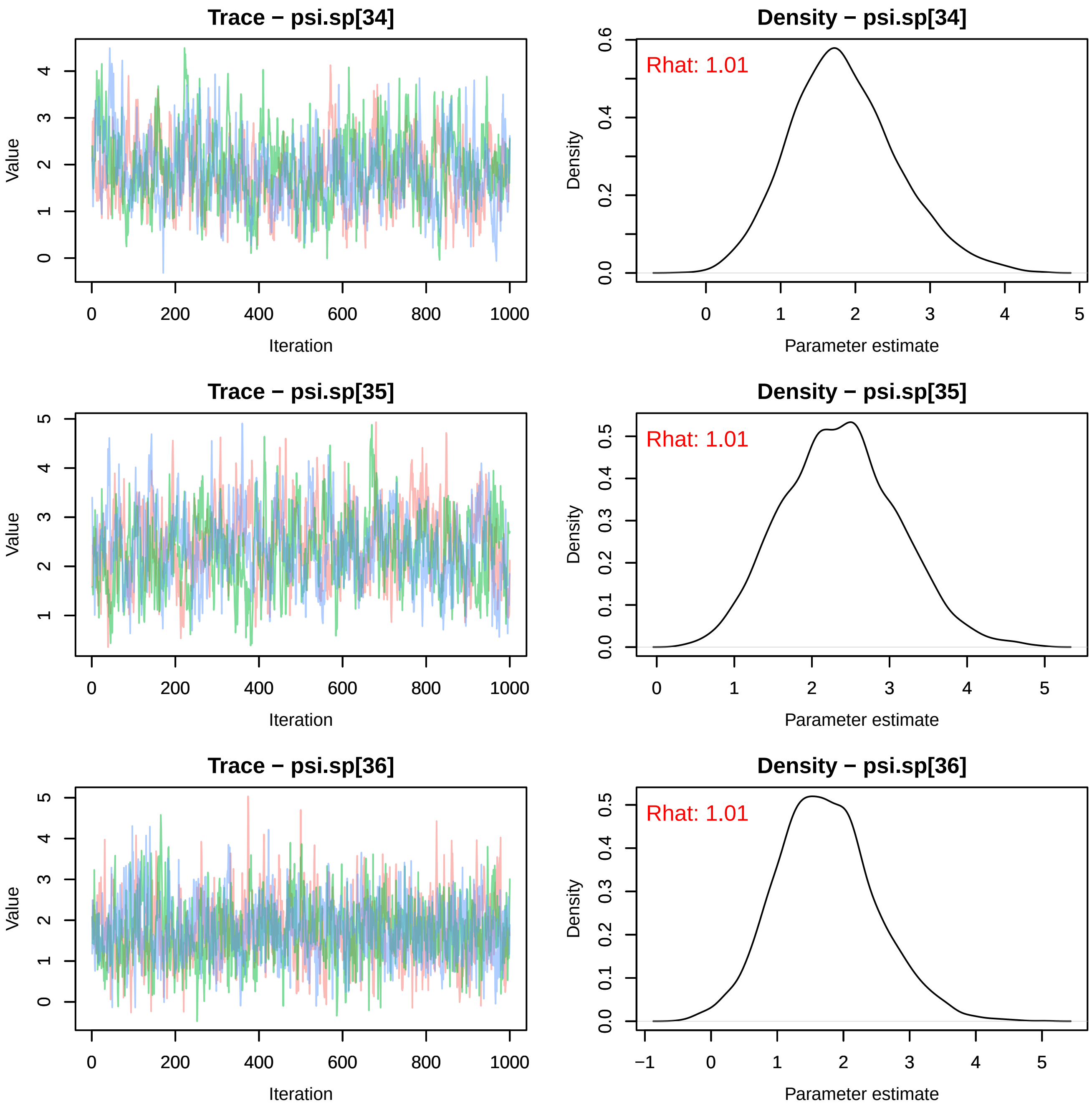
Trace − psi.sp[34]
Value
Iteration
Density − psi.sp[34]
Rhat: 1.01
Density
Parameter estimate
Trace − psi.sp[35]
Value
Iteration
Density − psi.sp[35]
Rhat: 1.01
Density
Parameter estimate
Trace − psi.sp[36]
Value
Iteration
Density − psi.sp[36]
Rhat: 1.01
Density
Parameter estimate

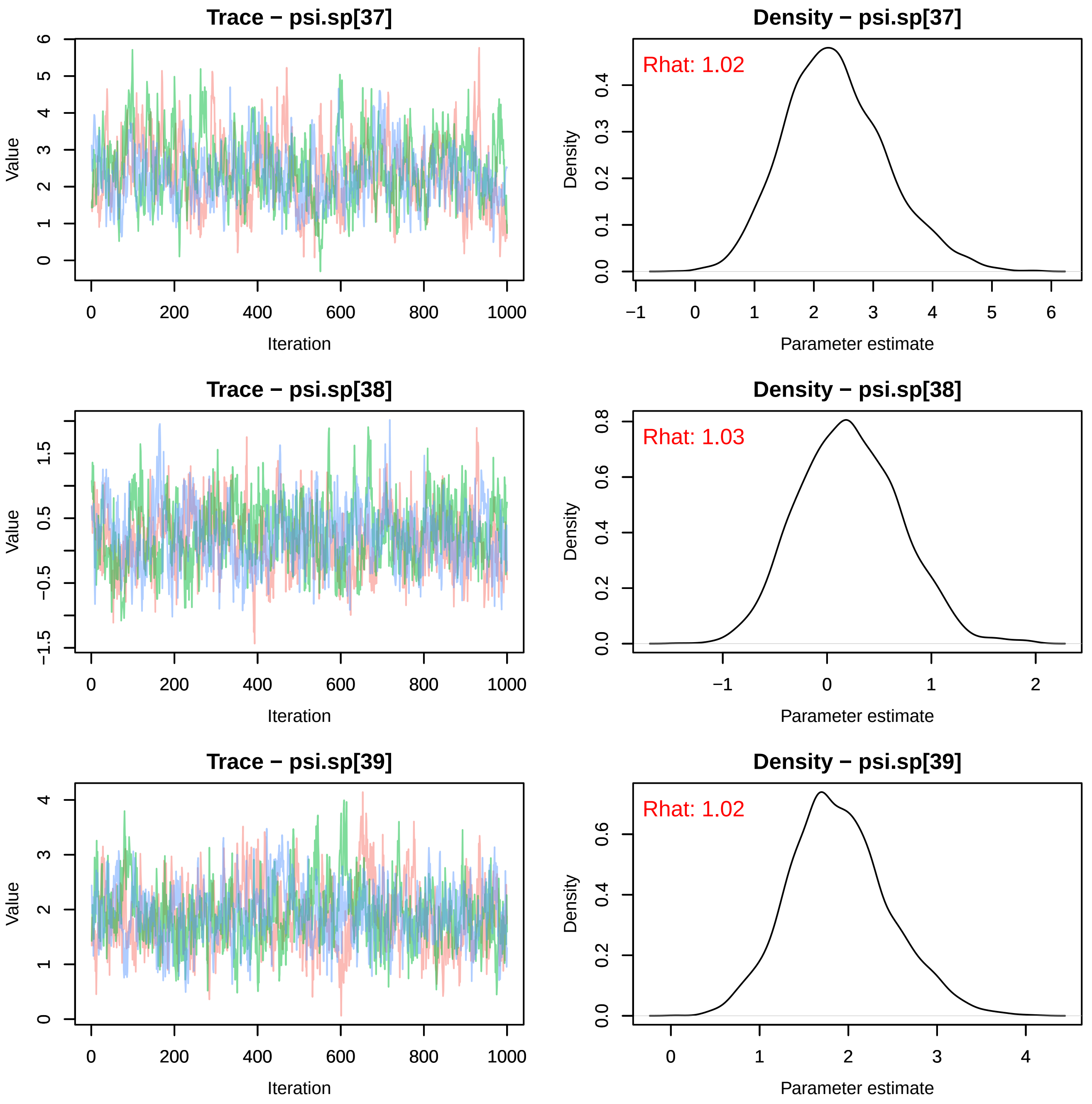

Trace − psi.sp[37]
Value
Iteration
Density − psi.sp[37]
Rhat: 1.02
Density
Parameter estimate
Trace − psi.sp[38]
Value
Iteration
Density − psi.sp[38]
Rhat: 1.03
Density
Parameter estimate
Trace − psi.sp[39]
Value
Iteration
Density − psi.sp[39]
Rhat: 1.02
Density
Parameter estimate

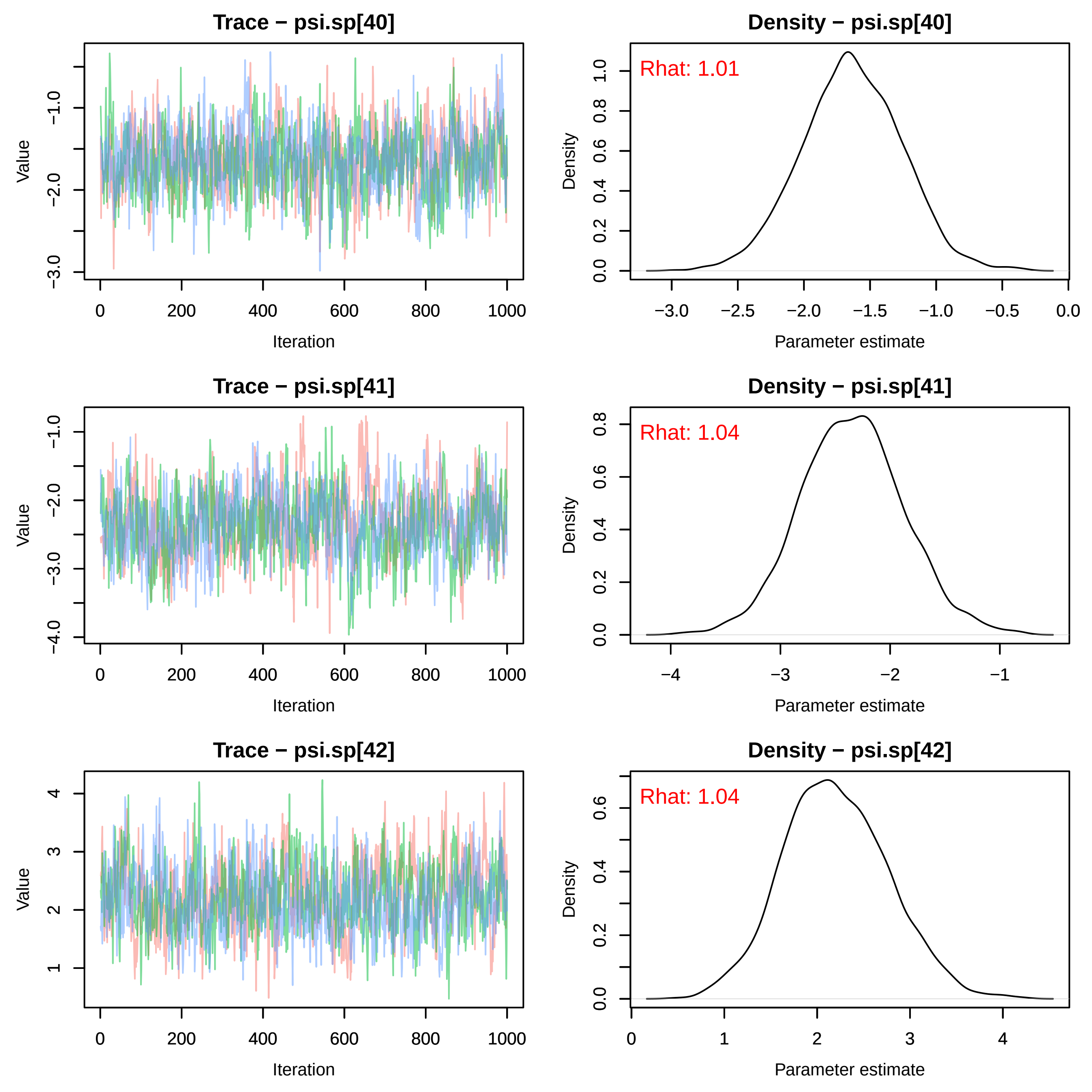
Trace − psi.sp[40]
Value
Iteration
Density − psi.sp[40]
Rhat: 1.01
Density
Parameter estimate
Trace − psi.sp[41]
Value
Iteration
Density − psi.sp[41]
Rhat: 1.04
Density
Parameter estimate
Trace − psi.sp[42]
Value
Iteration
Density − psi.sp[42]
Rhat: 1.04
Density
Parameter estimate

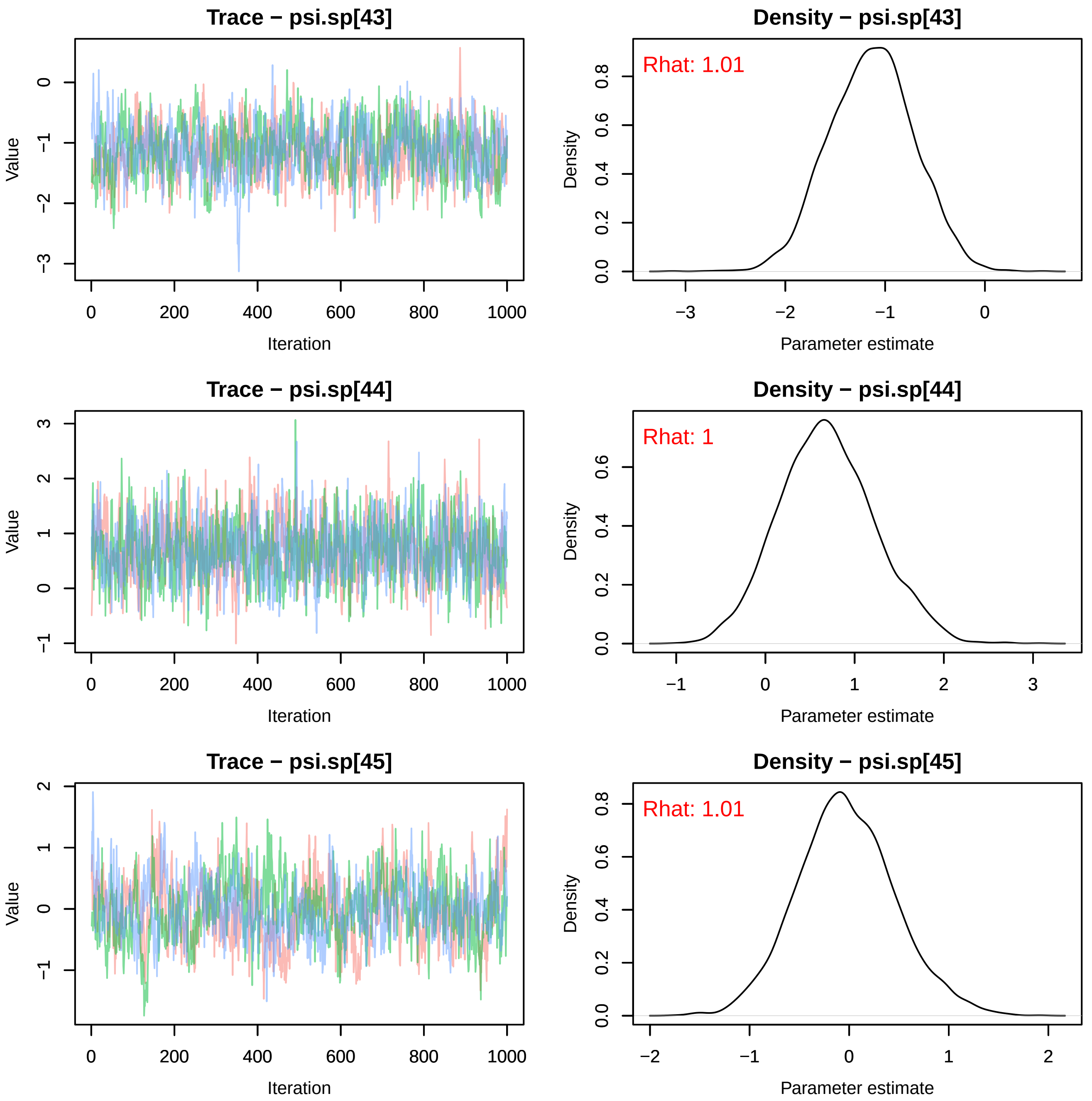
Trace − psi.sp[43]
Value
Iteration
Density − psi.sp[43]
Rhat: 1.01
Density
Parameter estimate
Trace − psi.sp[44]
Value
Iteration
Density − psi.sp[44]
Rhat: 1
Density
Parameter estimate
Trace − psi.sp[45]
Value
Iteration
Density − psi.sp[45]
Rhat: 1.01
Density
Parameter estimate

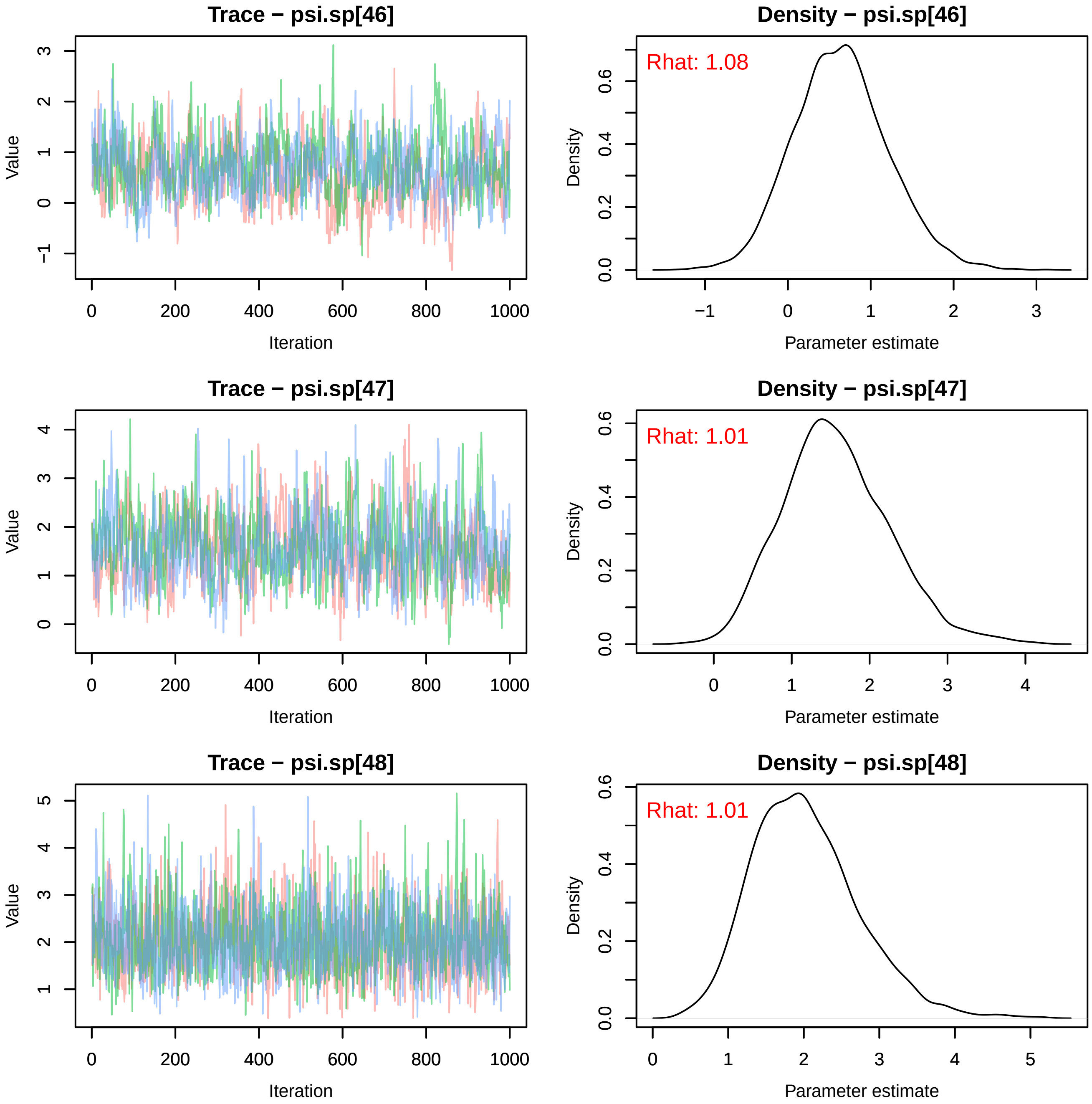
Trace − psi.sp[46]
Value
Iteration
Density − psi.sp[46]
Rhat: 1.08
Density
Parameter estimate
Trace − psi.sp[47]
Value
Iteration
Density − psi.sp[47]
Rhat: 1.01
Density
Parameter estimate
Trace − psi.sp[48]
Value
Iteration
Density − psi.sp[48]
Rhat: 1.01
Density
Parameter estimate

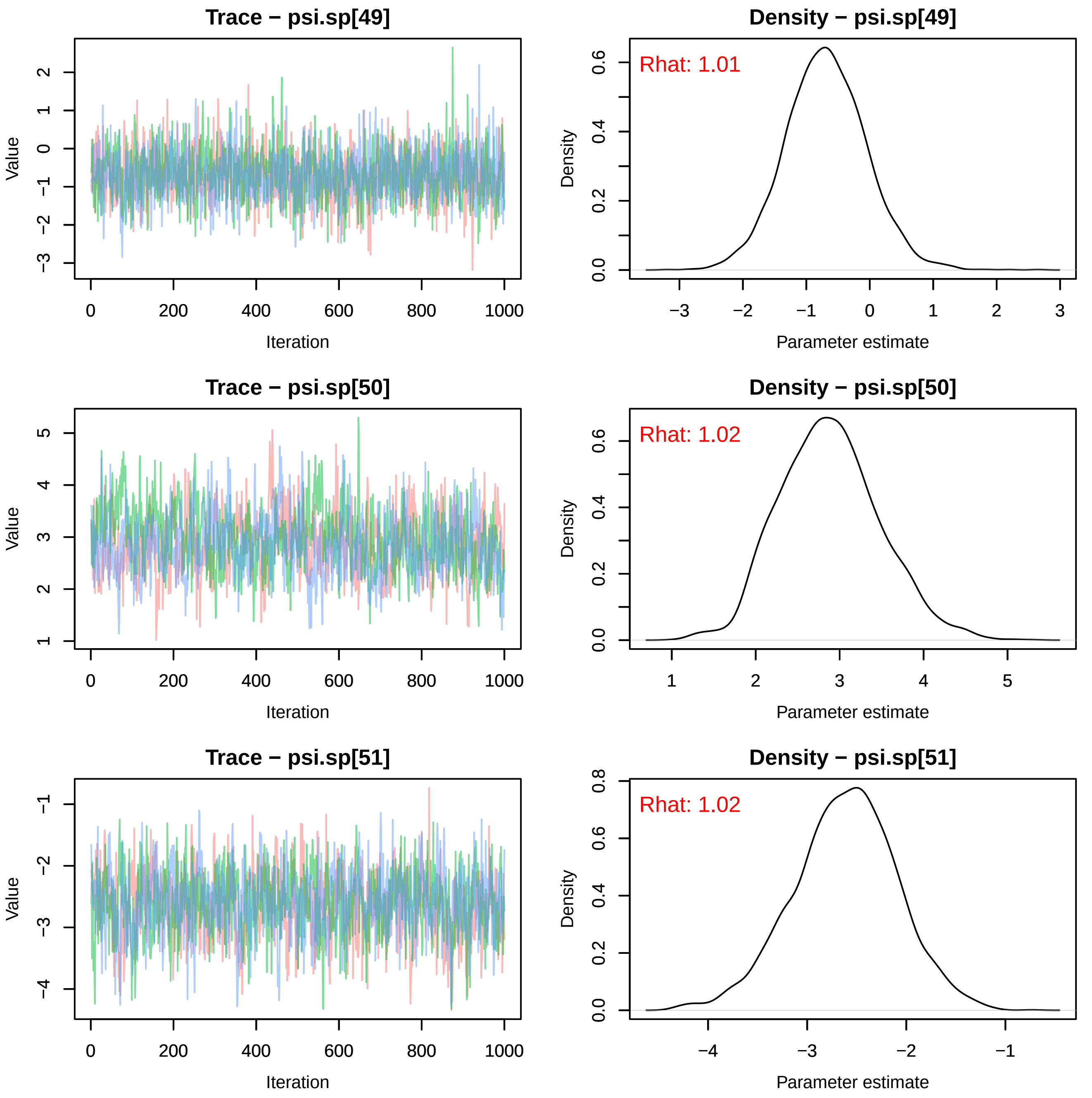

Trace − psi.sp[49]
Value
Iteration
Density − psi.sp[49]
Rhat: 1.01
Density
Parameter estimate
Trace − psi.sp[50]
Value
Iteration
Density − psi.sp[50]
Rhat: 1.02
Density
Parameter estimate
Trace − psi.sp[51]
Value
Iteration
Density − psi.sp[51]
Rhat: 1.02
Density
Parameter estimate

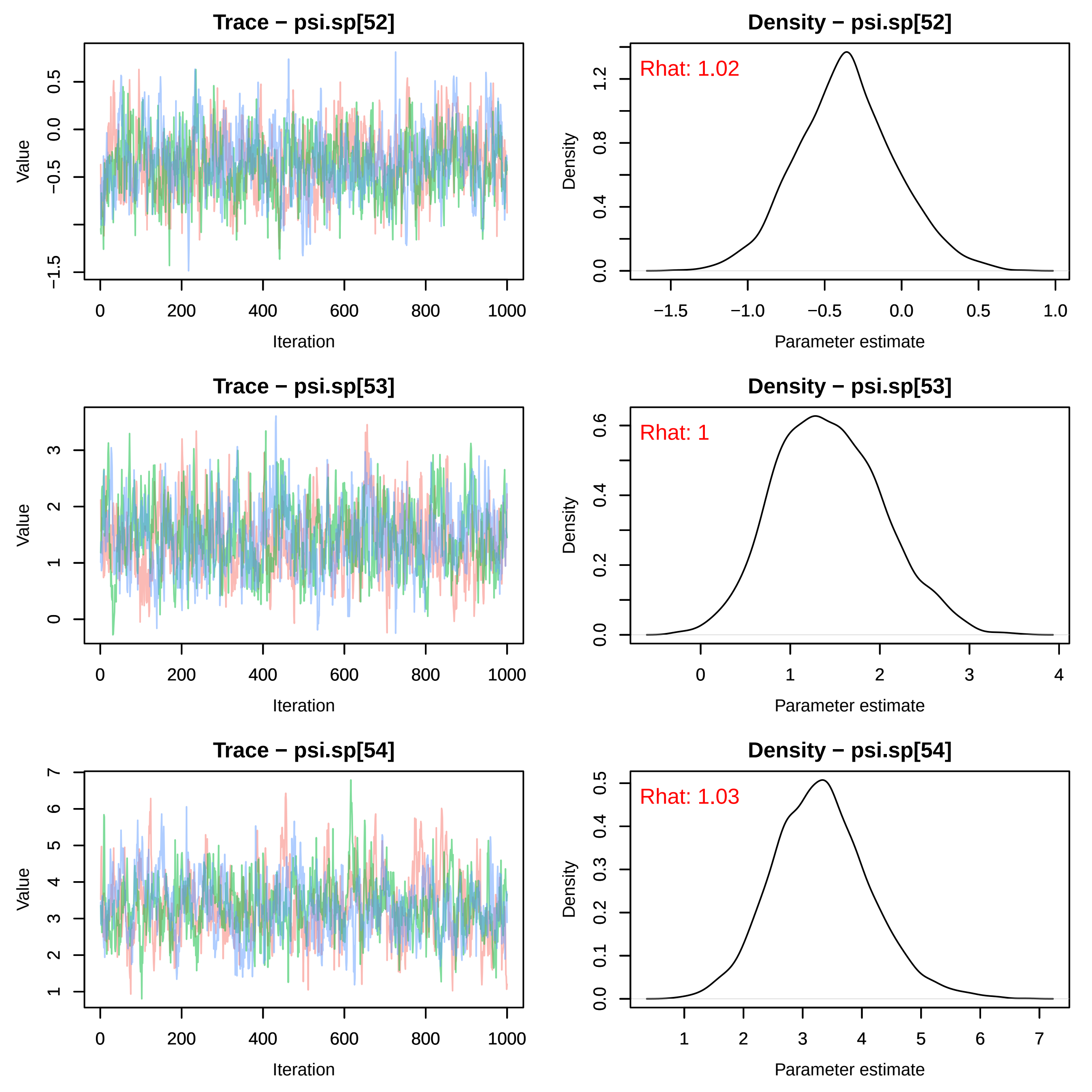
Trace − psi.sp[52]
Value
Iteration
Density − psi.sp[52]
Rhat: 1.02
Density
Parameter estimate
Trace − psi.sp[53]
Value
Iteration
Density − psi.sp[53]
Rhat: 1
Density
Parameter estimate
Trace − psi.sp[54]
Value
Iteration
Density − psi.sp[54]
Rhat: 1.03
Density
Parameter estimate

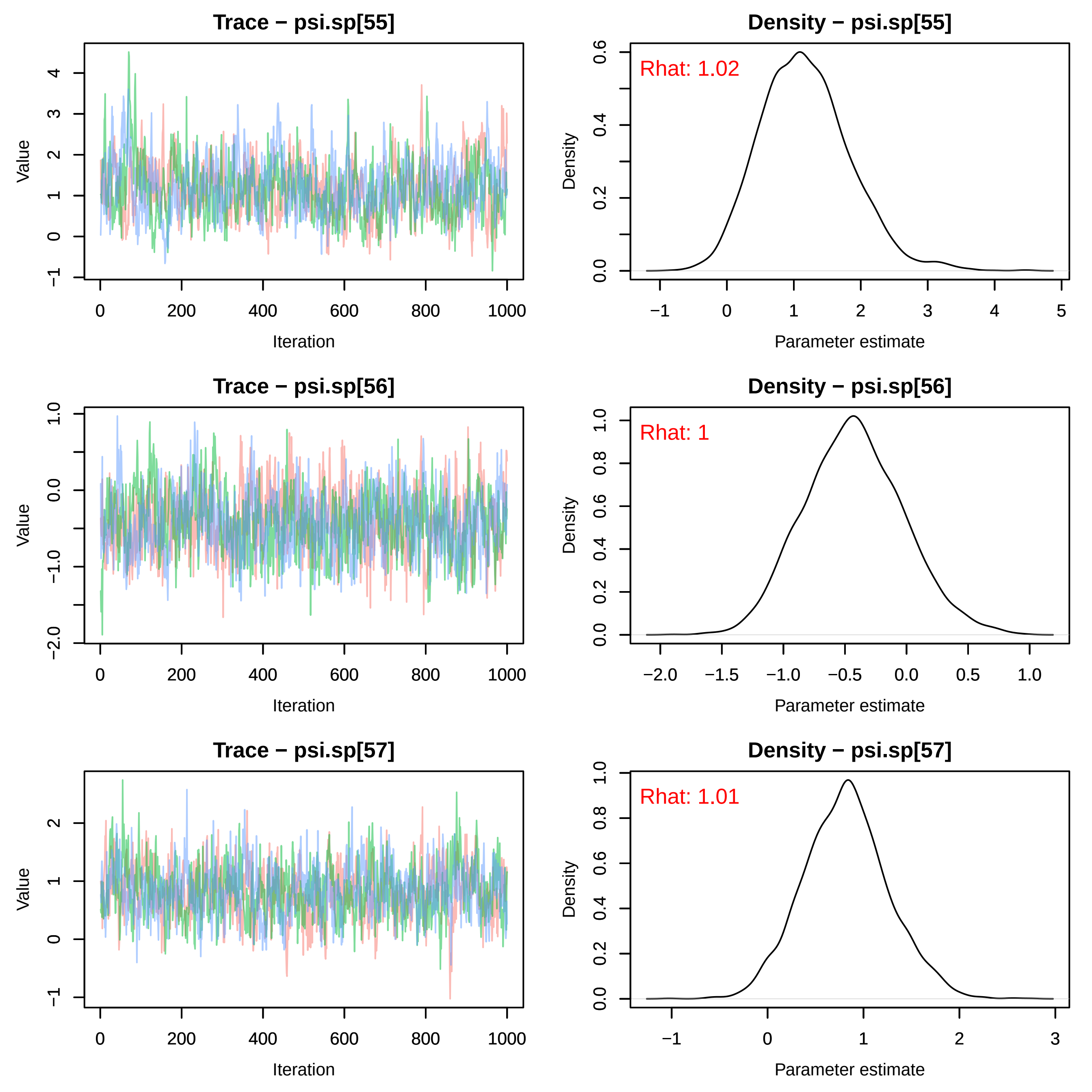
Trace − psi.sp[55]
Density − psi.sp[55]
Rhat: 1.02
Trace − psi.sp[56]
Density − psi.sp[56]
Rhat: 1
Trace − psi.sp[57]
Density − psi.sp[57]
Rhat: 1.01
Value
Iteration
Density
Parameter estimate

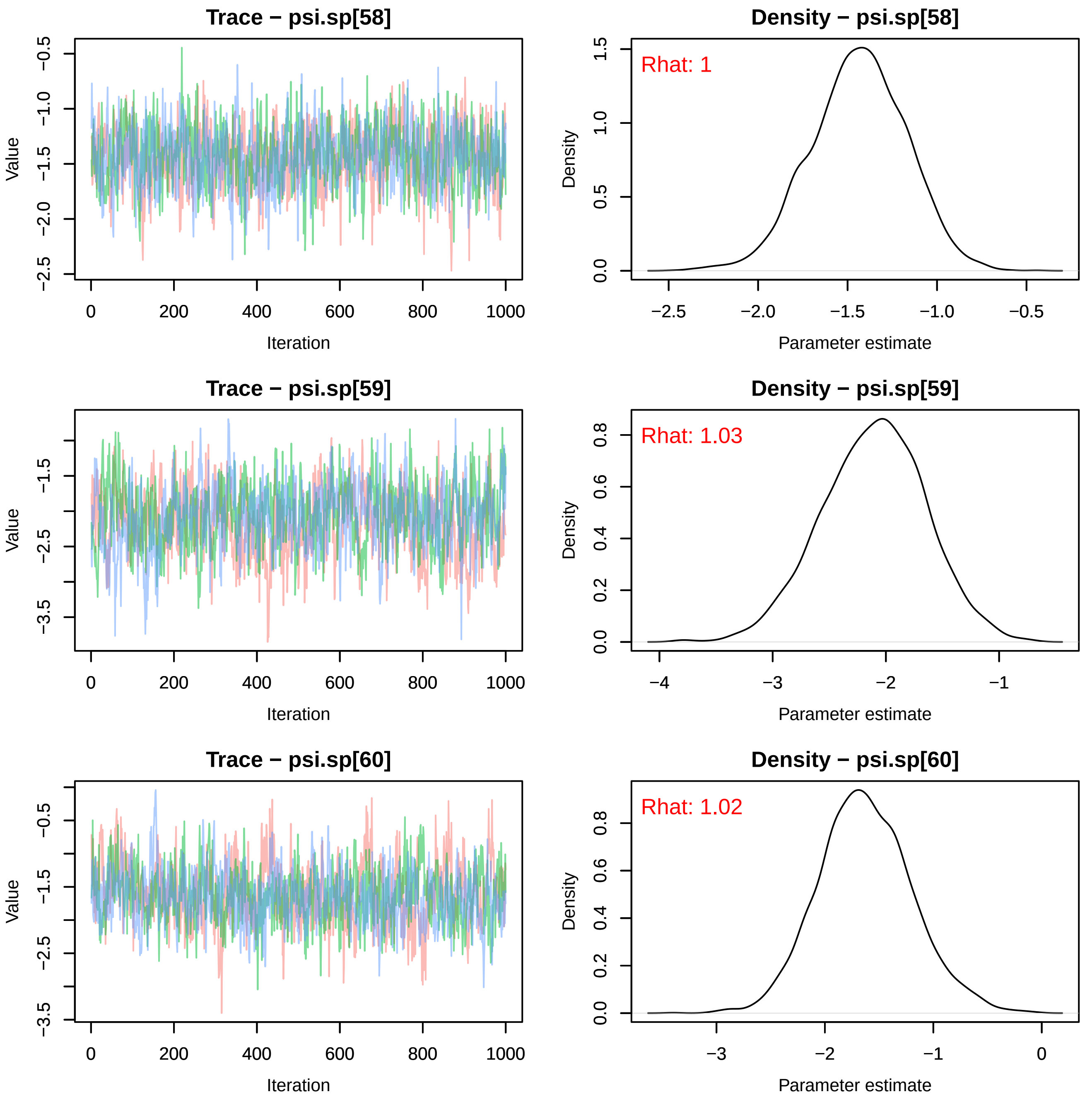
Trace − psi.sp[58]
Value
Iteration
Density − psi.sp[58]
Rhat: 1
Density
Parameter estimate
Trace − psi.sp[59]
Value
Iteration
Density − psi.sp[59]
Rhat: 1.03
Density
Parameter estimate
Trace − psi.sp[60]
Value
Iteration
Density − psi.sp[60]
Rhat: 1.02
Density
Parameter estimate

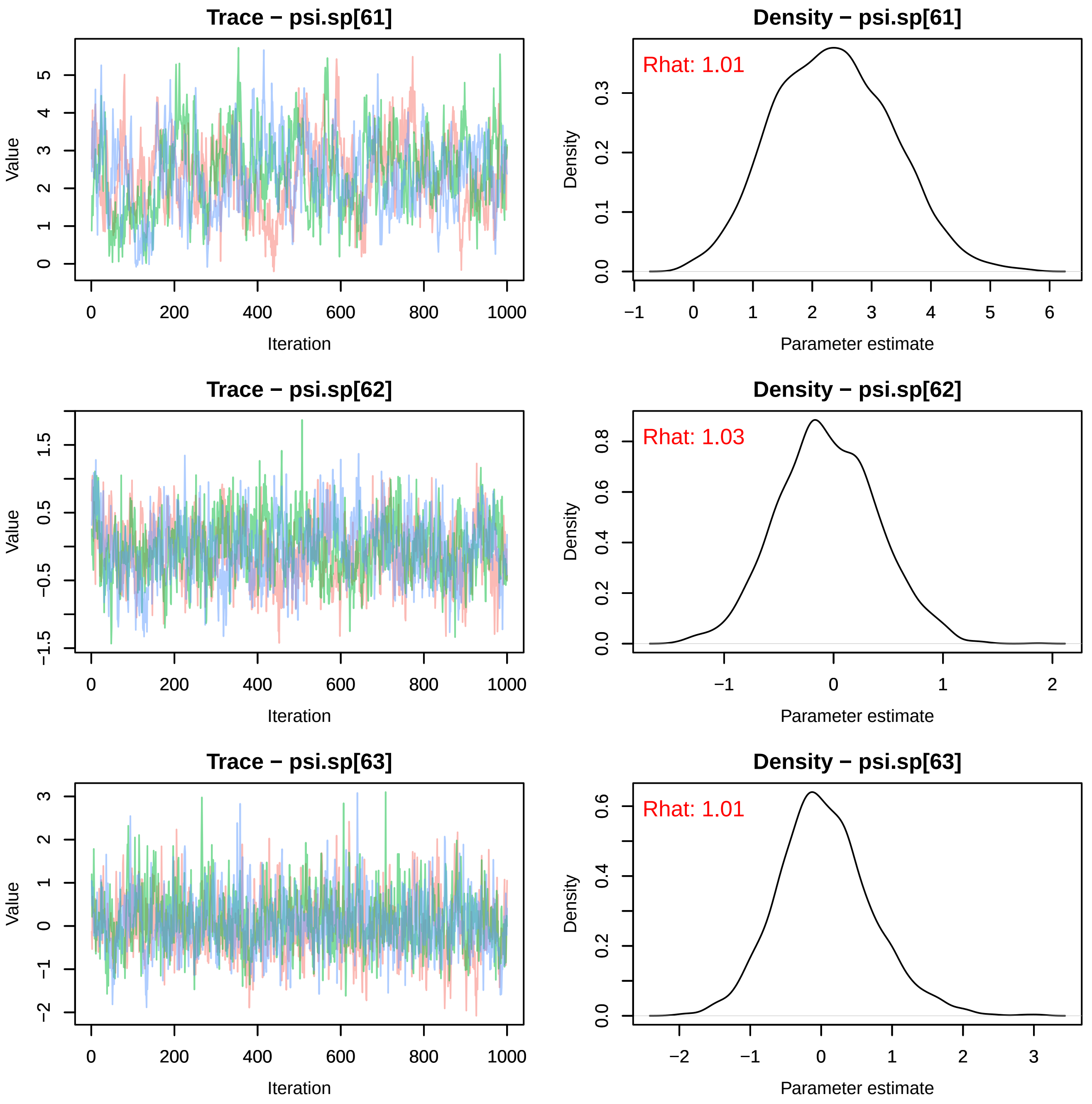
Trace − psi.sp[61]
Value
Iteration
Density − psi.sp[61]
Rhat: 1.01
Density
Parameter estimate
Trace − psi.sp[62]
Value
Iteration
Density − psi.sp[62]
Rhat: 1.03
Density
Parameter estimate
Trace − psi.sp[63]
Value
Iteration
Density − psi.sp[63]
Rhat: 1.01
Density
Parameter estimate

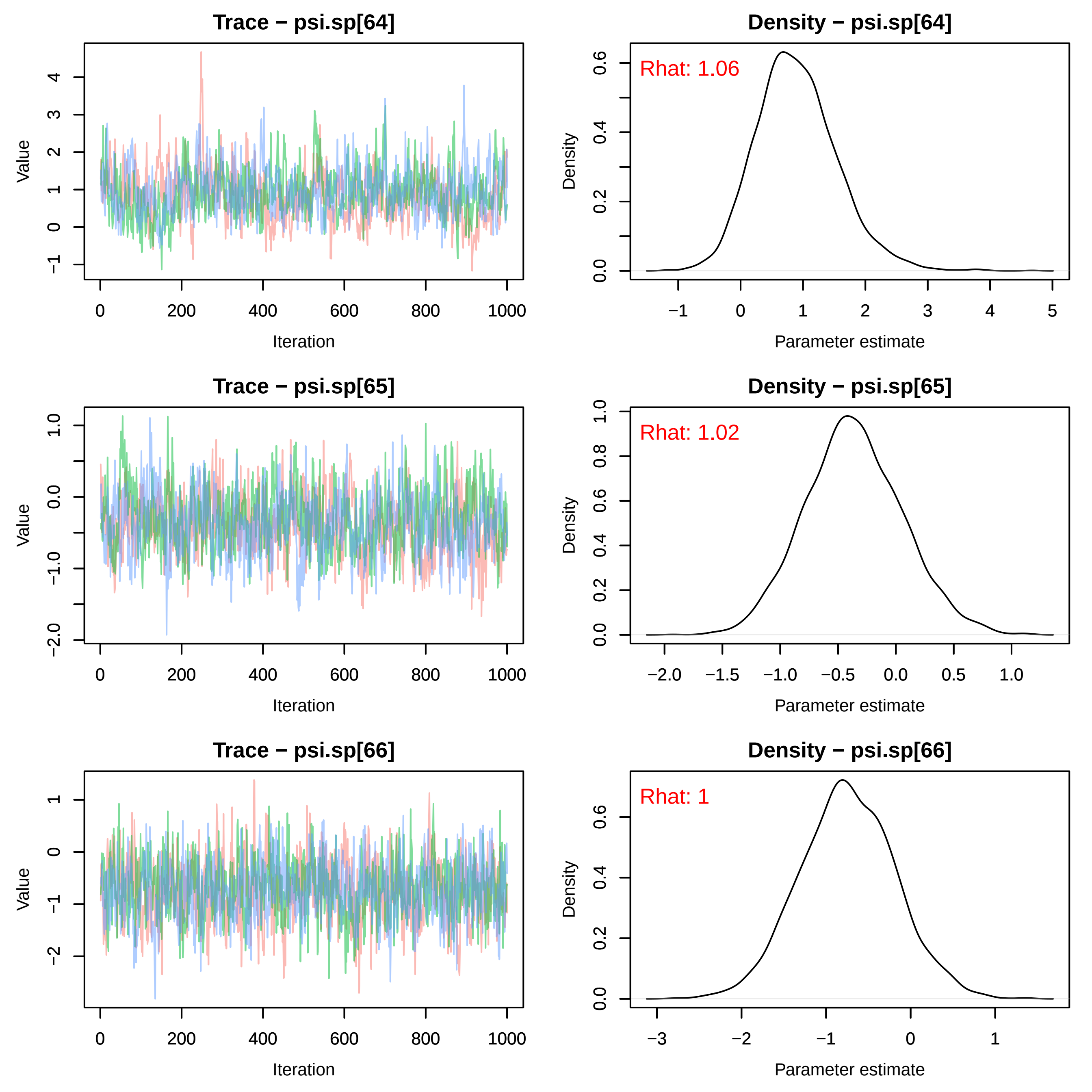
Trace − psi.sp[64]
Value
Iteration
Density − psi.sp[64]
Rhat: 1.06
Density
Parameter estimate
Trace − psi.sp[65]
Value
Iteration
Density − psi.sp[65]
Rhat: 1.02
Density
Parameter estimate
Trace − psi.sp[66]
Value
Iteration
Density − psi.sp[66]
Rhat: 1
Density
Parameter estimate

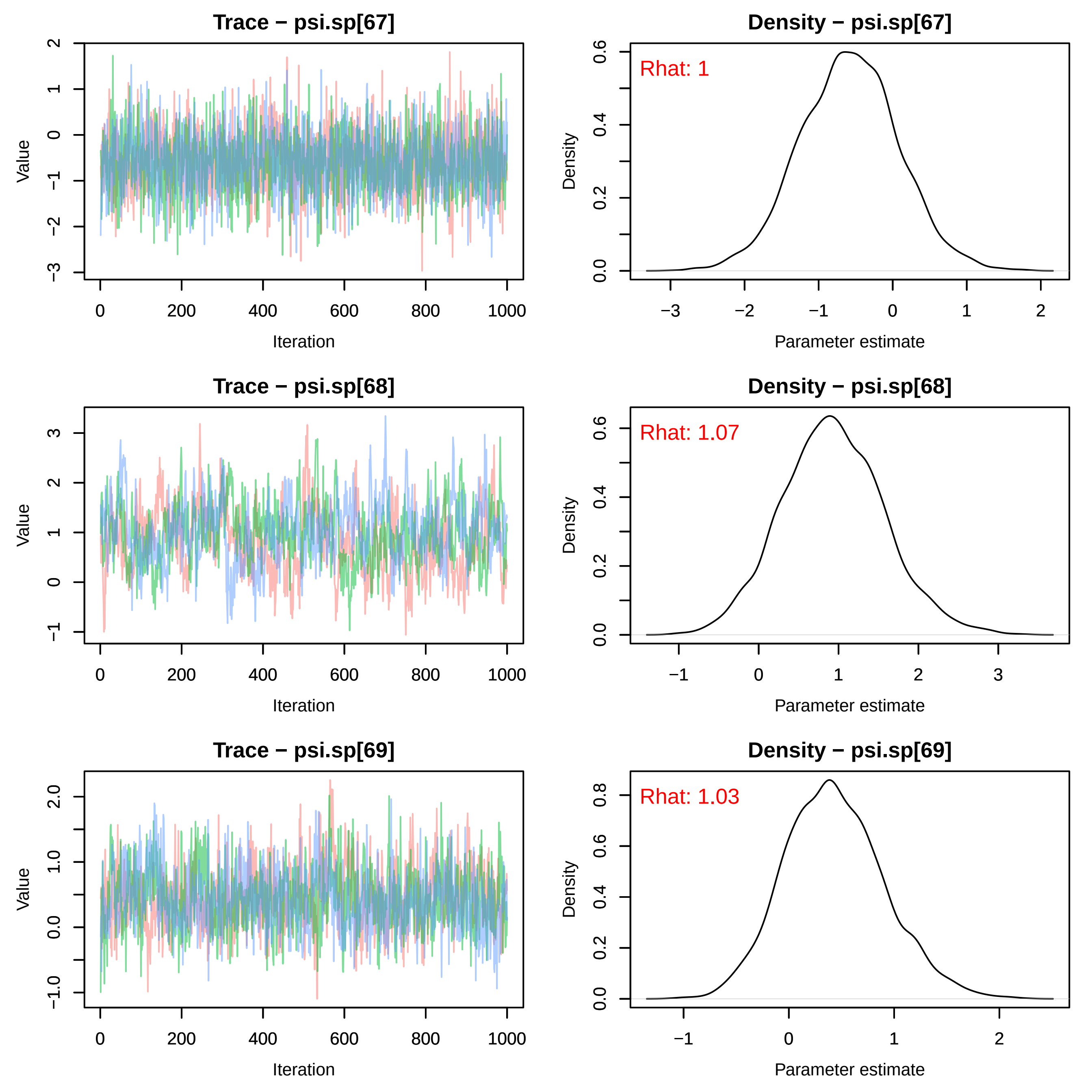
Trace − psi.sp[67]
Value
Iteration
Density − psi.sp[67]
Rhat: 1
Density
Parameter estimate
Trace − psi.sp[68]
Value
Iteration
Density − psi.sp[68]
Rhat: 1.07
Density
Parameter estimate
Trace − psi.sp[69]
Value
Iteration
Density − psi.sp[69]
Rhat: 1.03
Density
Parameter estimate

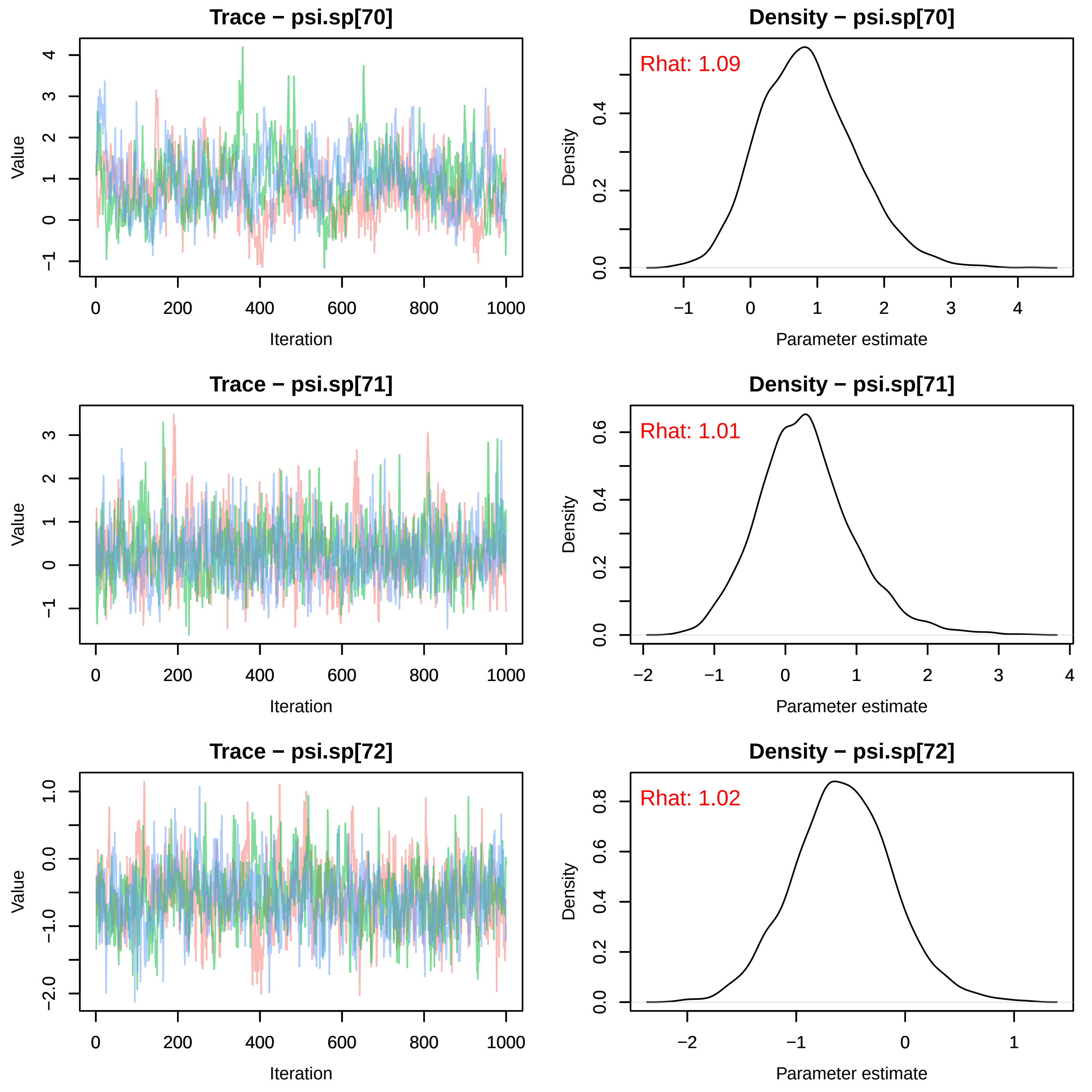

Trace − psi.sp[70]
Value
Iteration
Density − psi.sp[70]
Rhat: 1.09
Density
Parameter estimate
Trace − psi.sp[71]
Value
Iteration
Density − psi.sp[71]
Rhat: 1.01
Density
Parameter estimate
Trace − psi.sp[72]
Value
Iteration
Density − psi.sp[72]
Rhat: 1.02
Density
Parameter estimate

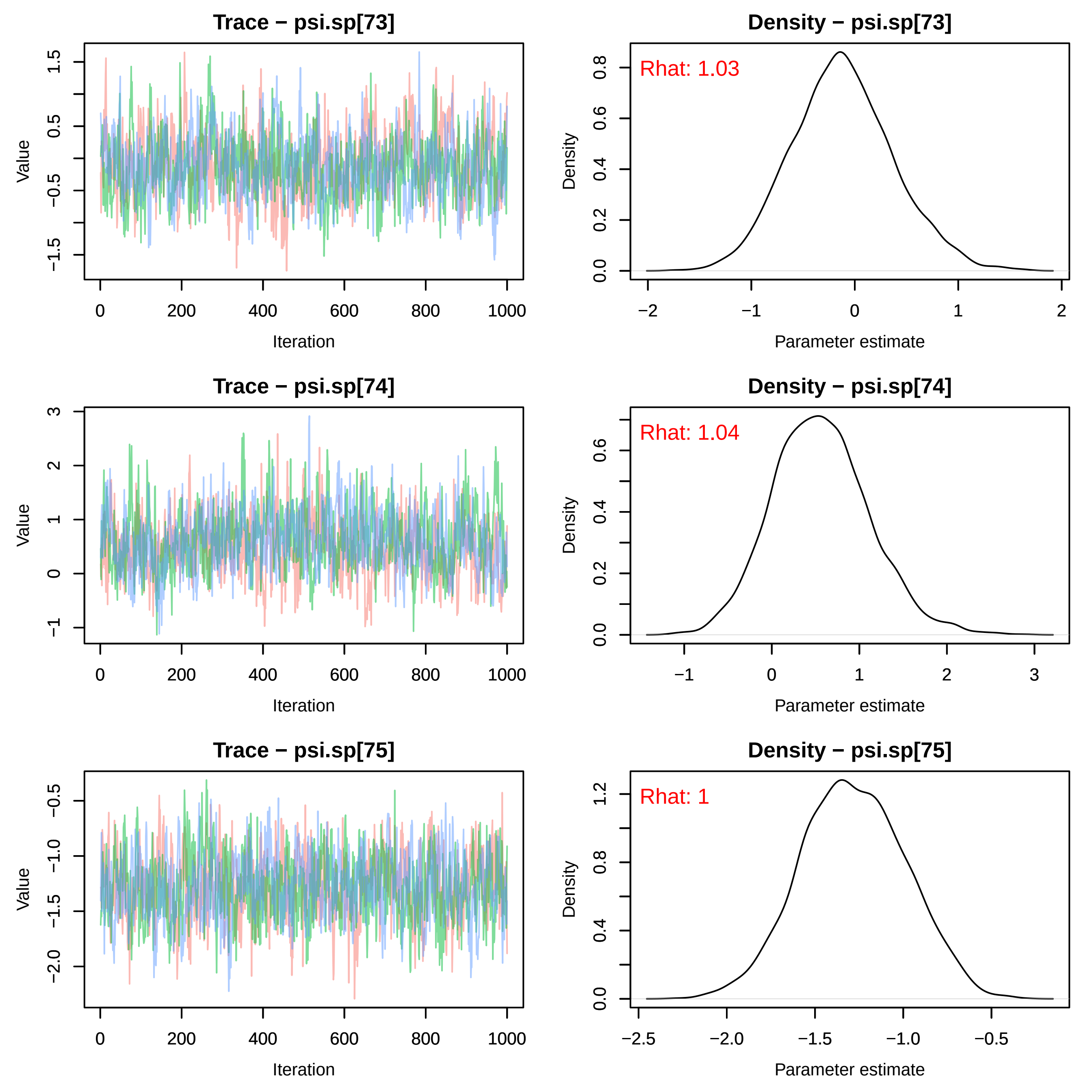

Trace − psi.sp[73]
Value
Iteration
Density − psi.sp[73]
Rhat: 1.03
Density
Parameter estimate
Trace − psi.sp[74]
Value
Iteration
Density − psi.sp[74]
Rhat: 1.04
Density
Parameter estimate
Trace − psi.sp[75]
Value
Iteration
Density − psi.sp[75]
Rhat: 1
Density
Parameter estimate

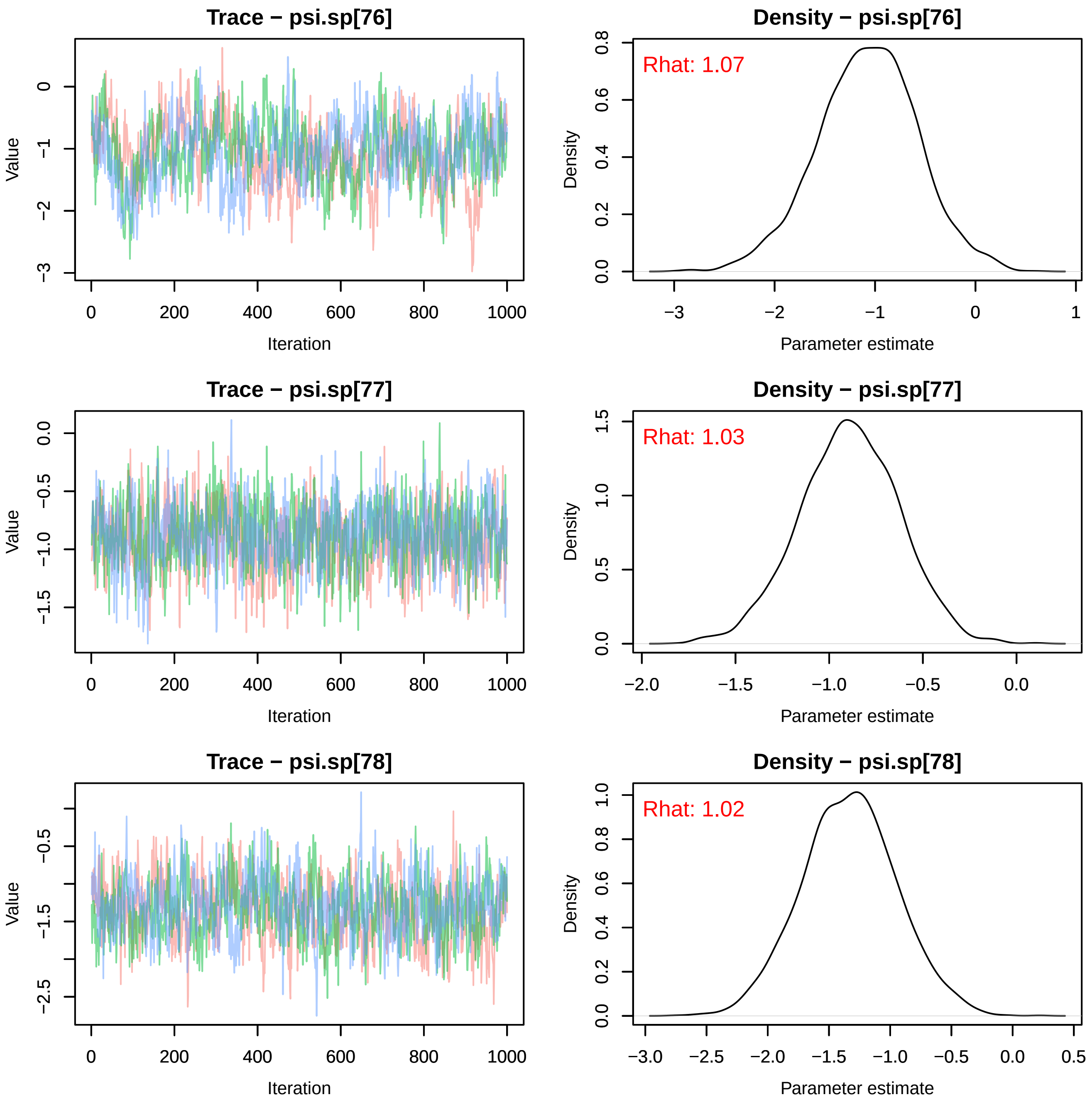
Trace − psi.sp[76]
Density − psi.sp[76]
Rhat: 1.07
Trace − psi.sp[77]
Density − psi.sp[77]
Rhat: 1.03
Trace − psi.sp[78]
Density − psi.sp[78]
Rhat: 1.02
Value
Iteration
Density
Parameter estimate

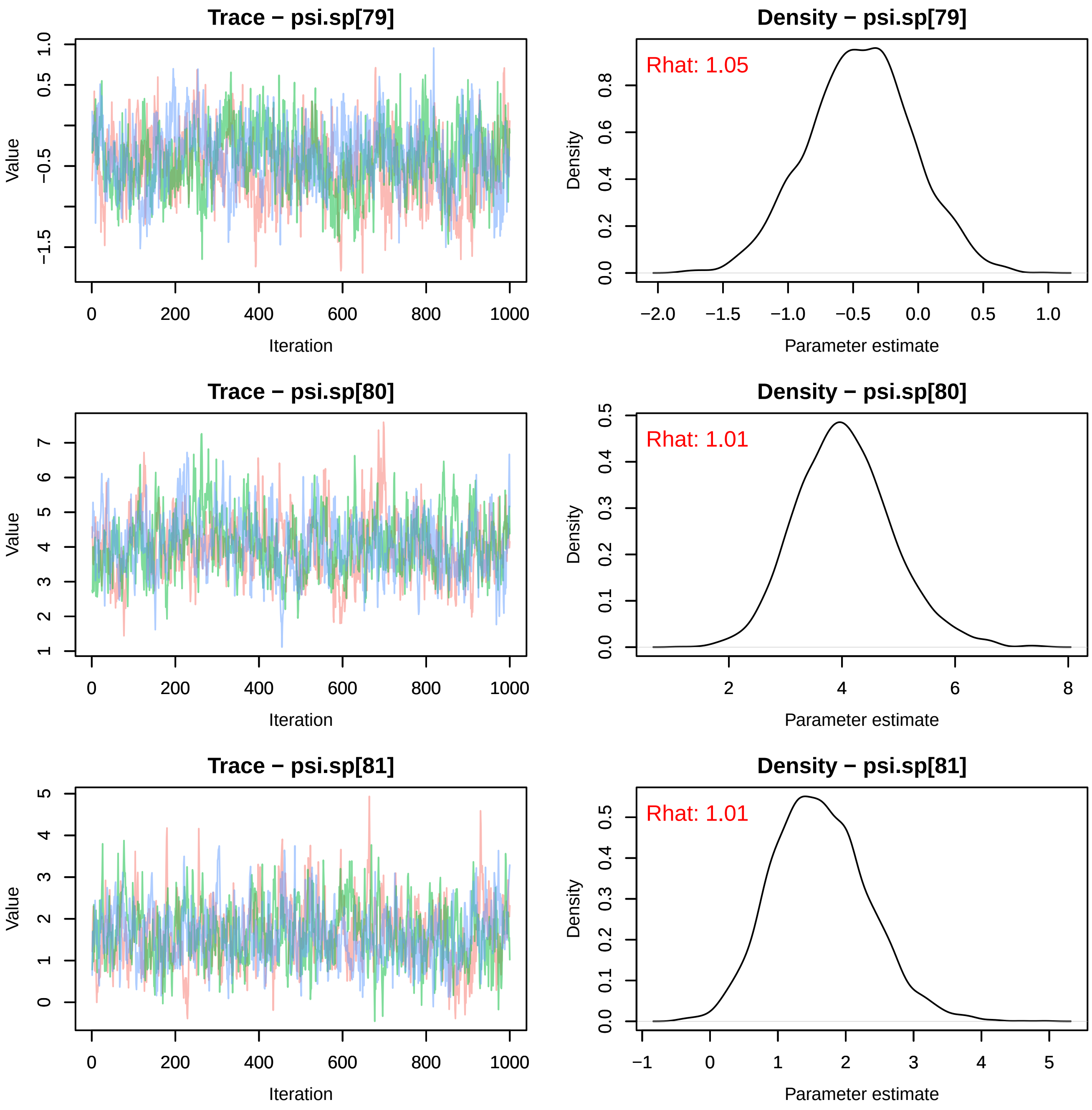
Trace − psi.sp[79]
Value
Iteration
Density − psi.sp[79]
Rhat: 1.05
Density
Parameter estimate
Trace − psi.sp[80]
Value
Iteration
Density − psi.sp[80]
Rhat: 1.01
Density
Parameter estimate
Trace − psi.sp[81]
Value
Iteration
Density − psi.sp[81]
Rhat: 1.01
Density
Parameter estimate

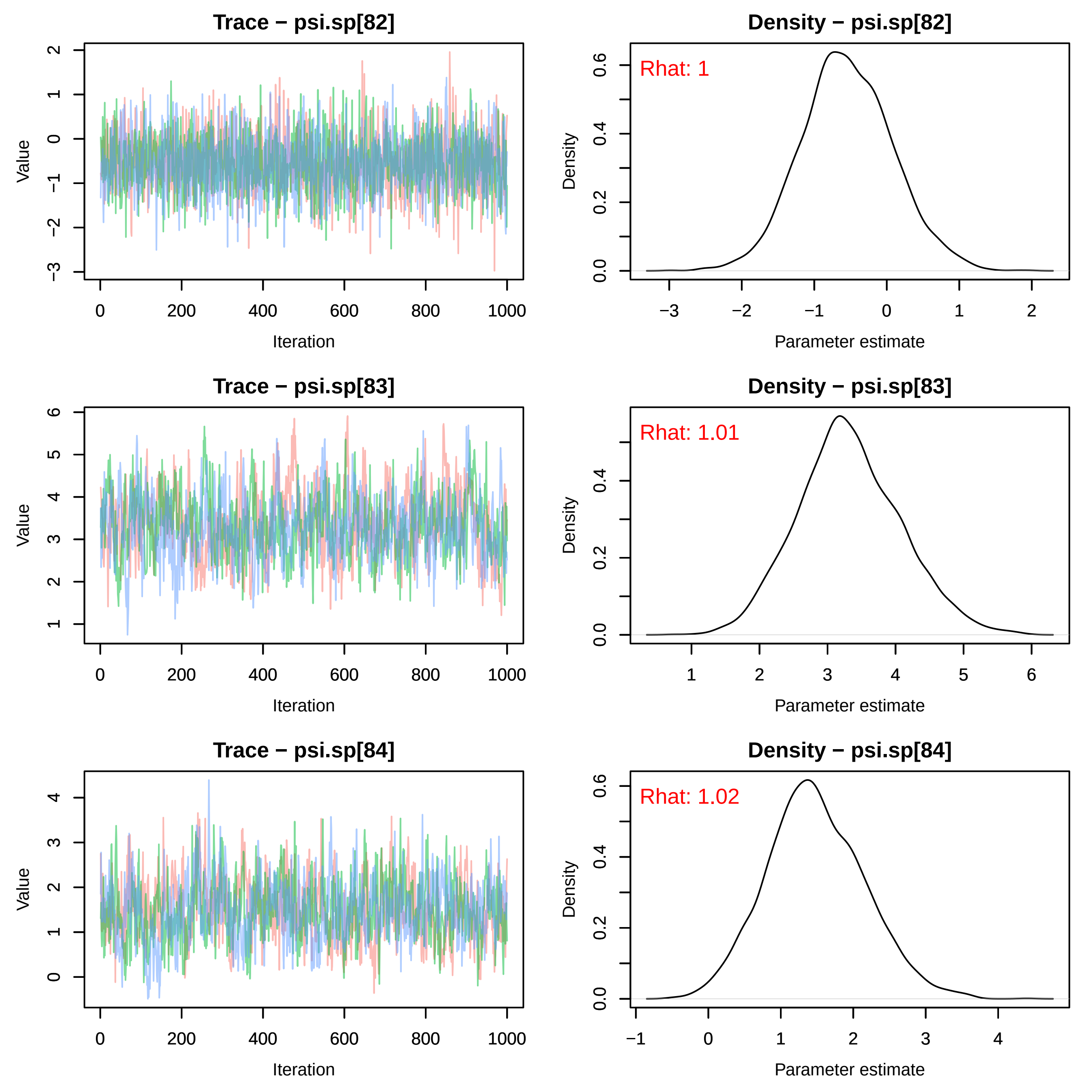
Trace − psi.sp[82]
Value
Iteration
Density − psi.sp[82]
Rhat: 1
Density
Parameter estimate
Trace − psi.sp[83]
Value
Iteration
Density − psi.sp[83]
Rhat: 1.01
Density
Parameter estimate
Trace − psi.sp[84]
Value
Iteration
Density − psi.sp[84]
Rhat: 1.02
Density
Parameter estimate

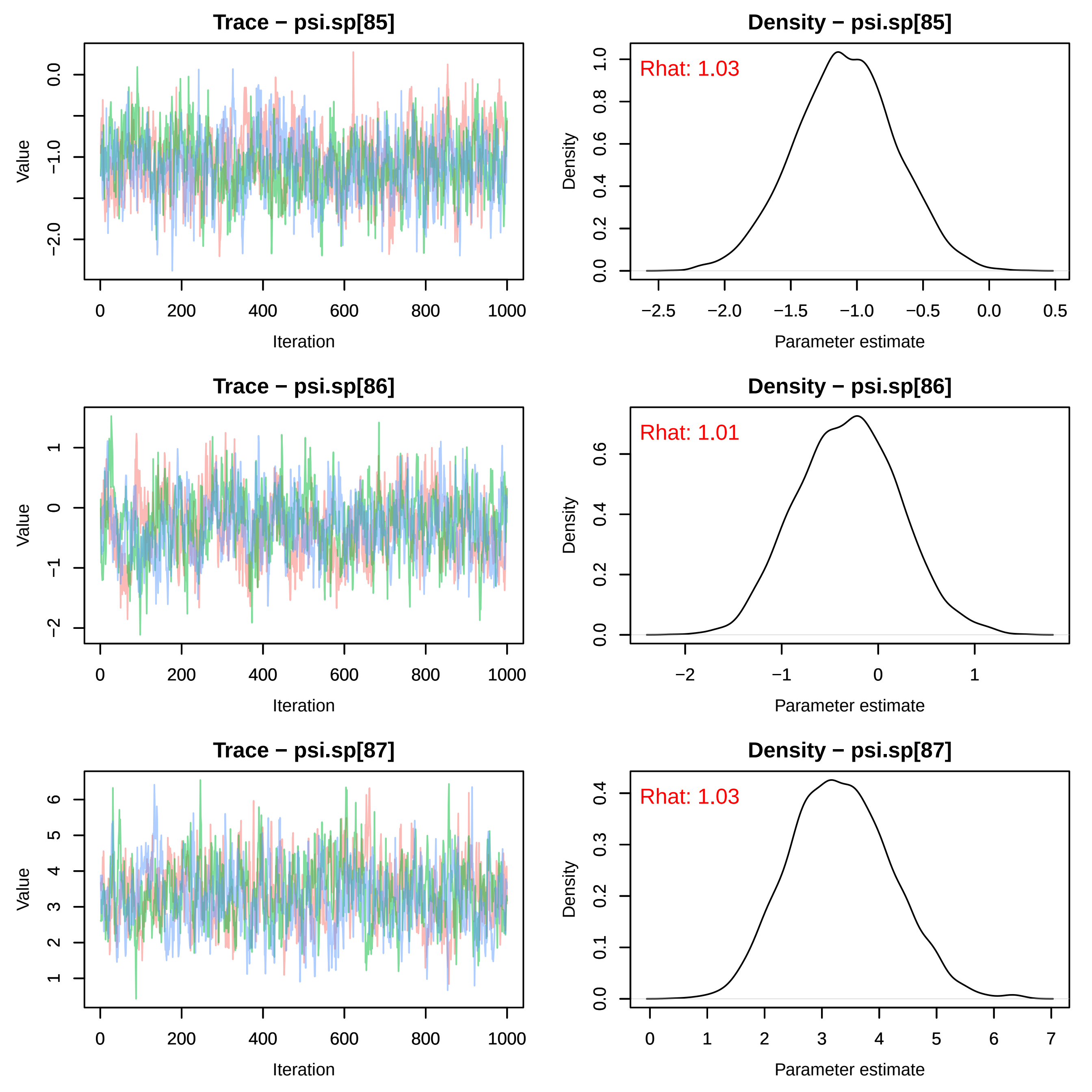

Trace − psi.sp[85]
Density − psi.sp[85]
Rhat: 1.03
Trace − psi.sp[86]
Density − psi.sp[86]
Rhat: 1.01
Trace − psi.sp[87]
Density − psi.sp[87]
Rhat: 1.03
Value
Iteration
Density
Parameter estimate

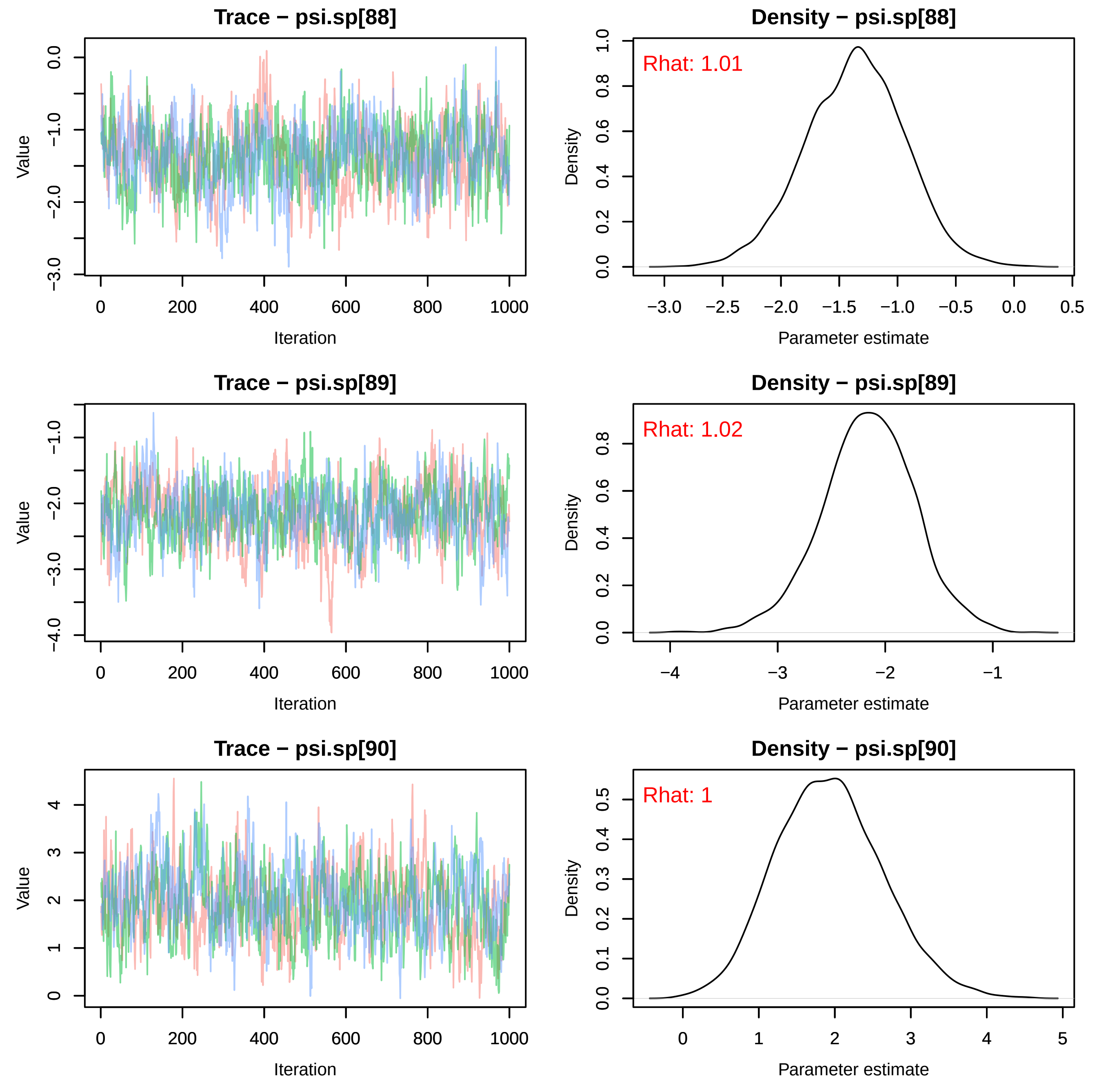
Trace − psi.sp[88]
Value
Iteration
Density − psi.sp[88]
Rhat: 1.01
Density
Parameter estimate
Trace − psi.sp[89]
Value
Iteration
Density − psi.sp[89]
Rhat: 1.02
Density
Parameter estimate
Trace − psi.sp[90]
Value
Iteration
Density − psi.sp[90]
Rhat: 1
Density
Parameter estimate

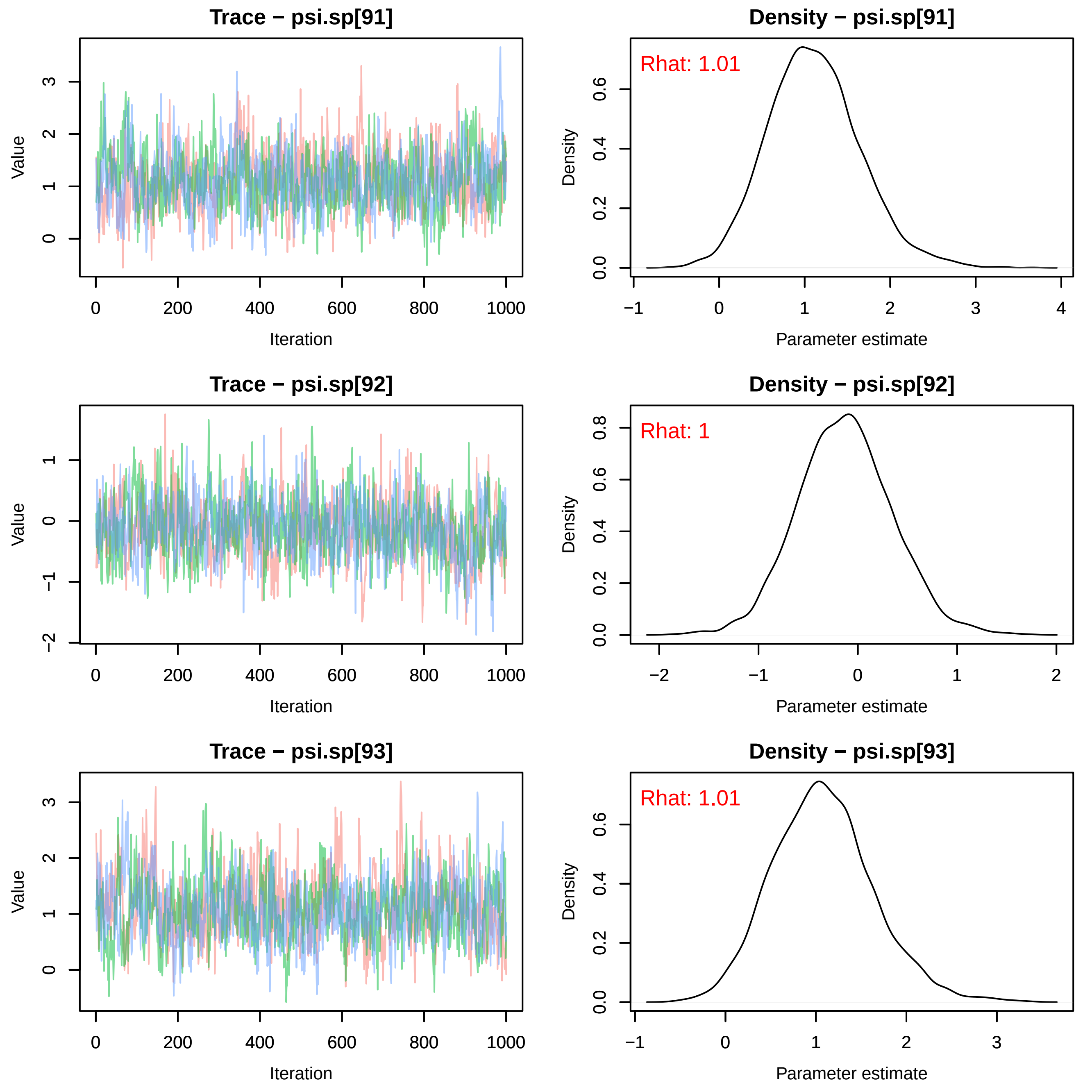
Trace − psi.sp[91]
Value
Iteration
0
200
400
600
800
1000
0
1
2
3
Density − psi.sp[91]
Rhat: 1.01
Density
Parameter estimate
0.0
0.2
0.4
0.6
−1
0
1
2
3
4
Trace − psi.sp[92]
Value
Iteration
0
200
400
600
800
1000
−2
−1
0
1
Density − psi.sp[92]
Rhat: 1
Density
Parameter estimate
0.0
0.2
0.4
0.6
0.8
−2
−1
0
1
2
Trace − psi.sp[93]
Value
Iteration
0
200
400
600
800
1000
0
1
2
3
Density − psi.sp[93]
Rhat: 1.01
Density
Parameter estimate
0.0
0.2
0.4
0.6
−1
0
1
2
3

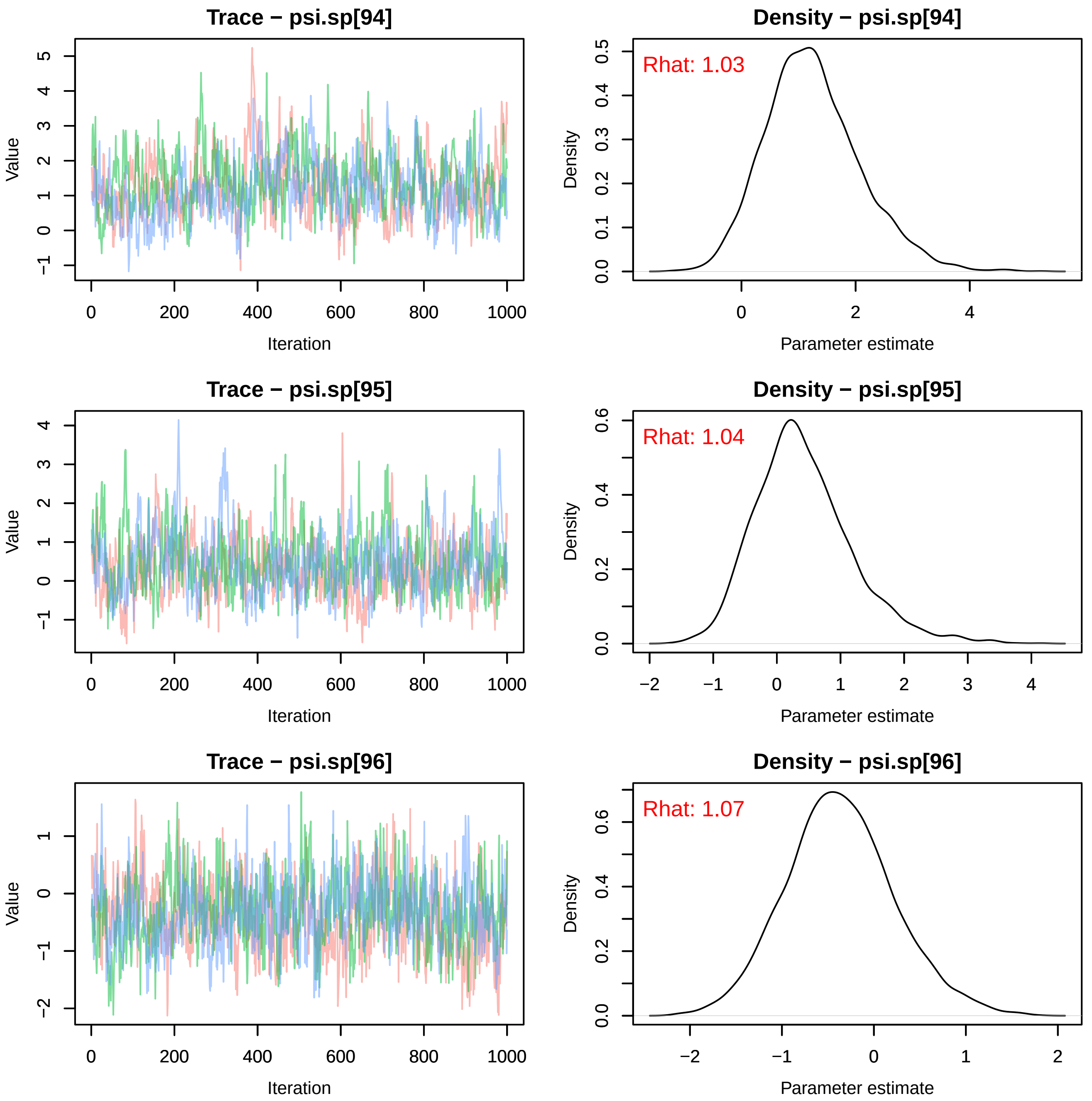
Trace − psi.sp[94]
Density − psi.sp[94]
Rhat: 1.03
Trace − psi.sp[95]
Density − psi.sp[95]
Rhat: 1.04
Trace − psi.sp[96]
Density − psi.sp[96]
Rhat: 1.07
Value
Iteration
Density
Parameter estimate

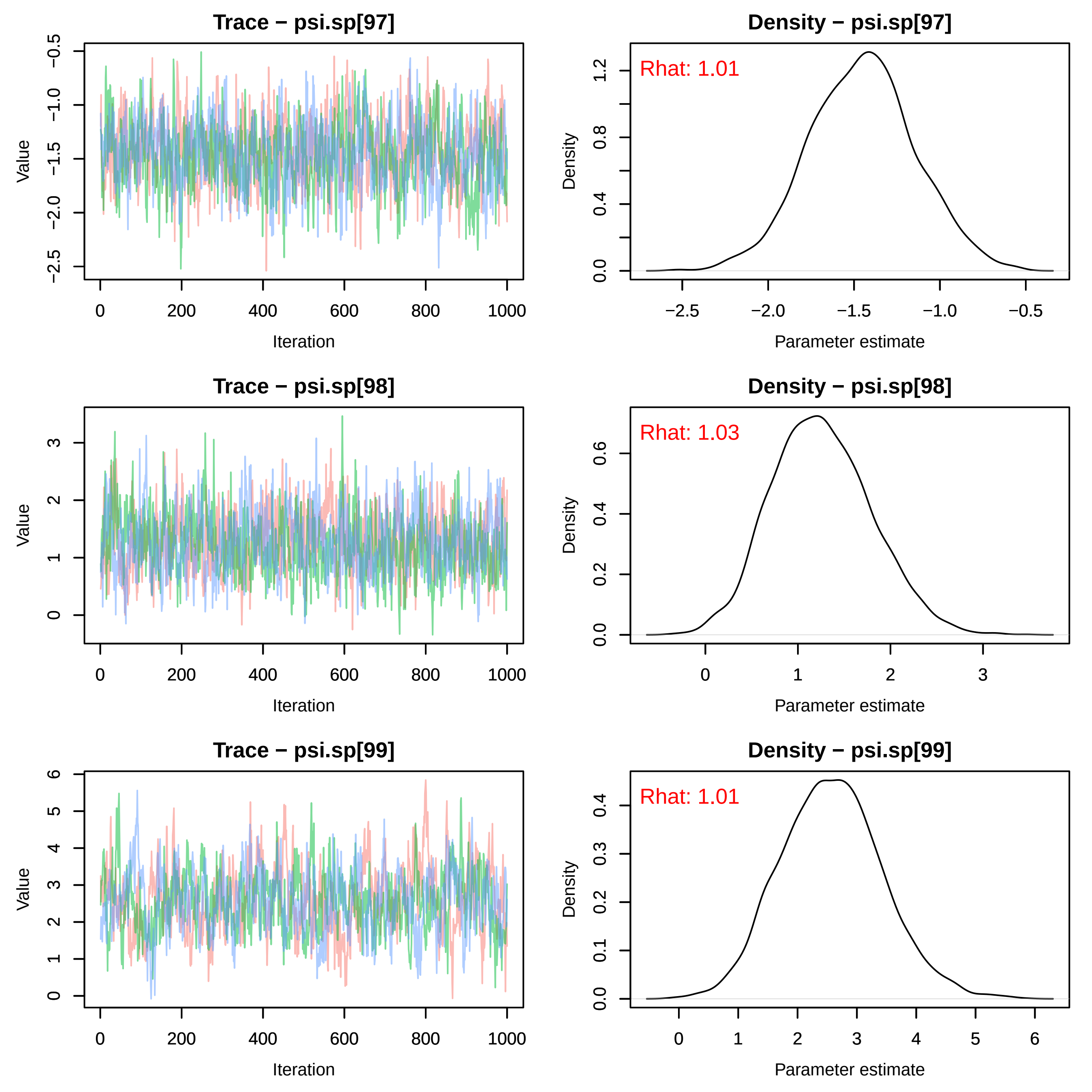

Trace − psi.sp[97]
Value
Iteration
Density − psi.sp[97]
Rhat: 1.01
Density
Parameter estimate
Trace − psi.sp[98]
Value
Iteration
Density − psi.sp[98]
Rhat: 1.03
Density
Parameter estimate
Trace − psi.sp[99]
Value
Iteration
Density − psi.sp[99]
Rhat: 1.01
Density
Parameter estimate

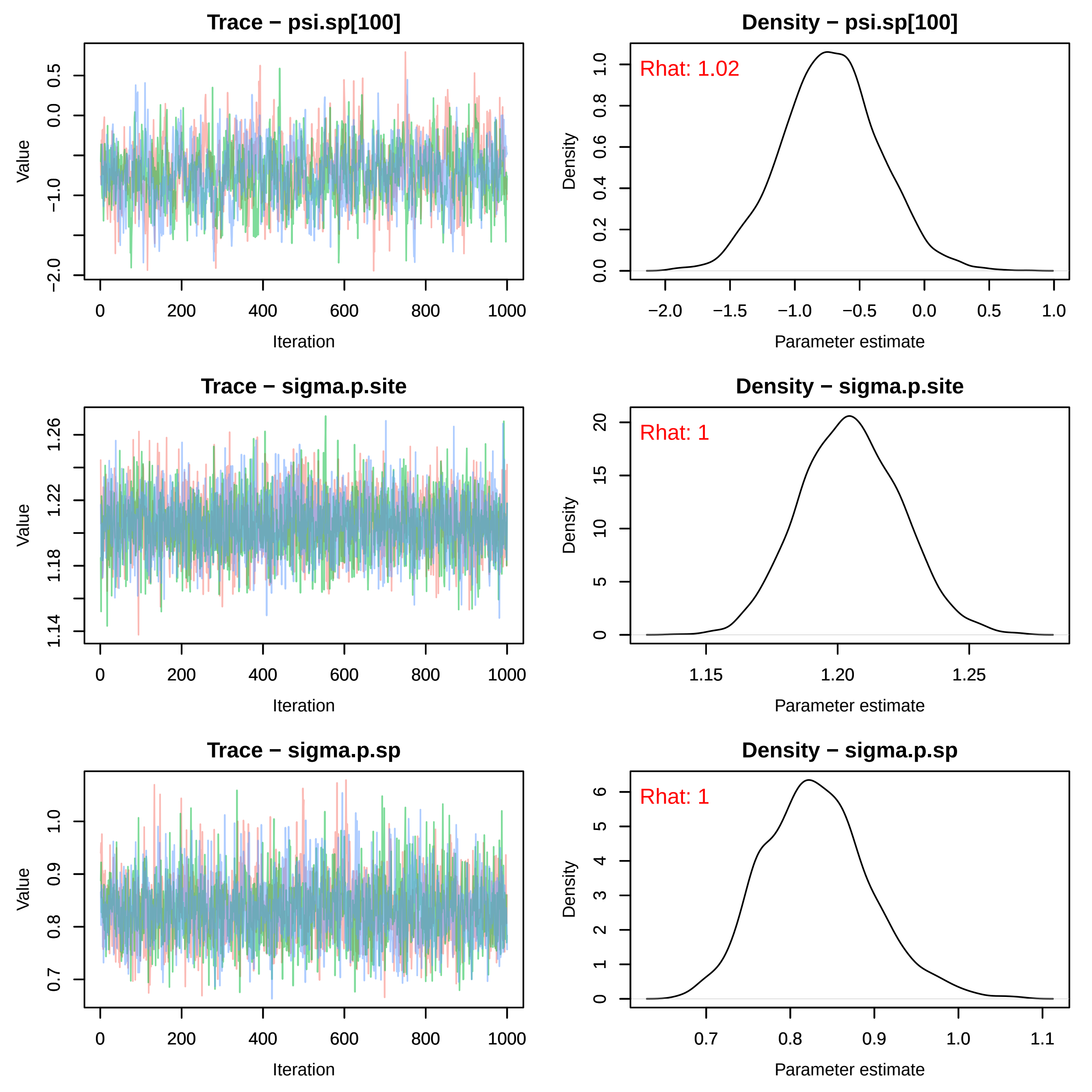
Trace − psi.sp[100]
Density − psi.sp[100]
Rhat: 1.02
Trace − sigma.p.site
Density − sigma.p.site
Rhat: 1
Trace − sigma.p.sp
Density − sigma.p.sp
Rhat: 1
Value
Iteration
Density
Parameter estimate

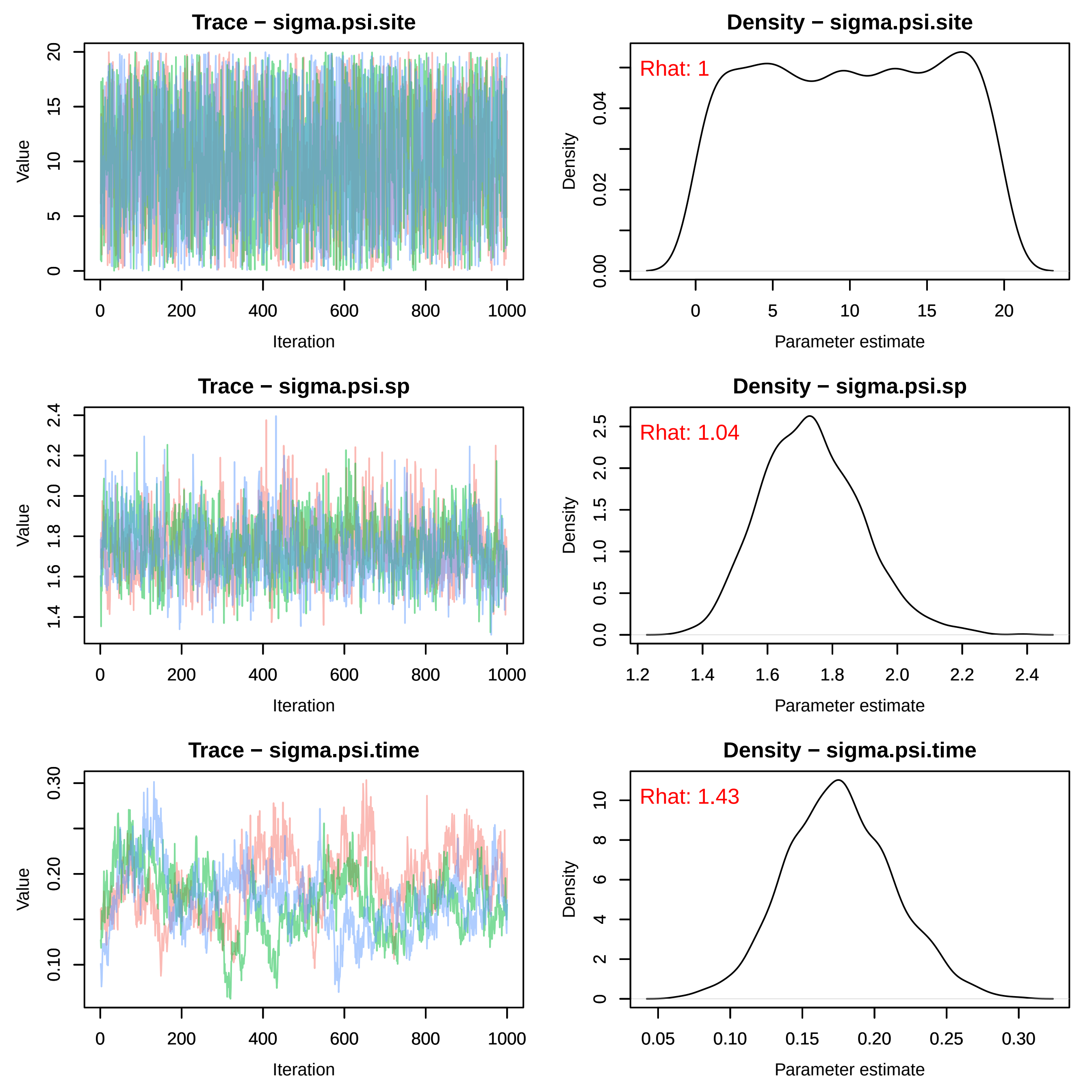

Trace − sigma.psi.site
Density − sigma.psi.site
Rhat: 1
Trace − sigma.psi.sp
Density − sigma.psi.sp
Rhat: 1.04
Trace − sigma.psi.time
Density − sigma.psi.time
Rhat: 1.43
Value
Density
Iteration
Parameter estimate

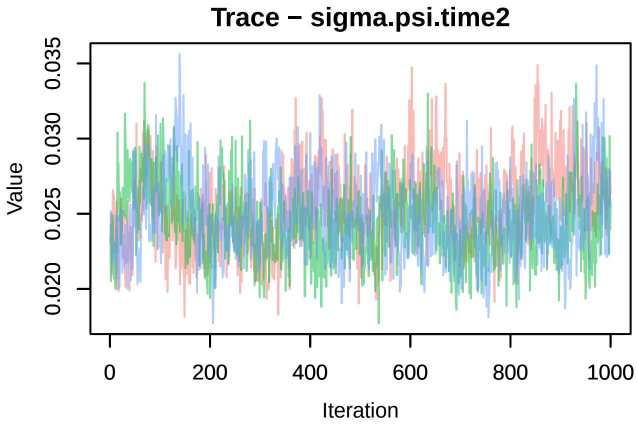
Trace − sigma.psi.time2
Value
0.020
0.025
0.030
0.035
0
200
400
600
800
1000
Iteration

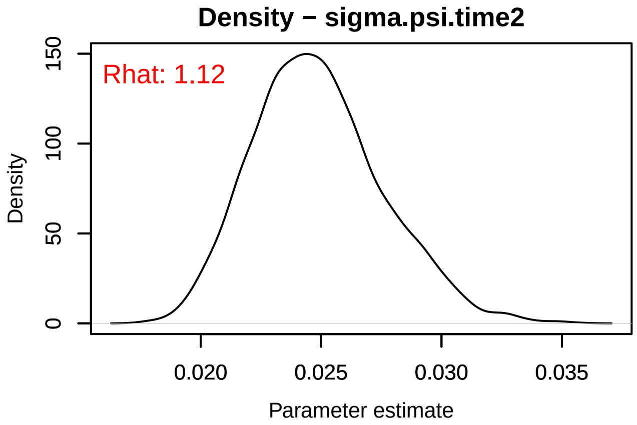
Density − sigma.psi.time2
Rhat: 1.12
Density
0
50
100
150
0.020
0.025
0.030
0.035
Parameter estimate
