## Supplementary material for "Winners and Losers Among European Arthropods over the Last Half Century of Global Change": hoverfly_envTrace.pdf

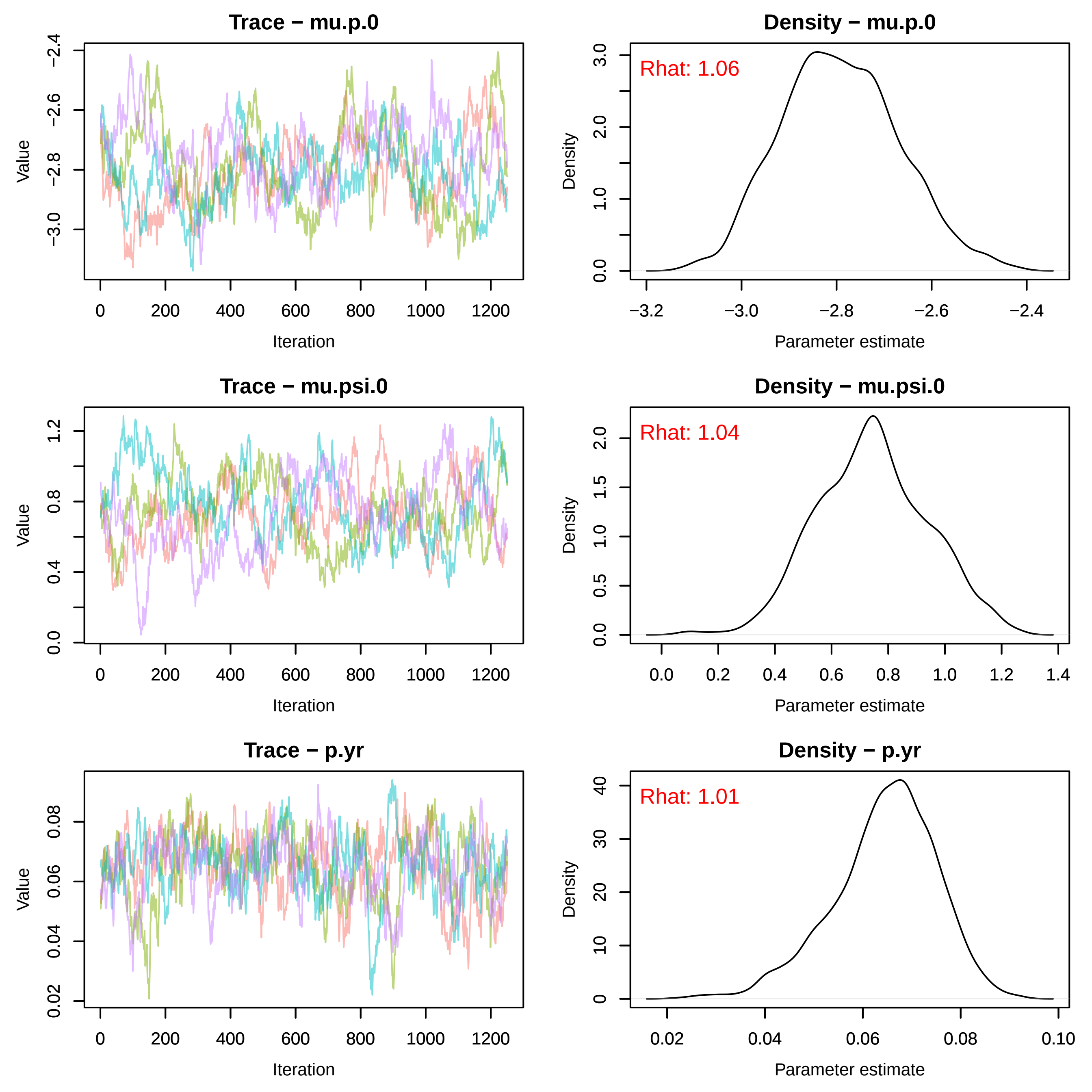

Trace − mu.p.0
Density − mu.p.0
Rhat: 1.06
Trace − mu.psi.0
Density − mu.psi.0
Rhat: 1.04
Trace − p.yr
Density − p.yr
Rhat: 1.01
Value
Iteration
Density
Parameter estimate

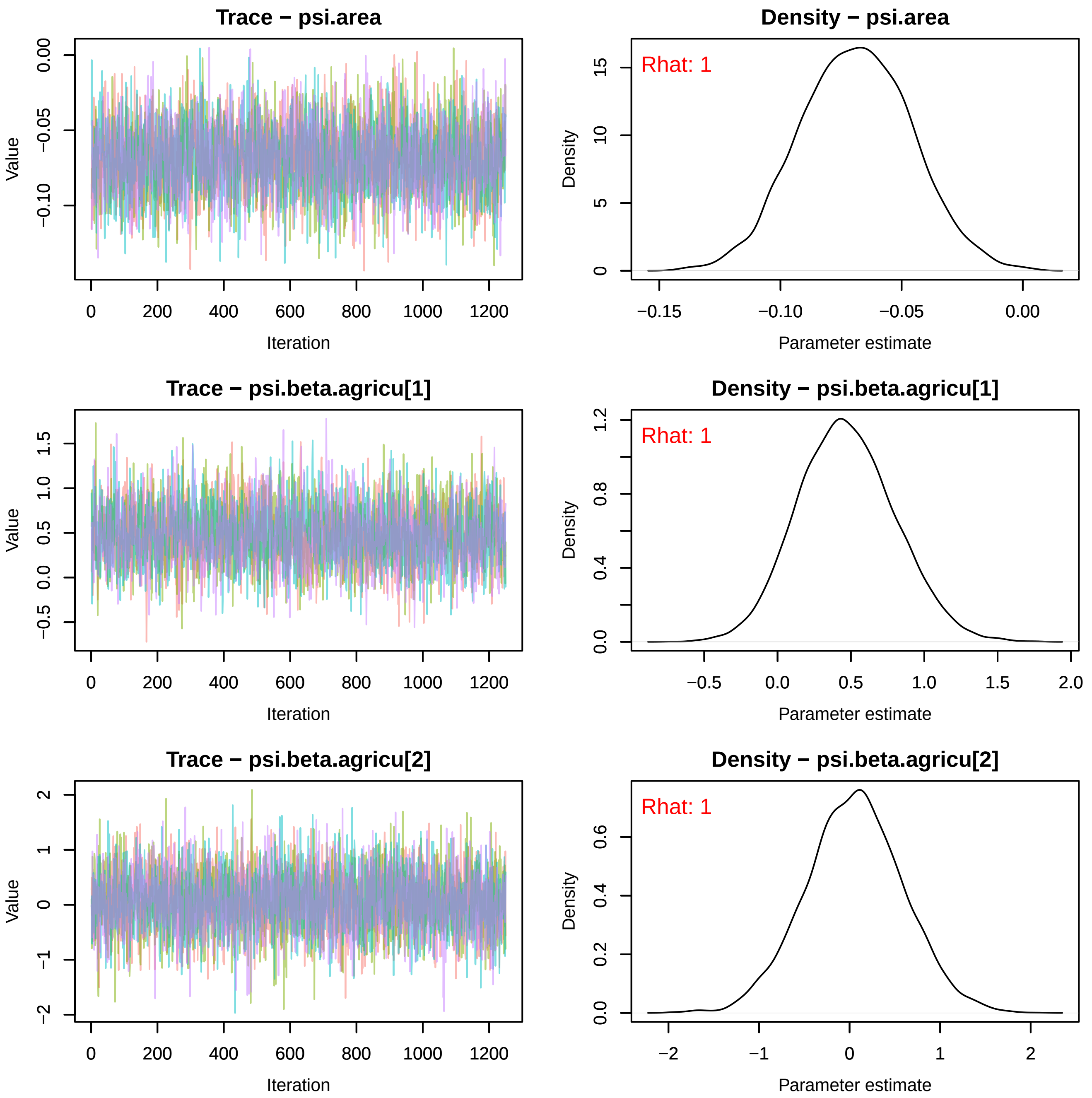
Trace − psi.area
Value
Iteration
0.00
−0.05
−0.10
0 200 400 600 800 1000 1200
Density − psi.area
Rhat: 1
Density
Parameter estimate
0 5 10 15
−0.15 −0.10 −0.05 0.00
Trace − psi.beta.agricu[1]
Value
Iteration
−0.5 0.0 0.5 1.0 1.5
0 200 400 600 800 1000 1200
Density − psi.beta.agricu[1]
Rhat: 1
Density
Parameter estimate
0.0 0.4 0.8 1.2
−0.5 0.0 0.5 1.0 1.5 2.0
Trace − psi.beta.agricu[2]
Value
Iteration
−2 −1 0 1 2
0 200 400 600 800 1000 1200
Density − psi.beta.agricu[2]
Rhat: 1
Density
Parameter estimate
0.0 0.2 0.4 0.6
−2 −1 0 1 2

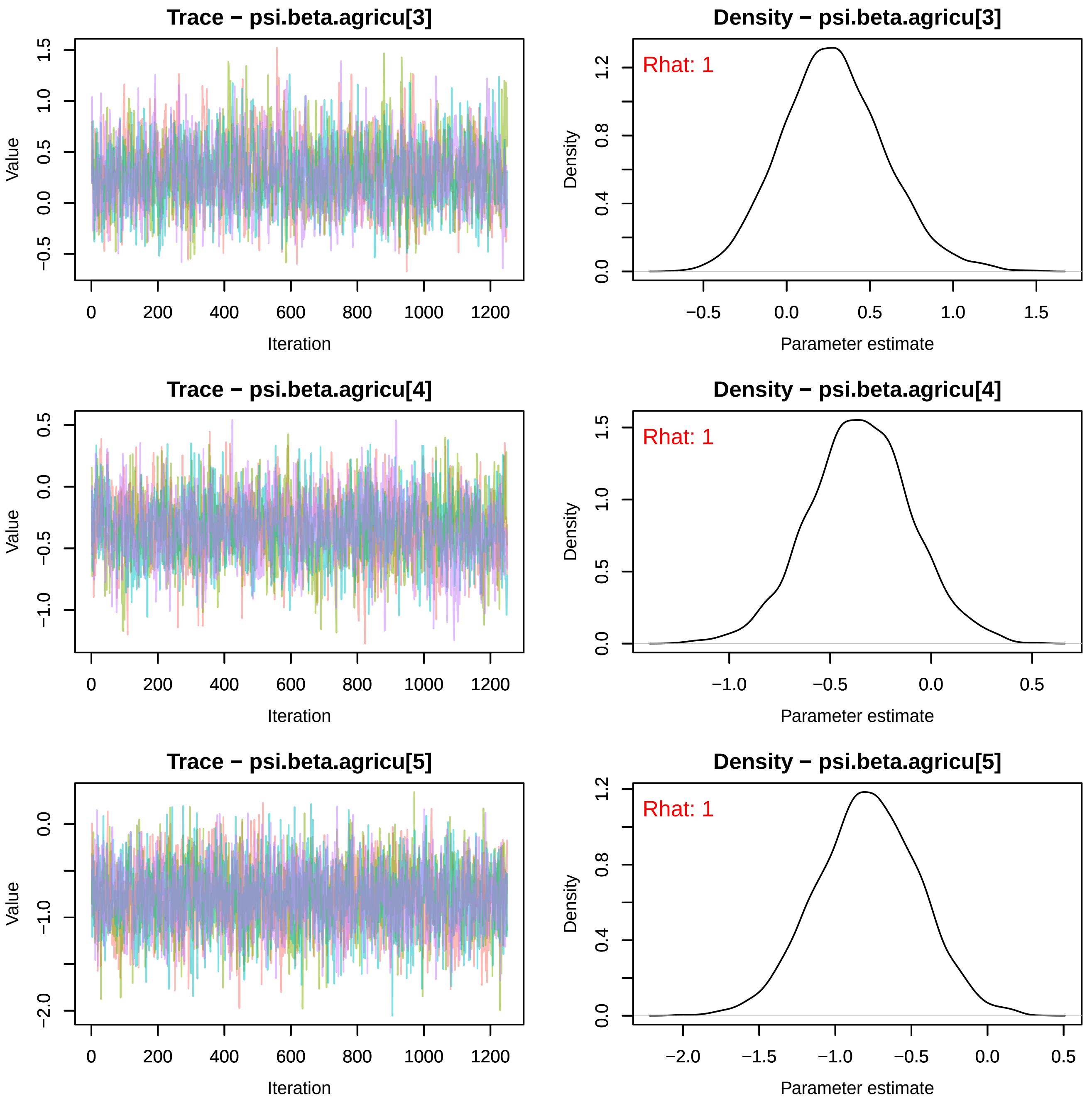
Trace − psi.beta.agricu[3]
Density − psi.beta.agricu[3]
Rhat: 1
Trace − psi.beta.agricu[4]
Density − psi.beta.agricu[4]
Rhat: 1
Trace − psi.beta.agricu[5]
Density − psi.beta.agricu[5]
Rhat: 1
Value
Iteration
Density
Parameter estimate
0
200
400
600
800
1000
1200

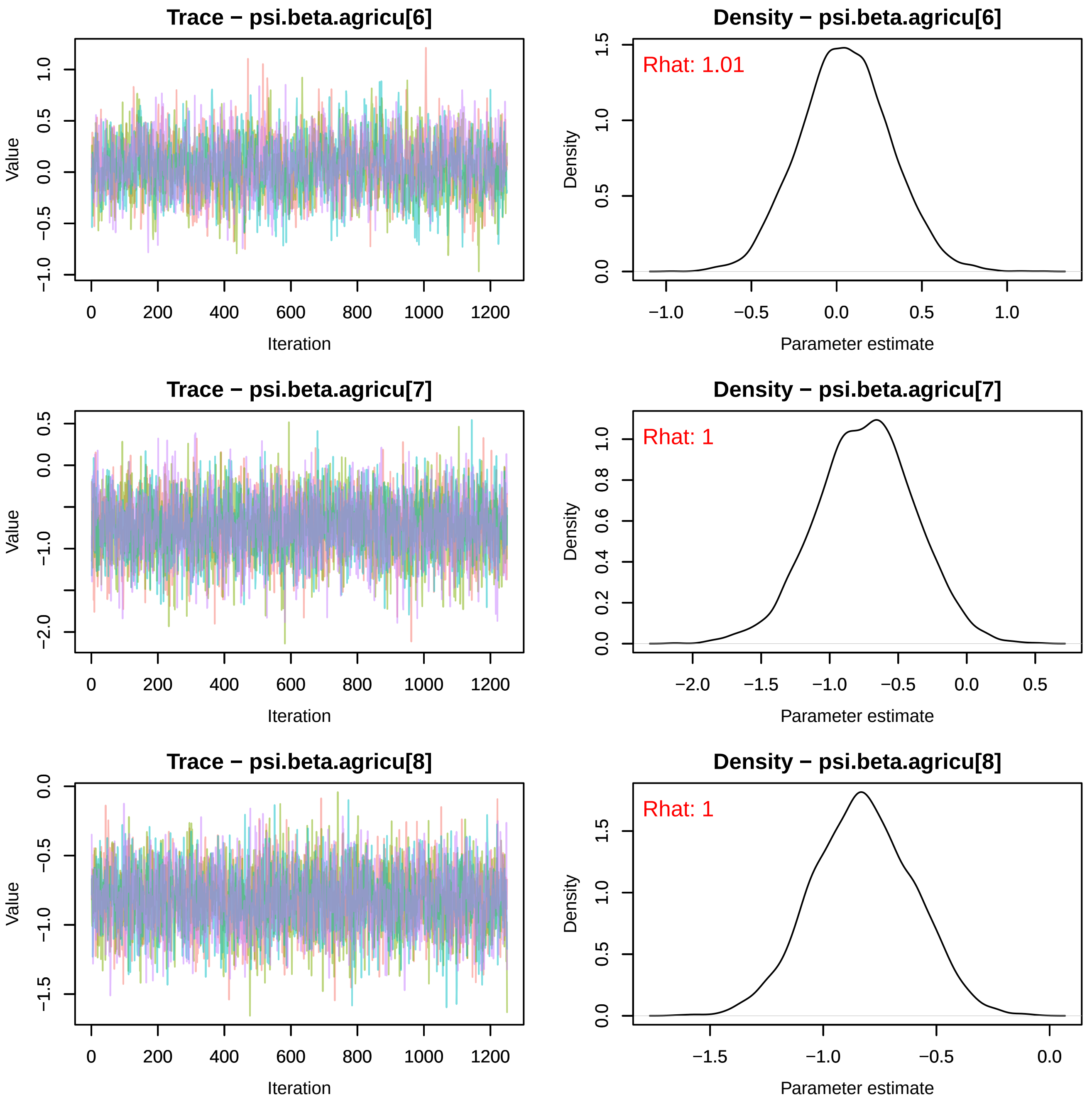
Trace − psi.beta.agricu[6]
Density − psi.beta.agricu[6]
Rhat: 1.01
Trace − psi.beta.agricu[7]
Density − psi.beta.agricu[7]
Rhat: 1
Trace − psi.beta.agricu[8]
Density − psi.beta.agricu[8]
Rhat: 1
Value
Iteration
Density
Parameter estimate

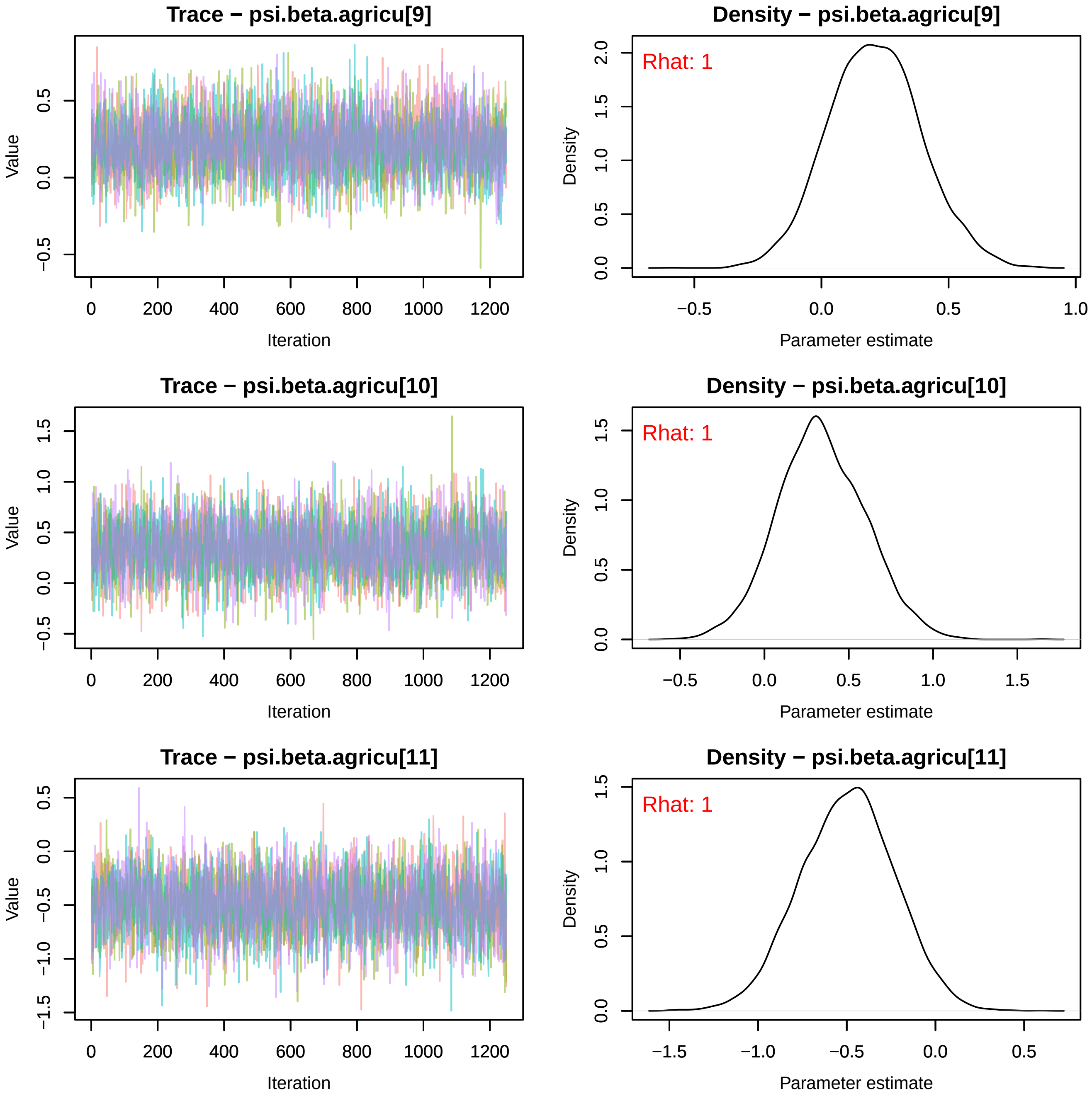
Trace − psi.beta.agricu[9]
Value
Iteration
Density − psi.beta.agricu[9]
Rhat: 1
Density
Parameter estimate
Trace − psi.beta.agricu[10]
Value
Iteration
Density − psi.beta.agricu[10]
Rhat: 1
Density
Parameter estimate
Trace − psi.beta.agricu[11]
Value
Iteration
Density − psi.beta.agricu[11]
Rhat: 1
Density
Parameter estimate

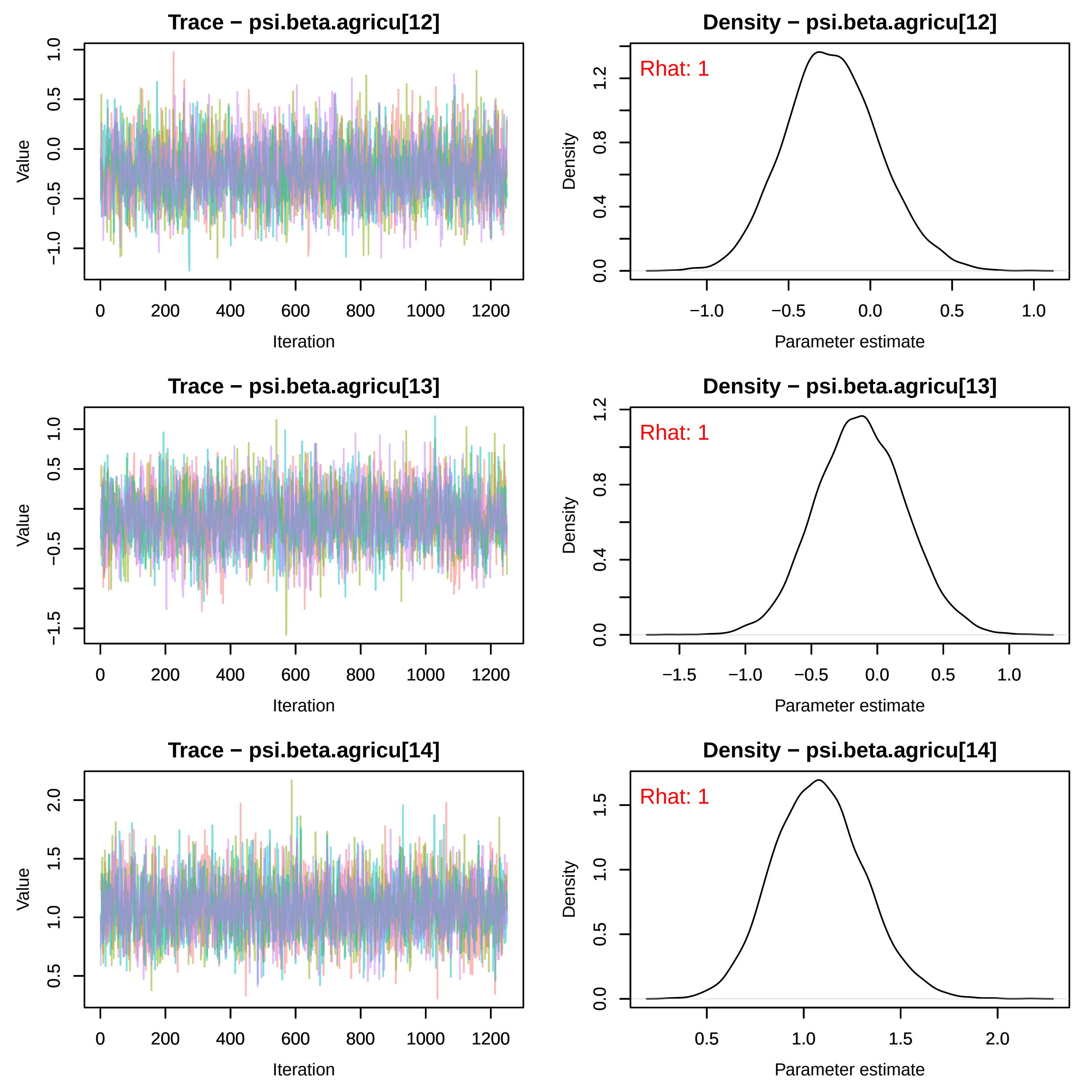
Trace − psi.beta.agricu[12]
Value
Iteration
Density − psi.beta.agricu[12]
Rhat: 1
Density
Parameter estimate
Trace − psi.beta.agricu[13]
Value
Iteration
Density − psi.beta.agricu[13]
Rhat: 1
Density
Parameter estimate
Trace − psi.beta.agricu[14]
Value
Iteration
Density − psi.beta.agricu[14]
Rhat: 1
Density
Parameter estimate

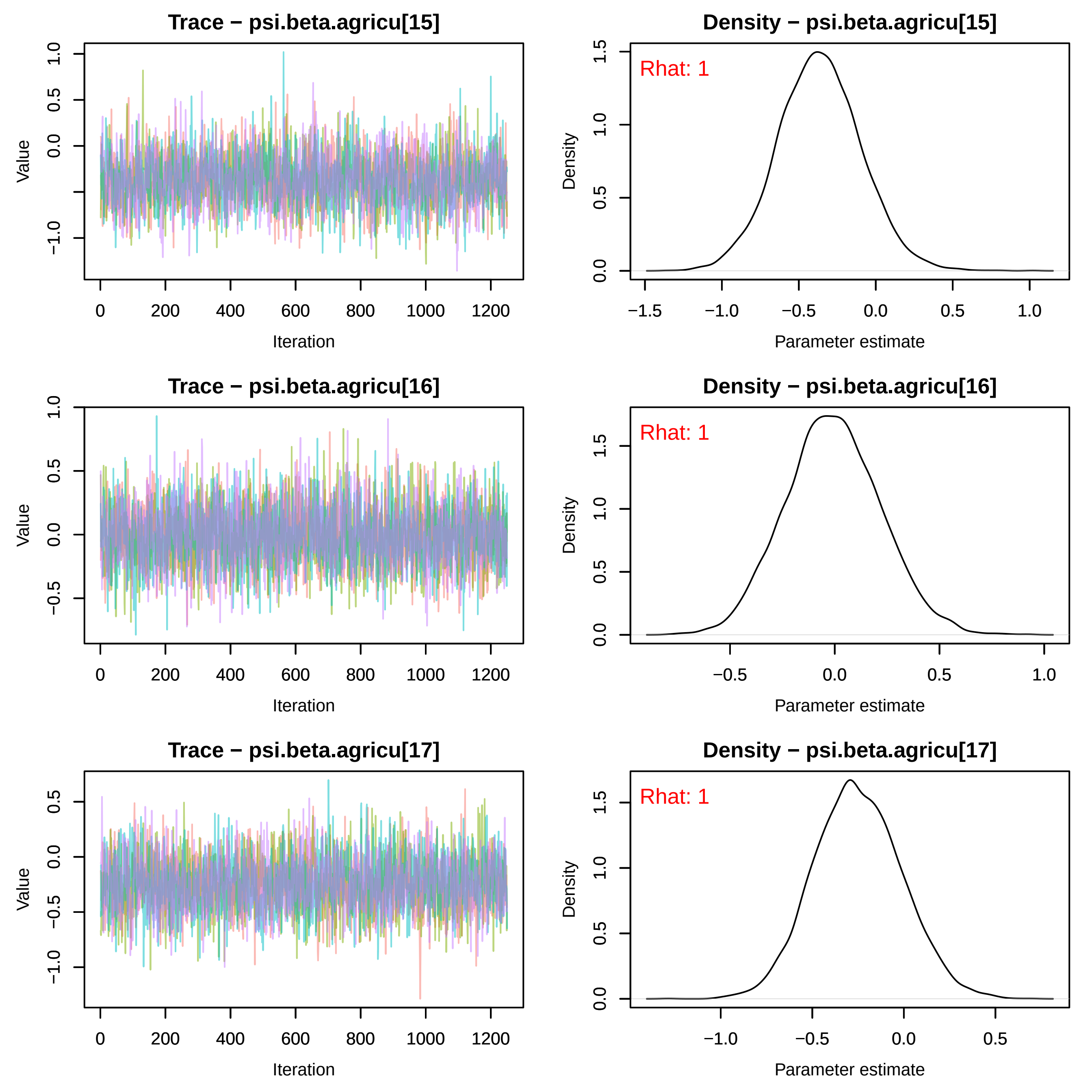
Trace − psi.beta.agricu[15]
Value
Iteration
Density − psi.beta.agricu[15]
Rhat: 1
Density
Parameter estimate
Trace − psi.beta.agricu[16]
Value
Iteration
Density − psi.beta.agricu[16]
Rhat: 1
Density
Parameter estimate
Trace − psi.beta.agricu[17]
Value
Iteration
Density − psi.beta.agricu[17]
Rhat: 1
Density
Parameter estimate

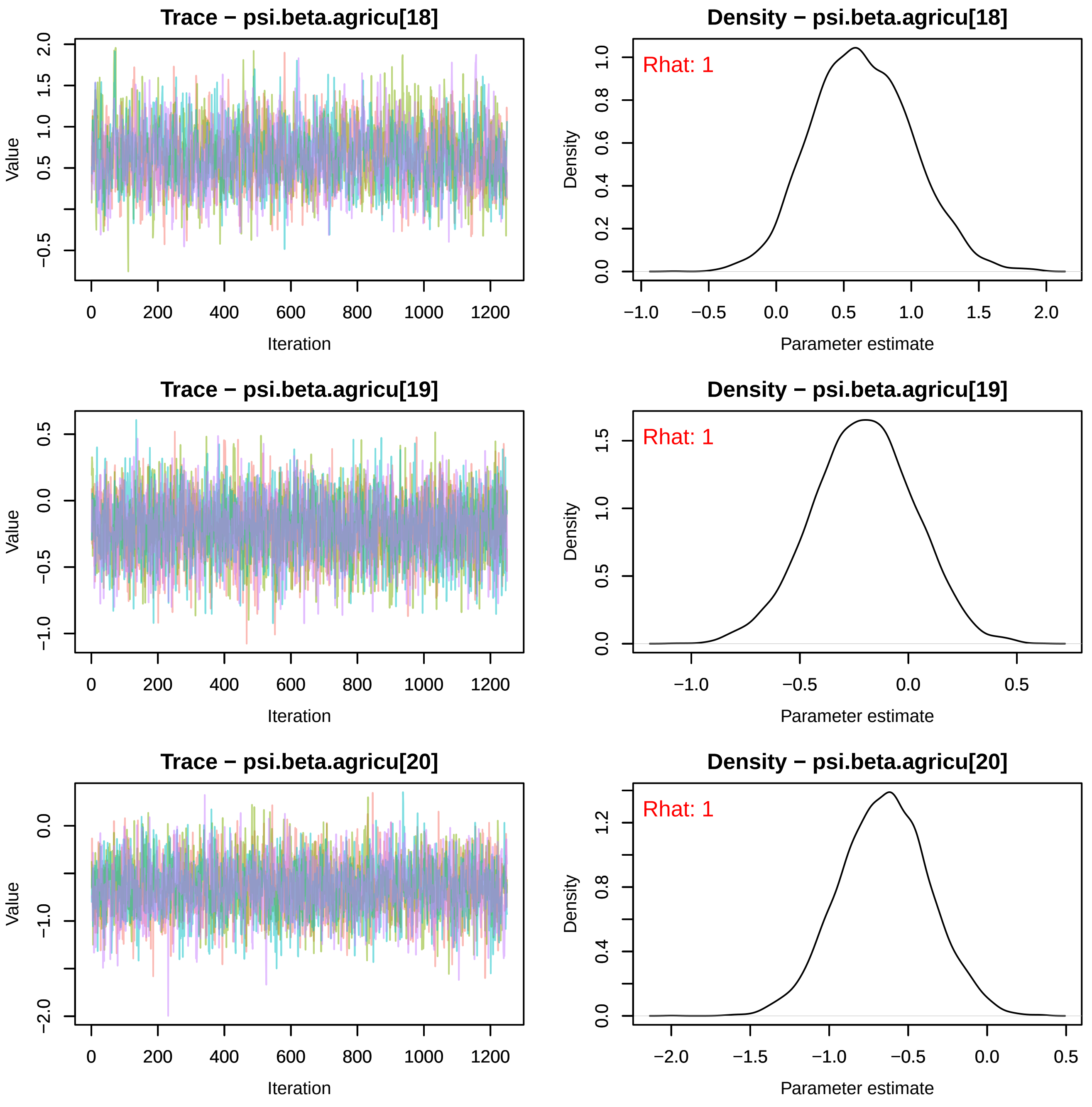
Trace − psi.beta.agricu[18]
Value
Iteration
Density − psi.beta.agricu[18]
Rhat: 1
Density
Parameter estimate
Trace − psi.beta.agricu[19]
Value
Iteration
Density − psi.beta.agricu[19]
Rhat: 1
Density
Parameter estimate
Trace − psi.beta.agricu[20]
Value
Iteration
Density − psi.beta.agricu[20]
Rhat: 1
Density
Parameter estimate

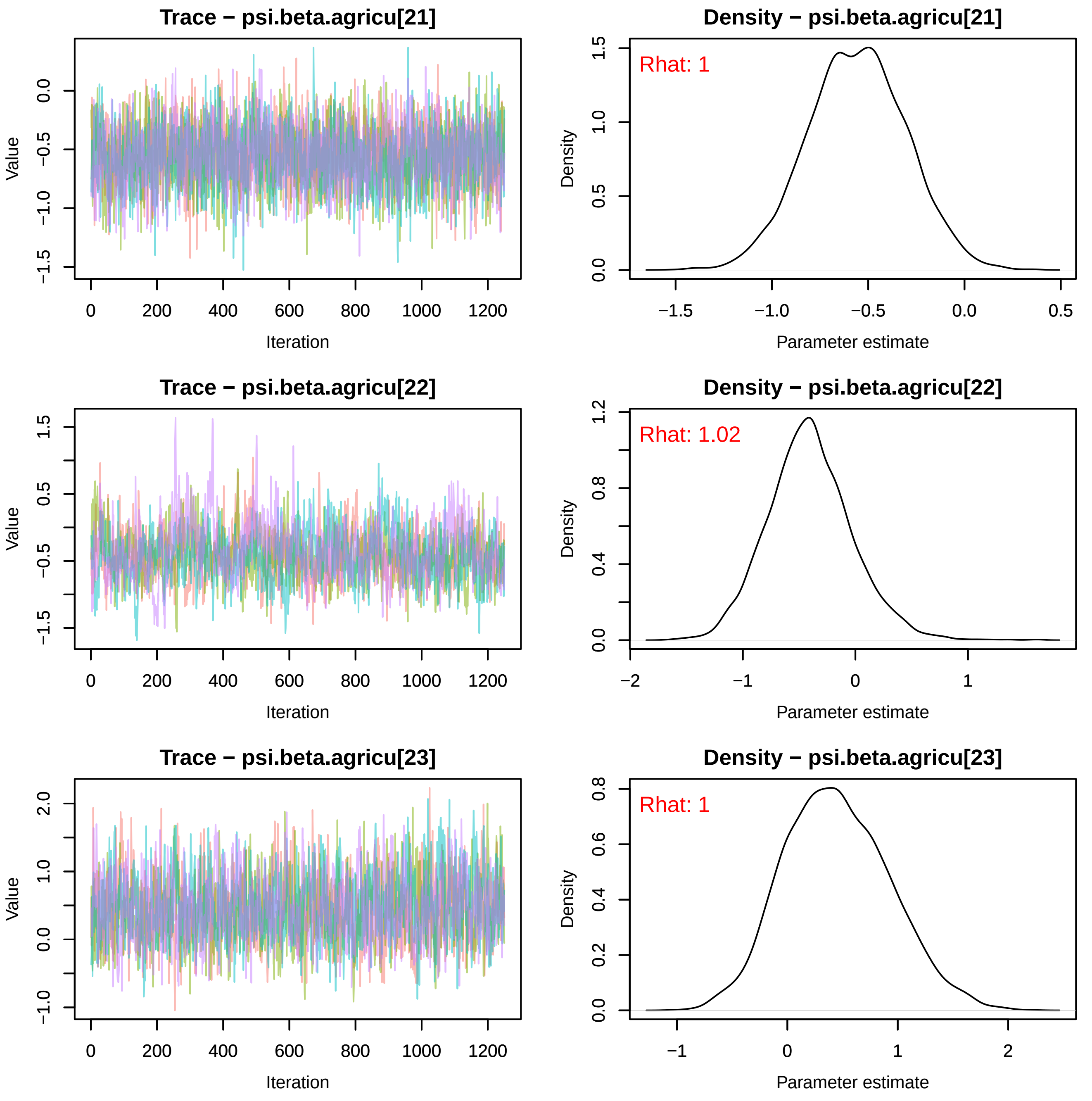
Trace − psi.beta.agricu[21]
Value
Iteration
Density − psi.beta.agricu[21]
Rhat: 1
Density
Parameter estimate
Trace − psi.beta.agricu[22]
Value
Iteration
Density − psi.beta.agricu[22]
Rhat: 1.02
Density
Parameter estimate
Trace − psi.beta.agricu[23]
Value
Iteration
Density − psi.beta.agricu[23]
Rhat: 1
Density
Parameter estimate

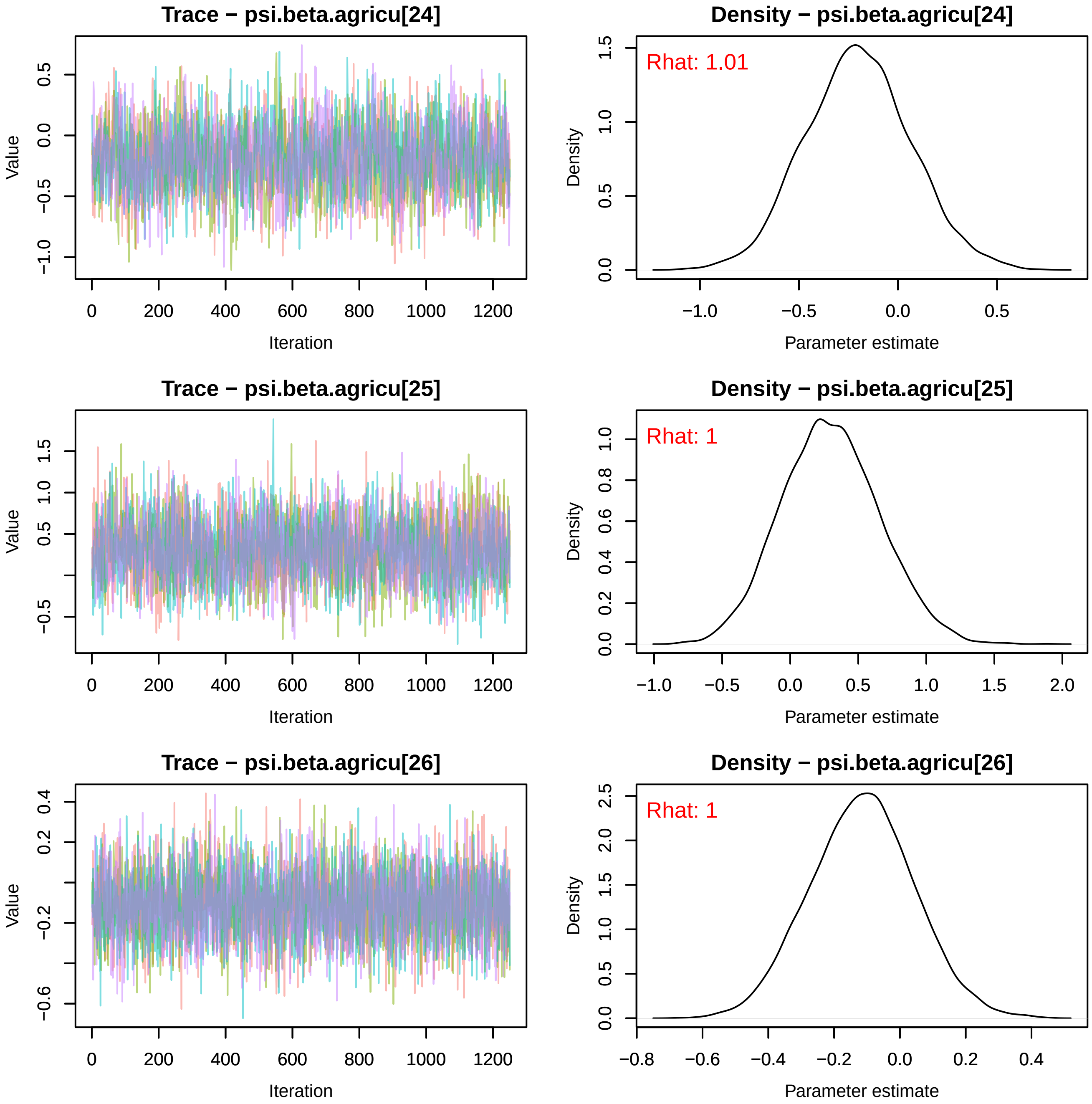
Trace − psi.beta.agricu[24]
Value
Iteration
Density − psi.beta.agricu[24]
Rhat: 1.01
Density
Parameter estimate
Trace − psi.beta.agricu[25]
Value
Iteration
Density − psi.beta.agricu[25]
Rhat: 1
Density
Parameter estimate
Trace − psi.beta.agricu[26]
Value
Iteration
Density − psi.beta.agricu[26]
Rhat: 1
Density
Parameter estimate

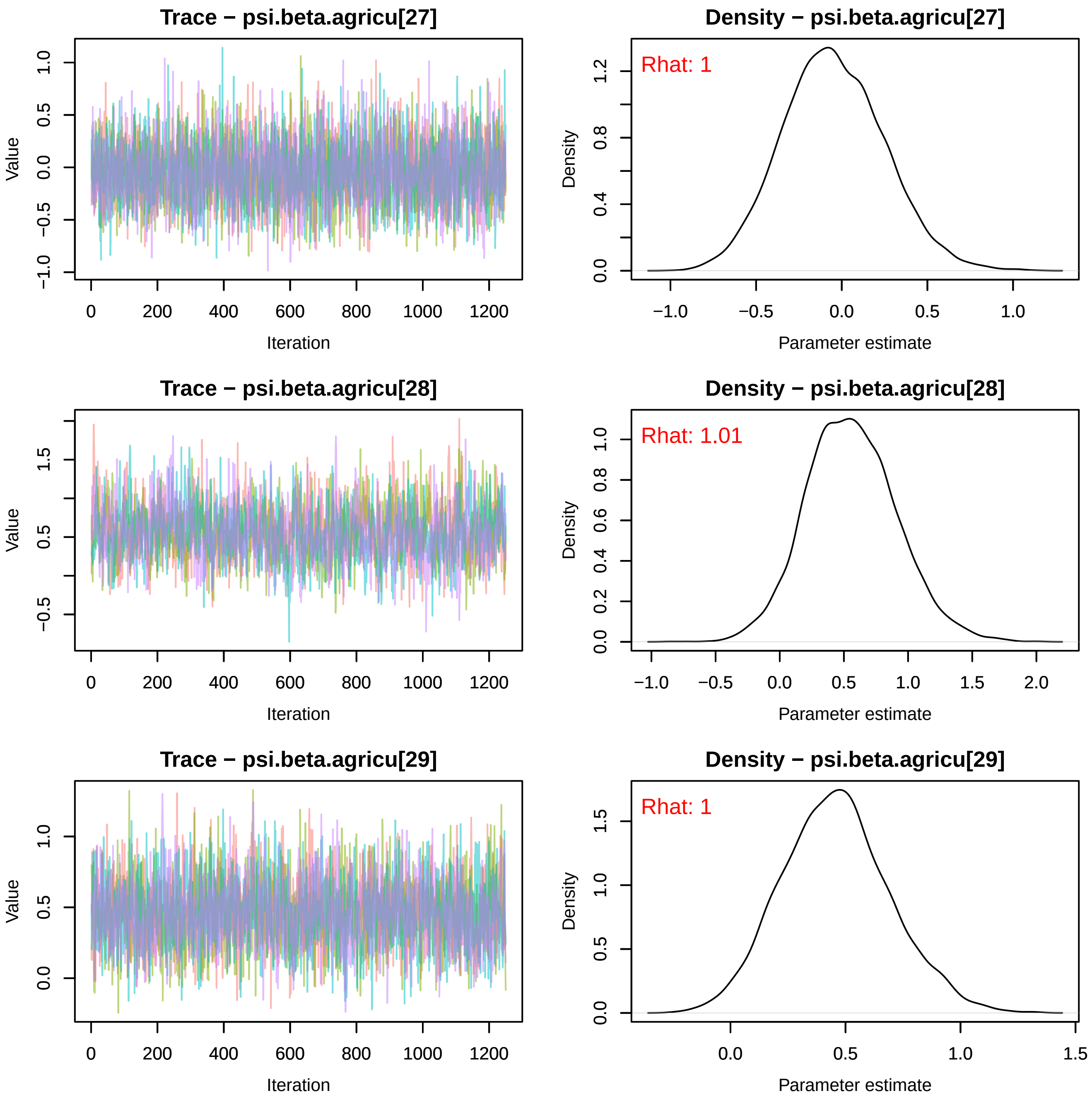
Trace − psi.beta.agricu[27]
Value
Iteration
Density − psi.beta.agricu[27]
Rhat: 1
Density
Parameter estimate
Trace − psi.beta.agricu[28]
Value
Iteration
Density − psi.beta.agricu[28]
Rhat: 1.01
Density
Parameter estimate
Trace − psi.beta.agricu[29]
Value
Iteration
Density − psi.beta.agricu[29]
Rhat: 1
Density
Parameter estimate

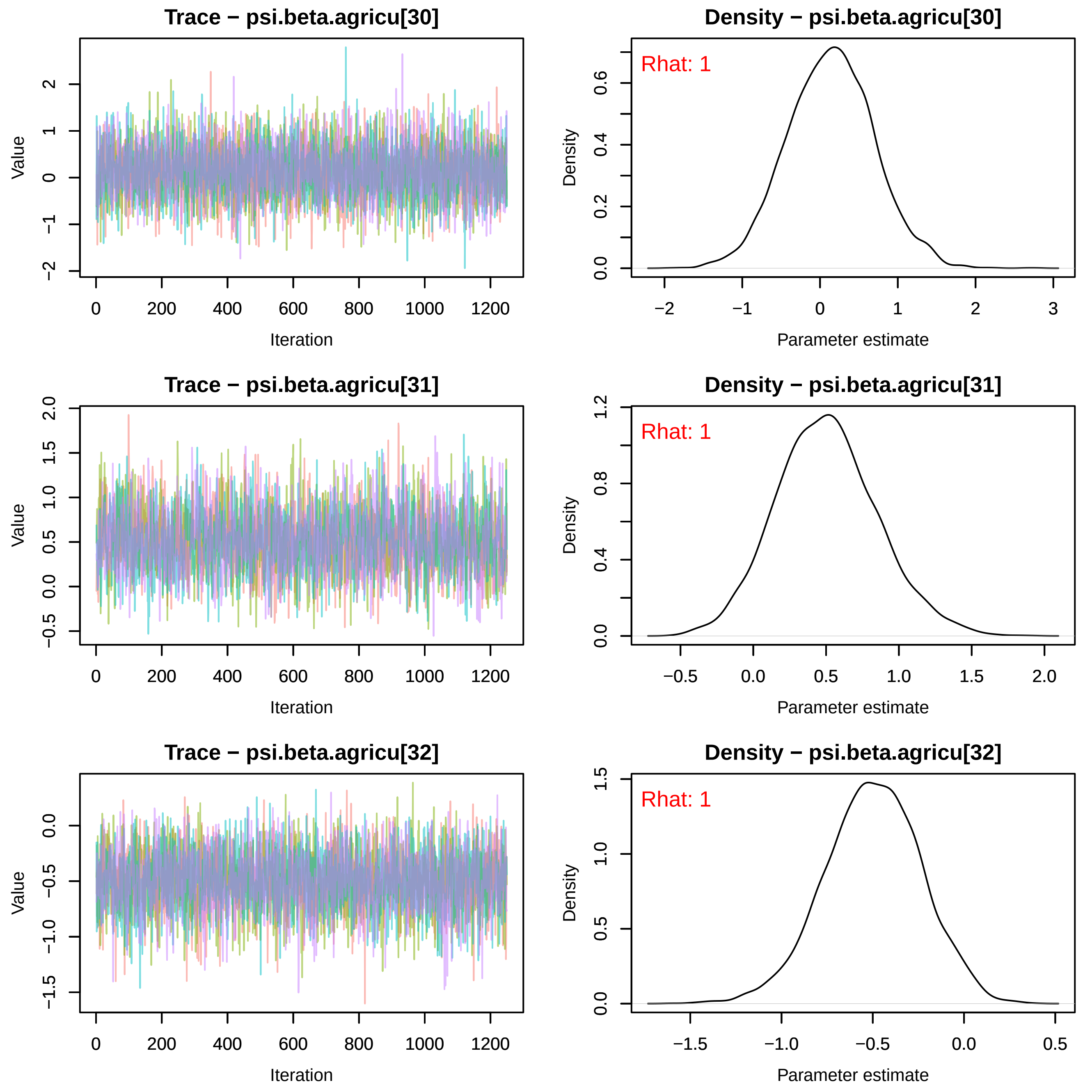
Trace − psi.beta.agricu[30]
Value
Iteration
0 200 400 600 800 1000 1200
−2 −1 0 1 2
Density − psi.beta.agricu[30]
Rhat: 1
Density
Parameter estimate
−2 −1 0 1 2 3
0.0 0.2 0.4 0.6
Trace − psi.beta.agricu[31]
Value
Iteration
0 200 400 600 800 1000 1200
−0.5 0.0 0.5 1.0 1.5 2.0
Density − psi.beta.agricu[31]
Rhat: 1
Density
Parameter estimate
−0.5 0.0 0.5 1.0 1.5 2.0
0.0 0.4 0.8 1.2
Trace − psi.beta.agricu[32]
Value
Iteration
0 200 400 600 800 1000 1200
−1.5 −1.0 −0.5 0.0
Density − psi.beta.agricu[32]
Rhat: 1
Density
Parameter estimate
−1.5 −1.0 −0.5 0.0 0.5
0.0 0.5 1.0 1.5

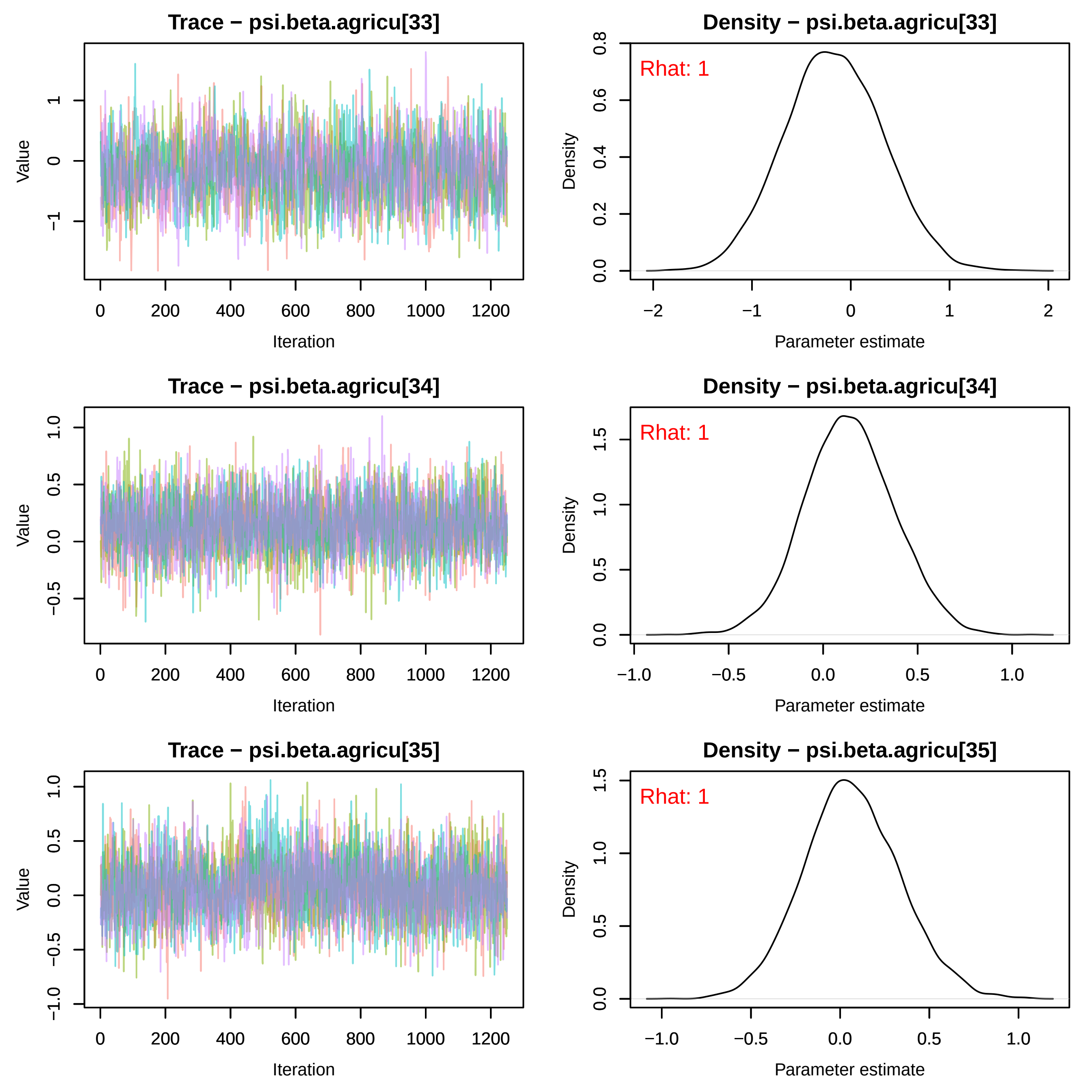
Trace − psi.beta.agricu[33]
Value
Iteration
Density − psi.beta.agricu[33]
Rhat: 1
Density
Parameter estimate
Trace − psi.beta.agricu[34]
Value
Iteration
Density − psi.beta.agricu[34]
Rhat: 1
Density
Parameter estimate
Trace − psi.beta.agricu[35]
Value
Iteration
Density − psi.beta.agricu[35]
Rhat: 1
Density
Parameter estimate

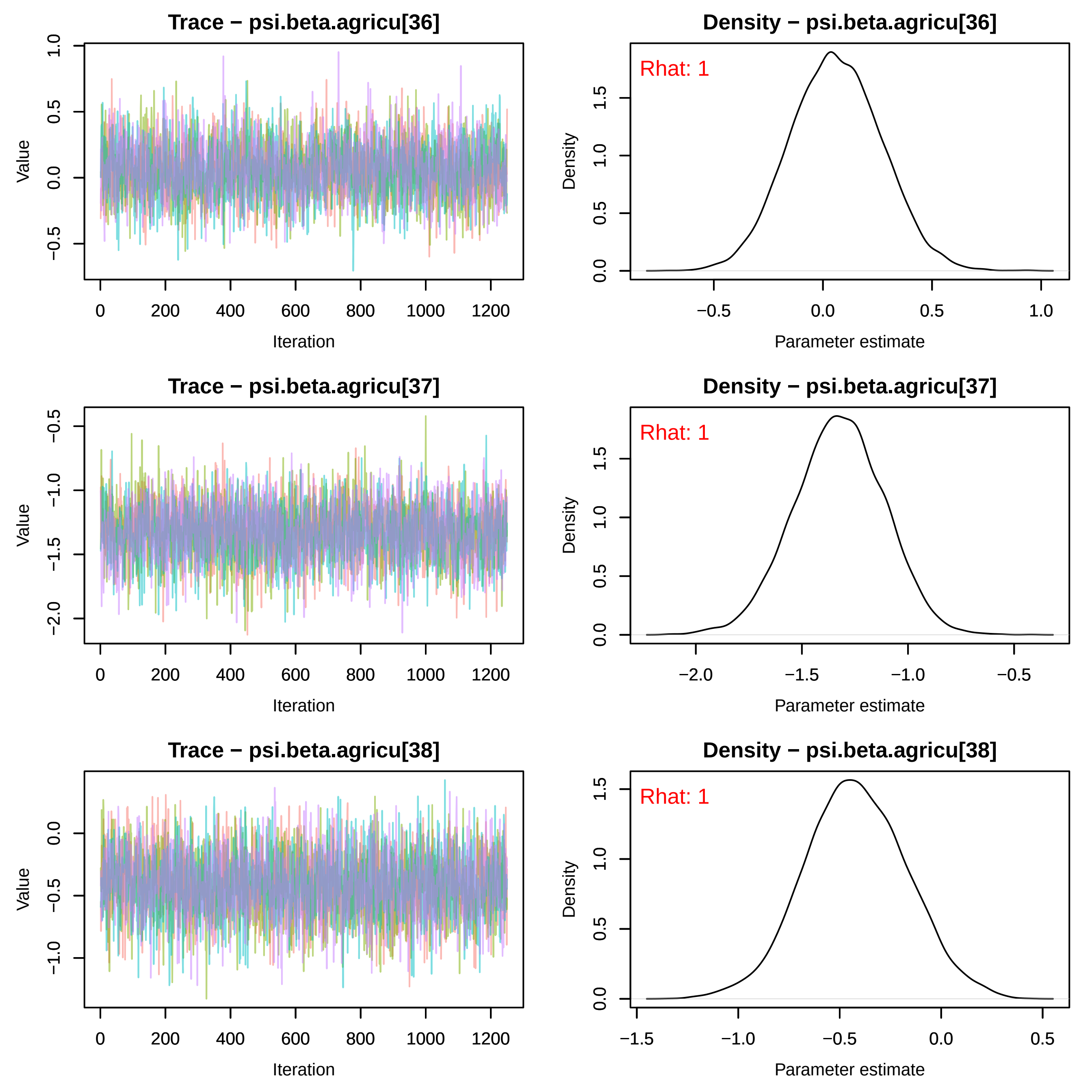
Trace − psi.beta.agricu[36]
Density − psi.beta.agricu[36]
Rhat: 1
Trace − psi.beta.agricu[37]
Density − psi.beta.agricu[37]
Rhat: 1
Trace − psi.beta.agricu[38]
Density − psi.beta.agricu[38]
Rhat: 1
Value
Iteration
Density
Parameter estimate

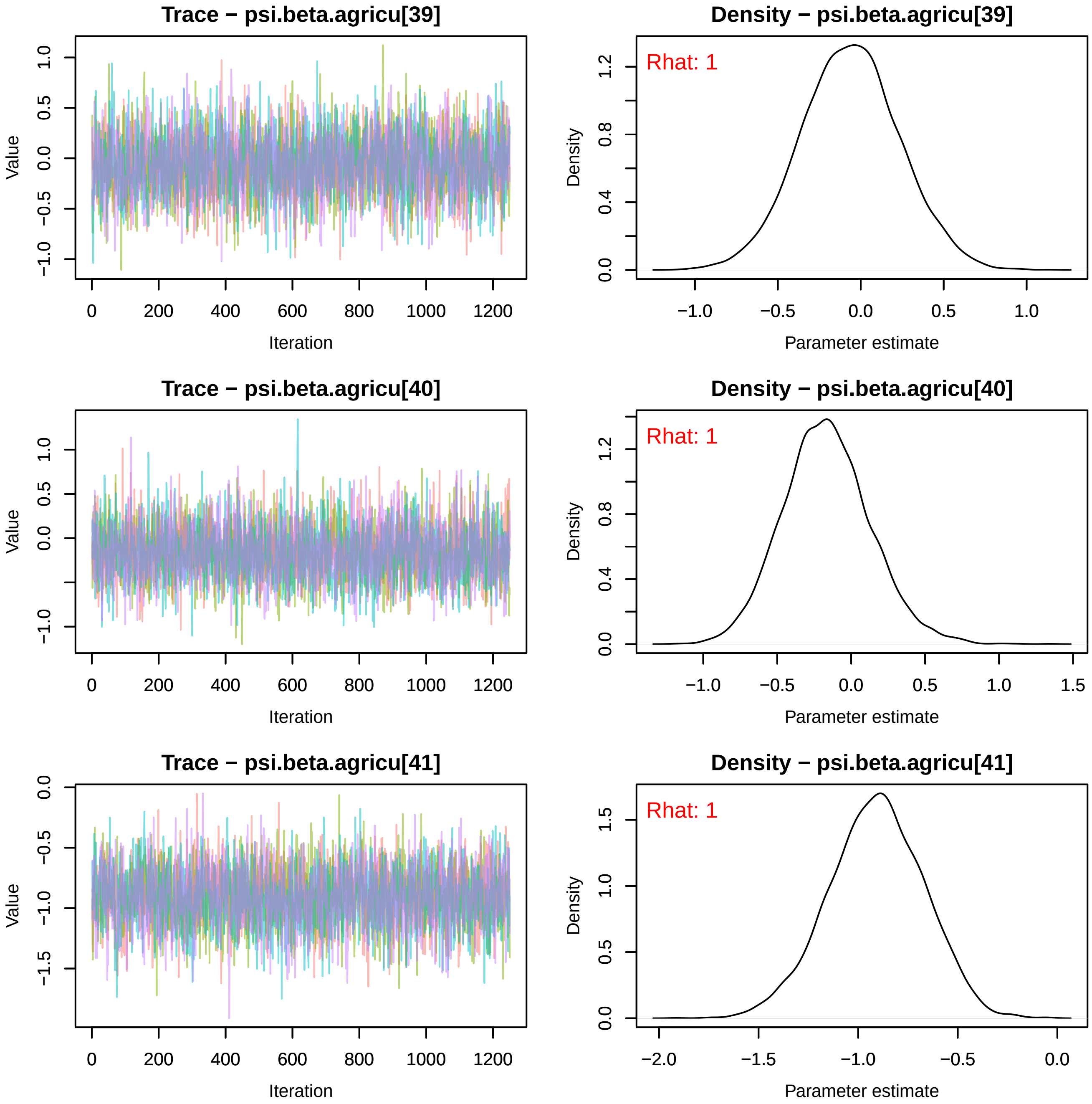
Trace − psi.beta.agricu[39]
Value
Iteration
0
200
400
600
800
1000
1200
−1.0
−0.5
0.0
0.5
1.0
Density − psi.beta.agricu[39]
Rhat: 1
Density
Parameter estimate
−1.0
−0.5
0.0
0.5
1.0
0.0
0.4
0.8
1.2
Trace − psi.beta.agricu[40]
Value
Iteration
0
200
400
600
800
1000
1200
−1.0
0.0
0.5
1.0
Density − psi.beta.agricu[40]
Rhat: 1
Density
Parameter estimate
−1.0
−0.5
0.0
0.5
1.0
1.5
0.0
0.4
0.8
1.2
Trace − psi.beta.agricu[41]
Value
Iteration
0
200
400
600
800
1000
1200
−1.5
−1.0
−0.5
0.0
Density − psi.beta.agricu[41]
Rhat: 1
Density
Parameter estimate
−2.0
−1.5
−1.0
−0.5
0.0
0.0
0.5
1.0
1.5

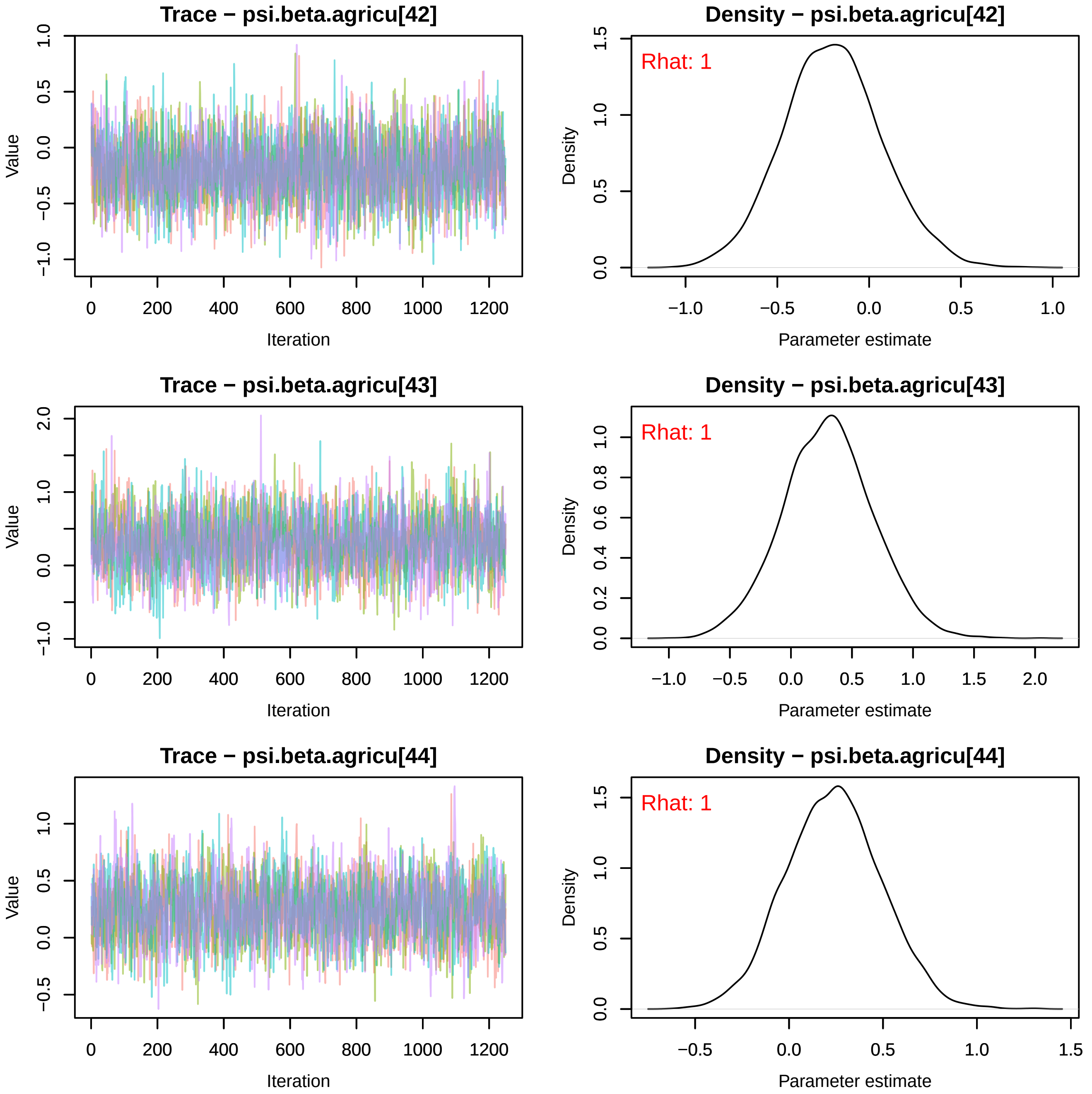
Trace − psi.beta.agricu[42]
Value
Iteration
Density − psi.beta.agricu[42]
Rhat: 1
Density
Parameter estimate
Trace − psi.beta.agricu[43]
Value
Iteration
Density − psi.beta.agricu[43]
Rhat: 1
Density
Parameter estimate
Trace − psi.beta.agricu[44]
Value
Iteration
Density − psi.beta.agricu[44]
Rhat: 1
Density
Parameter estimate

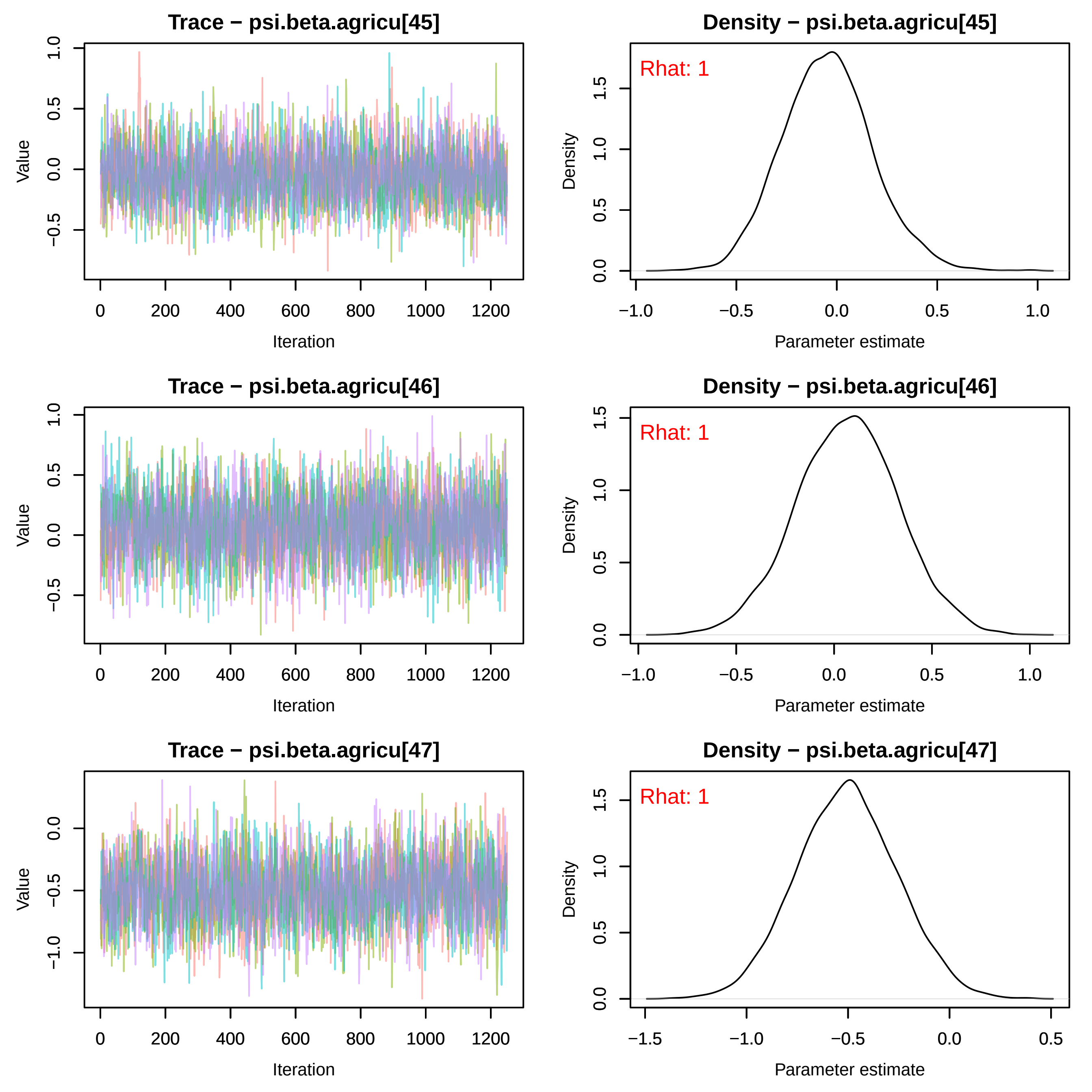

Trace − psi.beta.agricu[45]
Value
Iteration
Density − psi.beta.agricu[45]
Rhat: 1
Density
Parameter estimate
Trace − psi.beta.agricu[46]
Value
Iteration
Density − psi.beta.agricu[46]
Rhat: 1
Density
Parameter estimate
Trace − psi.beta.agricu[47]
Value
Iteration
Density − psi.beta.agricu[47]
Rhat: 1
Density
Parameter estimate

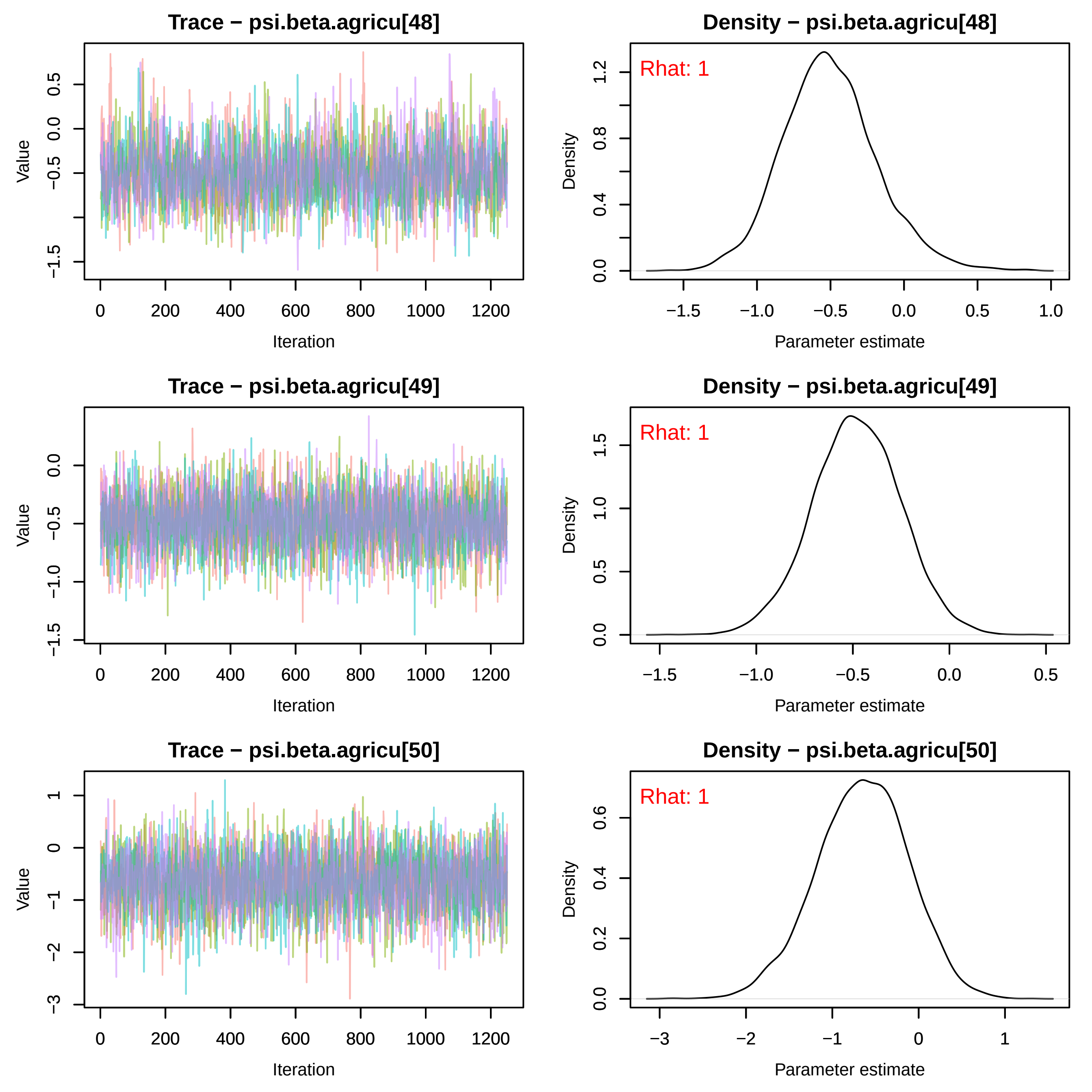
Trace − psi.beta.agricu[48]
Density − psi.beta.agricu[48]
Rhat: 1
Trace − psi.beta.agricu[49]
Density − psi.beta.agricu[49]
Rhat: 1
Trace − psi.beta.agricu[50]
Density − psi.beta.agricu[50]
Rhat: 1
Value
Iteration
Density
Parameter estimate
0
200
400
600
800
1000
1200
−1.5
−1.0
−0.5
0.0
0.5
1.0
−3
−2
−1
1
0.2
0.4
0.6
0.8
1.2
1.5

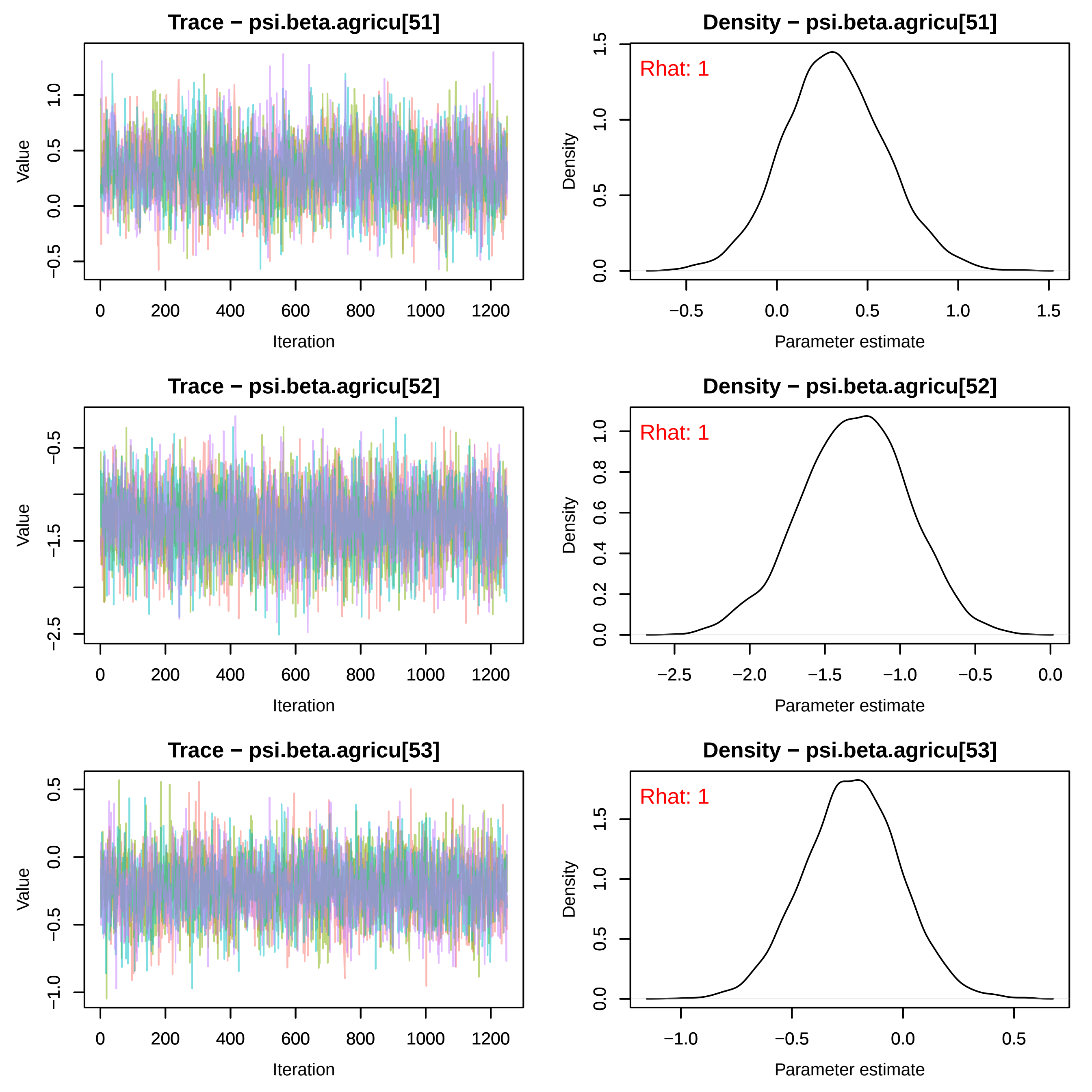

Trace − psi.beta.agricu[51]
Density − psi.beta.agricu[51]
Rhat: 1
Trace − psi.beta.agricu[52]
Density − psi.beta.agricu[52]
Rhat: 1
Trace − psi.beta.agricu[53]
Density − psi.beta.agricu[53]
Rhat: 1
Value
Iteration
Density
Parameter estimate

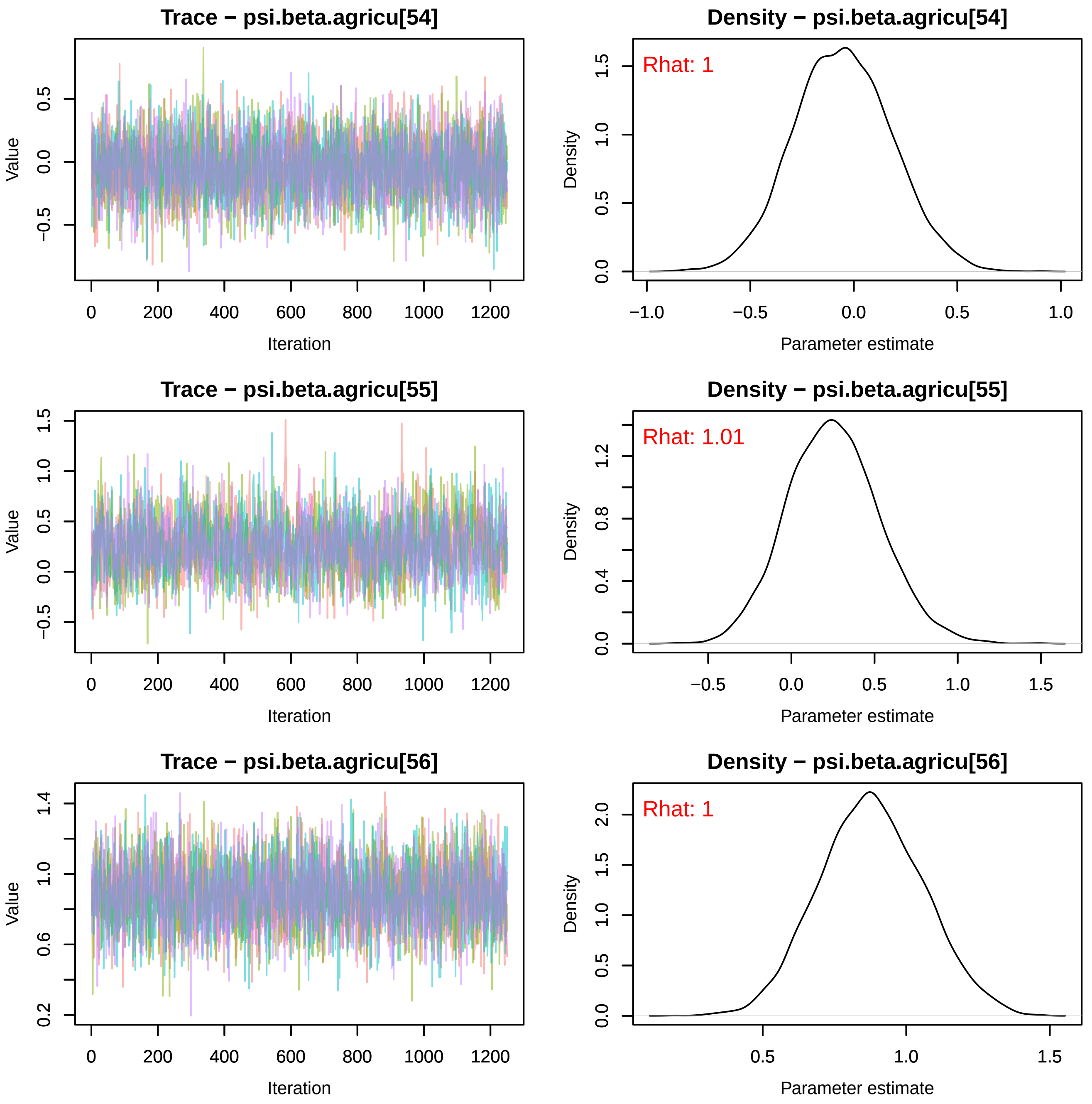

Trace − psi.beta.agricu[54]
Density − psi.beta.agricu[54]
Rhat: 1
Trace − psi.beta.agricu[55]
Density − psi.beta.agricu[55]
Rhat: 1.01
Trace − psi.beta.agricu[56]
Density − psi.beta.agricu[56]
Rhat: 1
Value
Iteration
Density
Parameter estimate

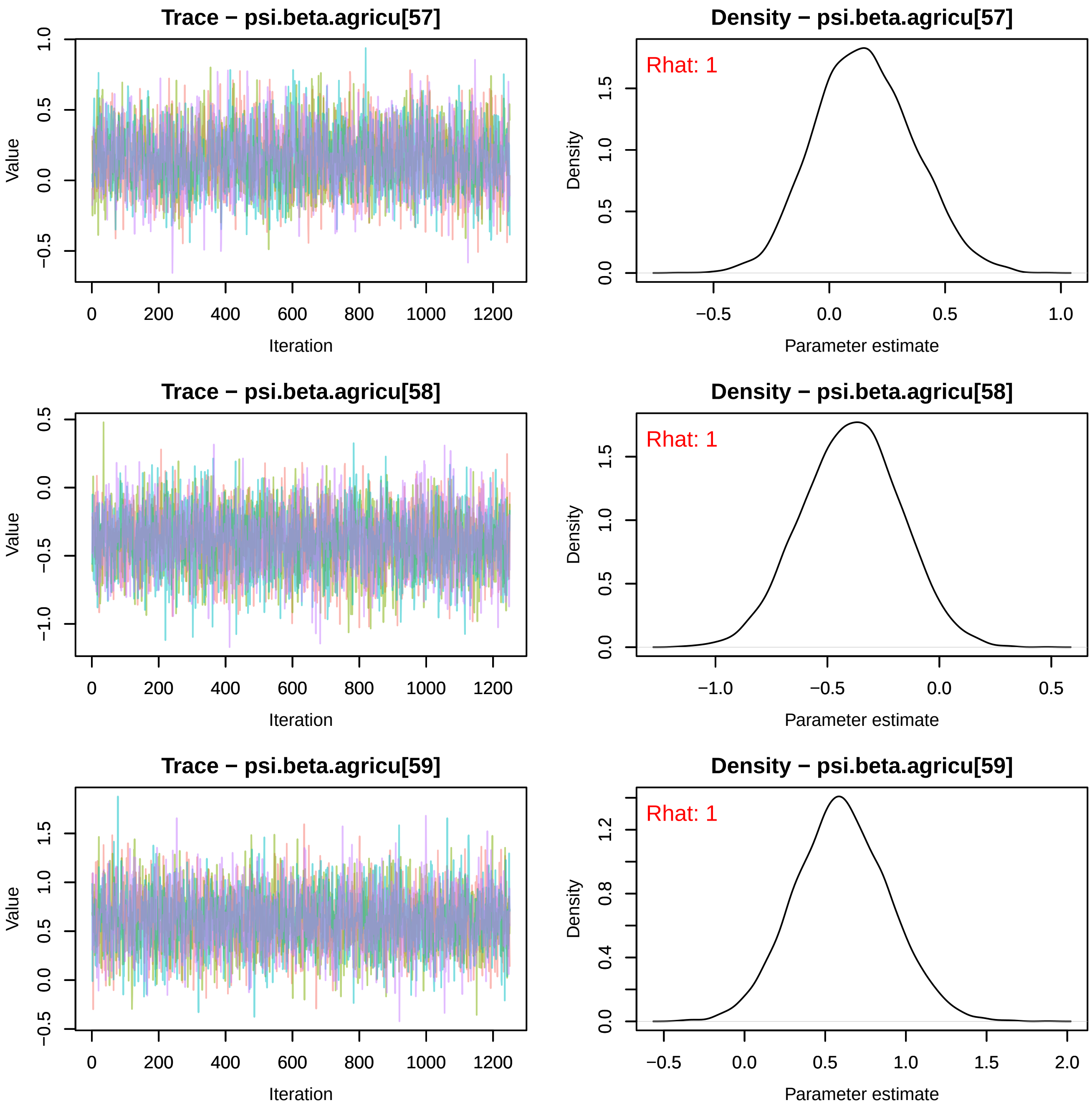

Trace − psi.beta.agricu[57]
Density − psi.beta.agricu[57]
Rhat: 1
Trace − psi.beta.agricu[58]
Density − psi.beta.agricu[58]
Rhat: 1
Trace − psi.beta.agricu[59]
Density − psi.beta.agricu[59]
Rhat: 1
Value
Iteration
Density
Parameter estimate

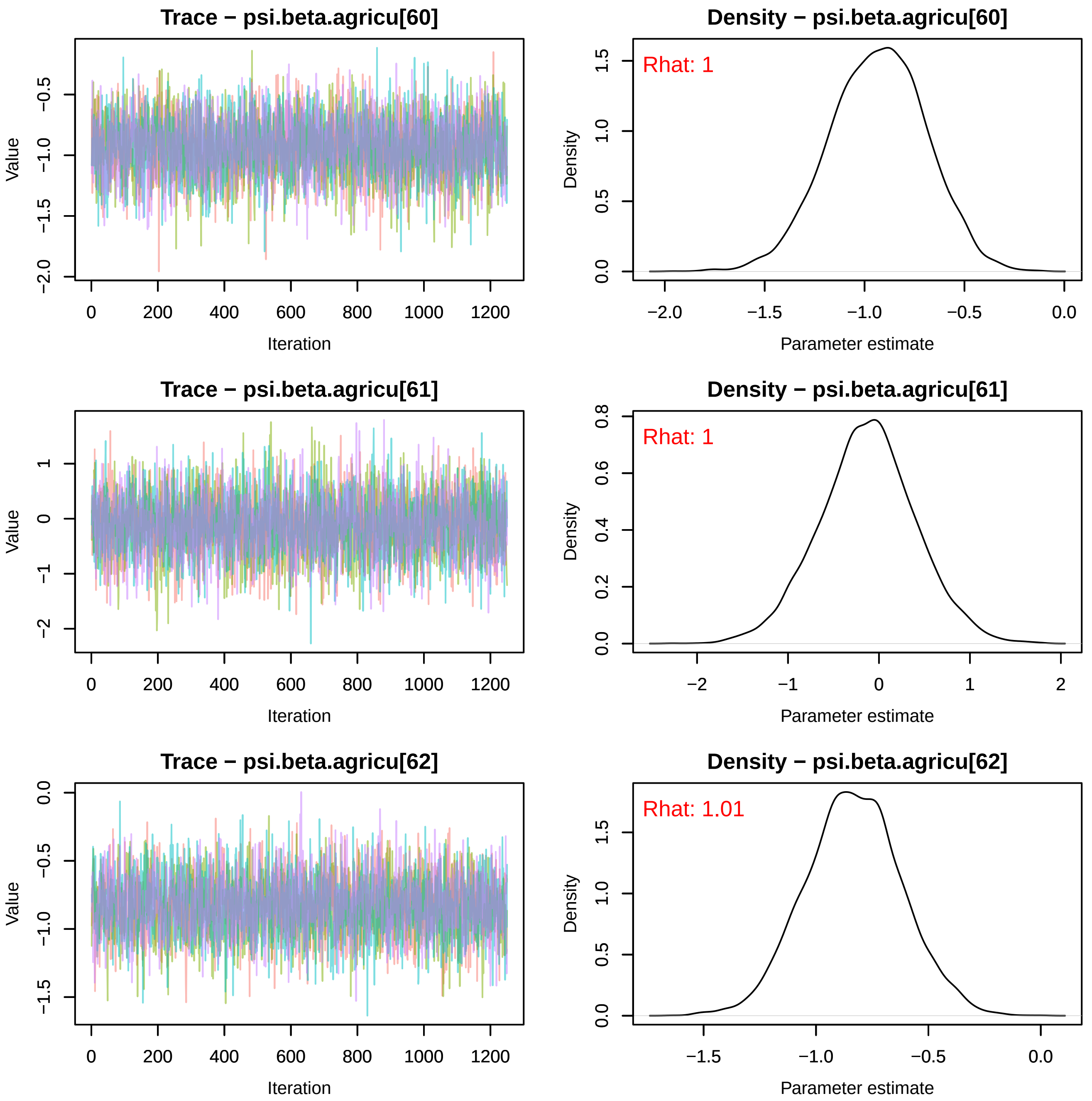
Trace − psi.beta.agricu[60]
Density − psi.beta.agricu[60]
Rhat: 1
Trace − psi.beta.agricu[61]
Density − psi.beta.agricu[61]
Rhat: 1
Trace − psi.beta.agricu[62]
Density − psi.beta.agricu[62]
Rhat: 1.01
Value
Iteration
Density
Parameter estimate

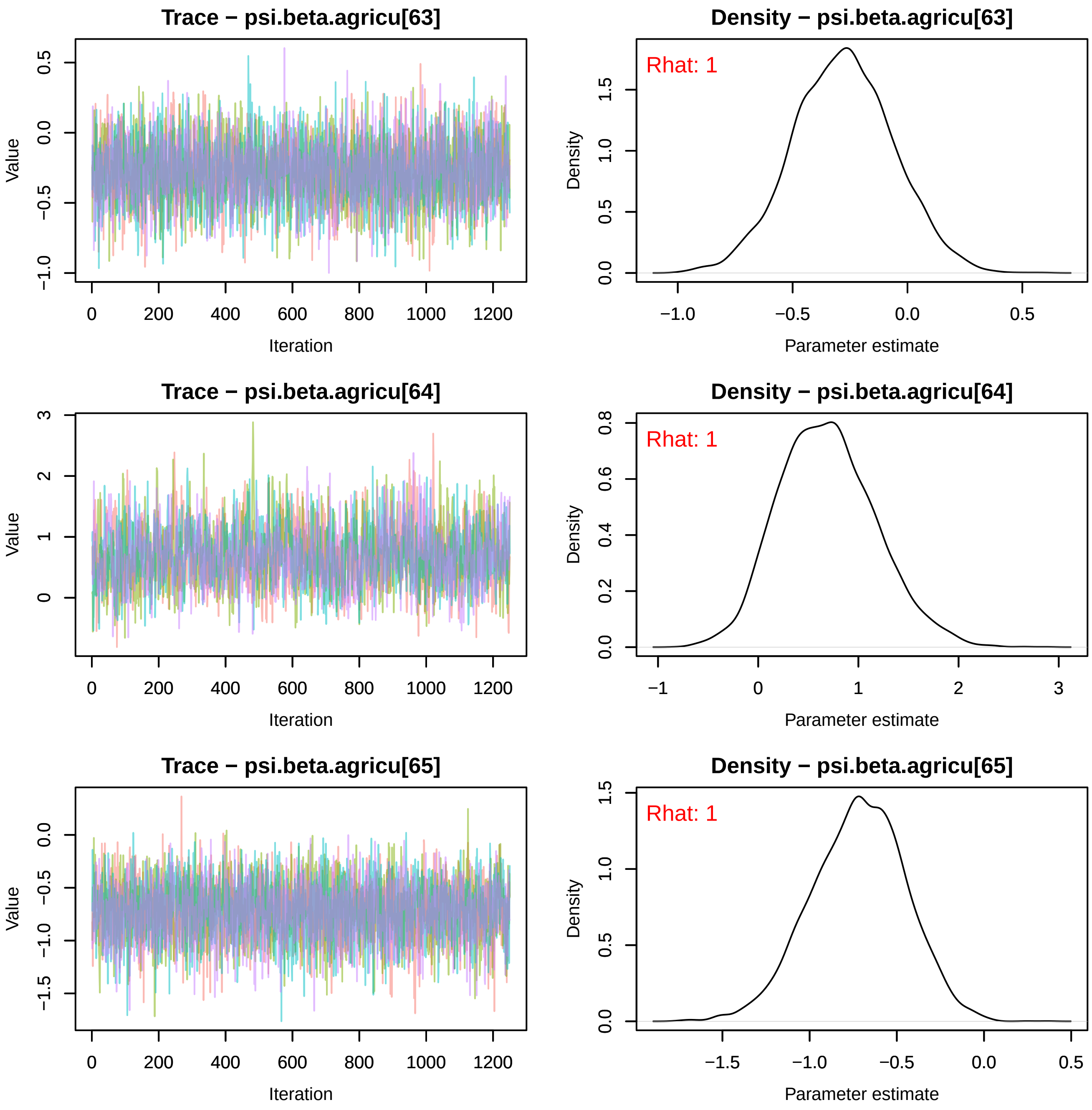
Trace − psi.beta.agricu[63]
Value
Iteration
Density − psi.beta.agricu[63]
Rhat: 1
Density
Parameter estimate
Trace − psi.beta.agricu[64]
Value
Iteration
Density − psi.beta.agricu[64]
Rhat: 1
Density
Parameter estimate
Trace − psi.beta.agricu[65]
Value
Iteration
Density − psi.beta.agricu[65]
Rhat: 1
Density
Parameter estimate

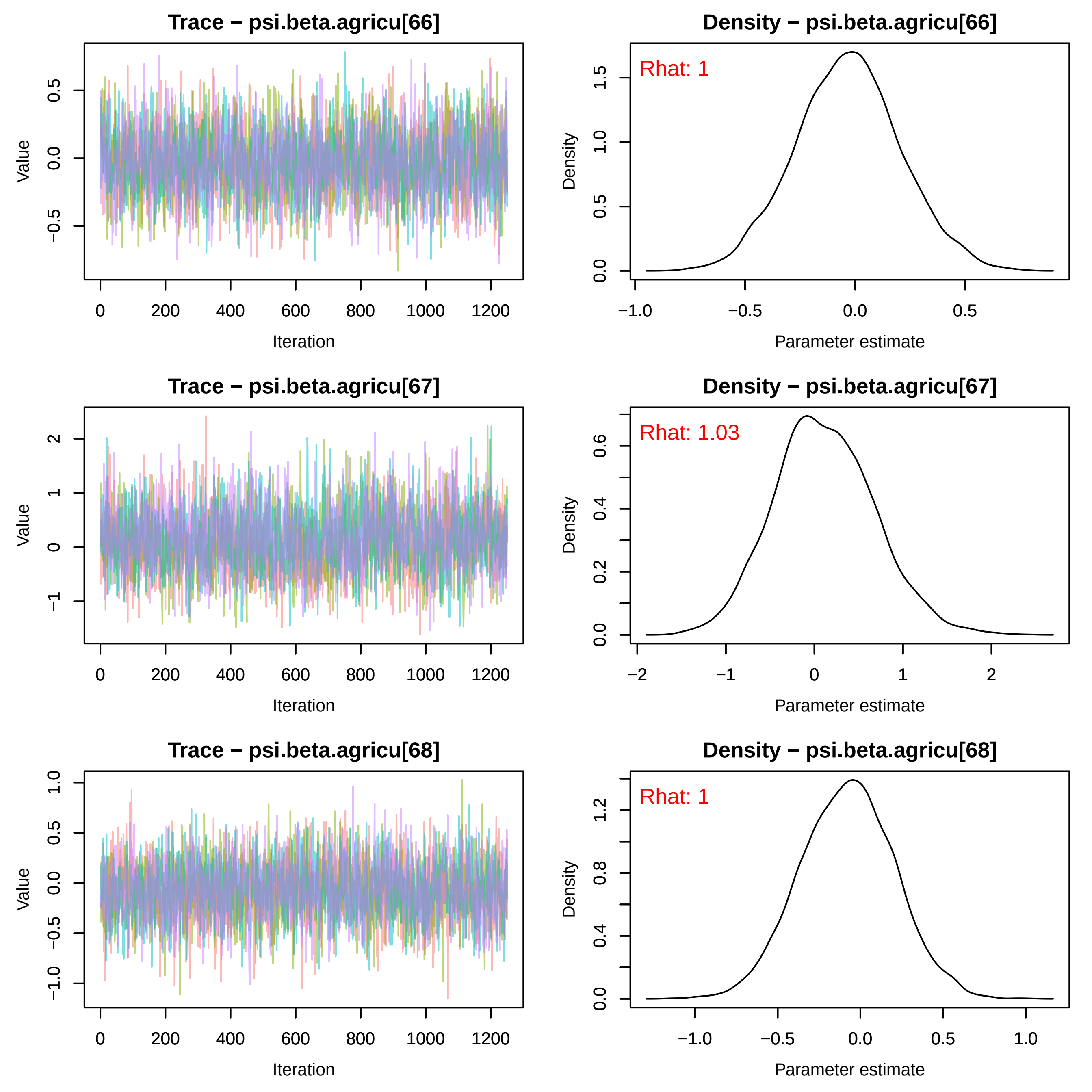
Trace − psi.beta.agricu[66]
Value
Iteration
Density − psi.beta.agricu[66]
Rhat: 1
Density
Parameter estimate
Trace − psi.beta.agricu[67]
Value
Iteration
Density − psi.beta.agricu[67]
Rhat: 1.03
Density
Parameter estimate
Trace − psi.beta.agricu[68]
Value
Iteration
Density − psi.beta.agricu[68]
Rhat: 1
Density
Parameter estimate

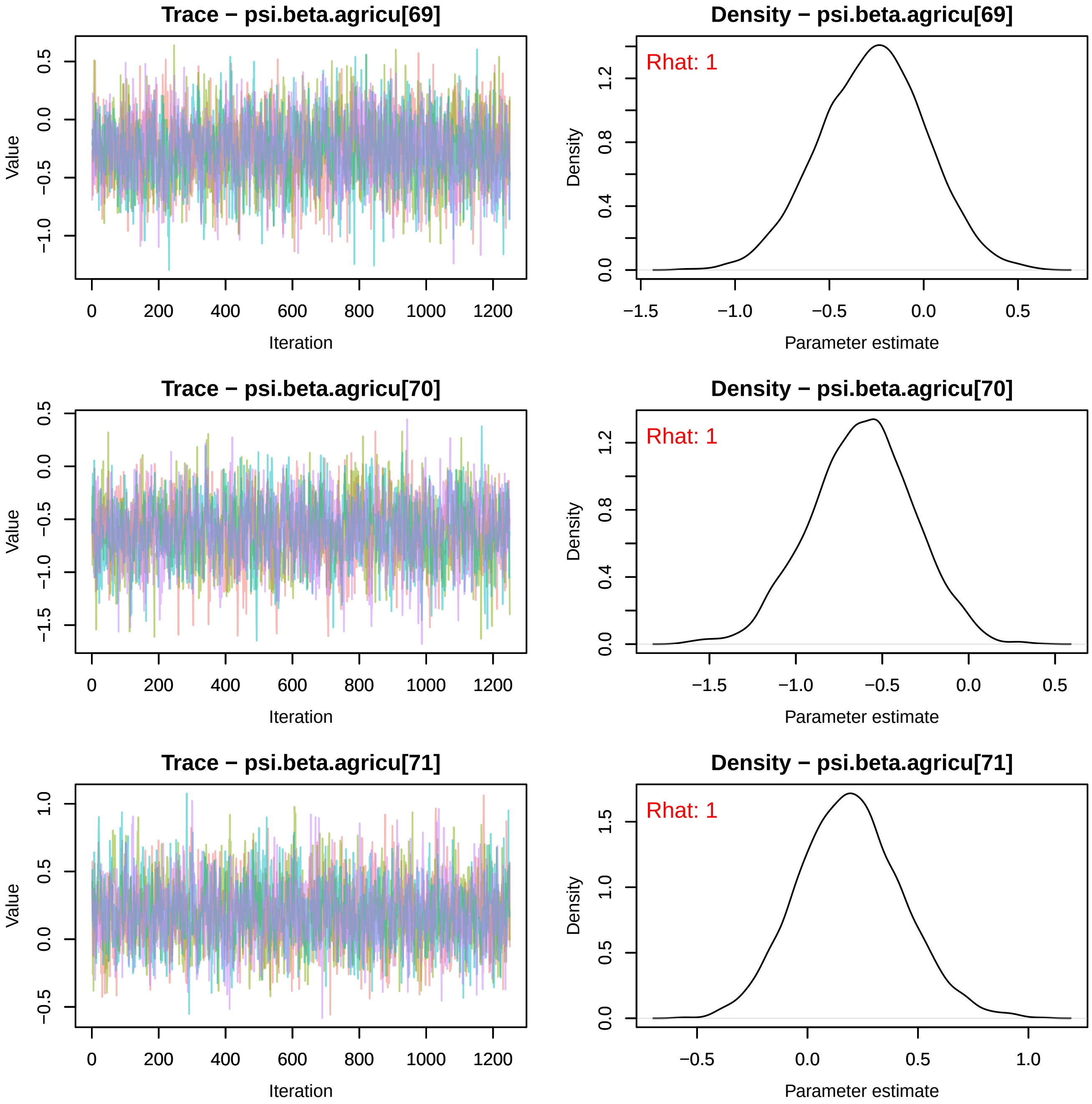

Trace − psi.beta.agricu[69]
Value
0.5
0.0
−0.5
−1.0
Iteration
0
200
400
600
800
1000
1200
Density − psi.beta.agricu[69]
Rhat: 1
Density
0.0
0.4
0.8
1.2
Parameter estimate
−1.5
−1.0
−0.5
0.0
0.5
Trace − psi.beta.agricu[70]
Value
0.5
0.0
−0.5
−1.0
−1.5
Iteration
0
200
400
600
800
1000
1200
Density − psi.beta.agricu[70]
Rhat: 1
Density
0.0
0.4
0.8
1.2
Parameter estimate
−1.5
−1.0
−0.5
0.0
0.5
Trace − psi.beta.agricu[71]
Value
1.0
0.5
0.0
−0.5
Iteration
0
200
400
600
800
1000
1200
Density − psi.beta.agricu[71]
Rhat: 1
Density
0.0
0.5
1.0
1.5
Parameter estimate
−0.5
0.0
0.5
1.0

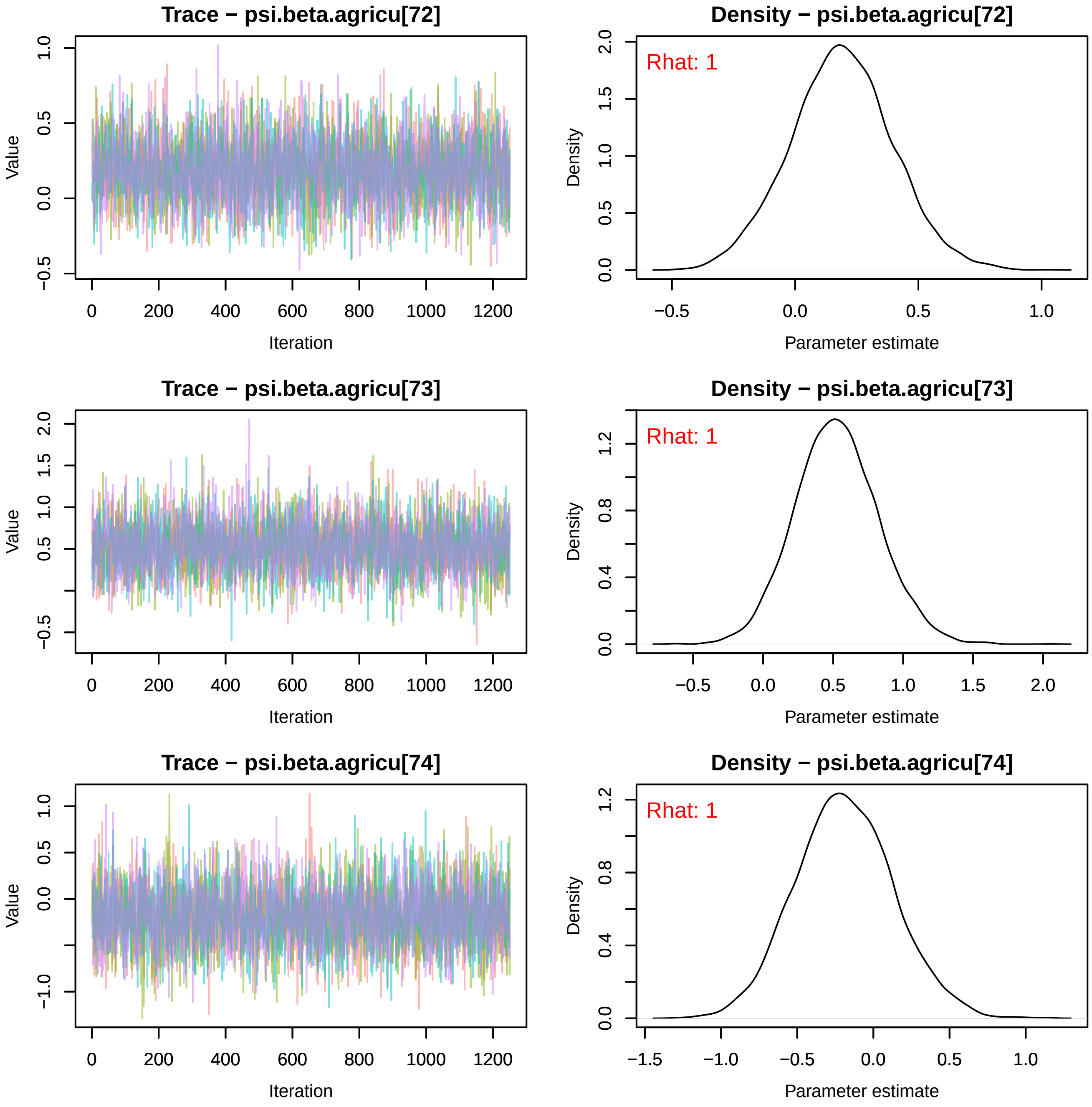
Trace − psi.beta.agricu[72]
Value
Iteration
Density − psi.beta.agricu[72]
Rhat: 1
Density
Parameter estimate
Trace − psi.beta.agricu[73]
Value
Iteration
Density − psi.beta.agricu[73]
Rhat: 1
Density
Parameter estimate
Trace − psi.beta.agricu[74]
Value
Iteration
Density − psi.beta.agricu[74]
Rhat: 1
Density
Parameter estimate

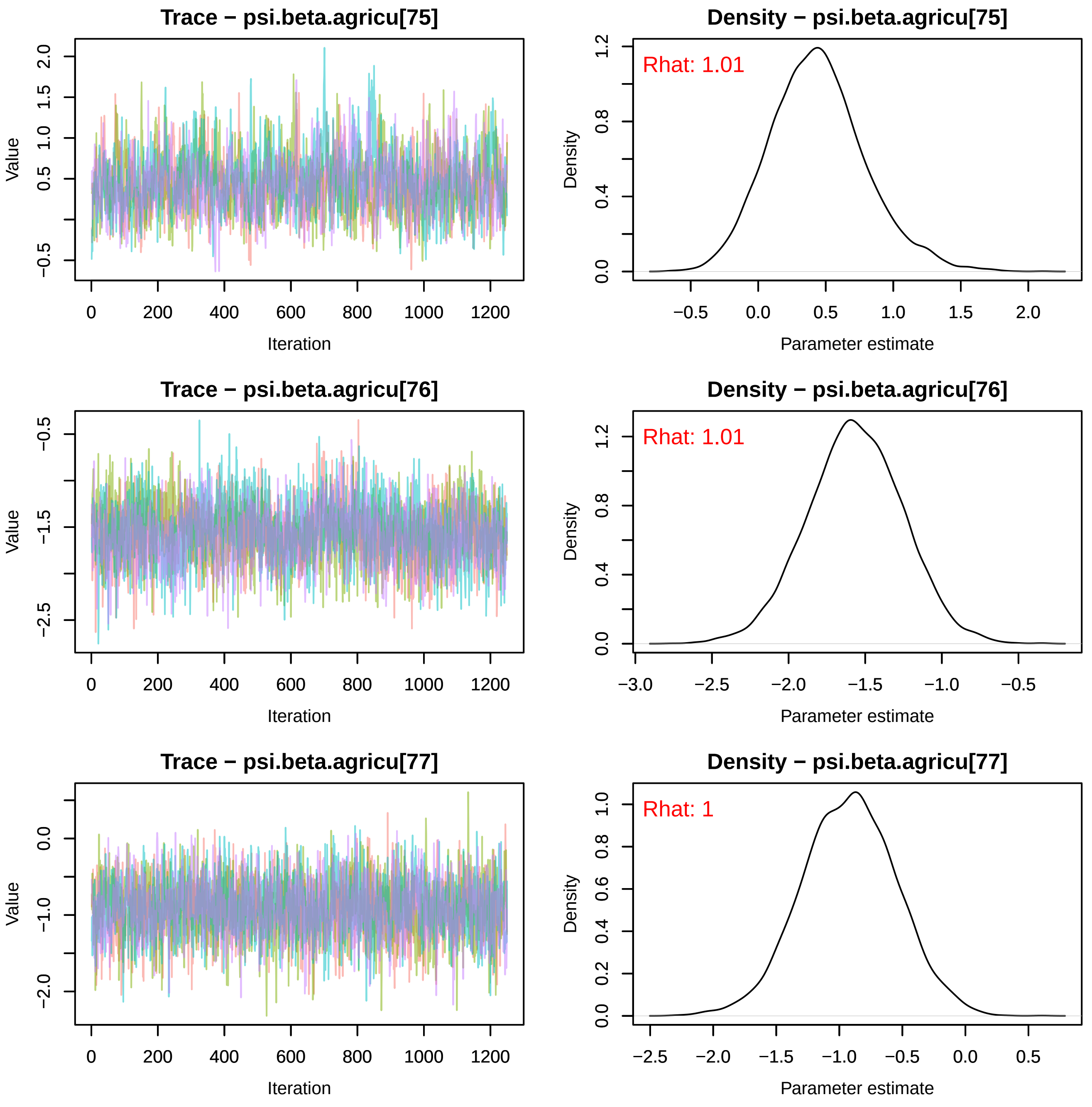
Trace − psi.beta.agricu[75]
Density − psi.beta.agricu[75]
Rhat: 1.01
Trace − psi.beta.agricu[76]
Density − psi.beta.agricu[76]
Rhat: 1.01
Trace − psi.beta.agricu[77]
Density − psi.beta.agricu[77]
Rhat: 1
Value
Iteration
Density
Parameter estimate

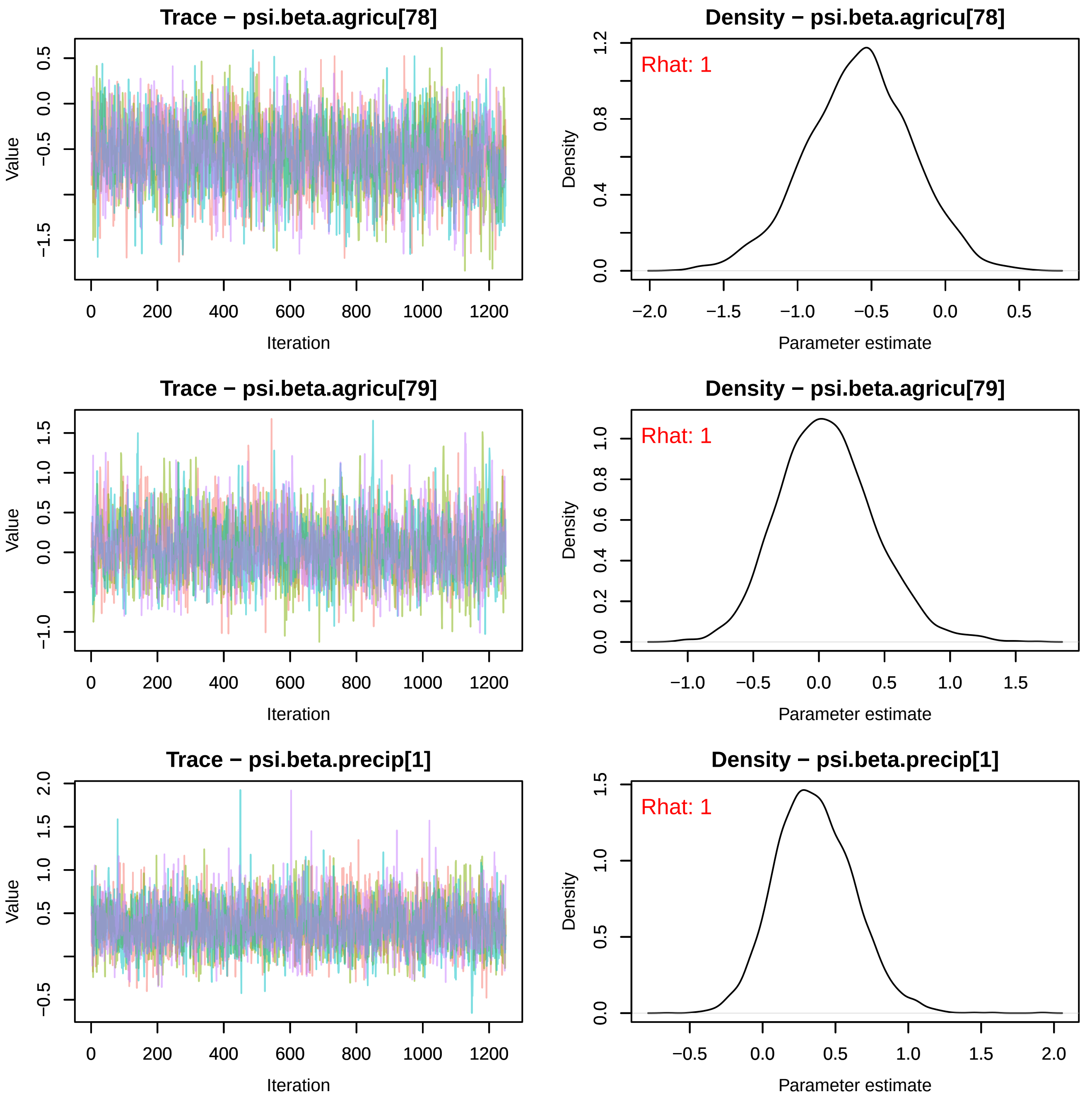
Trace − psi.beta.agricu[78]
Value
Iteration
Density − psi.beta.agricu[78]
Rhat: 1
Density
Parameter estimate
Trace − psi.beta.agricu[79]
Value
Iteration
Density − psi.beta.agricu[79]
Rhat: 1
Density
Parameter estimate
Trace − psi.beta.precip[1]
Value
Iteration
Density − psi.beta.precip[1]
Rhat: 1
Density
Parameter estimate

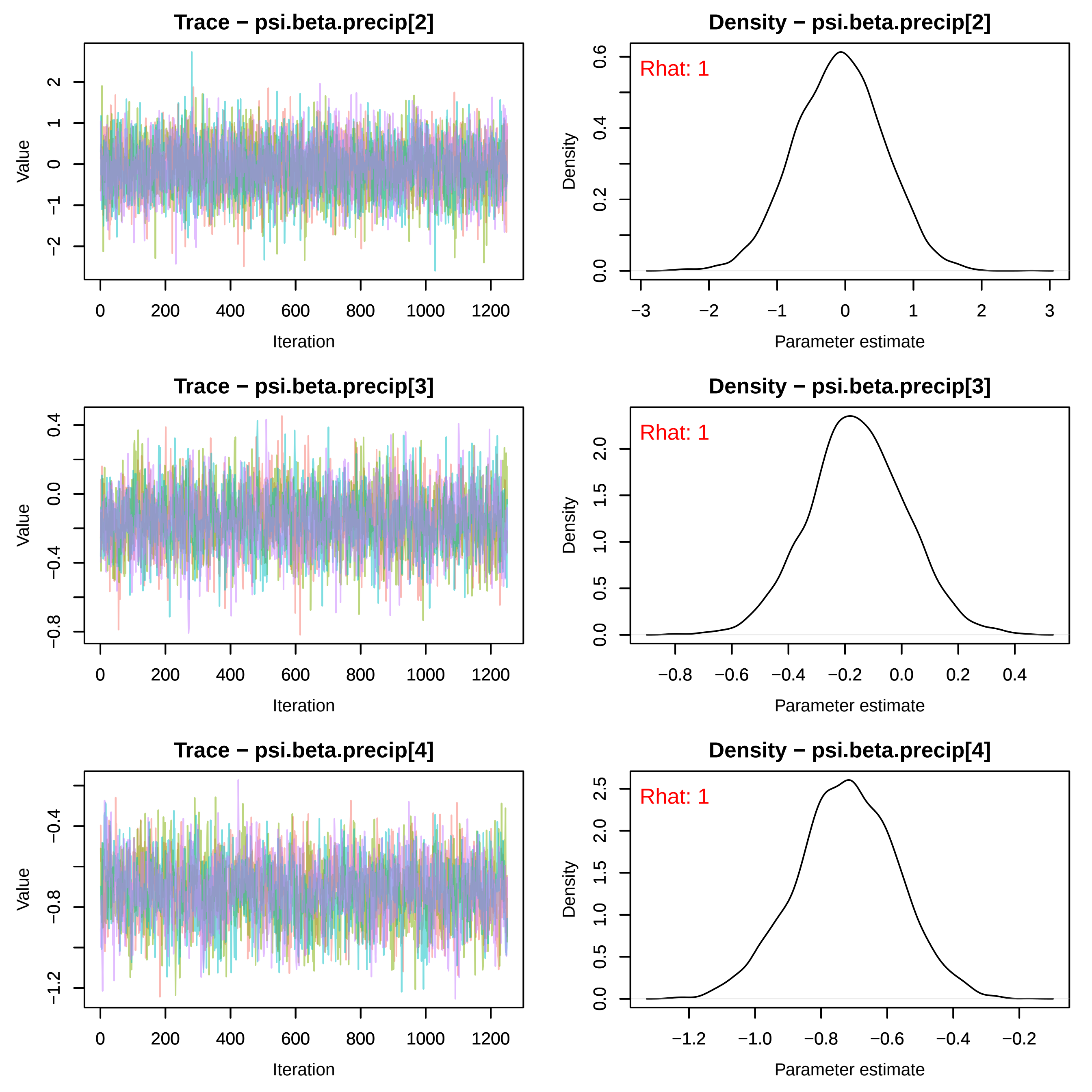

Trace − psi.beta.precip[2]
Value
Iteration
Density − psi.beta.precip[2]
Rhat: 1
Density
Parameter estimate
Trace − psi.beta.precip[3]
Value
Iteration
Density − psi.beta.precip[3]
Rhat: 1
Density
Parameter estimate
Trace − psi.beta.precip[4]
Value
Iteration
Density − psi.beta.precip[4]
Rhat: 1
Density
Parameter estimate

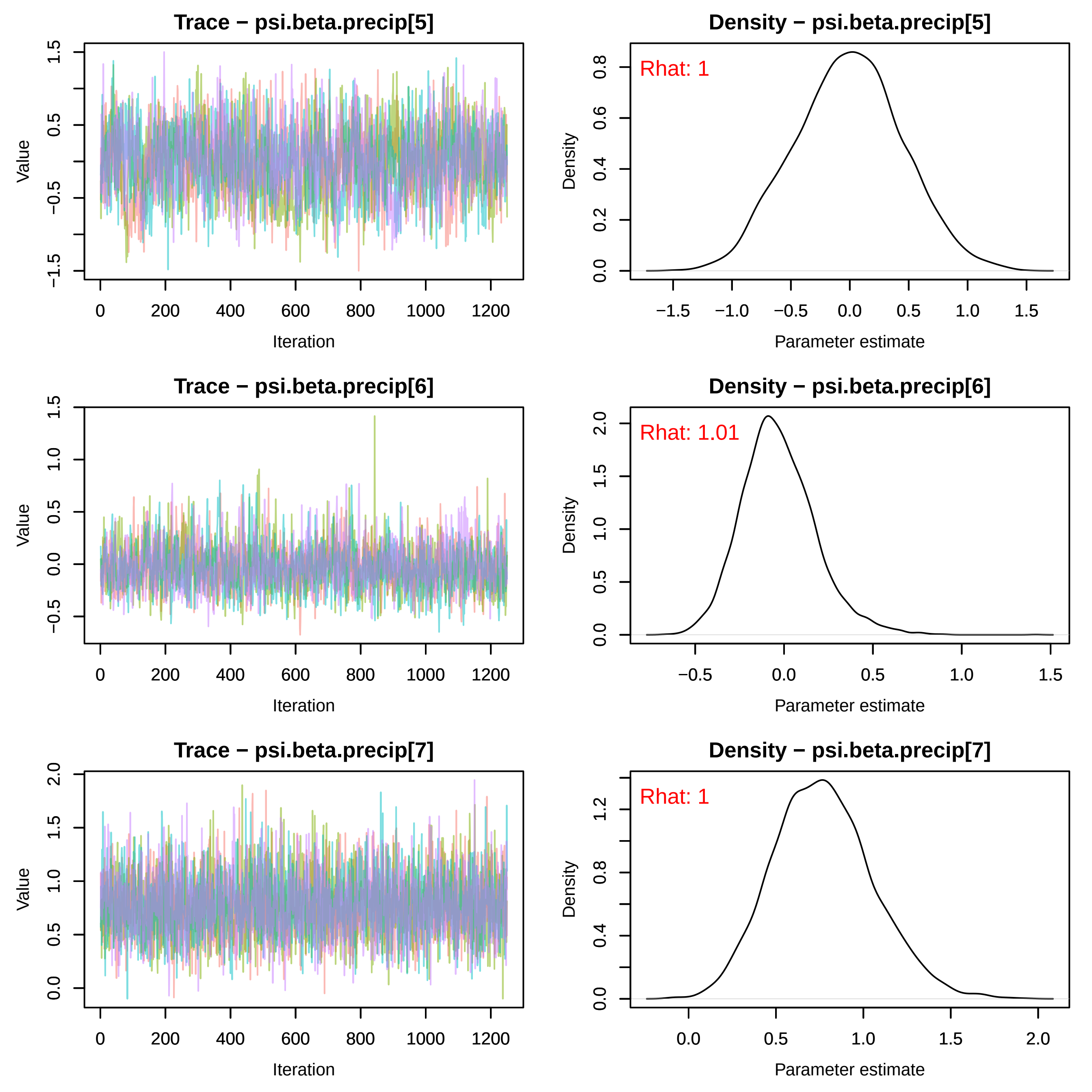
Trace − psi.beta.precip[5]
Value
Iteration
Density − psi.beta.precip[5]
Rhat: 1
Density
Parameter estimate
Trace − psi.beta.precip[6]
Value
Iteration
Density − psi.beta.precip[6]
Rhat: 1.01
Density
Parameter estimate
Trace − psi.beta.precip[7]
Value
Iteration
Density − psi.beta.precip[7]
Rhat: 1
Density
Parameter estimate

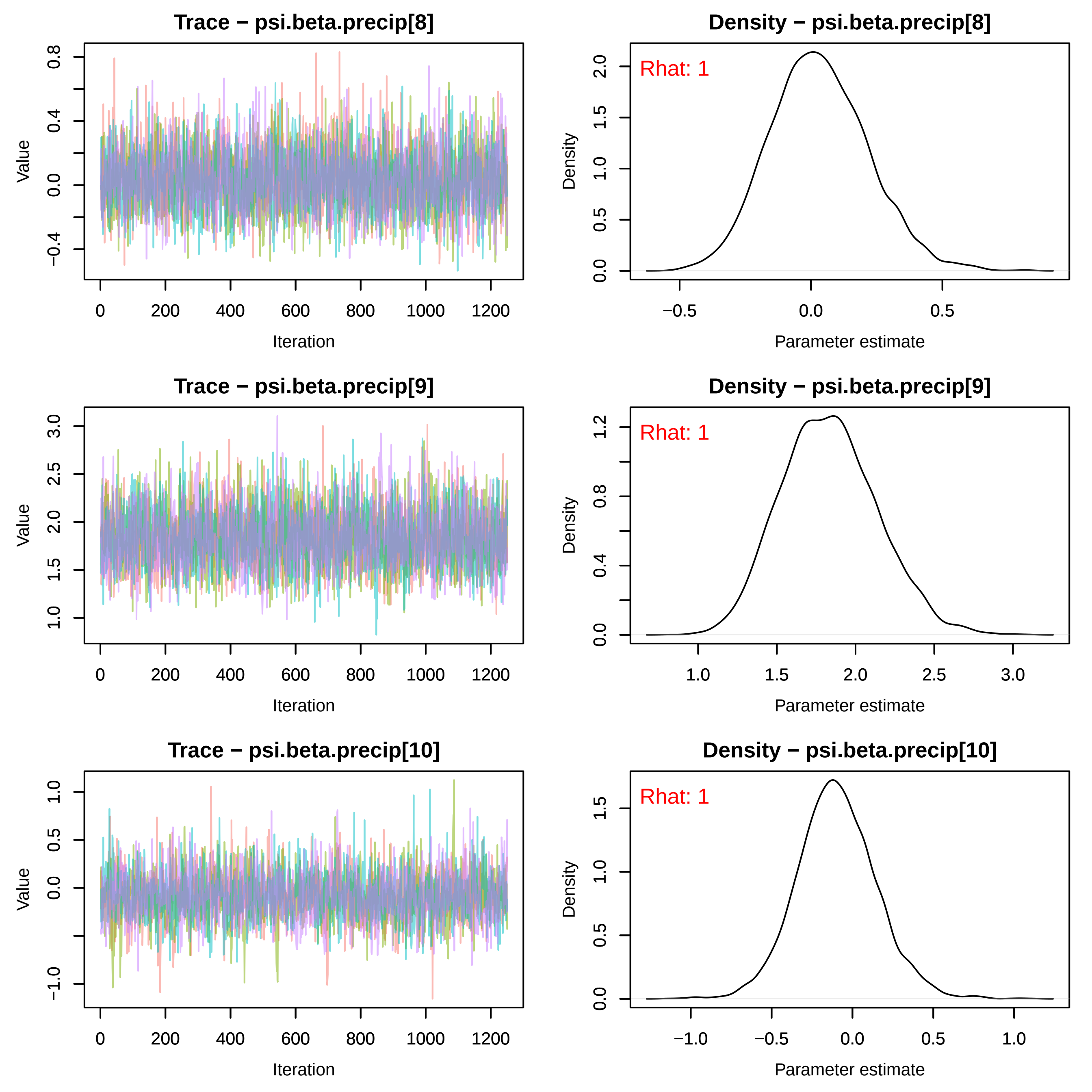
Trace − psi.beta.precip[8]
Value
Iteration
Density − psi.beta.precip[8]
Rhat: 1
Density
Parameter estimate
Trace − psi.beta.precip[9]
Value
Iteration
Density − psi.beta.precip[9]
Rhat: 1
Density
Parameter estimate
Trace − psi.beta.precip[10]
Value
Iteration
Density − psi.beta.precip[10]
Rhat: 1
Density
Parameter estimate

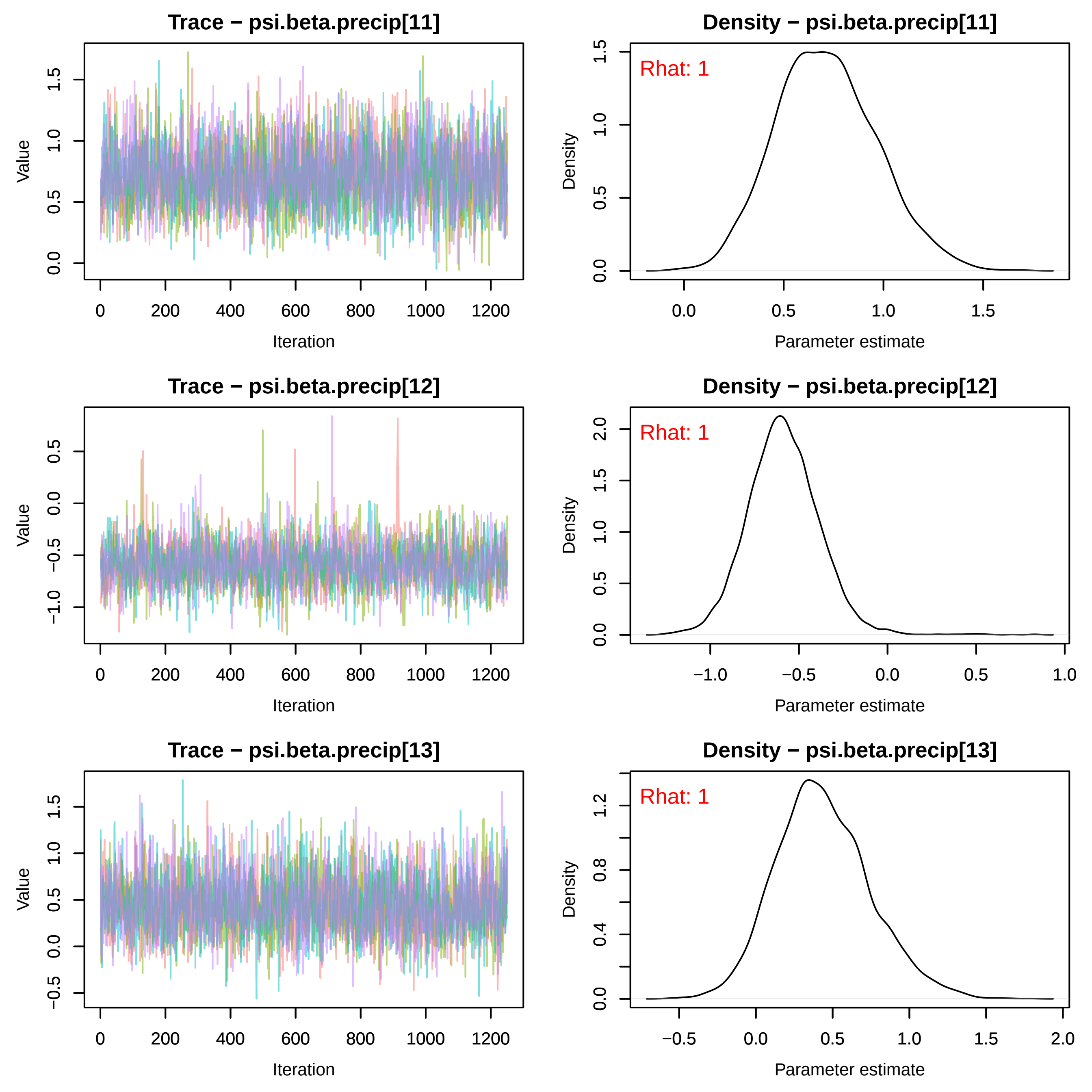

Trace − psi.beta.precip[11]
Value
Iteration
0 200 400 600 800 1000 1200
0.0 0.5 1.0 1.5
Density − psi.beta.precip[11]
Rhat: 1
Density
Parameter estimate
0.0 0.5 1.0 1.5
Trace − psi.beta.precip[12]
Value
Iteration
0 200 400 600 800 1000 1200
−1.0 −0.5 0.0 0.5
Density − psi.beta.precip[12]
Rhat: 1
Density
Parameter estimate
−1.0 −0.5 0.0 0.5 1.0
0.0 0.5 1.0 1.5 2.0
Trace − psi.beta.precip[13]
Value
Iteration
0 200 400 600 800 1000 1200
−0.5 0.0 0.5 1.0 1.5
Density − psi.beta.precip[13]
Rhat: 1
Density
Parameter estimate
−0.5 0.0 0.5 1.0 1.5 2.0
0.0 0.4 0.8 1.2

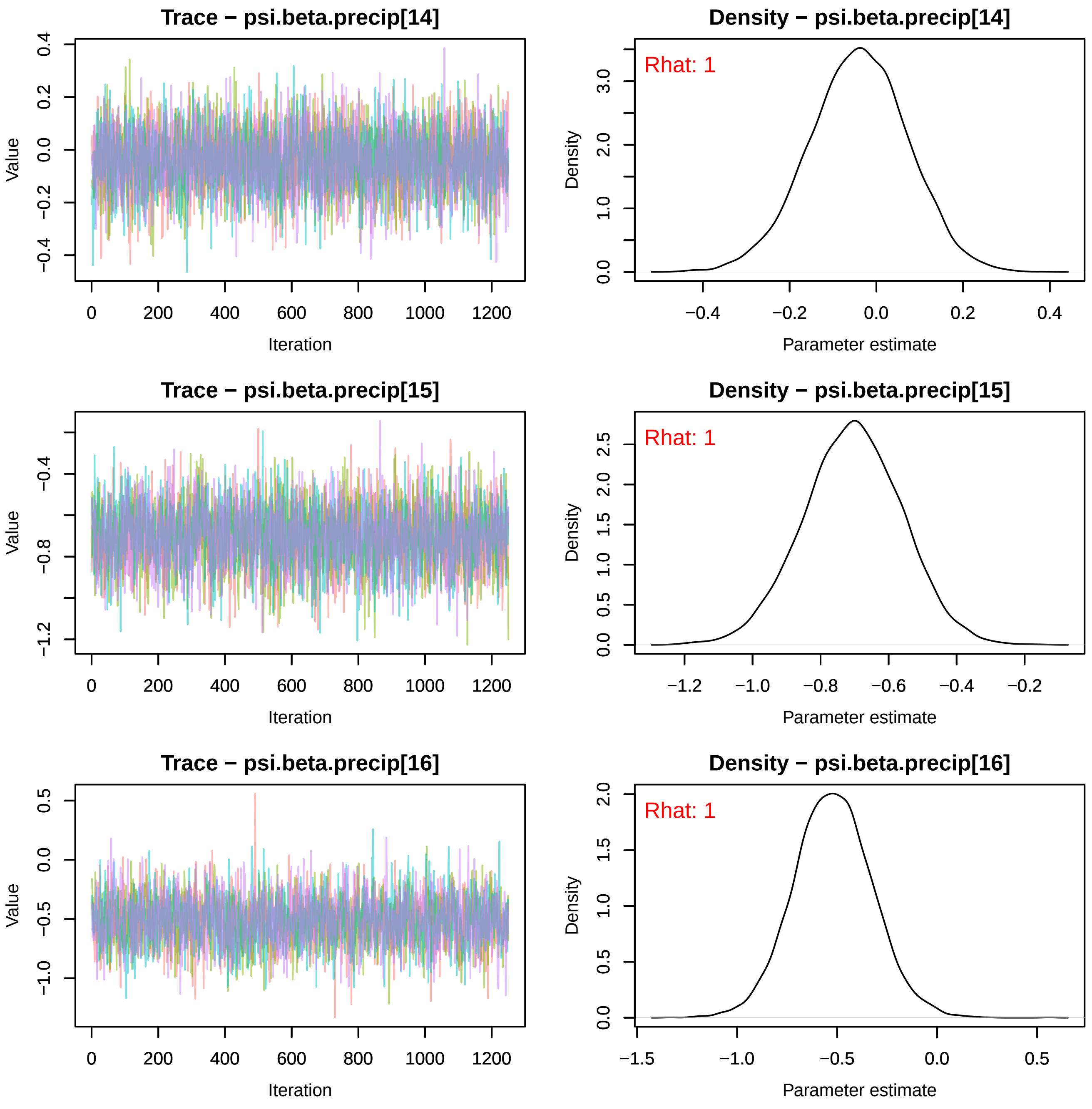
Trace − psi.beta.precip[14]
Value
Iteration
Density − psi.beta.precip[14]
Rhat: 1
Density
Parameter estimate
Trace − psi.beta.precip[15]
Value
Iteration
Density − psi.beta.precip[15]
Rhat: 1
Density
Parameter estimate
Trace − psi.beta.precip[16]
Value
Iteration
Density − psi.beta.precip[16]
Rhat: 1
Density
Parameter estimate

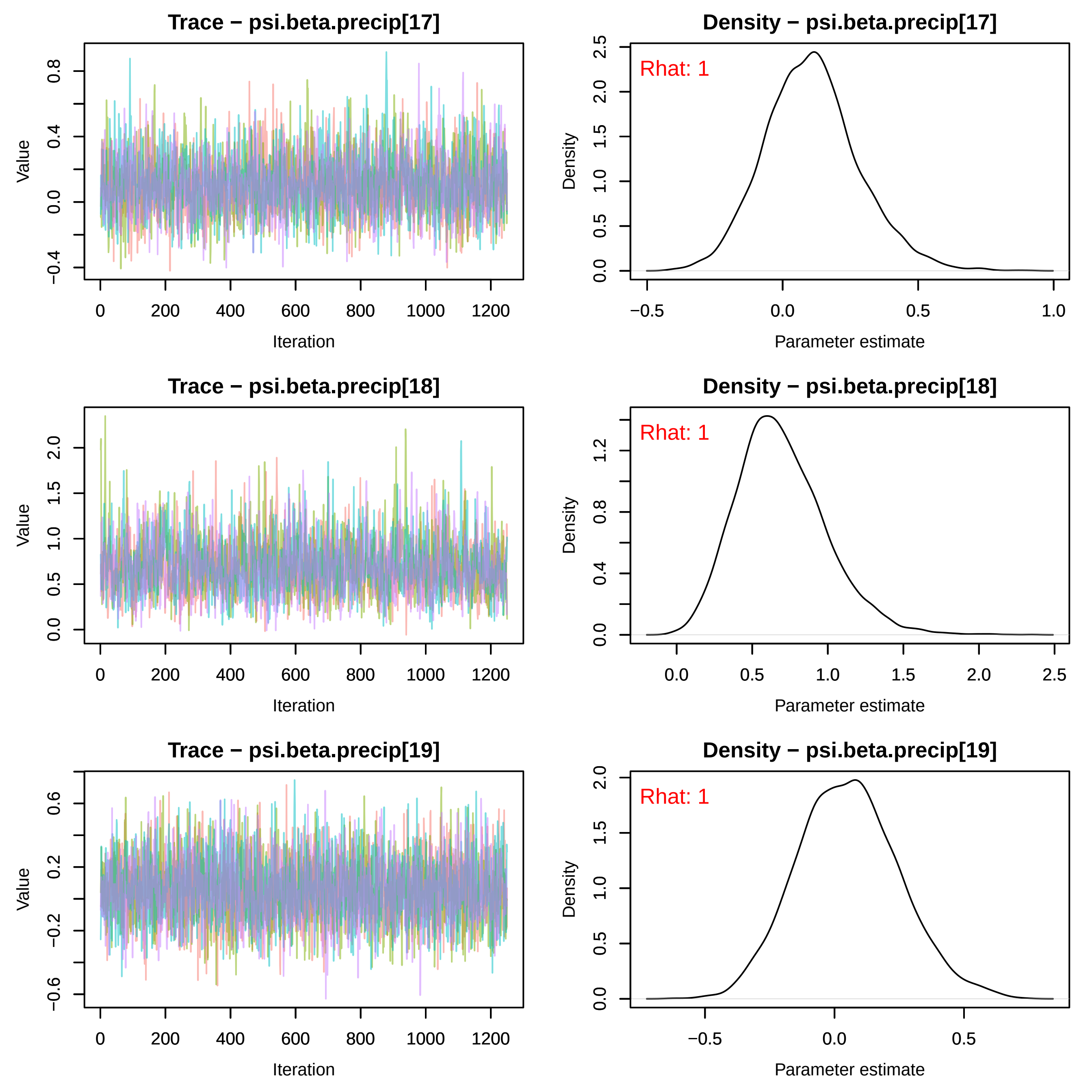
Trace − psi.beta.precip[17]
Density − psi.beta.precip[17]
Rhat: 1
Trace − psi.beta.precip[18]
Density − psi.beta.precip[18]
Rhat: 1
Trace − psi.beta.precip[19]
Density − psi.beta.precip[19]
Rhat: 1
Value
Iteration
Density
Parameter estimate

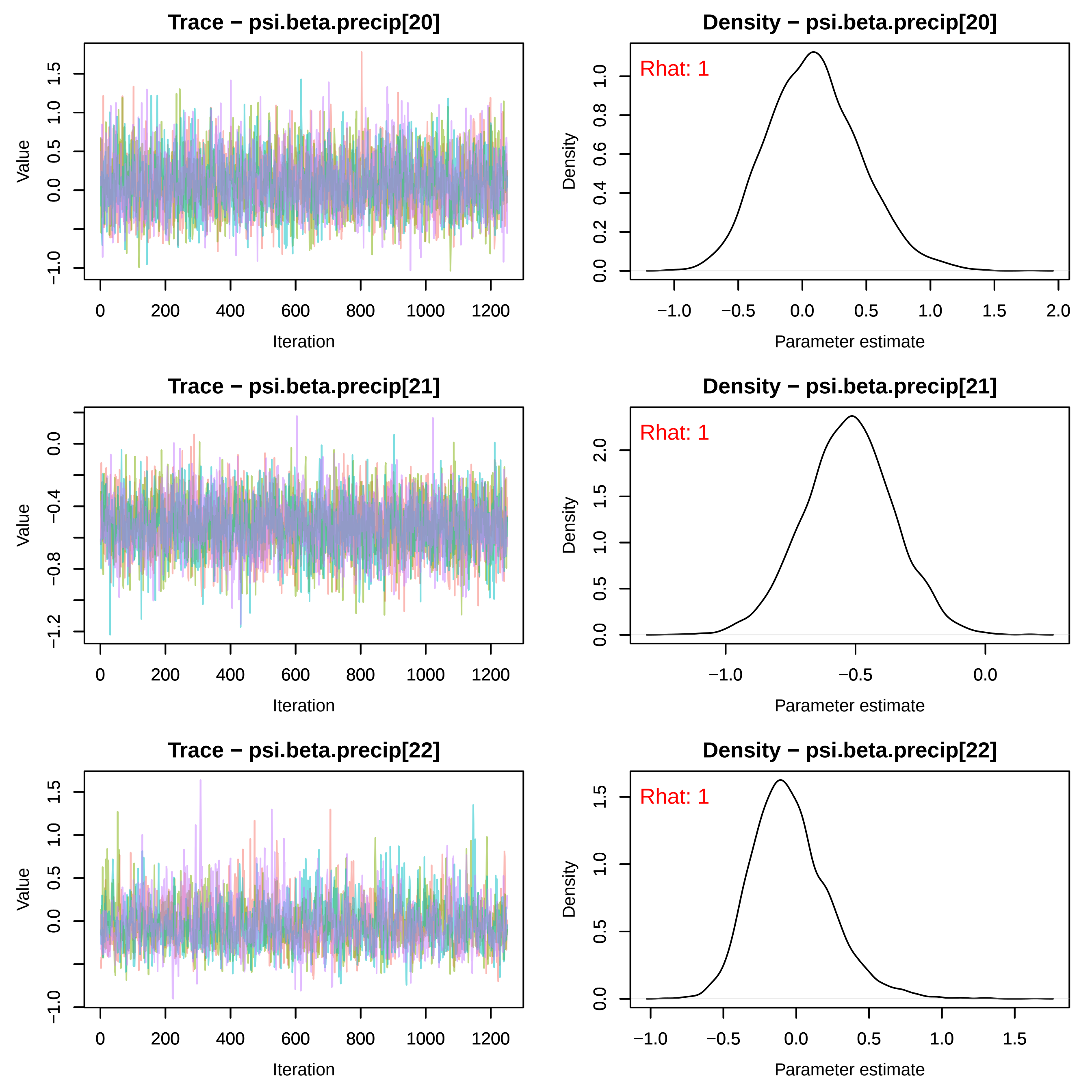

Trace − psi.beta.precip[20]
Value
Iteration
Density − psi.beta.precip[20]
Rhat: 1
Density
Parameter estimate
Trace − psi.beta.precip[21]
Value
Iteration
Density − psi.beta.precip[21]
Rhat: 1
Density
Parameter estimate
Trace − psi.beta.precip[22]
Value
Iteration
Density − psi.beta.precip[22]
Rhat: 1
Density
Parameter estimate

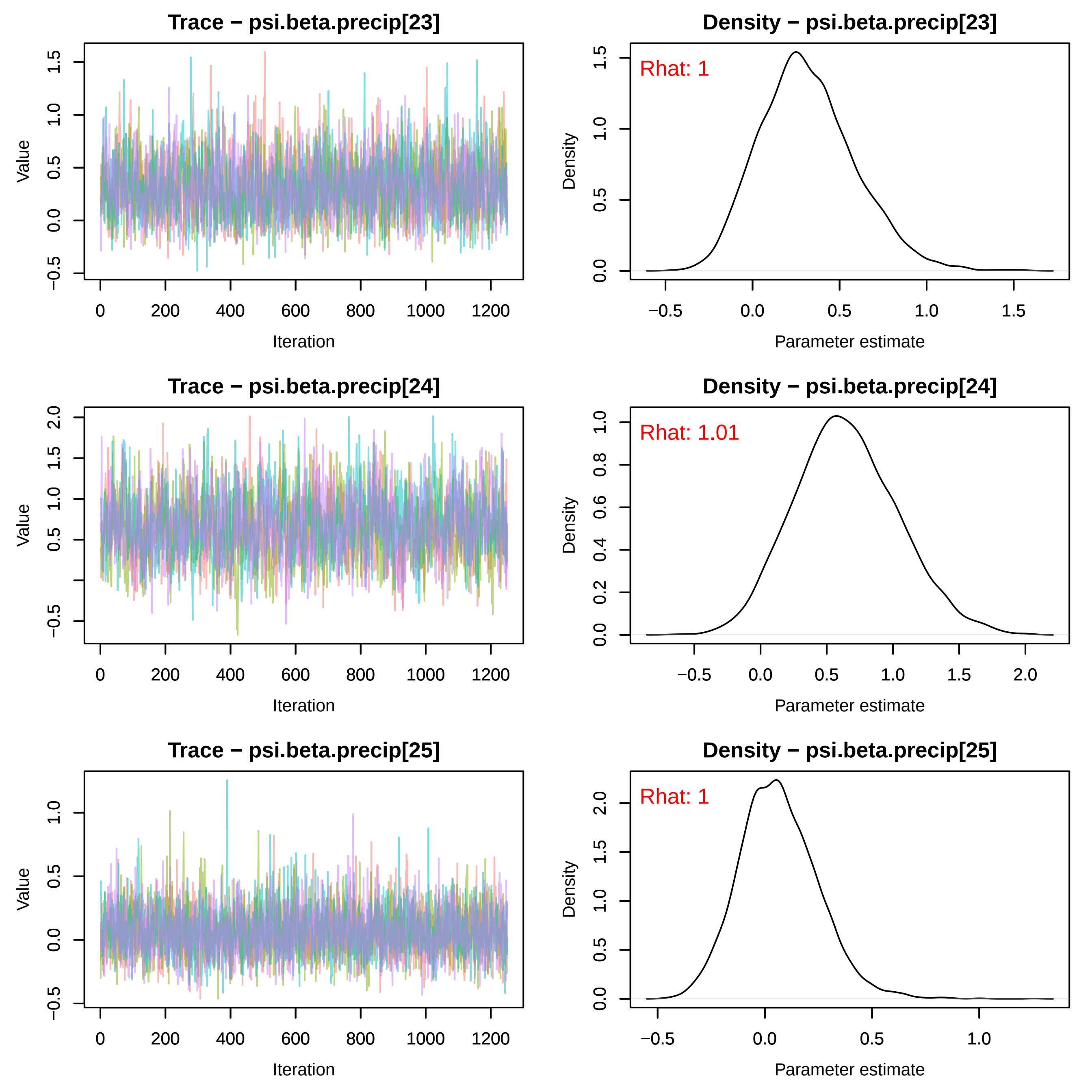
Trace − psi.beta.precip[23]
Value
Iteration
Density − psi.beta.precip[23]
Rhat: 1
Density
Parameter estimate
Trace − psi.beta.precip[24]
Value
Iteration
Density − psi.beta.precip[24]
Rhat: 1.01
Density
Parameter estimate
Trace − psi.beta.precip[25]
Value
Iteration
Density − psi.beta.precip[25]
Rhat: 1
Density
Parameter estimate

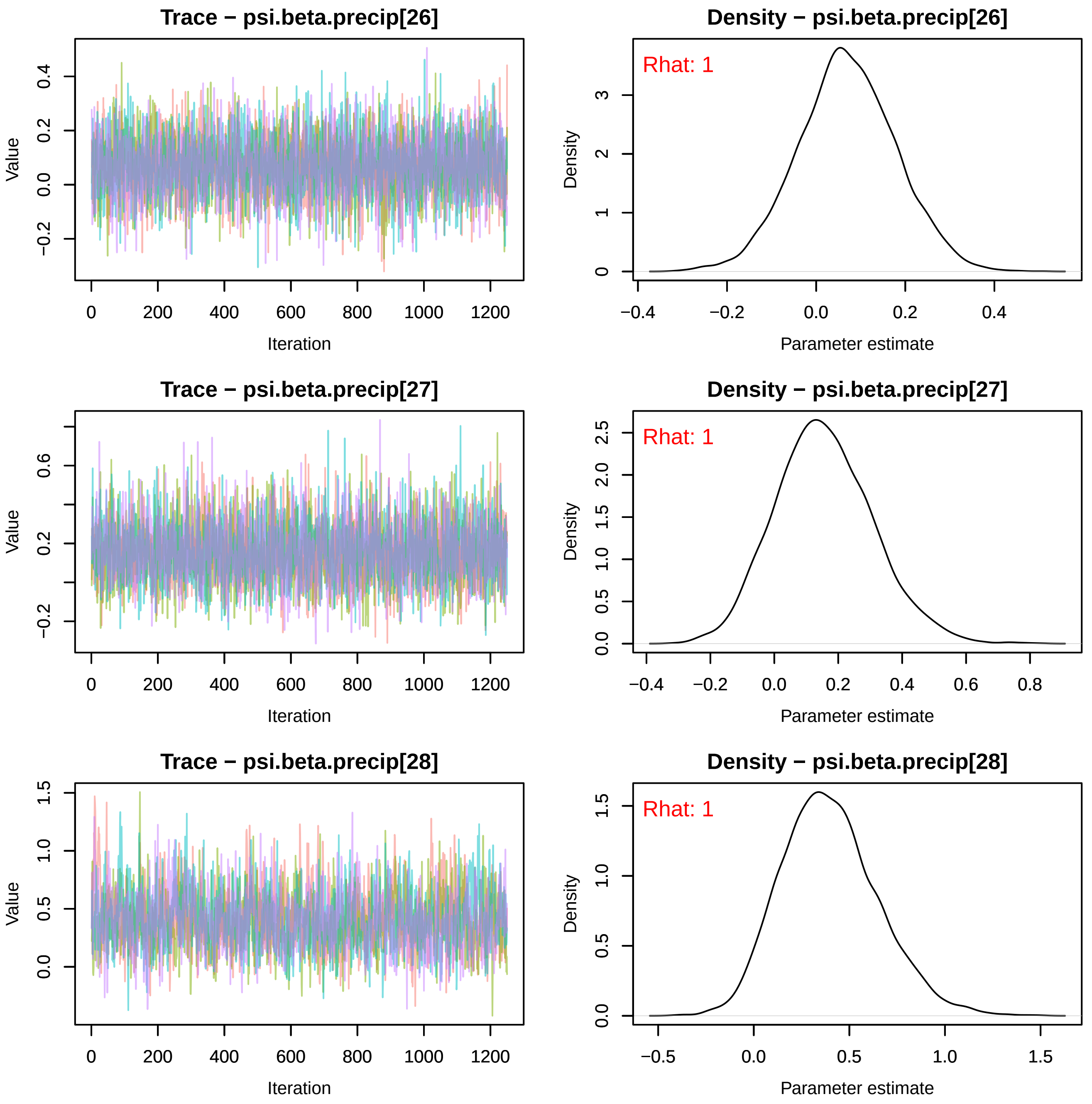
Trace − psi.beta.precip[26]
Density − psi.beta.precip[26]
Rhat: 1
Value
Iteration
Density
Parameter estimate
0
200
400
600
800
1000
1200
−0.2
0.0
0.2
0.4
0
1
2
3
−0.4
Trace − psi.beta.precip[27]
Density − psi.beta.precip[27]
Rhat: 1
Value
Iteration
Density
Parameter estimate
−0.2
0.2
0.6
0.0
0.5
1.0
1.5
2.0
2.5
0.8
Trace − psi.beta.precip[28]
Density − psi.beta.precip[28]
Rhat: 1
Value
Iteration
Density
Parameter estimate
0.0
0.5
1.0
1.5
−0.5

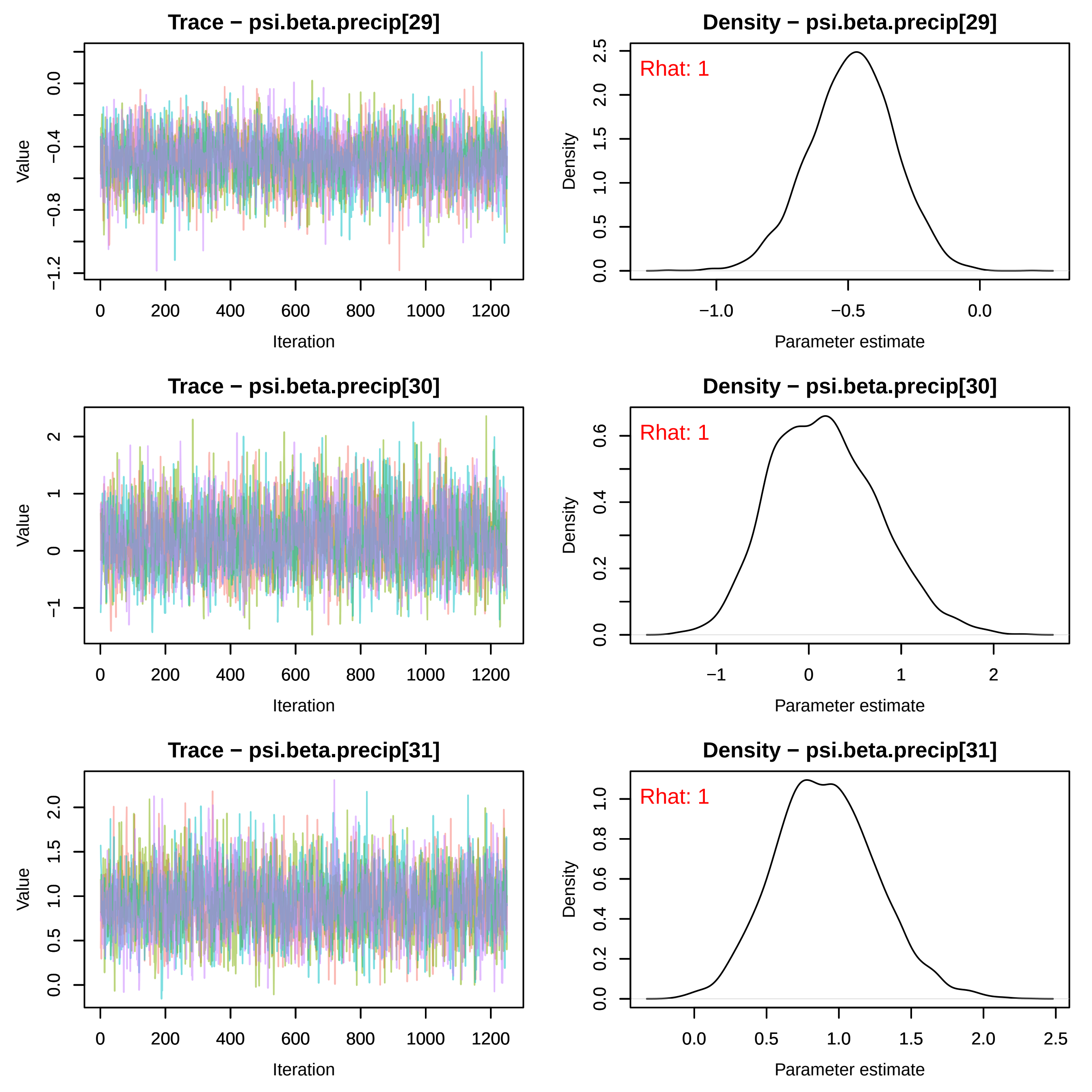
Trace − psi.beta.precip[29]
Density − psi.beta.precip[29]
Rhat: 1
Trace − psi.beta.precip[30]
Density − psi.beta.precip[30]
Rhat: 1
Trace − psi.beta.precip[31]
Density − psi.beta.precip[31]
Rhat: 1
Value
Iteration
Density
Parameter estimate

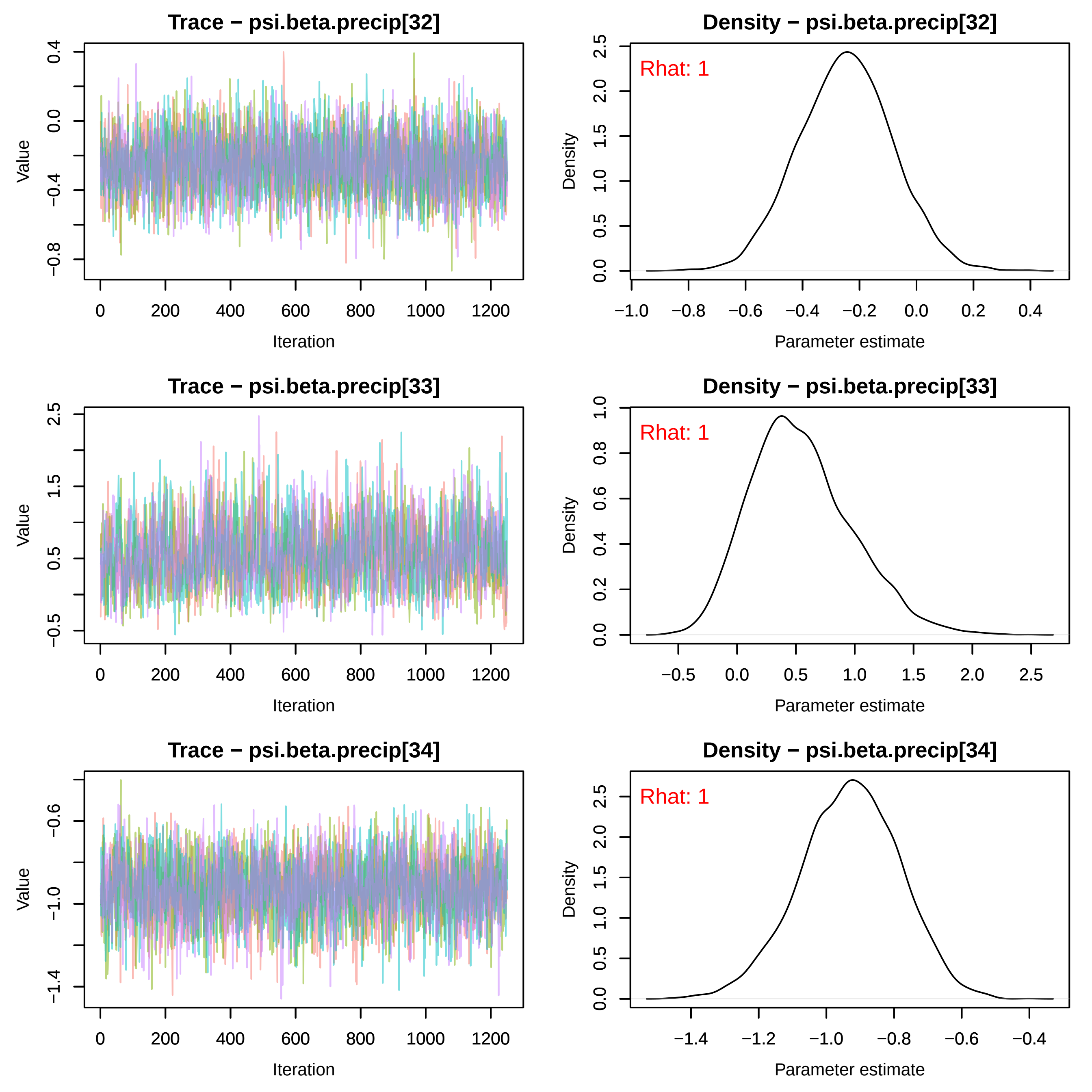

Trace − psi.beta.precip[32]
Value
Iteration
Density − psi.beta.precip[32]
Rhat: 1
Density
Parameter estimate
Trace − psi.beta.precip[33]
Value
Iteration
Density − psi.beta.precip[33]
Rhat: 1
Density
Parameter estimate
Trace − psi.beta.precip[34]
Value
Iteration
Density − psi.beta.precip[34]
Rhat: 1
Density
Parameter estimate

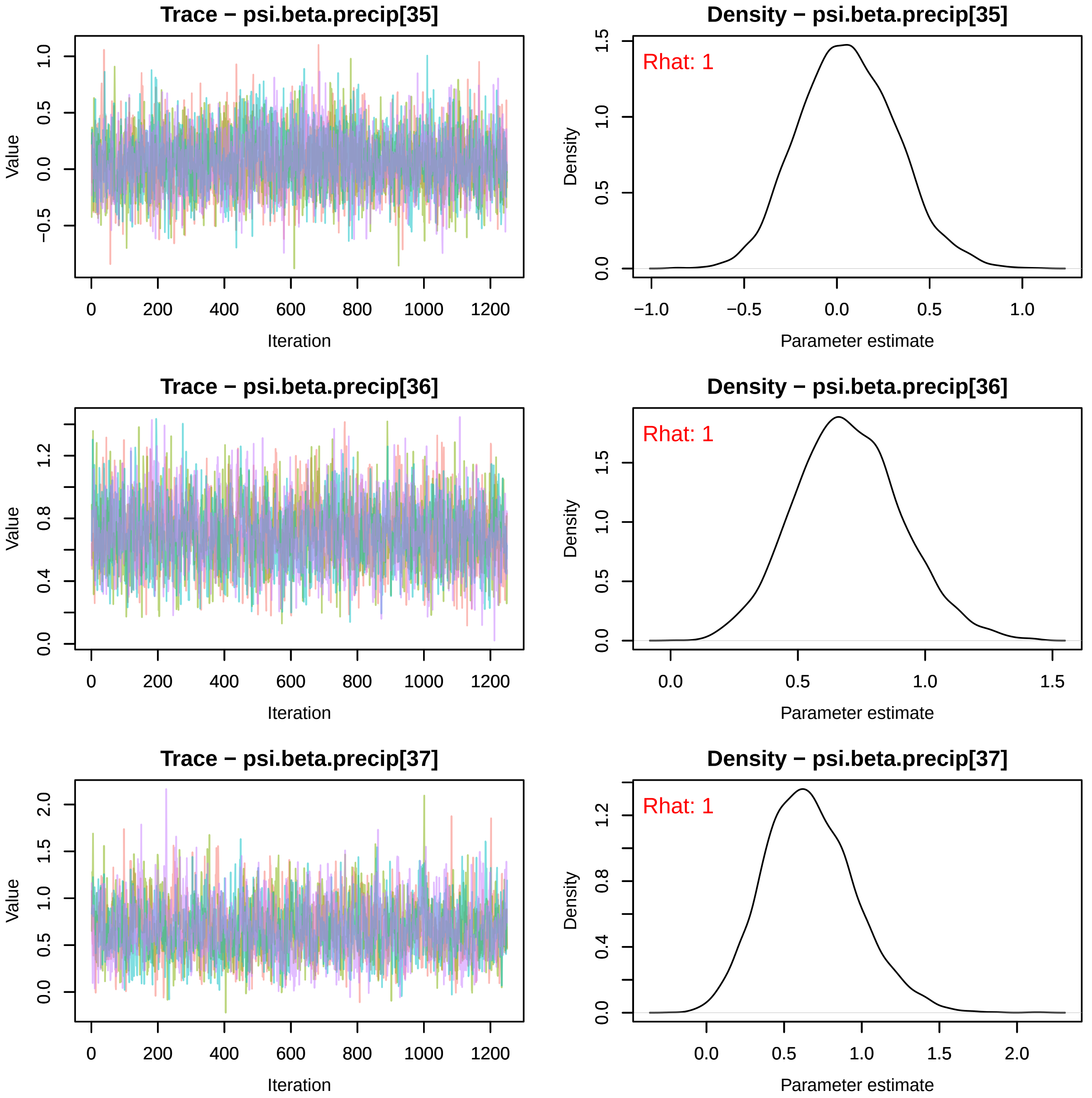

Trace − psi.beta.precip[35]
Value
−0.5
0.0
0.5
1.0
0
200
400
600
800
1000
1200
Iteration
Density − psi.beta.precip[35]
Rhat: 1
Density
0.0
0.5
1.0
1.5
−1.0
−0.5
0.0
0.5
1.0
Parameter estimate
Trace − psi.beta.precip[36]
Value
0.0
0.4
0.8
1.2
0
200
400
600
800
1000
1200
Iteration
Density − psi.beta.precip[36]
Rhat: 1
Density
0.0
0.5
1.0
1.5
0.0
0.5
1.0
1.5
Parameter estimate
Trace − psi.beta.precip[37]
Value
0.0
0.5
1.0
1.5
2.0
0
200
400
600
800
1000
1200
Iteration
Density − psi.beta.precip[37]
Rhat: 1
Density
0.0
0.4
0.8
1.2
0.0
0.5
1.0
1.5
2.0
Parameter estimate

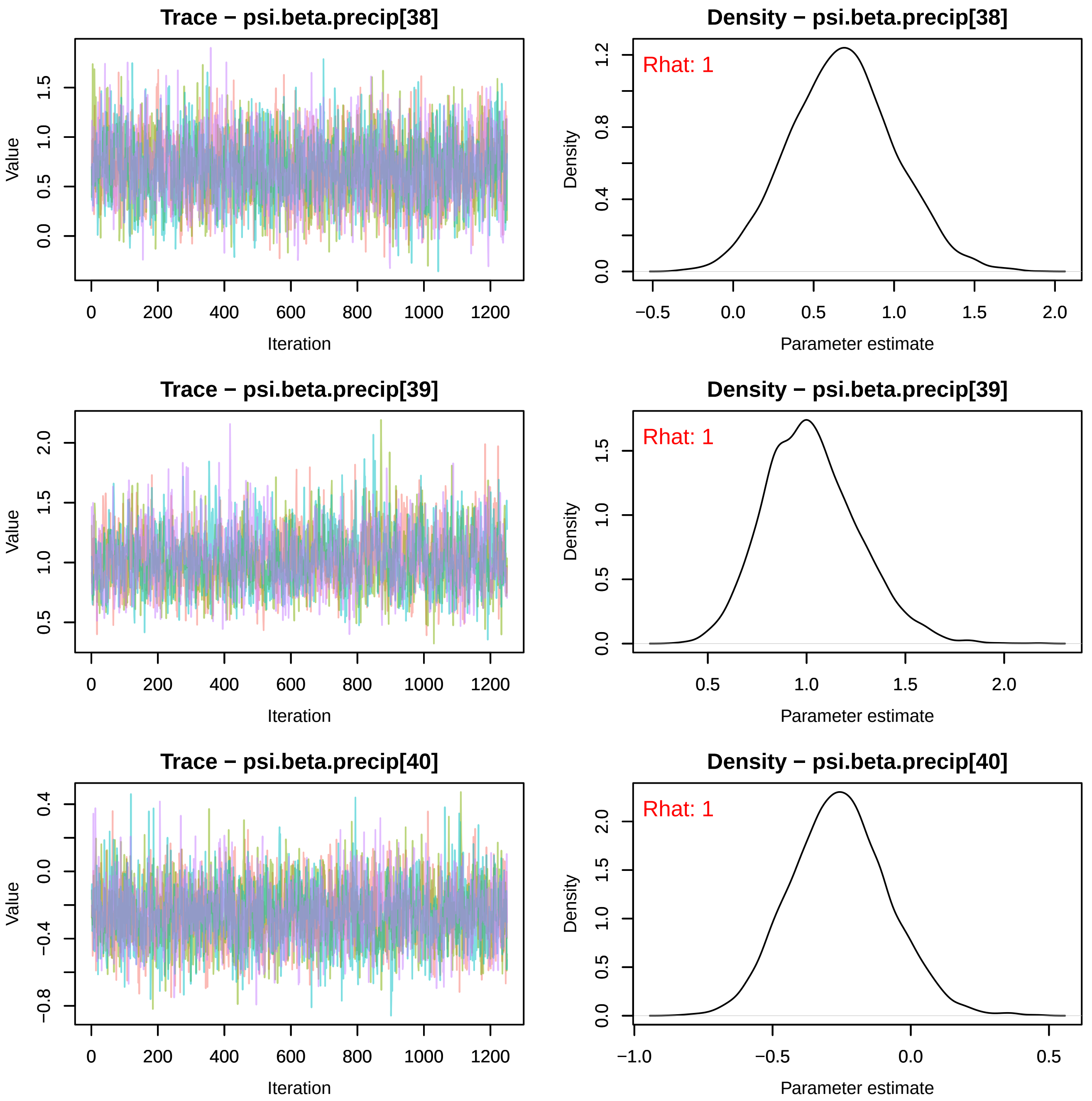
Trace − psi.beta.precip[38]
Value
Iteration
Density − psi.beta.precip[38]
Rhat: 1
Density
Parameter estimate
Trace − psi.beta.precip[39]
Value
Iteration
Density − psi.beta.precip[39]
Rhat: 1
Density
Parameter estimate
Trace − psi.beta.precip[40]
Value
Iteration
Density − psi.beta.precip[40]
Rhat: 1
Density
Parameter estimate

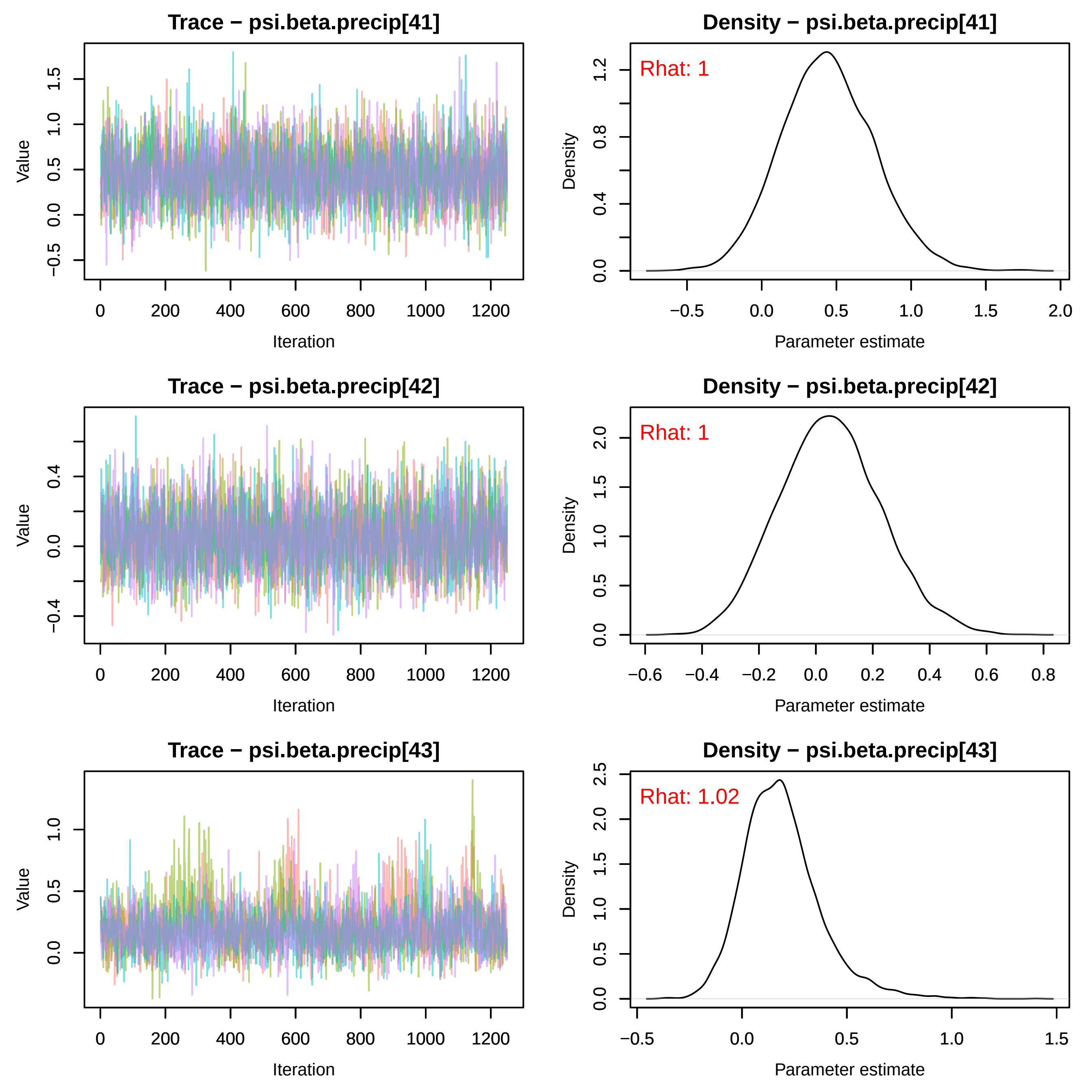
Trace − psi.beta.precip[41]
Value
Iteration
0 200 400 600 800 1000 1200
−0.5 0.0 0.5 1.0 1.5
Density − psi.beta.precip[41]
Rhat: 1
Density
Parameter estimate
−0.5 0.0 0.5 1.0 1.5 2.0
0.0 0.4 0.8 1.2
Trace − psi.beta.precip[42]
Value
Iteration
0 200 400 600 800 1000 1200
−0.4 0.0 0.4
Density − psi.beta.precip[42]
Rhat: 1
Density
Parameter estimate
−0.6 −0.4 −0.2 0.0 0.2 0.4 0.6 0.8
0.0 0.5 1.0 1.5 2.0
Trace − psi.beta.precip[43]
Value
Iteration
0 200 400 600 800 1000 1200
0.0 0.5 1.0
Density − psi.beta.precip[43]
Rhat: 1.02
Density
Parameter estimate
−0.5 0.0 0.5 1.0 1.5
0.0 0.5 1.0 1.5 2.0 2.5

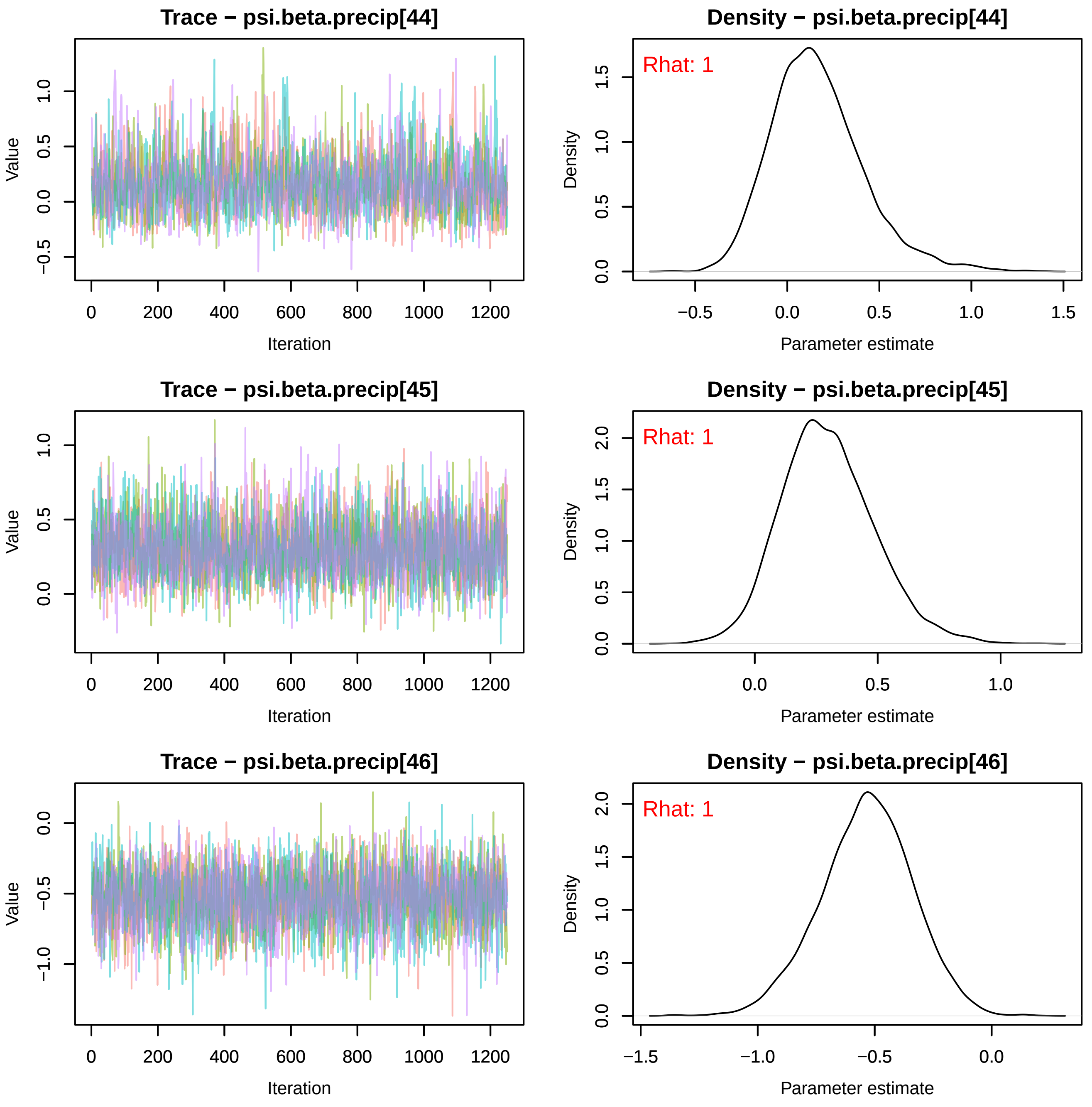
Trace − psi.beta.precip[44]
Value
Iteration
Density − psi.beta.precip[44]
Rhat: 1
Density
Parameter estimate
Trace − psi.beta.precip[45]
Value
Iteration
Density − psi.beta.precip[45]
Rhat: 1
Density
Parameter estimate
Trace − psi.beta.precip[46]
Value
Iteration
Density − psi.beta.precip[46]
Rhat: 1
Density
Parameter estimate

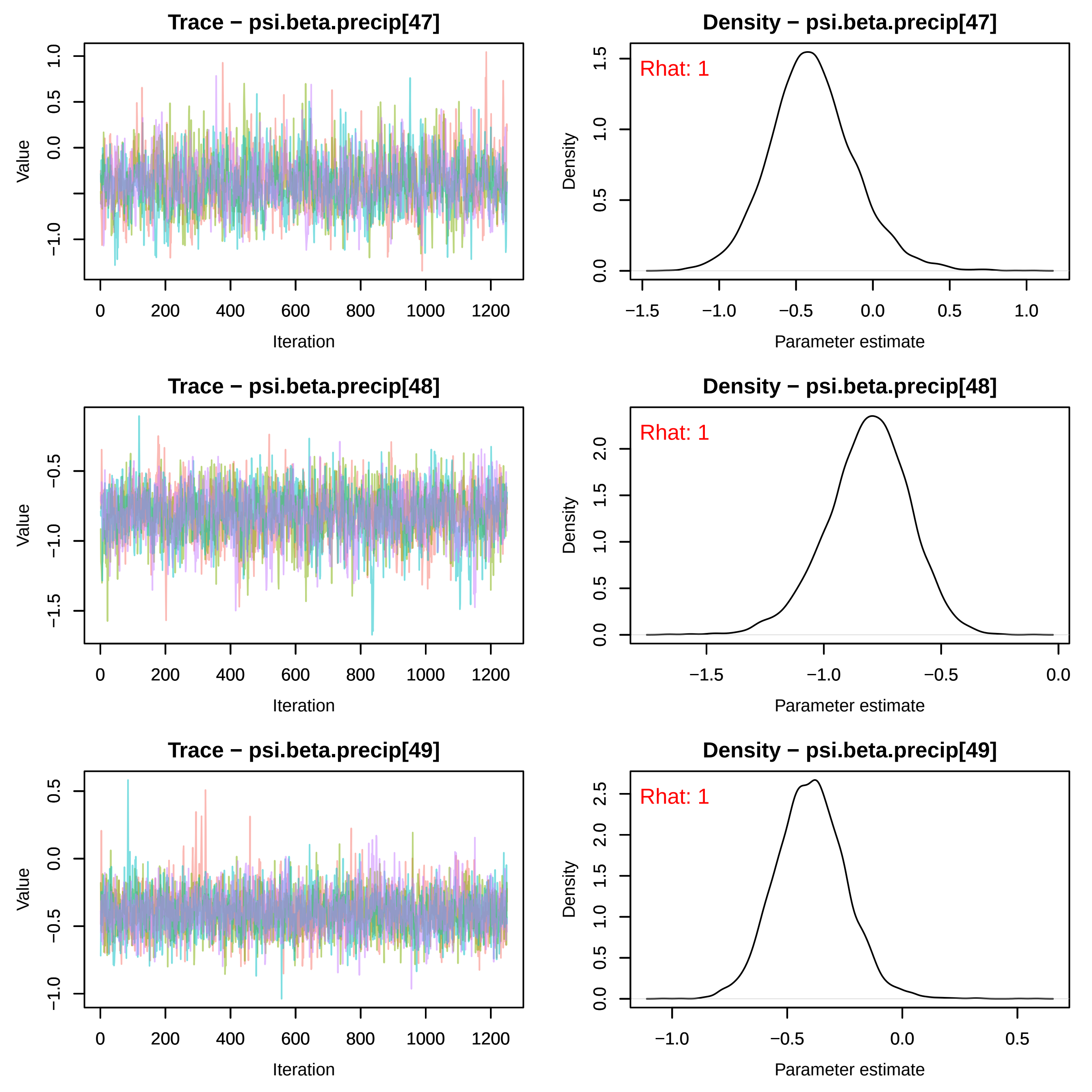
Trace − psi.beta.precip[47]
Density − psi.beta.precip[47]
Rhat: 1
Trace − psi.beta.precip[48]
Density − psi.beta.precip[48]
Rhat: 1
Trace − psi.beta.precip[49]
Density − psi.beta.precip[49]
Rhat: 1
Value
Iteration
Density
Parameter estimate

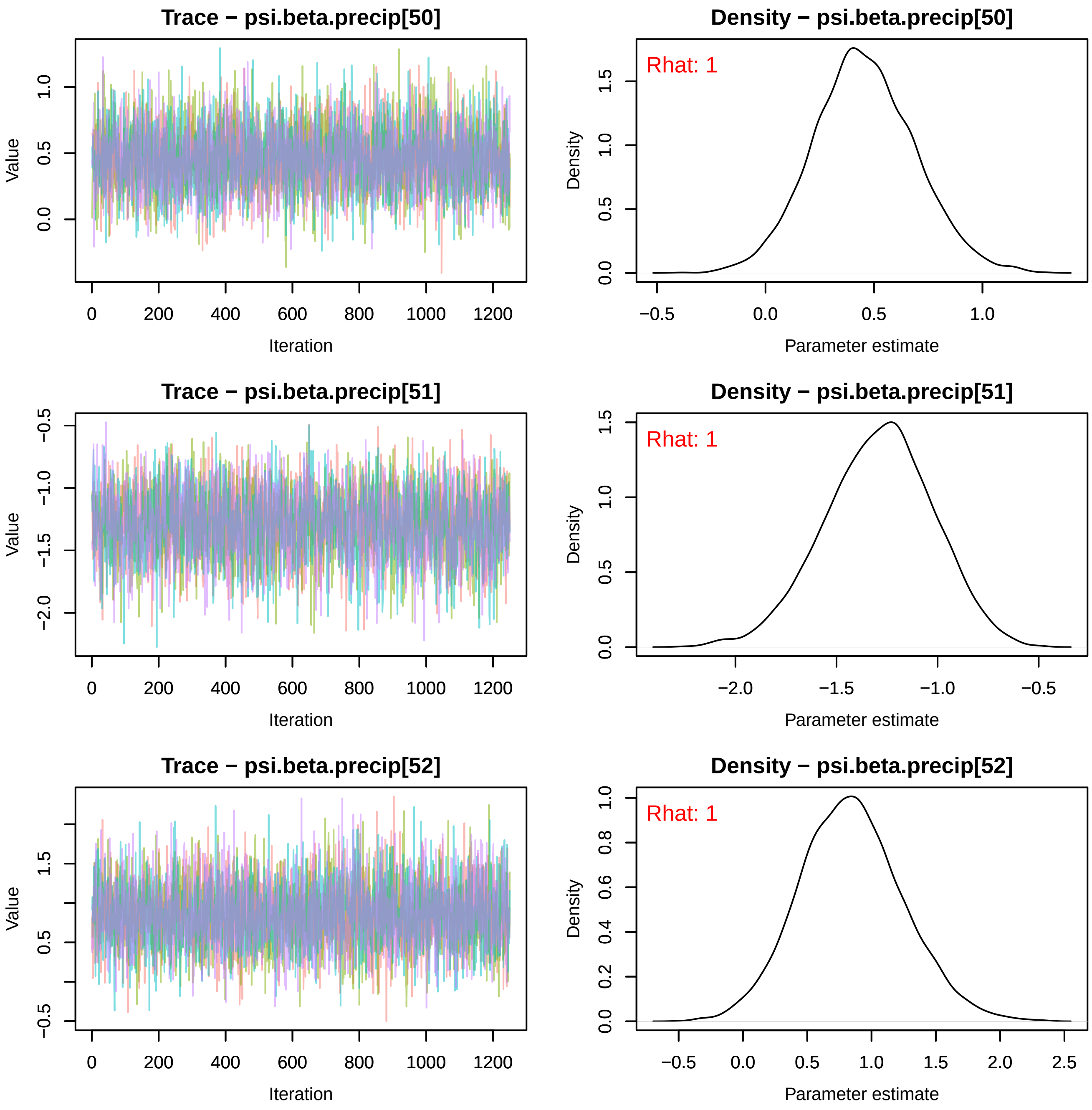
Trace − psi.beta.precip[50]
Density − psi.beta.precip[50]
Rhat: 1
Trace − psi.beta.precip[51]
Density − psi.beta.precip[51]
Rhat: 1
Trace − psi.beta.precip[52]
Density − psi.beta.precip[52]
Rhat: 1
Value
Iteration
Density
Parameter estimate

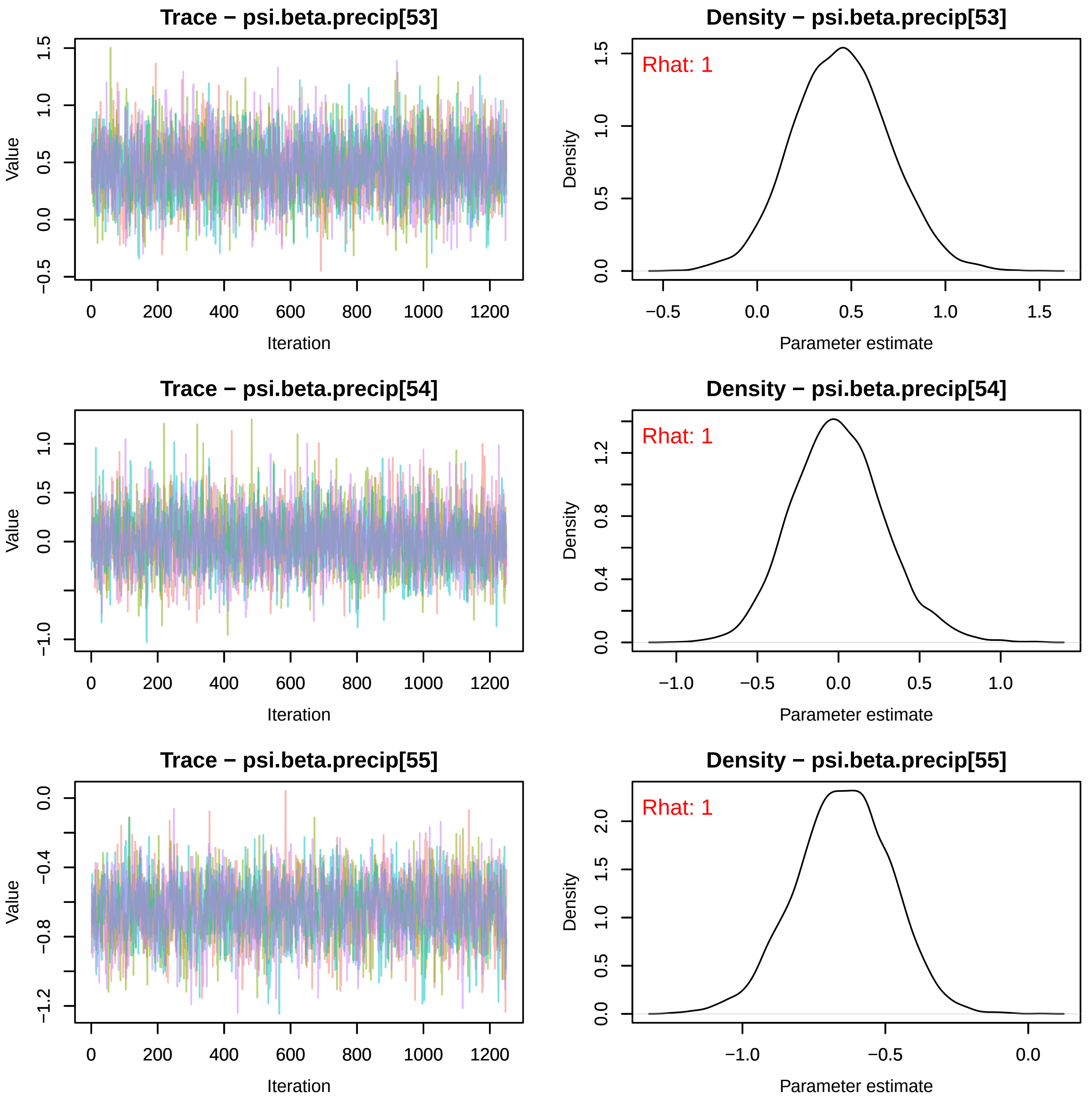
Trace − psi.beta.precip[53]
Value
Iteration
Density − psi.beta.precip[53]
Rhat: 1
Density
Parameter estimate
Trace − psi.beta.precip[54]
Value
Iteration
Density − psi.beta.precip[54]
Rhat: 1
Density
Parameter estimate
Trace − psi.beta.precip[55]
Value
Iteration
Density − psi.beta.precip[55]
Rhat: 1
Density
Parameter estimate

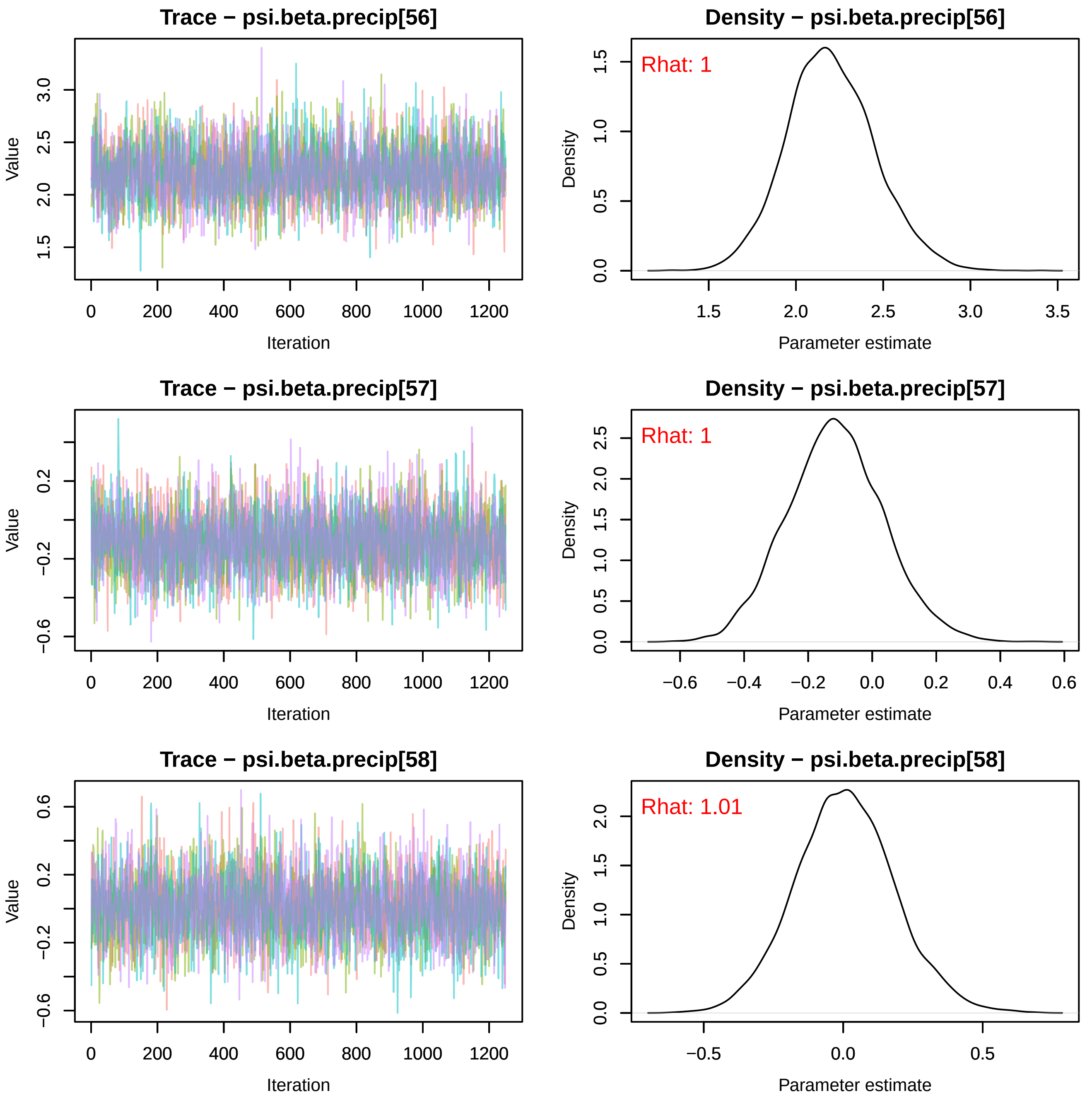
Trace − psi.beta.precip[56]
Density − psi.beta.precip[56]
Rhat: 1
Trace − psi.beta.precip[57]
Density − psi.beta.precip[57]
Rhat: 1
Trace − psi.beta.precip[58]
Density − psi.beta.precip[58]
Rhat: 1.01
Value
Iteration
Density
Parameter estimate

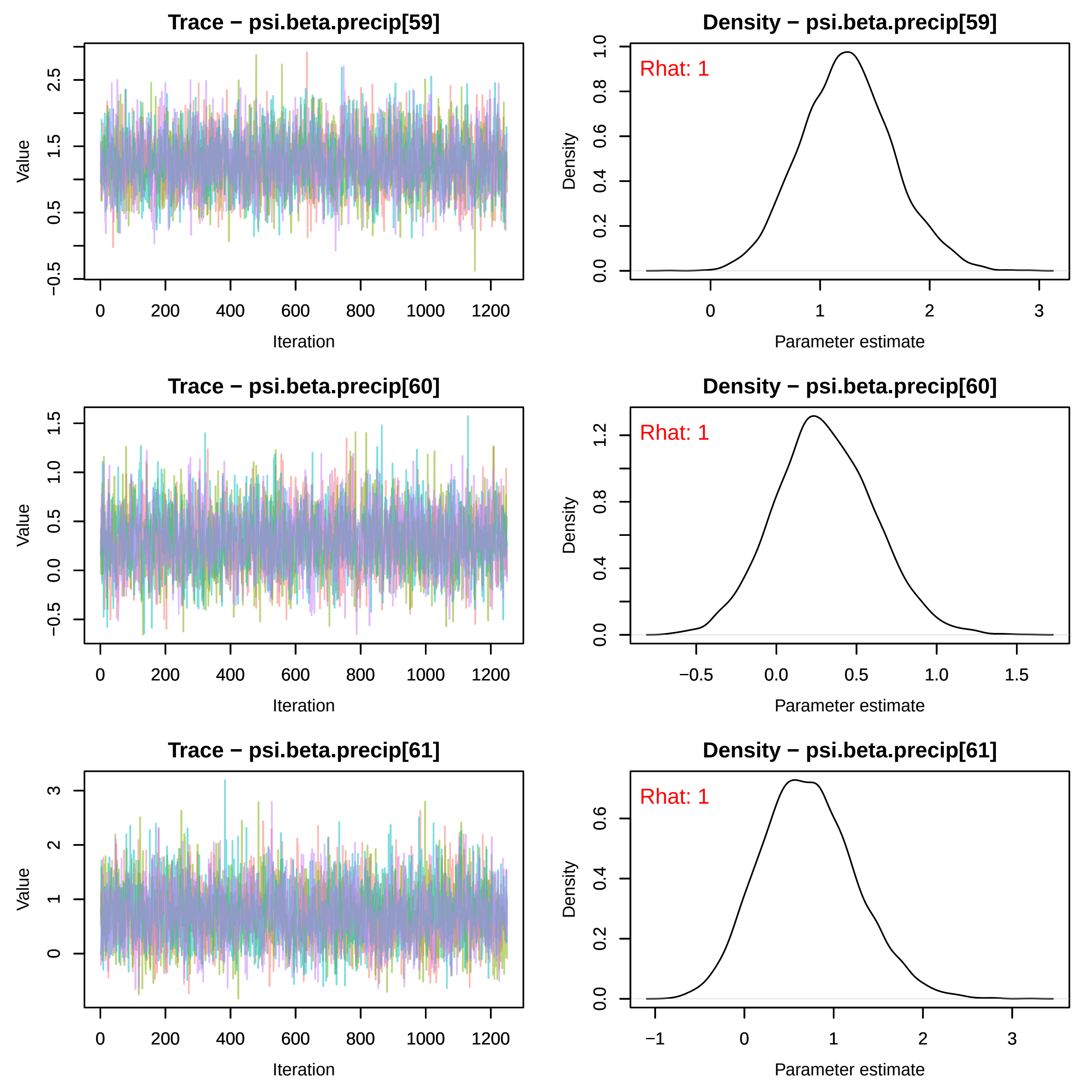
Trace − psi.beta.precip[59]
Value
Iteration
Density − psi.beta.precip[59]
Rhat: 1
Density
Parameter estimate
Trace − psi.beta.precip[60]
Value
Iteration
Density − psi.beta.precip[60]
Rhat: 1
Density
Parameter estimate
Trace − psi.beta.precip[61]
Value
Iteration
Density − psi.beta.precip[61]
Rhat: 1
Density
Parameter estimate

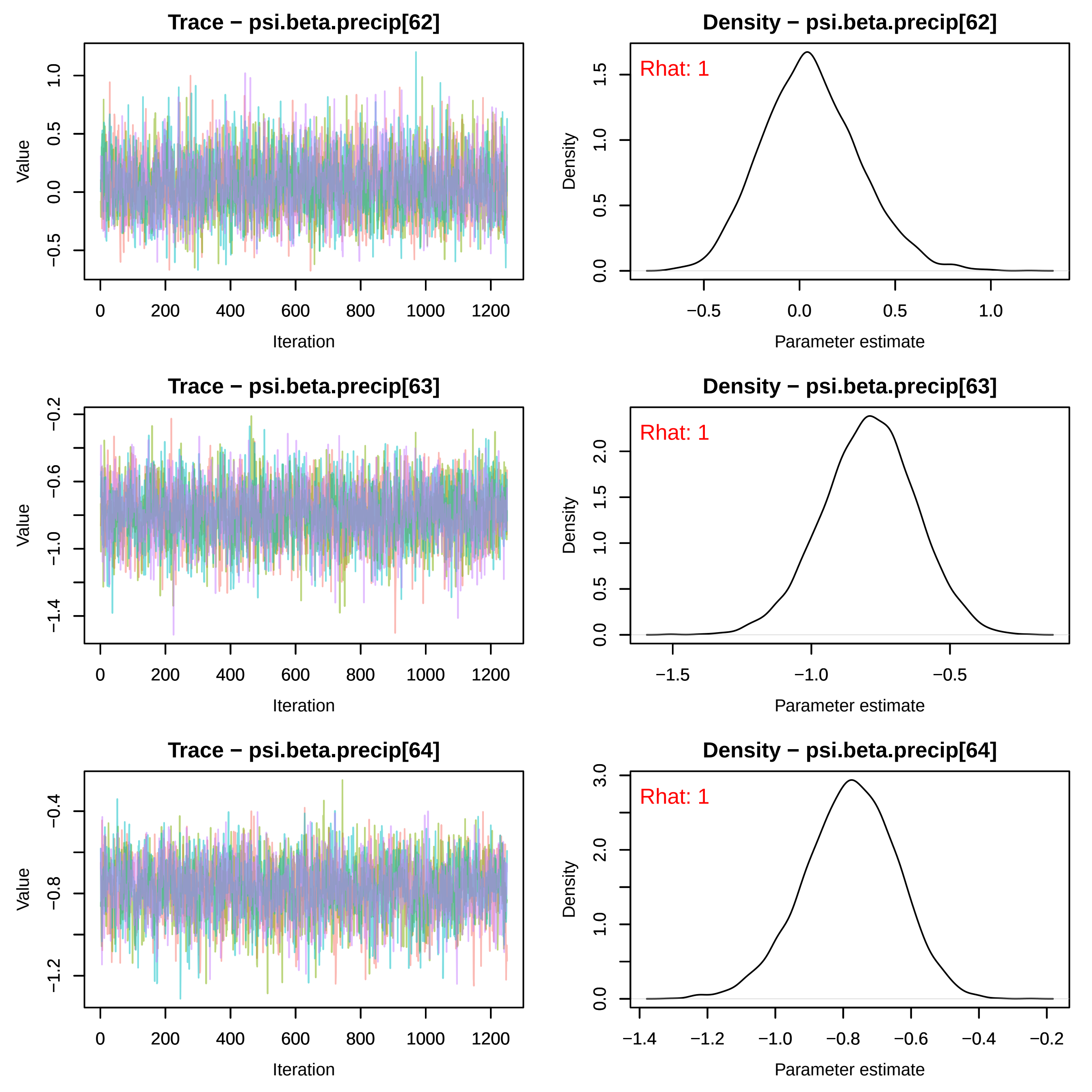
Trace − psi.beta.precip[62]
Value
Iteration
Density − psi.beta.precip[62]
Rhat: 1
Density
Parameter estimate
Trace − psi.beta.precip[63]
Value
Iteration
Density − psi.beta.precip[63]
Rhat: 1
Density
Parameter estimate
Trace − psi.beta.precip[64]
Value
Iteration
Density − psi.beta.precip[64]
Rhat: 1
Density
Parameter estimate

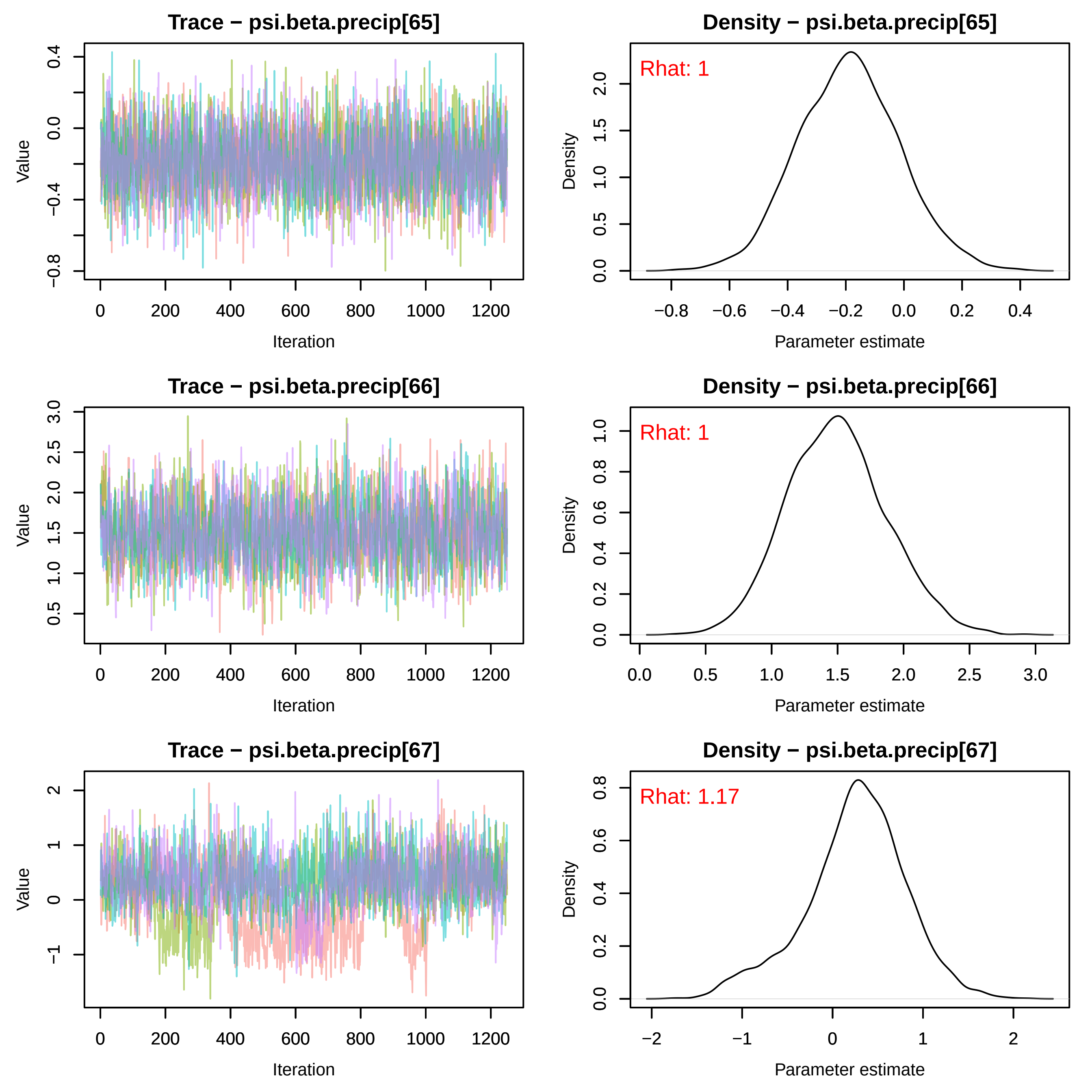
Trace − psi.beta.precip[65]
Density − psi.beta.precip[65]
Rhat: 1
Trace − psi.beta.precip[66]
Density − psi.beta.precip[66]
Rhat: 1
Trace − psi.beta.precip[67]
Density − psi.beta.precip[67]
Rhat: 1.17
Value
Iteration
Density
Parameter estimate

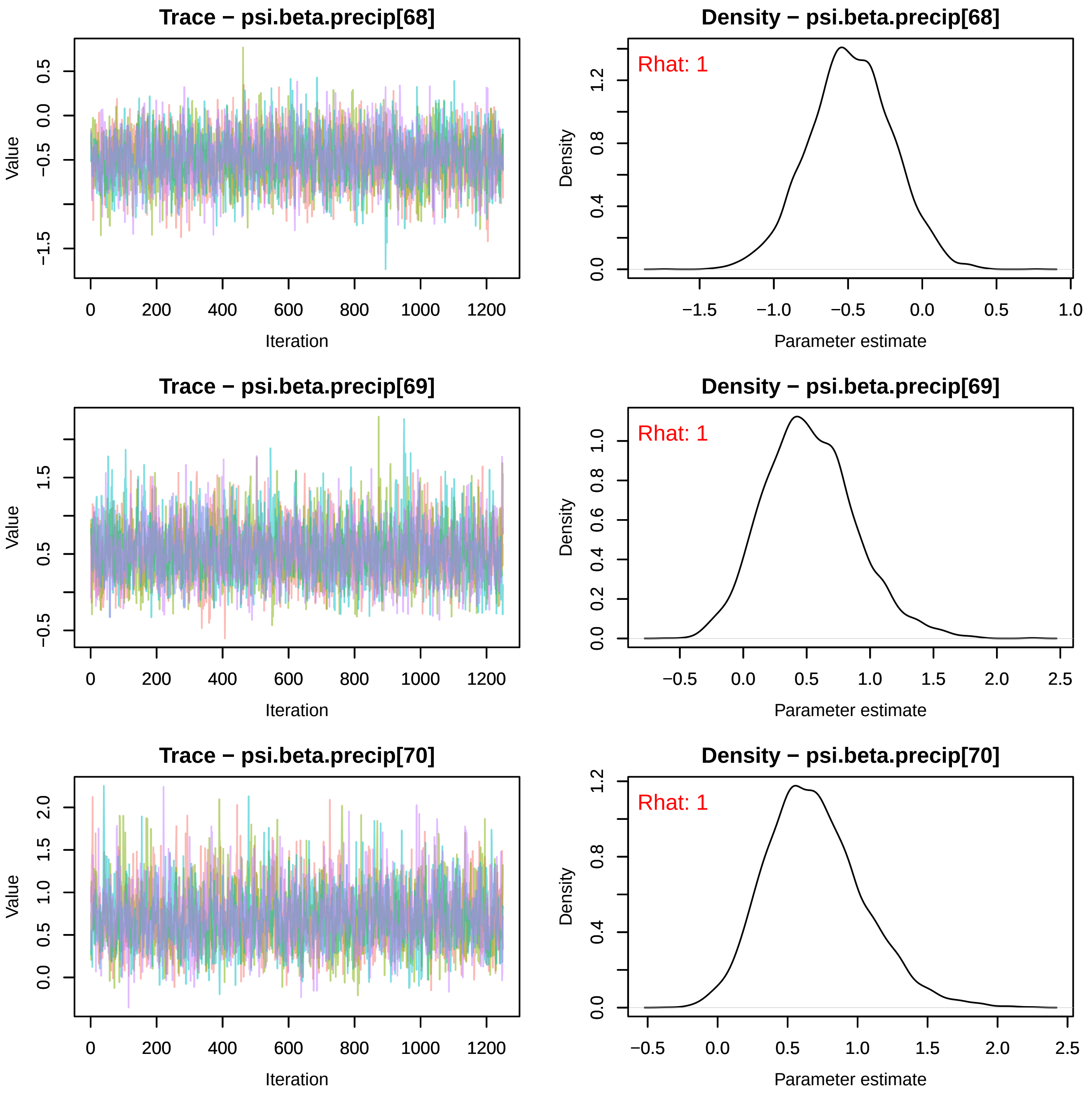
Trace − psi.beta.precip[68]
Density − psi.beta.precip[68]
Rhat: 1
Trace − psi.beta.precip[69]
Density − psi.beta.precip[69]
Rhat: 1
Trace − psi.beta.precip[70]
Density − psi.beta.precip[70]
Rhat: 1
Value
Iteration
Density
Parameter estimate

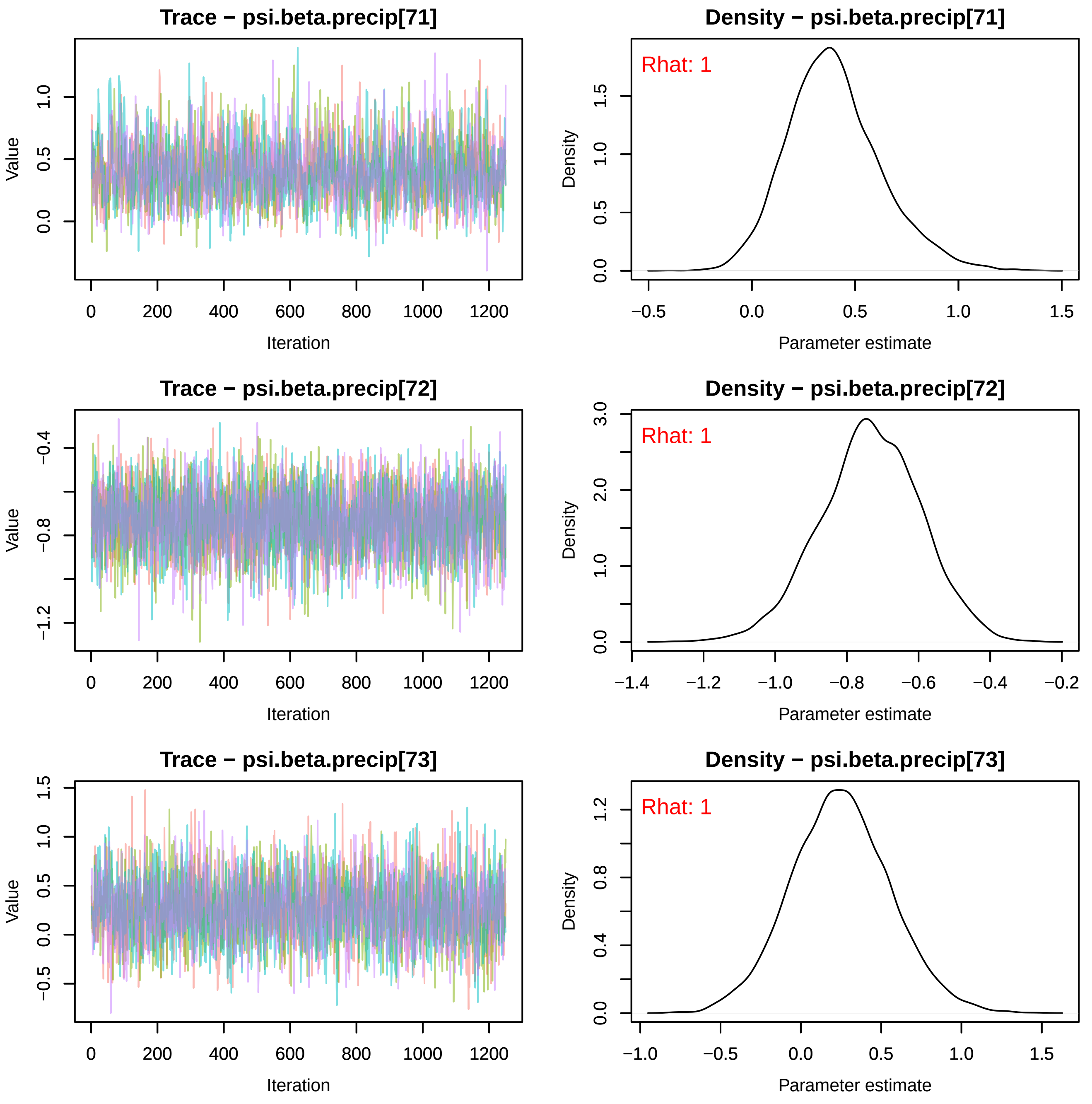
Trace − psi.beta.precip[71]
Value
Iteration
Density − psi.beta.precip[71]
Rhat: 1
Density
Parameter estimate
Trace − psi.beta.precip[72]
Value
Iteration
Density − psi.beta.precip[72]
Rhat: 1
Density
Parameter estimate
Trace − psi.beta.precip[73]
Value
Iteration
Density − psi.beta.precip[73]
Rhat: 1
Density
Parameter estimate

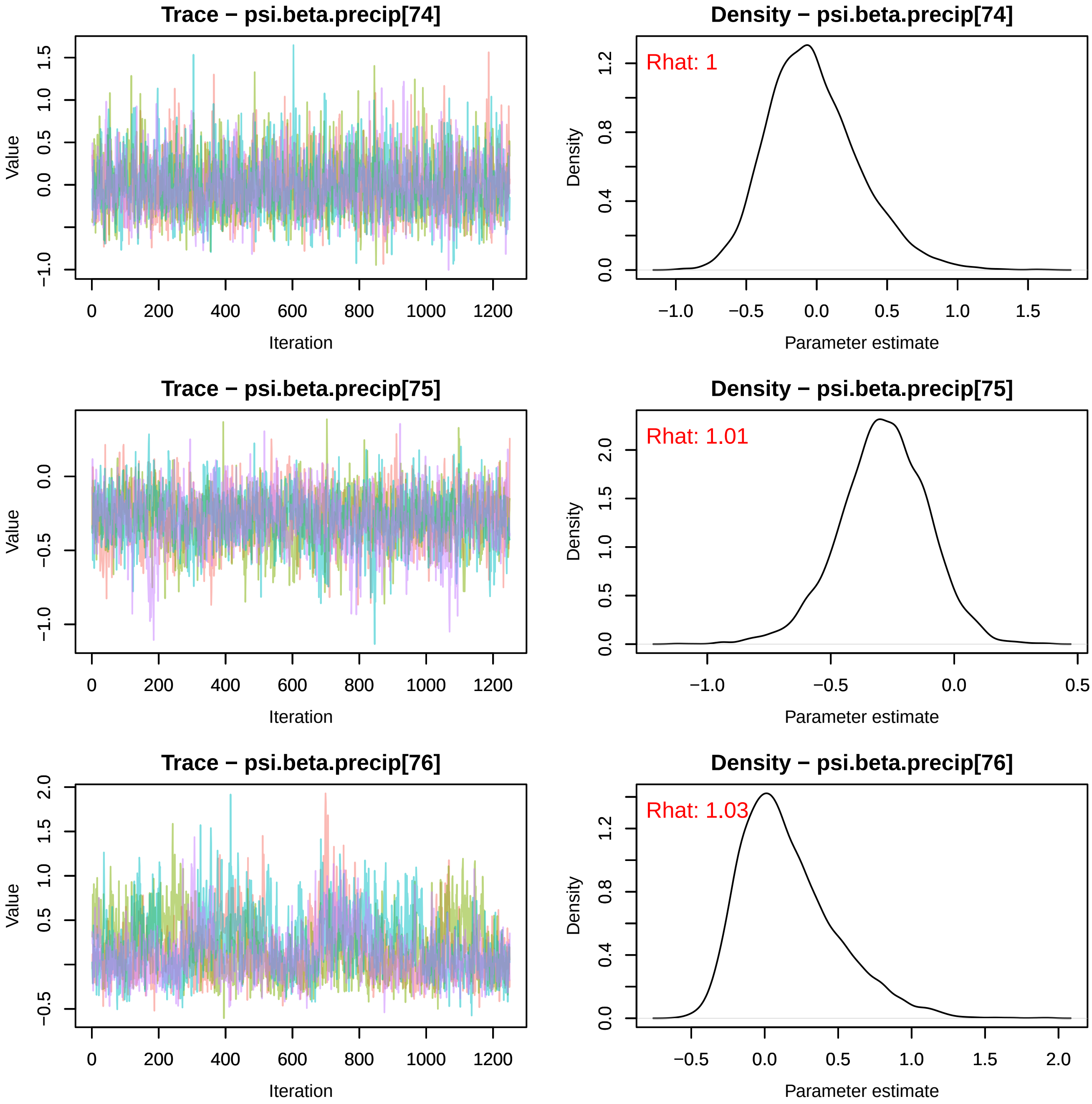

Trace − psi.beta.precip[74]
Density − psi.beta.precip[74]
Rhat: 1
Trace − psi.beta.precip[75]
Density − psi.beta.precip[75]
Rhat: 1.01
Trace − psi.beta.precip[76]
Density − psi.beta.precip[76]
Rhat: 1.03
Value
Iteration
Density
Parameter estimate

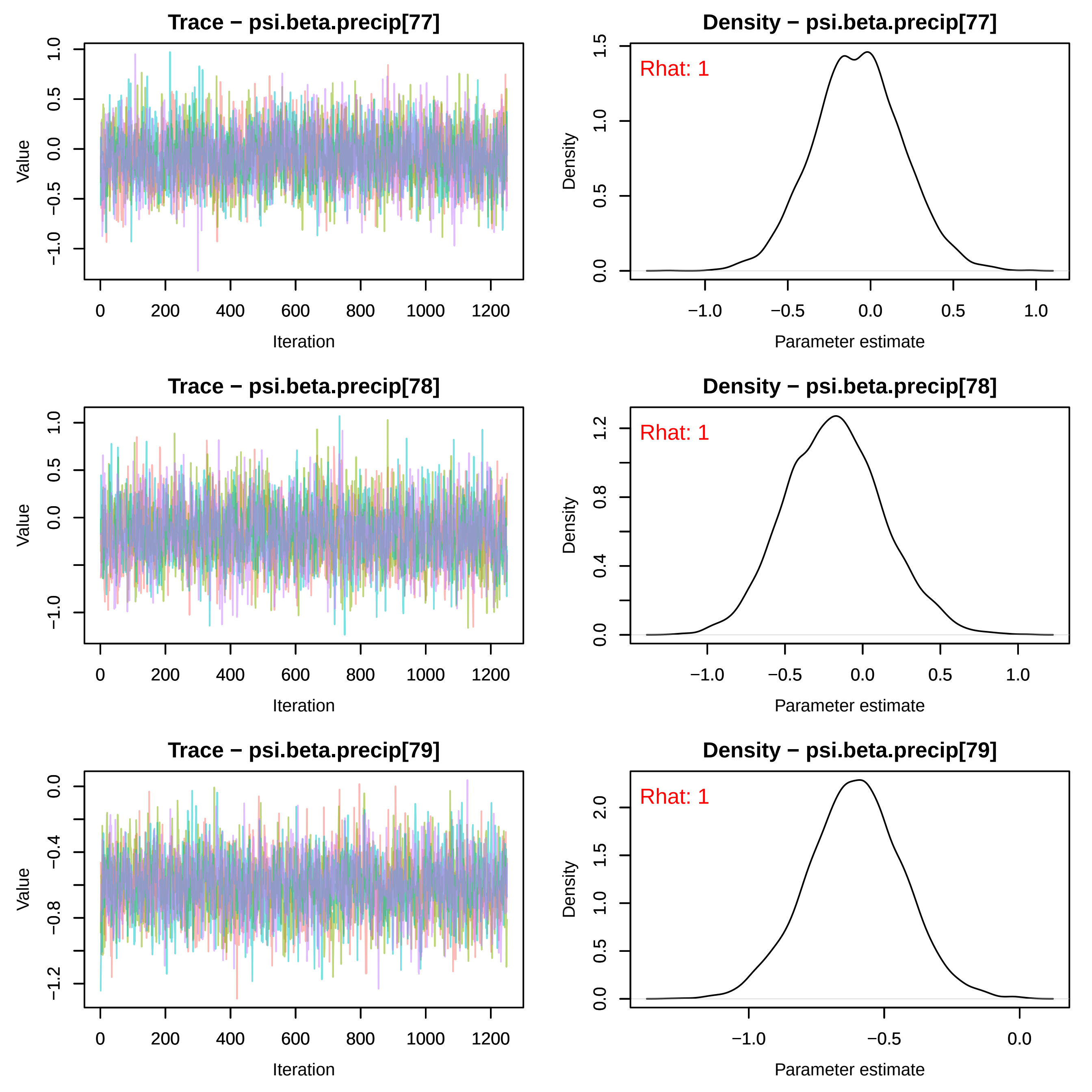
Trace − psi.beta.precip[77]
Value
Iteration
Density − psi.beta.precip[77]
Rhat: 1
Density
Parameter estimate
Trace − psi.beta.precip[78]
Value
Iteration
Density − psi.beta.precip[78]
Rhat: 1
Density
Parameter estimate
Trace − psi.beta.precip[79]
Value
Iteration
Density − psi.beta.precip[79]
Rhat: 1
Density
Parameter estimate

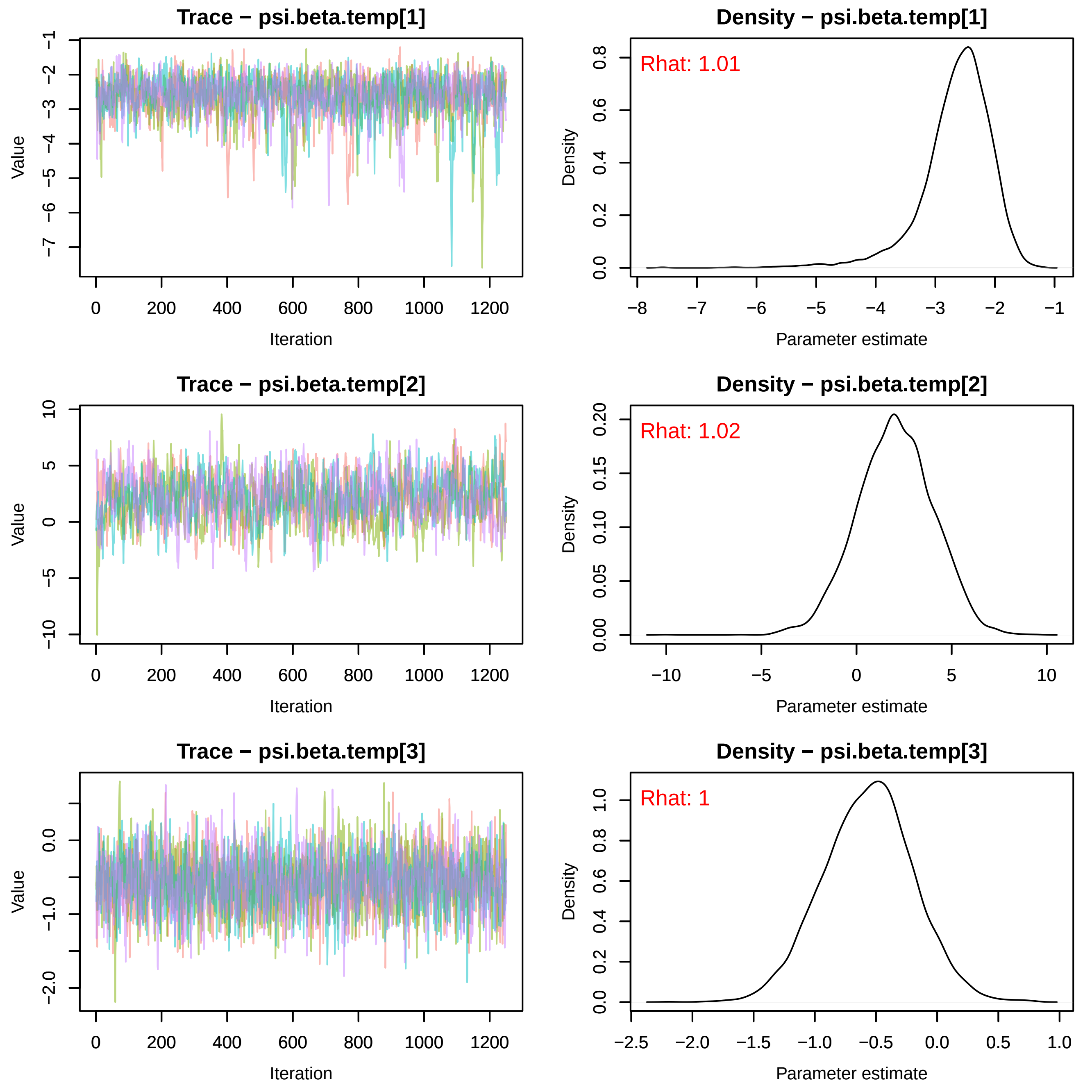
Trace − psi.beta.temp[1]
Value
Iteration
Density − psi.beta.temp[1]
Rhat: 1.01
Density
Parameter estimate
Trace − psi.beta.temp[2]
Value
Iteration
Density − psi.beta.temp[2]
Rhat: 1.02
Density
Parameter estimate
Trace − psi.beta.temp[3]
Value
Iteration
Density − psi.beta.temp[3]
Rhat: 1
Density
Parameter estimate

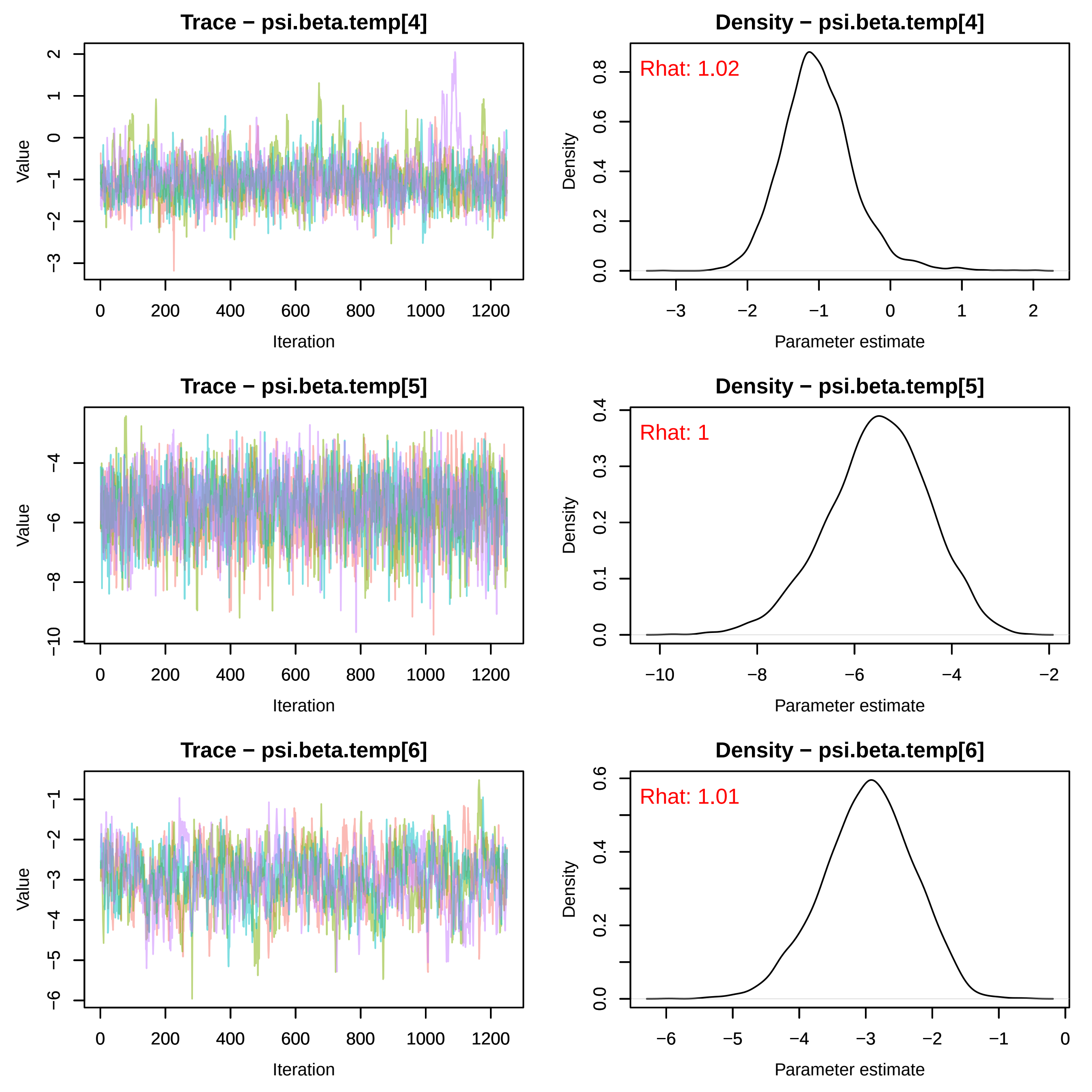

Trace − psi.beta.temp[4]
Value
Iteration
Density − psi.beta.temp[4]
Rhat: 1.02
Density
Parameter estimate
Trace − psi.beta.temp[5]
Value
Iteration
Density − psi.beta.temp[5]
Rhat: 1
Density
Parameter estimate
Trace − psi.beta.temp[6]
Value
Iteration
Density − psi.beta.temp[6]
Rhat: 1.01
Density
Parameter estimate

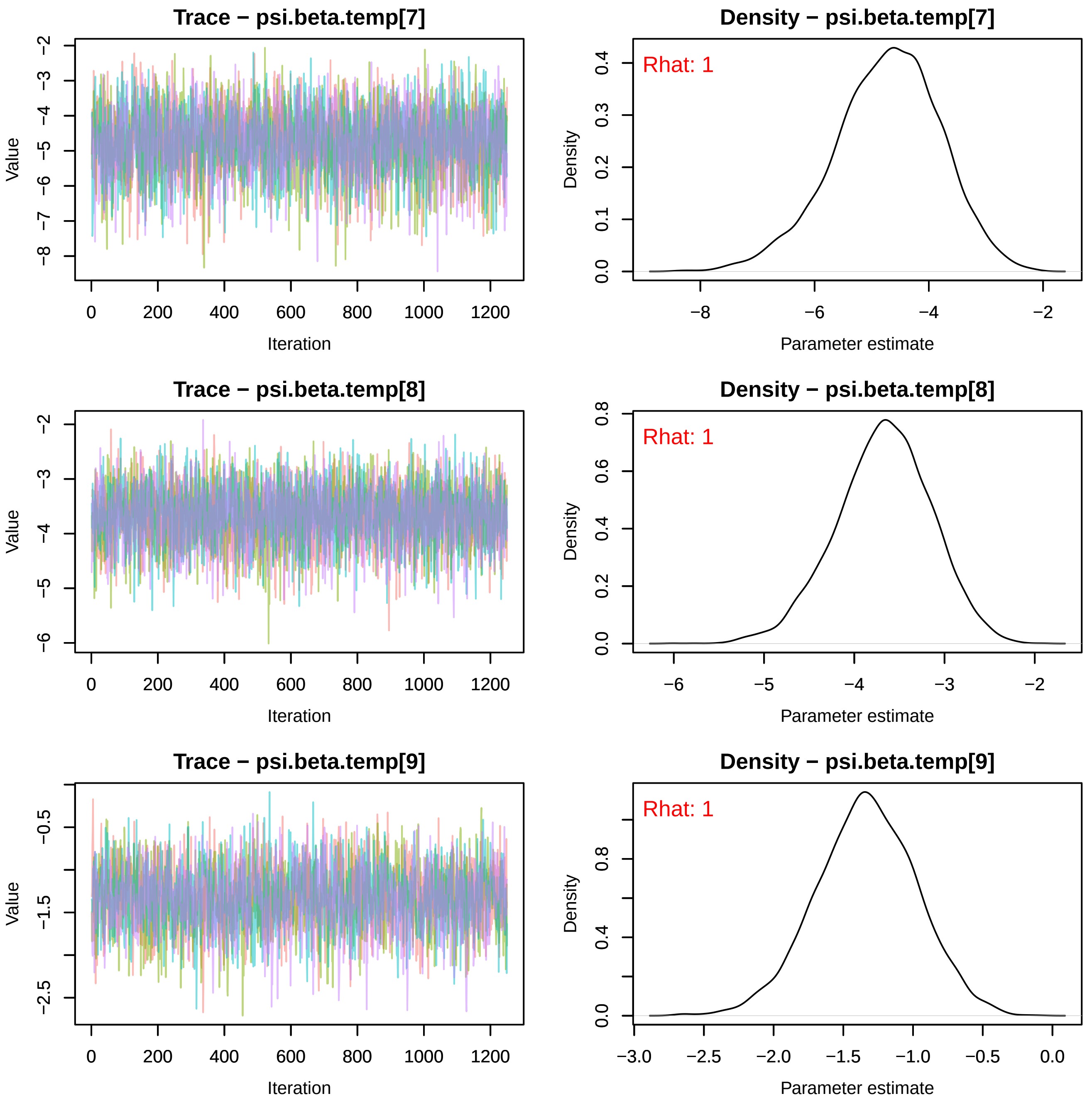

Trace − psi.beta.temp[7]
Density − psi.beta.temp[7]
Rhat: 1
Trace − psi.beta.temp[8]
Density − psi.beta.temp[8]
Rhat: 1
Trace − psi.beta.temp[9]
Density − psi.beta.temp[9]
Rhat: 1
Value
Iteration
Density
Parameter estimate

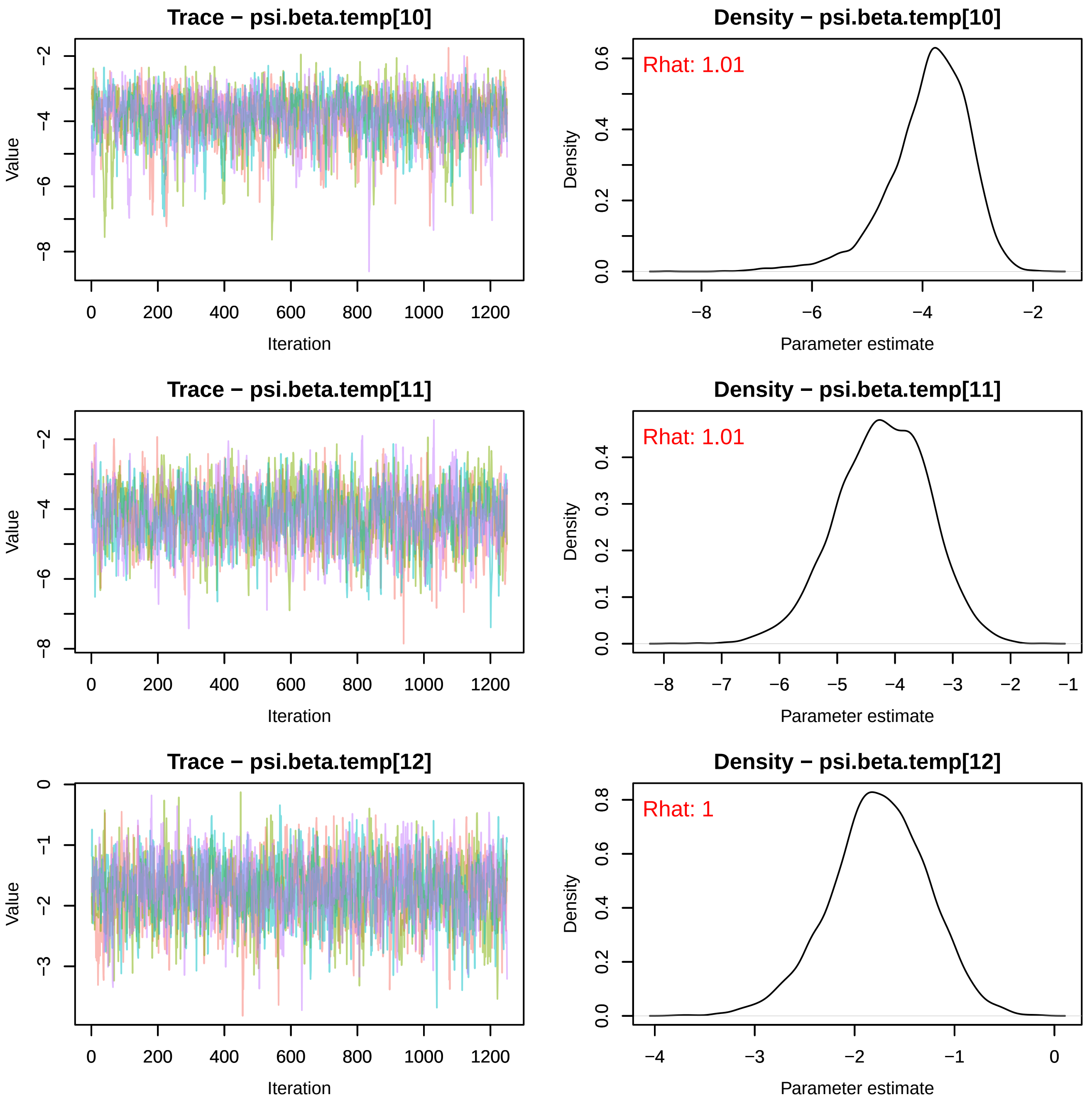

Trace − psi.beta.temp[10]
Value
Iteration
Density − psi.beta.temp[10]
Rhat: 1.01
Density
Parameter estimate
Trace − psi.beta.temp[11]
Value
Iteration
Density − psi.beta.temp[11]
Rhat: 1.01
Density
Parameter estimate
Trace − psi.beta.temp[12]
Value
Iteration
Density − psi.beta.temp[12]
Rhat: 1
Density
Parameter estimate

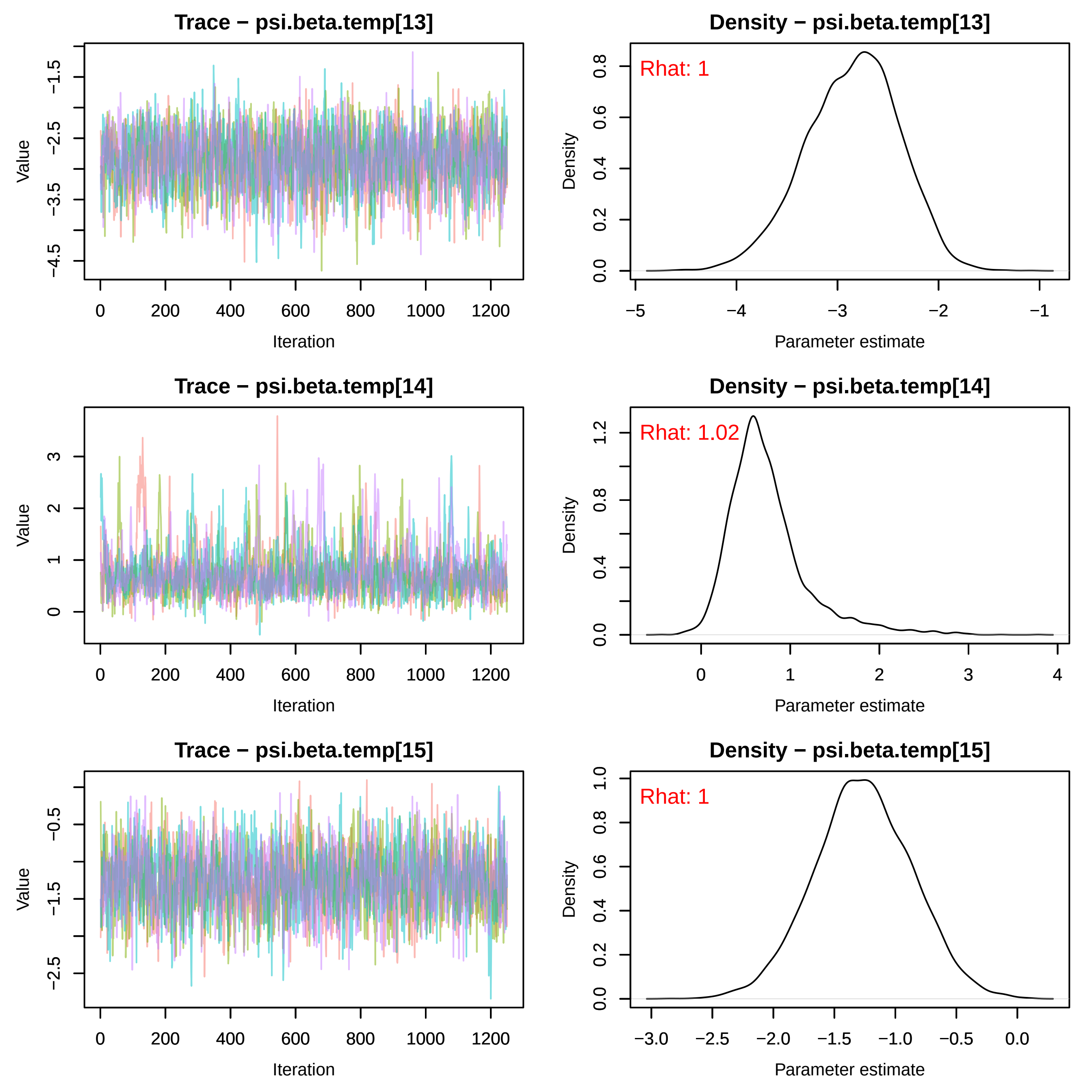
Trace − psi.beta.temp[13]
Value
Iteration
Density − psi.beta.temp[13]
Rhat: 1
Density
Parameter estimate
Trace − psi.beta.temp[14]
Value
Iteration
Density − psi.beta.temp[14]
Rhat: 1.02
Density
Parameter estimate
Trace − psi.beta.temp[15]
Value
Iteration
Density − psi.beta.temp[15]
Rhat: 1
Density
Parameter estimate

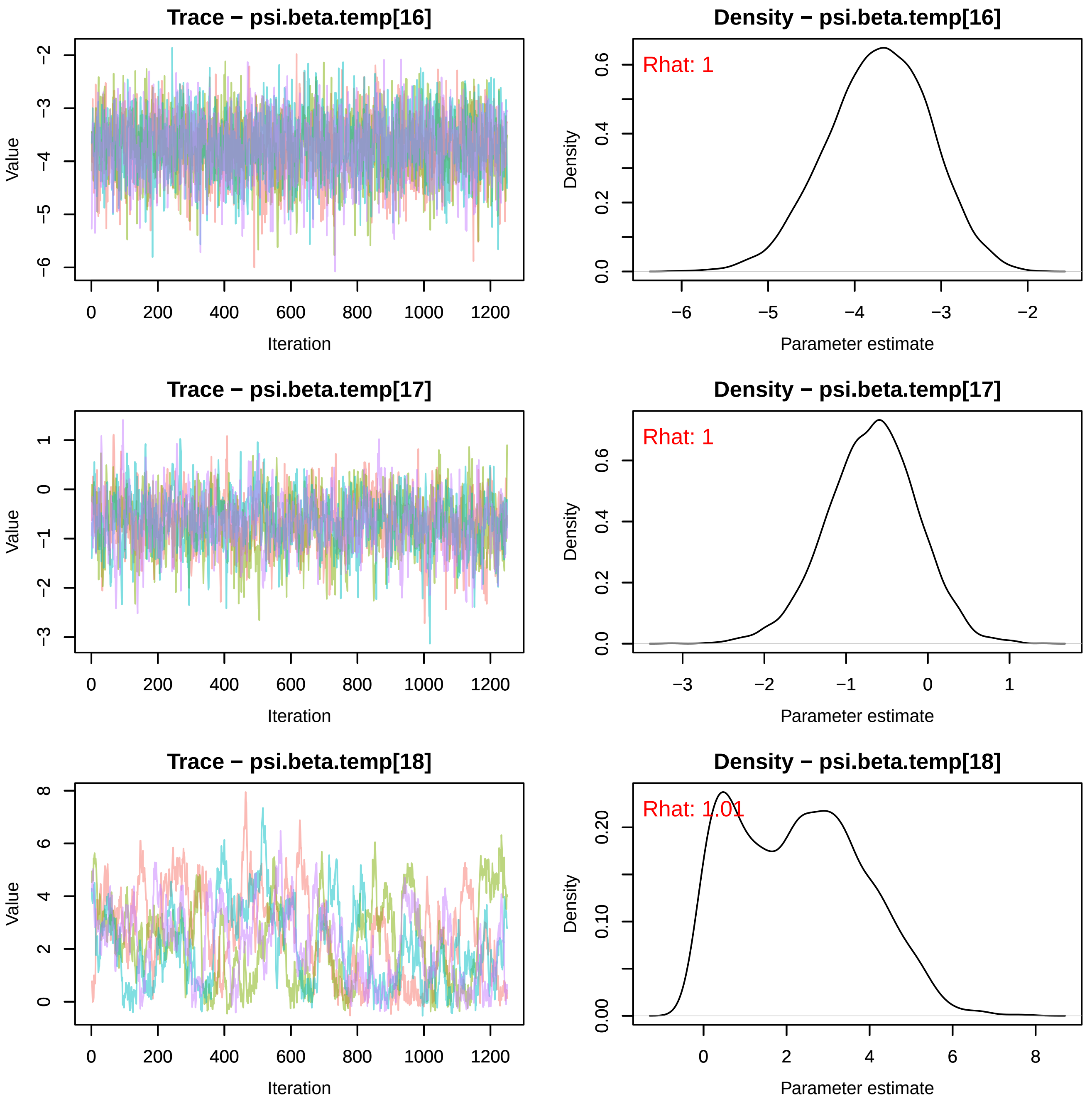
Trace − psi.beta.temp[16]
Density − psi.beta.temp[16]
Rhat: 1
Trace − psi.beta.temp[17]
Density − psi.beta.temp[17]
Rhat: 1
Trace − psi.beta.temp[18]
Density − psi.beta.temp[18]
Rhat: 1.01
Value
Iteration
Density
Parameter estimate

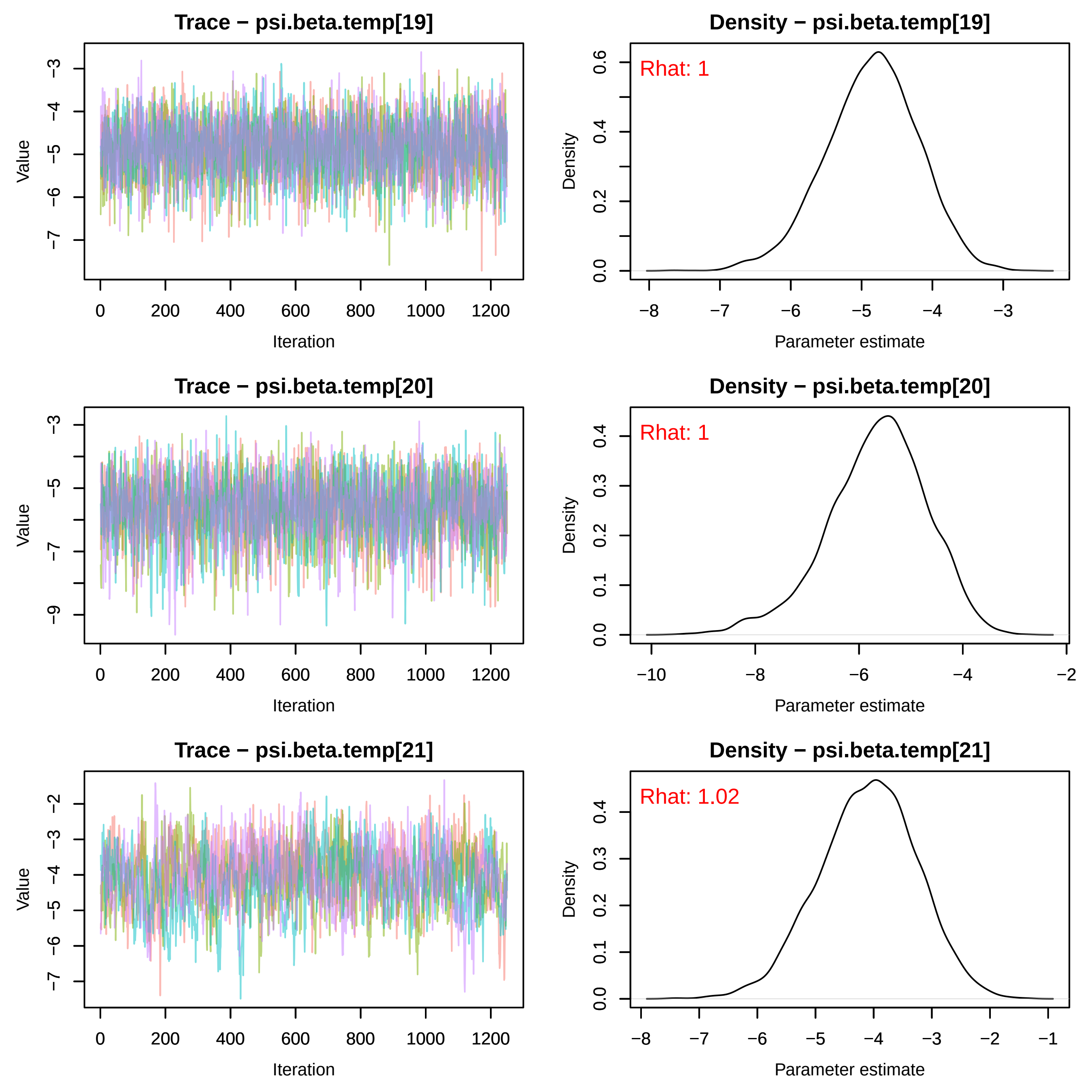
Trace − psi.beta.temp[19]
Value
Iteration
Density − psi.beta.temp[19]
Rhat: 1
Density
Parameter estimate
Trace − psi.beta.temp[20]
Value
Iteration
Density − psi.beta.temp[20]
Rhat: 1
Density
Parameter estimate
Trace − psi.beta.temp[21]
Value
Iteration
Density − psi.beta.temp[21]
Rhat: 1.02
Density
Parameter estimate

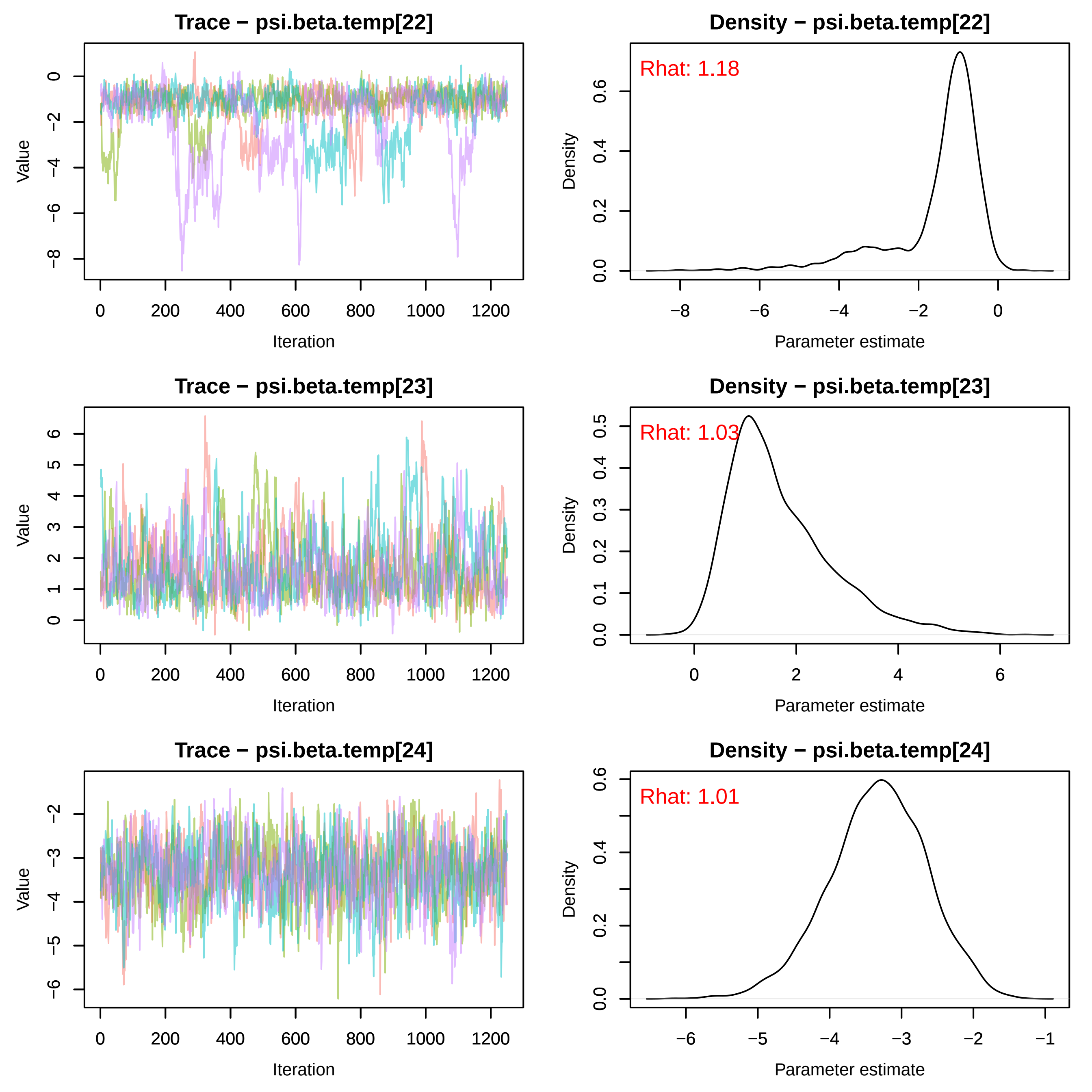
Trace − psi.beta.temp[22]
Density − psi.beta.temp[22]
Rhat: 1.18
Trace − psi.beta.temp[23]
Density − psi.beta.temp[23]
Rhat: 1.03
Trace − psi.beta.temp[24]
Density − psi.beta.temp[24]
Rhat: 1.01
Value
Iteration
Density
Parameter estimate

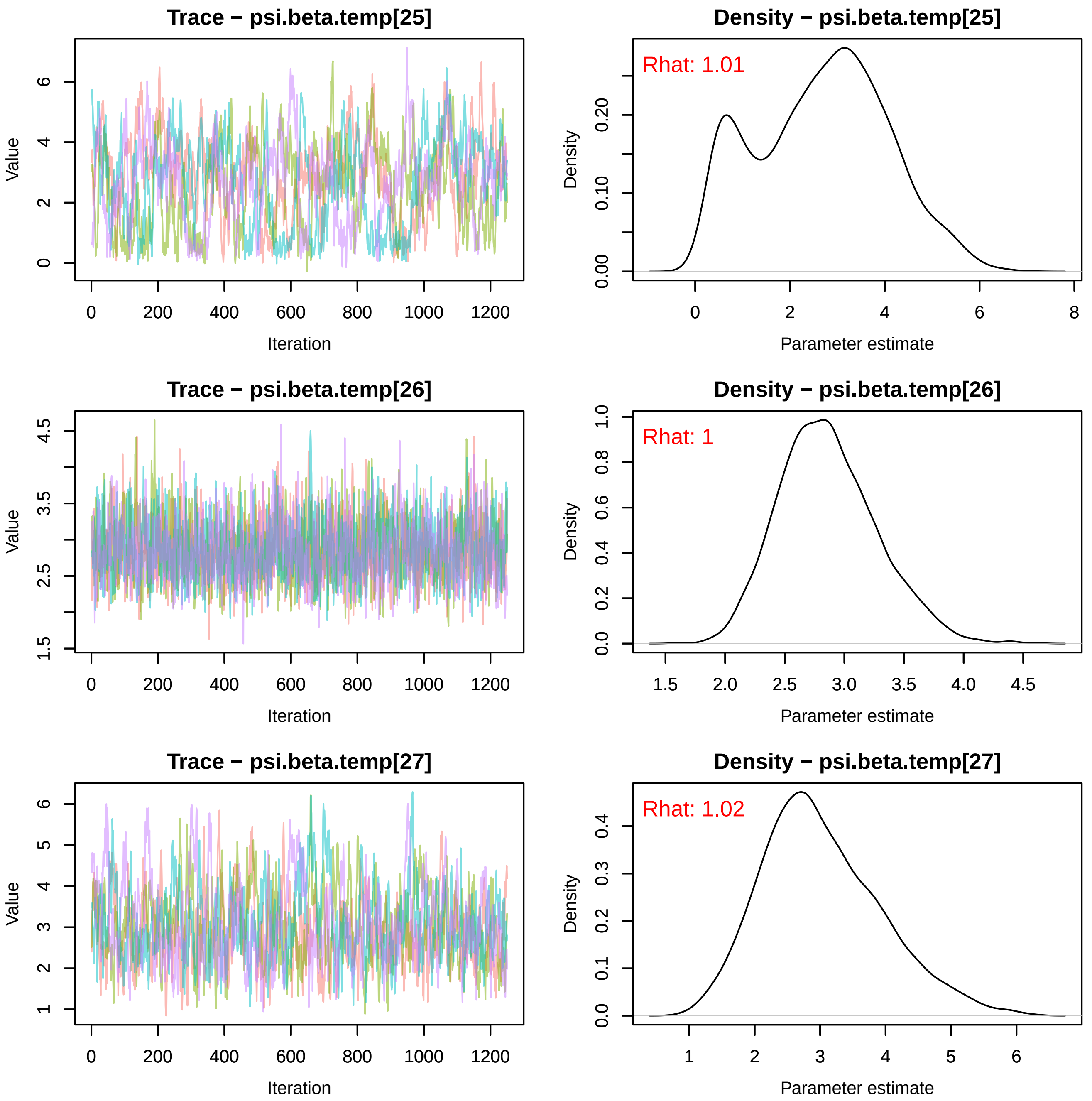
Trace − psi.beta.temp[25]
Density − psi.beta.temp[25]
Rhat: 1.01
Trace − psi.beta.temp[26]
Density − psi.beta.temp[26]
Rhat: 1
Trace − psi.beta.temp[27]
Density − psi.beta.temp[27]
Rhat: 1.02
Value
Iteration
Density
Parameter estimate

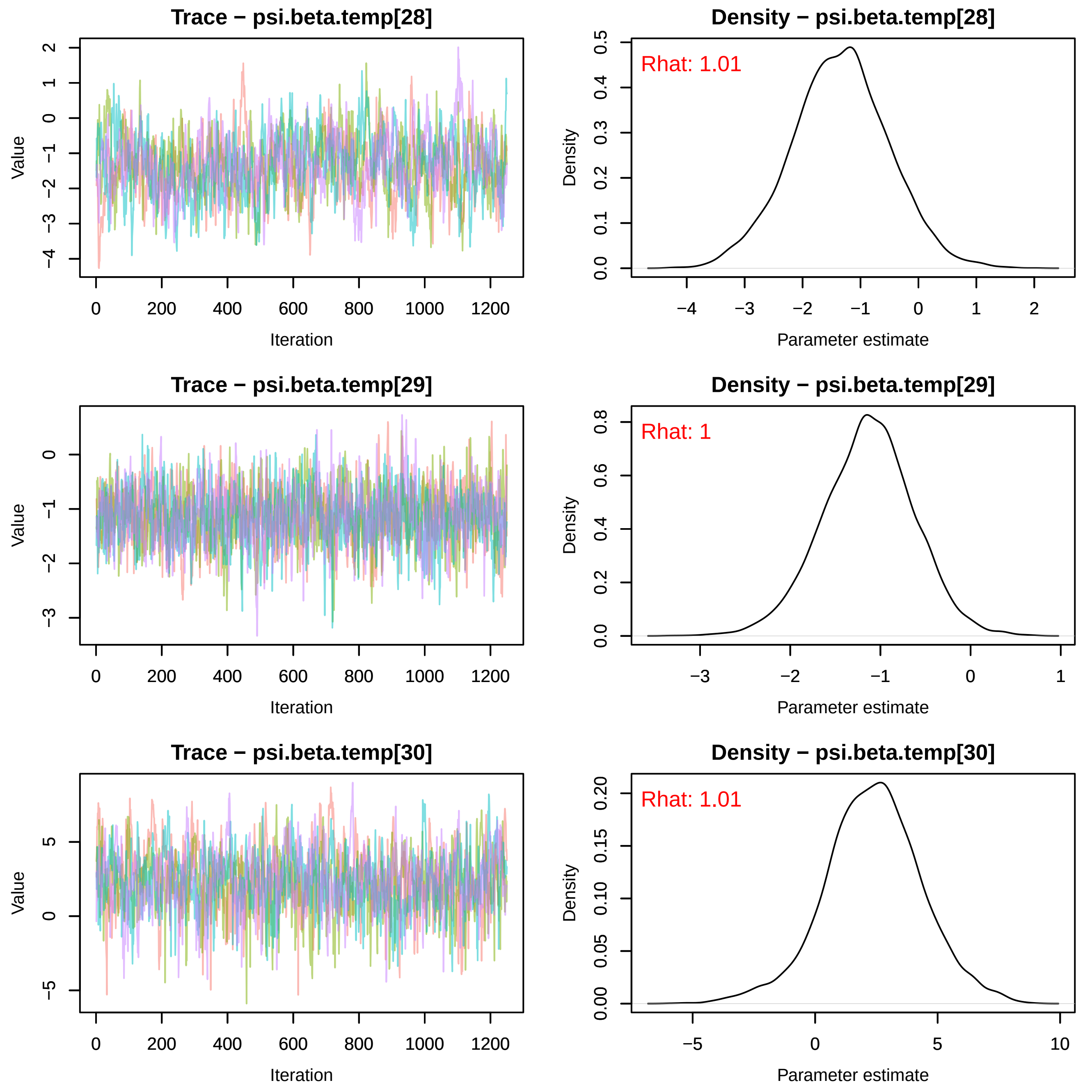

Trace − psi.beta.temp[28]
Value
Iteration
Density − psi.beta.temp[28]
Rhat: 1.01
Density
Parameter estimate
Trace − psi.beta.temp[29]
Value
Iteration
Density − psi.beta.temp[29]
Rhat: 1
Density
Parameter estimate
Trace − psi.beta.temp[30]
Value
Iteration
Density − psi.beta.temp[30]
Rhat: 1.01
Density
Parameter estimate

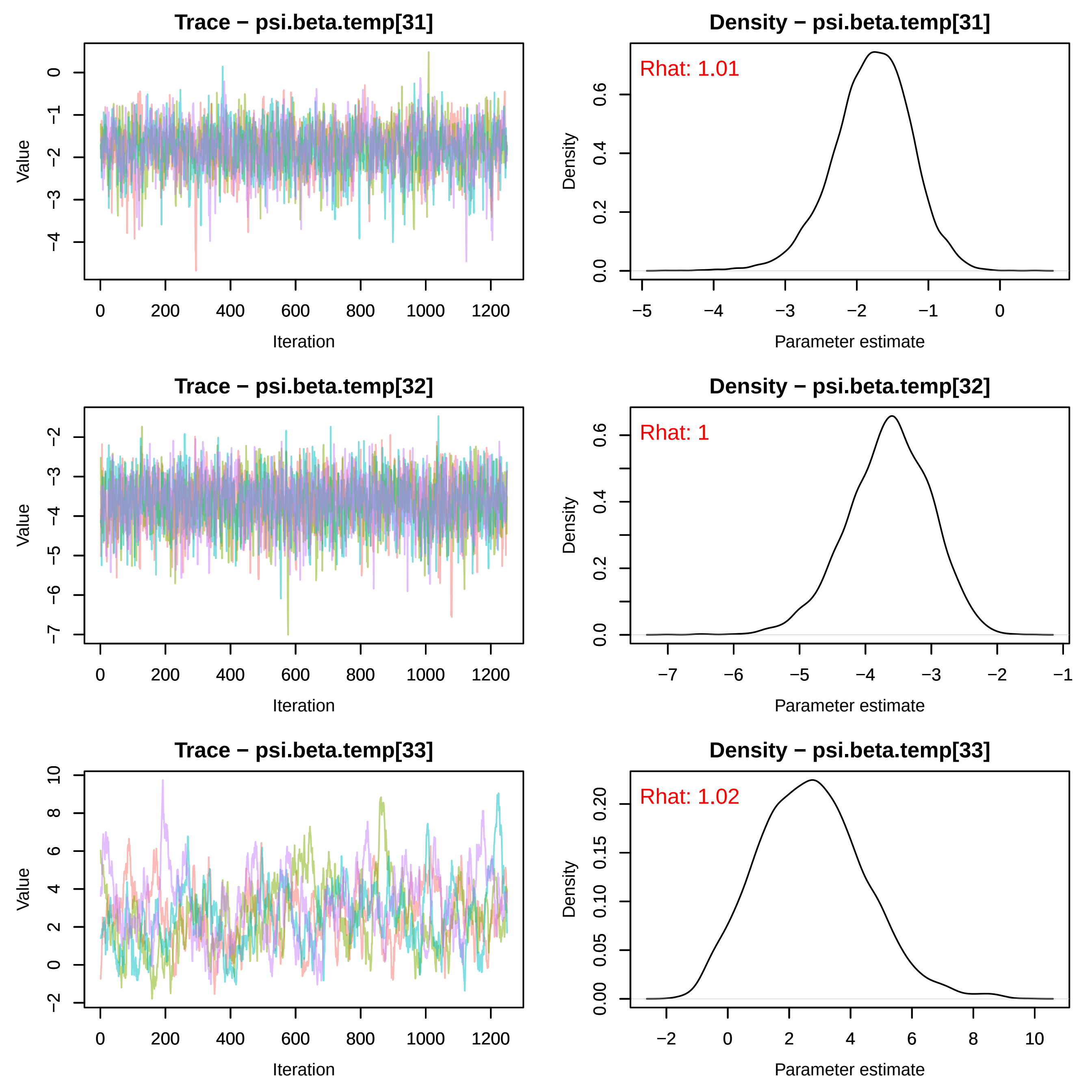
Trace − psi.beta.temp[31]
Density − psi.beta.temp[31]
Rhat: 1.01
Value
Iteration
Density
Parameter estimate
Trace − psi.beta.temp[32]
Density − psi.beta.temp[32]
Rhat: 1
Value
Iteration
Density
Parameter estimate
Trace − psi.beta.temp[33]
Density − psi.beta.temp[33]
Rhat: 1.02
Value
Iteration
Density
Parameter estimate

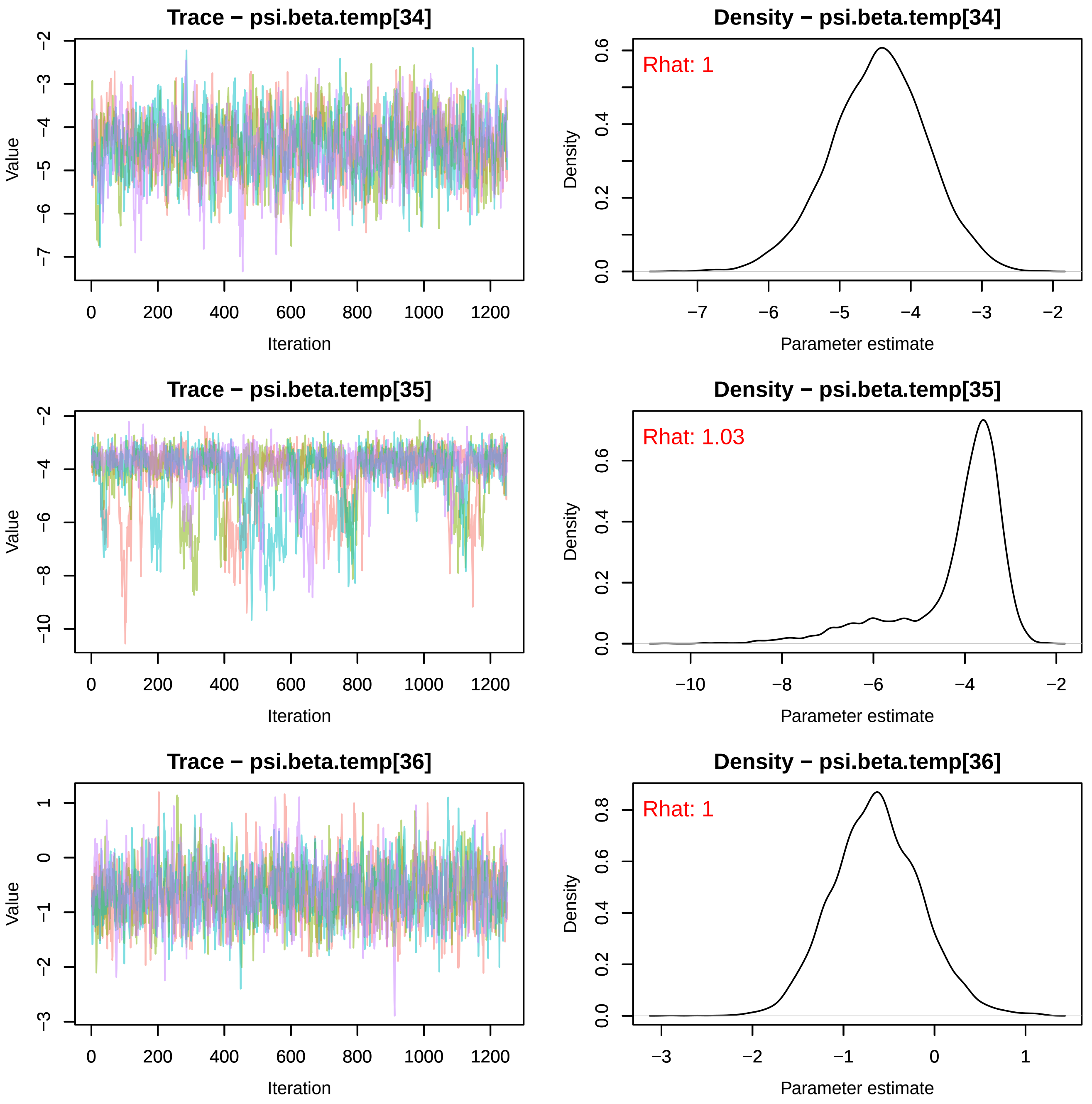
Trace − psi.beta.temp[34]
Value
Iteration
Density − psi.beta.temp[34]
Rhat: 1
Density
Parameter estimate
Trace − psi.beta.temp[35]
Value
Iteration
Density − psi.beta.temp[35]
Rhat: 1.03
Density
Parameter estimate
Trace − psi.beta.temp[36]
Value
Iteration
Density − psi.beta.temp[36]
Rhat: 1
Density
Parameter estimate

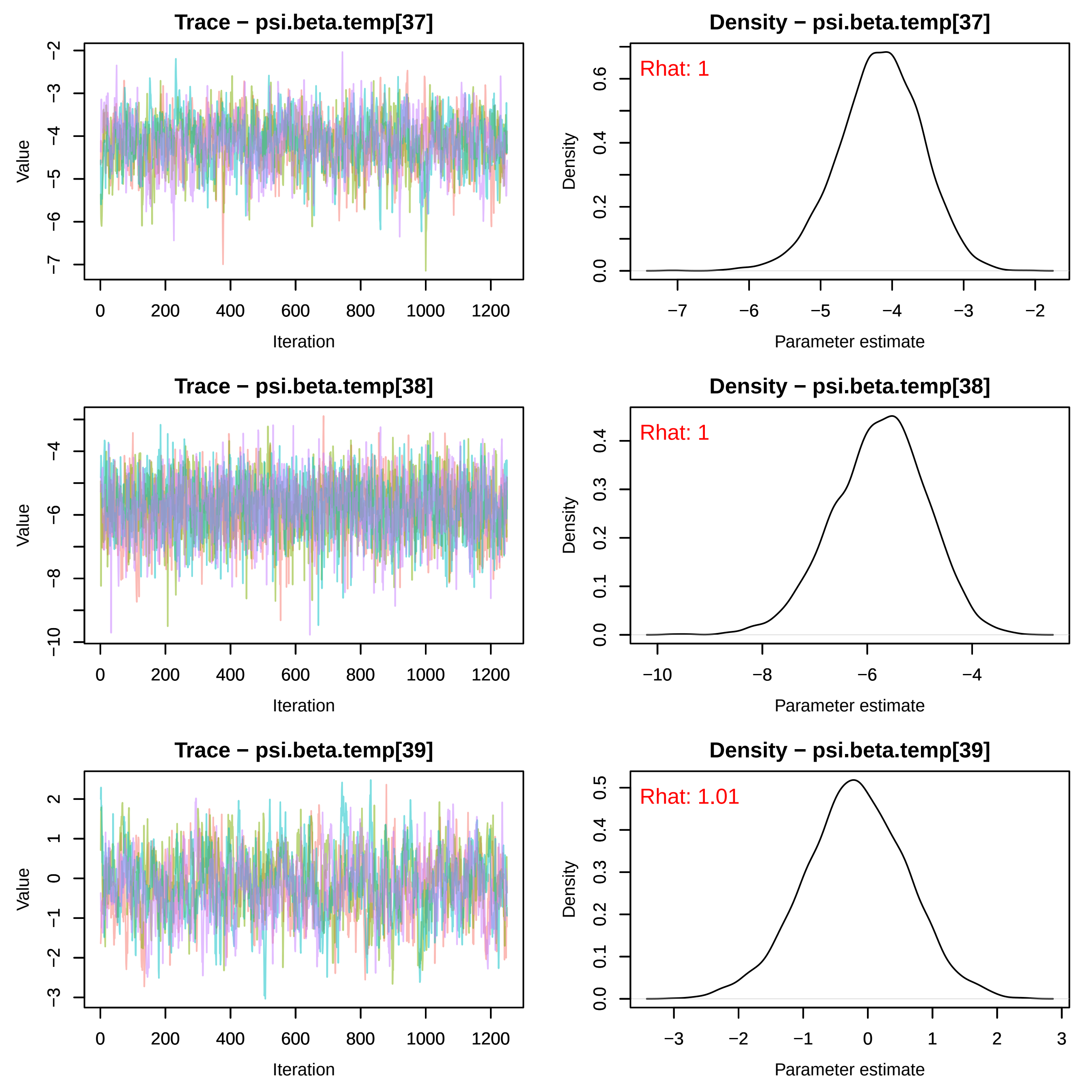
Trace − psi.beta.temp[37]
Density − psi.beta.temp[37]
Rhat: 1
Trace − psi.beta.temp[38]
Density − psi.beta.temp[38]
Rhat: 1
Trace − psi.beta.temp[39]
Density − psi.beta.temp[39]
Rhat: 1.01
Value
Iteration
Density
Parameter estimate

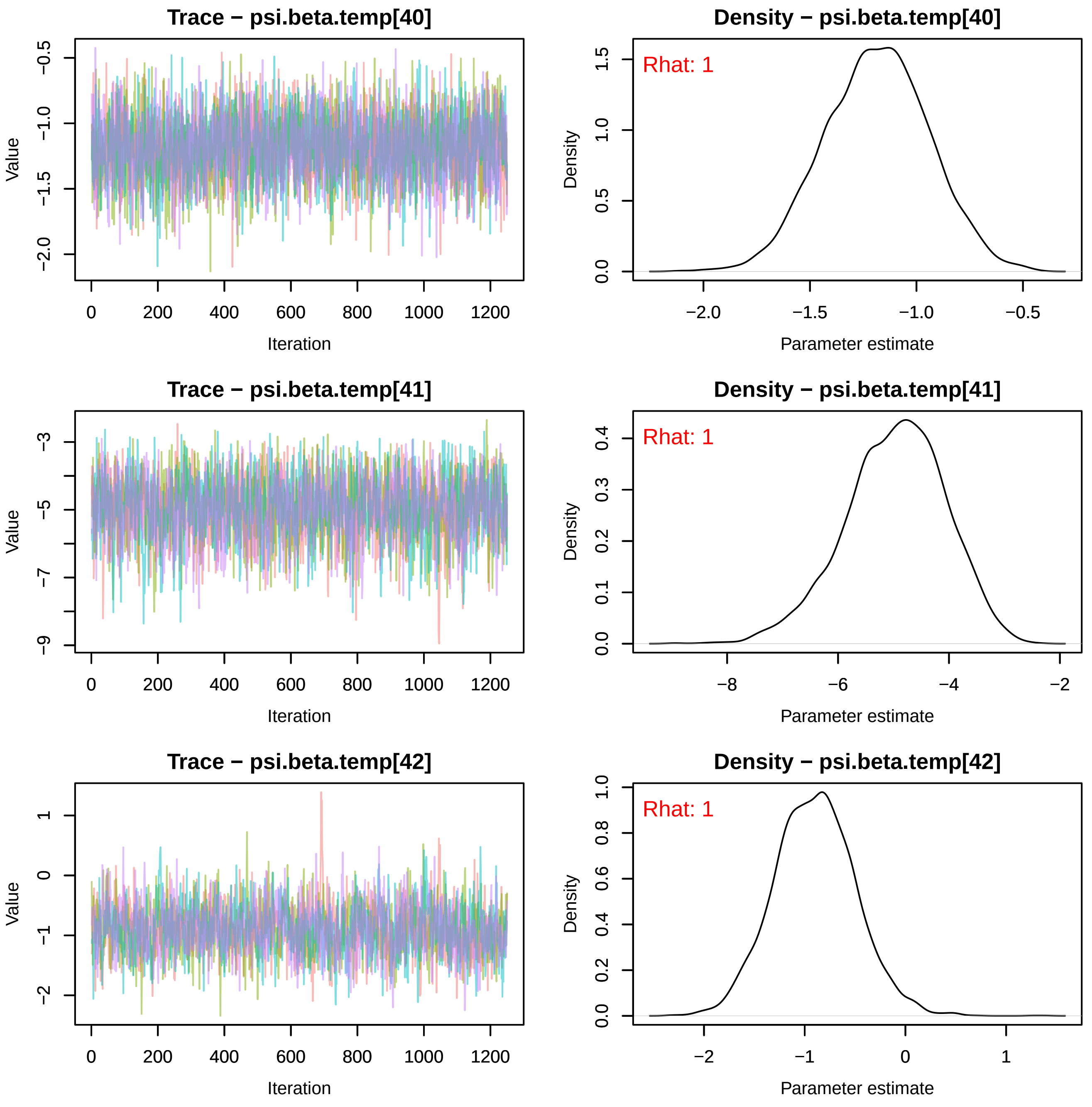
Trace − psi.beta.temp[40]
Value
Iteration
Density − psi.beta.temp[40]
Rhat: 1
Density
Parameter estimate
Trace − psi.beta.temp[41]
Value
Iteration
Density − psi.beta.temp[41]
Rhat: 1
Density
Parameter estimate
Trace − psi.beta.temp[42]
Value
Iteration
Density − psi.beta.temp[42]
Rhat: 1
Density
Parameter estimate

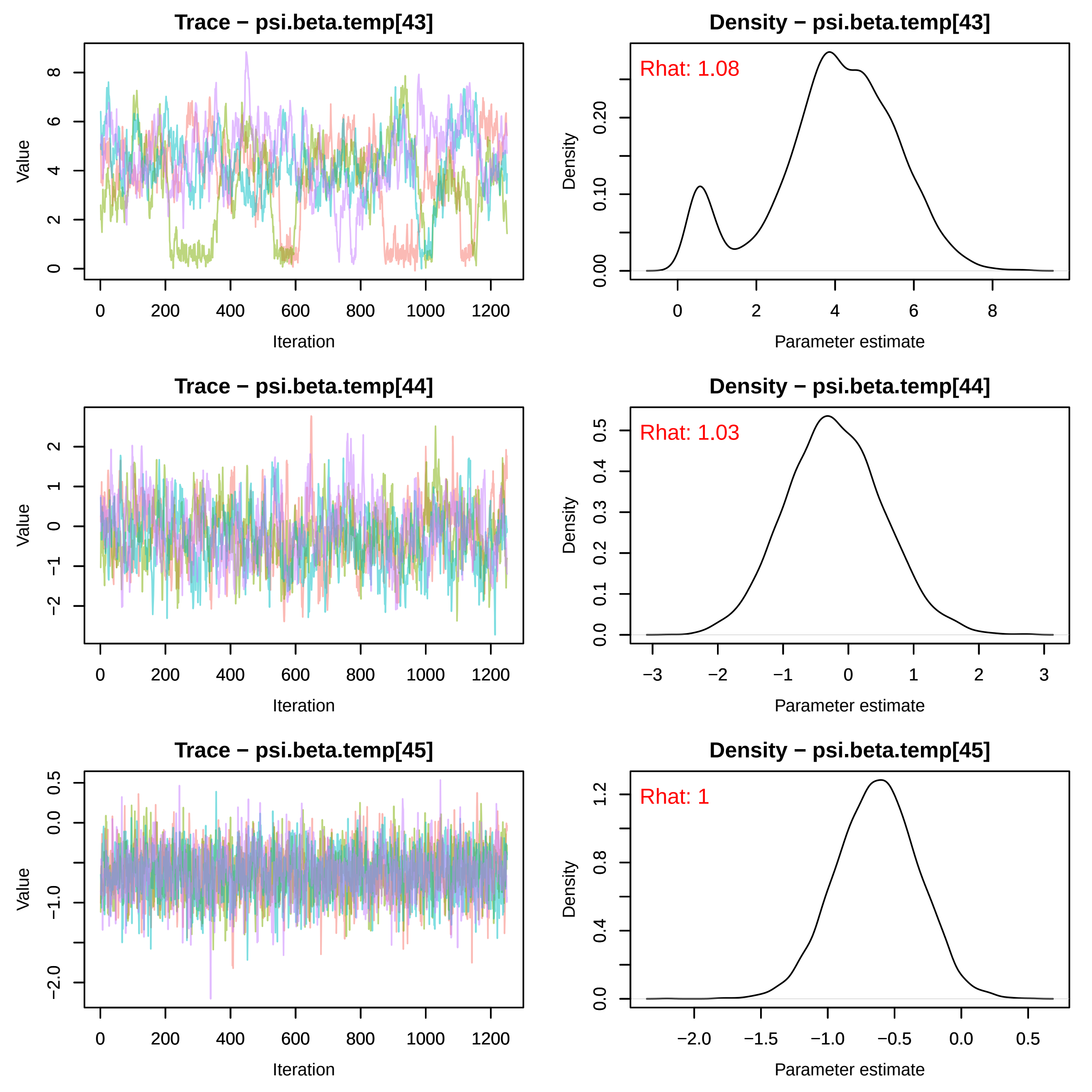
Trace − psi.beta.temp[43]
Value
Iteration
Density − psi.beta.temp[43]
Rhat: 1.08
Density
Parameter estimate
Trace − psi.beta.temp[44]
Value
Iteration
Density − psi.beta.temp[44]
Rhat: 1.03
Density
Parameter estimate
Trace − psi.beta.temp[45]
Value
Iteration
Density − psi.beta.temp[45]
Rhat: 1
Density
Parameter estimate

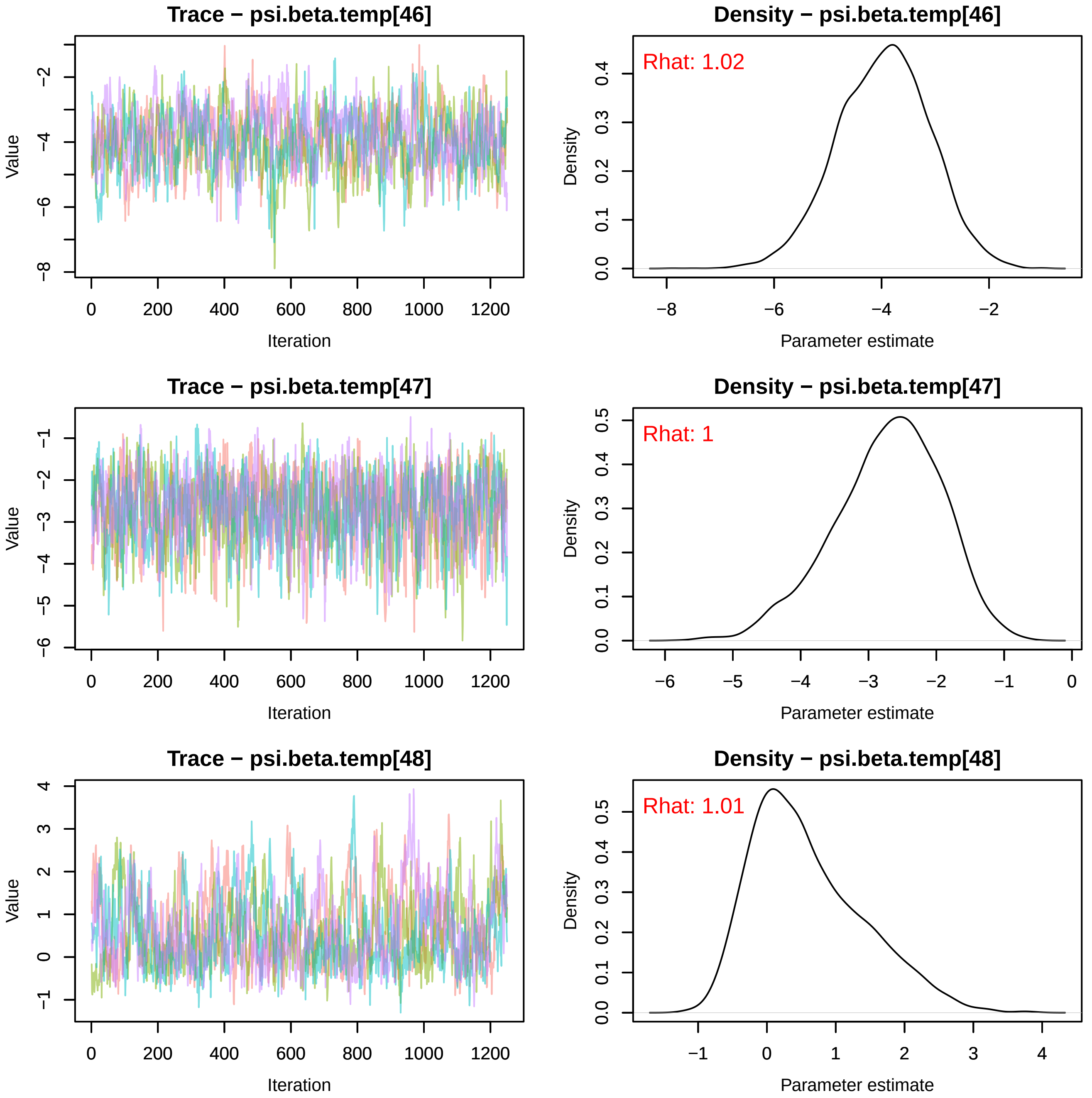
Trace − psi.beta.temp[46]
Value
Iteration
Density − psi.beta.temp[46]
Rhat: 1.02
Density
Parameter estimate
Trace − psi.beta.temp[47]
Value
Iteration
Density − psi.beta.temp[47]
Rhat: 1
Density
Parameter estimate
Trace − psi.beta.temp[48]
Value
Iteration
Density − psi.beta.temp[48]
Rhat: 1.01
Density
Parameter estimate

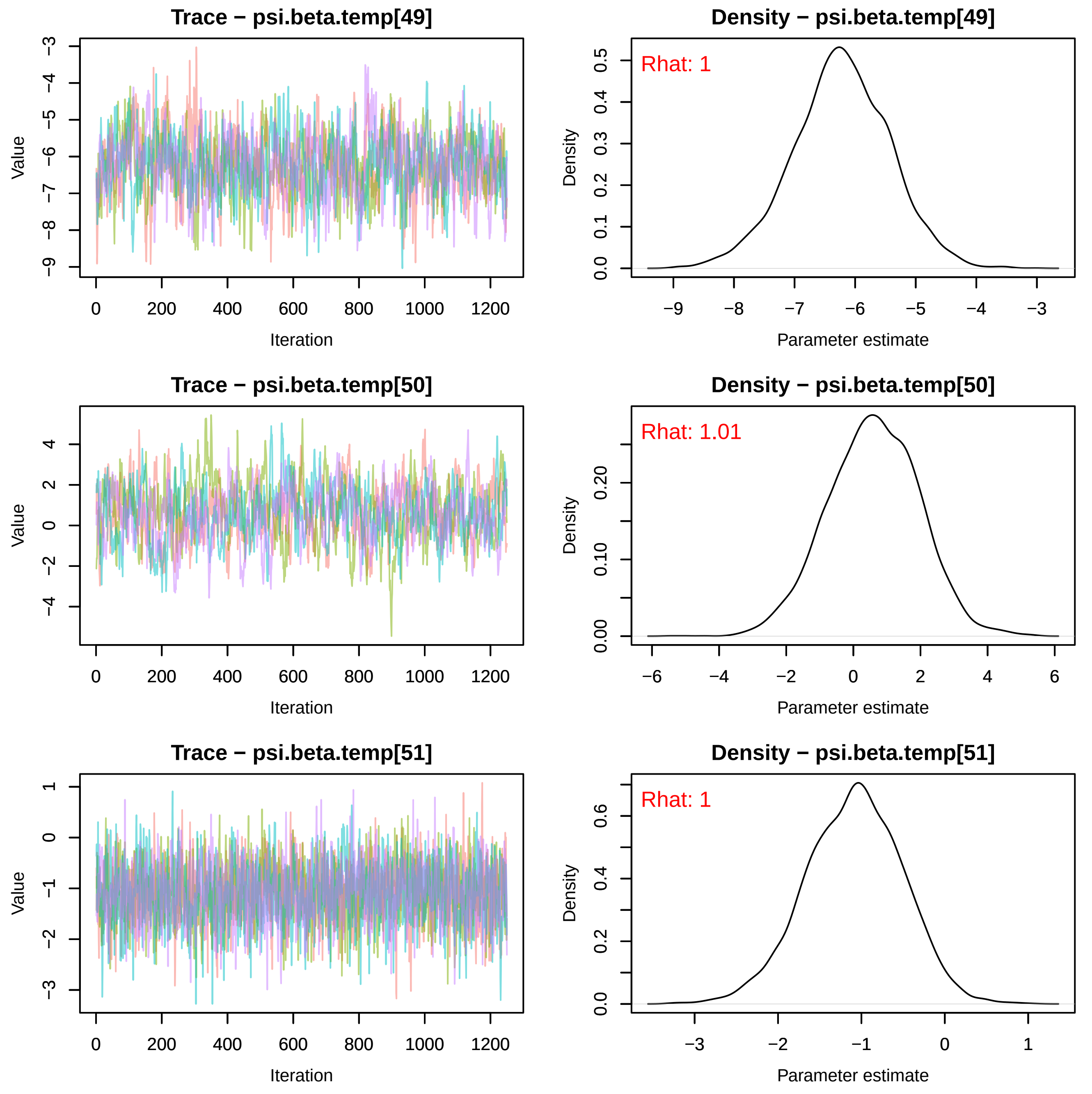
Trace − psi.beta.temp[49]
Value
−3
−4
−5
−6
−7
−8
−9
0
200
400
600
800
1000
1200
Iteration
Density − psi.beta.temp[49]
Rhat: 1
Density
0.0
0.1
0.2
0.3
0.4
0.5
−9
−8
−7
−6
−5
−4
−3
Parameter estimate
Trace − psi.beta.temp[50]
Value
4
2
0
−2
−4
0
200
400
600
800
1000
1200
Iteration
Density − psi.beta.temp[50]
Rhat: 1.01
Density
0.00
0.10
0.20
−6
−4
−2
0
2
4
6
Parameter estimate
Trace − psi.beta.temp[51]
Value
1
0
−1
−2
−3
0
200
400
600
800
1000
1200
Iteration
Density − psi.beta.temp[51]
Rhat: 1
Density
0.0
0.2
0.4
0.6
−3
−2
−1
0
1
Parameter estimate

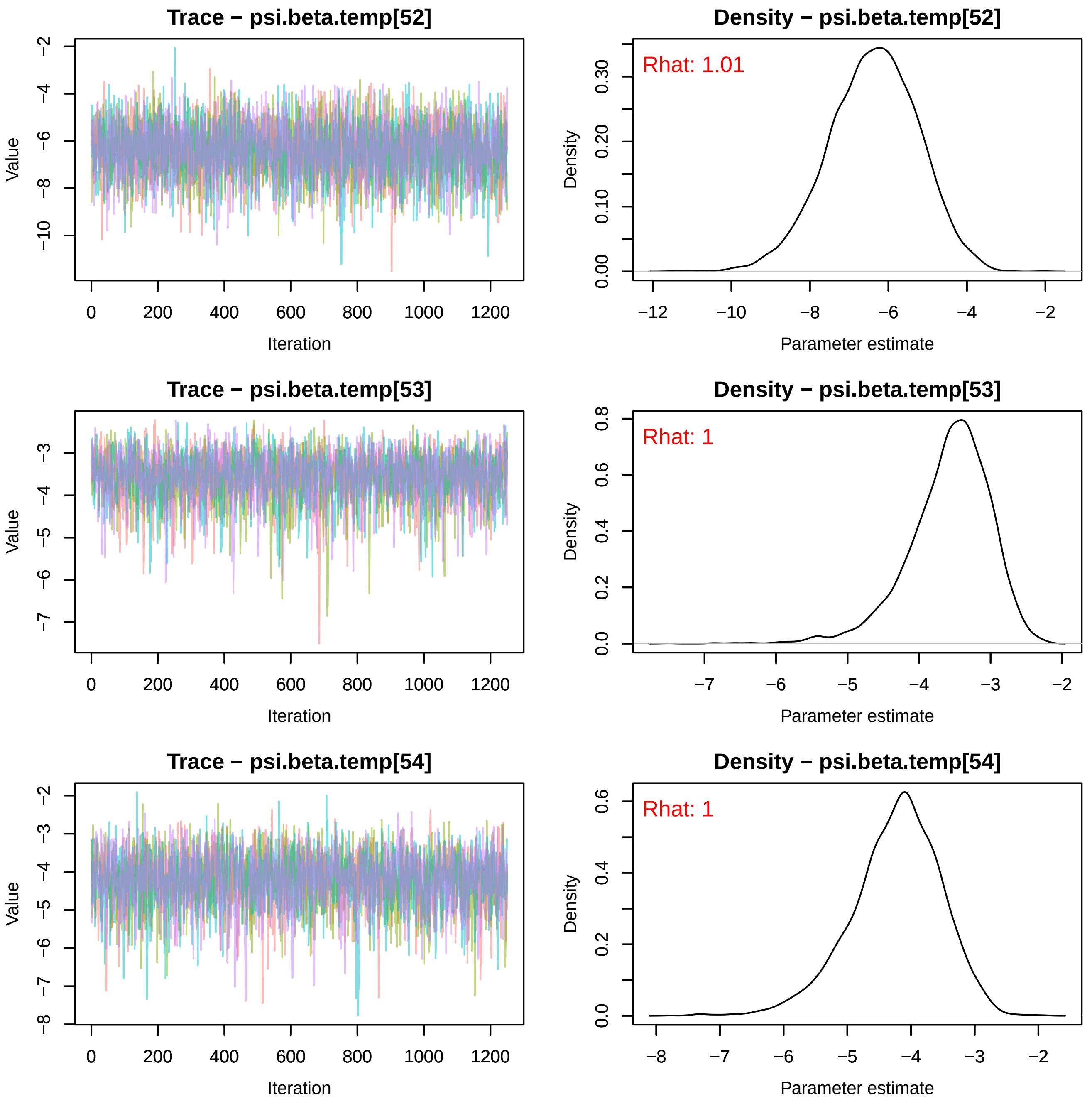
Trace − psi.beta.temp[52]
Value
Iteration
Density − psi.beta.temp[52]
Rhat: 1.01
Density
Parameter estimate
Trace − psi.beta.temp[53]
Value
Iteration
Density − psi.beta.temp[53]
Rhat: 1
Density
Parameter estimate
Trace − psi.beta.temp[54]
Value
Iteration
Density − psi.beta.temp[54]
Rhat: 1
Density
Parameter estimate

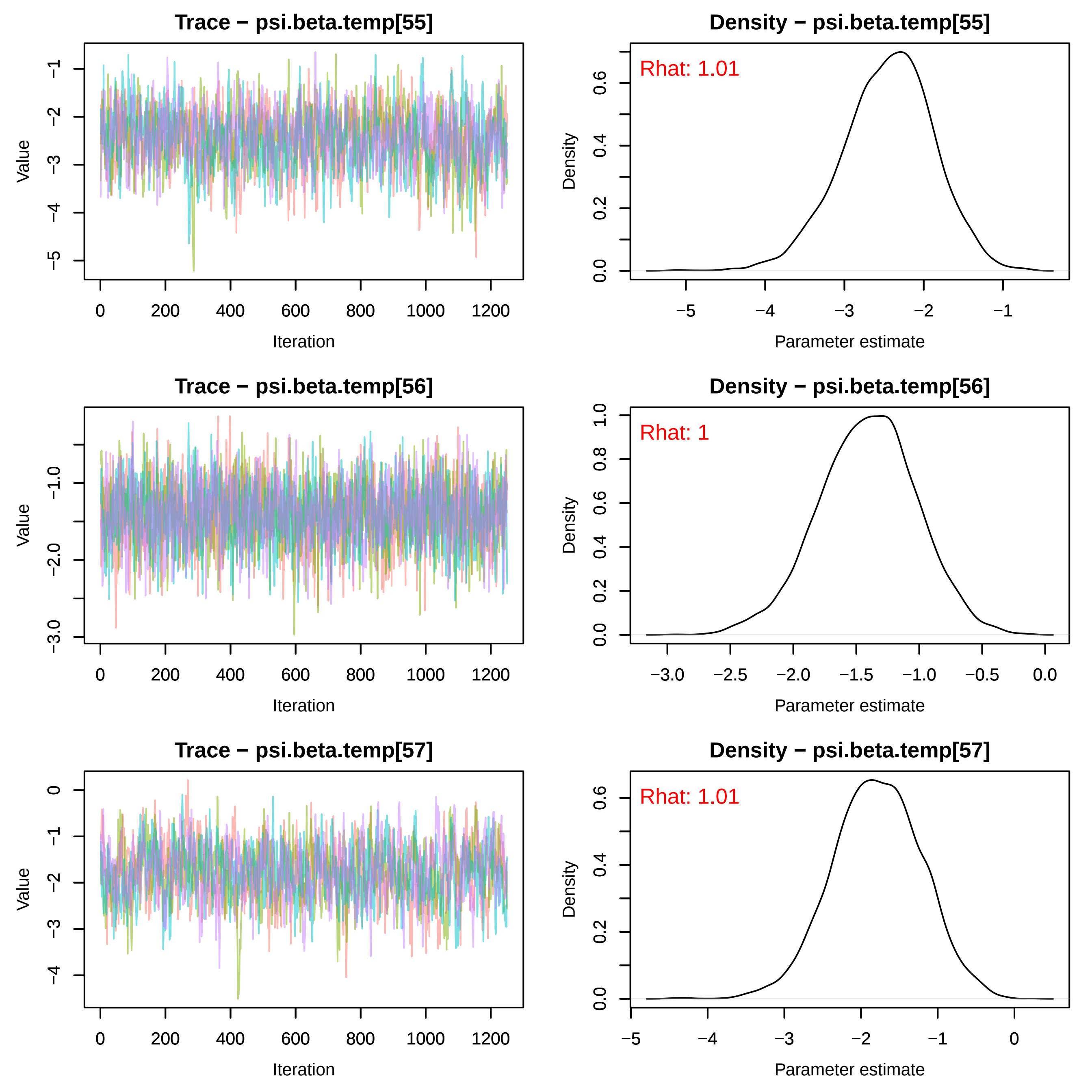
Trace − psi.beta.temp[55]
Value
Iteration
Density − psi.beta.temp[55]
Rhat: 1.01
Density
Parameter estimate
Trace − psi.beta.temp[56]
Value
Iteration
Density − psi.beta.temp[56]
Rhat: 1
Density
Parameter estimate
Trace − psi.beta.temp[57]
Value
Iteration
Density − psi.beta.temp[57]
Rhat: 1.01
Density
Parameter estimate

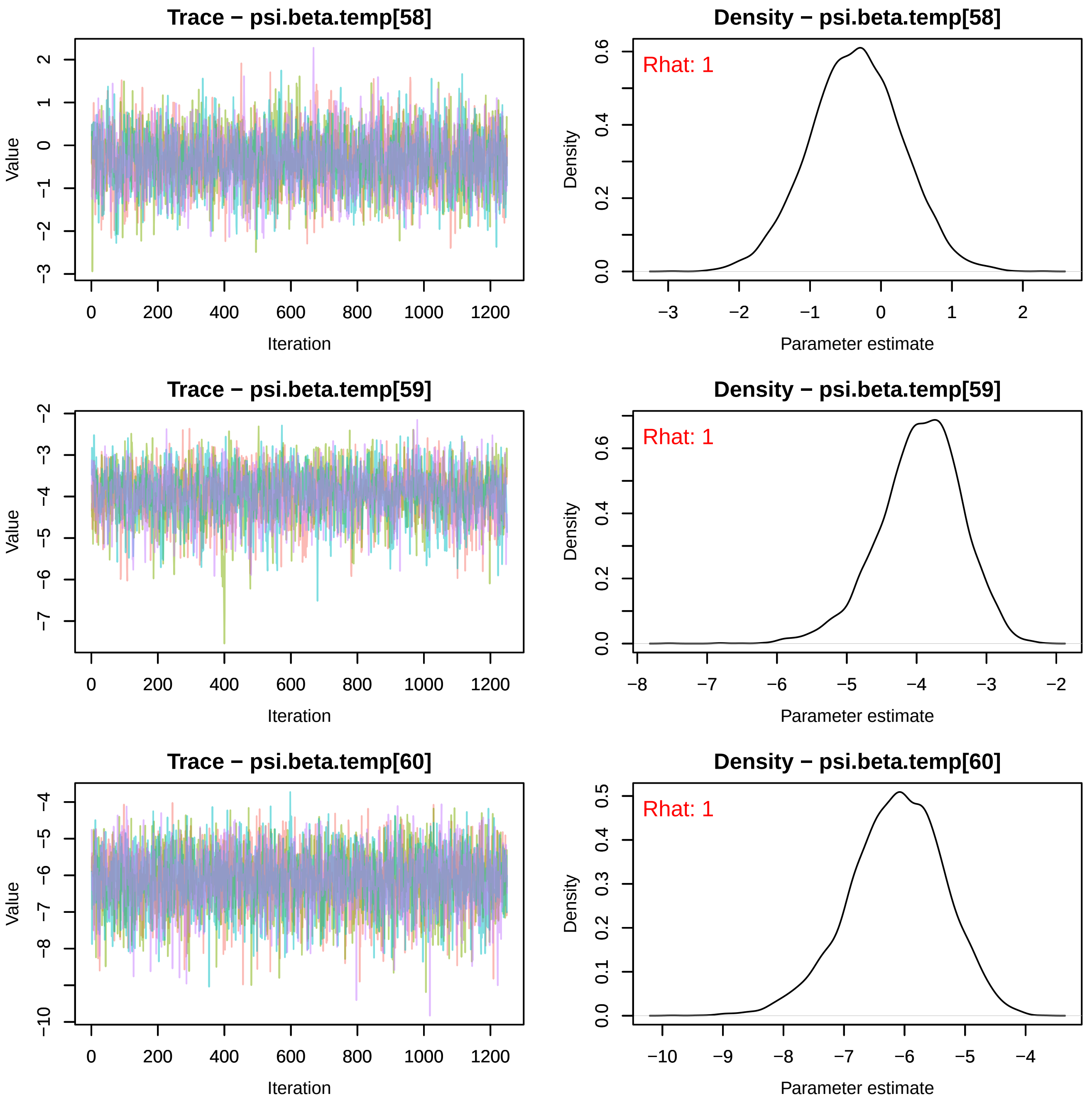

Trace − psi.beta.temp[58]
Value
Iteration
Density − psi.beta.temp[58]
Rhat: 1
Density
Parameter estimate
Trace − psi.beta.temp[59]
Value
Iteration
Density − psi.beta.temp[59]
Rhat: 1
Density
Parameter estimate
Trace − psi.beta.temp[60]
Value
Iteration
Density − psi.beta.temp[60]
Rhat: 1
Density
Parameter estimate

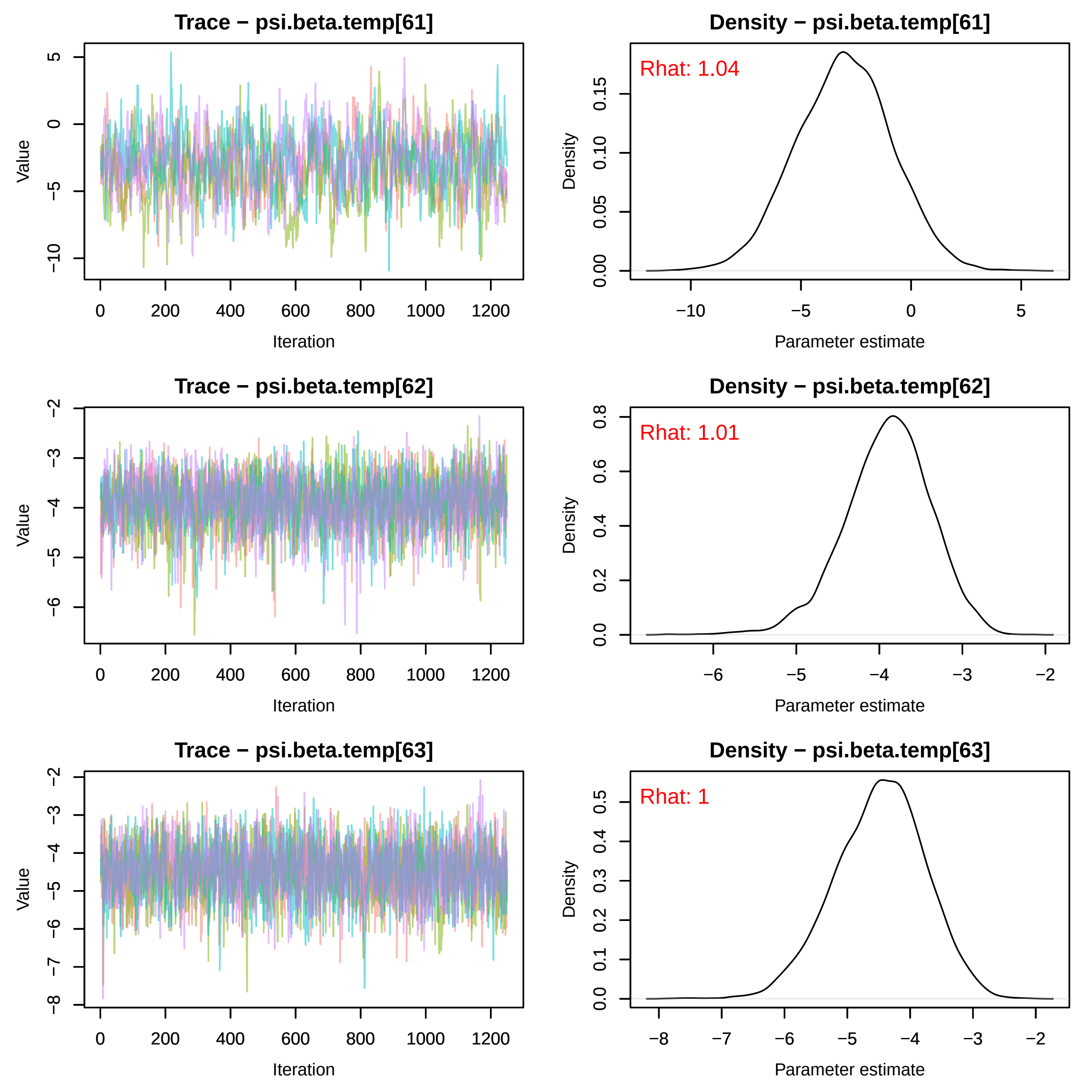
Trace − psi.beta.temp[61]
Value
Iteration
Density − psi.beta.temp[61]
Rhat: 1.04
Density
Parameter estimate
Trace − psi.beta.temp[62]
Value
Iteration
Density − psi.beta.temp[62]
Rhat: 1.01
Density
Parameter estimate
Trace − psi.beta.temp[63]
Value
Iteration
Density − psi.beta.temp[63]
Rhat: 1
Density
Parameter estimate

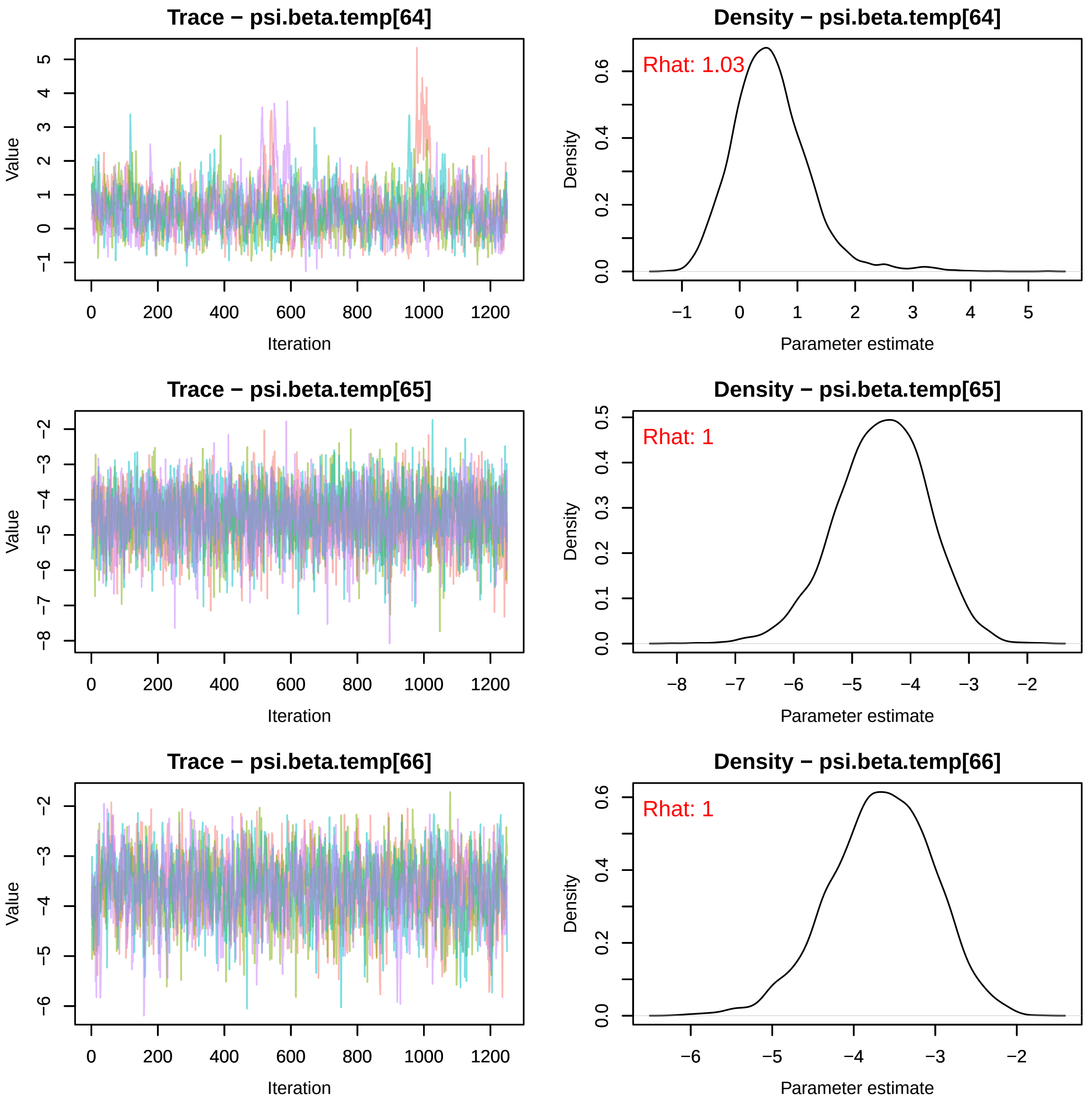
Trace − psi.beta.temp[64]
Value
Iteration
0 200 400 600 800 1000 1200
−1 0 1 2 3 4 5
Density − psi.beta.temp[64]
Rhat: 1.03
Density
Parameter estimate
−1 0 1 2 3 4 5
0.0 0.2 0.4 0.6
Trace − psi.beta.temp[65]
Value
Iteration
0 200 400 600 800 1000 1200
−8 −7 −6 −5 −4 −3 −2
Density − psi.beta.temp[65]
Rhat: 1
Density
Parameter estimate
−8 −7 −6 −5 −4 −3 −2
0.0 0.1 0.2 0.3 0.4 0.5
Trace − psi.beta.temp[66]
Value
Iteration
0 200 400 600 800 1000 1200
−6 −5 −4 −3 −2
Density − psi.beta.temp[66]
Rhat: 1
Density
Parameter estimate
−6 −5 −4 −3 −2
0.0 0.2 0.4 0.6

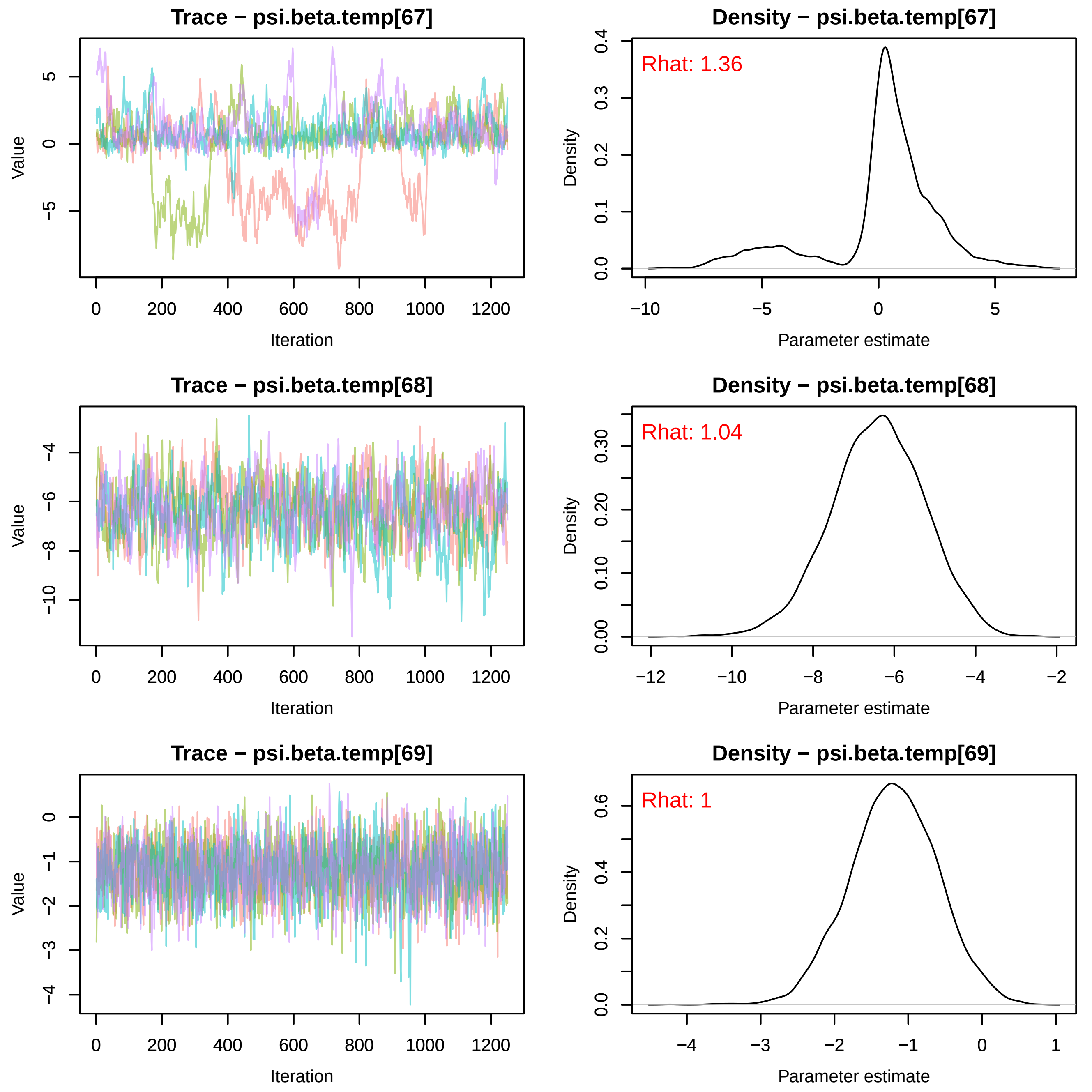
Trace − psi.beta.temp[67]
Value
Iteration
Density − psi.beta.temp[67]
Rhat: 1.36
Density
Parameter estimate
Trace − psi.beta.temp[68]
Value
Iteration
Density − psi.beta.temp[68]
Rhat: 1.04
Density
Parameter estimate
Trace − psi.beta.temp[69]
Value
Iteration
Density − psi.beta.temp[69]
Rhat: 1
Density
Parameter estimate

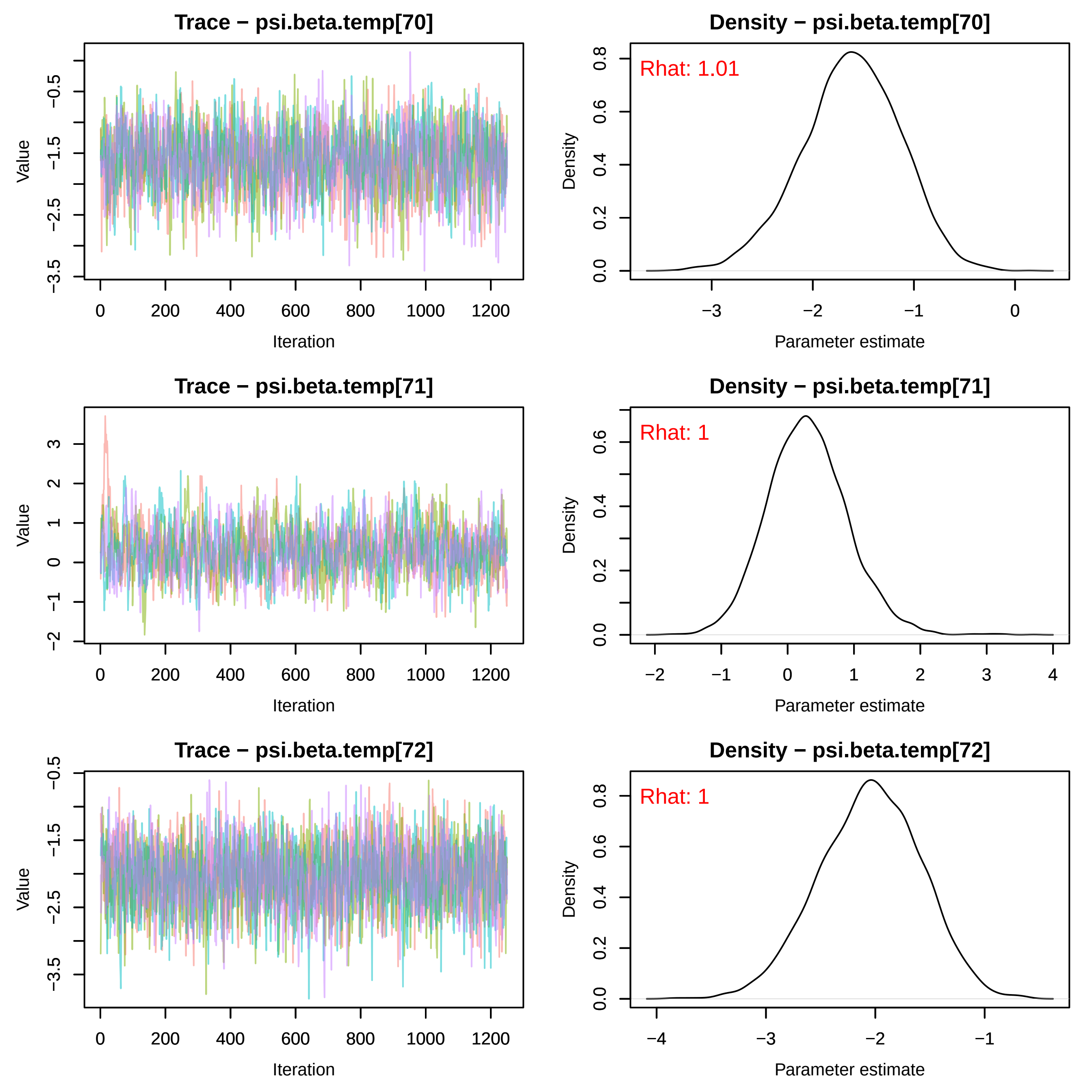
Trace − psi.beta.temp[70]
Value
Iteration
Density − psi.beta.temp[70]
Rhat: 1.01
Density
Parameter estimate
Trace − psi.beta.temp[71]
Value
Iteration
Density − psi.beta.temp[71]
Rhat: 1
Density
Parameter estimate
Trace − psi.beta.temp[72]
Value
Iteration
Density − psi.beta.temp[72]
Rhat: 1
Density
Parameter estimate

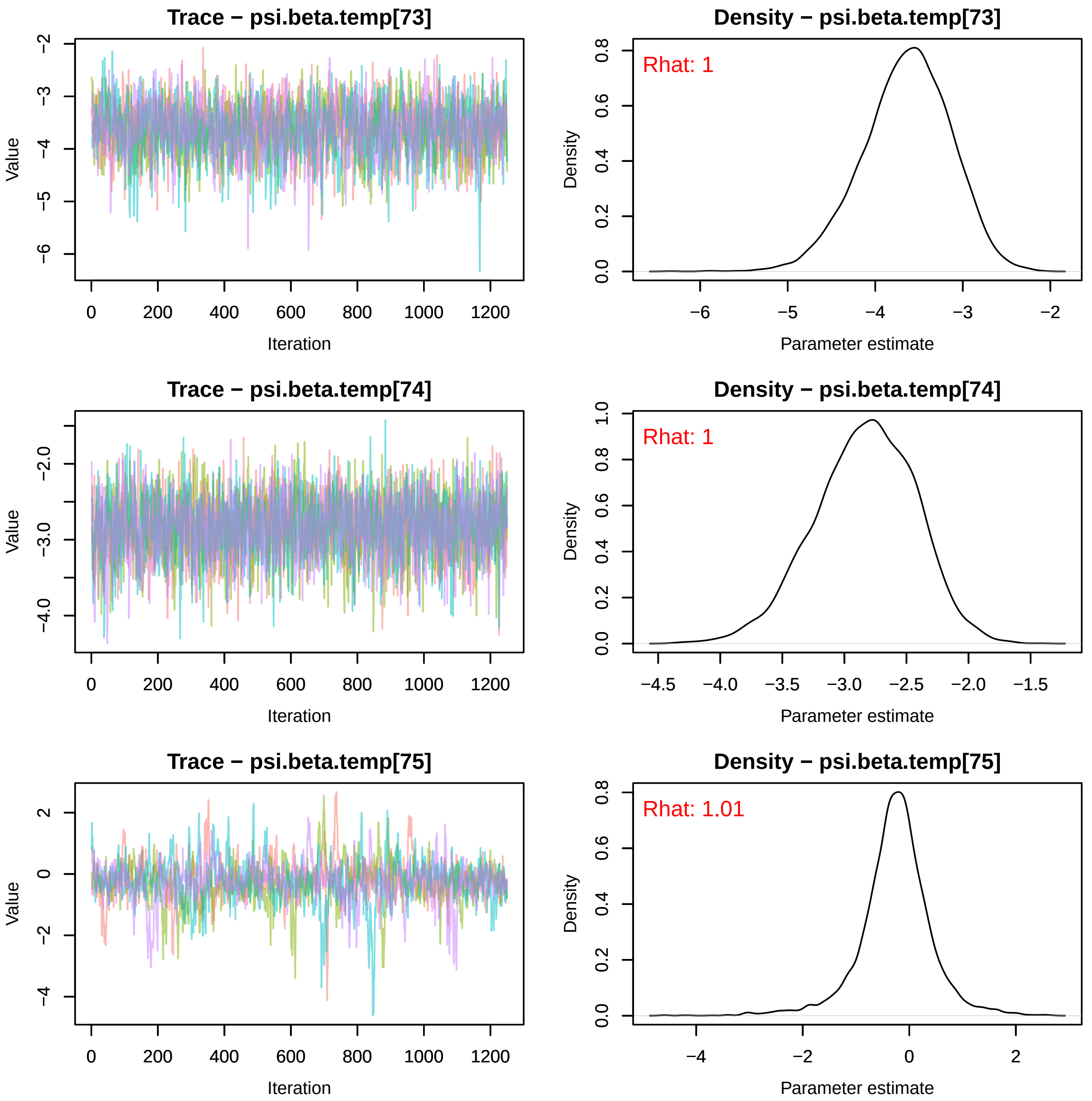
Trace − psi.beta.temp[73]
Value
Iteration
Density − psi.beta.temp[73]
Rhat: 1
Density
Parameter estimate
Trace − psi.beta.temp[74]
Value
Iteration
Density − psi.beta.temp[74]
Rhat: 1
Density
Parameter estimate
Trace − psi.beta.temp[75]
Value
Iteration
Density − psi.beta.temp[75]
Rhat: 1.01
Density
Parameter estimate

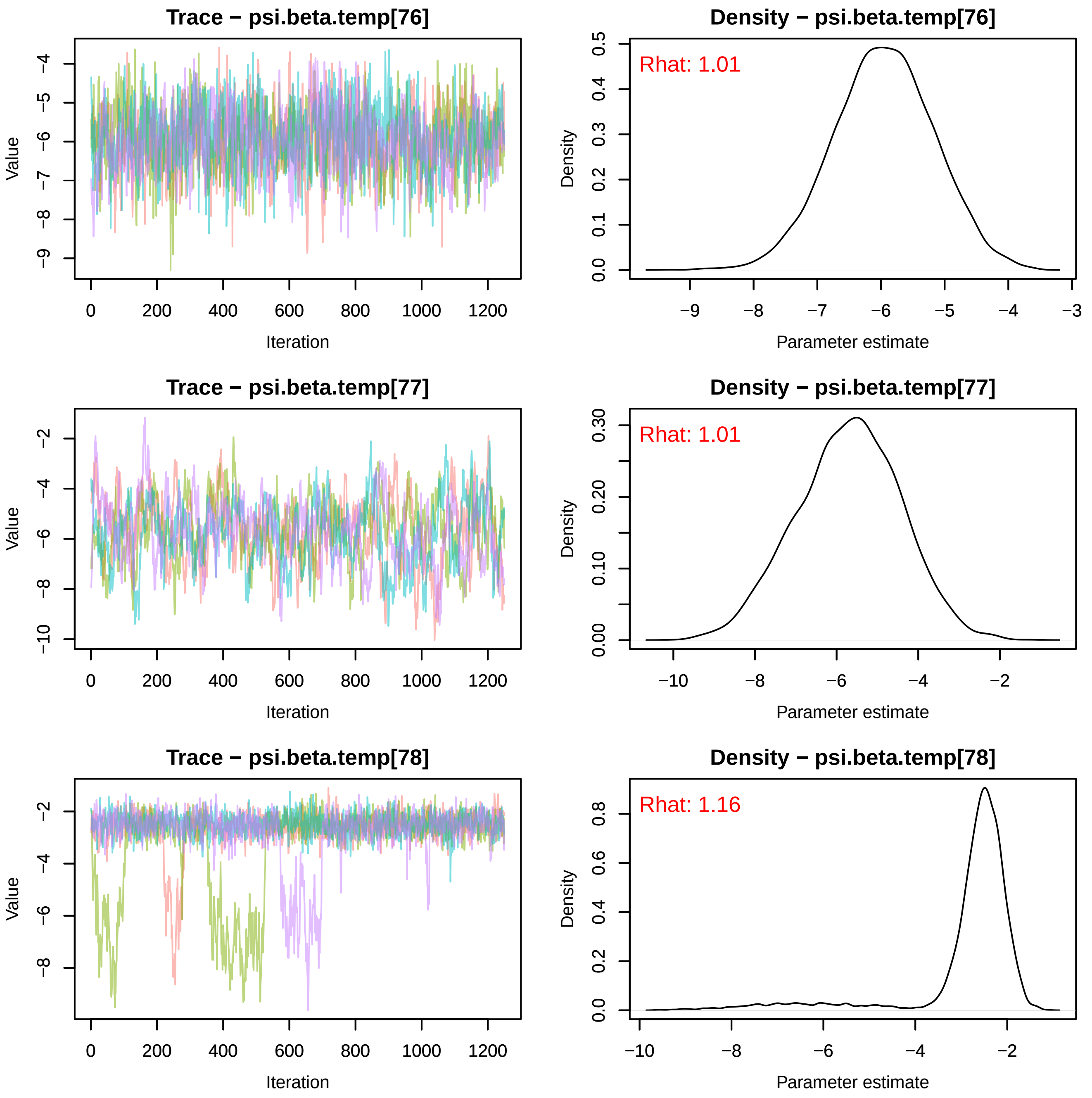
Trace − psi.beta.temp[76]
Value
Iteration
Density − psi.beta.temp[76]
Rhat: 1.01
Density
Parameter estimate
Trace − psi.beta.temp[77]
Value
Iteration
Density − psi.beta.temp[77]
Rhat: 1.01
Density
Parameter estimate
Trace − psi.beta.temp[78]
Value
Iteration
Density − psi.beta.temp[78]
Rhat: 1.16
Density
Parameter estimate

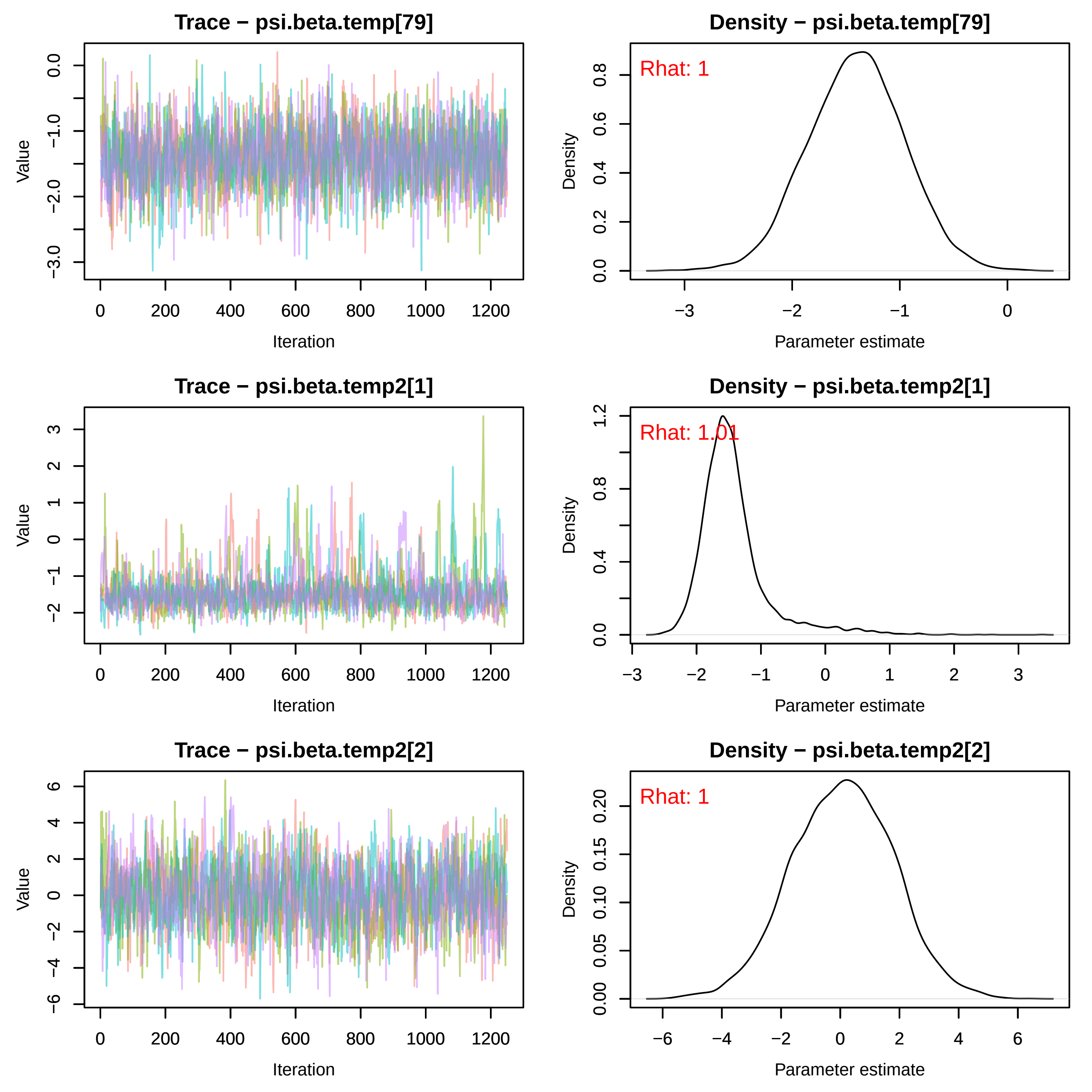
Trace − psi.beta.temp[79]
Value
Iteration
Density − psi.beta.temp[79]
Rhat: 1
Density
Parameter estimate
Trace − psi.beta.temp2[1]
Value
Iteration
Density − psi.beta.temp2[1]
Rhat: 1.01
Density
Parameter estimate
Trace − psi.beta.temp2[2]
Value
Iteration
Density − psi.beta.temp2[2]
Rhat: 1
Density
Parameter estimate

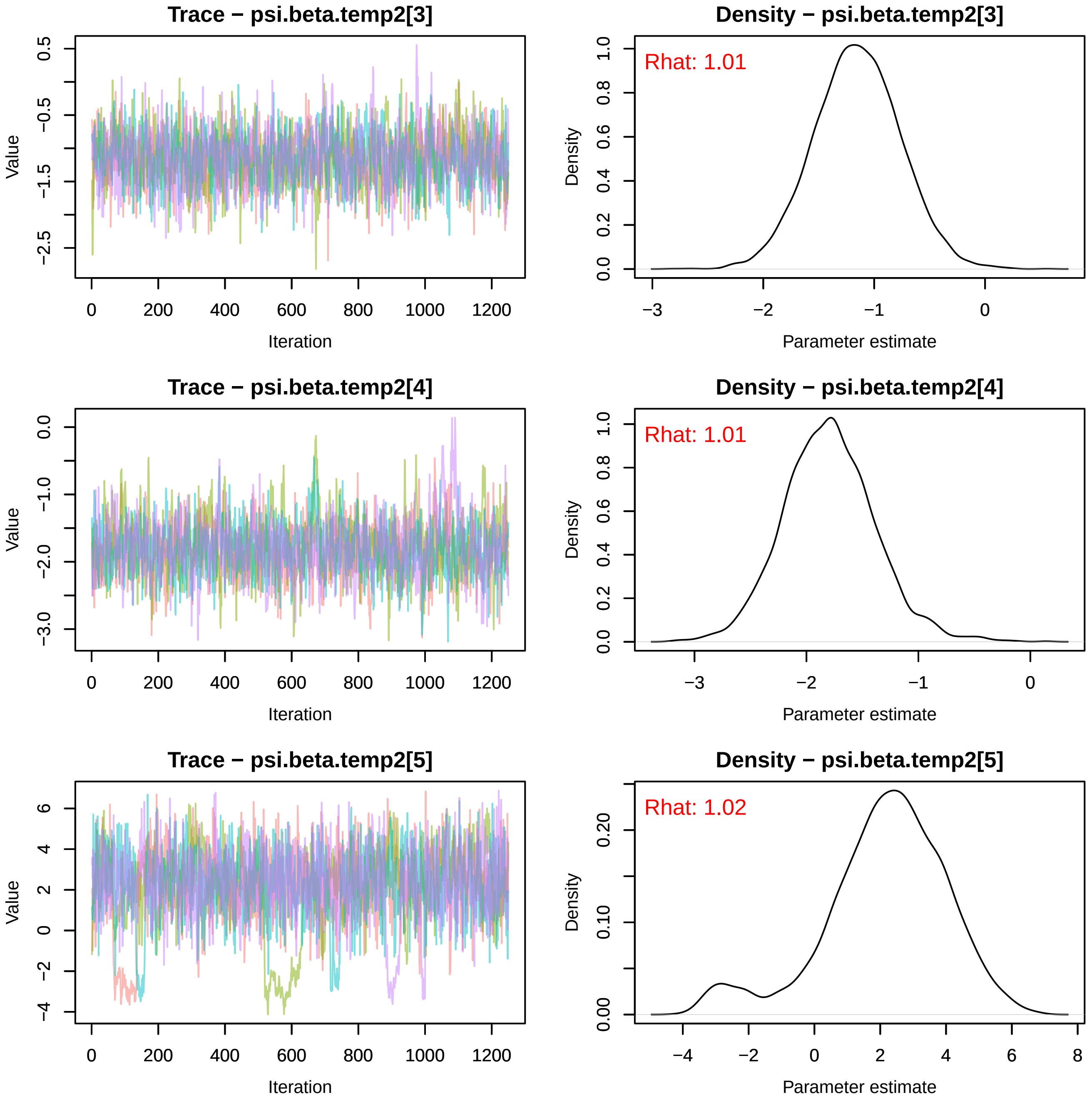
Trace − psi.beta.temp2[3]
Value
Iteration
0
200
400
600
800
1000
1200
0.5
−0.5
−1.5
−2.5
Density − psi.beta.temp2[3]
Rhat: 1.01
Density
Parameter estimate
0.0
0.2
0.4
0.6
0.8
1.0
−3
−2
−1
0
Trace − psi.beta.temp2[4]
Value
Iteration
0.0
−1.0
−2.0
−3.0
Density − psi.beta.temp2[4]
Rhat: 1.01
Density
Parameter estimate
Trace − psi.beta.temp2[5]
Value
Iteration
6
4
2
0
−2
−4
Density − psi.beta.temp2[5]
Rhat: 1.02
Density
Parameter estimate
0.00
0.10
0.20
−4
−2
0
2
4
6
8

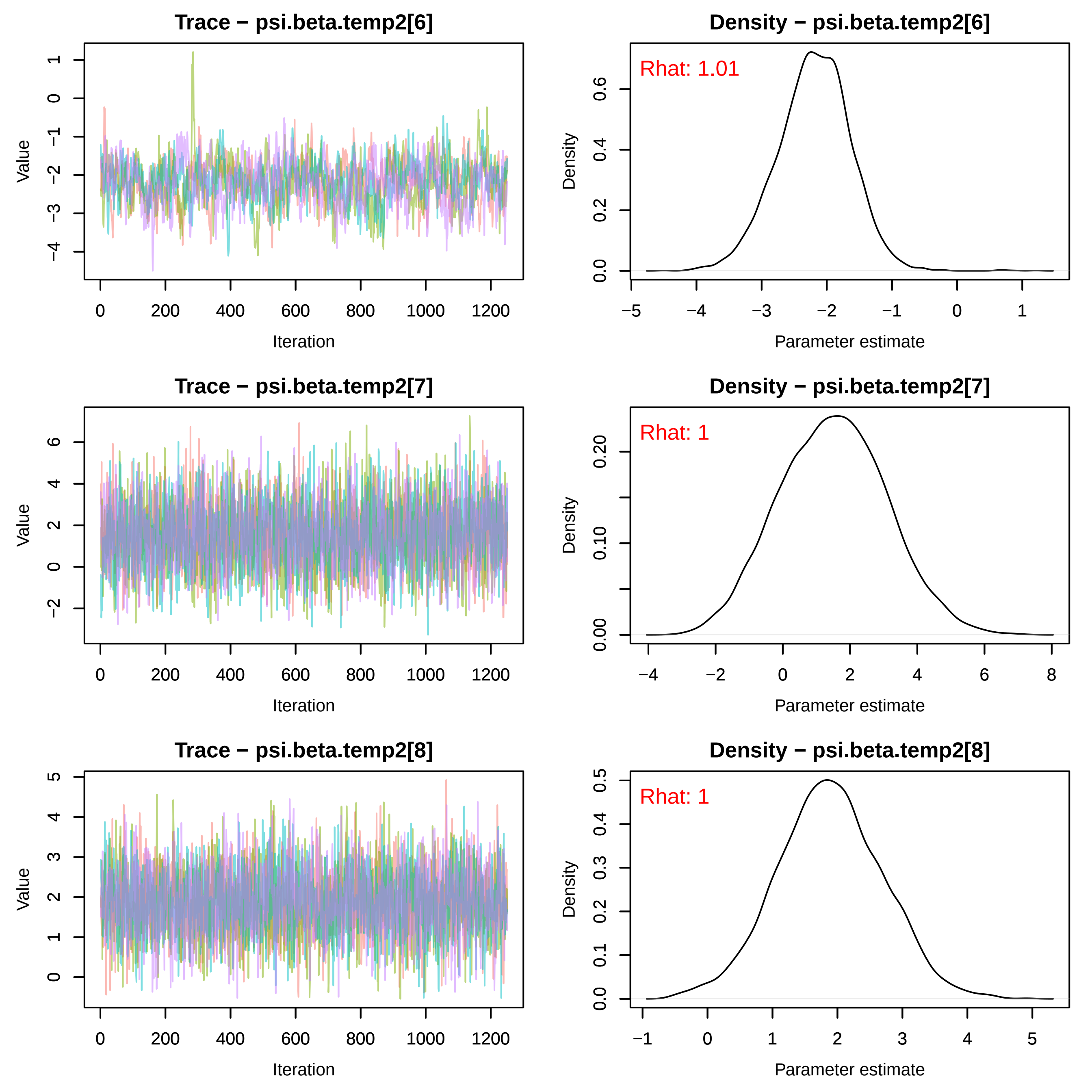
Trace − psi.beta.temp2[6]
Value
Iteration
0
200
400
600
800
1000
1200
1
0
−1
−2
−3
−4
Density − psi.beta.temp2[6]
Rhat: 1.01
Density
Parameter estimate
0.0
0.2
0.4
0.6
−5
−4
−3
−2
−1
0
1
Trace − psi.beta.temp2[7]
Value
Iteration
6
4
2
0
−2
Density − psi.beta.temp2[7]
Rhat: 1
Density
Parameter estimate
0.00
0.10
0.20
−4
−2
0
2
4
6
8
Trace − psi.beta.temp2[8]
Value
Iteration
5
4
3
2
1
0
Density − psi.beta.temp2[8]
Rhat: 1
Density
Parameter estimate
0.0
0.1
0.2
0.3
0.4
0.5
−1
0
1
2
3
4
5

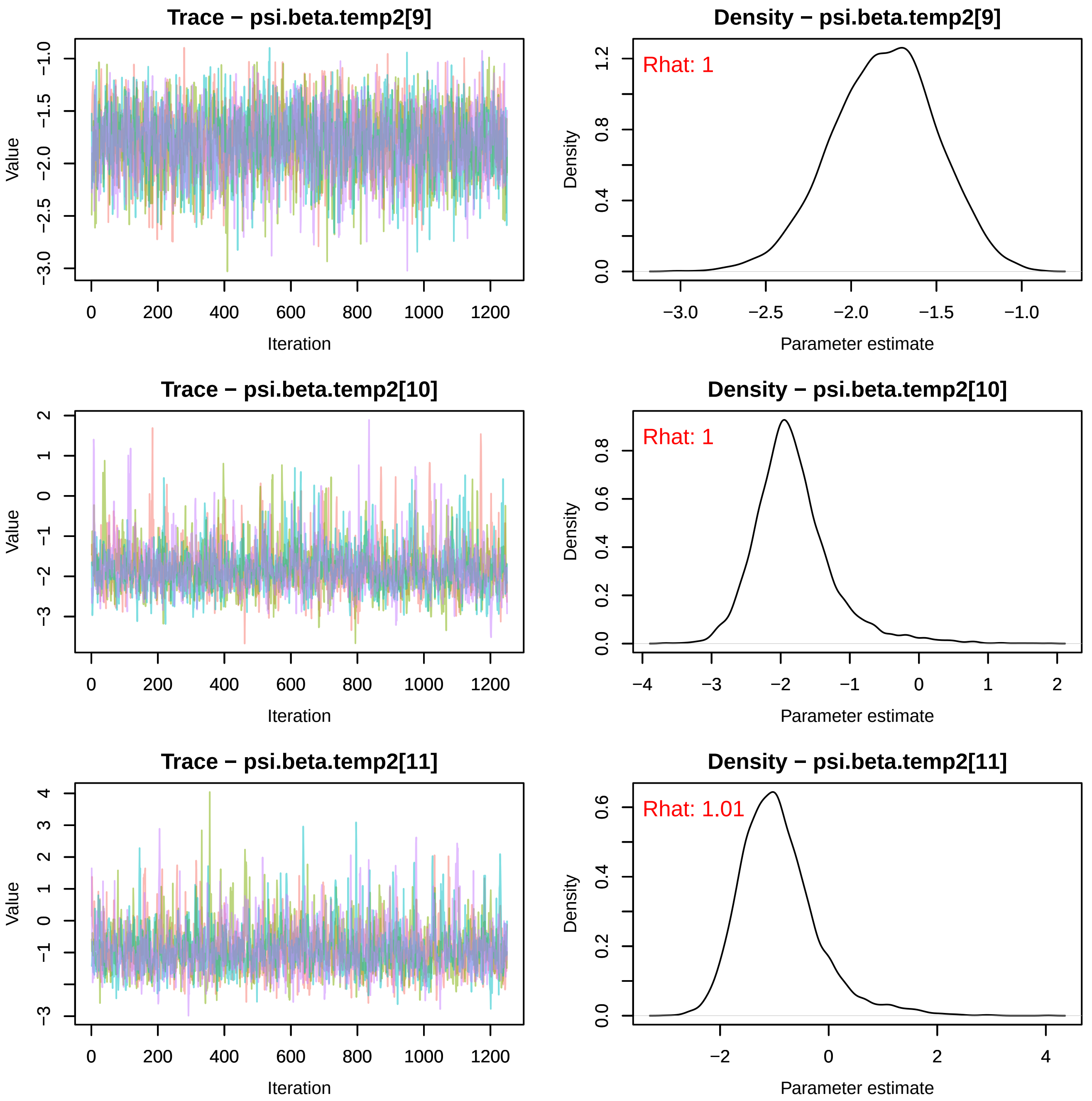
Trace − psi.beta.temp2[9]
Density − psi.beta.temp2[9]
Rhat: 1
Trace − psi.beta.temp2[10]
Density − psi.beta.temp2[10]
Rhat: 1
Trace − psi.beta.temp2[11]
Density − psi.beta.temp2[11]
Rhat: 1.01
Value
Iteration
Density
Parameter estimate
0
200
400
600
800
1000
1200

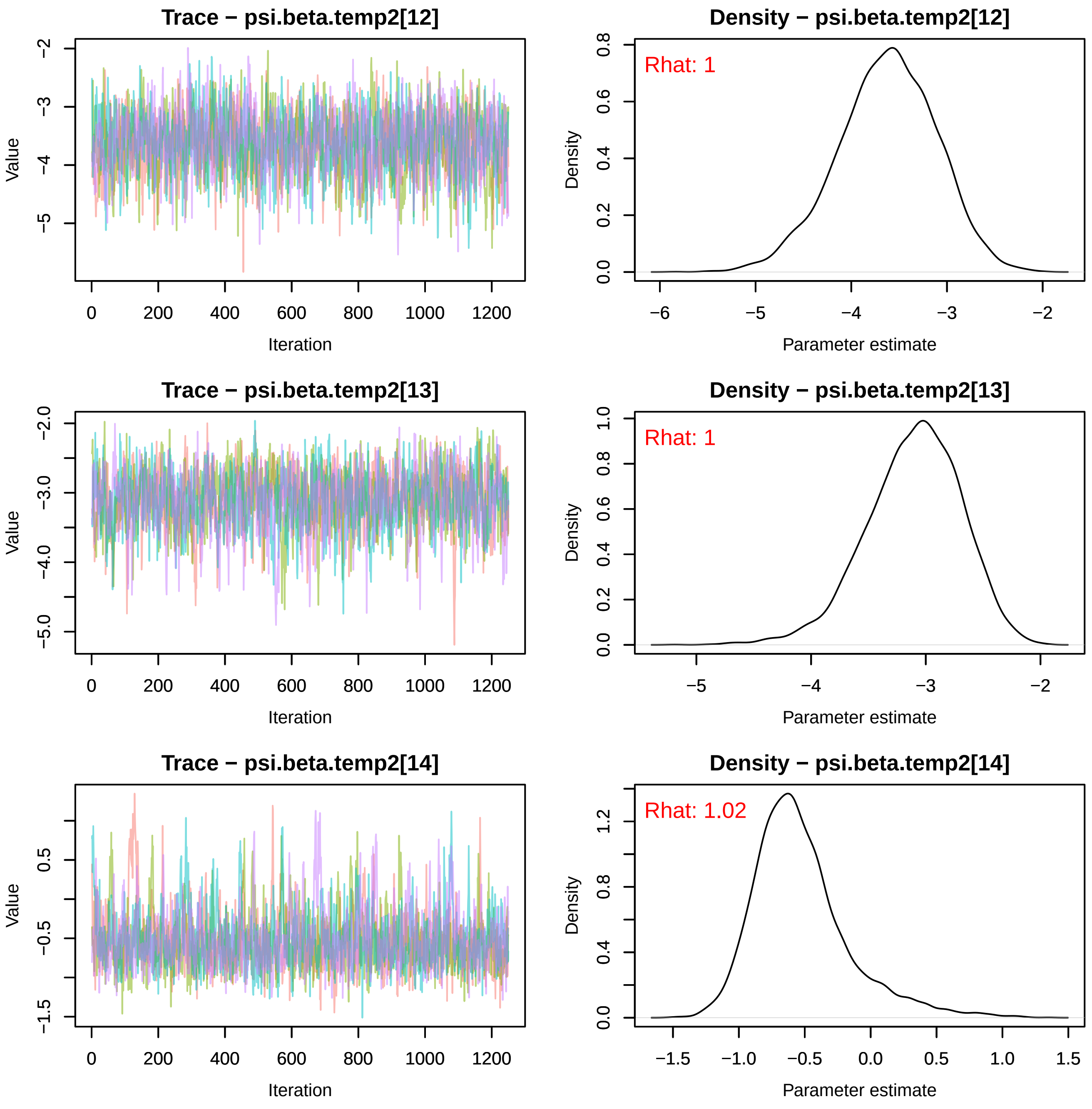
Trace − psi.beta.temp2[12]
Value
Iteration
0 200 400 600 800 1000 1200
−2 −3 −4 −5
Density − psi.beta.temp2[12]
Rhat: 1
Density
Parameter estimate
−6 −5 −4 −3 −2
0.0 0.2 0.4 0.6 0.8
Trace − psi.beta.temp2[13]
Value
Iteration
0 200 400 600 800 1000 1200
−2.0 −3.0 −4.0 −5.0
Density − psi.beta.temp2[13]
Rhat: 1
Density
Parameter estimate
−5 −4 −3 −2
0.0 0.2 0.4 0.6 0.8 1.0
Trace − psi.beta.temp2[14]
Value
Iteration
0 200 400 600 800 1000 1200
0.5 −0.5 −1.5
Density − psi.beta.temp2[14]
Rhat: 1.02
Density
Parameter estimate
−1.5 −1.0 −0.5 0.0 0.5 1.0 1.5
0.0 0.4 0.8 1.2

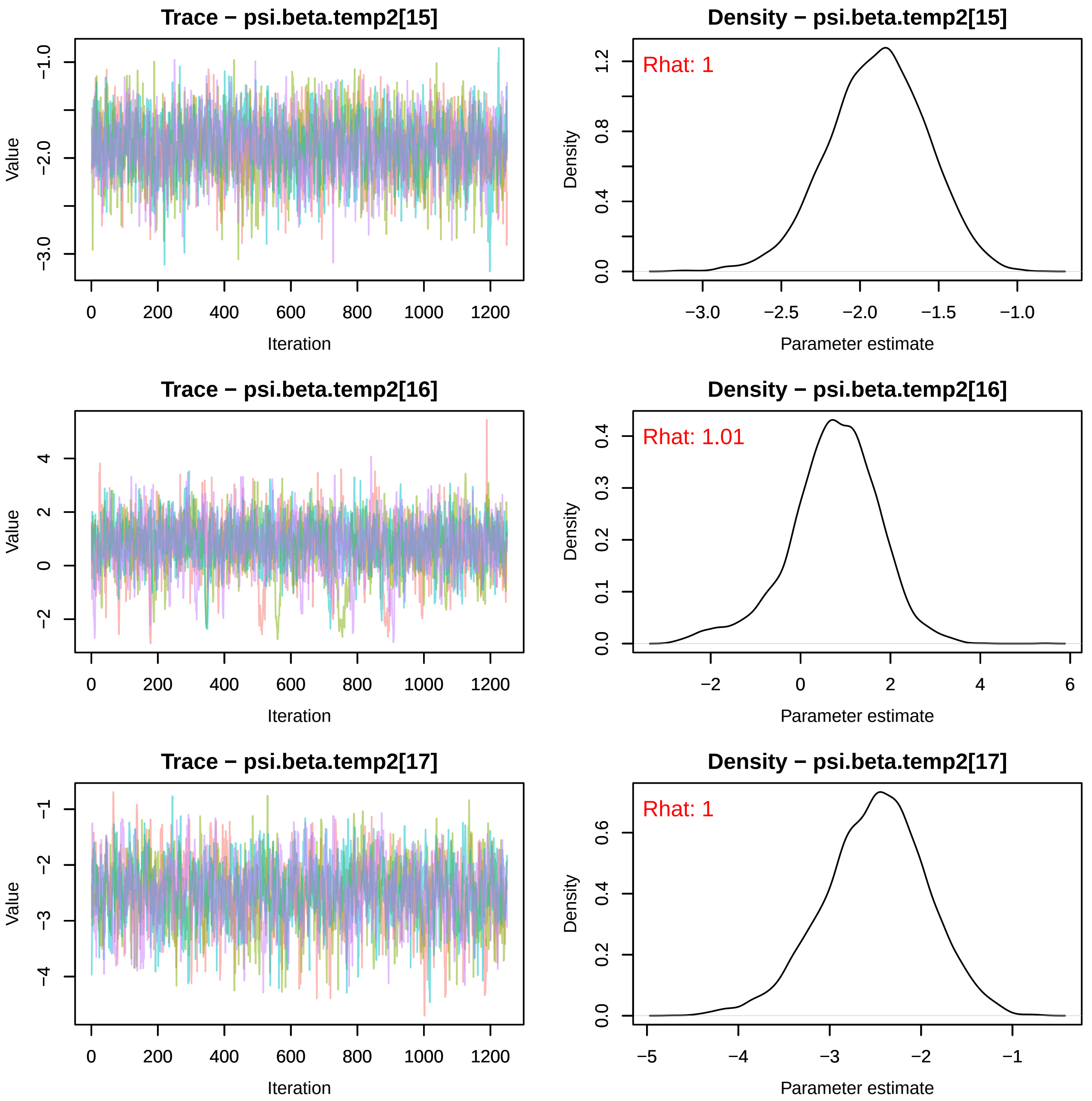
Trace − psi.beta.temp2[15]
Density − psi.beta.temp2[15]
Rhat: 1
Trace − psi.beta.temp2[16]
Density − psi.beta.temp2[16]
Rhat: 1.01
Trace − psi.beta.temp2[17]
Density − psi.beta.temp2[17]
Rhat: 1
Value
Iteration
Density
Parameter estimate

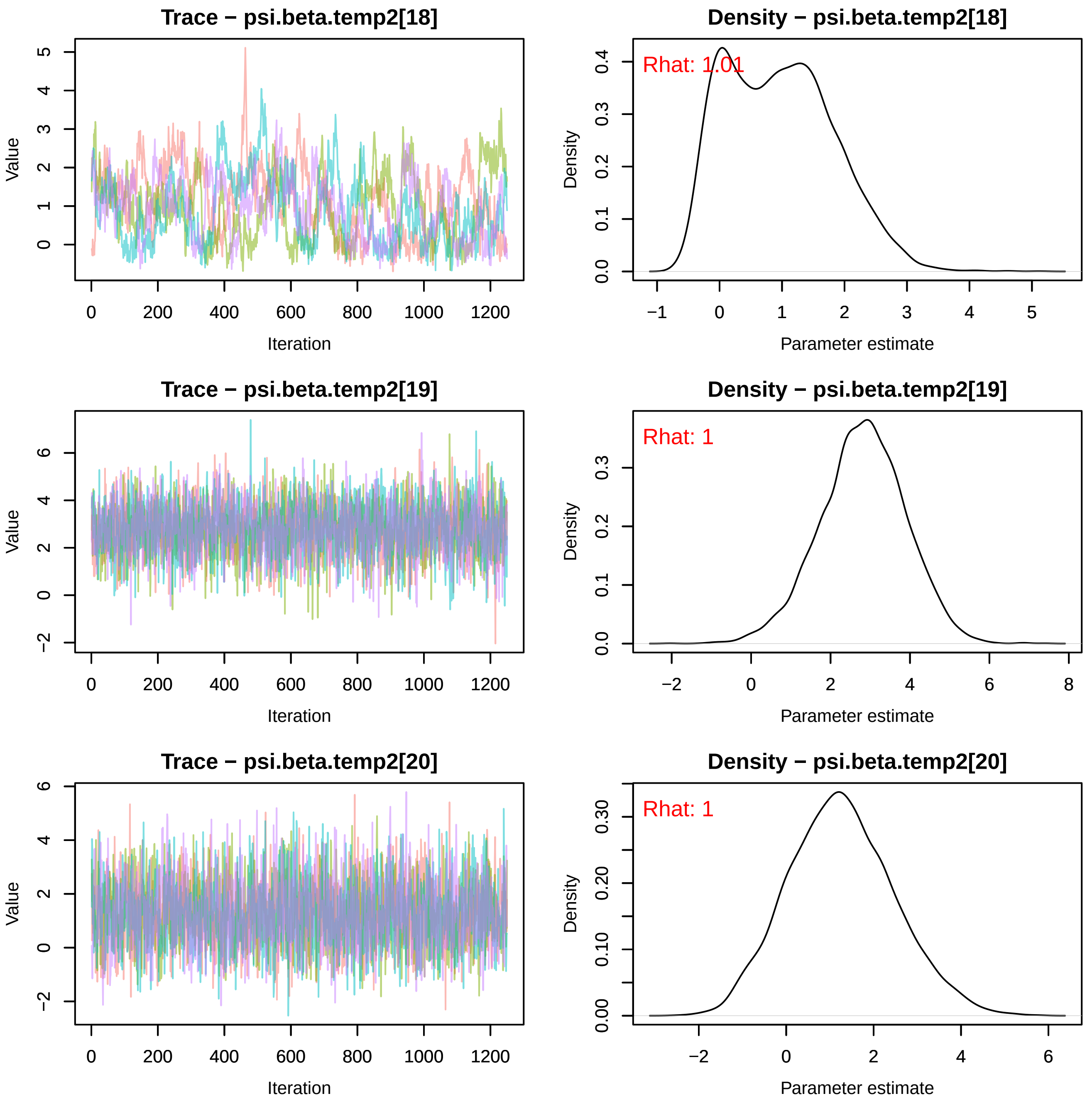
Trace − psi.beta.temp2[18]
Value
Iteration
Density − psi.beta.temp2[18]
Rhat: 1.01
Density
Parameter estimate
Trace − psi.beta.temp2[19]
Value
Iteration
Density − psi.beta.temp2[19]
Rhat: 1
Density
Parameter estimate
Trace − psi.beta.temp2[20]
Value
Iteration
Density − psi.beta.temp2[20]
Rhat: 1
Density
Parameter estimate

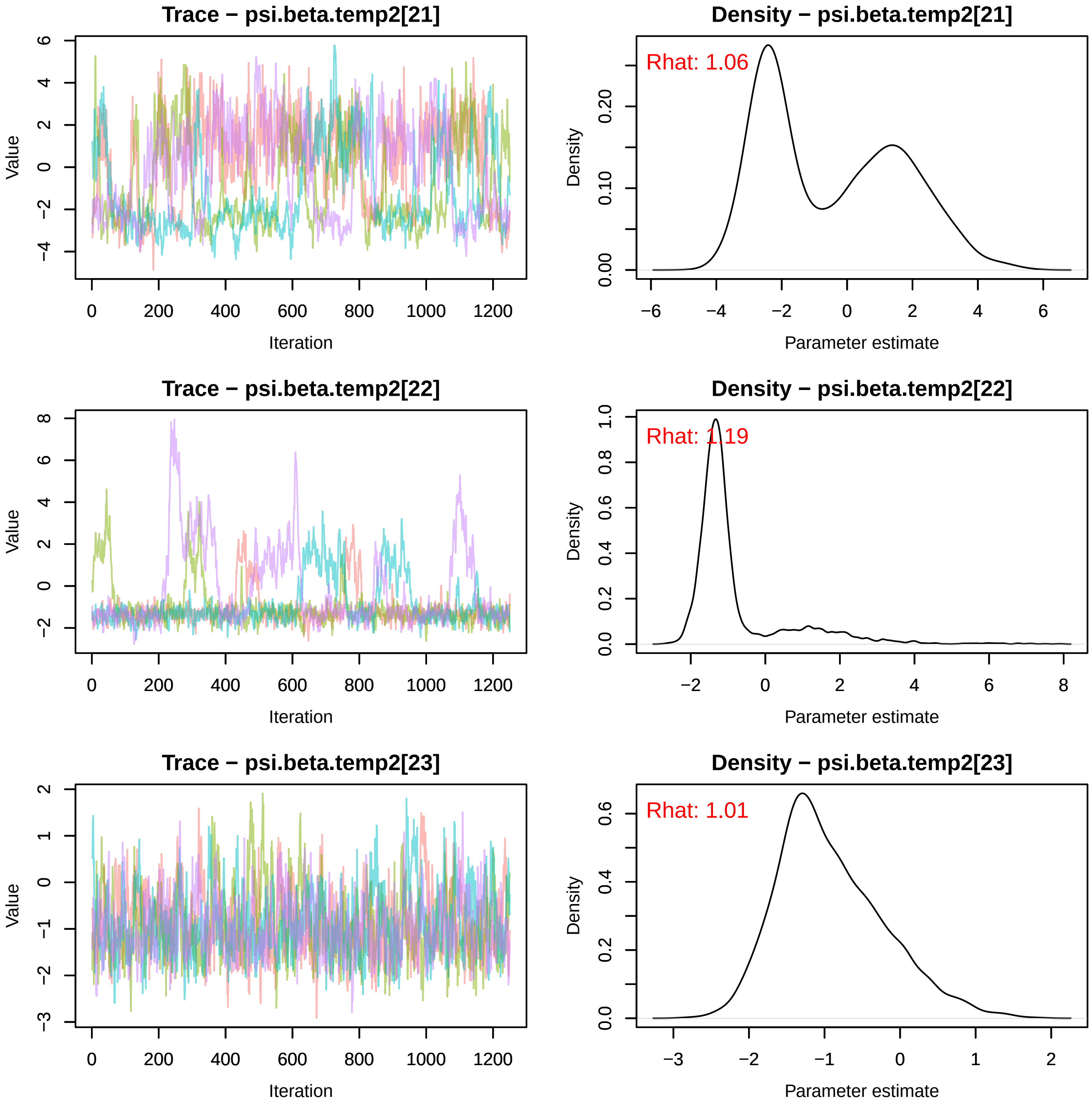
Trace − psi.beta.temp2[21]
Value
Iteration
0
200
400
600
800
1000
1200
−4
−2
0
2
4
6
Density − psi.beta.temp2[21]
Rhat: 1.06
Density
Parameter estimate
−6
−4
−2
0
2
4
6
0.00
0.10
0.20
Trace − psi.beta.temp2[22]
Value
Iteration
0
200
400
600
800
1000
1200
−2
0
2
4
6
8
Density − psi.beta.temp2[22]
Rhat: 1.19
Density
Parameter estimate
−2
0
2
4
6
8
0.0
0.2
0.4
0.6
0.8
1.0
Trace − psi.beta.temp2[23]
Value
Iteration
0
200
400
600
800
1000
1200
−3
−2
−1
0
1
2
Density − psi.beta.temp2[23]
Rhat: 1.01
Density
Parameter estimate
−3
−2
−1
0
1
2
0.0
0.2
0.4
0.6

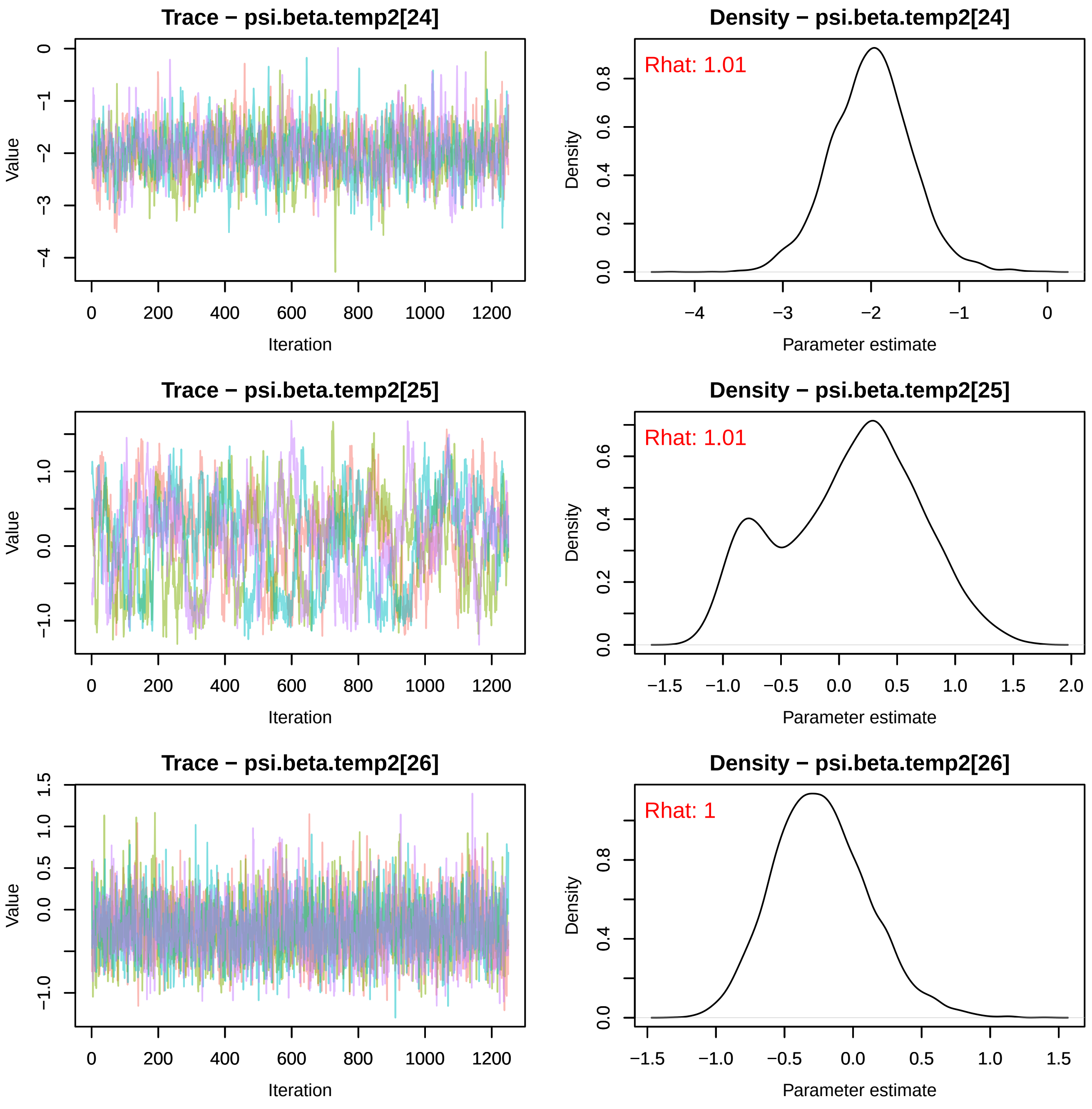
Trace − psi.beta.temp2[24]
Density − psi.beta.temp2[24]
Rhat: 1.01
Trace − psi.beta.temp2[25]
Density − psi.beta.temp2[25]
Rhat: 1.01
Trace − psi.beta.temp2[26]
Density − psi.beta.temp2[26]
Rhat: 1
Value
Iteration
Density
Parameter estimate
0
200
400
600
800
1000
1200
−4
−3
−2
−1
0
−1.5
−1.0
−0.5
0.0
0.5
1.0
1.5
2.0
0.2
0.4
0.6
0.8

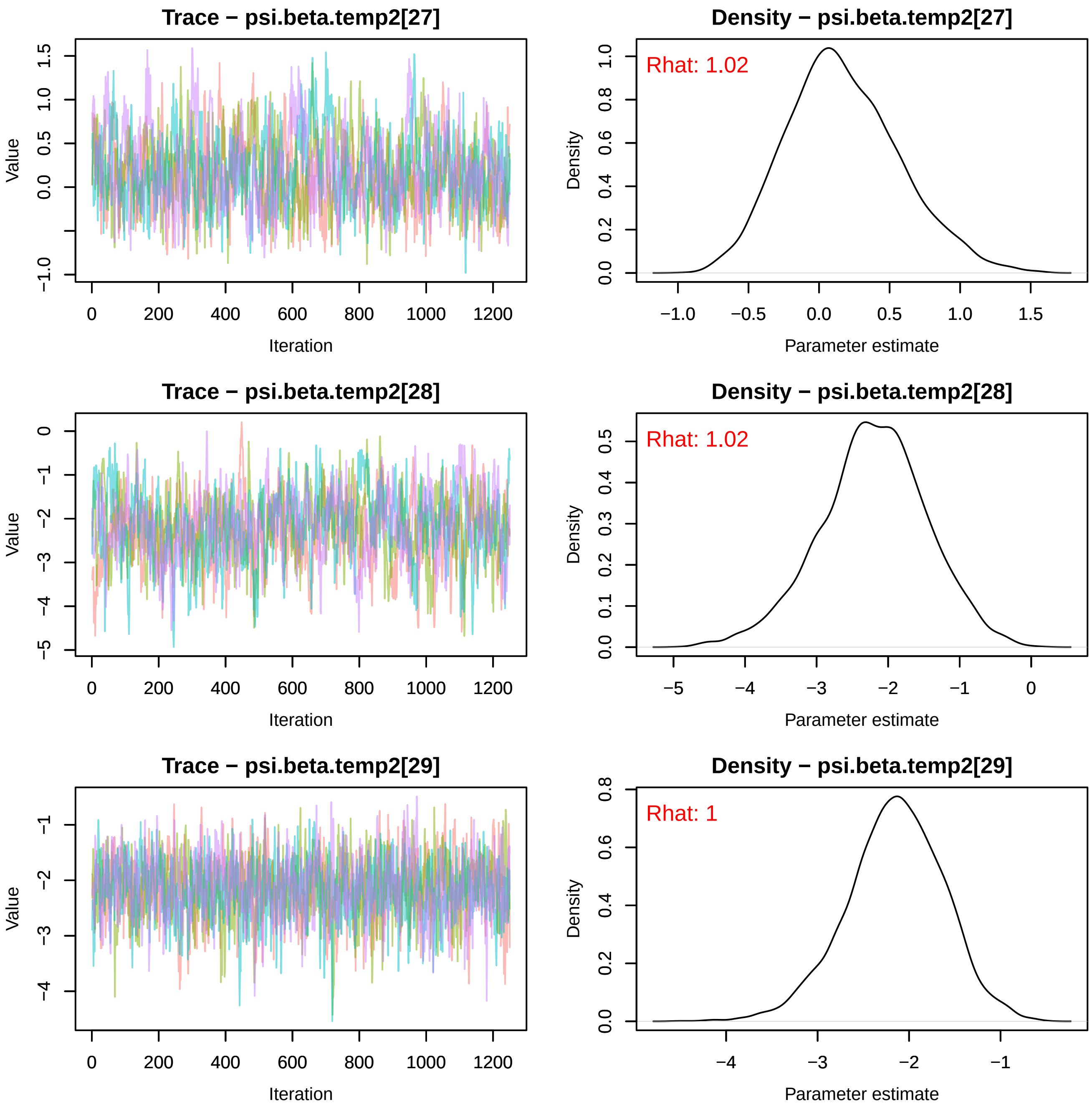

Trace − psi.beta.temp2[27]
Value
Iteration
Density − psi.beta.temp2[27]
Rhat: 1.02
Density
Parameter estimate
Trace − psi.beta.temp2[28]
Value
Iteration
Density − psi.beta.temp2[28]
Rhat: 1.02
Density
Parameter estimate
Trace − psi.beta.temp2[29]
Value
Iteration
Density − psi.beta.temp2[29]
Rhat: 1
Density
Parameter estimate

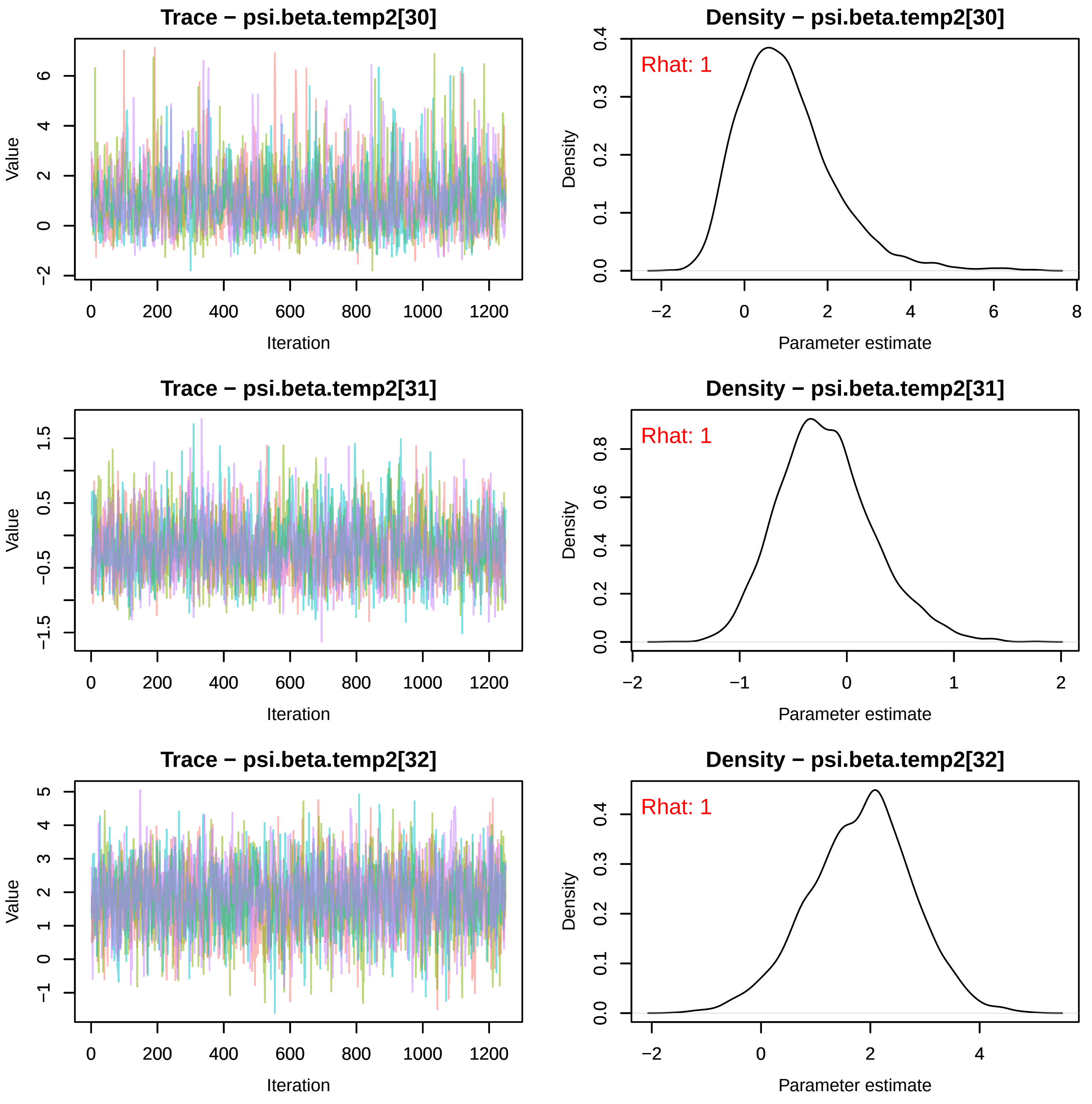
Trace − psi.beta.temp2[30]
Value
Iteration
Density − psi.beta.temp2[30]
Rhat: 1
Density
Parameter estimate
Trace − psi.beta.temp2[31]
Value
Iteration
Density − psi.beta.temp2[31]
Rhat: 1
Density
Parameter estimate
Trace − psi.beta.temp2[32]
Value
Iteration
Density − psi.beta.temp2[32]
Rhat: 1
Density
Parameter estimate

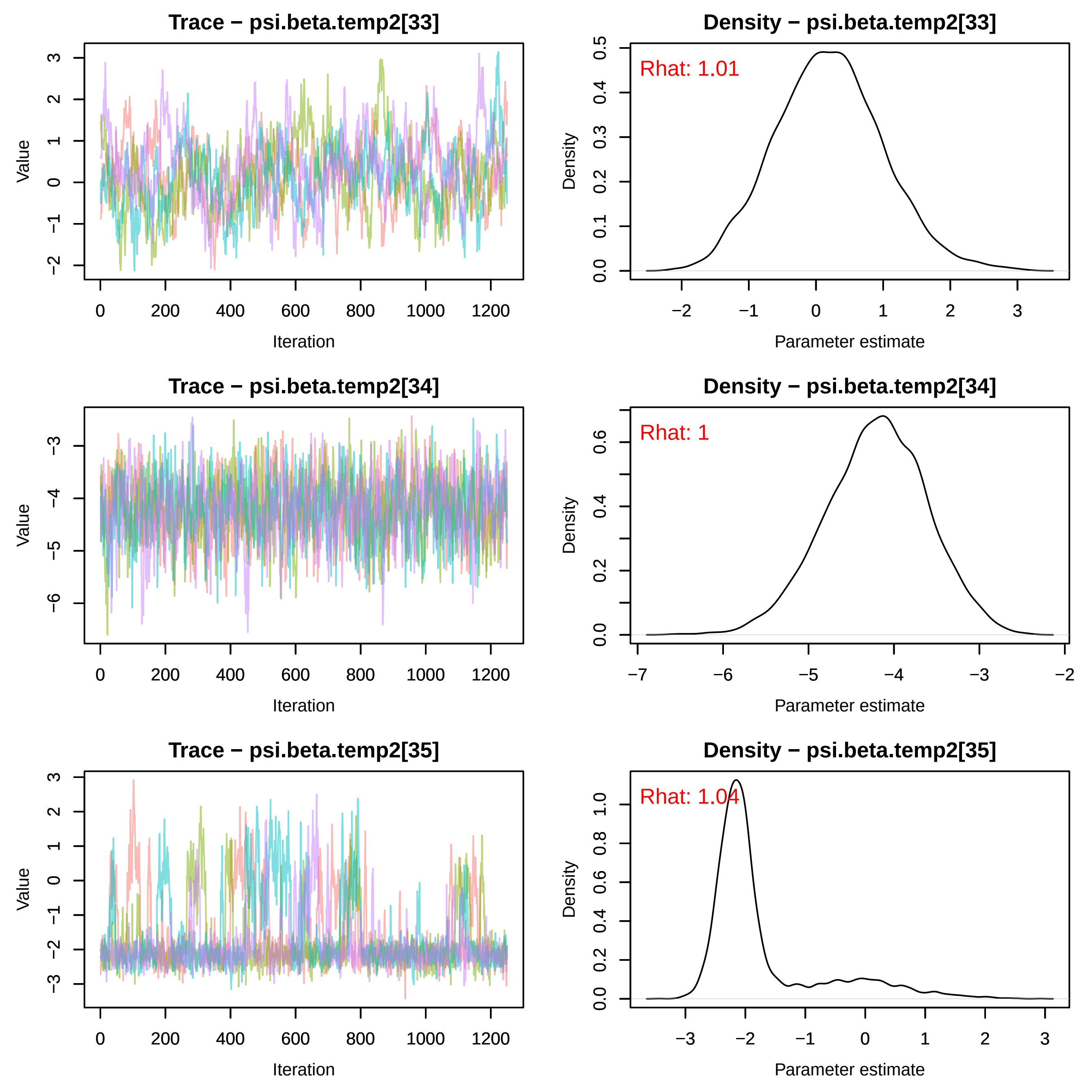
Trace − psi.beta.temp2[33]
Value
Iteration
Density − psi.beta.temp2[33]
Rhat: 1.01
Density
Parameter estimate
Trace − psi.beta.temp2[34]
Value
Iteration
Density − psi.beta.temp2[34]
Rhat: 1
Density
Parameter estimate
Trace − psi.beta.temp2[35]
Value
Iteration
Density − psi.beta.temp2[35]
Rhat: 1.04
Density
Parameter estimate

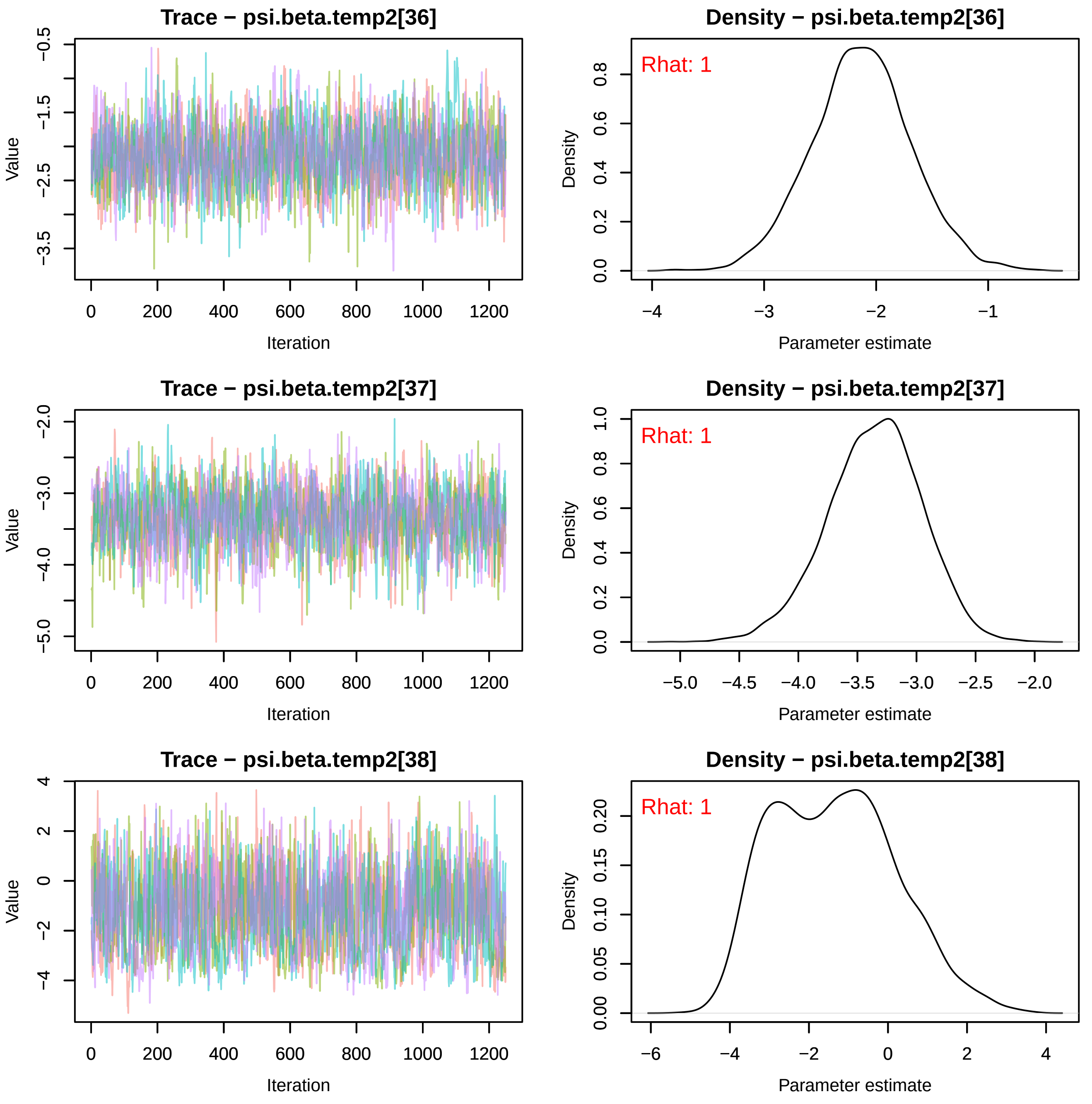
Trace − psi.beta.temp2[36]
Value
Iteration
Density − psi.beta.temp2[36]
Rhat: 1
Density
Parameter estimate
Trace − psi.beta.temp2[37]
Value
Iteration
Density − psi.beta.temp2[37]
Rhat: 1
Density
Parameter estimate
Trace − psi.beta.temp2[38]
Value
Iteration
Density − psi.beta.temp2[38]
Rhat: 1
Density
Parameter estimate

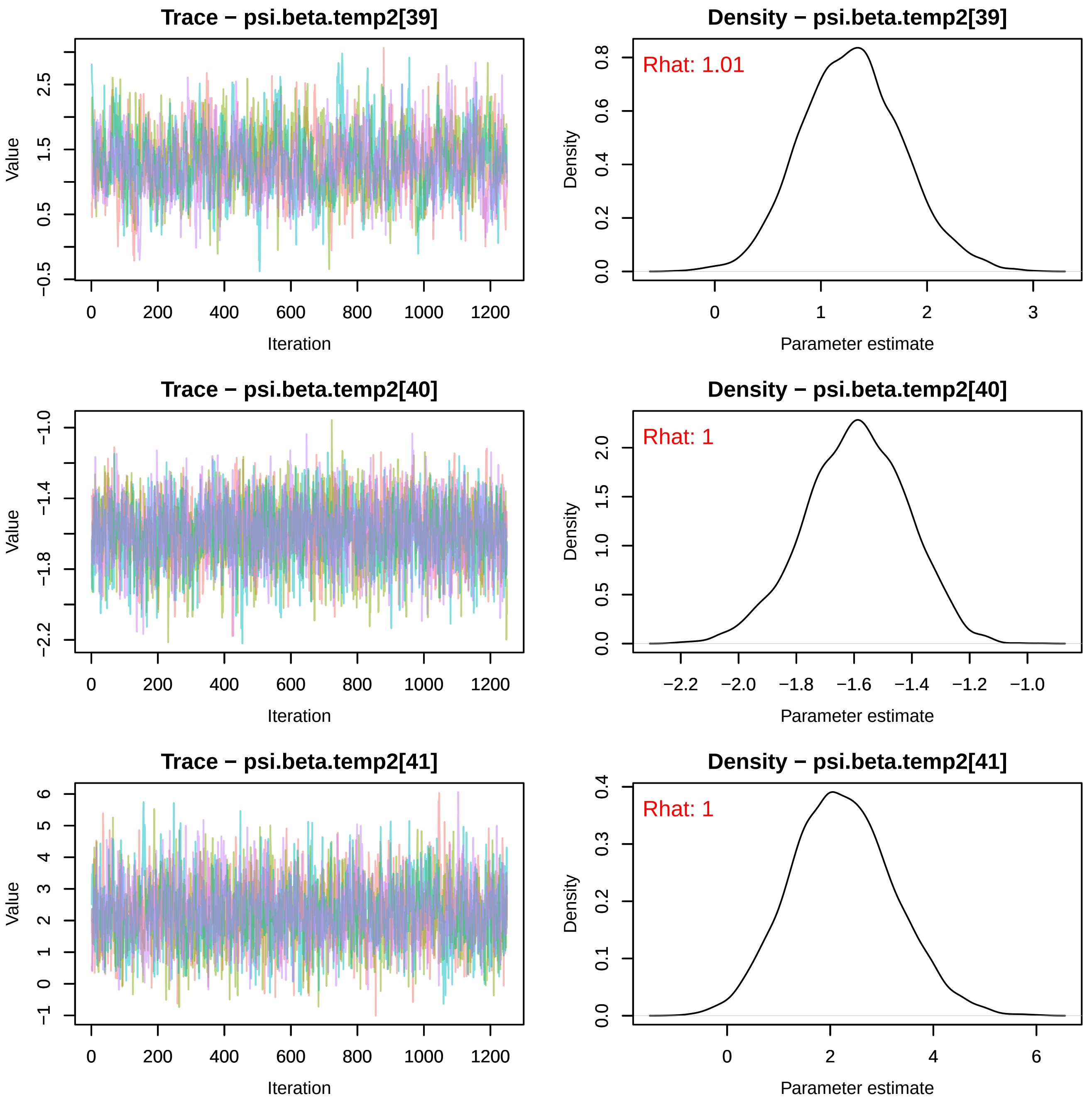
Trace − psi.beta.temp2[39]
Value
Iteration
Density − psi.beta.temp2[39]
Rhat: 1.01
Density
Parameter estimate
Trace − psi.beta.temp2[40]
Value
Iteration
Density − psi.beta.temp2[40]
Rhat: 1
Density
Parameter estimate
Trace − psi.beta.temp2[41]
Value
Iteration
Density − psi.beta.temp2[41]
Rhat: 1
Density
Parameter estimate

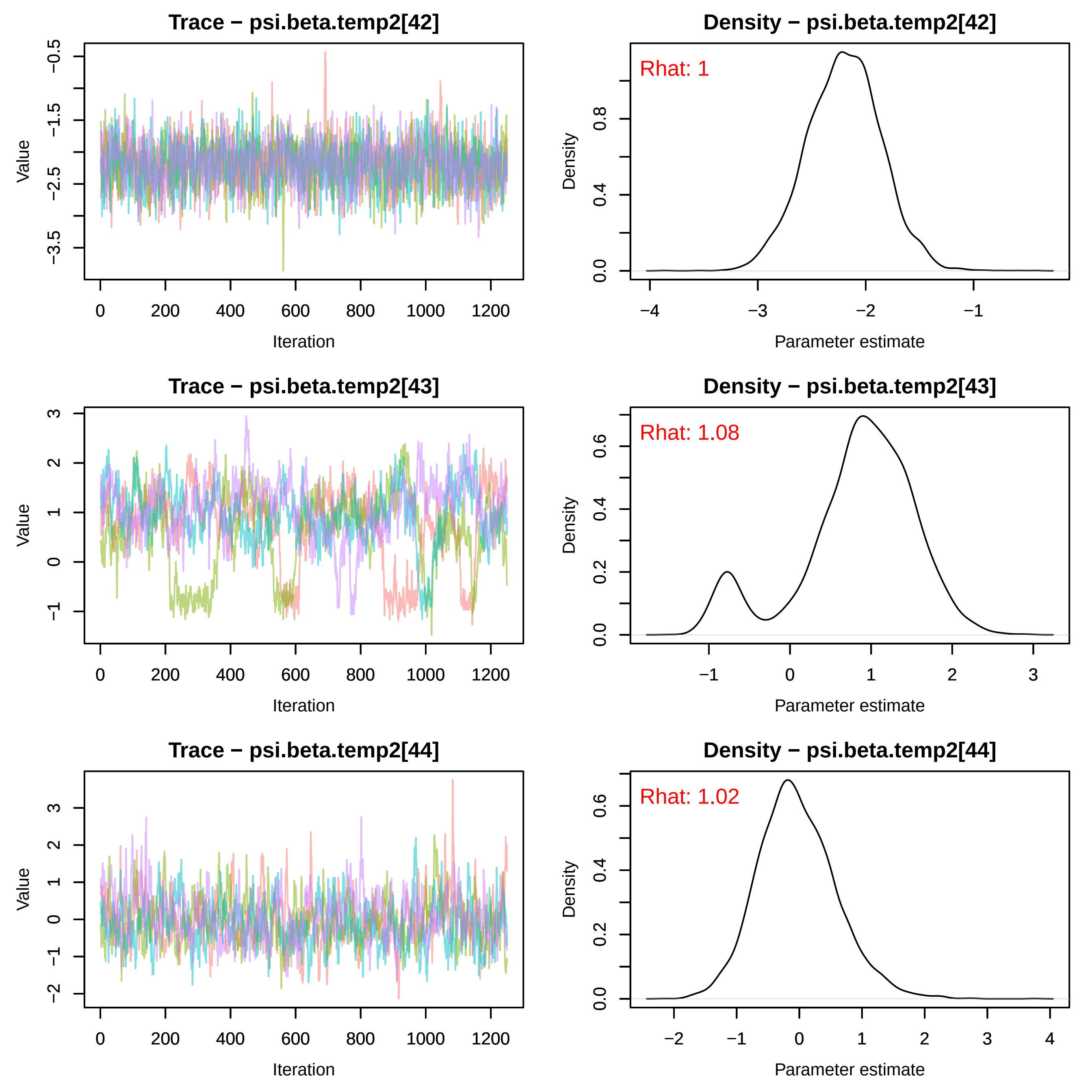
Trace − psi.beta.temp2[42]
Value
Iteration
Density − psi.beta.temp2[42]
Rhat: 1
Density
Parameter estimate
Trace − psi.beta.temp2[43]
Value
Iteration
Density − psi.beta.temp2[43]
Rhat: 1.08
Density
Parameter estimate
Trace − psi.beta.temp2[44]
Value
Iteration
Density − psi.beta.temp2[44]
Rhat: 1.02
Density
Parameter estimate

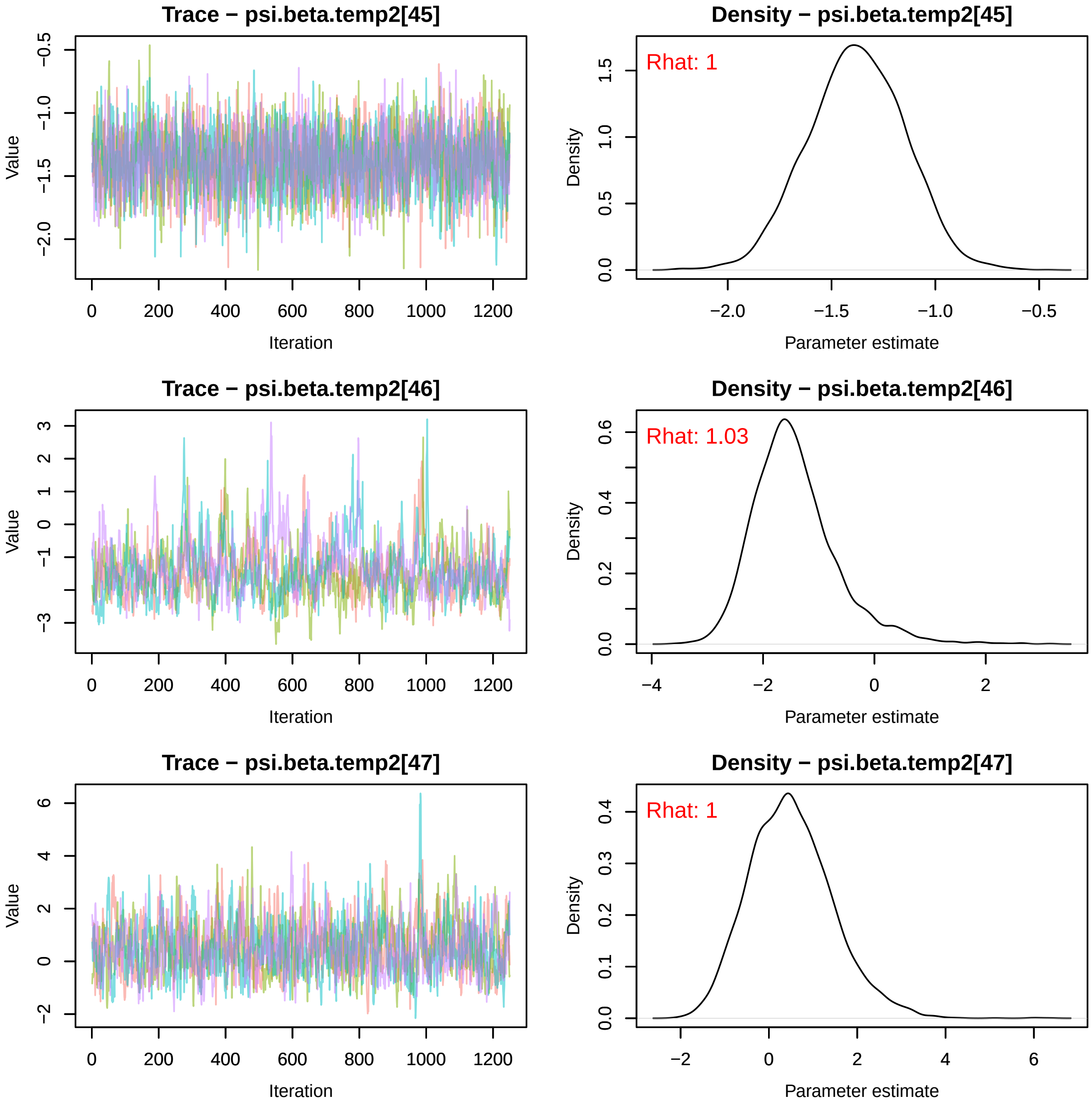
Trace − psi.beta.temp2[45]
Density − psi.beta.temp2[45]
Rhat: 1
Trace − psi.beta.temp2[46]
Density − psi.beta.temp2[46]
Rhat: 1.03
Trace − psi.beta.temp2[47]
Density − psi.beta.temp2[47]
Rhat: 1
Value
Iteration
Density
Parameter estimate

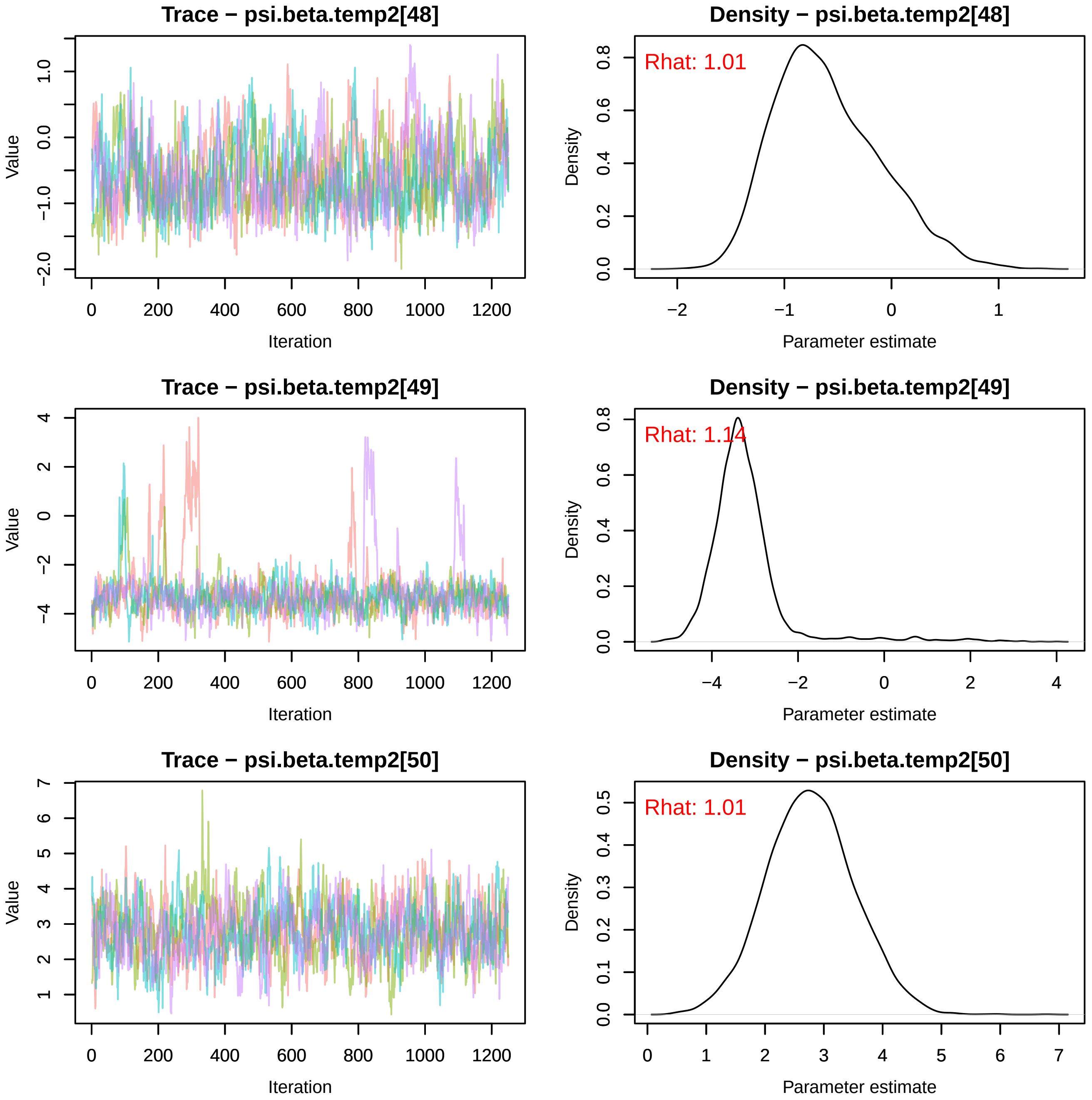
Trace − psi.beta.temp2[48]
Value
Iteration
Density − psi.beta.temp2[48]
Rhat: 1.01
Density
Parameter estimate
Trace − psi.beta.temp2[49]
Value
Iteration
Density − psi.beta.temp2[49]
Rhat: 1.14
Density
Parameter estimate
Trace − psi.beta.temp2[50]
Value
Iteration
Density − psi.beta.temp2[50]
Rhat: 1.01
Density
Parameter estimate

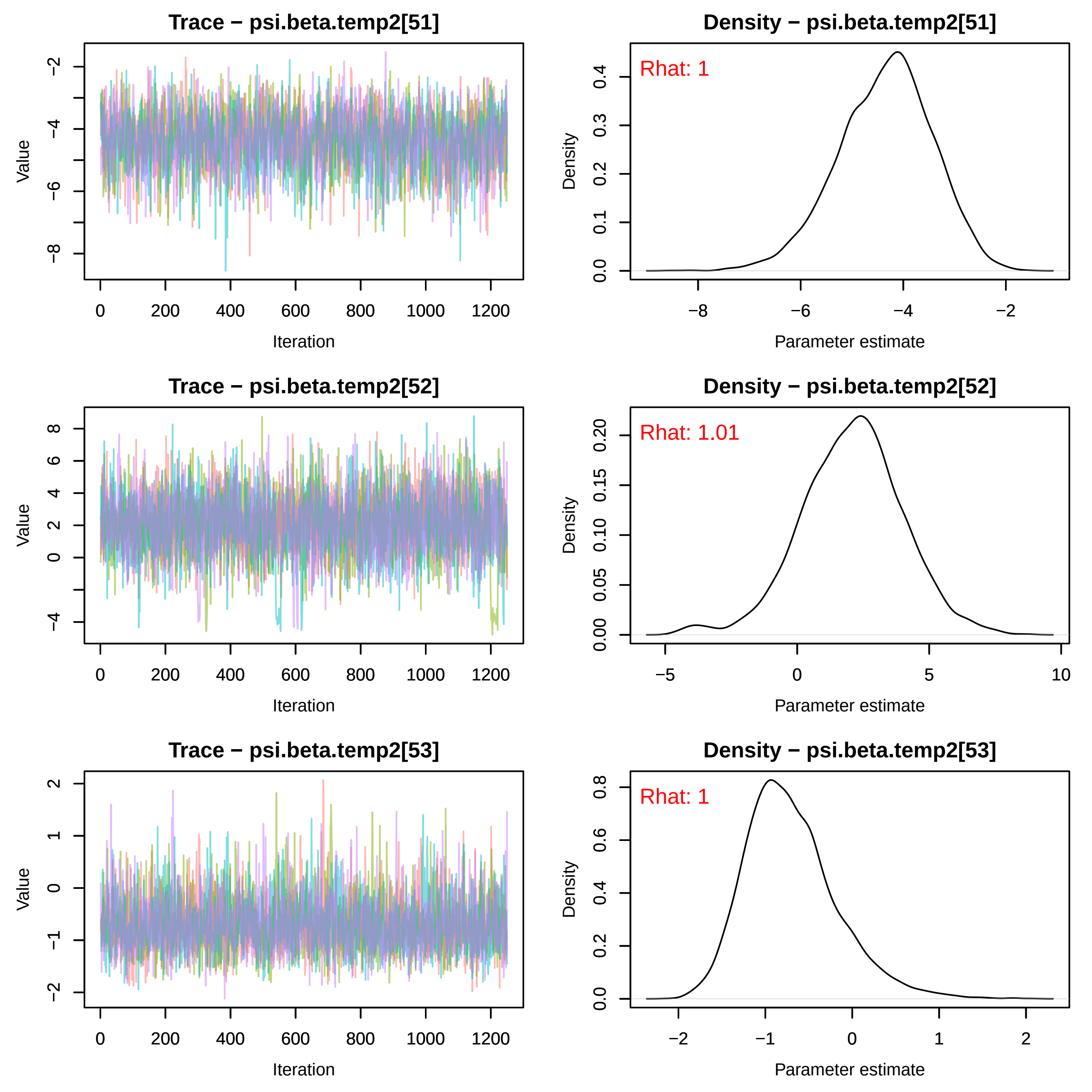
Trace − psi.beta.temp2[51]
Value
Iteration
Density − psi.beta.temp2[51]
Rhat: 1
Density
Parameter estimate
Trace − psi.beta.temp2[52]
Value
Iteration
Density − psi.beta.temp2[52]
Rhat: 1.01
Density
Parameter estimate
Trace − psi.beta.temp2[53]
Value
Iteration
Density − psi.beta.temp2[53]
Rhat: 1
Density
Parameter estimate

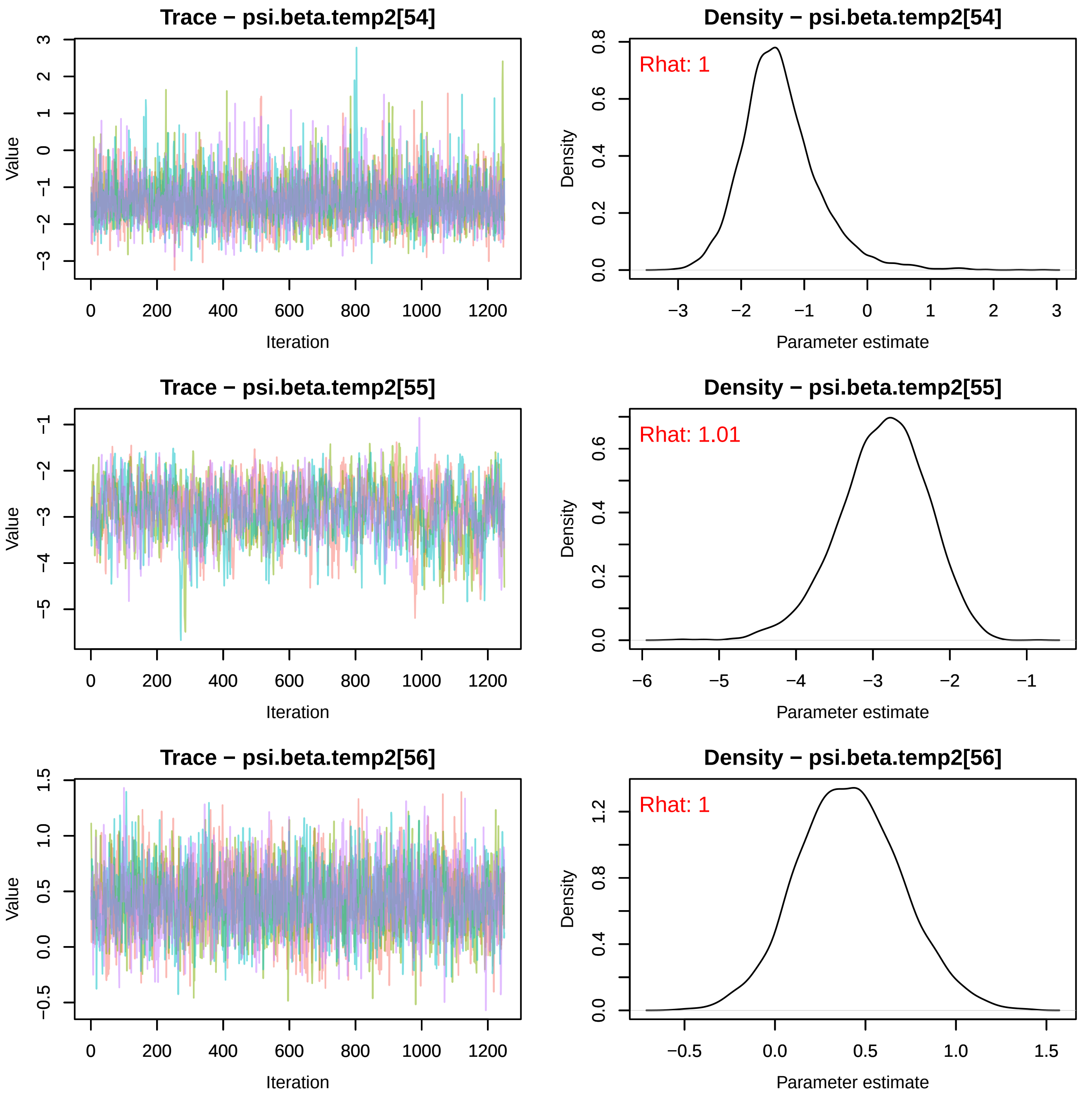
Trace − psi.beta.temp2[54]
Value
Iteration
Density − psi.beta.temp2[54]
Rhat: 1
Density
Parameter estimate
Trace − psi.beta.temp2[55]
Value
Iteration
Density − psi.beta.temp2[55]
Rhat: 1.01
Density
Parameter estimate
Trace − psi.beta.temp2[56]
Value
Iteration
Density − psi.beta.temp2[56]
Rhat: 1
Density
Parameter estimate

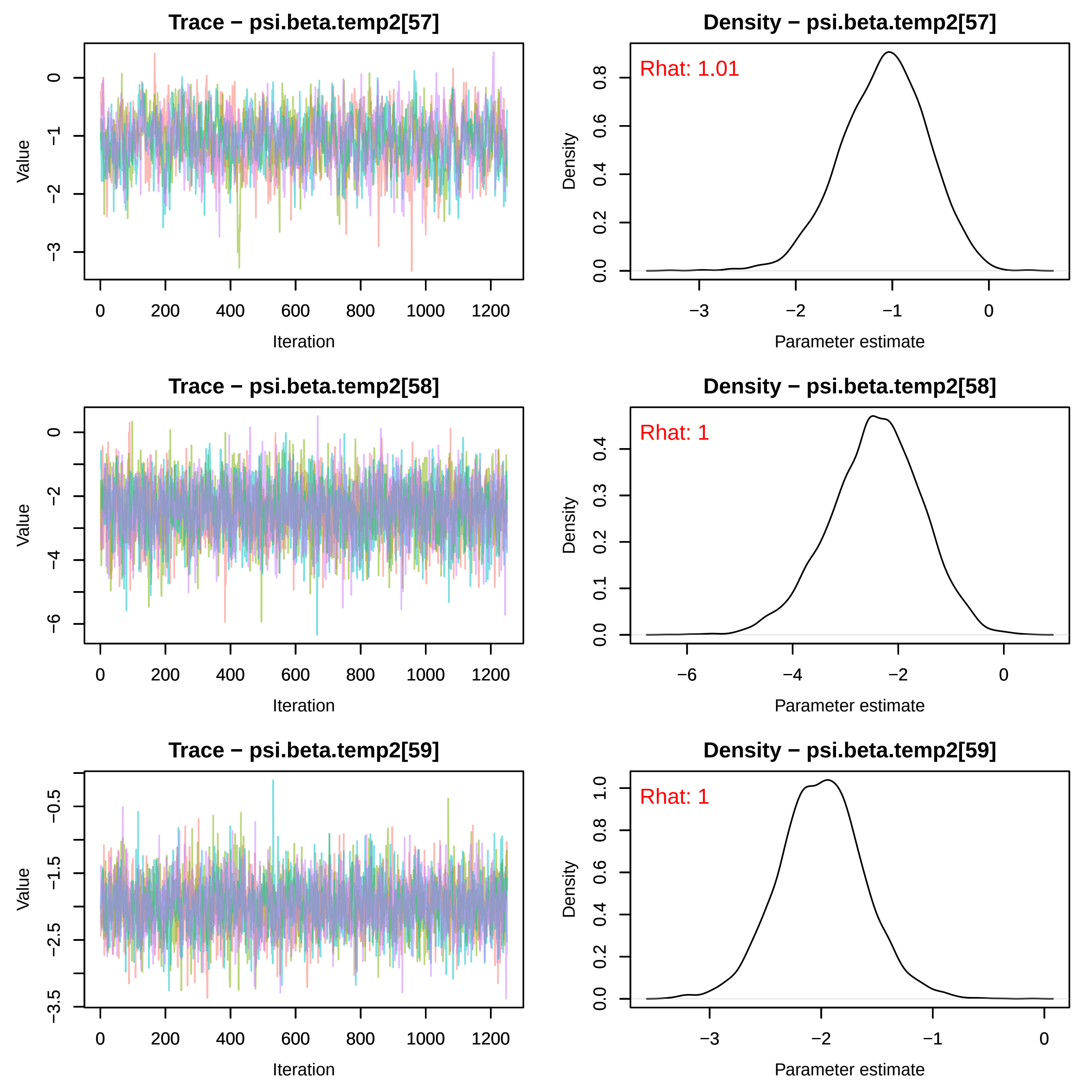

Trace − psi.beta.temp2[57]
Value
Iteration
Density − psi.beta.temp2[57]
Rhat: 1.01
Density
Parameter estimate
Trace − psi.beta.temp2[58]
Value
Iteration
Density − psi.beta.temp2[58]
Rhat: 1
Density
Parameter estimate
Trace − psi.beta.temp2[59]
Value
Iteration
Density − psi.beta.temp2[59]
Rhat: 1
Density
Parameter estimate

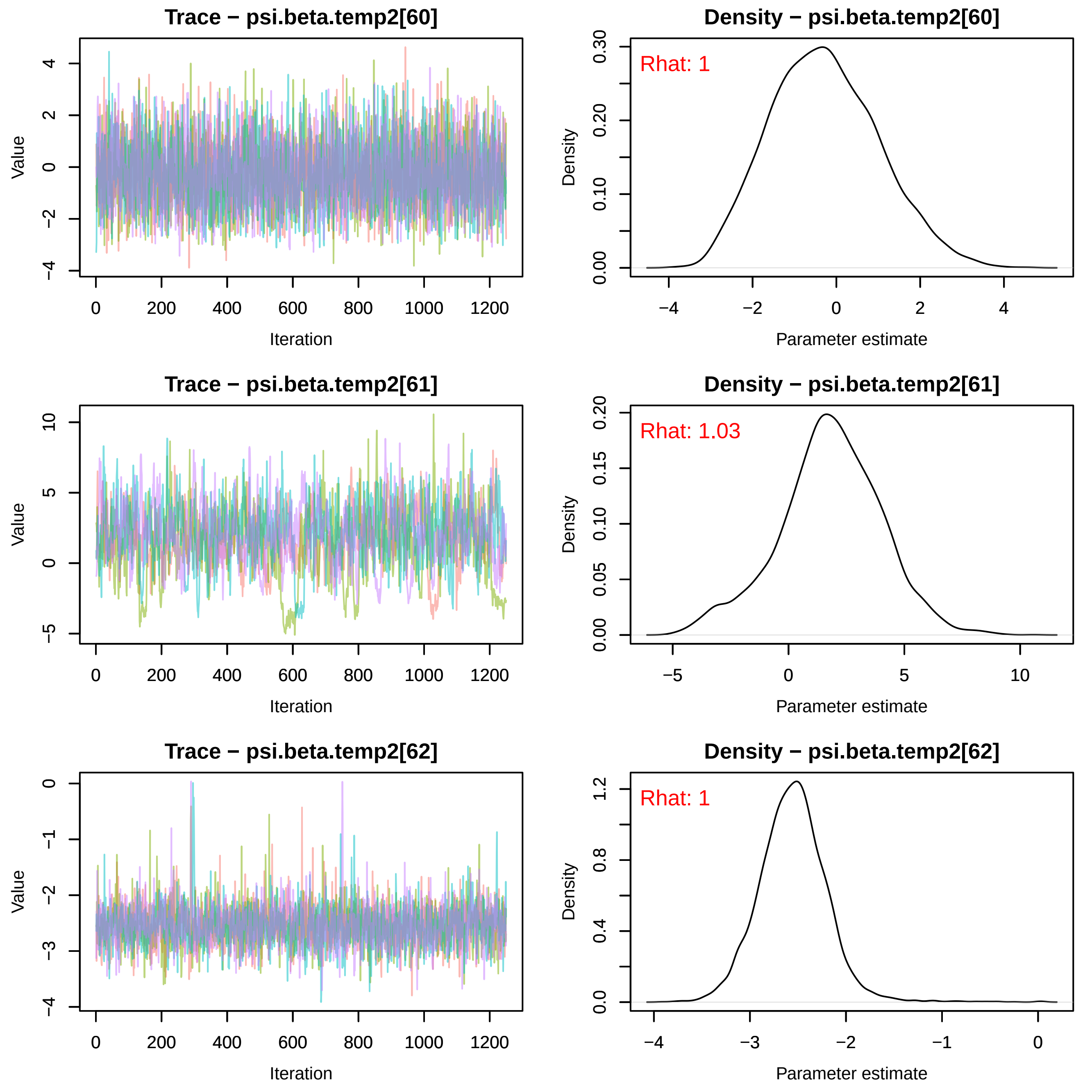
Trace − psi.beta.temp2[60]
Value
Iteration
Density − psi.beta.temp2[60]
Rhat: 1
Density
Parameter estimate
Trace − psi.beta.temp2[61]
Value
Iteration
Density − psi.beta.temp2[61]
Rhat: 1.03
Density
Parameter estimate
Trace − psi.beta.temp2[62]
Value
Iteration
Density − psi.beta.temp2[62]
Rhat: 1
Density
Parameter estimate

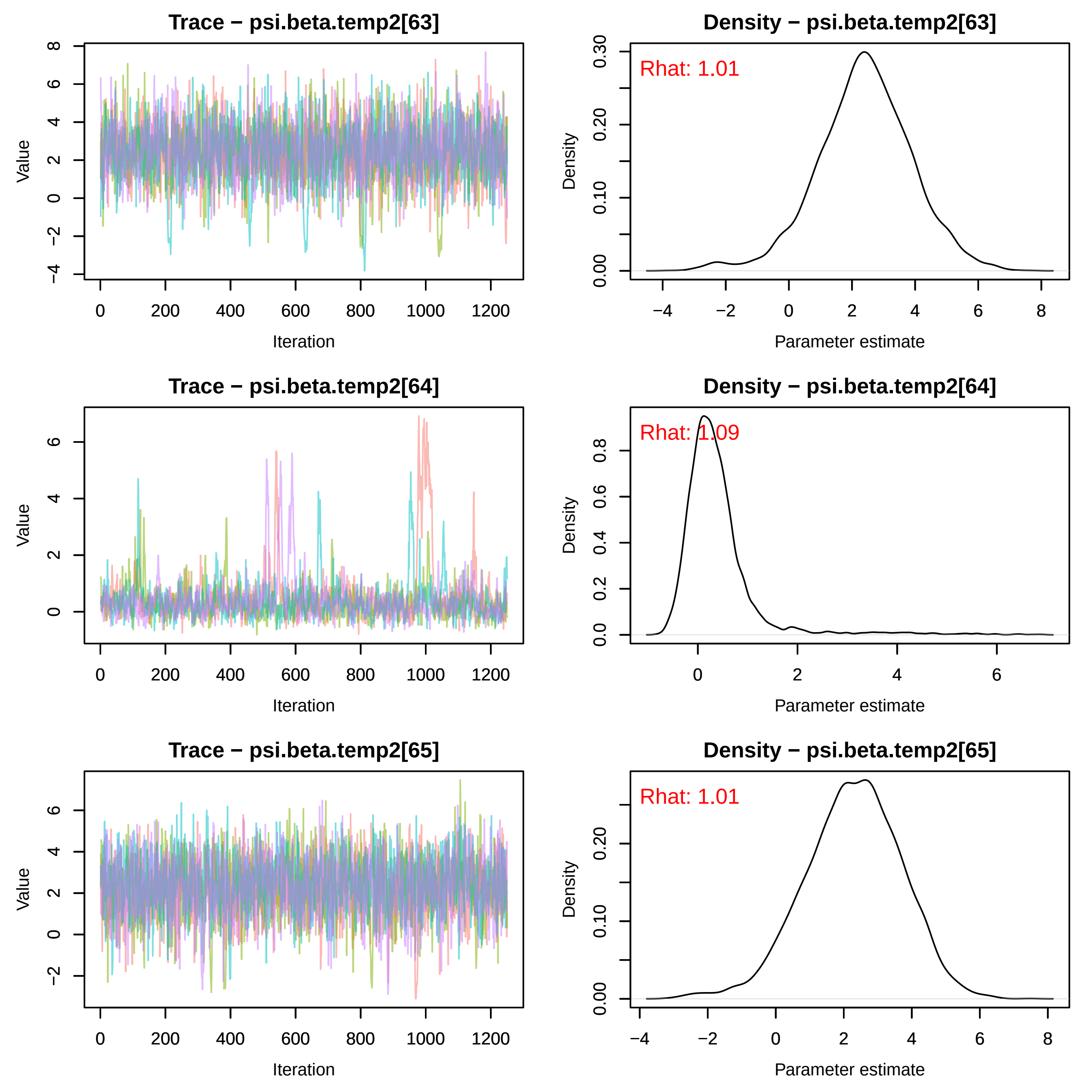
Trace − psi.beta.temp2[63]
Density − psi.beta.temp2[63]
Rhat: 1.01
Trace − psi.beta.temp2[64]
Density − psi.beta.temp2[64]
Rhat: 1.09
Trace − psi.beta.temp2[65]
Density − psi.beta.temp2[65]
Rhat: 1.01
Value
Iteration
Density
Parameter estimate

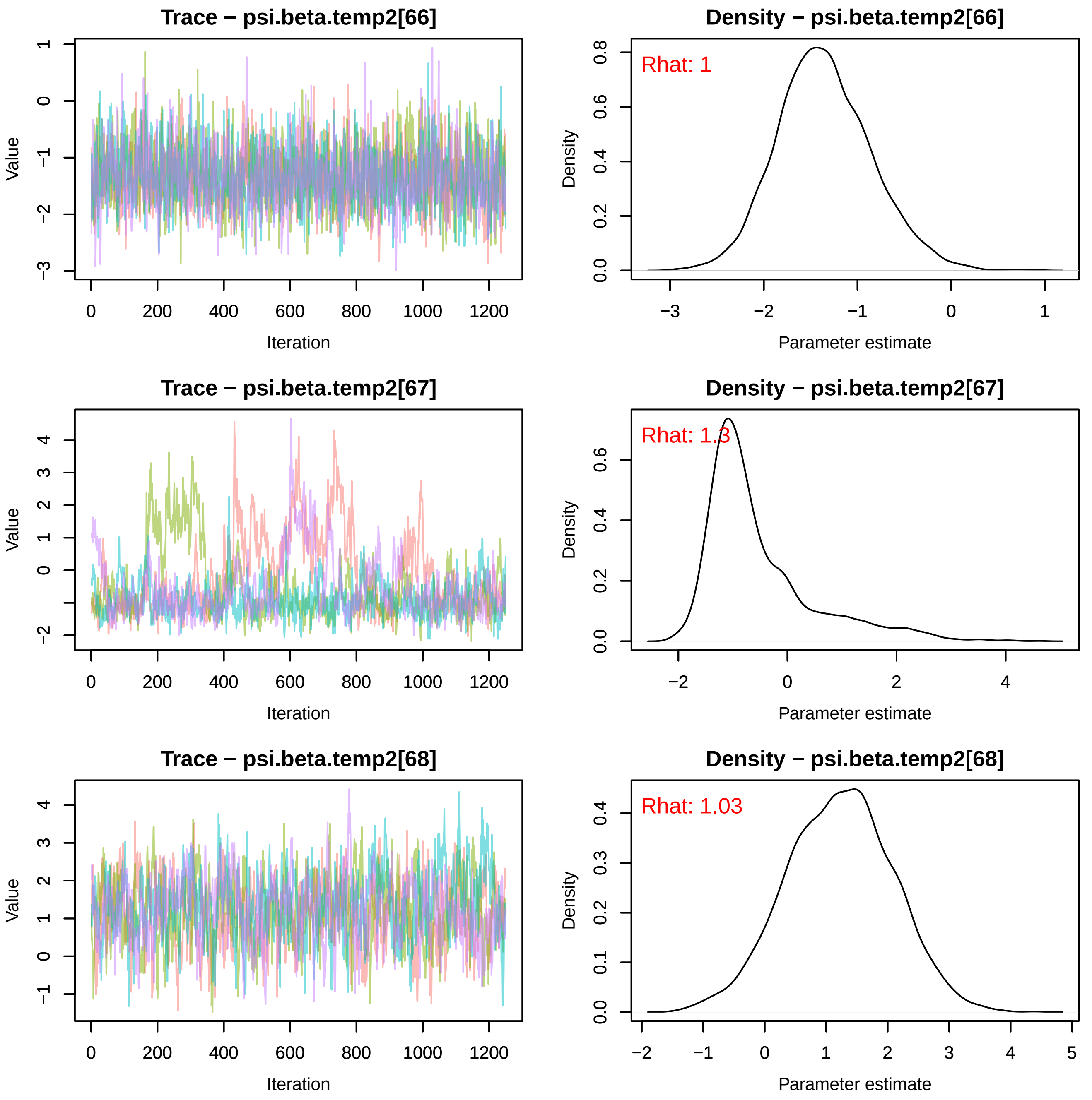
Trace − psi.beta.temp2[66]
Density − psi.beta.temp2[66]
Rhat: 1
Trace − psi.beta.temp2[67]
Density − psi.beta.temp2[67]
Rhat: 1.3
Trace − psi.beta.temp2[68]
Density − psi.beta.temp2[68]
Rhat: 1.03
Value
Iteration
Density
Parameter estimate

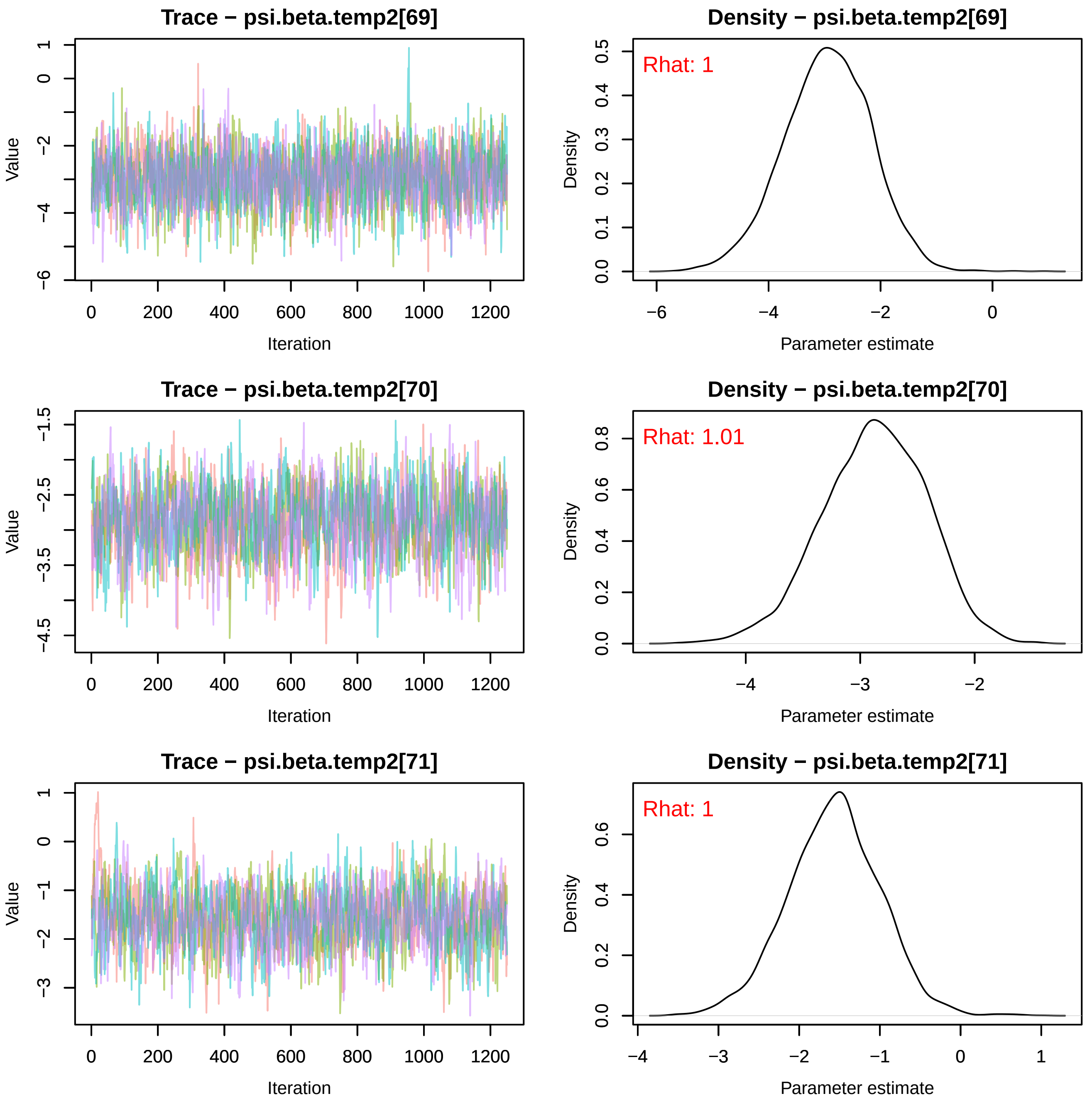
Trace − psi.beta.temp2[69]
Value
Iteration
Density − psi.beta.temp2[69]
Rhat: 1
Density
Parameter estimate
Trace − psi.beta.temp2[70]
Value
Iteration
Density − psi.beta.temp2[70]
Rhat: 1.01
Density
Parameter estimate
Trace − psi.beta.temp2[71]
Value
Iteration
Density − psi.beta.temp2[71]
Rhat: 1
Density
Parameter estimate

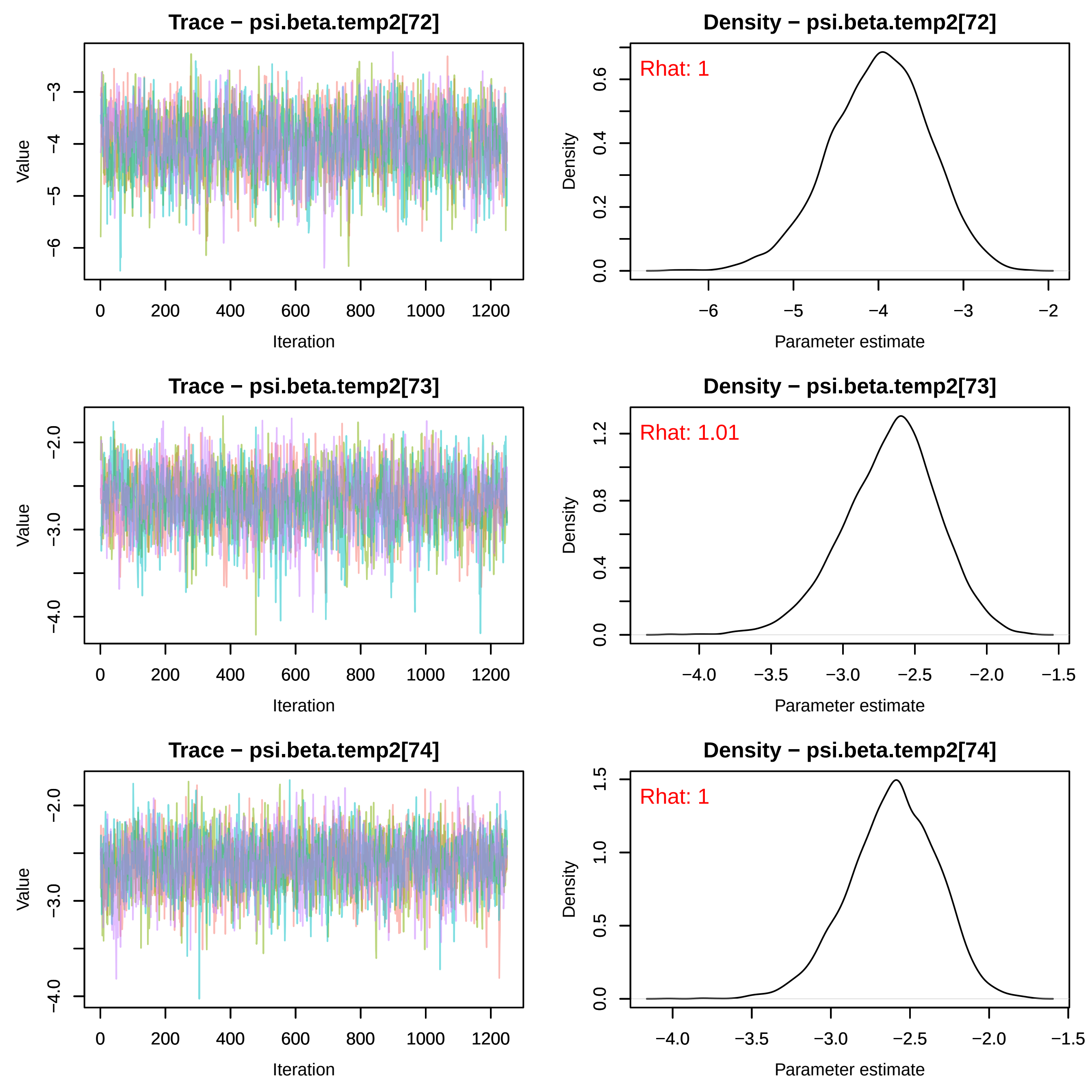

Trace − psi.beta.temp2[72]
Density − psi.beta.temp2[72]
Rhat: 1
Trace − psi.beta.temp2[73]
Density − psi.beta.temp2[73]
Rhat: 1.01
Trace − psi.beta.temp2[74]
Density − psi.beta.temp2[74]
Rhat: 1
Value
Iteration
Density
Parameter estimate

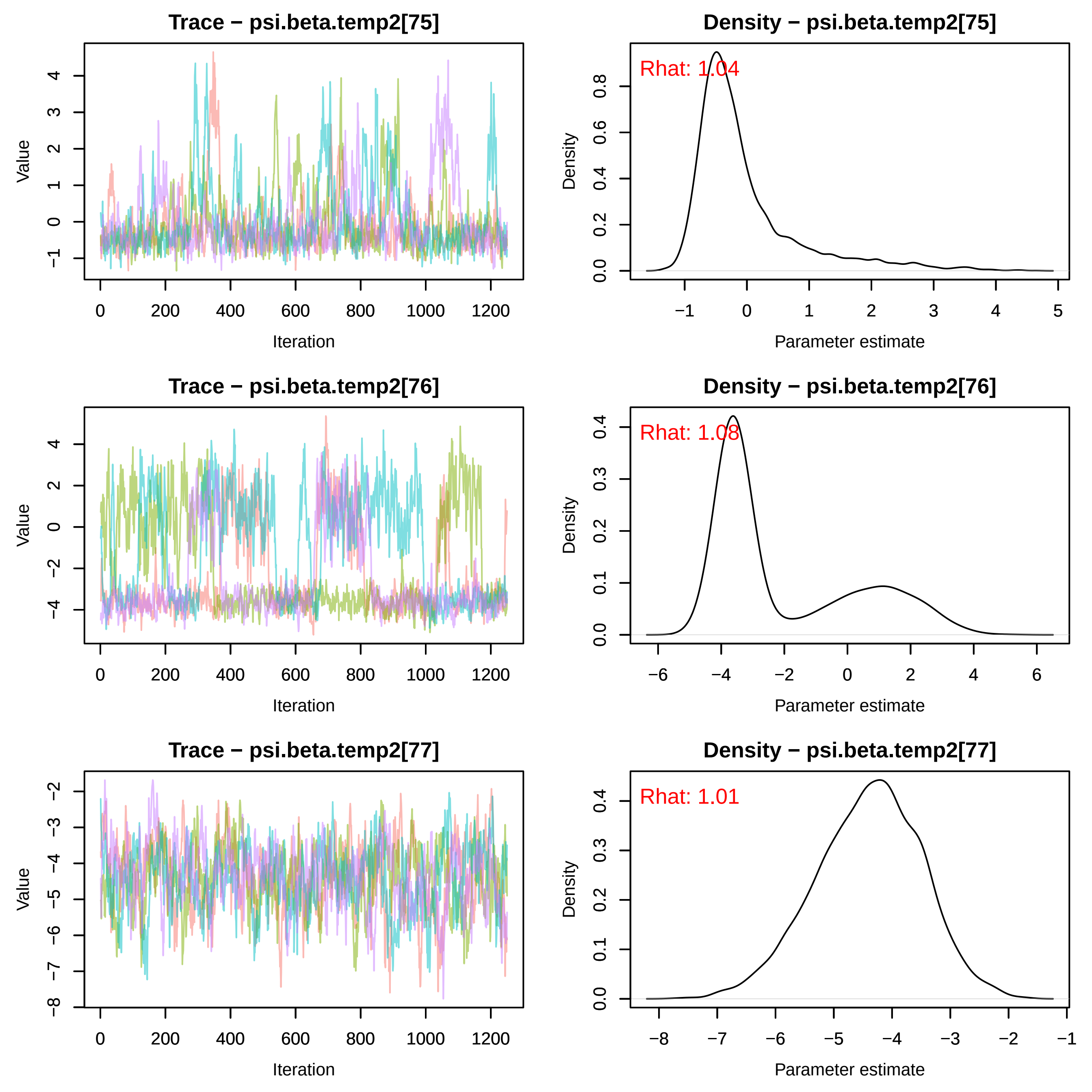

Trace − psi.beta.temp2[75]
Density − psi.beta.temp2[75]
Rhat: 1.04
Trace − psi.beta.temp2[76]
Density − psi.beta.temp2[76]
Rhat: 1.08
Trace − psi.beta.temp2[77]
Density − psi.beta.temp2[77]
Rhat: 1.01
Value
Iteration
Density
Parameter estimate

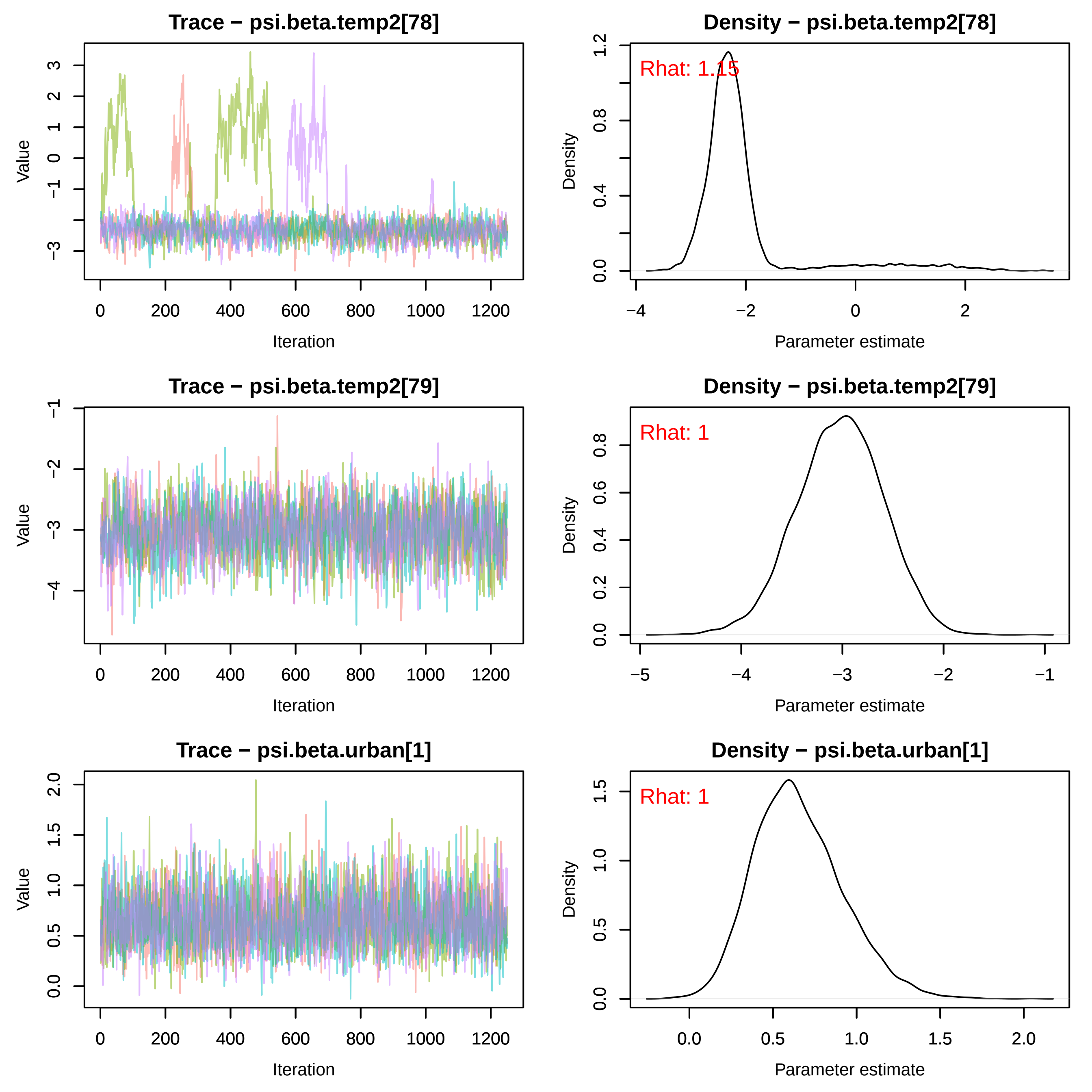

Trace − psi.beta.temp2[78]
Value
Iteration
0 200 400 600 800 1000 1200
3 2 1 0 −1 −3
Density − psi.beta.temp2[78]
Rhat: 1.15
Density
Parameter estimate
−4 −2 0 2
0.0 0.4 0.8 1.2
Trace − psi.beta.temp2[79]
Value
Iteration
0 200 400 600 800 1000 1200
−1 −2 −3 −4
Density − psi.beta.temp2[79]
Rhat: 1
Density
Parameter estimate
−5 −4 −3 −2 −1
0.0 0.2 0.4 0.6 0.8
Trace − psi.beta.urban[1]
Value
Iteration
0 200 400 600 800 1000 1200
0.0 0.5 1.0 1.5 2.0
Density − psi.beta.urban[1]
Rhat: 1
Density
Parameter estimate
0.0 0.5 1.0 1.5 2.0
0.0 0.5 1.0 1.5

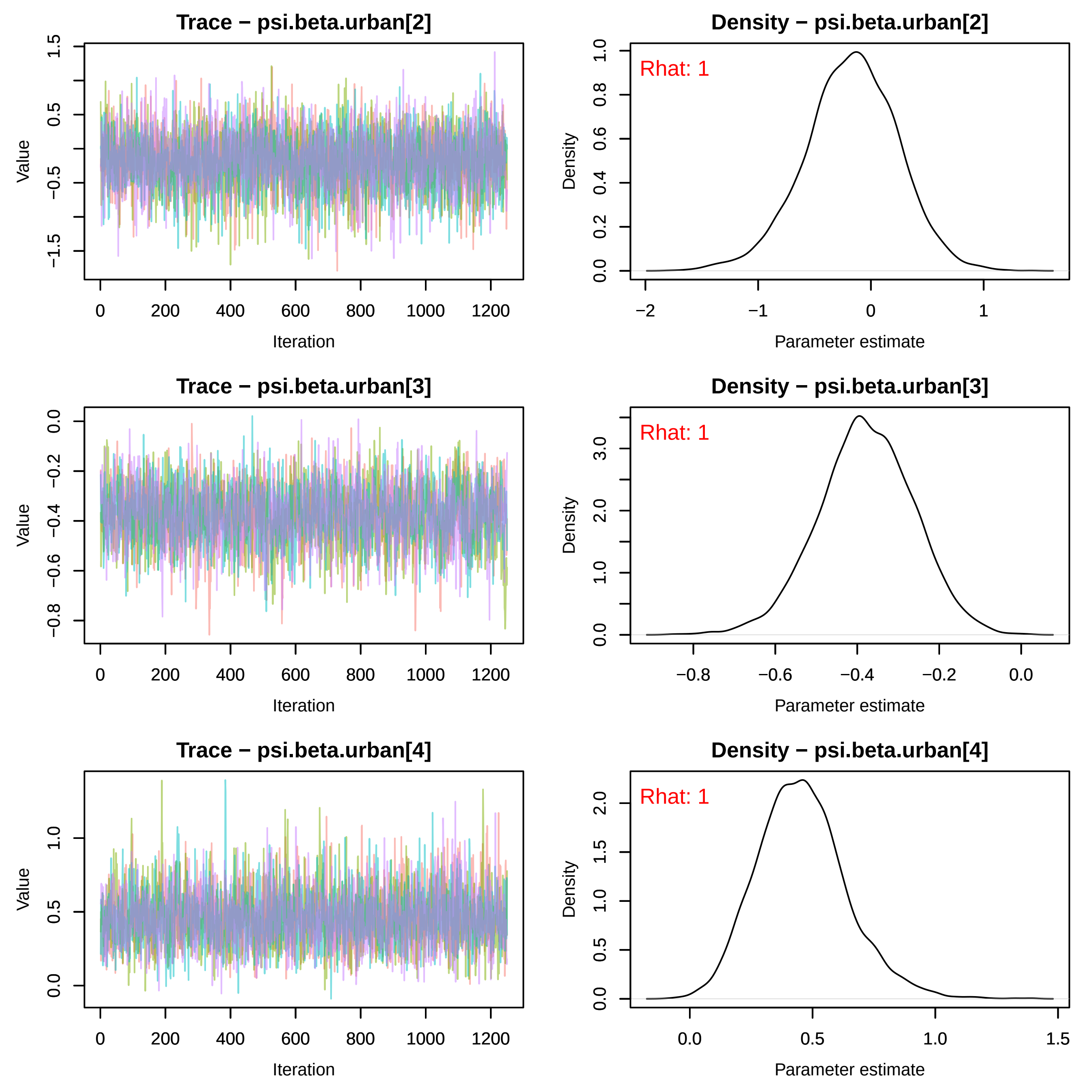
Trace − psi.beta.urban[2]
Density − psi.beta.urban[2]
Rhat: 1
Trace − psi.beta.urban[3]
Density − psi.beta.urban[3]
Rhat: 1
Trace − psi.beta.urban[4]
Density − psi.beta.urban[4]
Rhat: 1
Value
Iteration
Density
Parameter estimate

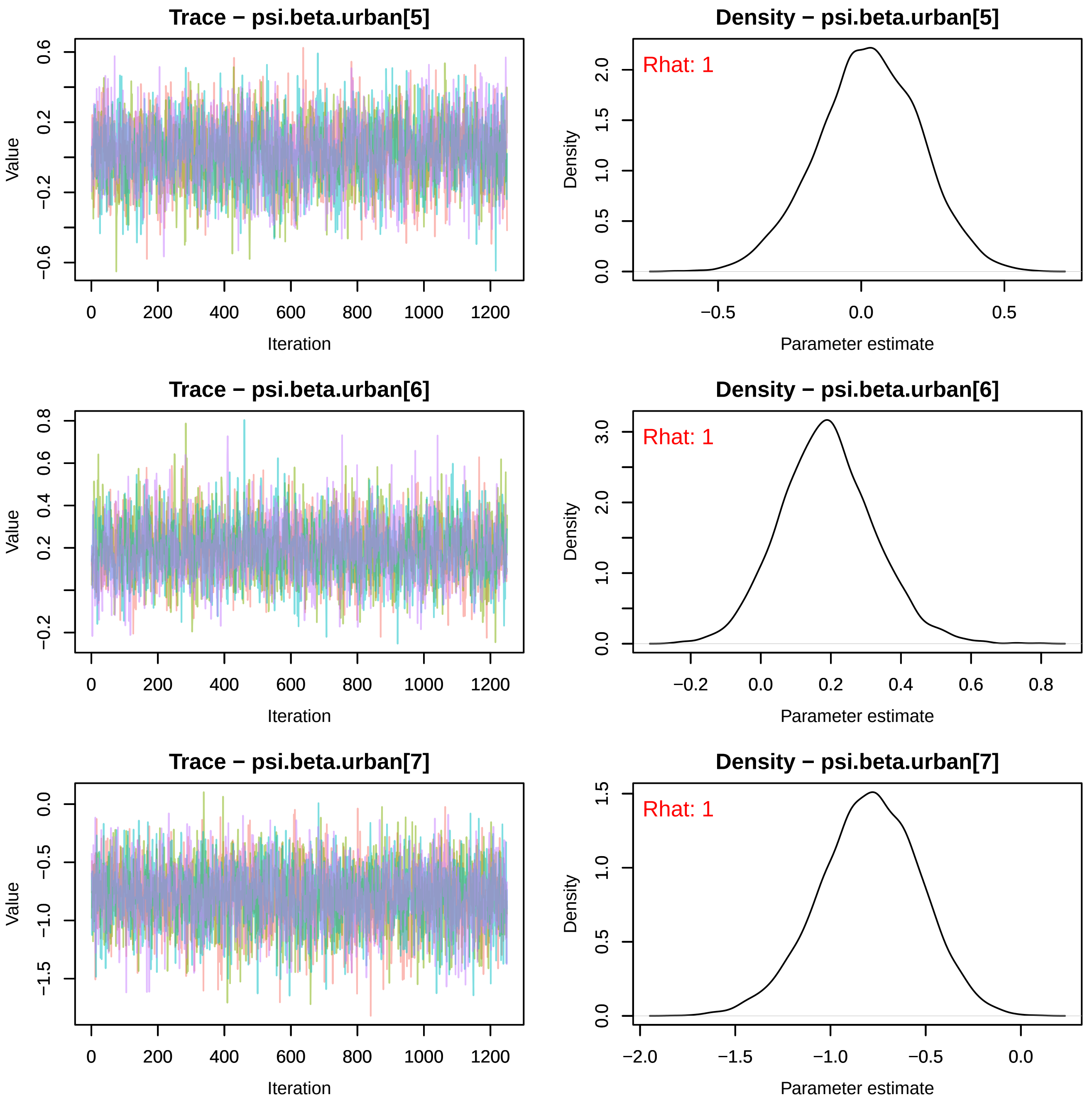
Trace − psi.beta.urban[5]
Value
Iteration
Density − psi.beta.urban[5]
Rhat: 1
Density
Parameter estimate
Trace − psi.beta.urban[6]
Value
Iteration
Density − psi.beta.urban[6]
Rhat: 1
Density
Parameter estimate
Trace − psi.beta.urban[7]
Value
Iteration
Density − psi.beta.urban[7]
Rhat: 1
Density
Parameter estimate

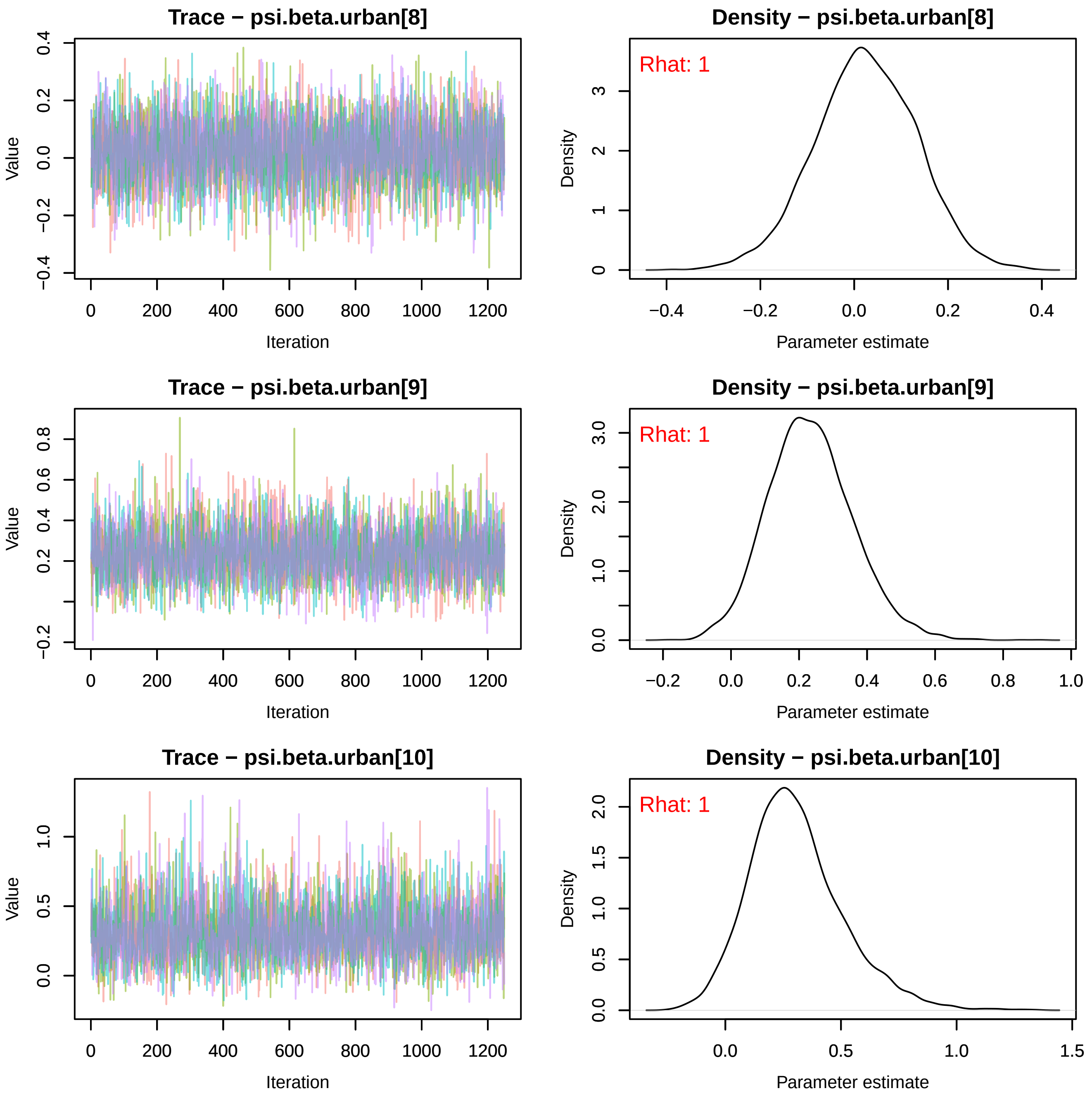
Trace − psi.beta.urban[8]
Value
Iteration
0 200 400 600 800 1000 1200
−0.4 −0.2 0.0 0.2 0.4
Density − psi.beta.urban[8]
Rhat: 1
Density
Parameter estimate
−0.4 −0.2 0.0 0.2 0.4
0 1 2 3
Trace − psi.beta.urban[9]
Value
Iteration
0 200 400 600 800 1000 1200
−0.2 0.2 0.4 0.6 0.8
Density − psi.beta.urban[9]
Rhat: 1
Density
Parameter estimate
−0.2 0.0 0.2 0.4 0.6 0.8 1.0
0.0 1.0 2.0 3.0
Trace − psi.beta.urban[10]
Value
Iteration
0 200 400 600 800 1000 1200
0.0 0.5 1.0
Density − psi.beta.urban[10]
Rhat: 1
Density
Parameter estimate
0.0 0.5 1.0 1.5
0.0 0.5 1.0 1.5 2.0

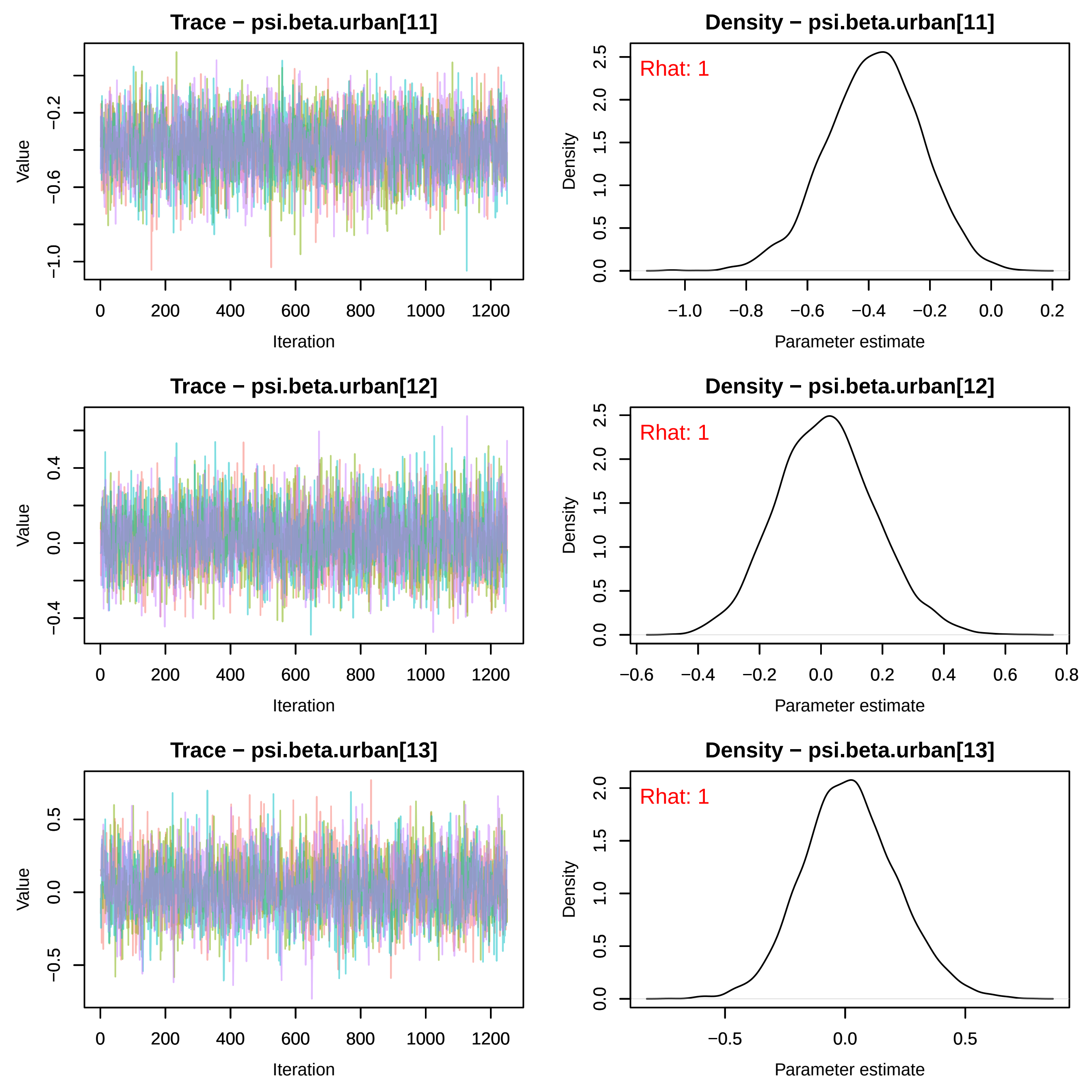
Trace − psi.beta.urban[11]
Density − psi.beta.urban[11]
Rhat: 1
Trace − psi.beta.urban[12]
Density − psi.beta.urban[12]
Rhat: 1
Trace − psi.beta.urban[13]
Density − psi.beta.urban[13]
Rhat: 1
Value
Iteration
Density
Parameter estimate
0
200
400
600
800
1000
1200
−0.2
−0.6
−1.0
0.4
0.0
−0.4
0.5
−0.5
0.0
0.5
1.0
1.5
2.0
2.5
−1.0
−0.8
−0.6
−0.4
−0.2
0.2
0.6
0.8

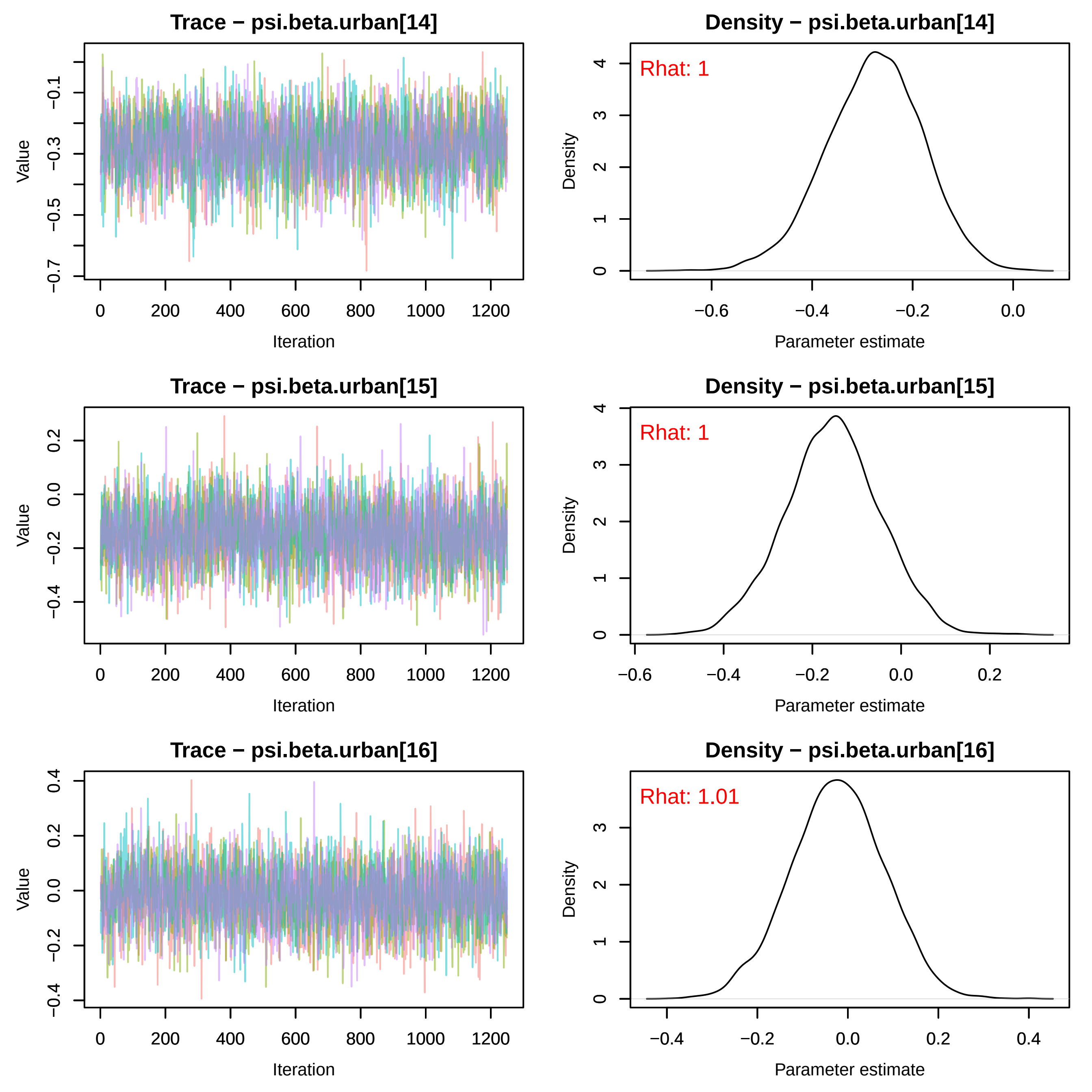
Trace − psi.beta.urban[14]
Value
Iteration
Density − psi.beta.urban[14]
Rhat: 1
Density
Parameter estimate
Trace − psi.beta.urban[15]
Value
Iteration
Density − psi.beta.urban[15]
Rhat: 1
Density
Parameter estimate
Trace − psi.beta.urban[16]
Value
Iteration
Density − psi.beta.urban[16]
Rhat: 1.01
Density
Parameter estimate

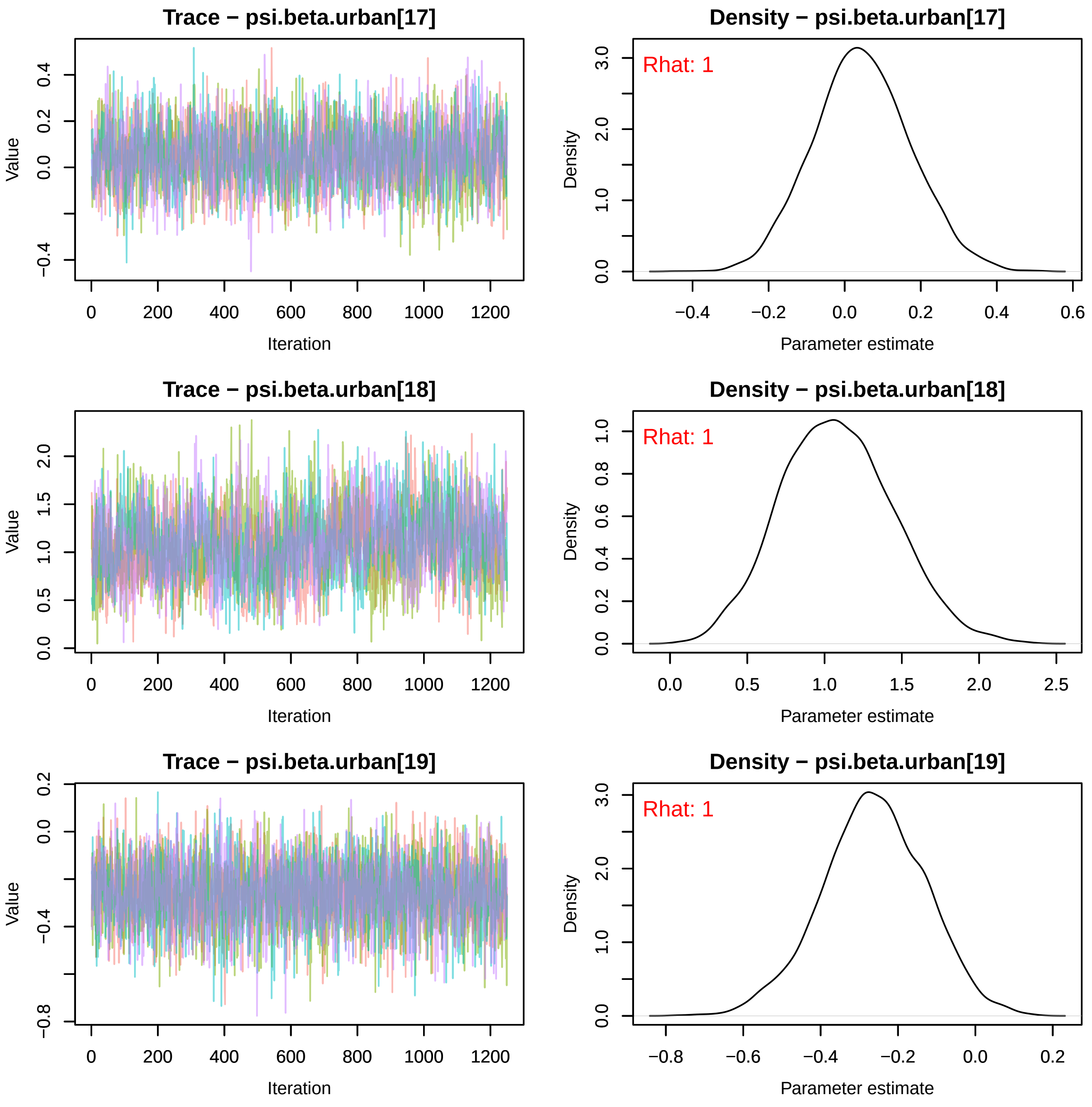

Trace − psi.beta.urban[17]
Value
Iteration
Density − psi.beta.urban[17]
Rhat: 1
Density
Parameter estimate
Trace − psi.beta.urban[18]
Value
Iteration
Density − psi.beta.urban[18]
Rhat: 1
Density
Parameter estimate
Trace − psi.beta.urban[19]
Value
Iteration
Density − psi.beta.urban[19]
Rhat: 1
Density
Parameter estimate

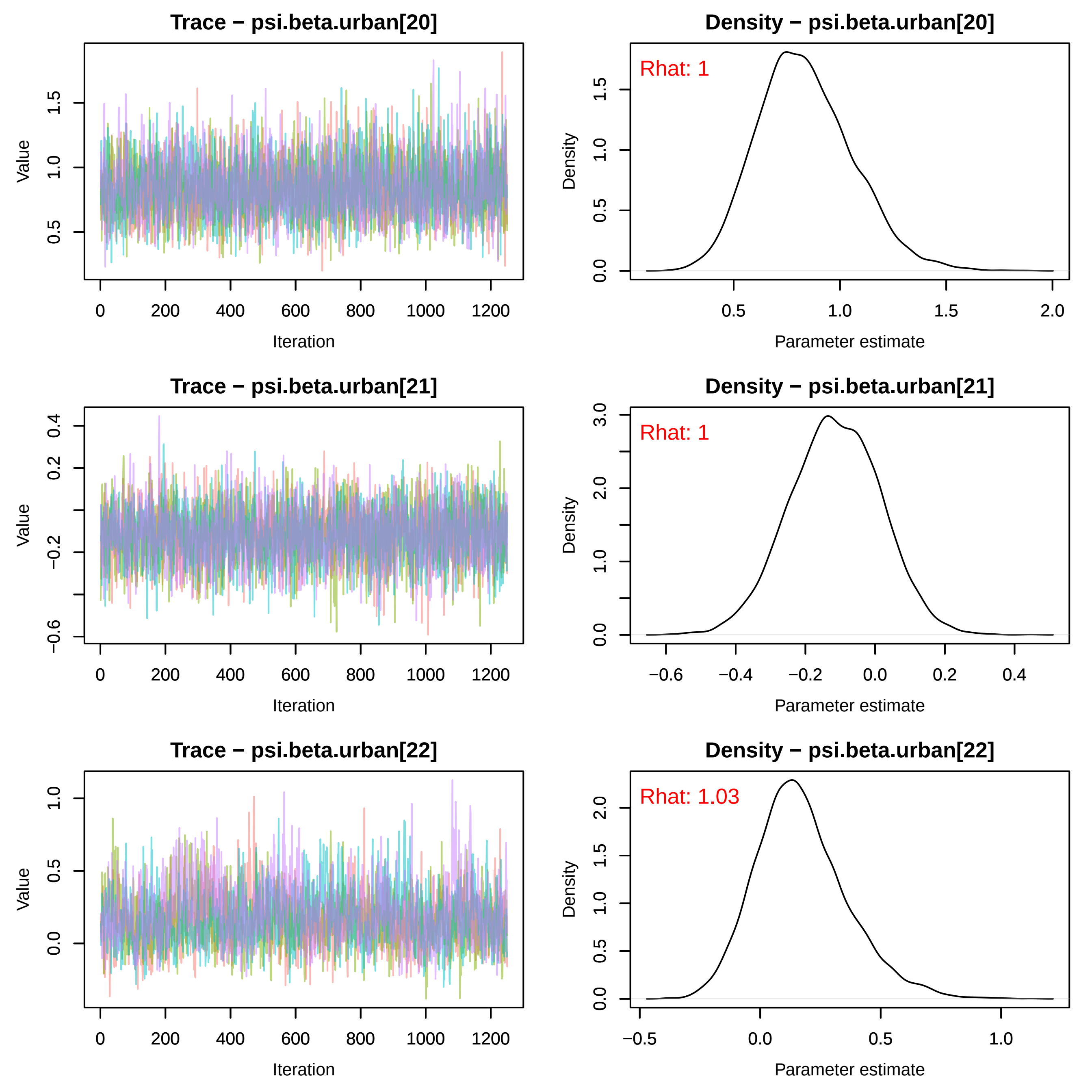

Trace − psi.beta.urban[20]
Value
Iteration
Density − psi.beta.urban[20]
Rhat: 1
Density
Parameter estimate
Trace − psi.beta.urban[21]
Value
Iteration
Density − psi.beta.urban[21]
Rhat: 1
Density
Parameter estimate
Trace − psi.beta.urban[22]
Value
Iteration
Density − psi.beta.urban[22]
Rhat: 1.03
Density
Parameter estimate

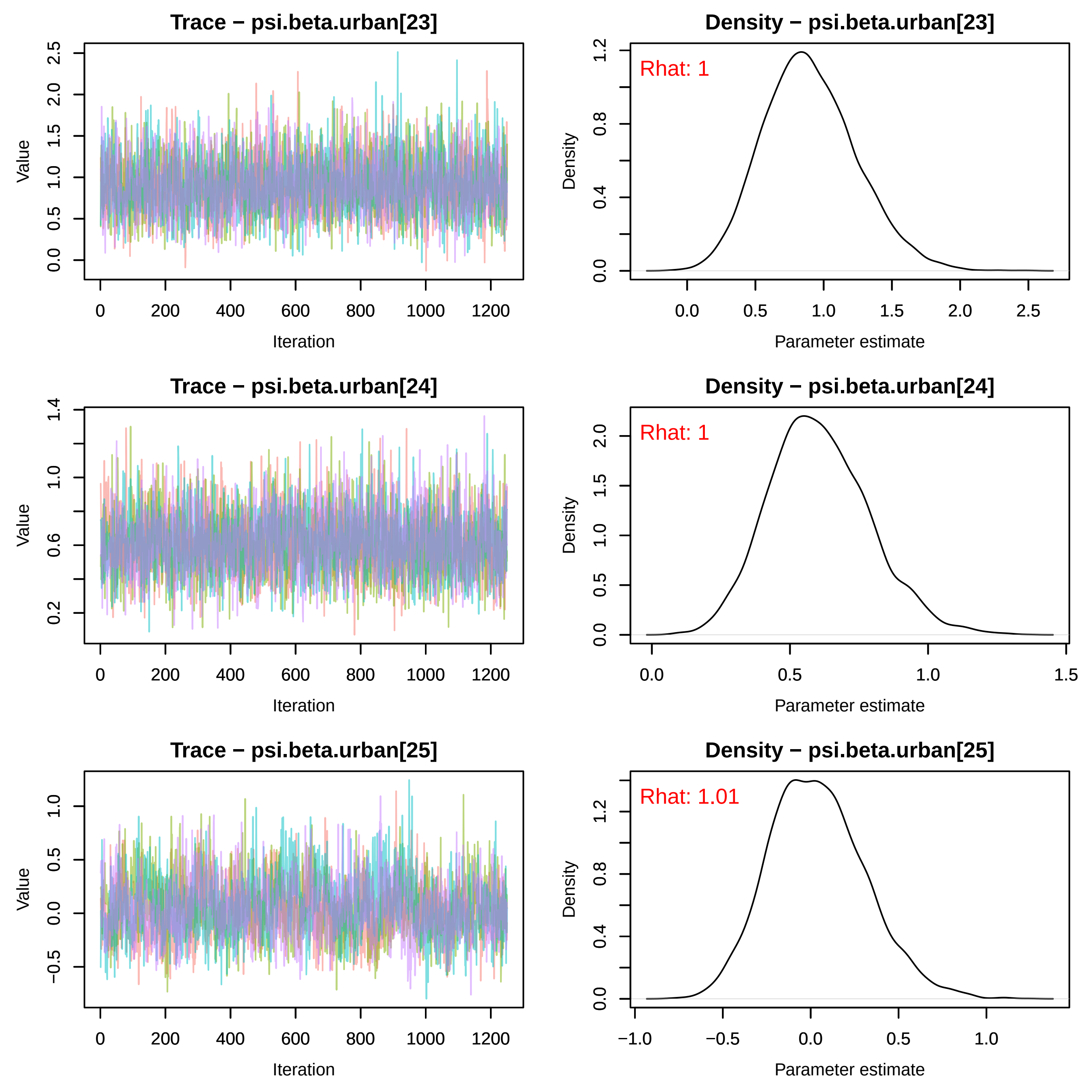
Trace − psi.beta.urban[23]
Value
Iteration
Density − psi.beta.urban[23]
Rhat: 1
Density
Parameter estimate
Trace − psi.beta.urban[24]
Value
Iteration
Density − psi.beta.urban[24]
Rhat: 1
Density
Parameter estimate
Trace − psi.beta.urban[25]
Value
Iteration
Density − psi.beta.urban[25]
Rhat: 1.01
Density
Parameter estimate

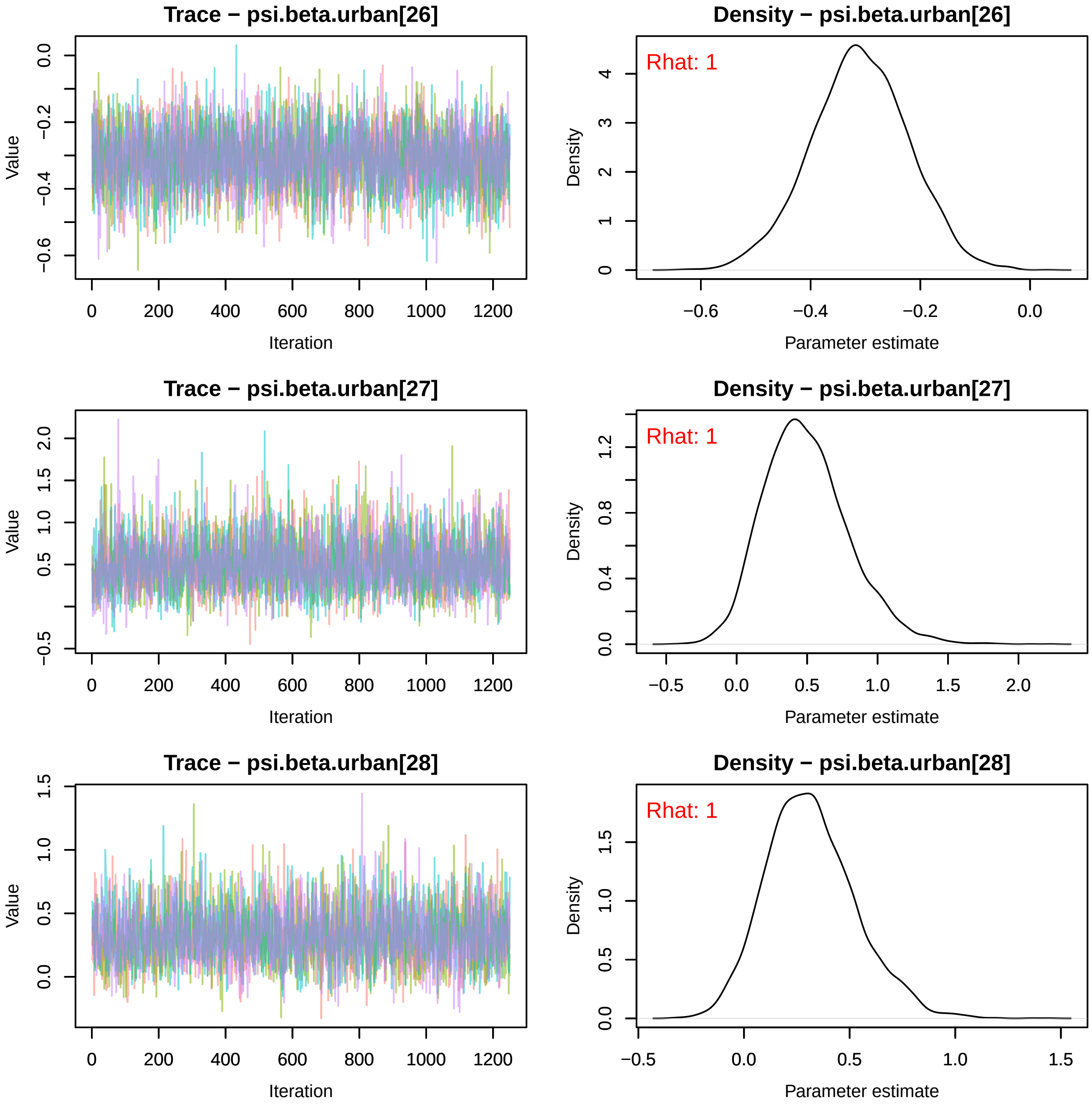
Trace − psi.beta.urban[26]
Value
Iteration
Density − psi.beta.urban[26]
Rhat: 1
Density
Parameter estimate
Trace − psi.beta.urban[27]
Value
Iteration
Density − psi.beta.urban[27]
Rhat: 1
Density
Parameter estimate
Trace − psi.beta.urban[28]
Value
Iteration
Density − psi.beta.urban[28]
Rhat: 1
Density
Parameter estimate

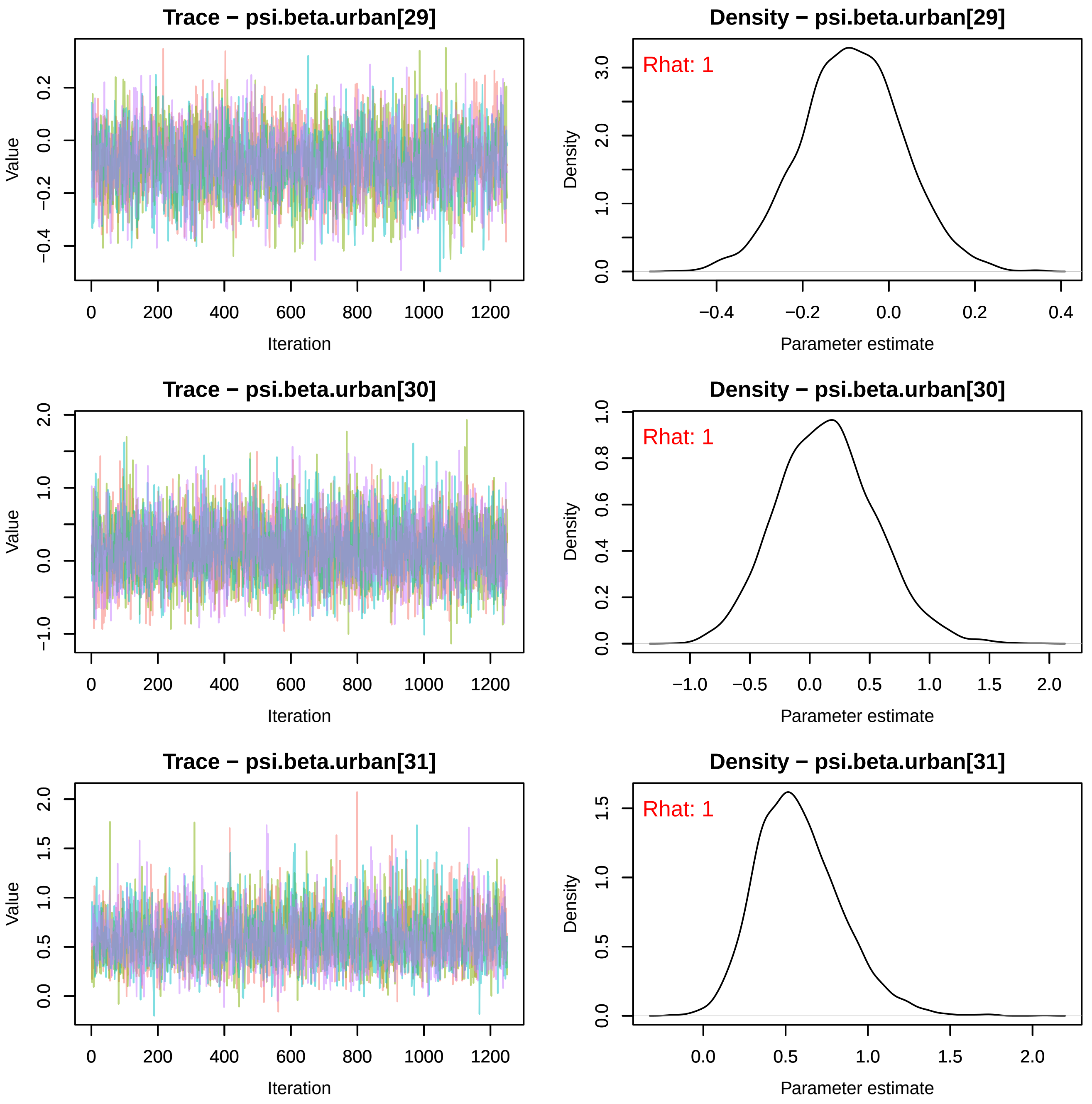
Trace − psi.beta.urban[29]
Density − psi.beta.urban[29]
Rhat: 1
Trace − psi.beta.urban[30]
Density − psi.beta.urban[30]
Rhat: 1
Trace − psi.beta.urban[31]
Density − psi.beta.urban[31]
Rhat: 1
Value
Iteration
Density
Parameter estimate

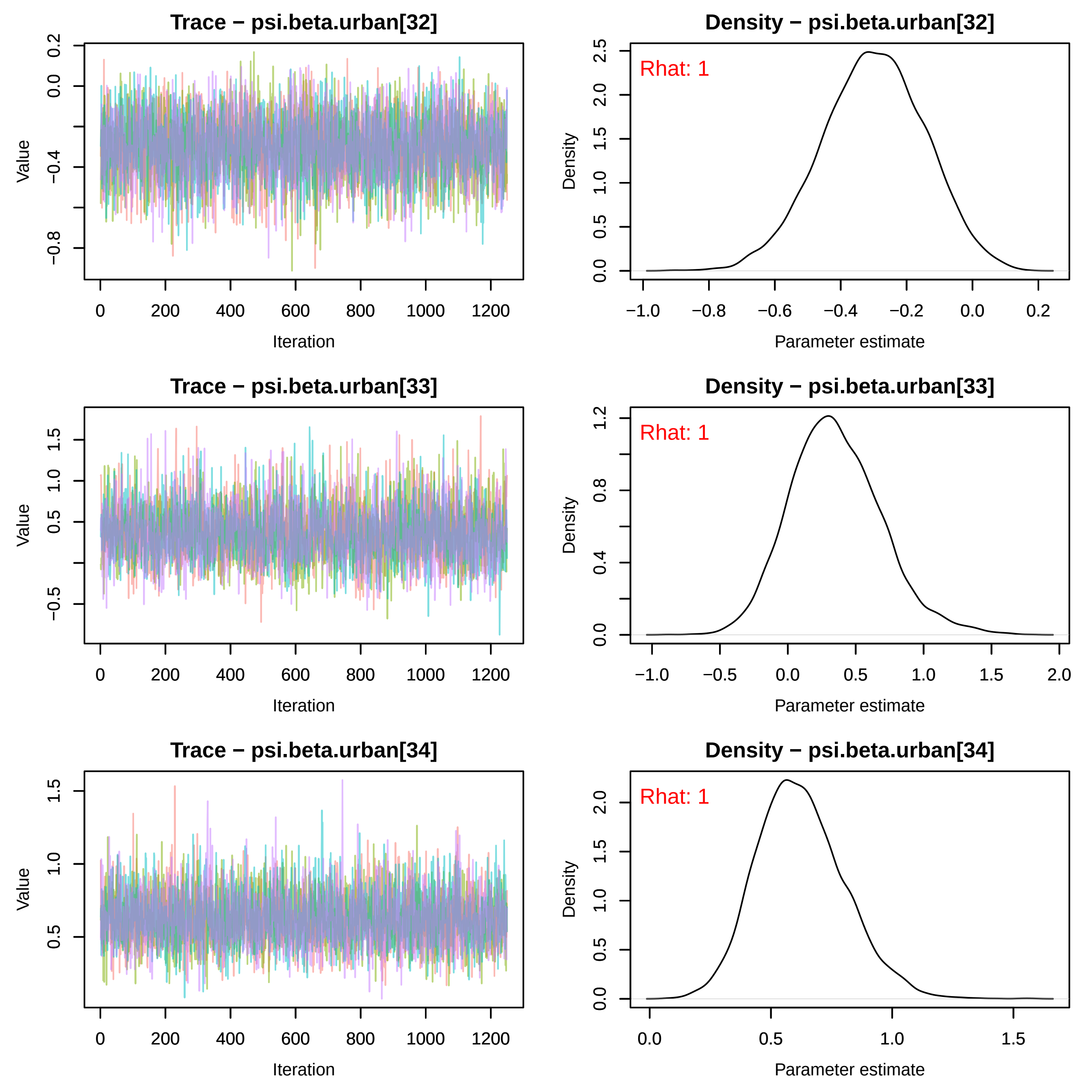
Trace − psi.beta.urban[32]
Value
0.2
0.0
−0.4
−0.8
0
200
400
600
800
1000
1200
Iteration
Density − psi.beta.urban[32]
Rhat: 1
Density
0.0
0.5
1.0
1.5
2.0
2.5
−1.0
−0.8
−0.6
−0.4
−0.2
0.0
0.2
Parameter estimate
Trace − psi.beta.urban[33]
Value
−0.5
0.5
1.0
1.5
0
200
400
600
800
1000
1200
Iteration
Density − psi.beta.urban[33]
Rhat: 1
Density
0.0
0.4
0.8
1.2
−1.0
−0.5
0.0
0.5
1.0
1.5
2.0
Parameter estimate
Trace − psi.beta.urban[34]
Value
0.5
1.0
1.5
0
200
400
600
800
1000
1200
Iteration
Density − psi.beta.urban[34]
Rhat: 1
Density
0.0
0.5
1.0
1.5
2.0
0.0
0.5
1.0
1.5
Parameter estimate

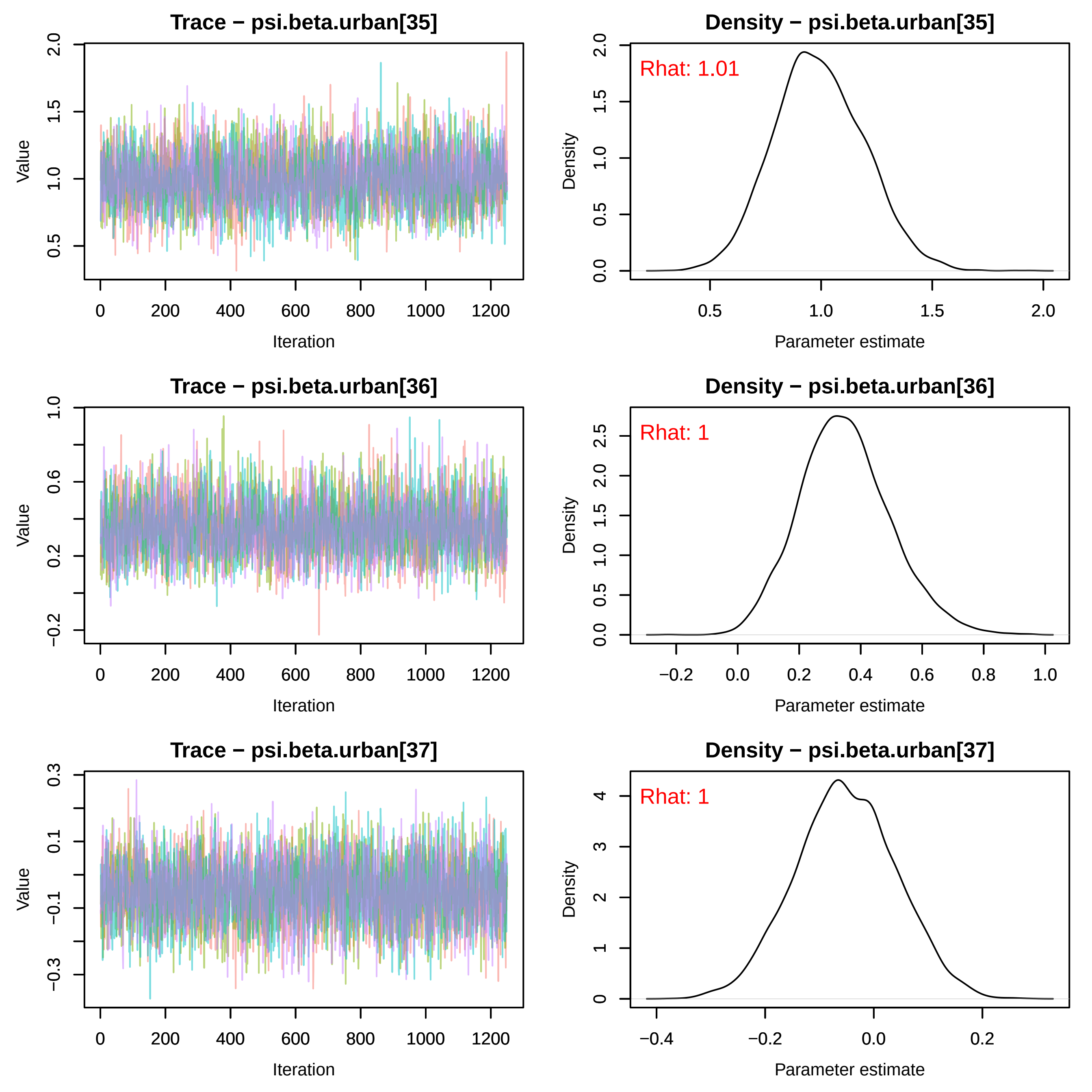
Trace − psi.beta.urban[35]
Density − psi.beta.urban[35]
Rhat: 1.01
Trace − psi.beta.urban[36]
Density − psi.beta.urban[36]
Rhat: 1
Trace − psi.beta.urban[37]
Density − psi.beta.urban[37]
Rhat: 1
Value
Density
Iteration
Parameter estimate
0
200
400
600
800
1000
1200

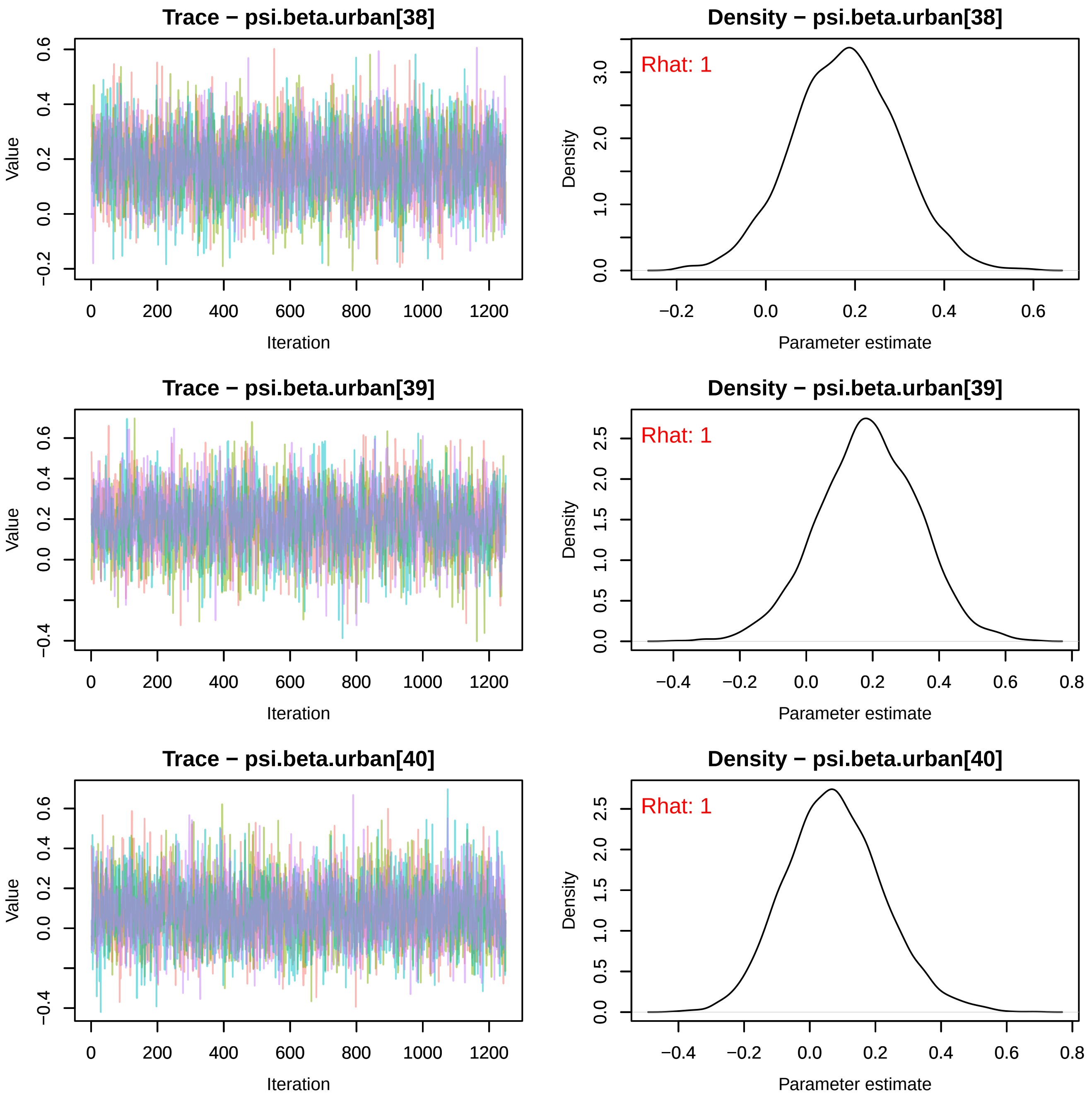
Trace − psi.beta.urban[38]
Density − psi.beta.urban[38]
Rhat: 1
Value
Iteration
Density
Parameter estimate
Trace − psi.beta.urban[39]
Density − psi.beta.urban[39]
Rhat: 1
Value
Iteration
Density
Parameter estimate
Trace − psi.beta.urban[40]
Density − psi.beta.urban[40]
Rhat: 1
Value
Iteration
Density
Parameter estimate

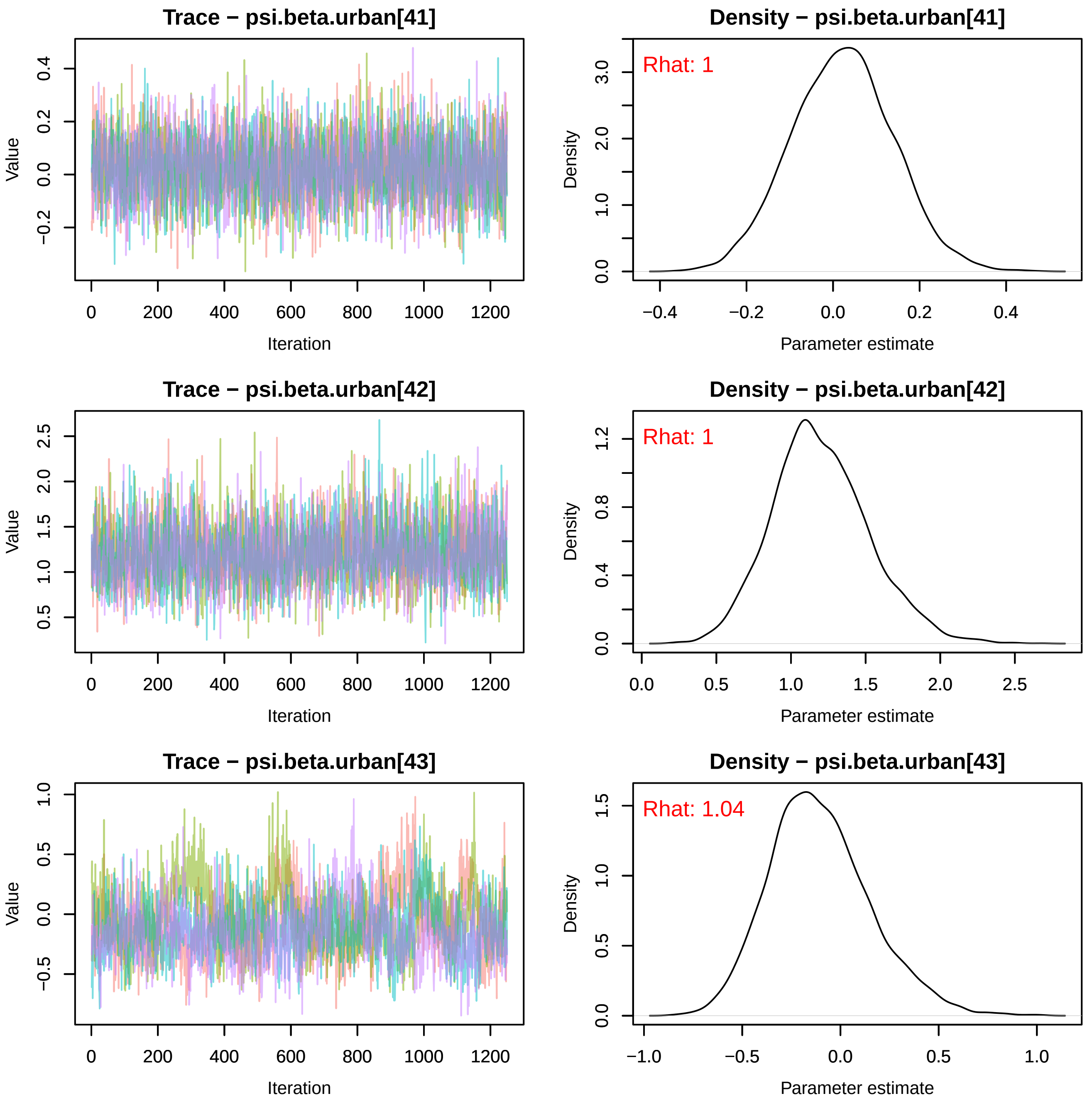

Trace − psi.beta.urban[41]
Value
Iteration
Density − psi.beta.urban[41]
Rhat: 1
Density
Parameter estimate
Trace − psi.beta.urban[42]
Value
Iteration
Density − psi.beta.urban[42]
Rhat: 1
Density
Parameter estimate
Trace − psi.beta.urban[43]
Value
Iteration
Density − psi.beta.urban[43]
Rhat: 1.04
Density
Parameter estimate

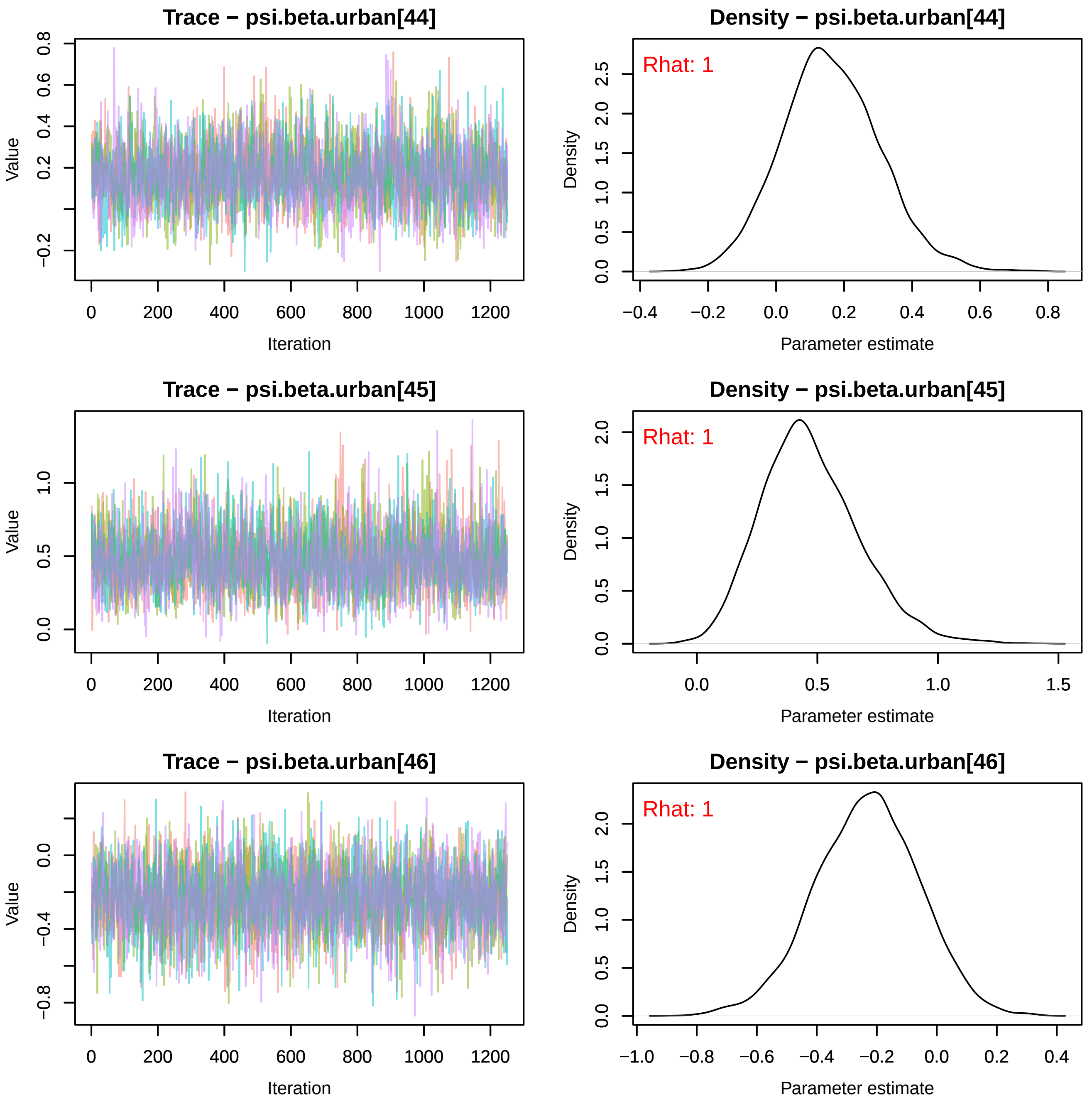

Trace − psi.beta.urban[44]
Value
−0.2
0.2
0.4
0.6
0.8
0
200
400
600
800
1000
1200
Iteration
Density − psi.beta.urban[44]
Rhat: 1
Density
0.0
0.5
1.0
1.5
2.0
2.5
−0.4
−0.2
0.0
0.2
0.4
0.6
0.8
Parameter estimate
Trace − psi.beta.urban[45]
Value
0.0
0.5
1.0
0
200
400
600
800
1000
1200
Iteration
Density − psi.beta.urban[45]
Rhat: 1
Density
0.0
0.5
1.0
1.5
2.0
0.0
0.5
1.0
1.5
Parameter estimate
Trace − psi.beta.urban[46]
Value
−0.8
−0.4
0.0
0
200
400
600
800
1000
1200
Iteration
Density − psi.beta.urban[46]
Rhat: 1
Density
0.0
0.5
1.0
1.5
2.0
−1.0
−0.8
−0.6
−0.4
−0.2
0.0
0.2
0.4
Parameter estimate

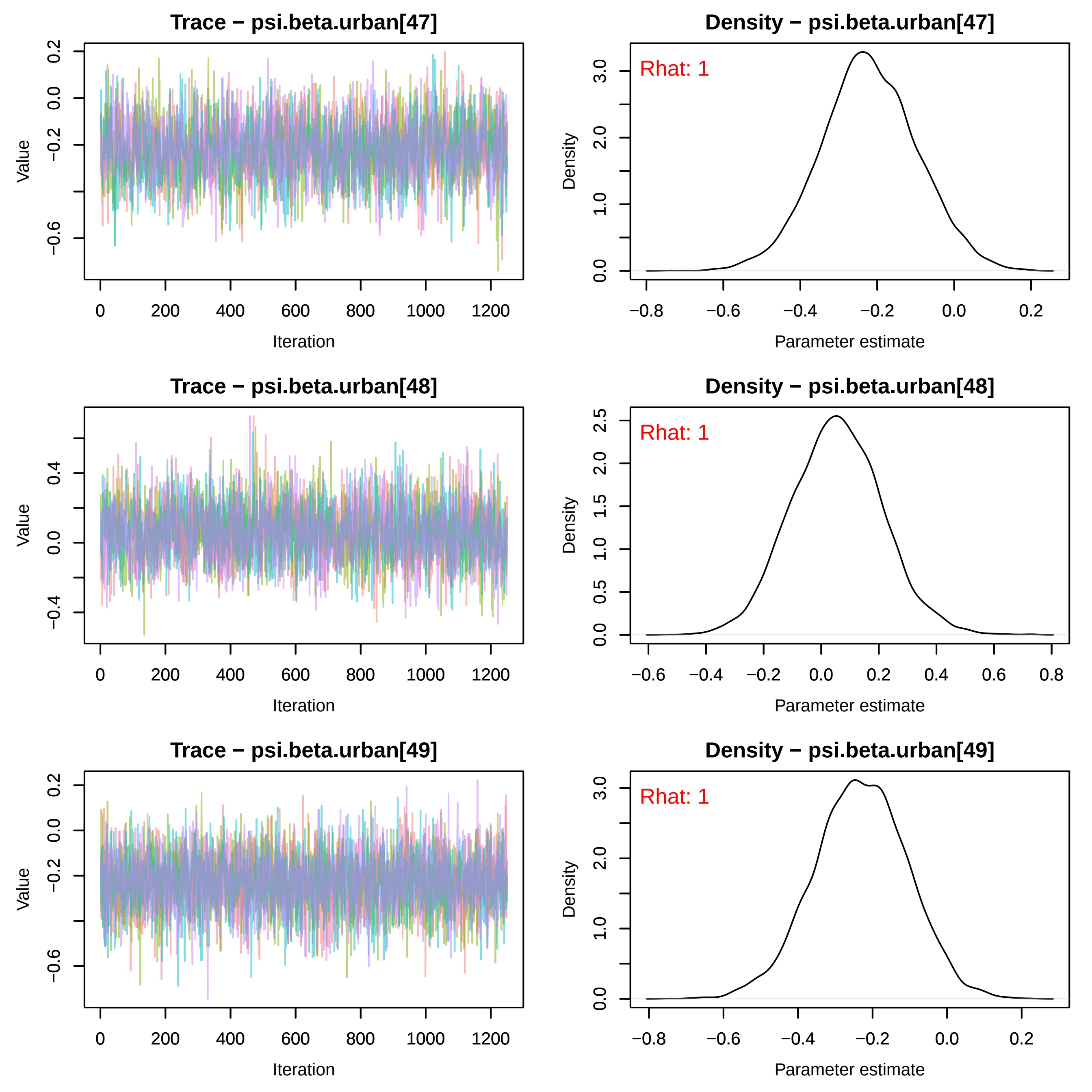
Trace − psi.beta.urban[47]
Value
Iteration
Density − psi.beta.urban[47]
Rhat: 1
Density
Parameter estimate
Trace − psi.beta.urban[48]
Value
Iteration
Density − psi.beta.urban[48]
Rhat: 1
Density
Parameter estimate
Trace − psi.beta.urban[49]
Value
Iteration
Density − psi.beta.urban[49]
Rhat: 1
Density
Parameter estimate

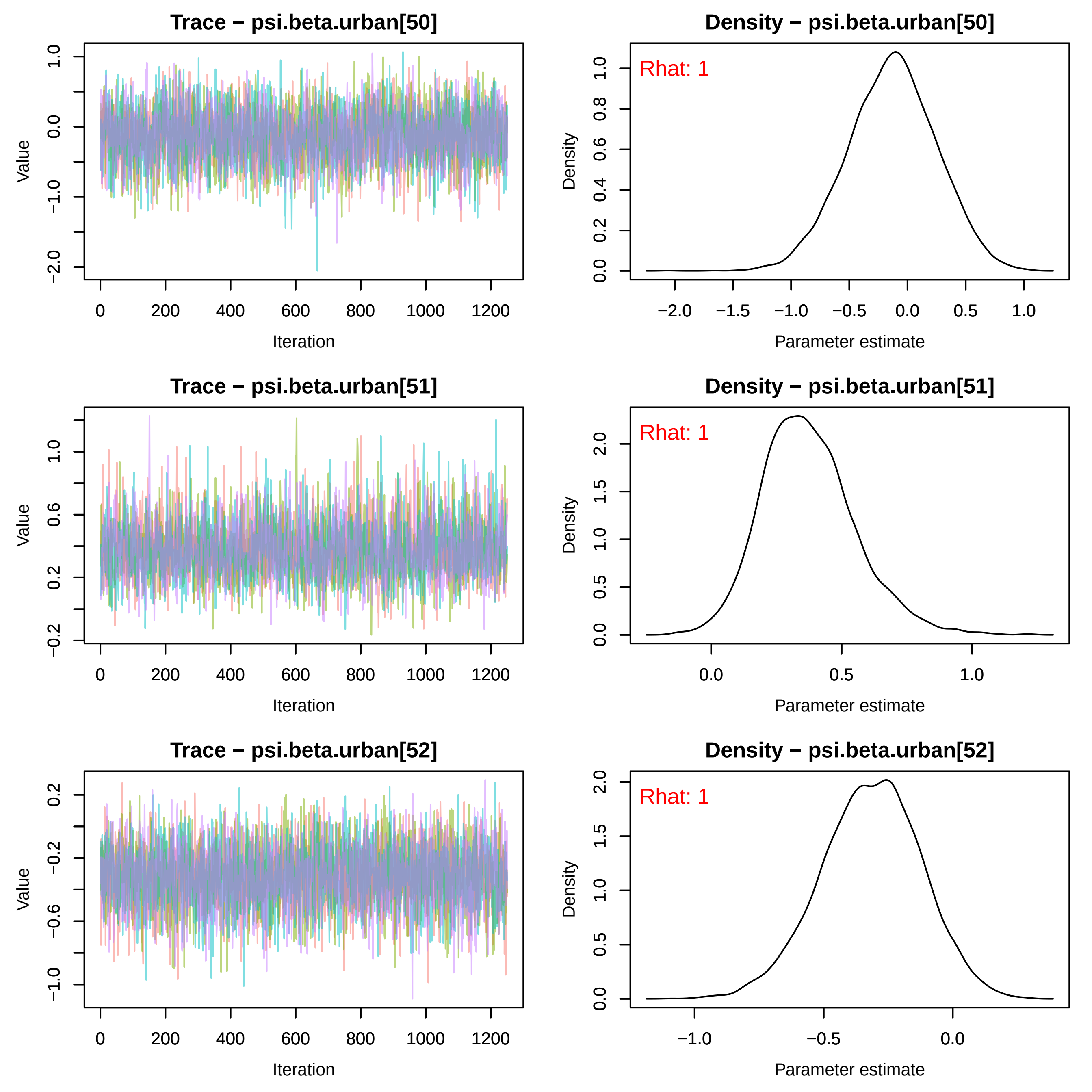
Trace − psi.beta.urban[50]
Density − psi.beta.urban[50]
Rhat: 1
Trace − psi.beta.urban[51]
Density − psi.beta.urban[51]
Rhat: 1
Trace − psi.beta.urban[52]
Density − psi.beta.urban[52]
Rhat: 1
Value
Iteration
Density
Parameter estimate

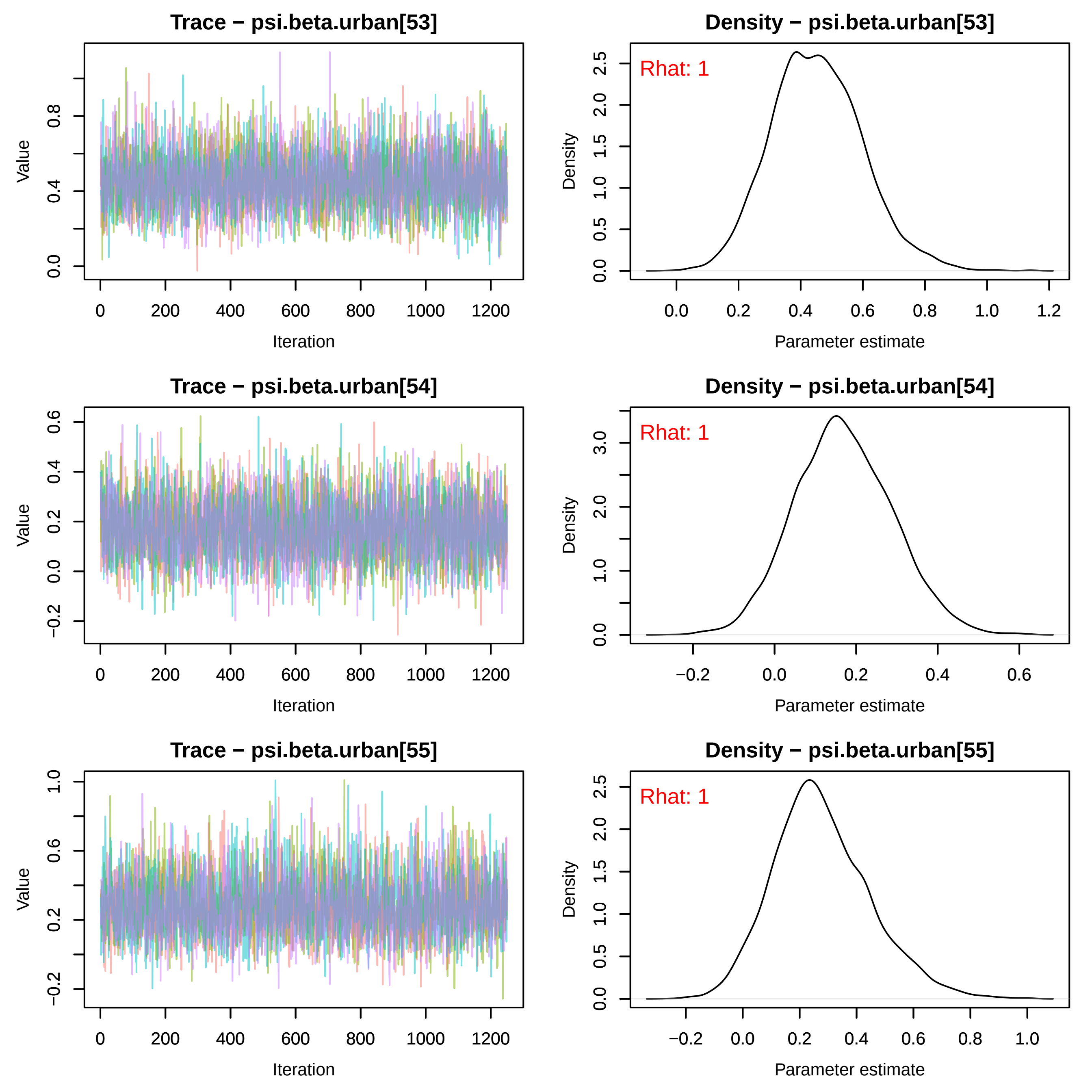
Trace − psi.beta.urban[53]
Value
Iteration
Density − psi.beta.urban[53]
Rhat: 1
Density
Parameter estimate
Trace − psi.beta.urban[54]
Value
Iteration
Density − psi.beta.urban[54]
Rhat: 1
Density
Parameter estimate
Trace − psi.beta.urban[55]
Value
Iteration
Density − psi.beta.urban[55]
Rhat: 1
Density
Parameter estimate

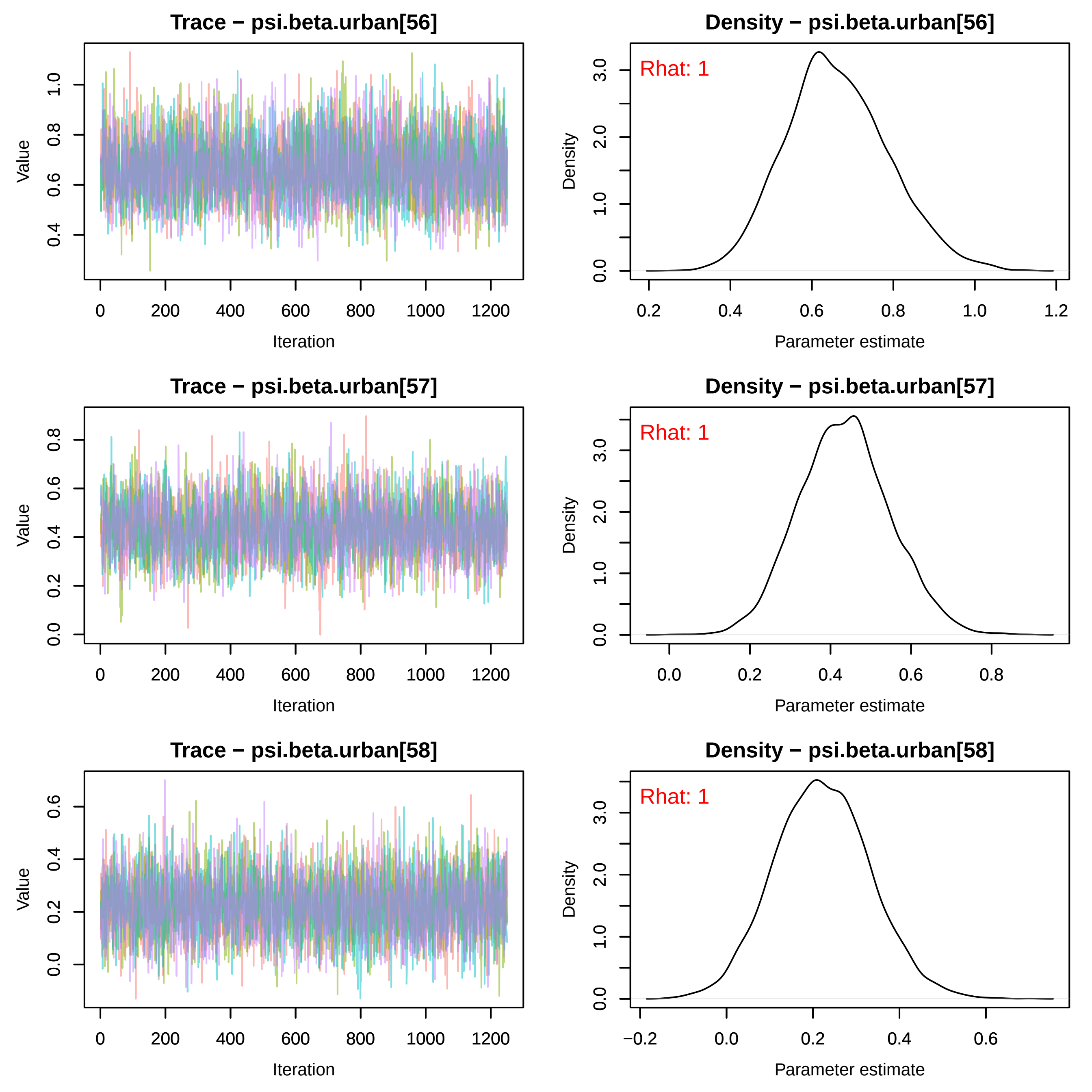
Trace − psi.beta.urban[56]
Value
Iteration
Density − psi.beta.urban[56]
Rhat: 1
Density
Parameter estimate
Trace − psi.beta.urban[57]
Value
Iteration
Density − psi.beta.urban[57]
Rhat: 1
Density
Parameter estimate
Trace − psi.beta.urban[58]
Value
Iteration
Density − psi.beta.urban[58]
Rhat: 1
Density
Parameter estimate

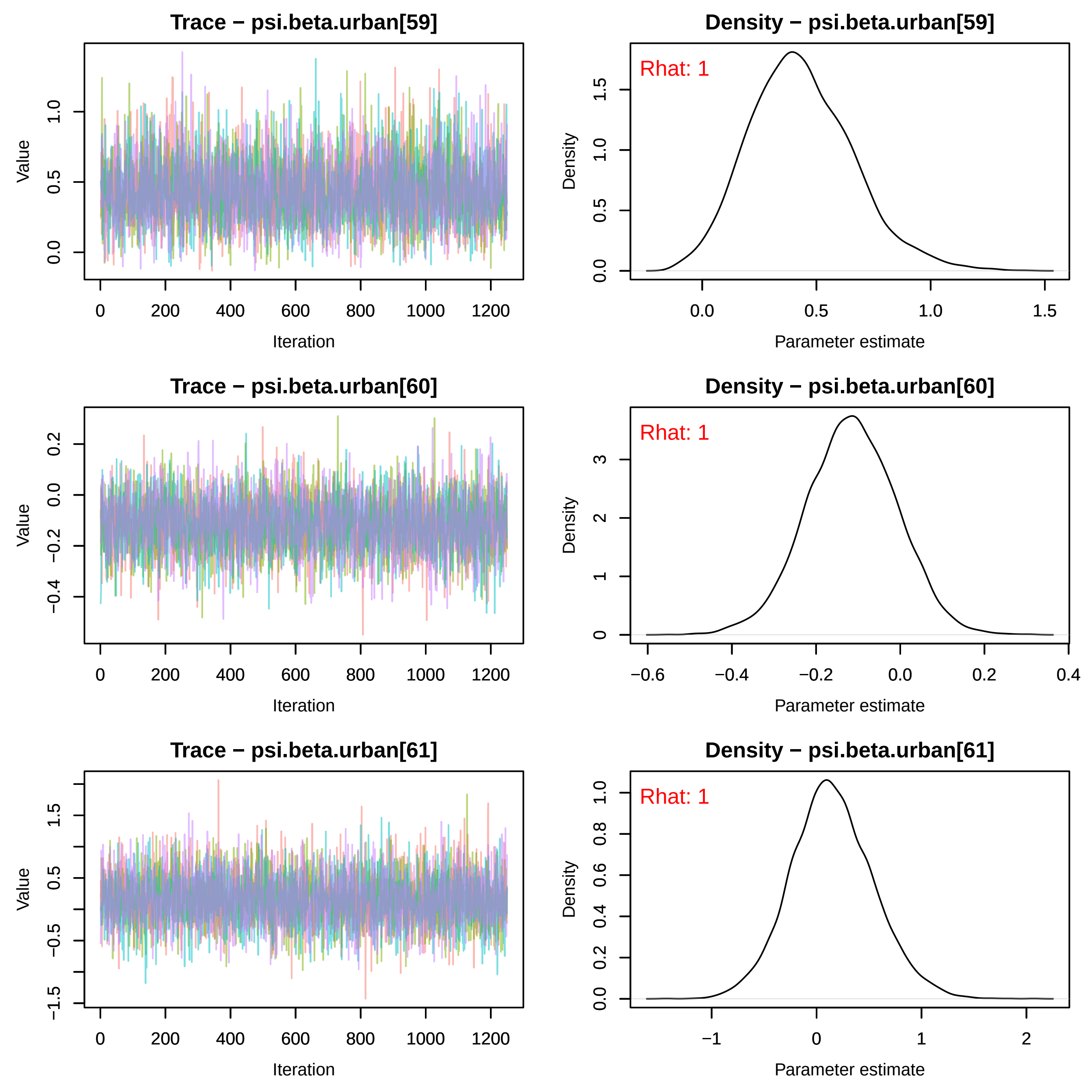
Trace − psi.beta.urban[59]
Density − psi.beta.urban[59]
Rhat: 1
Trace − psi.beta.urban[60]
Density − psi.beta.urban[60]
Rhat: 1
Trace − psi.beta.urban[61]
Density − psi.beta.urban[61]
Rhat: 1
Value
Iteration
Density
Parameter estimate
0
200
400
600
800
1000
1200

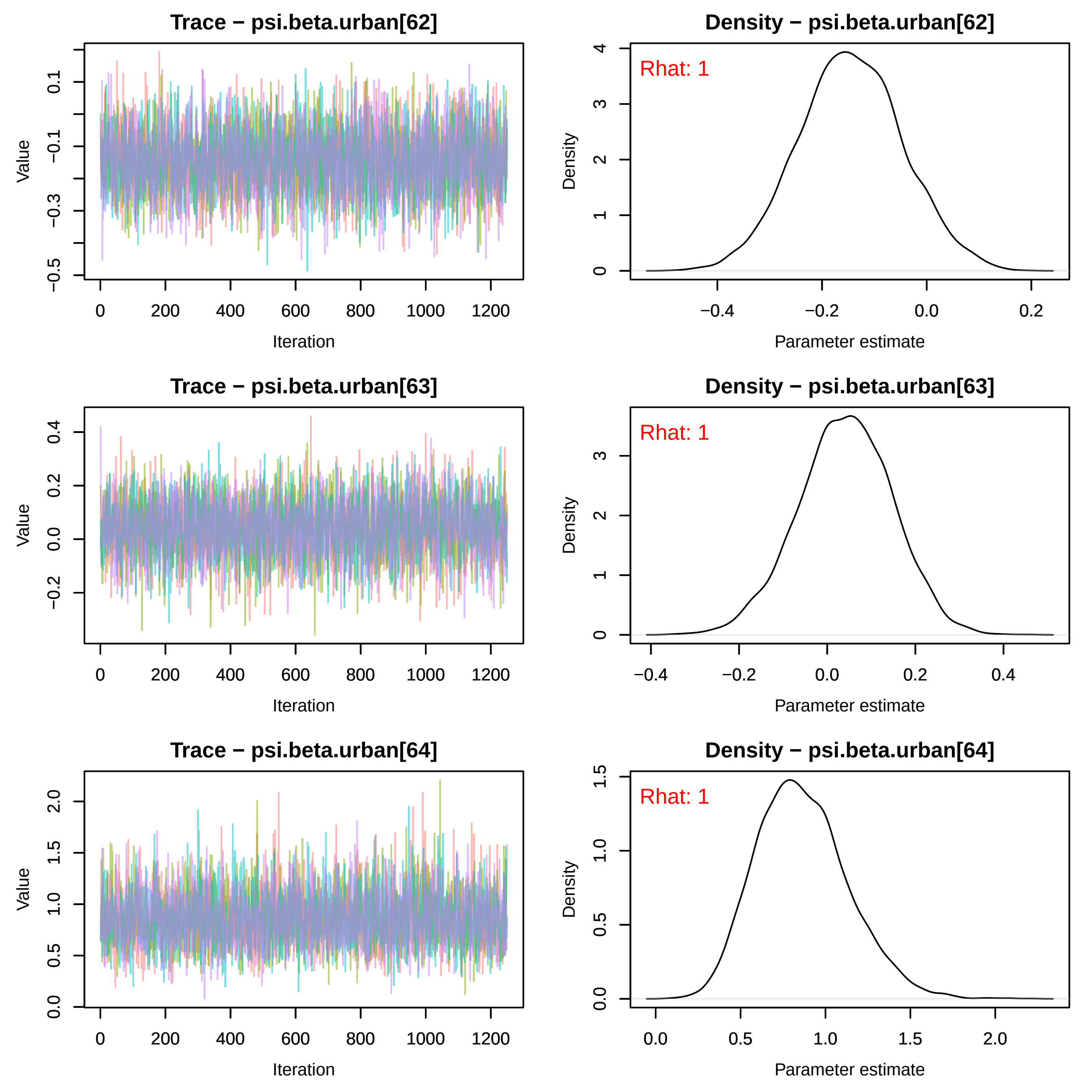
Trace − psi.beta.urban[62]
Density − psi.beta.urban[62]
Rhat: 1
Trace − psi.beta.urban[63]
Density − psi.beta.urban[63]
Rhat: 1
Trace − psi.beta.urban[64]
Density − psi.beta.urban[64]
Rhat: 1
Value
Iteration
Density
Parameter estimate

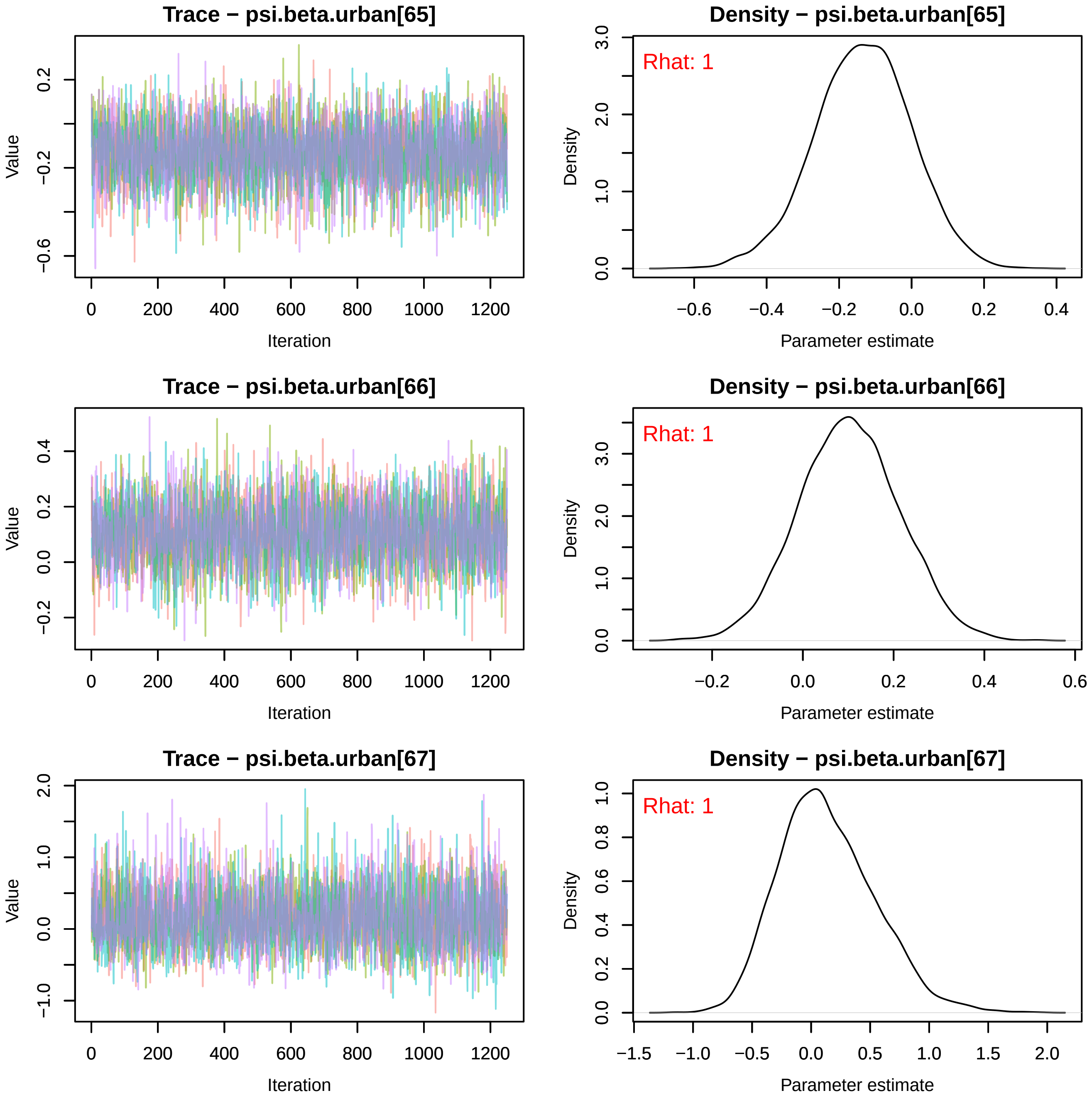
Trace − psi.beta.urban[65]
Value
Iteration
Density − psi.beta.urban[65]
Rhat: 1
Density
Parameter estimate
Trace − psi.beta.urban[66]
Value
Iteration
Density − psi.beta.urban[66]
Rhat: 1
Density
Parameter estimate
Trace − psi.beta.urban[67]
Value
Iteration
Density − psi.beta.urban[67]
Rhat: 1
Density
Parameter estimate

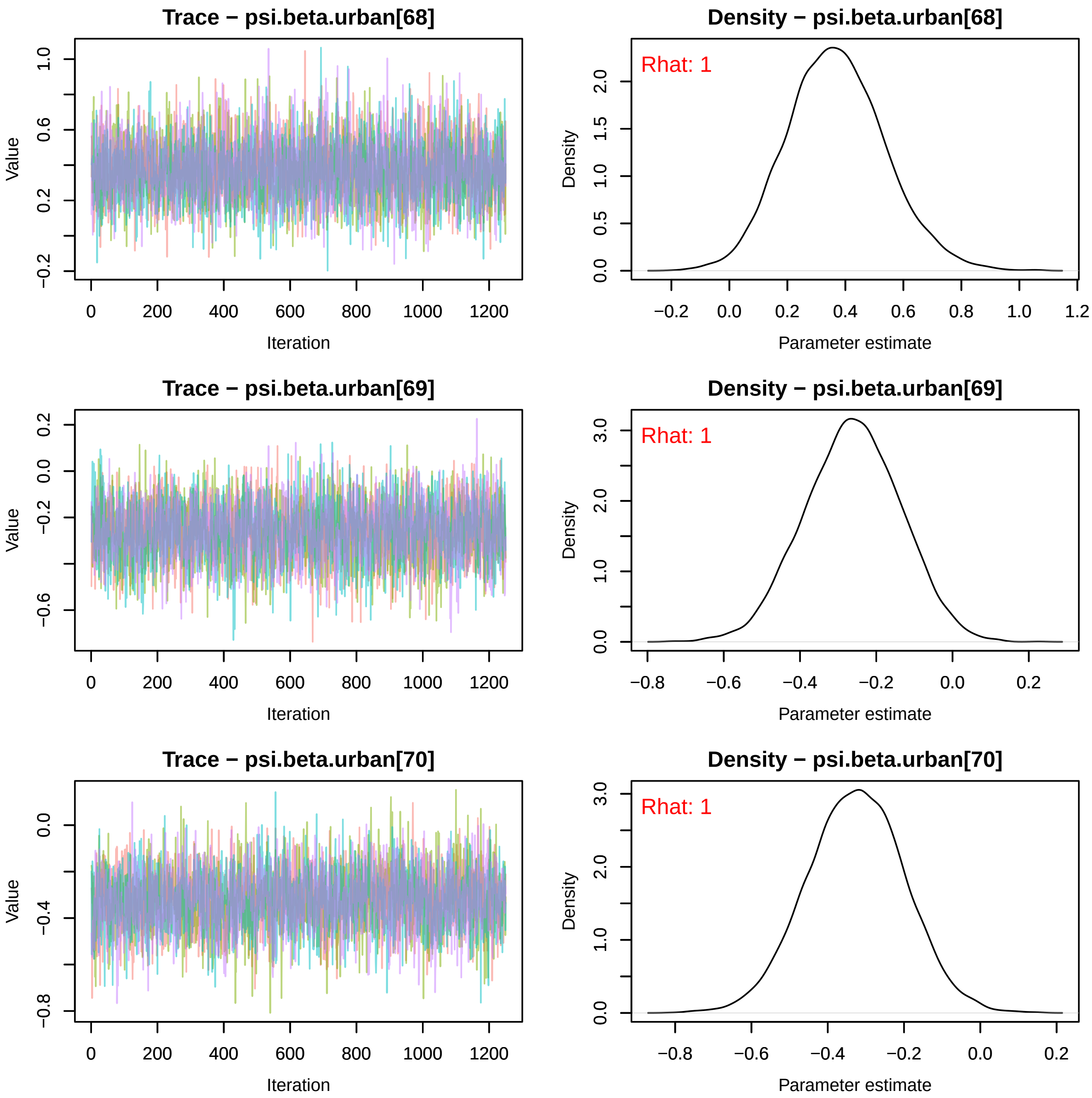
Trace − psi.beta.urban[68]
Value
Iteration
Density − psi.beta.urban[68]
Rhat: 1
Density
Parameter estimate
Trace − psi.beta.urban[69]
Value
Iteration
Density − psi.beta.urban[69]
Rhat: 1
Density
Parameter estimate
Trace − psi.beta.urban[70]
Value
Iteration
Density − psi.beta.urban[70]
Rhat: 1
Density
Parameter estimate

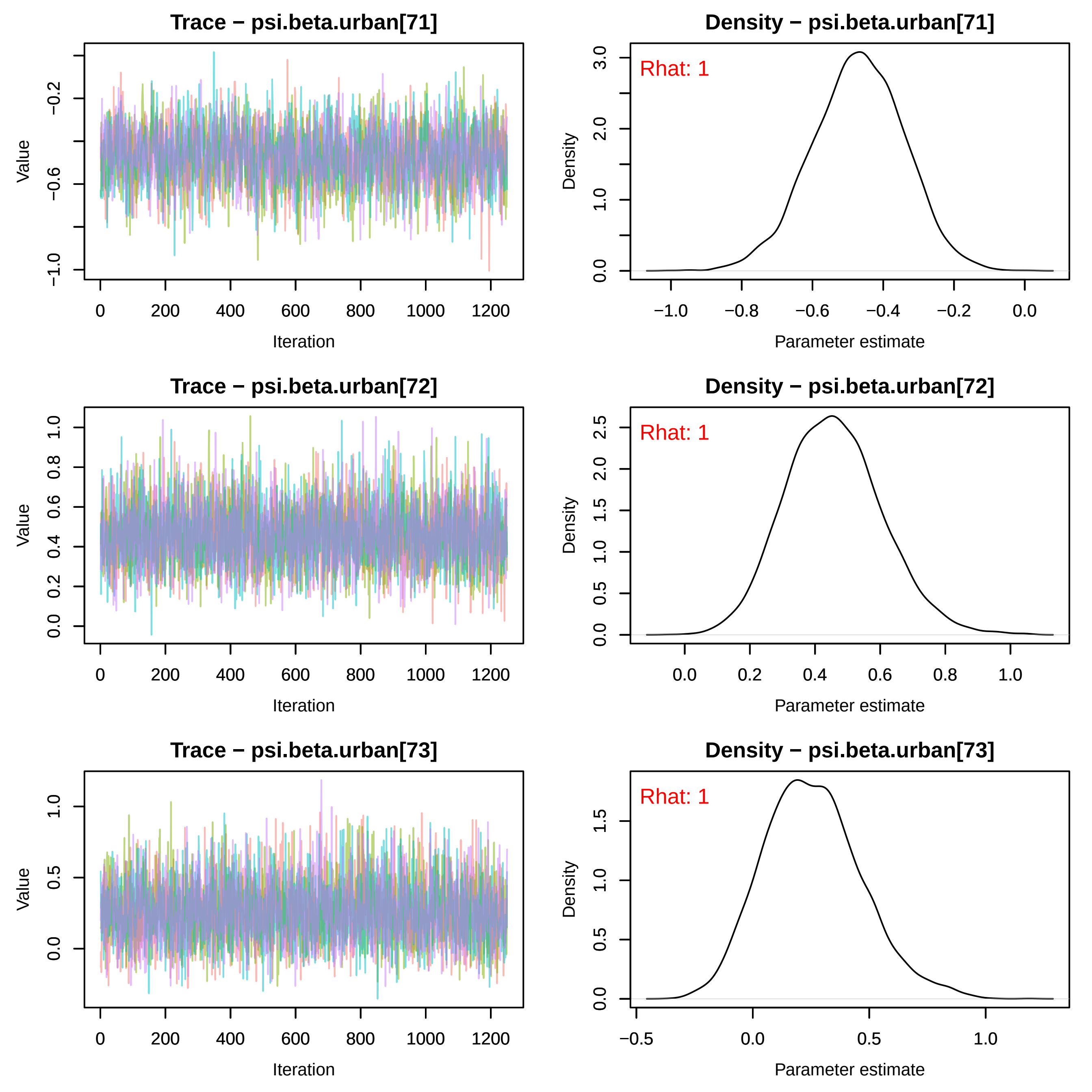
Trace − psi.beta.urban[71]
Value
Iteration
Density − psi.beta.urban[71]
Rhat: 1
Density
Parameter estimate
Trace − psi.beta.urban[72]
Value
Iteration
Density − psi.beta.urban[72]
Rhat: 1
Density
Parameter estimate
Trace − psi.beta.urban[73]
Value
Iteration
Density − psi.beta.urban[73]
Rhat: 1
Density
Parameter estimate

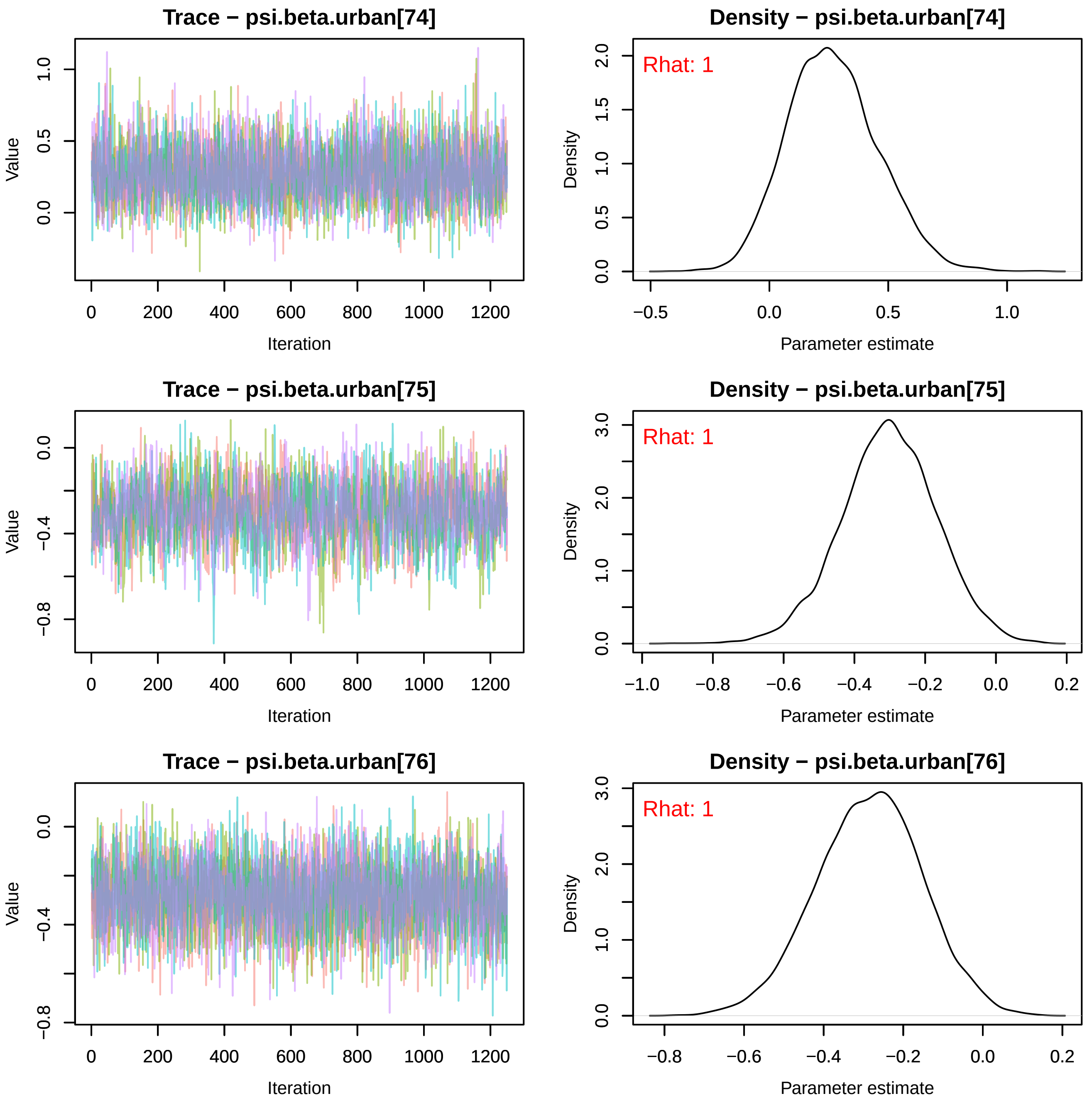
Trace − psi.beta.urban[74]
Density − psi.beta.urban[74]
Rhat: 1
Trace − psi.beta.urban[75]
Density − psi.beta.urban[75]
Rhat: 1
Trace − psi.beta.urban[76]
Density − psi.beta.urban[76]
Rhat: 1
Value
Iteration
Density
Parameter estimate

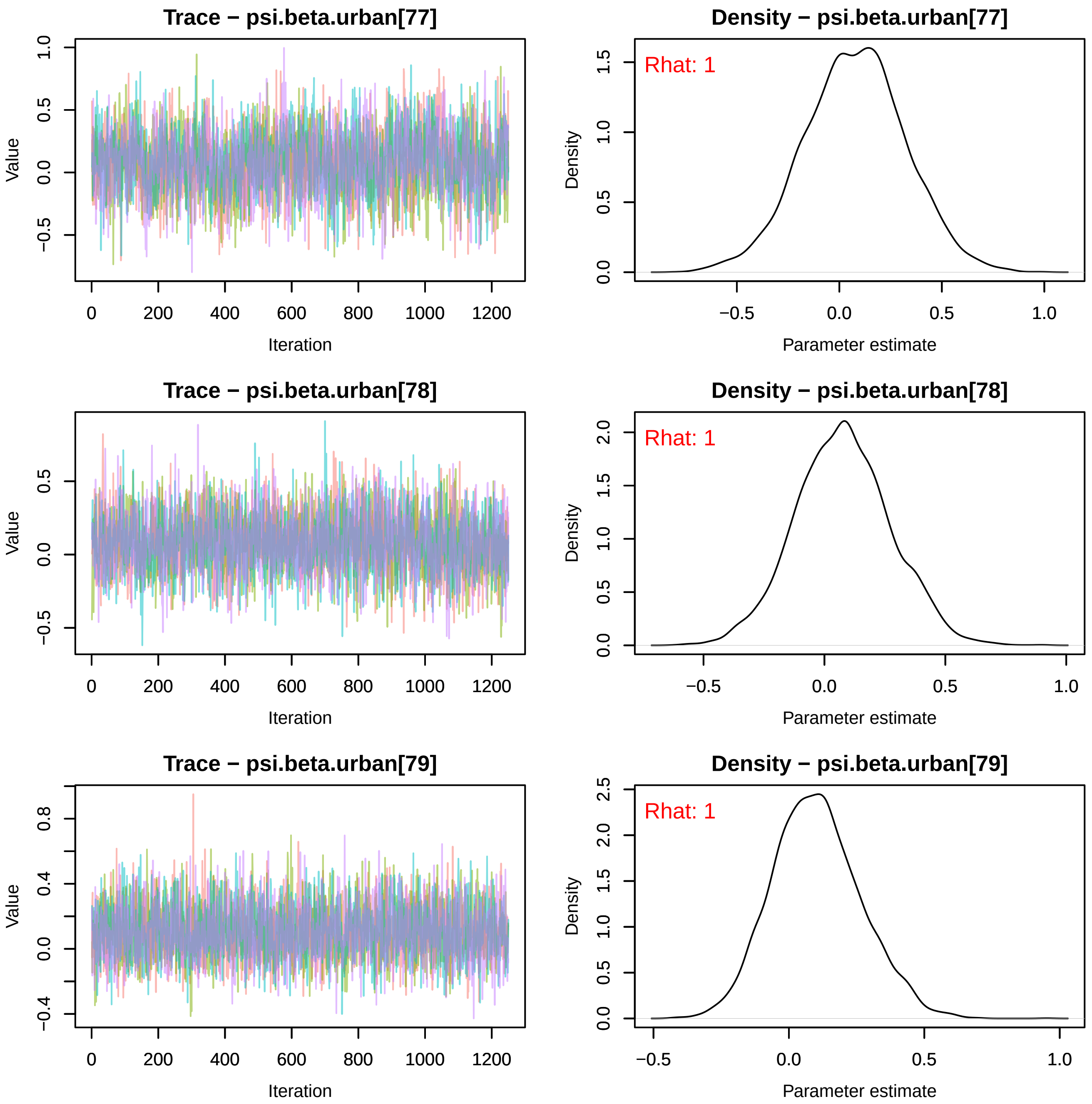
Trace − psi.beta.urban[77]
Value
Iteration
Density − psi.beta.urban[77]
Rhat: 1
Density
Parameter estimate
Trace − psi.beta.urban[78]
Value
Iteration
Density − psi.beta.urban[78]
Rhat: 1
Density
Parameter estimate
Trace − psi.beta.urban[79]
Value
Iteration
Density − psi.beta.urban[79]
Rhat: 1
Density
Parameter estimate

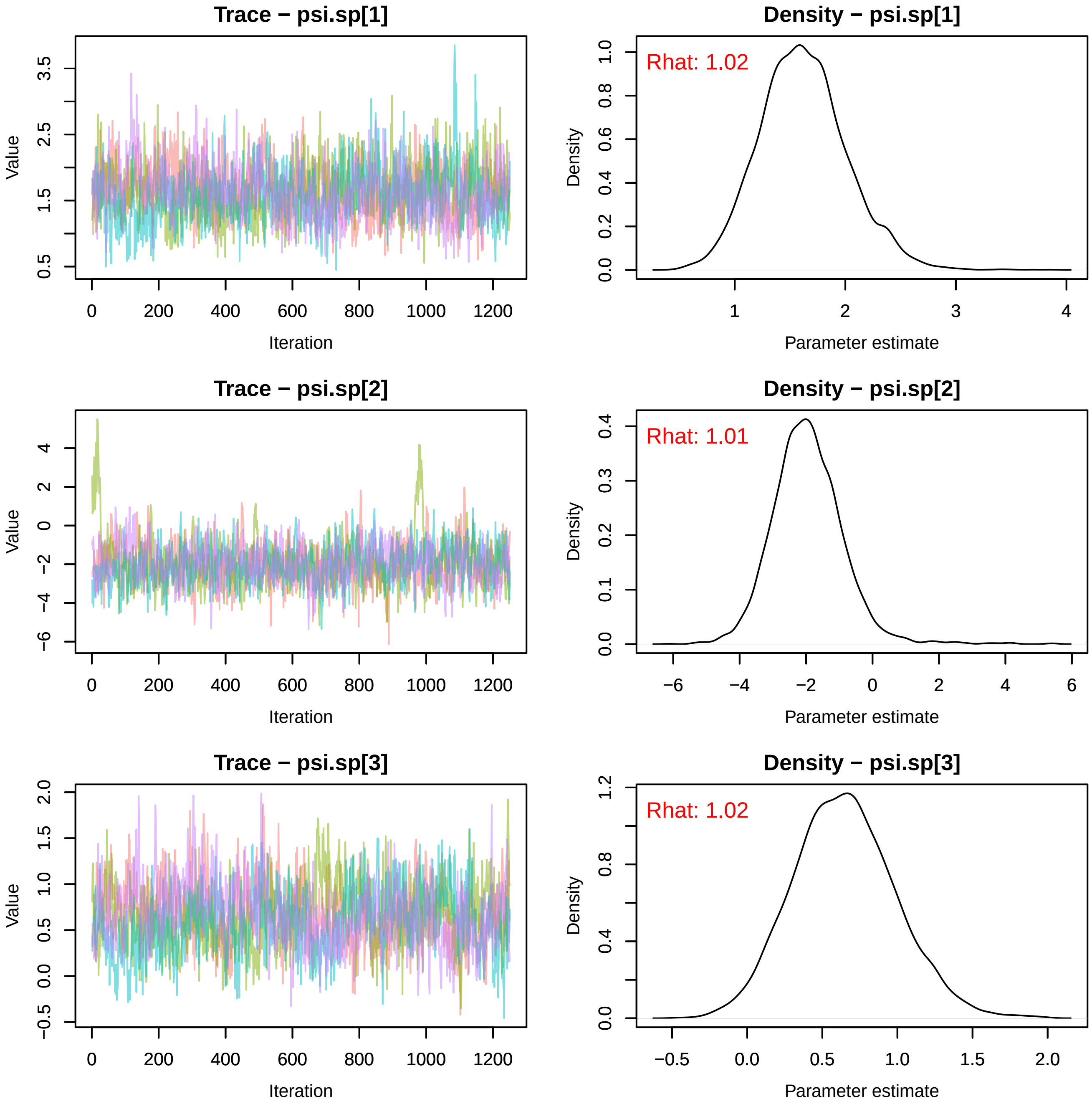
Trace − psi.sp[1]
Value
Iteration
Density − psi.sp[1]
Rhat: 1.02
Density
Parameter estimate
Trace − psi.sp[2]
Value
Iteration
Density − psi.sp[2]
Rhat: 1.01
Density
Parameter estimate
Trace − psi.sp[3]
Value
Iteration
Density − psi.sp[3]
Rhat: 1.02
Density
Parameter estimate

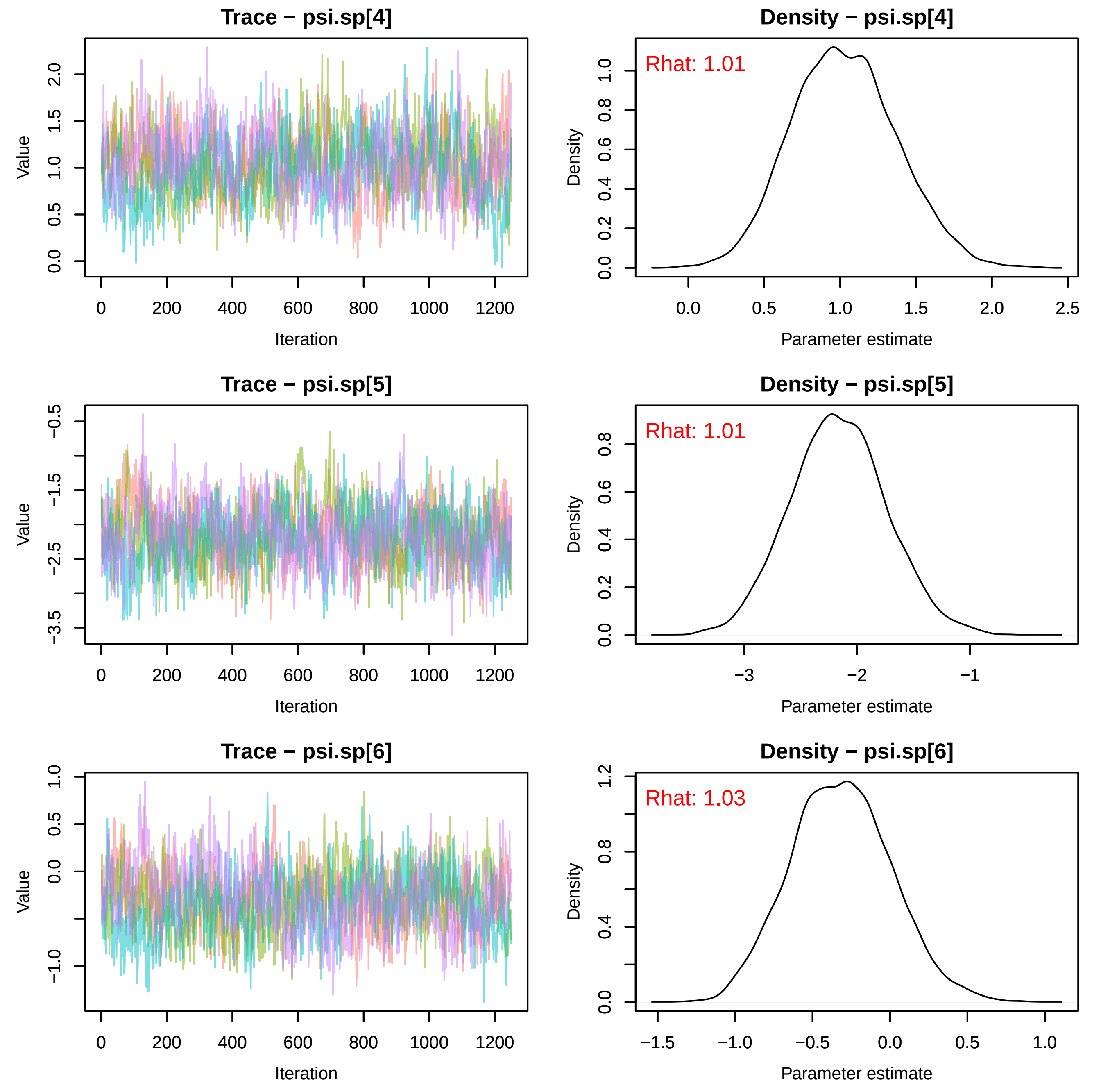

Trace − psi.sp[4]
Value
Iteration
Density − psi.sp[4]
Rhat: 1.01
Density
Parameter estimate
Trace − psi.sp[5]
Value
Iteration
Density − psi.sp[5]
Rhat: 1.01
Density
Parameter estimate
Trace − psi.sp[6]
Value
Iteration
Density − psi.sp[6]
Rhat: 1.03
Density
Parameter estimate

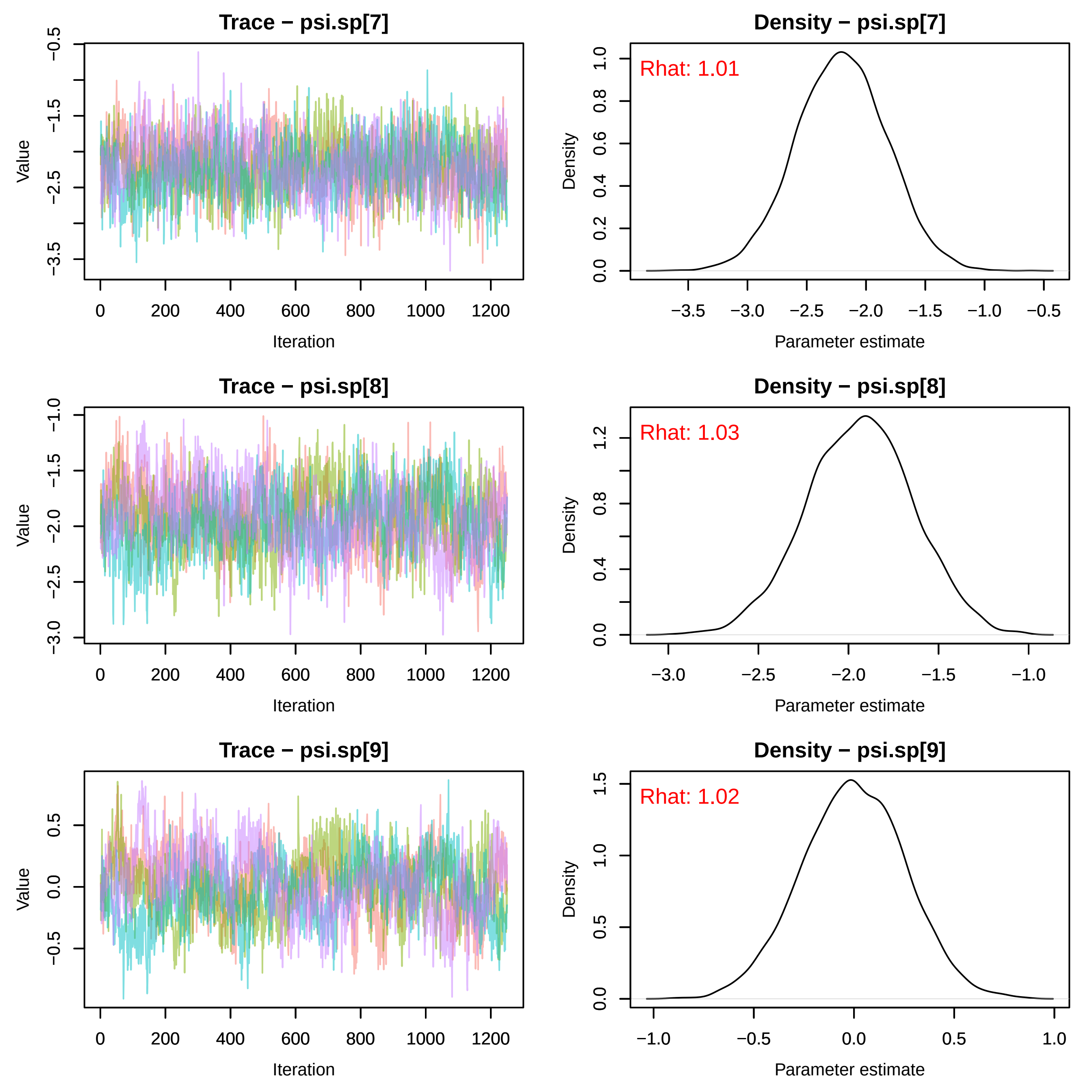

Trace − psi.sp[7]
Value
−0.5
−1.5
−2.5
−3.5
0
200
400
600
800
1000
1200
Iteration
Density − psi.sp[7]
Rhat: 1.01
Density
0.0
0.2
0.4
0.6
0.8
1.0
−3.5
−3.0
−2.5
−2.0
−1.5
−1.0
−0.5
Parameter estimate
Trace − psi.sp[8]
Value
−1.0
−1.5
−2.0
−2.5
−3.0
0
200
400
600
800
1000
1200
Iteration
Density − psi.sp[8]
Rhat: 1.03
Density
0.0
0.4
0.8
1.2
−3.0
−2.5
−2.0
−1.5
−1.0
Parameter estimate
Trace − psi.sp[9]
Value
0.5
0.0
−0.5
0
200
400
600
800
1000
1200
Iteration
Density − psi.sp[9]
Rhat: 1.02
Density
0.0
0.5
1.0
1.5
−1.0
−0.5
0.0
0.5
1.0
Parameter estimate

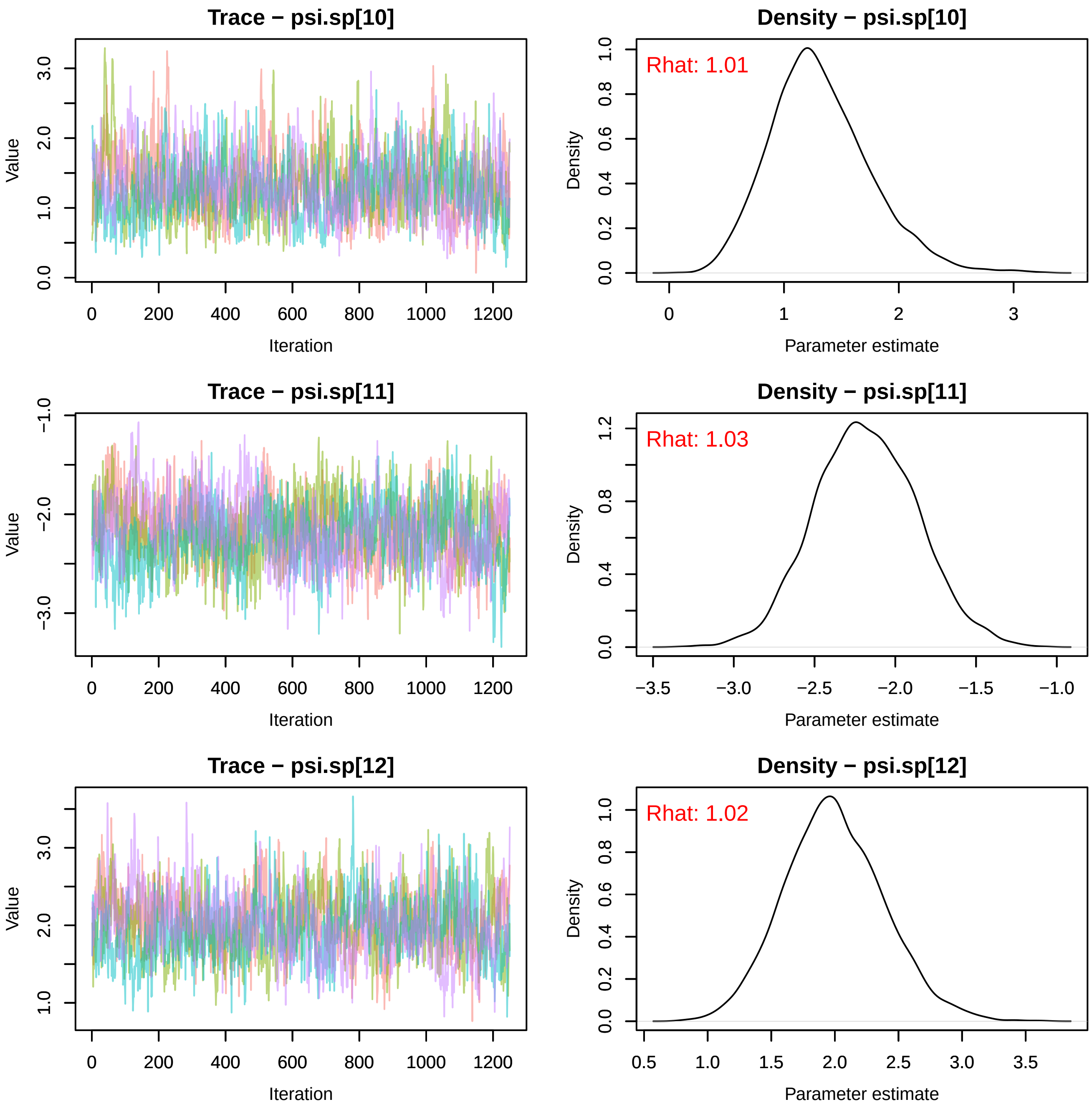
Trace − psi.sp[10]
Density − psi.sp[10]
Rhat: 1.01
Trace − psi.sp[11]
Density − psi.sp[11]
Rhat: 1.03
Trace − psi.sp[12]
Density − psi.sp[12]
Rhat: 1.02
Value
Iteration
Density
Parameter estimate
0
200
400
600
800
1000
1200

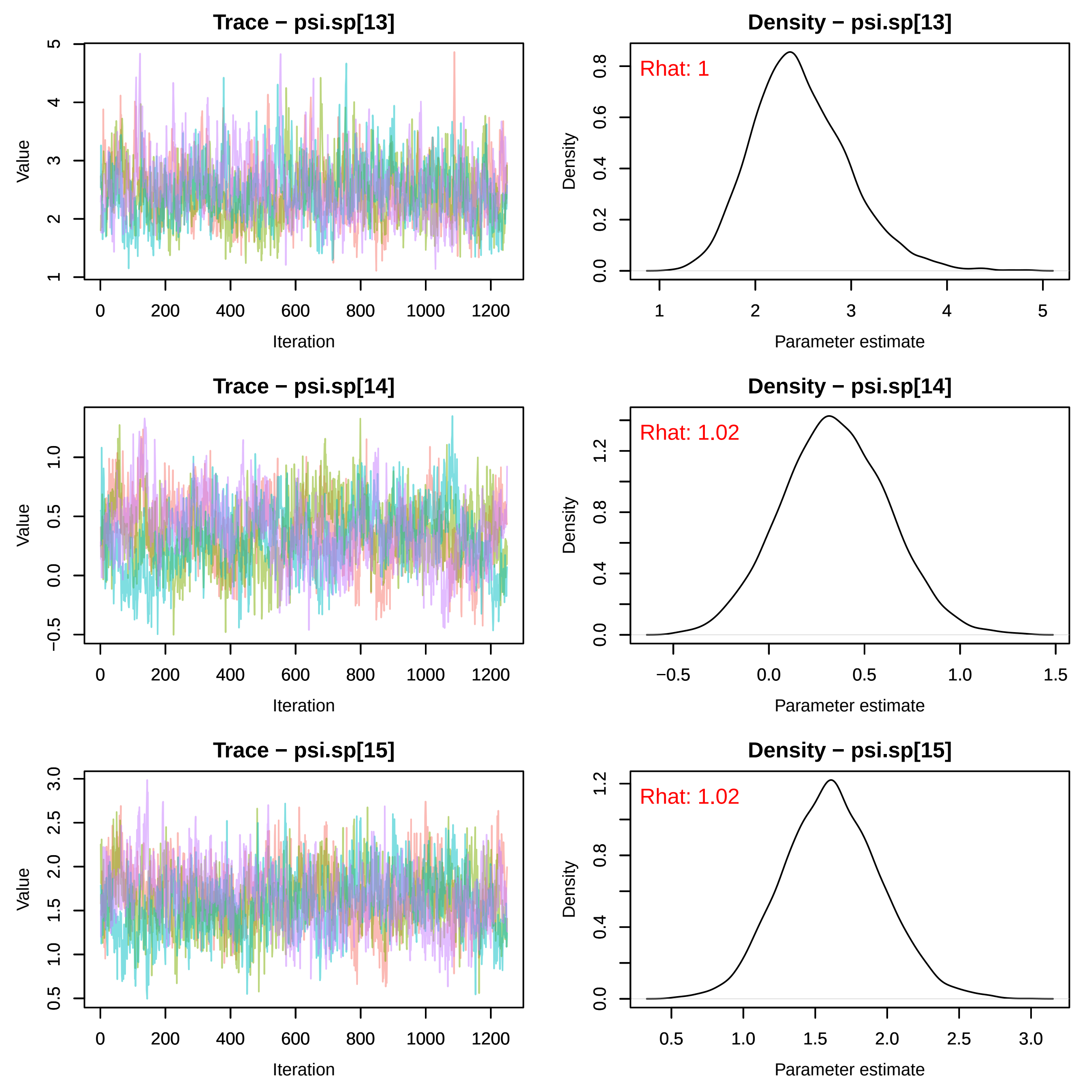
Trace − psi.sp[13]
Value
Iteration
0
200
400
600
800
1000
1200
1
2
3
4
5
Density − psi.sp[13]
Rhat: 1
Density
Parameter estimate
0.0
0.2
0.4
0.6
0.8
1
2
3
4
5
Trace − psi.sp[14]
Value
Iteration
−0.5
0.0
0.5
1.0
Density − psi.sp[14]
Rhat: 1.02
Density
Parameter estimate
0.0
0.4
0.8
1.2
−0.5
0.0
0.5
1.0
1.5
Trace − psi.sp[15]
Value
Iteration
0.5
1.0
1.5
2.0
2.5
3.0
Density − psi.sp[15]
Rhat: 1.02
Density
Parameter estimate
0.0
0.4
0.8
1.2
0.5
1.0
1.5
2.0
2.5
3.0

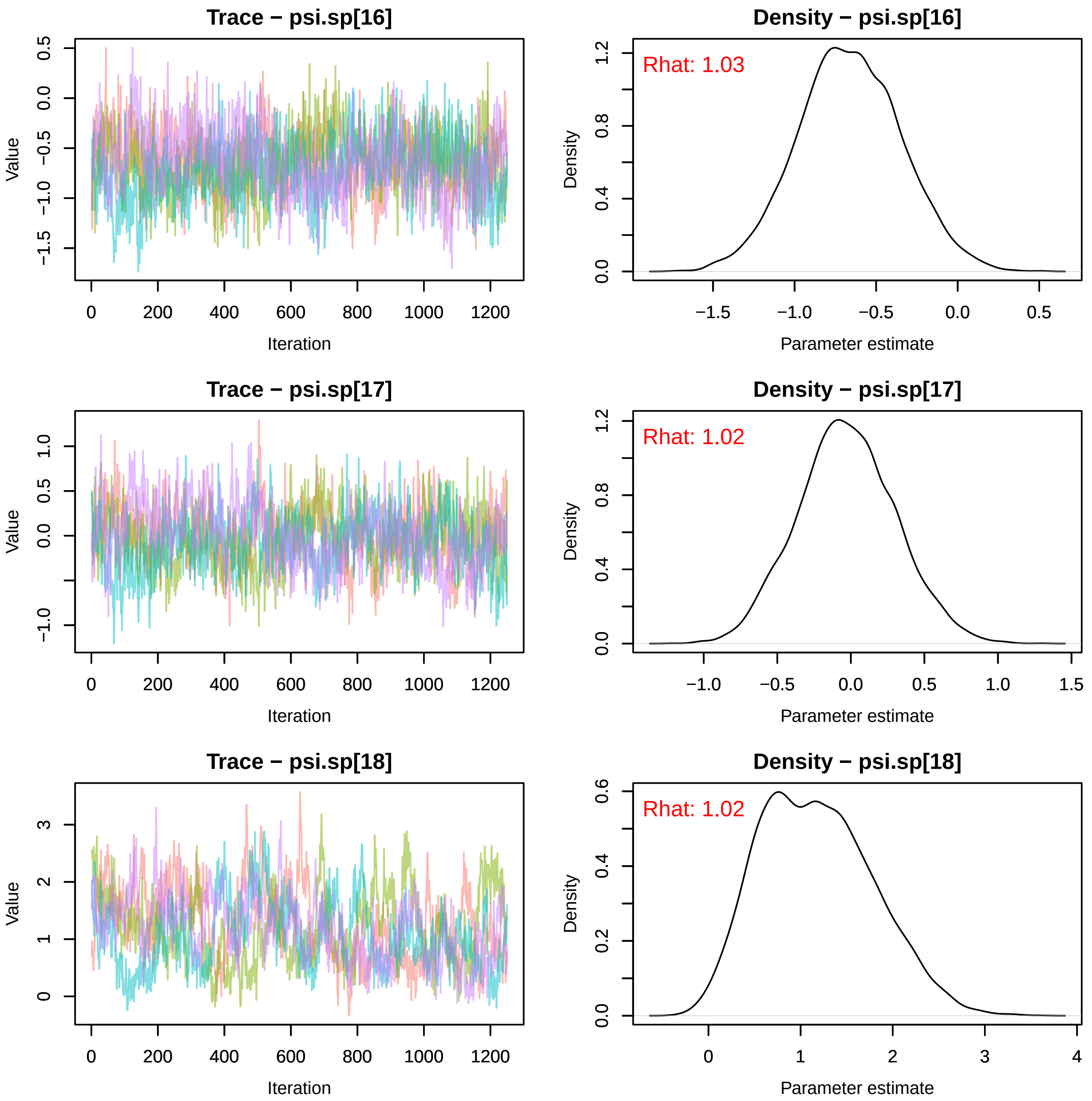
Trace − psi.sp[16]
Density − psi.sp[16]
Rhat: 1.03
Trace − psi.sp[17]
Density − psi.sp[17]
Rhat: 1.02
Trace − psi.sp[18]
Density − psi.sp[18]
Rhat: 1.02
Value
Iteration
Density
Parameter estimate

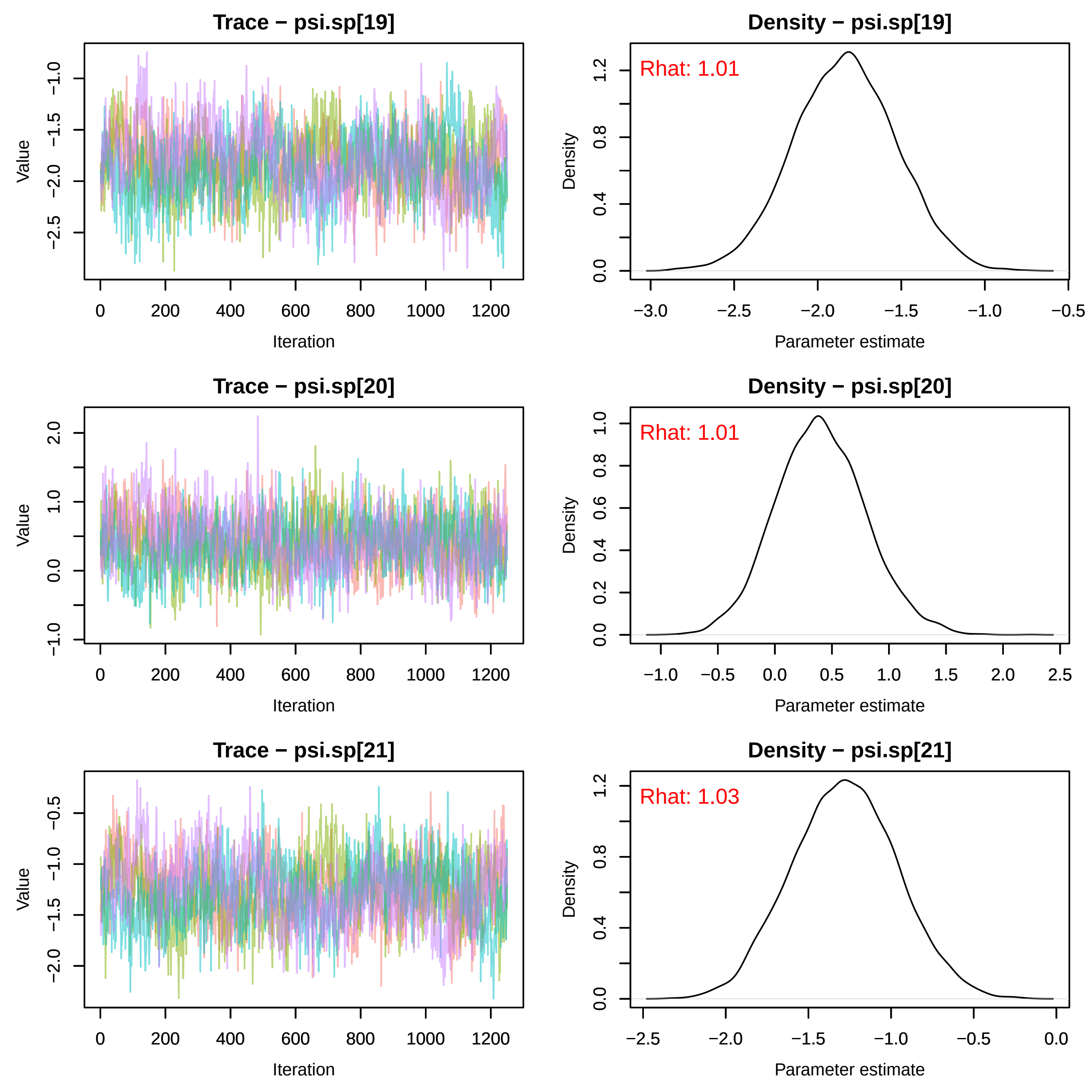

Trace − psi.sp[19]
Value
Iteration
Density − psi.sp[19]
Rhat: 1.01
Density
Parameter estimate
Trace − psi.sp[20]
Value
Iteration
Density − psi.sp[20]
Rhat: 1.01
Density
Parameter estimate
Trace − psi.sp[21]
Value
Iteration
Density − psi.sp[21]
Rhat: 1.03
Density
Parameter estimate

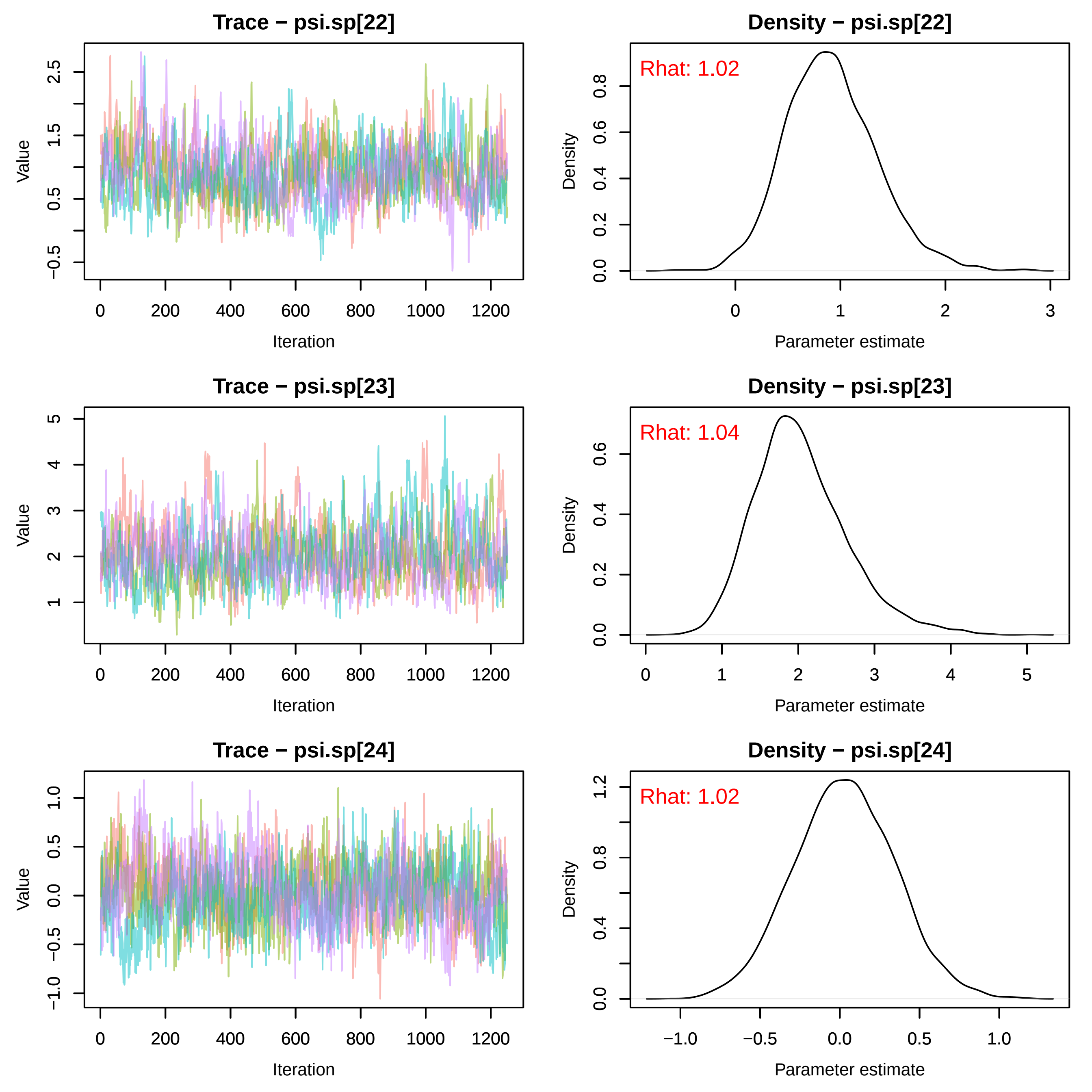
Trace − psi.sp[22]
Density − psi.sp[22]
Rhat: 1.02
Trace − psi.sp[23]
Density − psi.sp[23]
Rhat: 1.04
Trace − psi.sp[24]
Density − psi.sp[24]
Rhat: 1.02
Value
Iteration
Density
Parameter estimate

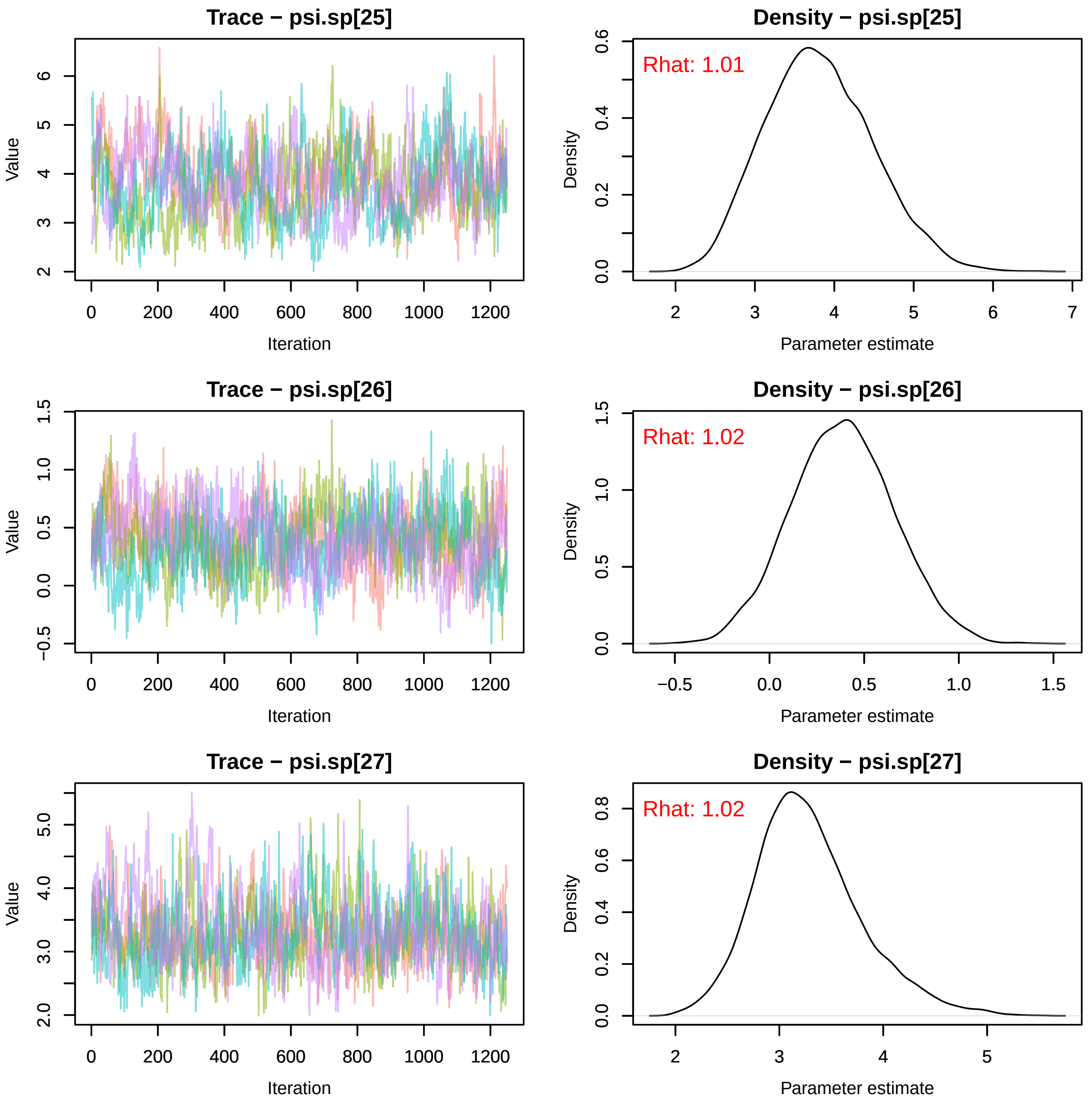
Trace − psi.sp[25]
Density − psi.sp[25]
Rhat: 1.01
Trace − psi.sp[26]
Density − psi.sp[26]
Rhat: 1.02
Trace − psi.sp[27]
Density − psi.sp[27]
Rhat: 1.02
Value
Iteration
Density
Parameter estimate

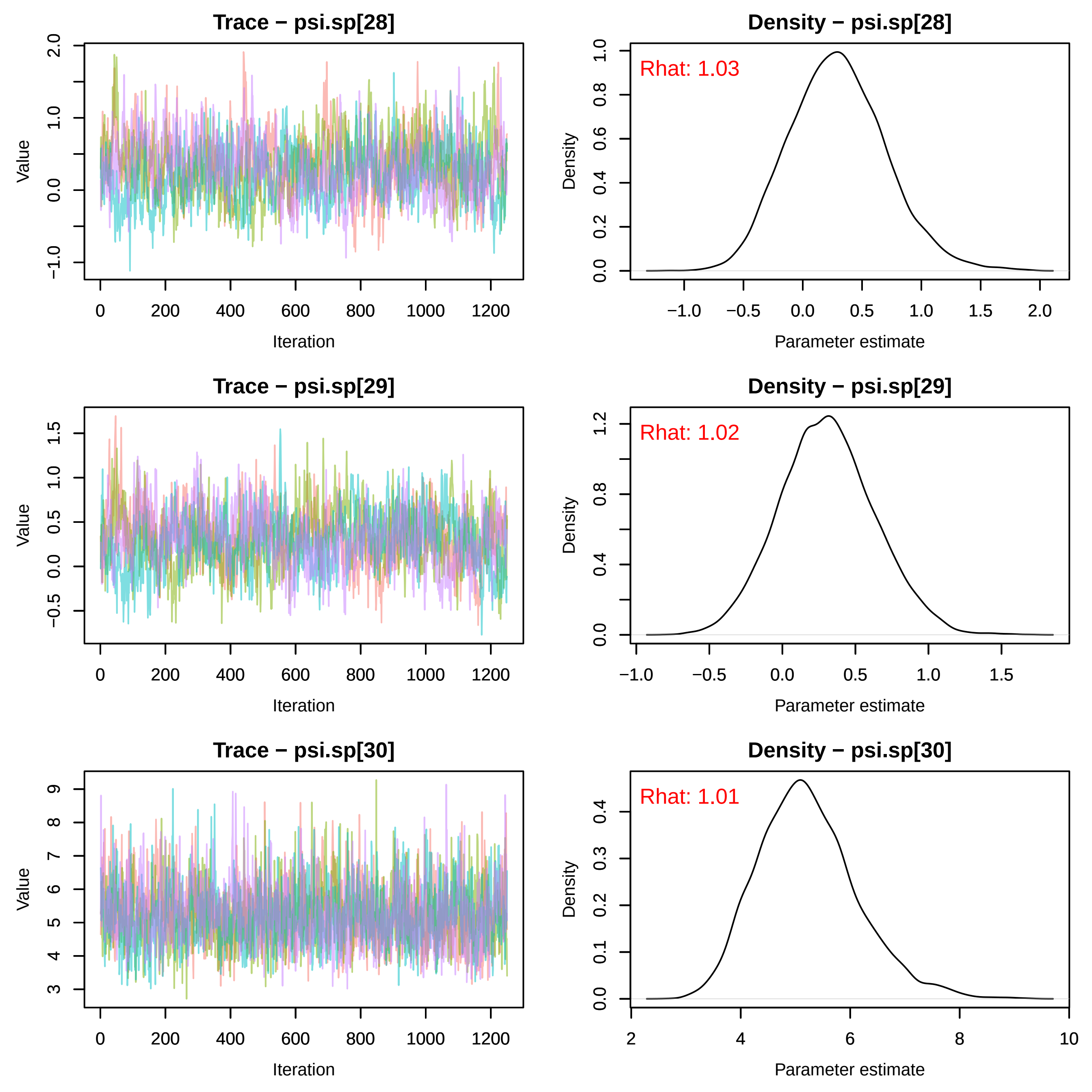

Trace − psi.sp[28]
Value
Iteration
Density − psi.sp[28]
Rhat: 1.03
Density
Parameter estimate
Trace − psi.sp[29]
Value
Iteration
Density − psi.sp[29]
Rhat: 1.02
Density
Parameter estimate
Trace − psi.sp[30]
Value
Iteration
Density − psi.sp[30]
Rhat: 1.01
Density
Parameter estimate

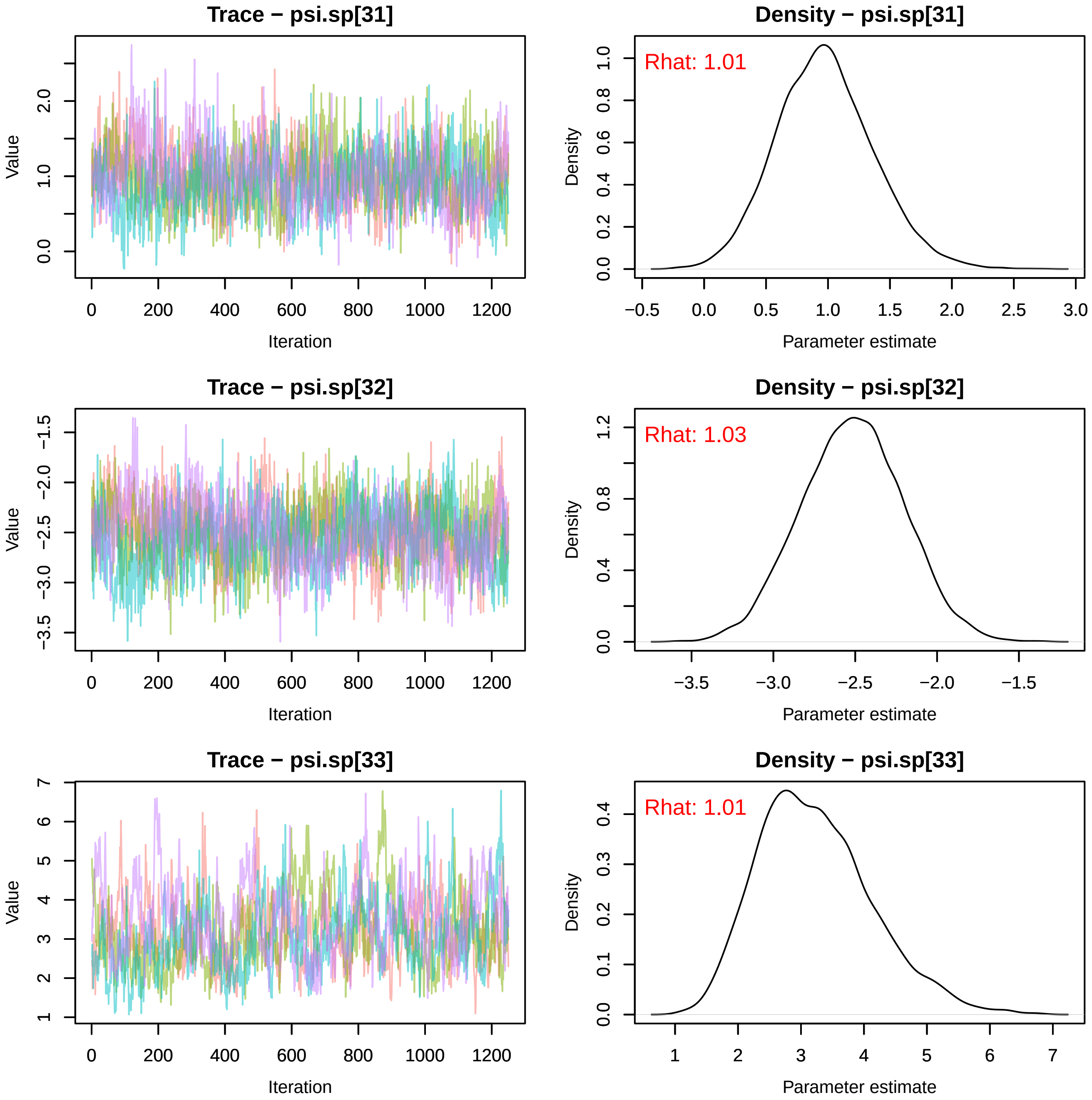
Trace − psi.sp[31]
Value
Iteration
Density − psi.sp[31]
Rhat: 1.01
Density
Parameter estimate
Trace − psi.sp[32]
Value
Iteration
Density − psi.sp[32]
Rhat: 1.03
Density
Parameter estimate
Trace − psi.sp[33]
Value
Iteration
Density − psi.sp[33]
Rhat: 1.01
Density
Parameter estimate

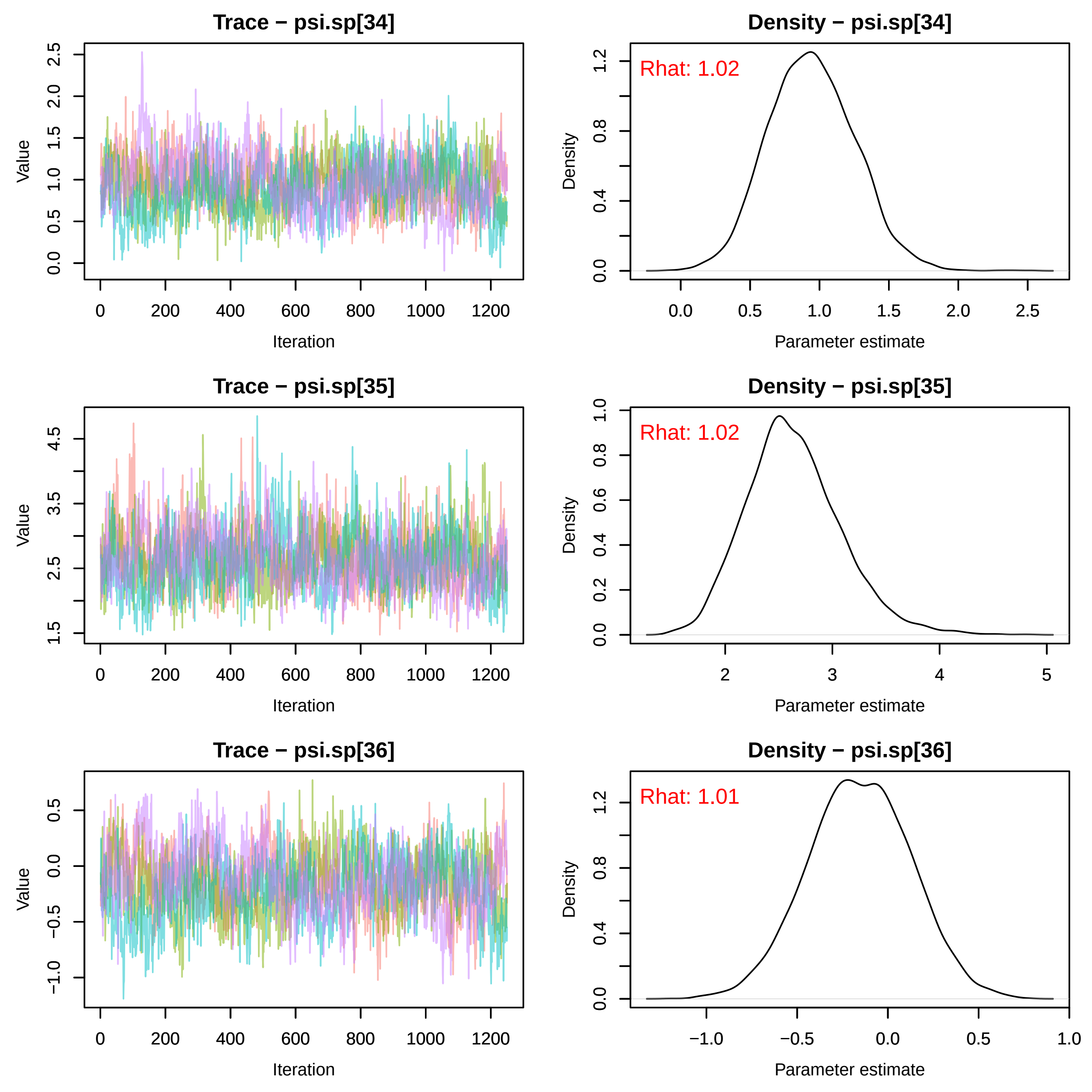
Trace − psi.sp[34]
Density − psi.sp[34]
Rhat: 1.02
Trace − psi.sp[35]
Density − psi.sp[35]
Rhat: 1.02
Trace − psi.sp[36]
Density − psi.sp[36]
Rhat: 1.01
Value
Iteration
Density
Parameter estimate

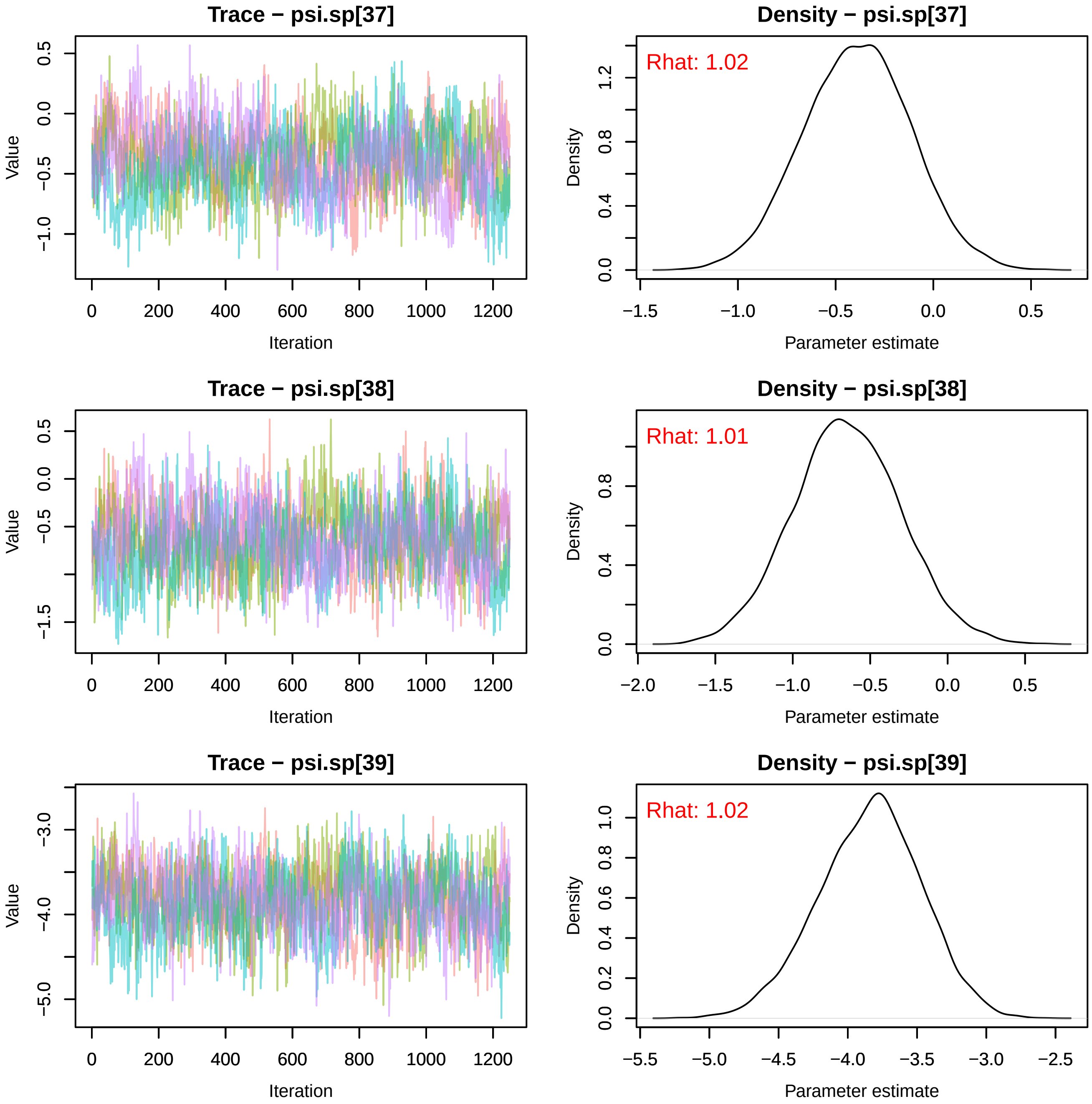
Trace − psi.sp[37]
Value
Iteration
Density − psi.sp[37]
Rhat: 1.02
Density
Parameter estimate
Trace − psi.sp[38]
Value
Iteration
Density − psi.sp[38]
Rhat: 1.01
Density
Parameter estimate
Trace − psi.sp[39]
Value
Iteration
Density − psi.sp[39]
Rhat: 1.02
Density
Parameter estimate

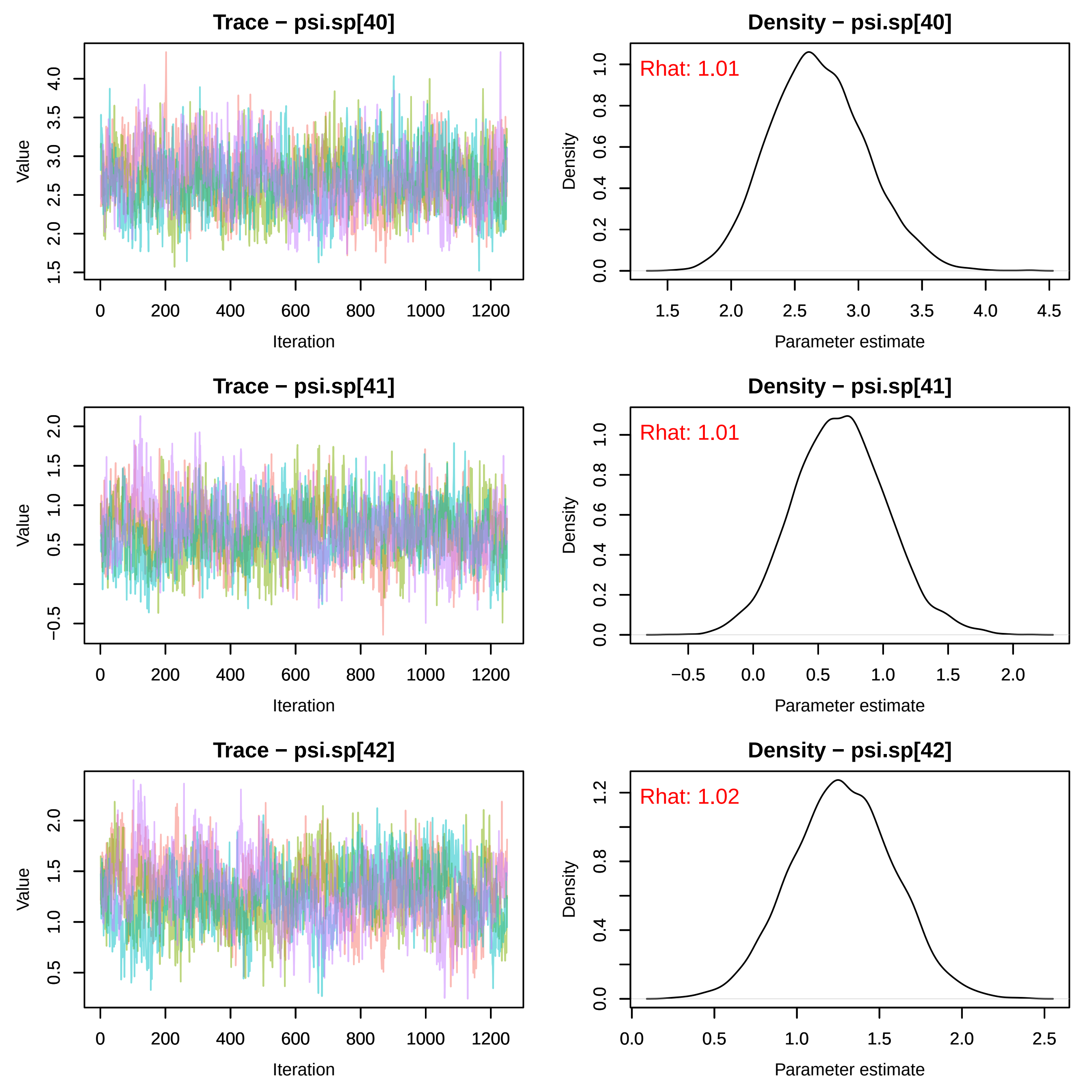
Trace − psi.sp[40]
Value
Iteration
Density − psi.sp[40]
Rhat: 1.01
Density
Parameter estimate
Trace − psi.sp[41]
Value
Iteration
Density − psi.sp[41]
Rhat: 1.01
Density
Parameter estimate
Trace − psi.sp[42]
Value
Iteration
Density − psi.sp[42]
Rhat: 1.02
Density
Parameter estimate

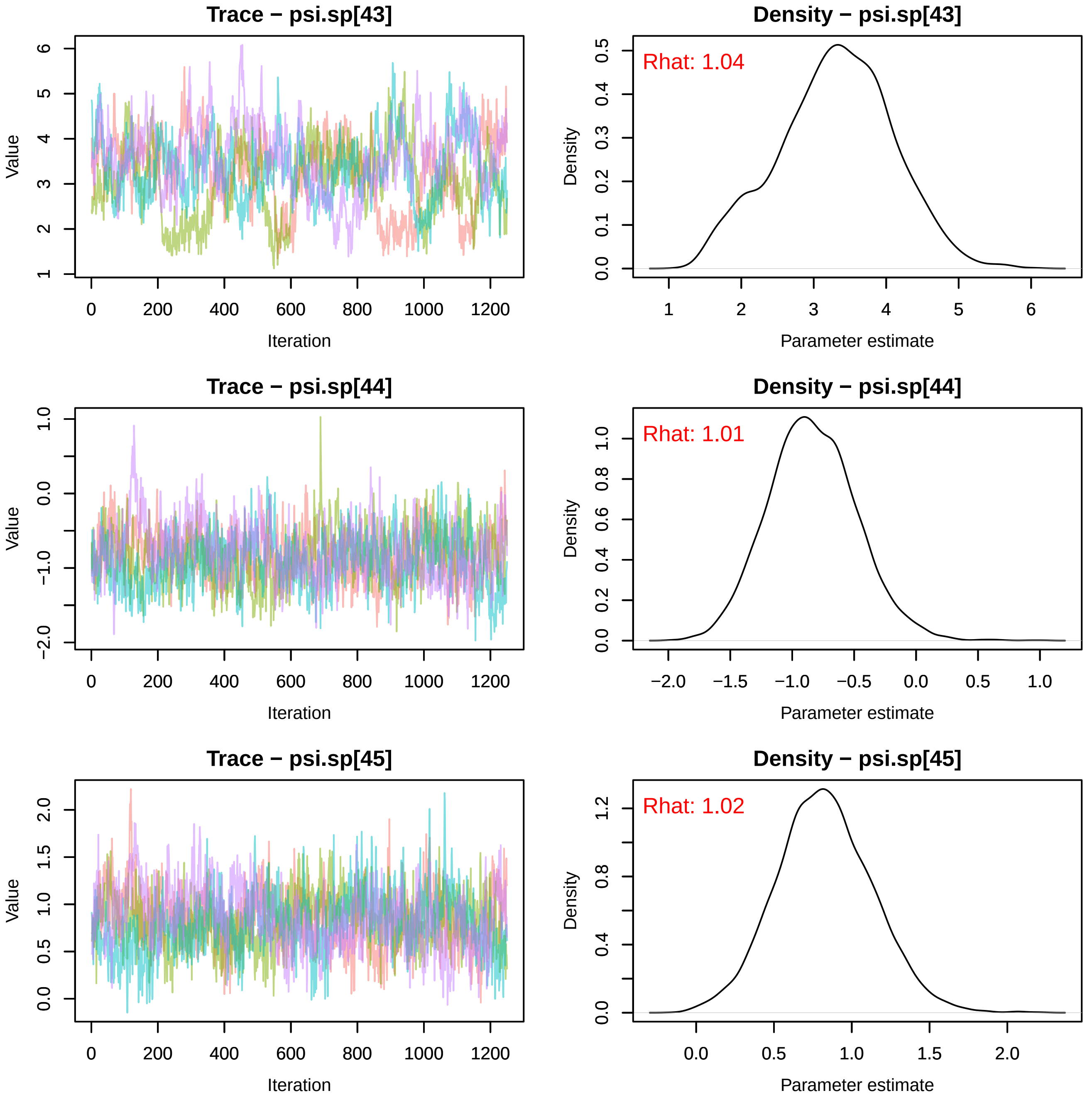

Trace − psi.sp[43]
Value
Iteration
Density − psi.sp[43]
Rhat: 1.04
Density
Parameter estimate
Trace − psi.sp[44]
Value
Iteration
Density − psi.sp[44]
Rhat: 1.01
Density
Parameter estimate
Trace − psi.sp[45]
Value
Iteration
Density − psi.sp[45]
Rhat: 1.02
Density
Parameter estimate

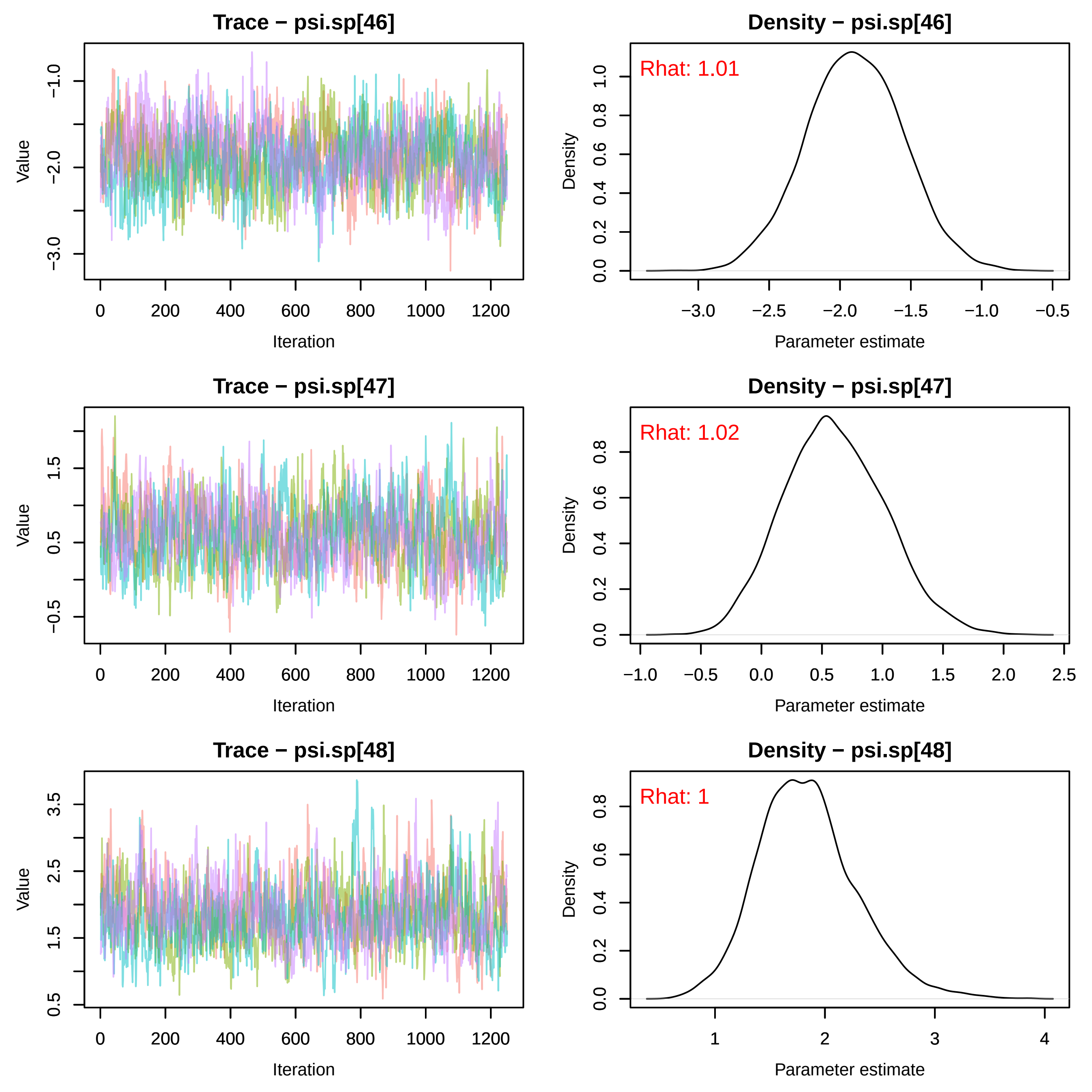
Trace − psi.sp[46]
Value
Iteration
Density − psi.sp[46]
Rhat: 1.01
Density
Parameter estimate
Trace − psi.sp[47]
Value
Iteration
Density − psi.sp[47]
Rhat: 1.02
Density
Parameter estimate
Trace − psi.sp[48]
Value
Iteration
Density − psi.sp[48]
Rhat: 1
Density
Parameter estimate

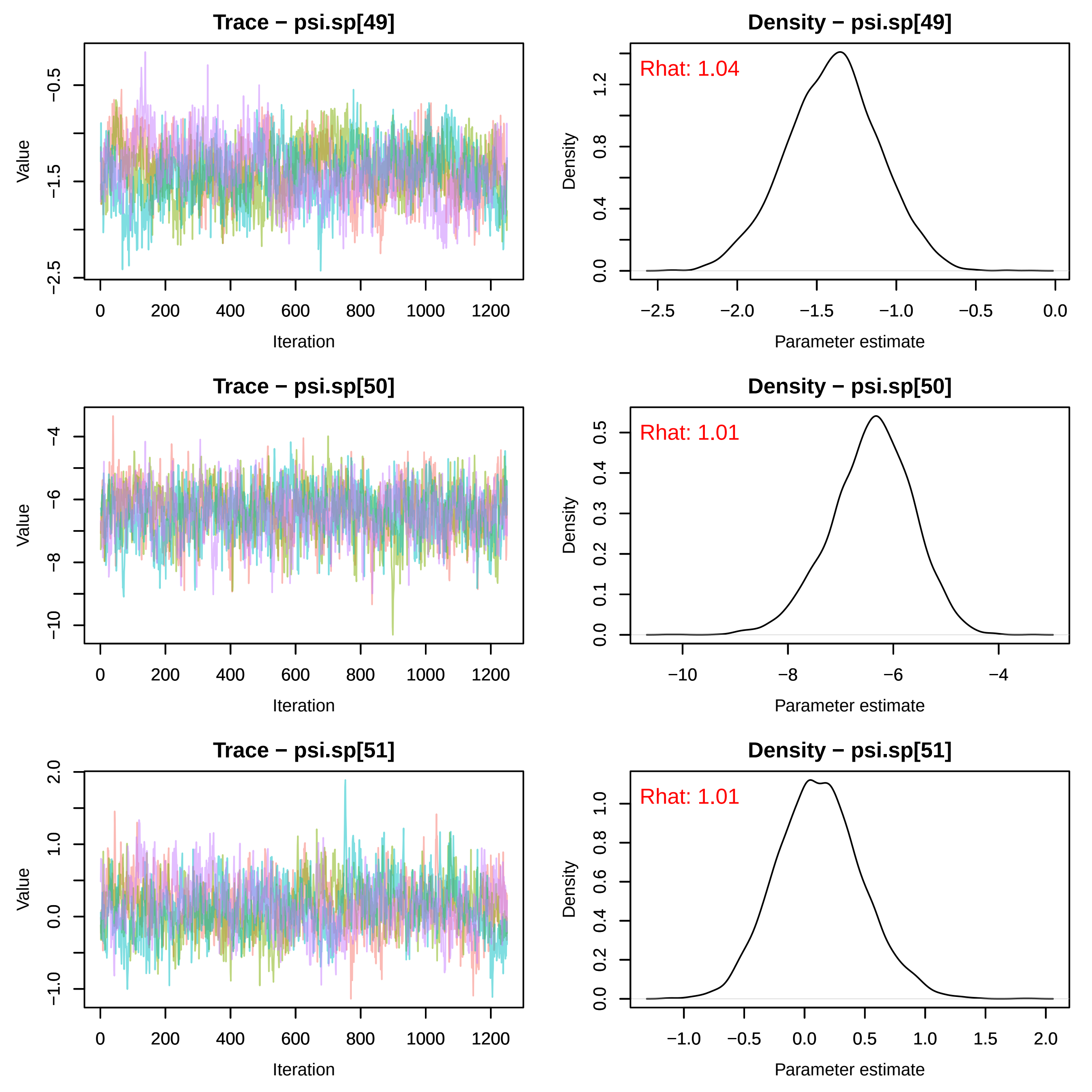
Trace − psi.sp[49]
Value
−2.5
−1.5
−0.5
0
200
400
600
800
1000
1200
Iteration
Density − psi.sp[49]
Rhat: 1.04
Density
0.0
0.4
0.8
1.2
−2.5
−2.0
−1.5
−1.0
−0.5
0.0
Parameter estimate
Trace − psi.sp[50]
Value
−10
−8
−6
−4
0
200
400
600
800
1000
1200
Iteration
Density − psi.sp[50]
Rhat: 1.01
Density
0.0
0.1
0.2
0.3
0.4
0.5
−10
−8
−6
−4
Parameter estimate
Trace − psi.sp[51]
Value
−1.0
0.0
1.0
2.0
0
200
400
600
800
1000
1200
Iteration
Density − psi.sp[51]
Rhat: 1.01
Density
0.0
0.2
0.4
0.6
0.8
1.0
−1.0
−0.5
0.0
0.5
1.0
1.5
2.0
Parameter estimate

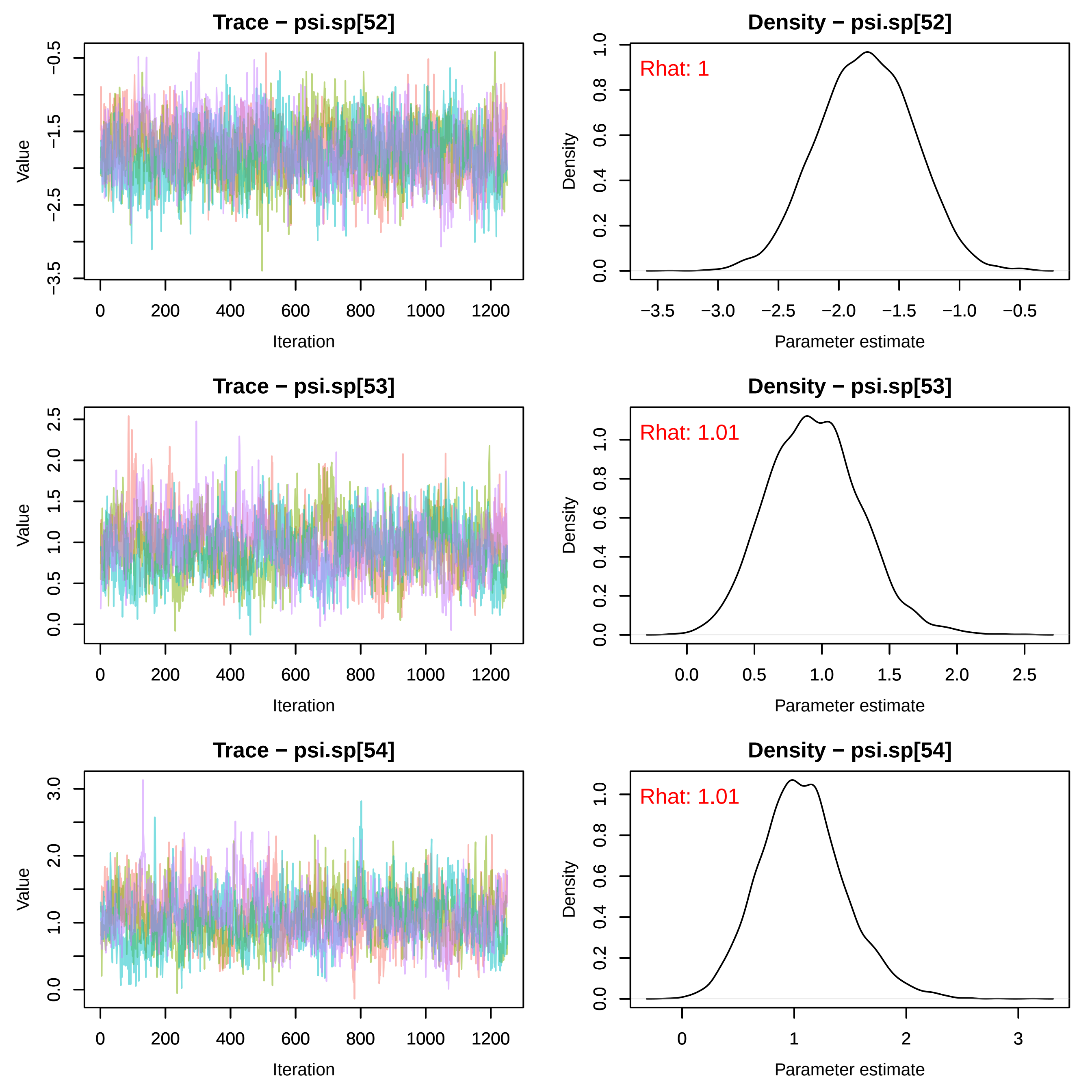

Trace − psi.sp[52]
Value
Iteration
Density − psi.sp[52]
Rhat: 1
Density
Parameter estimate
Trace − psi.sp[53]
Value
Iteration
Density − psi.sp[53]
Rhat: 1.01
Density
Parameter estimate
Trace − psi.sp[54]
Value
Iteration
Density − psi.sp[54]
Rhat: 1.01
Density
Parameter estimate

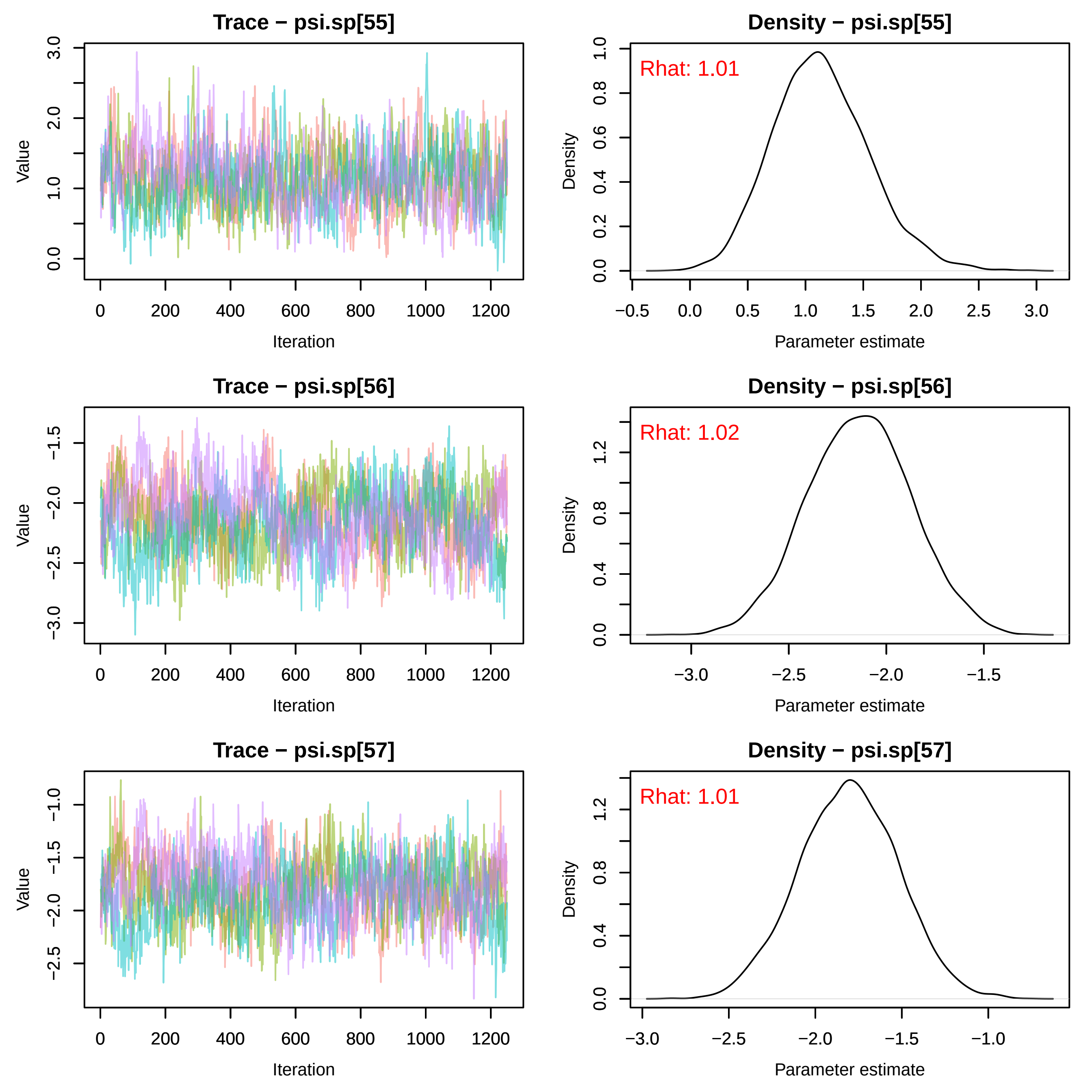
Trace − psi.sp[55]
Value
Iteration
Density − psi.sp[55]
Rhat: 1.01
Density
Parameter estimate
Trace − psi.sp[56]
Value
Iteration
Density − psi.sp[56]
Rhat: 1.02
Density
Parameter estimate
Trace − psi.sp[57]
Value
Iteration
Density − psi.sp[57]
Rhat: 1.01
Density
Parameter estimate

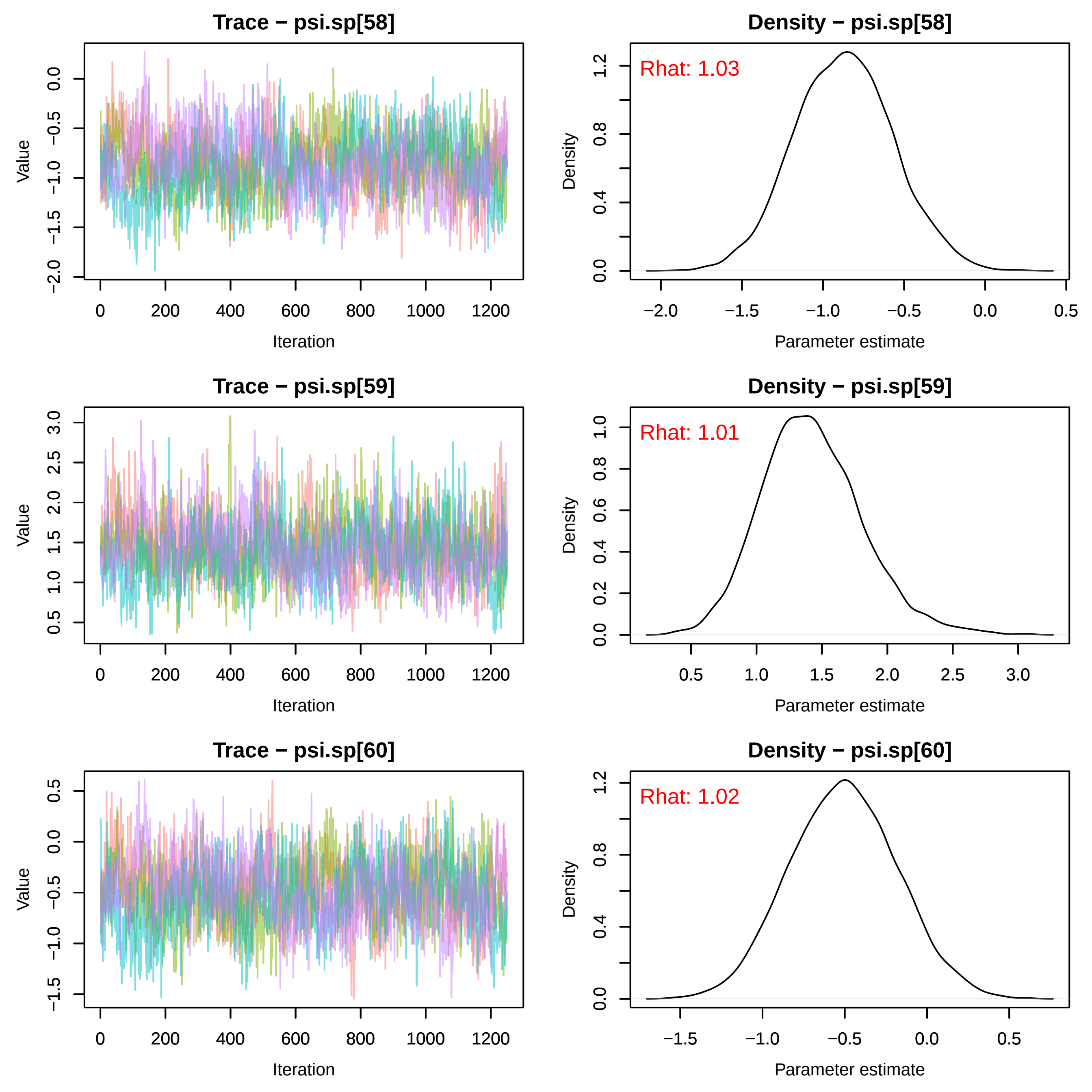
Trace − psi.sp[58]
Value
Iteration
Density − psi.sp[58]
Rhat: 1.03
Density
Parameter estimate
Trace − psi.sp[59]
Value
Iteration
Density − psi.sp[59]
Rhat: 1.01
Density
Parameter estimate
Trace − psi.sp[60]
Value
Iteration
Density − psi.sp[60]
Rhat: 1.02
Density
Parameter estimate

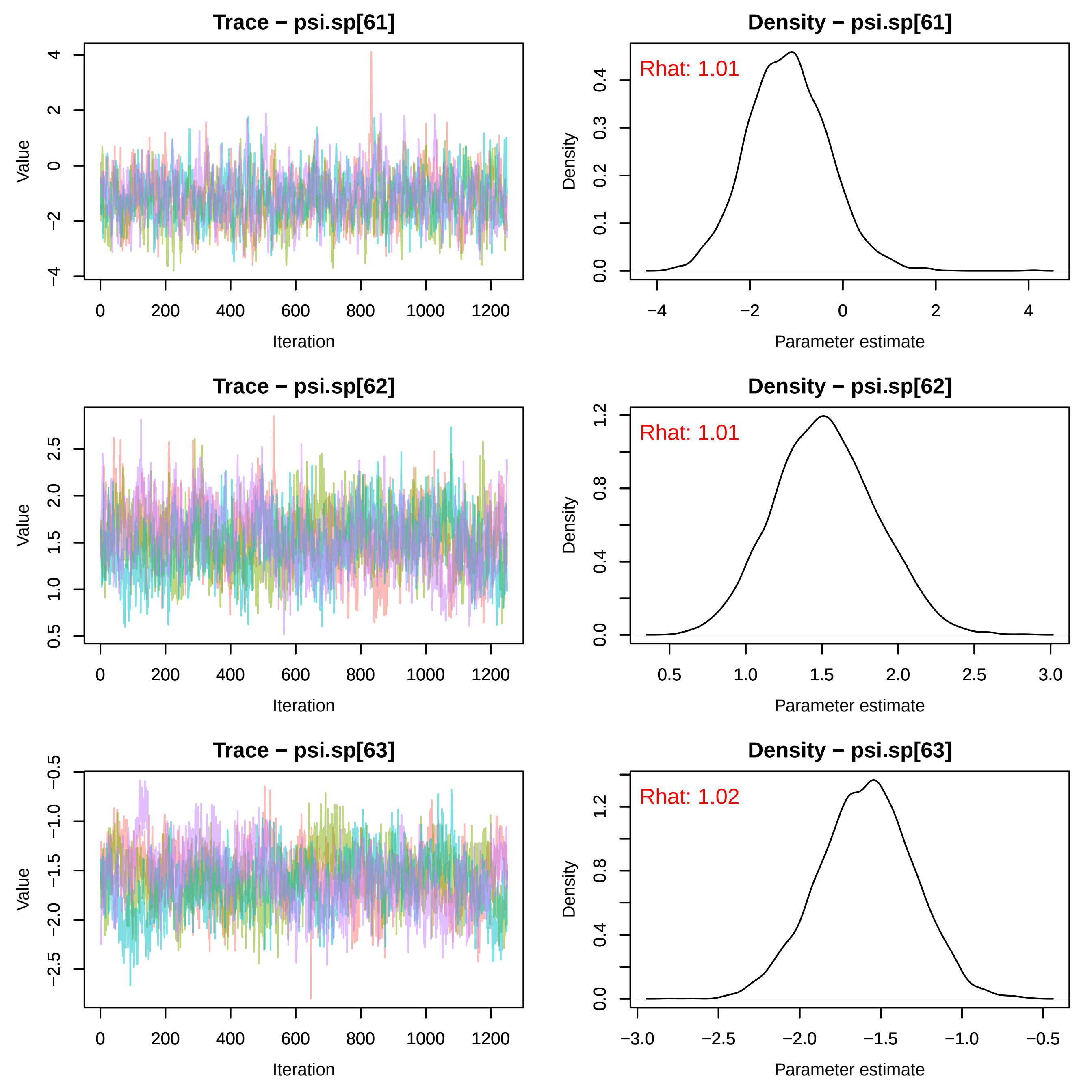
Trace − psi.sp[61]
Value
Iteration
Density − psi.sp[61]
Rhat: 1.01
Density
Parameter estimate
Trace − psi.sp[62]
Value
Iteration
Density − psi.sp[62]
Rhat: 1.01
Density
Parameter estimate
Trace − psi.sp[63]
Value
Iteration
Density − psi.sp[63]
Rhat: 1.02
Density
Parameter estimate

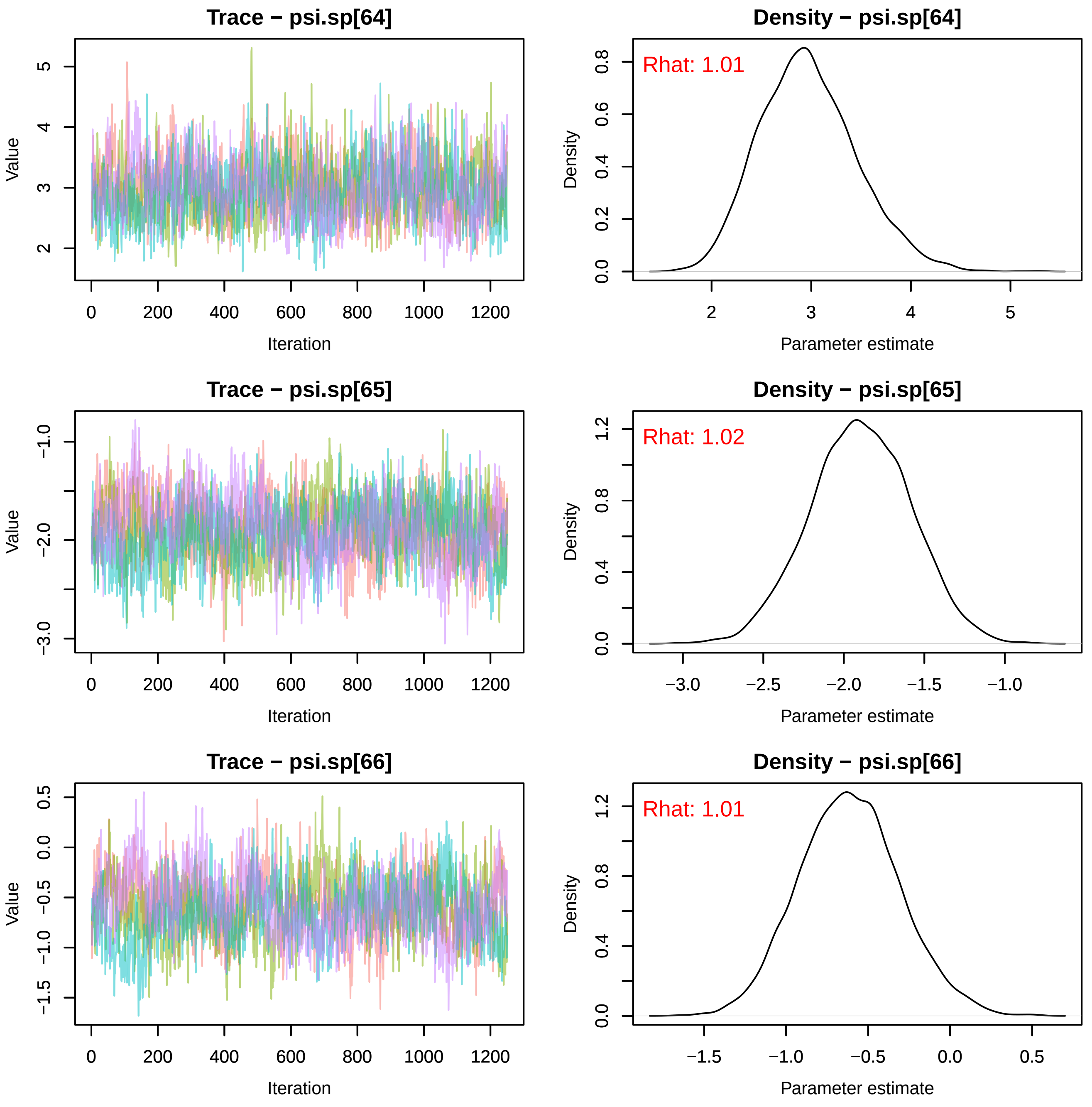

Trace − psi.sp[64]
Density − psi.sp[64]
Rhat: 1.01
Trace − psi.sp[65]
Density − psi.sp[65]
Rhat: 1.02
Trace − psi.sp[66]
Density − psi.sp[66]
Rhat: 1.01
Value
Iteration
Density
Parameter estimate

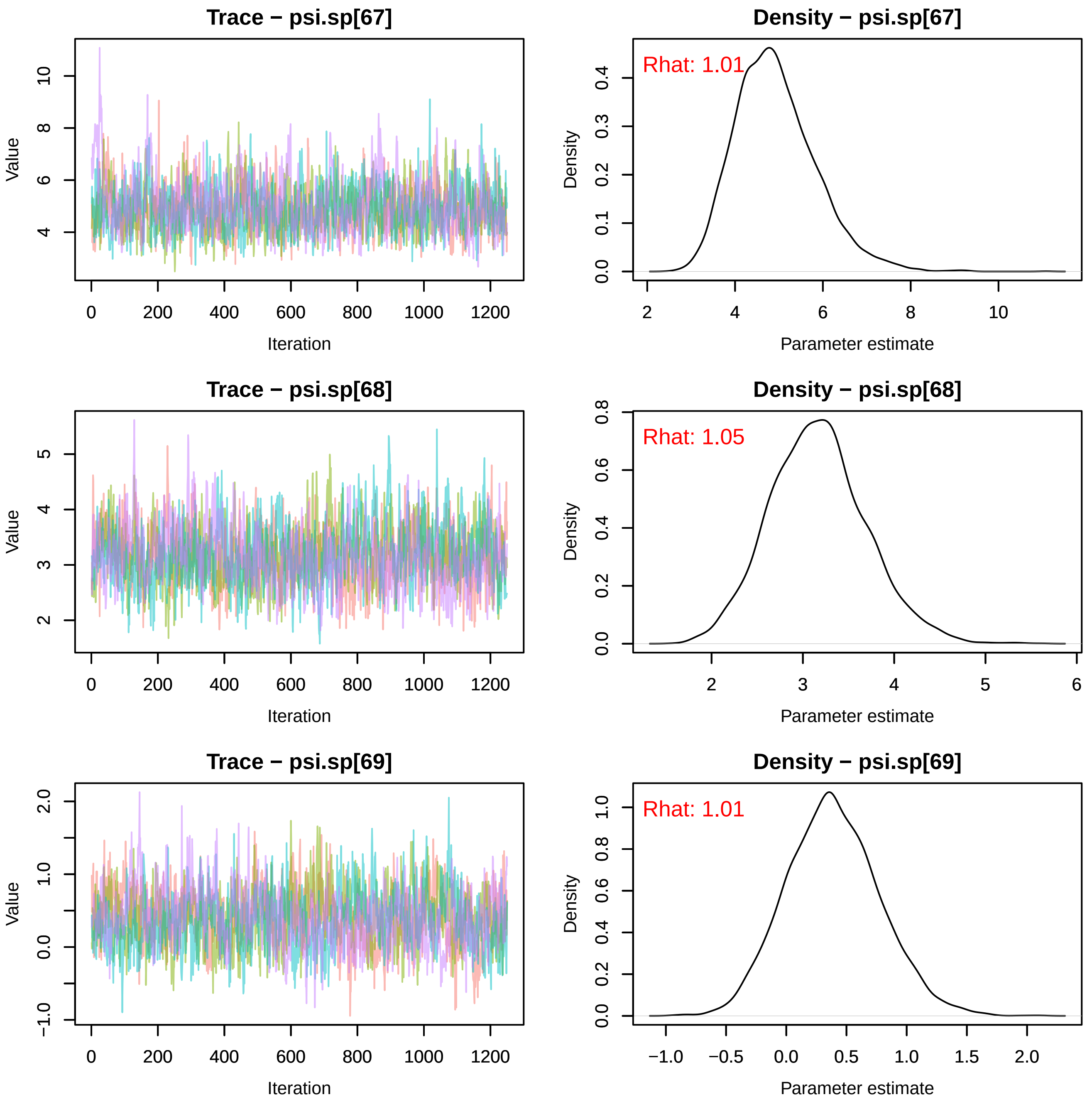

Trace − psi.sp[67]
Value
Iteration
Density − psi.sp[67]
Rhat: 1.01
Density
Parameter estimate
Trace − psi.sp[68]
Value
Iteration
Density − psi.sp[68]
Rhat: 1.05
Density
Parameter estimate
Trace − psi.sp[69]
Value
Iteration
Density − psi.sp[69]
Rhat: 1.01
Density
Parameter estimate

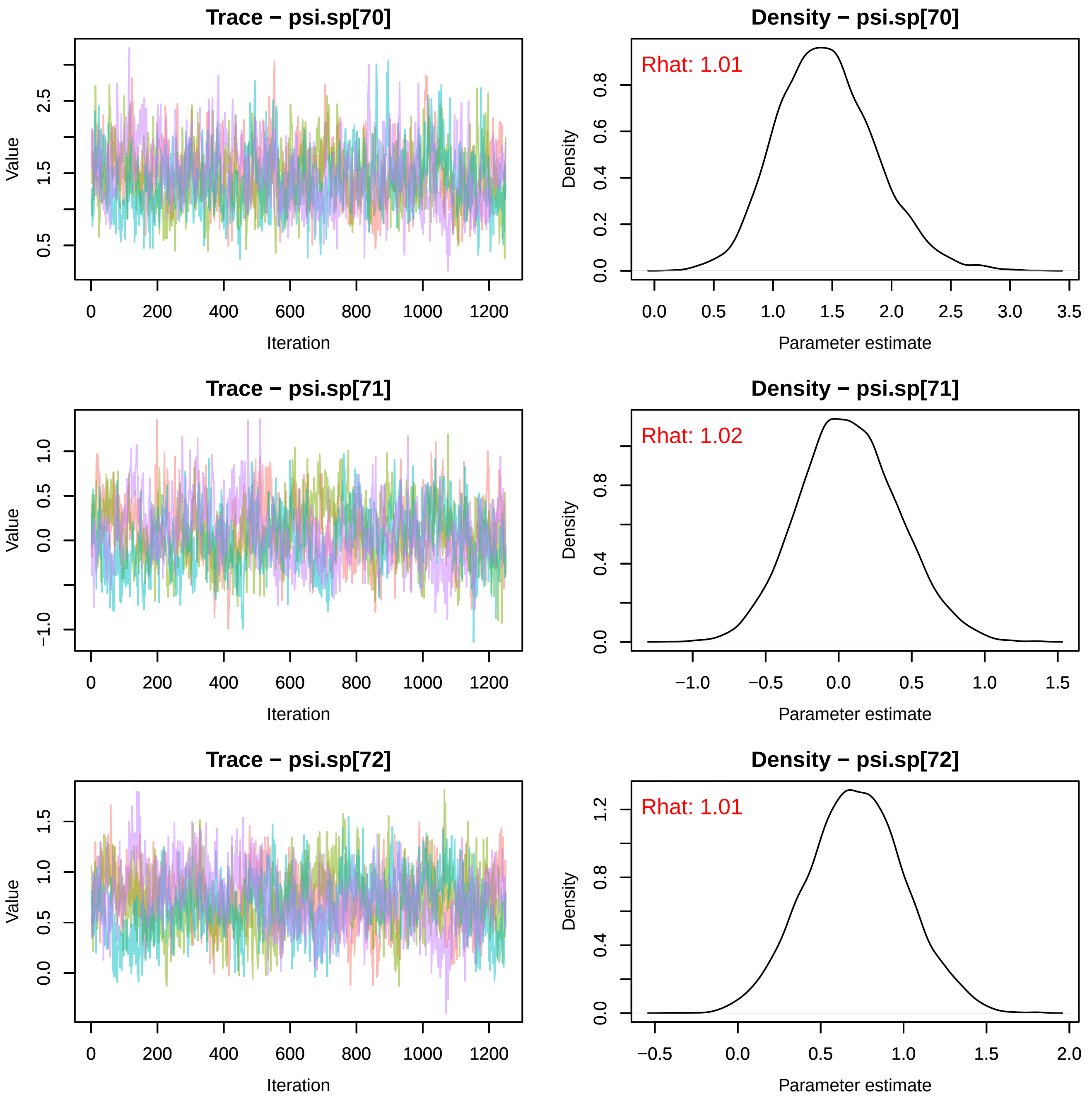
Trace − psi.sp[70]
Density − psi.sp[70]
Rhat: 1.01
Trace − psi.sp[71]
Density − psi.sp[71]
Rhat: 1.02
Trace − psi.sp[72]
Density − psi.sp[72]
Rhat: 1.01
Value
Iteration
Density
Parameter estimate

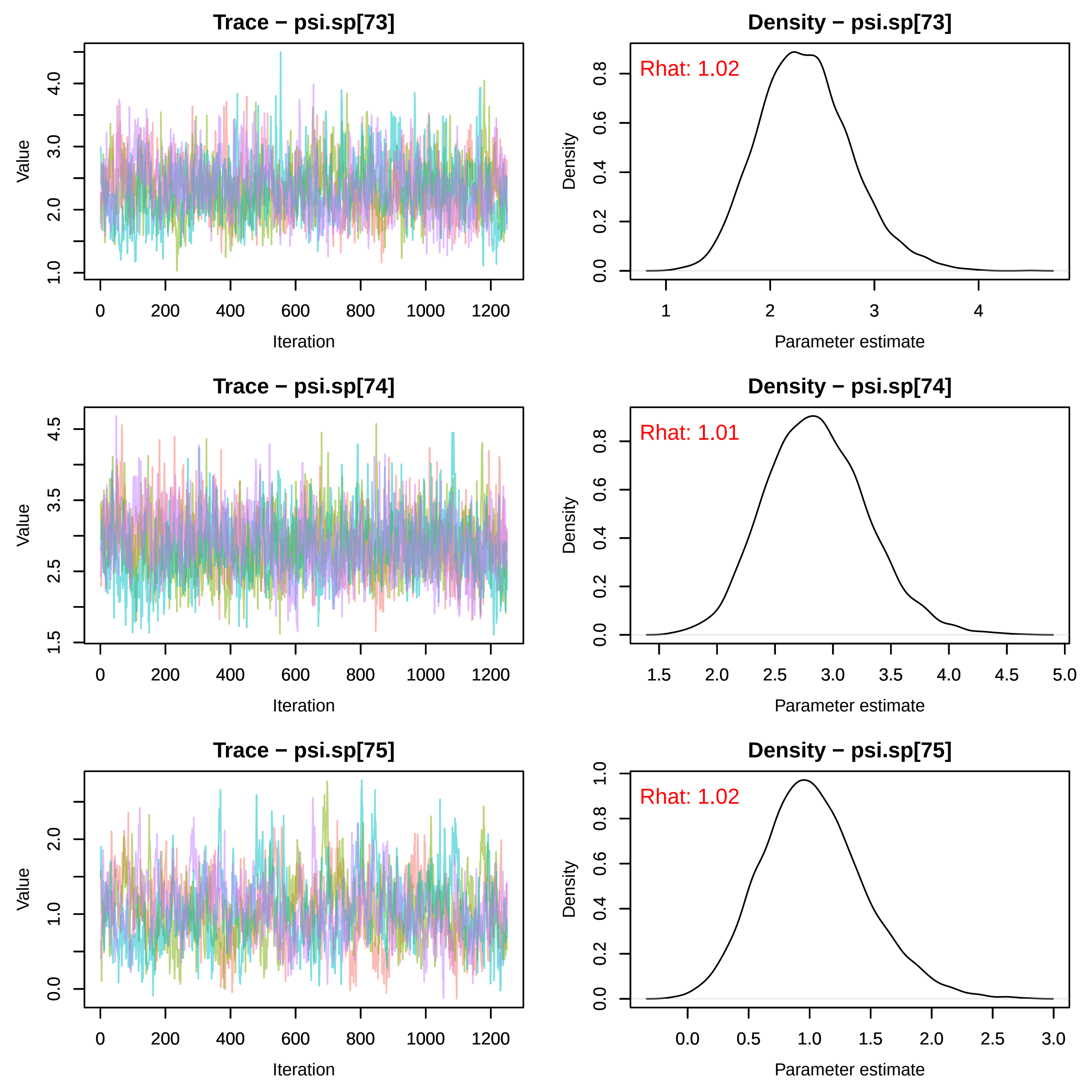
Trace − psi.sp[73]
Density − psi.sp[73]
Rhat: 1.02
Trace − psi.sp[74]
Density − psi.sp[74]
Rhat: 1.01
Trace − psi.sp[75]
Density − psi.sp[75]
Rhat: 1.02
Value
Iteration
Density
Parameter estimate
0
200
400
600
800
1000
1200
1.0
2.0
3.0
4.0
1.5
2.5
3.5
4.5
0.0
0.2
0.4
0.6
0.8
1
2
3
4
5.0
0.5

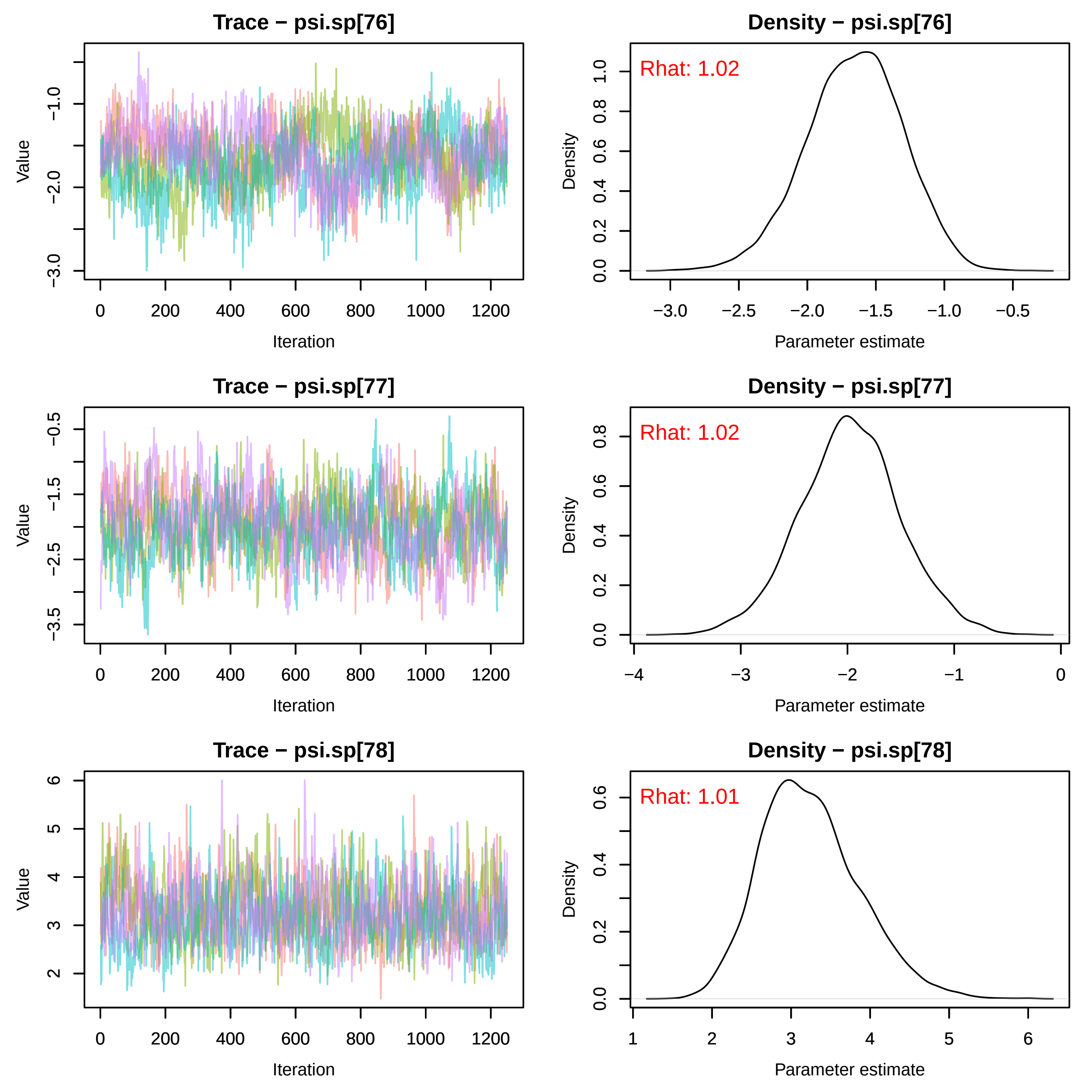
Trace − psi.sp[76]
Value
Iteration
Density − psi.sp[76]
Rhat: 1.02
Density
Parameter estimate
Trace − psi.sp[77]
Value
Iteration
Density − psi.sp[77]
Rhat: 1.02
Density
Parameter estimate
Trace − psi.sp[78]
Value
Iteration
Density − psi.sp[78]
Rhat: 1.01
Density
Parameter estimate

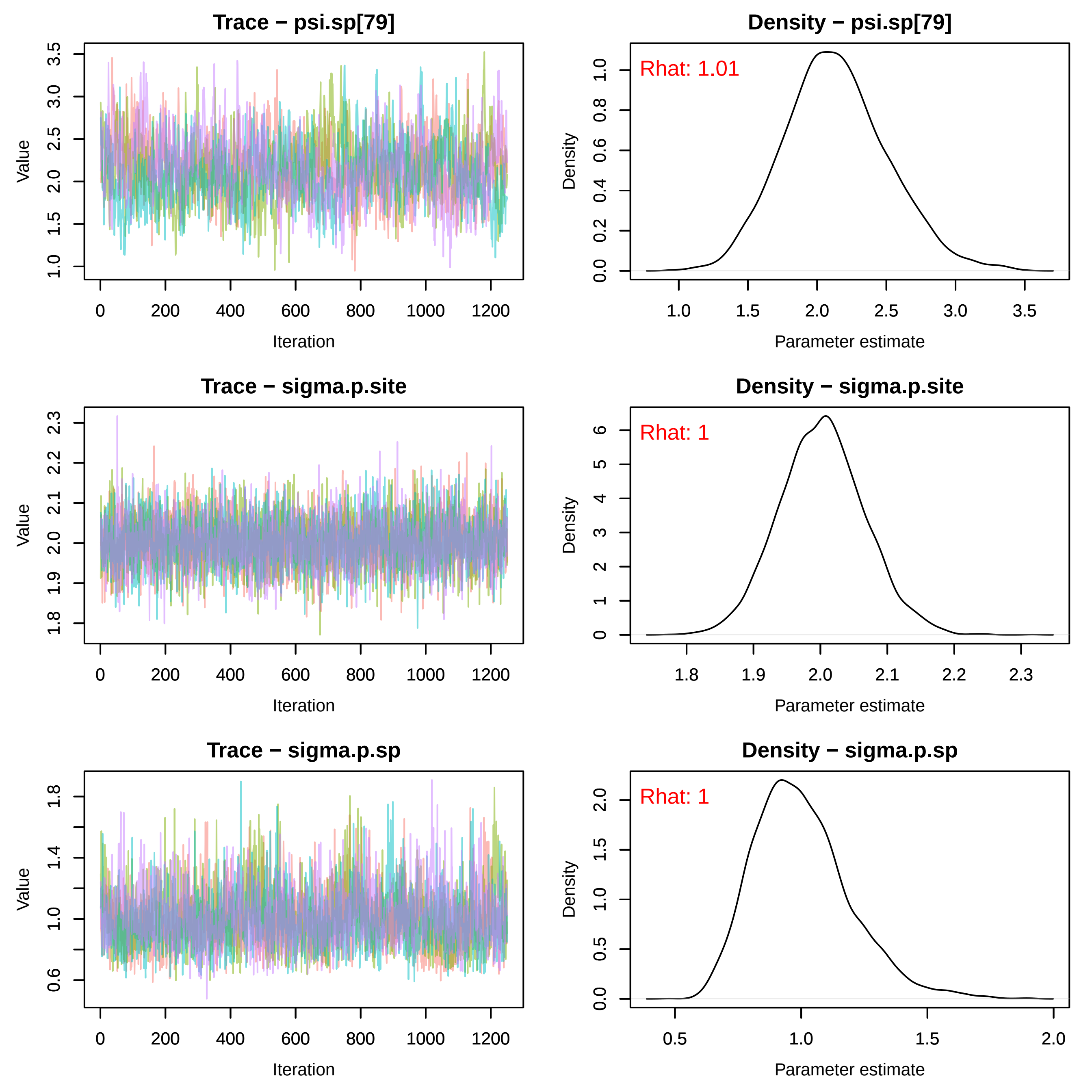

Trace − psi.sp[79]
Value
Iteration
Density − psi.sp[79]
Rhat: 1.01
Density
Parameter estimate
Trace − sigma.p.site
Value
Iteration
Density − sigma.p.site
Rhat: 1
Density
Parameter estimate
Trace − sigma.p.sp
Value
Iteration
Density − sigma.p.sp
Rhat: 1
Density
Parameter estimate

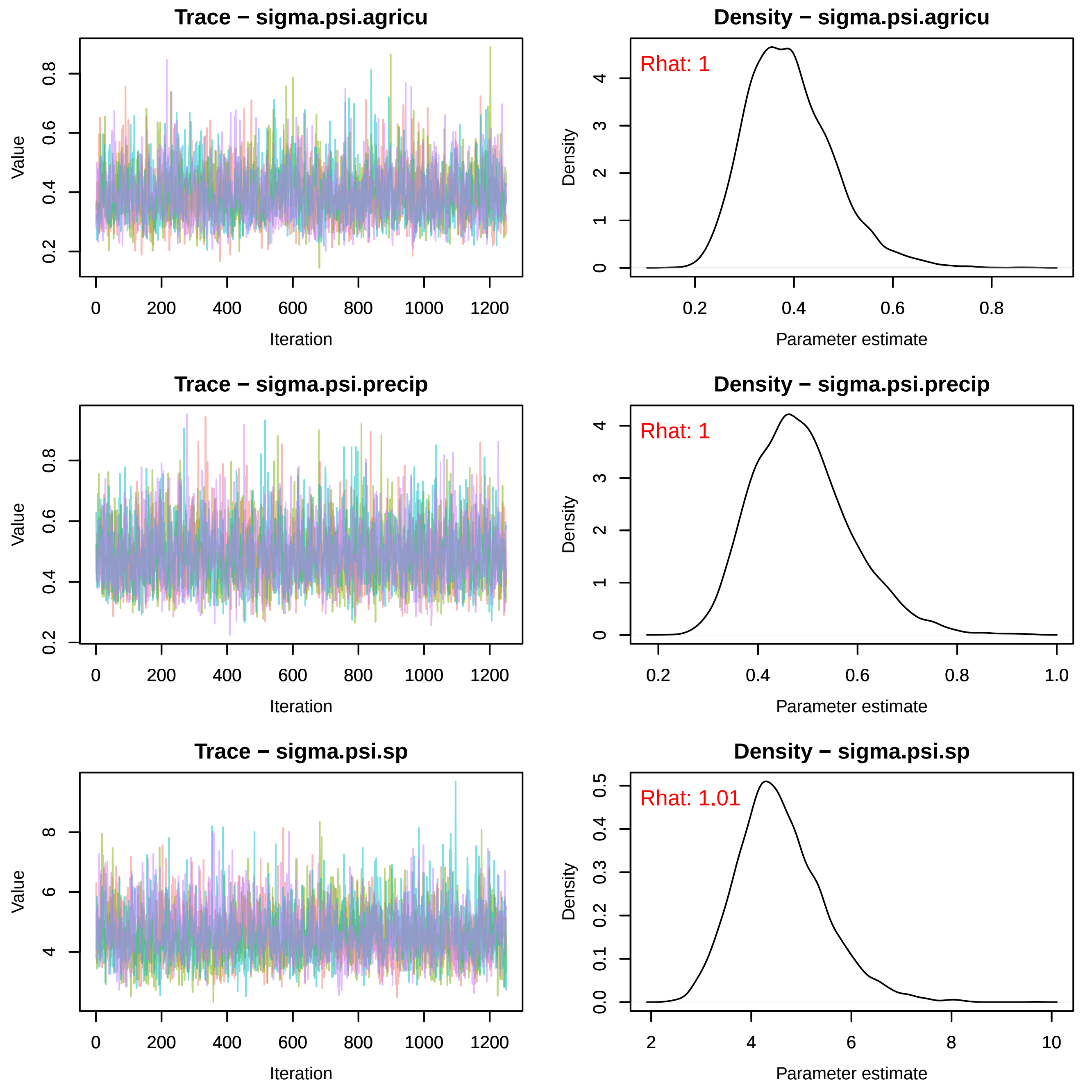
Trace − sigma.psi.agricu
Value
Iteration
0
200
400
600
800
1000
1200
0.2
0.4
0.6
0.8
Density − sigma.psi.agricu
Rhat: 1
Density
Parameter estimate
0.2
0.4
0.6
0.8
0
1
2
3
4
Trace − sigma.psi.precip
Value
Iteration
0.2
0.4
0.6
0.8
Density − sigma.psi.precip
Rhat: 1
Density
Parameter estimate
0.2
0.4
0.6
0.8
1.0
Trace − sigma.psi.sp
Value
Iteration
4
6
8
Density − sigma.psi.sp
Rhat: 1.01
Density
Parameter estimate
2
4
6
8
10
0.0
0.1
0.2
0.3
0.4
0.5

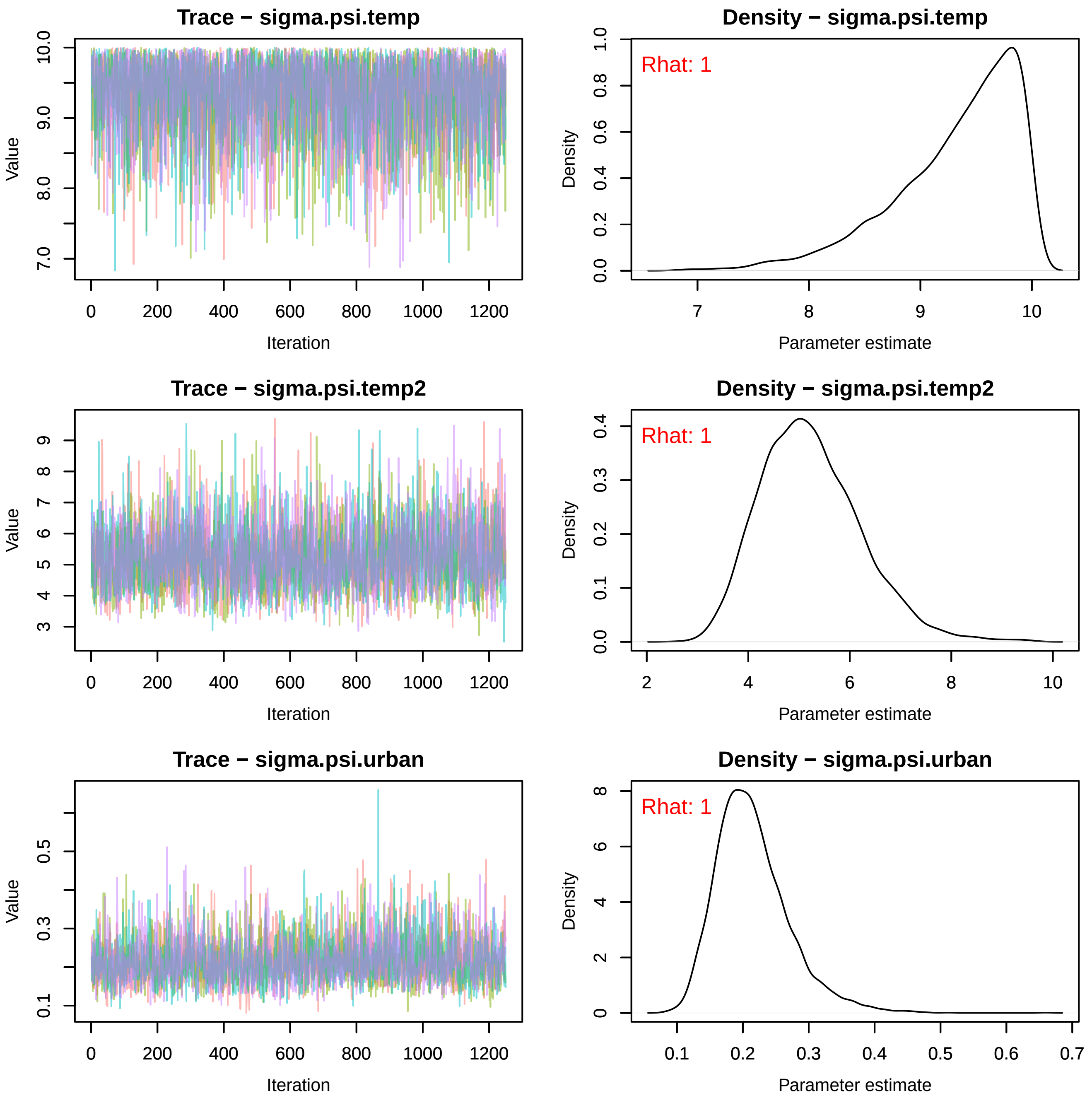

Trace − sigma.psi.temp
Value
Iteration
Density − sigma.psi.temp
Rhat: 1
Density
Parameter estimate
Trace − sigma.psi.temp2
Value
Iteration
Density − sigma.psi.temp2
Rhat: 1
Density
Parameter estimate
Trace − sigma.psi.urban
Value
Iteration
Density − sigma.psi.urban
Rhat: 1
Density
Parameter estimate
